# Exploitation of phylum-spanning omics resources reveals complexity in the nematode FLP signalling system and provides insights into *flp*-gene evolution

**DOI:** 10.1101/2024.08.25.609560

**Authors:** Ciaran J. McCoy, Christopher Wray, Laura Freeman, Bethany A. Crooks, Luca Golinelli, Nikki J. Marks, Liesbet Temmerman, Isabel Beets, Louise E. Atkinson, Angela Mousley

**Author notes:** Corresponding senior author. **Corresponding Author** Angela Mousley.

## Abstract

**Background:** Parasitic nematodes significantly undermine global human and animal health and productivity. Parasite control is reliant on anthelmintic administration however over-use of a limited number of drugs has resulted in escalating parasitic nematode resistance, threatening the sustainability of parasite control and underscoring an urgent need for the development of novel therapeutics. FMRFamide-like peptides (FLPs), the largest family of nematode neuropeptides, modulate nematode behaviours including those important for parasite survival, highlighting FLP receptors (FLP-GPCRs) as appealing putative novel anthelmintic targets. Advances in omics resources have enabled the identification of FLPs and neuropeptide-GPCRs in some parasitic nematodes, but remaining gaps in FLP-ligand libraries hinder the characterisation of receptor-ligand interactions, which are required to drive the development of novel control approaches.

**Results:** In this study we exploited recent expansions in nematode genome data to identify 2143 *flp*-genes in >100 nematode species across free-living, entomopathogenic, plant, animal and human lifestyles and representing 7 of the 12 major nematode clades (1). Our data reveal that: (i) the phylum-spanning *flps, flp*-1, −8, −14, and −18, may be representative of the *flp* profile of the last common ancestor of nematodes; (ii) the majority of parasitic nematodes have a reduced *flp* complement relative to free-living species; (iii) FLP prepropeptide architecture is variable within and between *flp*-genes and across nematode species; (iv) FLP prepropeptide signatures facilitate *flp*-gene discrimination; (v) FLP motifs display variable length, amino acid sequence, and conservation; (vi) CLANS analysis provides insight into the evolutionary history of *flp*-gene sequelogues and reveals putative *flp*-gene paralogues and, (viii) *flp* expression is upregulated in the infective larval stage of several nematode parasites.

**Conclusions:** These data provide the foundation required for phylum-spanning FLP-GPCR deorphanisation screens in nematodes to seed the discovery and development of novel parasite control approaches.

## Background

The next generation sequencing revolution has advanced helminth research through the generation of omics datasets for at least 130 nematode species, many of which have global scale impacts on international health and economy (2, 3). Nematode omics resources, which are publicly available through WormBase ParaSite (4), enable large scale *in silico* analyses that drive fundamental nematode research and underpin the development of novel parasite control strategies and diagnostic tools. Indeed, escalating reports of anthelmintic-resistant nematodes underscores the urgency in the requirement for novel therapeutics and diagnostics (5).

Nematode neuropeptide signalling is a key focus for novel drug target discovery in parasites, where neuropeptides and their receptors play prominent roles in modulating essential nematode behaviours (6). Indeed, nematode neuropeptide receptors are an untapped resource for anthelmintic control, however have not been exploited chemotherapeutically (6, 7). Realising the promise of novel drug targets within the nematode neuropeptide signalling system relies on pan-phylum data for receptor conservation, function, and deorphanisation. To date, receptor-peptide interaction data has been derived primarily from *Caenorhabditis elegans* where ∼40% of predicted nematode neuropeptide receptors [G-protein coupled receptors (GPCRs)] have been matched with their cognate neuropeptide ligands through heterologous expression approaches (8, 9). Deorphanisation of neuropeptide receptors in parasites has proven significantly more difficult, where only a few examples of heterologous and functional deorphanisation have been reported (10, 11).

In recent years there has been significant progress in expanding our knowledge of the nematode neuropeptide signalling system through the exploitation of omics datasets. Indeed, we now have neuropeptide ligand and GPCR profiles across a number of important nematode parasites (12–17). However, to successfully validate neuropeptide GPCRs as novel drug targets, we need to exploit omics data on a pan-phylum scale. These efforts will generate comprehensive libraries of GPCRs and ligands that enable the downstream interrogation of receptor-ligand interactions and prioritisation of neuropeptide GPCRs for therapeutic exploitation.

Here we report a phylum-spanning profile of the largest family of nematode neuropeptides (FMRFamide-like peptides, FLPs) across 109 nematode species, that will provide insights into the diversity and evolution of nematode *flp*-gene complements. The integration of the neuropeptide ligand profile data presented here with equivalent pan-phylum neuropeptide GPCR data paves the way for a new wave of innovative research to unravel neuropeptide signalling biology in nematodes that has broad reaching relevance to neurobiology, evolutionary biology, and drug target discovery.

## Results and Discussion

In recent years there has been significant progress in the scale and scope of helminth omics data; indeed, we now have a dedicated resource for helminth omics datasets, through WormBase Parasite, which has rapidly expanded to include genome and transcriptome data for >100 nematode species (4). These pan-phylum expansions in nematode omics data enable comprehensive analyses of *flp*-gene profiles across nematode species that represent diverse clades and lifestyles to transform our understanding of nematode FLP signalling systems. Previous analyses of *flp-*gene profiles in nematodes were significantly limited by species number and the quality of nematode omics resources (13, 15, 18) which challenges our ability to unravel nematode FLP ligand-receptor interactions that are critical to understanding FLP evolutionary and functional biology. In this study we significantly expanded analyses of *flp*-gene profiles across all publicly available nematode genomes (representing 108 species excluding C. elegans; 4) spanning 7 of the 12 major nematode clades and including free-living (FLN), entomopathogenic (EPN), plant- (PPN), animal- (APN) and human-parasitic species (HPN) (1, 19; see Figure 1).

**Figure 1.**
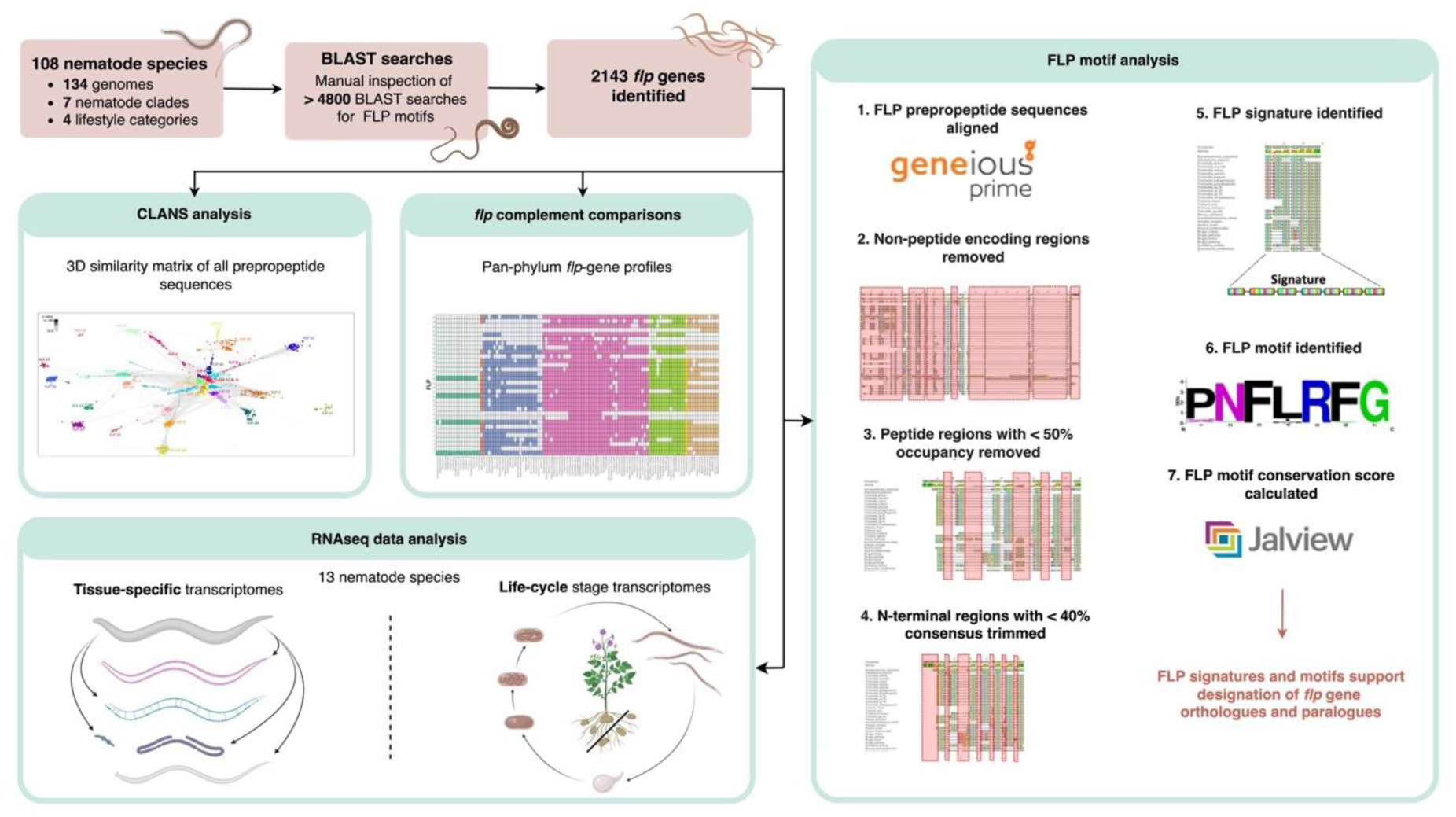
Bioinformatics pipeline for the identification and analysis of *flp*-genes, FLP signatures and FLP motifs. The bioinformatics pipeline enabled the identification of 2143 nematode *flp*-gene sequelogues using BLAST supported on the WormBase ParaSite server. Downstream analyses resulted in the identification of pan-phylum *flp*-gene profiles, FLP signatures and FLP motifs, and informed insights into *flp*-gene evolution and patterns in *flp*-gene expression.

Our BLAST-based approach has identified >2000 *flp*-gene sequelogues (see Figure 1) which, for the first time, captures pan-phylum *flp* diversity and reveals key insights into the complexity of nematode neuropeptidergic signalling. Through the analyses of significantly expanded datasets we have enhanced data to further inform several of the findings previously reported by McCoy, Atkinson (13) including: (i) nematode *flp*-gene profiles display both inter- and intra-clade diversities and similarities (Figure 2 and Figure 3A-C); (ii) the majority of parasitic nematodes have a reduced *flp* complement relative to free-living species (Figure 4A and B) and, (iii) several *flp*-genes are phylum-spanning whilst others display restricted profiles (Figure 2). Significantly, the generation of the phylum-spanning *flp*-gene profiles presented here, using all publicly available nematode genomics data, have facilitated new analyses that drill deeper to interrogate *flp*-gene expression, evolution and architecture. These novel detailed analyses advance our understanding of FLP neurosignalling in nematodes to reveal that: (iv) four *flp*-encoding genes (*flp*-1, −8, −14, and −18) display phylum spanning profiles (>87% conservation) and were likely represented in the *flp*-profile of the last common ancestor (LCA) of nematodes; (v) CLANS analyses support *flp*-gene sequelogue identification and provide insights into *flp* evolutionary history; (vi) FLP prepropeptide architecture is diverse within and between *flp*-genes and nematode species; (vii) phylum-spanning *flps* (*flp*-1, −8, − 14, and −18) display highly conserved motifs and, (viii) *flp*s with more restricted profiles (<50% conservation) display variability in motif conservation.

**Figure 2.**
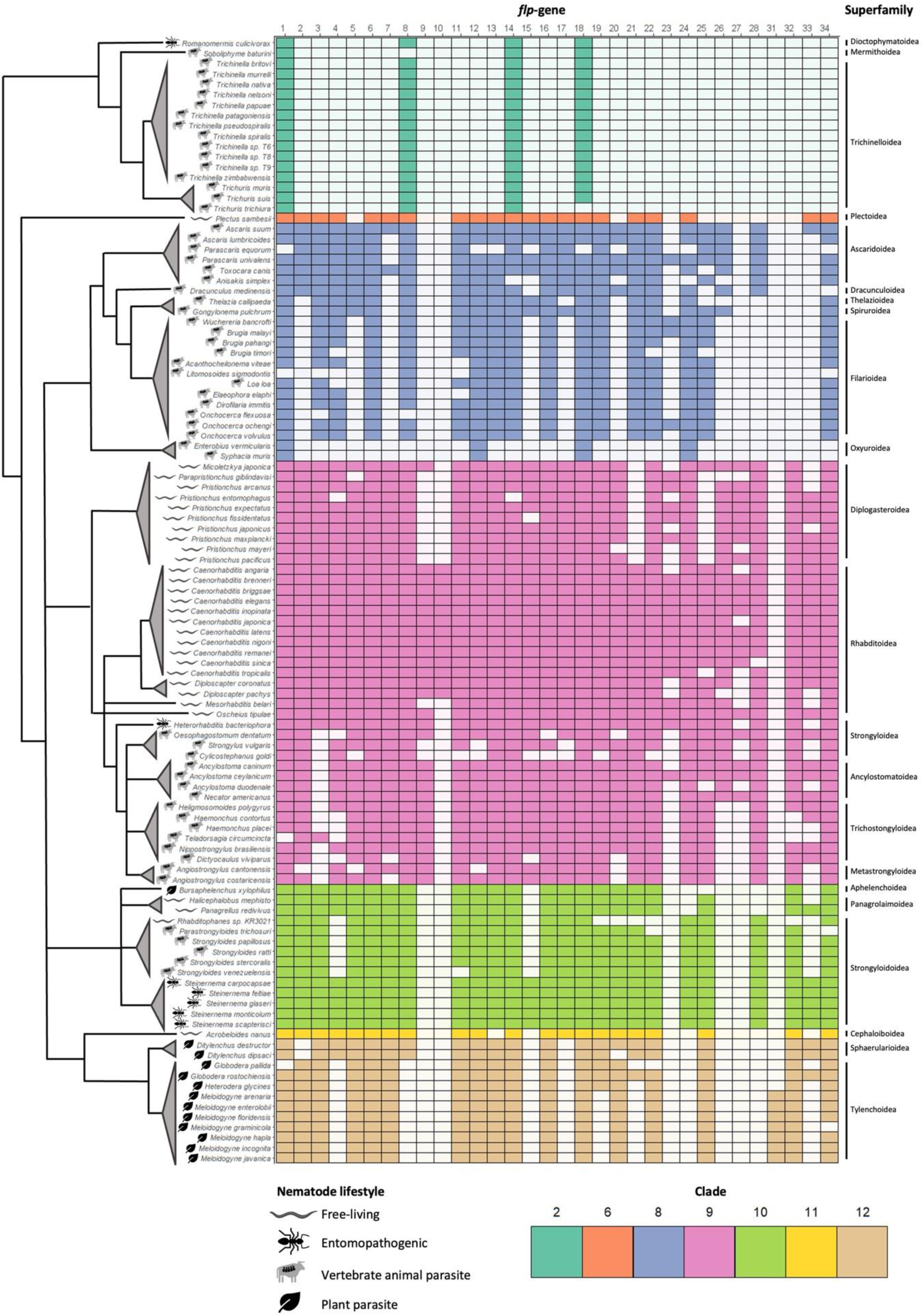
Pan-phylum *flp*-gene profiles. Coloured boxes indicate the presence of a *flp*-gene sequelogue in 109 nematode species, arranged according to phylogenetic clade and superfamily (Holterman, et al. 2006; van Megen, et al. 2009). Branch lengths are arbitrary. Nematode lifestyle categories (free-living, entomopathogenic, vertebrate animal parasite, plant parasite) are indicated by symbols displayed in the key.

**Figure 3.**
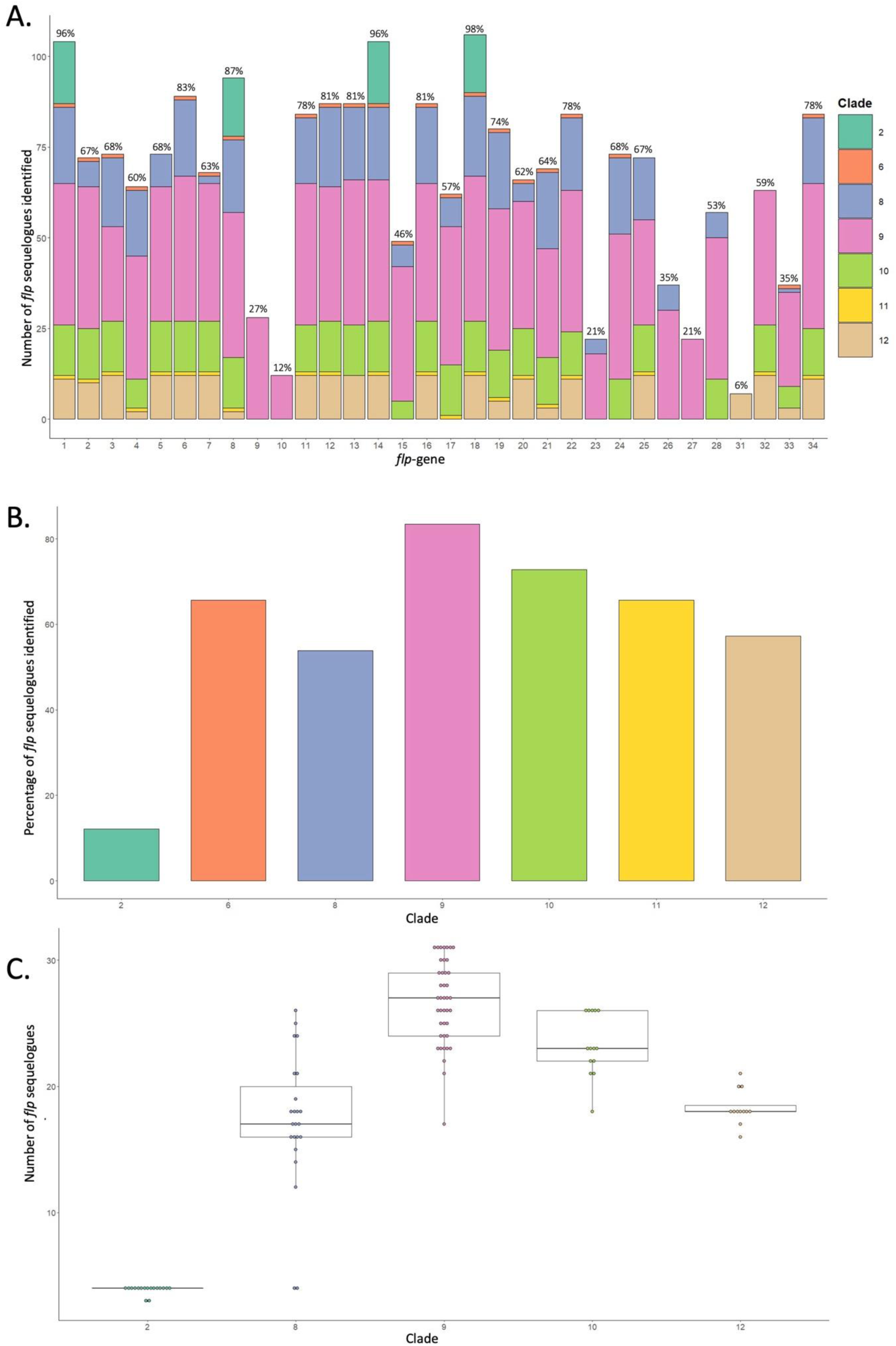
Distribution of *flp*-gene complements across nematode clades. (A) Total number of *flp-* gene sequelogues identified and their distribution within each clade. The percentage of nematode species that encode each *flp*-gene is noted above each bar. (B) Percentage of *flp*-gene sequelogues identified within each clade. (C) Number of *flp*-gene sequelogues identified within each clade; each data point represents an individual species. Data points are coloured according to clade (see legend). Note that clades 6 and 11 were excluded from analyses as they are represented by only one species each.

**Figure 4.**
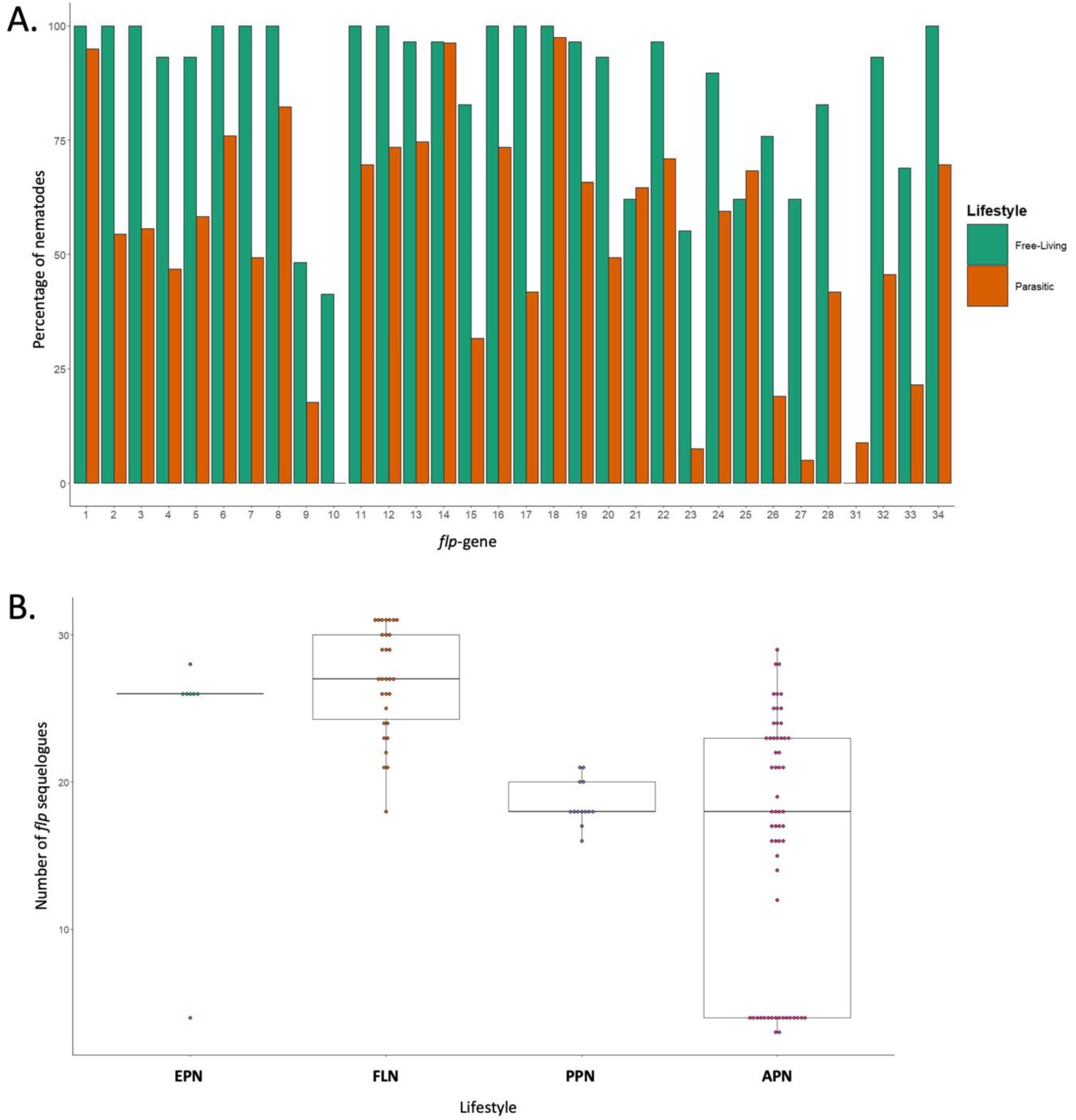
*flp*-gene profiles across nematode lifestyles. (A) Percentage of nematodes encoding each *flp*-gene in free-living and parasitic nematode species. (B) Number of *flp*-gene sequelogs identified within each lifestyle category [entomopathogenic nematode (EPN); free-living nematode (FLN); plant parasitic nematode (PPN); vertebrate animal parasitic nematode (APN)]. Each data point represents an individual species. Note that clades 6 and 11 were excluded from analyses as they are represented by only one species each.

### The phylum-spanning *flp*-genes*, flp*-1, −8, −14, and −18, represent the *flp* profile of the last common ancestor of nematodes

*flp*-1, −8, −14, and −18 were identified in >87% of species examined in this study (96%, 87%, 96%, and 98% of nematodes encoded *flp*-1, −8, −14, and −18 respectively) and in all nematode clades (Figure 2 and Figure 3A), highlighting their potential importance to nematode biology. Indeed, the peptides encoded on *C. elegans flp-*1, −8 and −14 are amongst the promiscuous nematode neuropeptides (interacting with up to 12 GPCRs; 8), supporting their involvement in multiple distinct signalling pathways.

These data suggest that the LCA of all nematodes is likely to have encoded at least four FLP-encoding genes (*flp*-1, −8, −14 and −18; see Figure 2) which appear to be conserved in almost all nematode species since the Precambrian period (20). However, it is also possible that the LCA of all nematodes displayed an expanded *flp* complement, beyond *flp*-1, −8, −14, and −18, and that the reduced *flp*-complement observed for clade 2 nematodes may reflect a loss of a proportion of the ancestral *flp* complement in this lineage; this is supported by the apparent loss of *flp*-34, an orthologue of the bilaterian-conserved NPY/NPF family of RFamide peptides, in clade 2 (21). Alternatively, the *flp*-gene complement may have expanded significantly within the lineages that led to the crown nematode clades 8-12 (1). Indeed, it is more likely that the current profiles represent a combination of these scenarios. Genome sequencing projects for representative nematodes from clades 1-5, as well as members of the sister phylum Nematomorpha (horse-hair worms), may enable expanded analyses to better understand the evolution of nematode *flp*-genes.

Clade 2 nematodes are considered more basal relative to the crown clades (clades 8-12; 1). Interestingly, some basal nematode species display more variable neuronal connectomes comprising larger numbers of neurons [e.g. >1000 neurons in *Pontonema spp*. (Clade 1) and *Mermis negrescens* (Clade 2)] compared to crown clade species [e.g. *Ascaris suum* (Clade 8; 298 neurons) and *C. elegans* (Clade 9; 302 neurons)] which display anatomically similar connectomes (22). It is possible that molecular diversification of neuropeptide-encoding genes within the crown clade neuronal genomes may compensate for the relative anatomical simplicity of crown clade nematode nervous systems (22), however expansion in the number of complete neuronal maps for nematodes (including the clade 2 species examined here) will enhance interpretation of these data. Interestingly, Han, Boas (23) revealed variation in the neuroanatomy of several crown clade nematode species that underscores the importance of expanding nematode connectome data to enable the direct comparison between the variable complexities of anatomical architectures and chemical compositions.

Our data indicate that the LCA of all crown clade nematodes broadly displayed a *flp* complement similar to *C. elegans* [at least 28 *flp*-genes encoded across crown clade nematodes; *flp*-1-8, 11-28, - 33 and −34 (Figure 2)]. Note that *flp*-23 and −26 appear to have been lost from clade 10-12 species, whilst *flp*-24 and −28 are also absent in clade 11 and 12 nematodes (Figure 2 and Figure 3). The only clade 6 species examined here, *Plectus sambesi*, also exhibits a *flp* complement largely similar to the LCA of the crown clades (21 *flp*-genes; Figure 2). A small proportion of the nematode *flp* complement appears to have evolved relatively recently, where *flp*-9, −10 and −27 are only present in clade 9 nematodes, and *flp*-31 is *Meloidogyne spp*. specific (Figure 2 and Figure 3A).

McCoy, Atkinson (13) indicated that *flp*-gene profiles broadly display inter-clade diversity and intra-clade similarity; however, expansion of these analyses has now revealed differences in *flp*-gene profiles within clades that may reflect *flp*-gene losses within specific nematode lineages (Figure 2). Indeed, this is clearly evidenced in clade 8 where: (i) the Ascaridoidea (≤26 *flp*-gene sequelogues) display broader *flp* complements that appear to be more ancestral; (ii) the Filarioidea (≤18 *flp*-gene sequelogues) exhibit reduced *flp*-complements relative to Ascaridoidea, and (iii) the pinworms (Oxyuroidea) appear to have lost all but four of the ancestral *flp*-genes (Figure 2). Distinct *flp*-gene loss events are also evident within clade 9-12 species; for example, *Meloidogyne spp*. have lost several genes (*flp*-4, −8, −19, −21 and −33) relative to the LCA of all clade 12 PPNs; Figure 2). Although some nematode clades appear to display reduced intra-clade diversity compared to others [e.g. clade 2 species (average 3.9 ± 0.3 *flp*-genes) vs clade 8 (average 17.2 ± 5.5 *flp*-genes); see Figure 3C], the variability in the number of nematode genomes available that represent genera and evolutionary lineages for each clade caveats conclusive comparative analyses at this stage.

### The majority of parasitic nematodes have a reduced *flp* complement relative to free-living species

On average, parasitic nematode genomes encode fewer *flp*-genes relative to *C. elegans* and most other FLNs (Figure 4A). However, not all parasites display reduced *flp* complements; for example, EPNs (e.g. *Steinernema* spp.) appear to display *flp* complements similar to FLNs with the exception of the EPN *Romanomermis culicuvorax* (Figure 2 and Figure 4B). In contrast, both PPNs and APNs tend to display reduced *flp* complements relative to FLNs, where a subset of APNs have much more restricted *flp* profiles [clade 2 species and Oxyuroidea (pinworms; Clade 8); Figure 2]. While the available data point towards these emerging trends they are caveated by limitations including differential representation across nematode lifestyles, clade, and genera, where the species analysed here are biased towards economically, medically and scientifically relevant nematodes and may include many species within the same genus (e.g. 11 *Caenorhabditis* spp.). In addition, these broad lifestyle categories do not necessarily reflect the significant divergence between specific parasitic lifecycles and host niches, or the independent evolution of the representative parasites in these groups (20), that may collectively impact *flp*-gene profiles. Despite the reduced *flp* complement within parasitic nematodes relative to FLNs, there are several *flp*-genes that appear to be equally distributed across both lifestyles (*flp*-1, −14 and −18; see Figure 4A), underscoring their potential importance to nematode biology.

### FLP prepropeptide architecture is variable within and between *flp*-genes and across nematode species

FLP prepropeptide architecture is characterised by the number and relative position of FLP peptides. Until now, limited numbers of publicly available nematode genomes prevented pan-phylum analyses of FLP prepropeptide architecture. Here we show that nematode *flp*-genes display diversity in the numbers of peptides encoded, ranging between 1-13 FLPs (Figure 5A). Several *flp*-genes (*flp*-10, −12, −21, −23, −27, −28, −31, −32, and −33) encode a single peptide in all species that encode these genes, while other *flp*-genes display a broader range in the number of peptides encoded, for example *flp*-1, −3, −7, −13, and −18 (see Figure 5A). Interestingly, *flp*-genes that display high mean peptide numbers also display a high range in the number of peptides encoded (Figure 5A).

**Figure 5.**
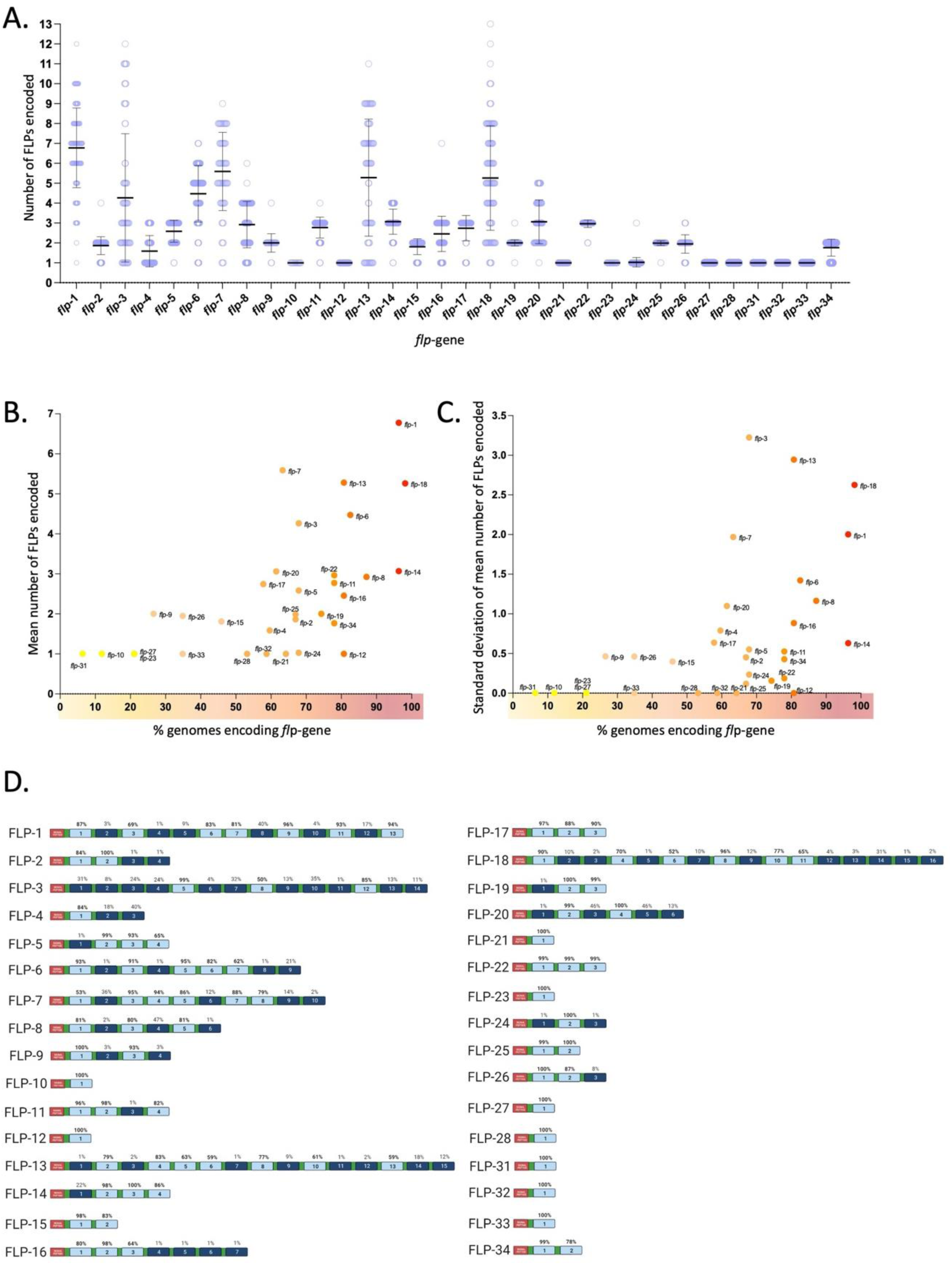
Analysis of FLPs encoded by *flp*-gene sequelogues. (A) The number of FLPs encoded on each *flp*-gene, by each individual species, is represented by a single data point (blue circle). The average number of FLPs encoded for each *flp*-gene is represented by a black bar. (B) Correlation of the mean number of FLPs encoded by each *flp*-gene and the percentage of nematode genomes that encode that *flp*-gene. Each data point is coloured according to the percentage of genomes encoding that specific *flp*-gene, where yellow represents a low percentage of genomes and red represents a high percentage of genomes. (C) Correlation of the standard deviation of the mean number of FLPs encoded by each *flp*-gene and the percentage of nematode genomes that encode that *flp*-gene. Each data point is coloured according to the percentage of genomes encoding that specific *flp*-gene, where yellow represents a low percentage of genomes and red represents a high percentage of genomes. (D) FLP prepropeptide architecture as represented by a schematic which provides an overview of the maximum number of peptides that could be encoded by each *flp*-gene. Note that the arrangement of peptides encoded will vary according to species. FLPs are numbered sequentially from the N- to C-terminus and the percentage of nematode species that encode each FLP peptide are noted above each bar. Peptide regions with >50% peptide occupancy are coloured light blue, peptide regions with <50% peptide occupancy are coloured dark blue. Signal peptides are coloured red. Peptide flanking cleavage sites are coloured green. Full multiple sequence alignments relevant to this schematic are provided in Supplementary File 2.

Our data also reveal a positive correlation between the degree of *flp*-gene conservation and (i) the mean number of peptides encoded (Spearman’s rho = 0.66; p<0.0001; Figure 5B), and (ii) the range in the number of peptides encoded (represented by the standard deviation; Spearman’s rho = 0.59; p<0.001; Figure 5C). These data indicate that there may be a selection pressure for genes encoding higher numbers of peptides, which could facilitate the production of higher quantities of mature peptides, and/or greater diversification in peptide sequence that have the potential to relay distinct receptor interactions and functions.

Interrogation of FLP prepropeptide architecture, in the context of the relative position of the encoded FLP peptides, is challenged by the ability to generate accurate multiple sequence alignments (MSAs) for prepropeptides that display diversity in FLP peptide number and motif conservation. Despite this, it is possible to curate MSAs to generate representative consensus prepropeptide architectures for a specific *flp*-gene across all species where all of the potential peptides encoded by that gene are numbered sequentially from the N- to C-terminus (Figure 5D and Supplementary Figure 1). These analyses reveal that FLP prepropeptides display variable FLP peptide position conservation where the relative position of some peptides are highly conserved, e.g. peptide at position 2 on the FLP-2 prepropeptide is present in 100% of the species that encode *flp*-2, compared to others, e.g. peptides at positions 3 and 4 on the FLP-2 prepropeptide are only present in one of the species that encode *flp*-2 (Figure 5D and Supplementary Figure 1). These data have the potential to facilitate the identification of peptide duplication and loss events that can be mapped to phylogenetic ancestors providing novel insights into *flp*-gene evolution. For example, for *flp*-2, the LCA of *Meloidogyne*, *Heterodera* and *Globodera* spp. appears to have lost one of the two more ancestral FLP-2 peptides (see Supplementary Figure 1). Other examples include: (i) *flp-11*, where the LCA of all Clade 8 species possessed the more divergent peptide (at peptide position 4; NGAPQPFVRFamide), which was subsequently lost by the LCA of the clade 8 filarid nematodes, *Gongylonema pulchrum,* and *Thelazia callippaeda* (Supplementary Figure 1); and (ii) *flp-15*, where Clade 8 spp. that possess *flp-15* have lost one of the two more ancestral peptides (at peptide position 2; Supplementary Figure 1). For other *flp*-genes, it is more difficult to identify peptide expansions and/or losses challenging our ability to define the evolutionary history within specific nematode lineages. This observation may support rapid evolutionary change of peptide numbers between species and lineages for some *flp*-genes.

### FLP prepropeptide signatures facilitate *flp*-gene discrimination

The enhanced breadth of this study enables detailed analyses of FLP prepropeptide signatures and FLP motif conservation on a pan-phylum scale. Defining peptide motifs based on amino acid sequence conservation is challenging (24); however, in this study highly conserved FLP peptides were used to characterise prepropeptide signatures and FLP motifs. Specific FLP prepropeptide signatures (see Figure 6A and Supplementary Figure 1) were derived from prepropeptide MSAs and represent the most highly conserved FLP peptides as defined by both peptide occupancy (present in >50% of species that encode specific *flp*) and conservation (displays >40% AA consensus) (see methods; Figure 6A and Supplementary Figure 1). Characterisation of FLP prepropeptide signatures provides an enhanced level of sequence-level information to aid the discrimination of *flp*-gene sequelogues. Here we report FLP prepropeptide signatures for each *flp*-encoding gene that represent the core set of peptides most commonly encoded (>50% of species) by the specific *flp*-gene across the nematode phylum. Consequently, FLP prepropeptide signatures may display a subset of the maximum number of peptides that a specific *flp*-gene can encode (see Figure 5D, Supplementary Figure 1). In addition, the level of amino acid conservation in most FLP peptides reduces from the highly conserved dipeptide C-terminal sequence (RFamide) to the N-terminal residues (see Figure 6A), such that many of the FLP prepropeptide signatures reported here are derived from only a proportion of the full length encoded FLP peptides (see Figure 6A and Supplementary Figure 1). Indeed, this aligns with the significance of the highly conserved FLP C-terminus as demonstrated through physiology studies indicating that N-terminally positioned amino acids are less important for receptor binding and activity (25).

**Figure 6.**
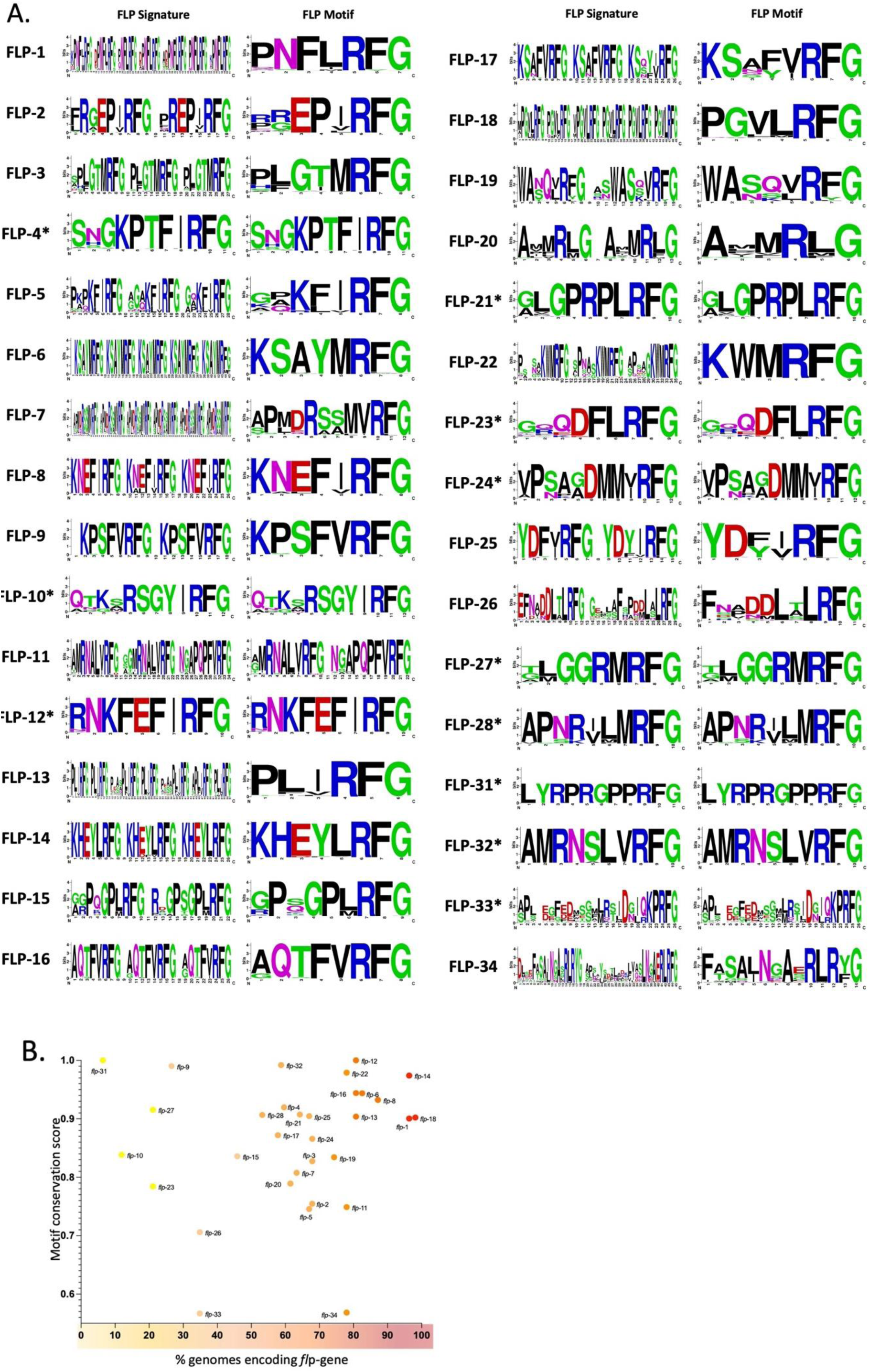
FLP signatures and motifs. (A) WebLogo representation of amino acid sequence and occupancy within FLP signatures and FLP motifs. Identical FLP signatures and motifs are highlighted by an asterix. Amino acid conservation at each position is measured in bits and amino acids are numbered sequentially from the N- to C-terminus. Amino acids are coloured according to chemical properties where polar amino acids are green, basic are blue, acidic are red, and hydrophobic are black. (B) Correlation of the FLP motif conservation score and the percentage of nematode genomes that encode that *flp*-gene. Each data point is coloured according to the percentage of genomes encoding that specific *flp*-gene, where yellow represents a low percentage of genomes and red represents a high percentage of genomes.

### FLP motifs display variable length, amino acid sequence, and conservation

A FLP motif, as defined here, is a unique single short amino acid sequence logo that is representative of the peptides encoded by a specific *flp*-gene. The FLP motif is derived from a MSA of the individual FLP peptides that constitute a specific FLP prepropeptide signature (see Figure 6 and Supplementary Figure 1). For example, the FLP-1 motif (PNFLRFG) is derived from the FLP-1 prepropeptide signature (Figure 6A) which contains seven conserved (>50% occupancy, >40% AA consensus) peptides (of a possible 13; see Figure 5D and Supplementary Figure 1). Our data reveal variability in FLP motif length, where FLP motifs range from 6 (e.g. FLP-13, −20 and −22) to 26 (e.g. FLP-33) amino acids (see Figure 6). Further, variation within the canonical dipeptide C-terminal RFamide signature is also evident for several *flp*-genes; for example, RYamide (e.g. FLP-19, −34), RLamide (e.g. FLP-6, - 19, −20, −25) or RMamide (FLP-20) C-terminal dipeptides (see Figure 6A and Supplementary Figure 1).

Although the majority of FLP motifs are highly conserved (for example, 53% of *flp*-genes encode a motif with a conservation score ≥0.9: *flp-*1, −4, −6, −8, −9, −12, −13, −14, −16, −18, −21, −22, −25, −27, −28, −31, −32; Figure 6B), only 28% of these *flp*-genes are highly conserved across nematode species (>80% of species; *flp*-1, −6, −8, −12, ,-13, −14, −16, −18, −21). Indeed, while several very highly conserved FLP motifs are encoded by highly conserved *flp*-genes, statistical analysis does not support a positive correlation between motif conservation and *flp*-gene conservation (Spearman’s rho = 0.145; Figure 6B).

The majority of multi-peptide encoding *flp*-genes encode peptides that display high AA conservation such that they can be represented by a single highly conserved motif (e.g. FLP-6, −8, −9, −12, −14, −16, −22, −31 and −32; see Figure 5D, 6, and Supplementary Figure 1). In contrast, other FLP encoding genes display a higher degree of FLP peptide AA diversity such that they are represented by a more diverse single motif (e.g. FLP-2, −5, −26, −33 and −34) or, in the case of FLP-11, two distinct peptide motifs (NGAPQPFVRFG and MRNALVRFG; see Figure 6A, and Supplementary Figure 1). Diversity in AA sequence of FLP peptides encoded by the same gene may result in differential GPCR activation and contribute to FLP promiscuity. Indeed, the *Ascaris* FLP-11 peptides NGAPQPFVRFamide and AMRNALVRFamide induce distinct muscle-based response types *in vitro* (26), and the homologous *C. elegans* peptides [Ce-FLP-11-3 (NGAPQPFVRFamide) and Ce-FLP-11-2 (ASGGMRNALVRFamide)] display notably distinct potencies against specific receptors, with Ce-FLP-11-3 representing the most potent ligand of DMSR-1 and DMSR-7, and Ce-FLP-11-2 more strongly activating FRPR-8 and NPR-22 (8). Pan-phylum FLP peptide AA diversity analysis will direct physiology and reverse pharmacology experiments to reveal the influence of peptide sequence on GPCR activation, peptide-receptor promiscuity, and peptide-receptor binding pocket co-evolution.

The pan-phylum scale of this study has also enabled detailed analyses of prepropeptide sequences to reveal that the majority of sequence conservation is restricted to FLP-encoding regions as predicted using existing models (13, 15). However, closer inspection of interpeptide regions reveals short, conserved, stretches of AA sequences within (e.g. FLP-1, −6, −12, −14, −17; Supplementary File 2) and between (e.g. FLP*-*6 and FLP-17; see Supplementary Figure 2) several FLP prepropeptides that could represent novel non-RFamide peptides, or unknown structural or functional domains. Notably, several of these regions, in multiple FLP prepropeptides, display conserved cysteines or stretches of multiple hydrophobic residues (Supplementary File S2, S7, S13, S15 and S18), suggesting that these regions may be related to prepropeptide structure or processing and may regulate neuropeptide processing, protein-prepropeptide interactions, or subcellular propeptide trafficking. Interestingly, several alpha-helical prohormone domains have been shown to be important to endoplasmic reticulum (ER) translocation or secretion of mammalian neuropeptides (27, 28). To reveal the structure of the conserved propeptide regions identified here, AlphaFold was used to model the structure of FLP-1, −12, −14 and 17 from *C. elegans* (Clade 9), *A. suum* (Clade 8) and *Bursaphelenchus xylophilus* (Clade 10); these analyses demonstrate that the conserved propeptide regions form alpha helices as part of the structural model (Supplementary Figure 3; https://alphafold.com/). It is possible that these alpha helical prohormone domains are important to FLP ER translocation or secretion from dense core vesicles in nematodes.

### CLANS analysis provides insight into the evolutionary history of *flp*-gene sequelogues

MSA based neuropeptide phylogenetics are challenged by distinct patterns of evolution and functional constraints displayed by neuropeptides, for example: (i) the short and often rapidly evolving neuropeptide sequences, (ii) the variable numbers of mature neuropeptides encoded on homologous genes, which can be gained or be lost over the course of evolution; and (iii) the often repetitive sequence of neuropeptides encoded on the same prepropeptide (29). To overcome these challenges, we examined the 3D clustering of FLP prepropeptide sequences based on all-against-all sequence similarity analyses where specific *flp*-gene sequelogue groups were represented in CLANS-generated clusters (see Figure 7). Note that the FLP prepropeptides that display lower sequence similarity to other FLP sequences are positioned on the periphery of the CLANs output; e.g. FLP-21, - 23, −31 and −33 clusters do not display connections within any other FLP clusters (E-value cut off = 1E-5; Figure 7A). Other sequence divergent FLP prepropeptides (e.g. FLP-10, −12, −13, −19, −20 −24, - 26, −27, −28 and −34) are indirectly connected to the central FLP clusters through a more limited network of transitive BLAST linkages (Figure 7A). These peripheral clusters predominantly represent FLP prepropeptides that typically encode a single FLP peptide (e.g. FLP-10, −12, −21, −23, −24, −27, −28, −31 and −33; see Figure 8A) and often display more limited conservation across the phylum (e.g. *flp-* 10, −23, −26, −27, −31 and −33 have conservation ranging from 6% (*flp*-31) to 35% (*flp*-26 and *flp*-33); see Figure 3A). Conversely, FLP prepropeptide sequences that encode a higher number of FLPs appeared to cluster more centrally and display a greater network of connections with other FLP clusters (Figure 7A). Removal of peripheral clusters (e.g. those that display no connection with any other FLP clusters; FLP-2, −4, −10, −15, −19, −20, −24, −25, −26, −27, −28, −32, and −34) and increasing the stringency of the E-value cut-off to 1E-10 improved resolution of the relationship between more tightly clustered FLP sequences (Figure 7B). This sequential clustering analysis demonstrated that closely related FLP clusters are split largely into two lobes; (i) lobe 1 including FLP-1, −3, −7, −13, −18, and (ii) lobe 2 including FLP-6, −8, −14, −16, and –17; see Figure 7B). Significantly, all of the most broadly conserved *flp*-genes (*flp-*1, −8, −14 and −18) are represented within lobe 1 or lobe 2; this may support their status as ancestral nematode *flp*-genes from which a proportion of the other *flp*-genes may have arisen via gene duplication.

**Figure 7.**
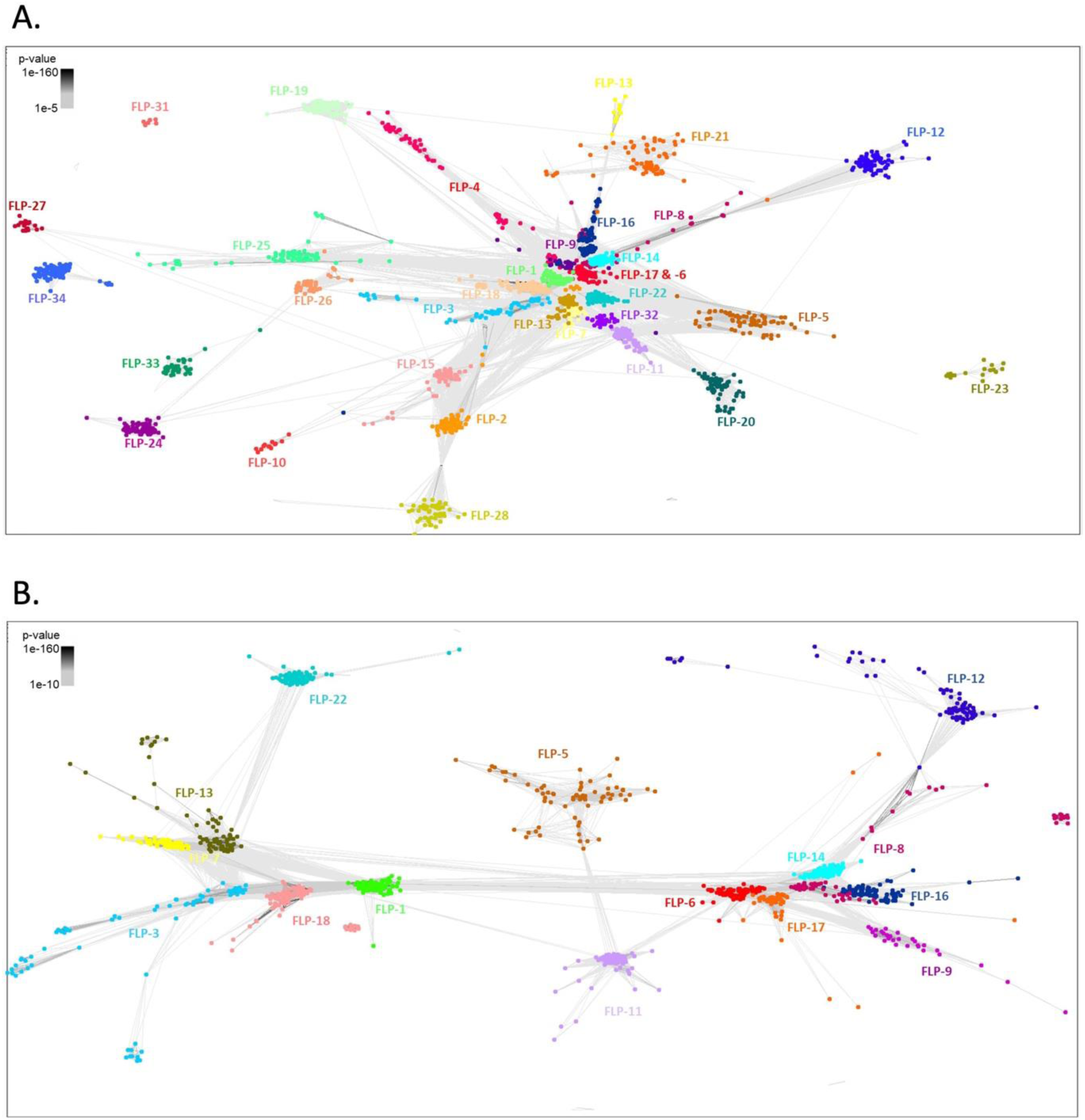
FLP prepropeptide CLANS analysis. (A) Similarity matrix derived from all-against-all BLASTp comparisons between all nematode FLP prepropeptide sequences, E-value limit: 1e-5. (B) Similarity matrix derived from all-against-all BLASTp comparisons between nematode FLP prepropeptide sequences for more similar sequelogue clusters, E-value limit: 1e-10. Individual FLP prepropeptides are coloured according to *flp* gene.

**Figure 8.**
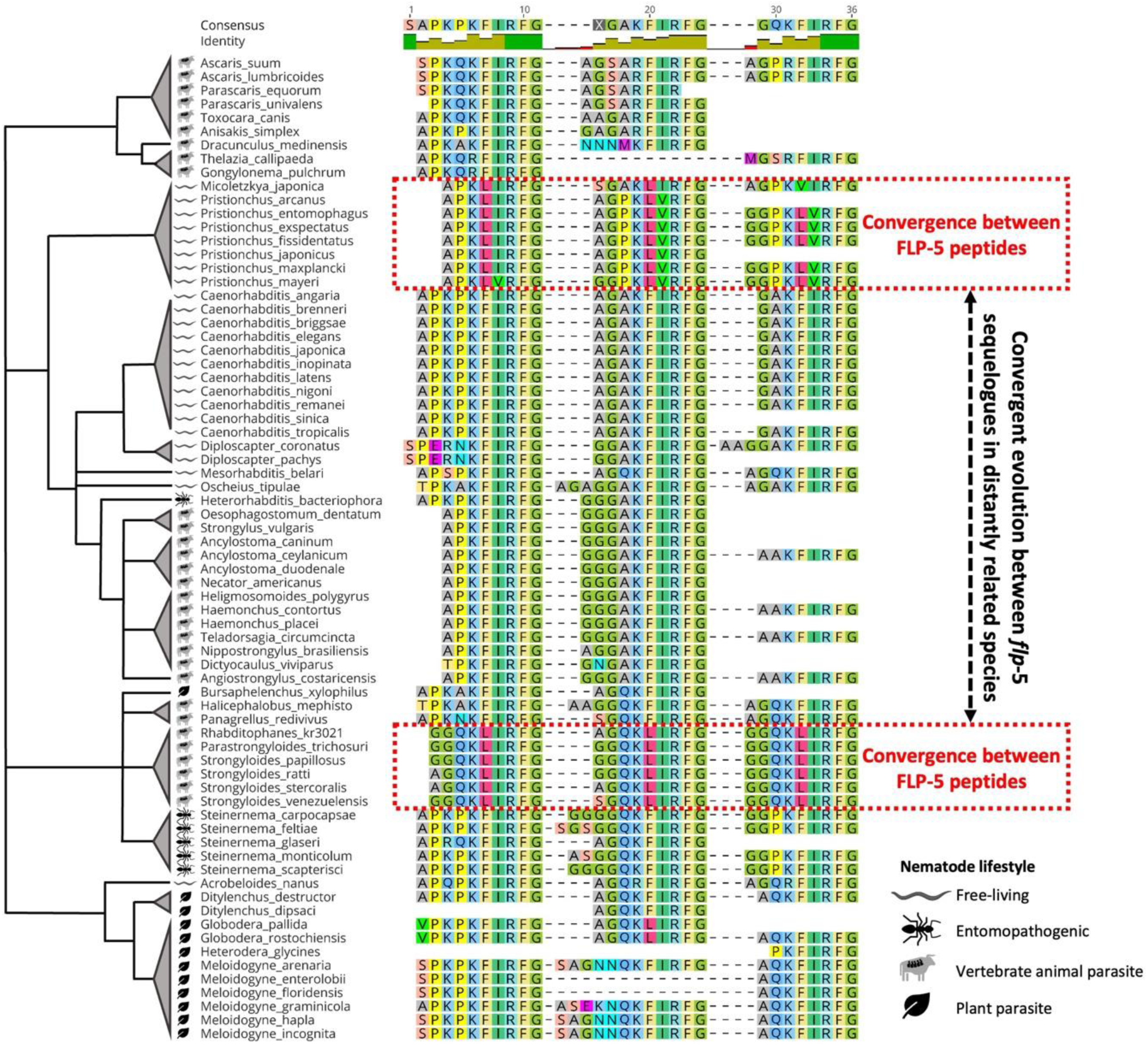
FLP-5 sequence convergence. Multiple sequence alignment of FLP-5 peptides across all *flp-5* sequelogues. Red boxes highlight examples of convergent evolution between FLP-5 peptides encoded by the same *flp*-5 sequelogues. Black dashed arrow highlights convergence of FLP-5 peptides encoded by *flp*-5 sequelogues in distantly related nematode lineages. Nematode species arranged according to phylogenetic clade (Holterman, et al. 2006; van Megen, et al. 2009). Branch lengths are arbitrary. Nematode lifestyle categories (free-living, entomopathogenic, vertebrate animal parasite, plant parasite) are indicated by symbols displayed in the key.

### Integration of CLANS and FLP-motif analyses reveals putative *flp*-gene paralogues

FLP-encoding gene sequelogues that share more similar FLP motifs (see Figure 6A), for example FLP-11 and −32, FLP-8 and −12, FLP-6 and −17, and FLP-4 and −9, and FLP-27 and −34, display biochemically similar amino acids at equivalent positions within their FLP motifs, and also cluster closely through CLANS (see Figure 7 and Figure 2). These data suggest that these FLP-encoding genes may represent paralogues that have the potential to activate the same or closely related GPCRs. However, in these specific cases, the available *C. elegans* deorphanisation data suggest that the majority of these putative paralogous pairs encode peptides that do not activate the same sets of GPCRs (beyond promiscuous FLP-GPCRs), except for FLP-27 and −34, both of which activate NPR-11 (8). This may suggest that many of these paralogues may have adopted distinct roles and/or divergent signalling pathways. However, this analysis is caveated by the absence of deorphanisation datasets beyond *C. elegans*, the promiscuous nature of many nematode GPCRs, and the potential for undiscovered FLP peptides. Further support for *flp*-gene paralogues is demonstrated by FLP-6 and FLP-17 which share sequence similarity beyond the motif; indeed, a region within the prepropeptide sequence between the N-terminal signal peptide and C-terminal neuropeptides is conserved (FCXXFPXLXXC) in the majority of *flp*-6 and −17 sequelogues. (Supplementary Figure 2) bolstering their candidacy as paralogous genes.

In contrast, although lacking similar FLP motifs, we know that *flp*-2, −3, and −28 likely evolved via gene duplication, given that these genes are found clustered together within the intron of a conserved glutamate synthase gene, and display conserved synteny between multiple nematode species (13). The divergent FLP-2, −3 and −28 motifs do not display conserved motif features beyond the C-terminal RFamide (see Figure 6A), which may indicate that *flp*-2, −3 and −28 may have evolved rapidly as a result of reduced functional constraint following gene duplication. This example highlights the difficulties associated with *flp*-gene phylogenetic analysis, where related neuropeptides can display significant sequence divergence, masking their common ancestry. This is also likely true for other divergent *flp*-gene paralogues that can no longer be readily discerned or grouped together.

### Pan-phylum analyses indicate FLP sequence convergence within and between *flp*-genes

Several FLP peptide MSAs display putative convergent evolution between FLPs encoded on the same gene and, in some cases, between FLP motifs across distantly related nematode species. For example, in *Micoletzka japonica* and *Pristionchus* spp three of the peptides encoded on *flp*-5 appear to have diverged from the ancestral KFIRFamide motif, and independently converged to a shared KL(V/I)RFamide motif (Figure 8). A similar pattern of peptide convergence is evident in *Parastongyloides* and *Strongyloides* spp., where *flp*-5 has independently acquired three shared KLIRFamide motifs that are also divergent from the ancestral KFIRFamide motif (see Figure 8).

Several other examples of putative convergent evolution in FLP peptide sequence are evident including: (i) *flp*-2 (*Pristionchus* spp. encode two peptides with EPVRFamide motifs in contrast to the consensus EPIRFamide); (ii) *flp-*6 (in *Meloidogyne* spp. the motifs of two of the four encoded FLPs have diverged from KSAYMRFamide and converged to KSAYMRLamide); (iii) *flp-*11 (in *Meloidogyne* spp. the second *flp-*11 peptide has diverged from a MRNALVRFamide motif and become more similar to the FLP-7 motif RSAMVRFamide); (iv) *flp*-13 (*Strongyloides* and *Parastrongyloides* spp. display multiple peptides that have converged to a consensus PLVRFamide motif and diverged from PLIRFamide); (v) *flp-*15 (*Micoletzkya japonica* and *Pristionchus* spp. peptides have diverged from the GPLRFamide motif into two divergent GPMRFamide-encoding peptides); (vi) *flp*-17 (*Pristionchus* spp. encode three convergent peptides with a KSNFVRFamide motif that have independently diverged from KSAFVRFamide and KSQYRFamide motifs); (vii) *flp*-18 (*Pristionchus spp.* encodes a peptide that has diverged significantly from the PGVLRFamide motif to become more similar to the FLP-16 peptide motif AQAFVRFamide) (see Supplementary Figure 1). Together these data suggest multiple independent peptide sequence convergence events between peptides encoded on the same gene and, in some cases, between *flp*-genes. It would be interesting to explore the NP-GPCR interactions for these examples, to determine if convergent evolution correlates with receptor binding.

### *flp* expression is upregulated in the infective larval stage of several nematode parasites

The most recent meta-analysis of *flp*-gene expression (in *C. elegans*, *S. stercoralis*, *Ancylostoma ceylanicum*, *G. pallida* and *Brugia malayi*) indicated general up-regulation of *flp* transcripts in the dauer-like infective juveniles of the majority of parasites examined (30), suggesting a role for nematode *flps* in host finding and/or infection. Here we extended these analyses to 13 nematode parasites, and show that infective juveniles/dauer stage *flp* up-regulation extends to *Onchocerca volvulus* (L3), *Ancylostoma caninum* (L3), *Teladorsagia circumcinta* (L3), *M. incognita* (J2), and *B. xylophilus* (insect associated dauer) (see Supplementary Figure 4). In contrast, this trend is not evident in *Trichuris muris* (L3) or *Haemonchus contortus* (L3) infective juveniles (Supplementary Figure 4). *flp* upregulation in *Dictyocaulus viviparus* L1 larvae was noted however the infective L3 stage RNASeq data were not available to analyse. Where tissue level RNASeq data exist (primarily from *A. suum*), variable *flp* expression was noted across all tissues types preventing preliminary hypotheses on function (Supplementary Figure 4). In addition, comparing relative expression profiles between *flp*-genes and across all species can begin to reveal trends that inform the most highly expressed *flp*-genes at the species-, clade-, and lifestyle-level. Although variability in *flp*-gene expression across species and lifestyles is evident, the data presented here suggest that *flp*-14 represents the most highly expressed *flp*-gene in the nematode species and lifecycle stages examined. Indeed, *flp*-14 represents: (i) the most or second most highly expressed *flp*-gene in 76% of the species examined here; (ii) the most highly expressed *flp*-gene within any individual lifestage examined here (see Supplementary Figure 5). This is interesting given the observations made by McVeigh, Leech (15) who suggested that *flp*-14 may represent one the most highly expressed nematode *flp*-genes, along with *flp*-1 and −11; whilst our data concur with flp-14 observations, they do not suggest that *flp*-1 and −11 are also among the most highly expressed *flp*-genes. These data demonstrate the value and utility of expansions and improvements in RNAseq datasets for multiple species and lifestages (4).

### Pan-phylum analyses provide new insights into the evolution and diversity of nematode *flp*-gene complements that will seed novel drug target discovery pipelines

In this study we generated pan-phylum profiles for FLPs across all available nematode genomes (>100 species) and have identified >2000 putative FLP-encoding genes. These analyses uncover new pan-phylum insights to reveal that: (i) the phylum-spanning *flp*-genes*, flp*-1, −8, −14, and −18, may be representative of the *flp* profile of the LCA of nematodes; (ii) the majority of parasitic nematodes have a reduced *flp* complement relative to free-living species; (iii) FLP prepropeptide architecture is variable within and between *flp*-genes and across nematode species; (iv) FLP prepropeptide signatures facilitate *flp*-gene discrimination; (v) FLP motifs display variable length, amino acid sequence, and conservation; (vi) CLANS analysis provides insight into the evolutionary history of *flp*-gene sequelogues and reveals putative *flp*-gene paralogues and, (viii) *flp* expression is upregulated in the infective larval stage of several nematode parasites.

### Conclusions

This research represents a significant advance in the scale of available nematode FLP neurosignalling data that can be exploited to unravel the complexity of nematode neuropeptide interactomics. Integration of our pan-phylum FLP profiles with equivalent pan-phylum NP-GPCR datasets will seed innovative approaches to the *in silico* prediction of receptor-ligand interactions, deorphanisation, functional biology, and more integrated analysis on the coevolution of neuropeptides and receptor binding pockets. Significantly, this work has enhanced understanding of neuropeptide-signalling systems in therapeutically relevant parasitic nematodes to inform future drug discovery programmes for nematode parasite control.

## Methods

### Datasets analysed

A pan-phylum BLAST-based approach was employed to identify sequelogues of 32 *C. elegans flp*-genes from 108 nematode species (represented by 134 genomes; 4). The genome data mined in this study were derived from all publicly available nematode genome assemblies with predicted protein datasets at the time of study (WormBase ParaSite versions 14-16; see Supplementary Table 1). Note that the quality of genome assembly varies between species as represented by BUSCO score (31; see Supplementary Table 1, 32). In cases where multiple genome assemblies for a single species were available (19 species; see Table S1), the highest quality assembly (based on BUSCO score) was prioritised. If a positive hit was not identified, additional assemblies were then searched until exhausted. For RNAseq analysis, transcriptome data for 13 nematode species were examined in this study; transcriptome datasets were selected based on the number of lifecycle stages/tissues available per species, the number of replicates sequenced per sample (typically ≥3) and the phylogenetic position of each species relative to other available transcriptome datasets.

### FLP-encoding gene sequelogue identification

FLP-encoding gene sequelogues were identified using a BLAST-based approach via WormBase ParaSite (https://parasite.wormbase.org/Multi/Tools/Blast; 4) as previously described (13, 16). Briefly, *C. elegans* FLP prepropeptide sequences for *flp*-1-28 and *flp*-32-34 were employed as BLASTp query sequences; note that *Meloidogyne incognita flp*-31 was employed as a query for *flp*-31 sequelogues because *C. elegans* does not encode *flp-*31. Default BLAST settings were used; expect values were set to >1000 to limit false negative returns. All BLASTp returns were examined for C-terminal FLP motifs flanked by typical mono/dibasic cleavage sites to eliminate false positive hits (15). Negative BLASTp returns were cross-checked using translated nucleotide BLAST (tBLASTn) searches to identify unannotated *flp*-genes. Where tBLASTn revealed a putative *flp*-gene sequelogue, the genome region encoding the sequence (+/- 100 nucleotide base pairs) was translated using the ExPASy translate tool for sequence verification (https://web.expasy.org/translate; 33). All BLASTp and tBLASTn return sequences are provided in Supplementary File 1.

### Post-BLAST FLP signature and motif analysis

FLP prepropeptides were aligned using the MUSCLE alignment tool (34) in Geneious Prime (https://www.geneious.com/) using default settings. Mature FLP peptides were predicted using the relative postions of the signal peptide and mono/dibasic cleavage sites as previously reported (13–16). Sequence alignments were edited to remove sequences flanking the predicted mature neuropeptide. Aligned mature peptide regions were analysed in Jalview (35) to characterise the FLP prepropeptide signature and subsequently the FLP motif across all of the sequelogues for each FLP-encoding gene. To define the FLP prepropeptide signature, only peptides that were present in >50% of species in the MSA were analysed (50% occupancy; see Figure 1). Subsequently each animo residue within each peptide region (that displayed >50% occupancy) was analysed sequentially in the C- to N-terminal direction. Any amino acid in the predicted peptide that displayed >40 % consensus amino acid conservation was included in the FLP prepropeptide signature. To generate a single representative FLP motif for each FLP-encoding gene a MSA of the individual peptides comprising the FLP prepropeptide signature was created (Figure 1). Within each FLP motif individual amino acid residue conservation scores were calculated using the Jalview built in conservation “AACon” calculator tool under the Karlin scoring option (35). Average amino acid conservation scores were also calculated for each individual FLP motif. WebLogo (https://weblogo.berkeley.edu/logo.cgi; 36) was used to produce a graphical representation of amino acid conservation within each FLP signature and motif.

### Clustered Analysis of Sequences (CLANS)

The Clustered Analysis of Sequences (CLANS) algorithm (https://toolkit.tuebingen.mpg.de/#/tools/clans; 37) was used to perform all-against-all BLASTp comparisons between all identified putative FLP-encoding prepropeptide sequences and to generate a three-dimensional similarity matrix. All parameters were set as default, with the exception of the E-value which was set to either 1E-5 or 1E-10 to facilitate sufficient cluster separation as required. A lower E-value (1E-10) was employed to better separate the highly clustered prepropeptide sequences concentrated at the centre of the higher E-value (1E-5) plot. The CLANS file outputs were examined and coloured after 10,000 clustering rounds using the Java-based desktop software.

### RNAseq analysis

Publicly available lifestage and tissue specific transcriptome datasets, representing 13 nematode species, were collated from WormBase ParaSite v14 Gene Expression database (4) and published literature (see Supplementary Table 1). Transcriptome data included metadata, raw counts, transcripts per million (TPM) and DESeq2 differential expression data (in log2foldchange and adjusted p value formats). An expression threshold of 1.5 TPM was applied (38, 39). Datasets were mined using species-specific *flp*-gene IDs generated through BLAST analyses in this study and z-scores calculated from log2averageTPMs; note that expression of unannotated genes could not be analysed. Heatmaps displaying z-scores were generated using GraphPad Prism.

## Availability of data and materials

Data produced in this study are reported in this paper and will be shared by the corresponding authors upon request.

## Authors and Affiliations

School of Biological Sciences, Queen’s University Belfast, 19 Chlorine Gardens, Belfast BT9 5DL, United Kingdom

Ciaran J. McCoy, Christopher Wray, Laura Freeman, Bethany A. Crooks, Louise E. Atkinson, Angela Mousley

Animal Physiology and Neurobiology, Department of Biology, University of Leuven (KU Leuven), Naamsestraat 59, 3000 Leuven, Belgium.

Luca Golinelli, Liesbet Temmerman, Isabel Beets

## Contributions

This study was conceived by AM, LA and CMCC. Investigation including experiments and analyses were performed by CMCC, CW, LF, BC and LG. The manuscript was written by AM, LA and CMCC with contributions from IB, LT, LG and NM. All authors read and approved the final manuscript.

## Supplementary Information

**Supplementary Table 1: Genome and transcriptome datasets used in this study.**

**Supplementary Table 2: Putative *flp*-gene paralogues.**

**Supplementary File 1: FLP prepropeptide sequences identified via BLAST.**

**Supplementary File 2: FLP prepropeptide alignments.** Pages 1-28 represent an individual FLP prepropeptide alignment per page where page 1 represent FLP-1 and page 28 represents FLP-28, and each page between pages 29-32 represent FLP-31-34. Dashed black boxes highlight relatively conserved propeptide regions of FLP-1, −6, −12, −14 and −17 sequelogues.

**Supplementary Figure 1: FLP peptide alignments, signatures and motifs.** Note that only highly conserved aligned peptide regions (>50% occupancy) were used to generate the peptide signatures and motifs.

**Supplementary Figure 2: FLP-6 and FLP-17 prepropeptide alignment.** The conserved prepropeptide motif is highlighted by a red box and the black box (dashed line) highlights where this is not conserved.

**Supplementary Figure 3. AlphaFold derived structures of FLP prepropeptides.** Structures were downloaded (https://alphafold.com) and annotated. Red box indicates signal peptide; black arrows indicate conserved regions of the propeptide that are predicted to form alpha helices; * indicates a truncated predicted protein that does not encode a signal peptide; ** indicates an N-terminal extension of the predicted protein.

**Supplementary Figure 4. Lifestage and tissue-specific *flp*-gene expression profiles.** Heatmaps generated by mapping mean *flp* transcript Z-scores from lifestage and/or tissue-specific RNA-seq libraries. Y-axis denotes *flp*-gene. X-axis denotes lifestages or tissue types. Unannotated genes or those falling below the expression inclusion threshold of 1.5 TPM were omitted. *Trichuris muris*: life stages include second stage larvae (L2), third stage larvae (L3), adult female (AF), adult male (AM), and mixed adult (AMix Sex); tissues include adult female anterior (AF Ant), adult female posterior (AF Post), adult male anterior (AM Ant), adult male posterior (AM Post). *Ascaris suum*: tissues include anterior intestine, female intestine, female pharynx, female head, male head, male intestine, male pharynx, mid intestine, ovary, posterior intestine, seminal vesicle, testis, uterus, whole intestine, whole worm. *Dirofilaria immitis*: tissues include adult female body wall (AF Bodywall), adult female head (AF Head), adult female intestine (AF Intestine), adult female uterus (AF Uterus). *Brugia malayi*: life stages include eggs and embryos, microfilariae (MF), stage three larvae (L3), stage four larvae (L4), adult female (AF) and adult male (AM). *Onchocerca volvulus*: life stages include eggs and embryos, microfilariae (MF), stage two larvae (L2), stage three larvae (L3), adult female (AF), adult male (AM). *Ancylostoma caninum*: life stages include eggs, stage one larvae (L1), stage two larvae (L2), activated stage three larvae (L3 (A)), non-activated stage three larvae (L3 (NA)), untreated stage three larvae (L3 (UT)), adult female and adult male. *Dictyocaulus viviparus*: life stages include egg, stage one larvae (L1), mixed stage one and two larvae (L1+L2), stage two larvae (L2), stage three larvae (L3), mixed stage five larvae and adult females (L5AF), mixed stage five larvae and adult males (L5AM), mixed gender stage five larvae L5(Mixed), mixed stage five larvae and adults L5(Adult), hypobiotic larvae (LHyp), adult female (AF), adult male (AM); *Haemonchus contortus*: life stages include eggs, stage one larvae (L1), stage four larvae (L4), adult female (AF), adult male (AM), adult female gut (AF gut). *Teladorsagia circumcinta*: life stages include stage three larvae (L3), adult female, adult male. *Bursaphelenchus xylophilus*: life stages include mixed embryo, stage two larvae and 3 hour pooled male and females (L2+3h mixed), stage two larvae and 6 hour pooled male and females (L2+6h mixed), stage two larvae and pooled male and females (L2 mixed), stage three dispersal juvenile (S3 DJ), dauer, stage three larvae pooled male and female (L3 mixed), stage four larvae pooled male and female (L4 mixed), adult female, adult male, mixed propagative. *Strongyloides stercoralis*: life stages include post free-living stage one larvae (PFL L1), post parasitic stage one larvae (PP L1), post parasitic stage three larvae (PP L3), activated stage three larvae (L3+), infective stage three larvae (L3i), free-living females (FL Females), parasitic females (P Females). *Globodera pallida*: life stages include egg, stage two juveniles (J2), 7 days post infection (7dpi), 14 days post infection (14dpi), 21 days post infection (21dpi), 28 days post infection (28dpi), 35 days post infection (35dpi) and adult male (AM); *Meloidogyne incognita*: life stages include egg, stage two juvenile larvae (J2), stage three juvenile larvae (J3), stage four juvenile larvae (J4), adult female (AF). All datasets used are detailed in Supplementary Table 1.

**Supplementary Figure 5. Average *flp* transcripts per million values derived from lifestage and tissue specific RNASeq datasets.** Each data point (grey circle) represents a specific *flp* TPM for a lifestage or tissue. All datasets used are detailed in Supplementary Table 1.

## References

1. Holterman M, van der Wurff A, van den Elsen S, van Megen H, Bongers T, Holovachov O, et al. Phylum-wide analysis of SSU rDNA reveals deep phylogenetic relationships among nematodes and accelerated evolution toward crown Clades. Mol Biol Evol. 2006;23(9):1792–800.

2. Mendoza-de Gives P. Soil-Borne Nematodes: Impact in Agriculture and Livestock and Sustainable Strategies of Prevention and Control with Special Reference to the Use of Nematode Natural Enemies. Pathogens. 2022;11(6).

3. Hotez PJ, Lo NC. Neglected tropical diseases: public health control programs and mass drug administration. Hunter’s Tropical Medicine and Emerging Infectious Diseases: Elsevier; 2020. p. 209–13.

4. Howe KL, Bolt BJ, Shafie M, Kersey P, Berriman M. WormBase ParaSite - a comprehensive resource for helminth genomics. Mol Biochem Parasitol. 2017;215:2–10.

5. Kaplan RM, Vidyashankar AN. An inconvenient truth: global worming and anthelmintic resistance. Vet Parasitol. 2012;186(1-2):70–8.

6. McVeigh P, Atkinson L, Marks NJ, Mousley A, Dalzell JJ, Sluder A, et al. Parasite neuropeptide biology: Seeding rational drug target selection? Int J Parasitol Drugs Drug Resist. 2012;2:76–91.

7. Maule AG, Mousley A, Marks NJ, Day TA, Thompson DP, Geary TG, et al. Neuropeptide signaling systems - potential drug targets for parasite and pest control. Curr Top Med Chem. 2002;2(7):733–58.

8. Beets I, Zels S, Vandewyer E, Demeulemeester J, Caers J, Baytemur E, et al. System-wide mapping of peptide-GPCR interactions in C. elegans. Cell Rep. 2023;42(9):113058.

9. Gershkovich MM, Groß VE, Kaiser A, Prömel S. Pharmacological and functional similarities of the human neuropeptide Y system in C. elegans challenges phylogenetic views on the FLP/NPR system. Cell Communication and Signaling. 2019;17(1):123.

10. Anderson RC, Newton CL, Millar RP, Katz AA. The Brugia malayi neuropeptide receptor-4 is activated by FMRFamide-like peptides and signals via Galphai. Mol Biochem Parasitol. 2014;195(1):54–8.

11. Atkinson LE, Stevenson M, McCoy CJ, Marks NJ, Fleming C, Zamanian M, et al. flp-32 Ligand/Receptor Silencing Phenocopy Faster Plant Pathogenic Nematodes. Plos Pathogens. 2013;9(2).

12. Atkinson LE, McCoy CJ, Crooks BA, McKay FM, McVeigh P, McKenzie D, et al. Phylum-Spanning Neuropeptide GPCR Identification and Prioritization: Shaping Drug Target Discovery Pipelines for Nematode Parasite Control. Frontiers in Endocrinology. 2021;12.

13. McCoy CJ, Atkinson LE, Zamanian M, McVeigh P, Day TA, Kimber MJ, et al. New insights into the FLPergic complements of parasitic nematodes: Informing deorphanisation approaches. EuPA Open Proteom. 2014;3:262–72.

14. McVeigh P, Alexander-Bowman S, Veal E, Mousley A, Marks NJ, Maule AG. Neuropeptide-like protein diversity in phylum Nematoda. Int J Parasitol. 2008;38(13):1493–503.

15. McVeigh P, Leech S, Mair GR, Marks NJ, Geary TG, Maule AG. Analysis of FMRFamide-like peptide (FLP) diversity in phylum Nematoda. Int J Parasitol. 2005;35(10):1043–60.

16. McKay FM, McCoy CJ, Marks NJ, Maule AG, Atkinson LE, Mousley A. In silico analyses of neuropeptide-like protein (NLP) profiles in parasitic nematodes. bioRxiv. 2021:2021.03.03.433794.

17. Coghlan A, Tyagi R, Cotton JA, Holroyd N, Rosa BA, Tsai IJ, et al. Comparative genomics of the major parasitic worms. Nature Genetics. 2019;51(1):163-+.

18. Peymen K, Watteyne J, Frooninckx L, Schoofs L, Beets I. The FMRFamide-like peptide family in nematodes. Frontiers in Endocrinology. 2014;5.

19. van Megen H, van den Elsen S, Holterman M, Karssen G, Mooyman P, Bongers T, et al. A phylogenetic tree of nematodes based on about 1200 full-length small subunit ribosomal DNA sequences. Nematology. 2009;11:927–S27.

20. Blaxter M, Koutsovoulos G. The evolution of parasitism in Nematoda. Parasitology. 2015;142 Suppl 1(Suppl 1):S26–39.

21. Fadda M, De Fruyt N, Borghgraef C, Watteyne J, Peymen K, Vandewyer E, et al. NPY/NPF-related neuropeptide FLP-34 signals from serotonergic neurons to modulate aversive olfactory learning in Caenorhabditis elegans. Journal of Neuroscience. 2020;40(31):6018–34.

22. Schafer W. Nematode nervous systems. Curr Biol. 2016;26(20):R955–R9.

23. Han Z, Boas S, Schroeder NE. Unexpected Variation in Neuroanatomy among Diverse Nematode Species. Front Neuroanat. 2016;9:162.

24. Valdar WS. Scoring residue conservation. Proteins. 2002;48(2):227–41.

25. Mousley A, Novozhilova E, Kimber MJ, Day TA, Maule AG. Neuropeptide Physiology in Helminths. In: Geary TG, Maule AG, editors. Neuropeptide Systems as Targets for Parasite and Pest Control. Boston, MA: Springer US; 2010. p. 78–97.

26. Moffett CL, Beckett AM, Mousley A, Geary TG, Marks NJ, Halton DW, et al. The ovijector of Ascaris suum: multiple response types revealed by Caenorhabditis elegans FMRFamide-related peptides. International Journal for Parasitology. 2003;33(8):859–76.

27. Dirndorfer D, Seidel RP, Nimrod G, Miesbauer M, Ben-Tal N, Engelhard M, et al. The α-helical structure of prodomains promotes translocation of intrinsically disordered neuropeptide hormones into the endoplasmic reticulum. J Biol Chem. 2013;288(20):13961–73.

28. Garcia AL, Han SK, Janssen WG, Khaing ZZ, Ito T, Glucksman MJ, et al. A prohormone convertase cleavage site within a predicted alpha-helix mediates sorting of the neuronal and endocrine polypeptide VGF into the regulated secretory pathway. J Biol Chem. 2005;280(50):41595–608.

29. Jékely G. Global view of the evolution and diversity of metazoan neuropeptide signaling. Proc Natl Acad Sci U S A. 2013;110(21):8702–7.

30. Lee JS, Shih PY, Schaedel ON, Quintero-Cadena P, Rogers AK, Sternberg PW. FMRFamide-like peptides expand the behavioral repertoire of a densely connected nervous system. Proc Natl Acad Sci U S A. 2017;114(50):E10726–e35.

31. Simao FA, Waterhouse RM, Ioannidis P, Kriventseva EV, Zdobnov EM. BUSCO: assessing genome assembly and annotation completeness with single-copy orthologs. Bioinformatics. 2015;31(19):3210–2.

32. Parra G, Bradnam K, Korf I. CEGMA: a pipeline to accurately annotate core genes in eukaryotic genomes. Bioinformatics. 2007;23(9):1061–7.

33. Gasteiger E, Gattiker A, Hoogland C, Ivanyi I, Appel RD, Bairoch A. ExPASy: The proteomics server for in-depth protein knowledge and analysis. Nucleic acids research. 2003;31(13):3784–8.

34. Edgar RC. MUSCLE: a multiple sequence alignment method with reduced time and space complexity. BMC Bioinformatics. 2004;5(1):113.

35. Waterhouse AM, Procter JB, Martin DMA, Clamp M, Barton GJ. Jalview Version 2—a multiple sequence alignment editor and analysis workbench. Bioinformatics. 2009;25(9):1189–91.

36. Crooks GE, Hon G, Chandonia JM, Brenner SE. WebLogo: A sequence logo generator. Genome Research. 2004;14(6):1188–90.

37. Frickey T, Lupas A. CLANS: a Java application for visualizing protein families based on pairwise similarity. Bioinformatics. 2004;20(18):3702–4.

38. Wagner GP, Kin K, Lynch VJ. A model based criterion for gene expression calls using RNA-seq data. Theory Biosci. 2013;132(3):159–64.

39. Soneson C, Robinson MD. Bias, robustness and scalability in single-cell differential expression analysis. Nature Methods. 2018;15(4):255-+.

