## Supplementary File 1 for "Exploitation of phylum-spanning omics resources reveals complexity in the nematode FLP signalling system and provides insights into *flp*-gene evolution"

```
>FLP-1_Romanomermis_culicivorax_FLP_1_nRc.2.0.1.t41952_RA
MGVSSTLLAATVLFFASTILTRRVVNGECCENDETSVFCQSYSKKTMDEKAKIRSILGEN
CQLLVDIPQKFSTDVRPNFIRFGRPSAMSPNFVRFGRSALTPNFVRFGRQQLPNFVRFGK
RQLEIEDDIDQNYDENHDDNDEESFRRLTRKANFIRFGKRRSFSGPNFIRFGRNGMNPNF
IRFG
>FLP-1_Soboliphyme_baturini_FLP_1_SBAD_0001333901_mRNA_1
MRKPNFIRFGRADPWSETRPNFLRFGRSRDTHQPEQFFRLTRNSEDDRRPNFIRFGKSGK
PDPFEMSDRKPNIIRFGRSVNPKFQQYERKESIEAPAKGFAA
>FLP-1_Trichinella_britovi_FLP_1_T03_6970.1
MTVIWKIRSDVSTSANIRMLHLERTYKREHSVNEITDVEETANEEAGGYGYWSPQQANKW
FVNCSFQLRLNHHFPVDHHHNAEIMTIFIPTSVLSKNEDSSGLQTIGVYDDGEYSDDYFE
LTKRDLEPVYTDLRSVPSFLRFGRNMNNPLMRSVLEPNFIRFSRNSPNFLRFGRAAAANF
LRFGRASPNFLRFGKANPNFLRFGRDSDLNRQYRAAPNFLRFG
>FLP-1_Trichinella_murrelli_FLP_1_T05_2477.1
MTVIWKIRSDVSTSANIRMLHLERTYKREHSVNEITDVEETANEEAGGYGYWSPQQANKW
FVNCSFQLRLNHHFPVDHHHNAEIMTIFIPTSVLSKNEDSSGLQTIGVYDDGEYSDDYFE
LTKRDLEPVYTDLRSVPSFLRFGRNMNNPLMRSVLEPNFIRFSRNSPNFLRFGRAAAANF
LRFGRASPNFLRFGKANPNFLRFGRDSDLNRQYRAAPNFLRFG
>FLP-1_Trichinella_nativa_FLP_1_T02_7924.1
MTVIWKIRSDVSTSANIRMLHLERTYKREHSVNEITDVEETANEEAGGYGYWSPQQANKW
FVNCSFQLRLNHHFPVDHHHNAEIMTIFIPTSVLSKNEDSSGLQTIGVYDDGEYSDDYFE
LTKRDLEPVYTDLRSVPSFLRFGRNMNNPLMRSVLEPNFIRFSRNSPNFLRFGRAAAANF
LRFGRASPNFLRFGKANPNFLRFGRDSDLNRQYRAAPNFLRFG
>FLP-1_Trichinella_nelsoni_FLP_1_T07_5623.1
MAFIPKLITLLSITVTIFIPTSVLSKNEDSSGLQTIGVYDDGEYSDDYFELTKRDLEPVY
TDLRSVPSFLRFGRNMNNPLMRSVLEPNFIRFSRNSPNFLRFGRAAAANFLRFGRASPNF
LRFGKANPNFLRFGRDSDLNRQYRAAPNFLRFG
>FLP-1_Trichinella_papuae_FLP_1_T10_8944.1
MTVICKIRSDLSTSAIKYECYILSERISASTVLMKLQMWRKPPTRRRWTLAIGHHSRLIS
GLSTVPFNCDITTIFLSIVIITLKSMTICIPTSVLSKTEDSSGFETIGVYDDGEYLDDYF
ELTKRDLEPIYTDLRSVPSFLRFGRSMNNPFMRSVLEPNFIRFGRNSPNFLRFGRAAAAN
FLRFGRASPNFLRFGKASPNFLRFGRDLDLNRQYRAAPNFLRFG
>FLP-1_Trichinella_patagoniensis_FLP_1_T12_993.1
MTVIWKIRSDVSTSANIRMLHLERTYKREHSVNEITDVEETANEEAGGYGYWSPQQANKW
FVNCSFQLRLNHHFPVDHHHNAEIMTIFIPTSVLSKNEDSSGLQTIGVYDDGEYSDDYFE
LTKRDLEPVYTDLRSVPSFLRFGRNMNNPLMRSVLEPNFIRFSRNSPNFLRFGRAAAANF
LRFGRASPNFLRFGKANPNFLRFGRDSDLNRQYRAAPNFLRFG
>FLP-1_Trichinella_pseudospiralis_T4A_12377.1
SVNEITDVEETANEEAVDSGNWSPQQANKCKTEDSSGFETIGVYDDGDYLDDYYELTKRD
LEPIYTDLRSVPSFLRFGRSMNNPFMRSVLEPNFIRFGRNSPNFLRFGRAAAANFLRFGR
ASPNFLRFGKASPNFLRFGRDSDLNRQYRAAPNFLRFG
>FLP-1_Trichinella_sp.T6_FLP_1_T06_832.1
MTVIWKIRSDVSTSANIRMLHLERTYKREHSVNEITDVEETANEEAGGYGYWSPQQANKW
FVNCSFQLRLNHHFPVDHHHNAEIMTIFIPTSVLSKNEDSSGLQTIGVYDDGEYSDDYFE
LTKRDLEPVYTDLRSVPSFLRFGRNMNNPLMRSVLEPNFIRFSRNSPNFLRFGRAAAANF
LRFGRASPNFLRFGKANPNFLRFGRDSDLNRQYRAAPNFLRFG
>FLP-1_Trichinella_sp.T8_FLP_1_T08_7337.1
MTVIWKIRSDVSTSANIRMLHLERTYKREHSVNEITDVEETANEEAGGYGYWSPQQANKW
FVNCSFQLRLNHHFPVDHHHNAEIMTIFIPTSVLSKNEDSSGLQTIGVYDDGEYSDDYFE
LTKRDLEPVYTDLRSVPSFLRFGRNMNNPLMRSVLEPNFIRFSRNSPNFLRFGRAAAANF
LRFGRASPNFLRFGKANPNFLRFGRDSDLNRQYRAAPNFLRFG
>FLP-1_Trichinella_sp.T9_FLP_1_T09_2530.1
MTVIWKIRSDVSTSANIRMLHLERTYKREHSVNEITDVEETANEEAGGYGYWSPQQANKW
FVNCSFQLRLNHHFPVDHHHNAEIMTIFIPTSVLSKNEDSSGLQTIGVYDDGEYSDDYFE
```

LTKRDLEPVYTDLRSVPSFLRFGRNMNNPLMRSVLEPNFIRFSRNSPNFLRFGRAAAANF
LRFGRASPNFLRFGKANPNFLRFGRDSDLNRQYRAAPNFLRFG
>FLP-1_Trichinella_zimbabwensis_FLP_1_T11_12361.1
MTICIPTSVLSKTEDSSGFETIGVYDDGEYLDDYFELTKRDLEPIYTDLRSVPSFLRFGR
SMNNPFMRSVLEPNFIRFGRNSPNFLRFGRAAAANFLRFGRASPNFLRFGKASPNFLRFG
RDLDLNRQYRAAPNFLRFG
>FLP-1_Trichuris_muris_FLP_1_TMUE_3000013417.1
MEDAGRLVMFPEEEDLIEEYPDISEADDEWYARIQRAGKNGEDWISKKRVNFLRFGRKMT
NPNFLRFGRAPNSPNFLRFGRLAGNPNFLRFGRQPYSNRQDRAVGNFLRFG
>FLP-1_Trichuris_suis_FLP_1_D918_08076
MVNSSGMTLPAKGMDDAGALLAVPEEEDLFDGTLLICFGDFEIRTAYAFAEYADISGTDD
ELYPRMQRSGRSDEDWLSKKRVNFLRFGRKMSNPNFLRFGRASNSPNFLRFGRRAVNPNF
LRFGRKLYSNRQDRAVGNFLRFG
>FLP-1_Trichuris_trichiura_FLP_1_TTRE_0000091401_mRNA_1
MIKQYAVGSFLLLQCIILLSGVHSKIENTQMVNSSGITLPAKAMDDAGTLLVLPEEDDFF
DGPTFTFRKEVAFRISFFARCRVSKKRVNFLRFGRKTSNPNFLRFGRAPNSPNFLRFGRA
PNSPNFLRFGRQIGNPNFLRFGRKPYSNRQDRAVGNFLRFG
>FLP-1_Tricinella_spiralis_T01_3205.1
MAFIPKLITLLSITVTIFIPTSVLSKNEDSSGLQTIGVYDDGEYLDDYFELTKRDLEPVY
TDLRSVPSFLRFGRNMNNPVMRSVLEPNFIRFGRNSPNFLRFGRAAAANFLRFGRASPNF
LRFGKANPNFLRFGRDSDLNRHYRAAPNFLRFG
>FLP-1_Plectus_sambesii_PSAMB.scaffold93size81355.g1693.t1
MAKVGCVLVQSTLLVAWILAVSLTQVDAECCENEDADNESAFCTVYNILSVPEKAEVRLY
LGENCEGDADEAMNKMEKRKPNFIRFGRSGPNFLRFGKRGADPNFLRFGRASDPNFLRFG
KKDRNFLRFGRNTPNFLRFGRSSDPNFLRFGKPDPNFLRFGKSAPDSDPNFLRFGKRSDS
DPKFLRFGKRKPNFLRFGRSADDLERFDREYRKPNFLRFG
>FLP-1_Acanthocheilonema_viteae_nAv.1.0.1.t00761-RA
MLSSSQQAEIRQHFRDDCQGDADEGAKKPEKRKPNFIRFGRTASIMYGKKDADPNFLQFE
RLSSAFGGQNFFRFERAAEPNFLRFGRVTDPNFLRFGKSTEPNFLRFGKRFEMRDPNFLR
FGRNNSLQPSQEHKERFNREDRKPNFLRFGK
>FLP-1_Ascaris_lumbricoides_FLP_1_ALUE_0002073601_mRNA_1
MMRQSHSVMSPLHVALFLIFCSSQVLGECCNDGQTSDFCAVFNMLSPTEQAEVRSYLGDN
CDGDADEAVRKIEKRKPNFIRFGRTAPPLTFGKKGSDPNFLRFGRTPSNNFLRFGKSNQA
QNFLRFGRNAEPNFLRFGRPADPNFLRFGKSAEPNFLRFGKRSDIGISEPNFLRFGRNNF
LRFGRNDQFDREYRKPNFLRFGK
>FLP-1_Ascaris_suum_FLP_1_GS_07036
MINRFAAPPLTFGKKGSDPNFLRFGRTPSNNFLRFGKSNQAQNFLRFGRNAEPNFLRFAD
PNFLRFGKSAEPNFLRFGKRSDIGISEPNFLRFGRNNFLRFGRNDQFDREYRKPNFLRFG
K
>FLP-1_Brugia_malayi_Bm3533.1
MVLIHAILLICSLTQVSSECCRNGITSDYCIIFNMLSSSQQAEIRQYFGHDCQDVDEATR
KIEKRKPNFIRFGRTAPPAMYGKKDVDPKFLQFGHSSSAFIPSGQNFLRFGRAAEPNFLH
LGRVTDPNFLRFGKSAEPNFLRFGKRTEVGDPNFLRFGKNSSSQSTPDNEGFSRQDRKPN
FLRFGK
>FLP-1_Brugia_pahangi_BPAG_0001004901-mRNA-1
MVLIHAILLICSLTQRLAYTHVTLYISHVRKFAFPLLQFLIPYVSGVDVSSECCRNGITS
DYCIIFNMLSSSQQAEIRQYFRHDCQDVDEATRKIEKRKPNFIRFGRTAPPAMHGKKDVD
PKFLQFGHSSSAFTPSGQNFLRFGRAAEPNFLHLGRVTDPNFLRFGKSAEPNFLRFGKRT
EVGDPNFLRFGKNSSFQSTPEYNEGFSRQDRKPNFLRFGK
>FLP-1_Brugia_timori_BTMF_0001517201-mRNA-1
MLSSSQQAEIRQYFGHDCQDVDEATRKIEKRKPNFIRFGRTAPPAMYGKKDVDPKFLQFG
HSSSAFIPSGQNFLRFGRAAEPNFLHLGRVTDPNFLRFGKSAEPNFLRFGKRTEVGDPNF
LRFGKN
>FLP-1_Dirofilaria_immitis_nDi.2.2.2.t01041

MVLVHAVLLICSLTQVSSECCRNAITLDFCTMFNMLSSSQQAEVRQHFGDNCDEDANEAS
KKIEKRKPNFIRFGRDASSIMYGKKDNDPNILQFGRSSSIFTPGGQNFLRFGRAVEPNFL
RFGRVIDPKFLRSGKSAEPNFLRFGKRIEVNEPNFLRLGRNNFLQTDQKYNEGFSRQNRK
PNFLRFADYCTARSSAKDVPDDMGMQILKILVWSQGRPPAVYTSASII
>FLP-1_Dracunculus_medinensis_DME_0001006201-mRNA-1
LLPIKKQEEINRFMGNICEQHPSNKMLKLMENGVKIKRKPNFMRFGRSGRPNRITSYFFG
PMNLIKGRQIDPIFLRFGKSAEPNFLRFGKRSQERNFFVRYGDELDRESRRPNFLRFGK
>FLP-1_Elaeophora_elaphi_EEL_0000075801-mRNA-1
MVLVSSECCRNAITSDFCTIYNMLSSSQQAEIRQHFGDDCQGDADEADTKIEKRKPNFIR
FGRTDADPKFLQFGRSSSAFAPSGQNFLRFGRASEPNFLRFGRVTDPNFLRFGKSAEPNF
LRFGKRIEVRDPNFLRFGRNSSLQPNGEYNEGFSREGRKPNFLRFGK
>FLP-1_Enterobius_vermicularis_EVEC_0000490501-mRNA-1
MRLLYVAPLLILYFSQVHGECCYEGNESLFCKVFQSASKAKQDAIRYYFNDDCSELENEF
MQELANAKDLSVIPITKEAALDSQYDPGYDENVRLNRRAVIASKKDYYNTENKMRNFLRF
GKRADGTFYRYNRDGQTLAFLRDVKSAHGSKPNFLRFGKRQFTRQGASGIPFNPALIPDH
KGGPNFLRFG
>FLP-1_Gongylonema_pulchrum_GPUH_0000186401-mRNA-1
MSGLNNSKRKNFEVSSDCCTGGSTSDLCQLLKLLSSSEQAEPRQYLGEDCEEYASEPAGK
TEKRKPNFIRFGRAAASPTVFREKGVDPNFLRFGRASGAFFSHGTERSWIGSSLLFSRTM
DPNFLRFGKSTEPNFLRFGRRMELSEPNFLRFGREPNEEFNRQYRKPNFLRFG
>FLP-1_Litomosoides_sigmodontis_FLP-3_nLs.2.1.2.t04733-RA
MSSQIENFNTGKLLGRGRKCVCHLMNGHTEHRSGKWDQQELYRSIMQALEGRCSTRSNLA
KEFEIQIREFTATLECHLVDLLRKRLEELNEPDRLLISRLNRALCELGDSNSSVLSRPPY
HMLSPNRPAHRVWTMCTKDPERLRNLVSEVCHMANIACDNGDEQNSCQIKFSAALPSAWN
AADCAKYLIEKPEIEDGLTGQRTQHFGSCRSVKNGFKLSLTCVSGVIHHFVVSEIVKDLS
KLYAQSFSLMKSASVTLERAATSLPTLISADDPAGCSENDPSIKSRPFSELAKEKSNPDK
MSDKQLSSEENRTVIPLEPSVFVREMKKQSFRESSFTDESDSACALKEEAFNSTSPSAAS
SIPLCQTDSSYDSRDMQAPLFEQTMSSCNLSTSTTVTLGNLSTEPNFLRFGERLFSRPID
IISLPNIFTKKQLLAVSDSCGGKLSQDCLGVYIVTCKGEVLHHLPTRDGSASSLTVDPTN
WRLLVSVMHSKGRSIYAFDITNRFKKVEVIPCPKEPKIELSRTRWITVSPRGELFAVSGD
NNRSAIWMYNHSRKGWKILKESRKTRYQYLQVAEDQAEYKAVVLLTCDAAQNRILLFVAD
HTGTLINEYDLTKTYKLYEYIRNPASAIVDSLGNLLVLDYATSRLWALLSNTKGMRHVKE
IIIPNPLGPQEALGIAAHGDWIYMTCFARREVICVRYLRNGVLALPCTTTLESPKSDRRA
TSLPRPTRKPNQV
>FLP-1_Onchocerca_flexuosa_X798_01891
MFNMLSSSQQEEIRRYFVDDCEGDADEAARKIEKRKPNFIGFGRAASPIMYGKKDTDPNF
LQFERSSSAFTPSGQNFFRFGRAAEPNFLRFGRVKDSNFLRFDKSAEPNFLRFGKRIETS
KPNFLRFGRNVSSQNDQEYREGFSRQDRKPNFLRFGK
>FLP-1_Onchocerca_ochengi_OOCN_0000477801-mRNA-1
MFNMLSSSQQAEIRQYFVDDCEGDADEAPKKIEKKKPNFIRFGRAASPIMHGKKDTDPNF
LQFERSSSAFTPSGQNFLRFGRAAEPNFLRFGRVRDSNFLRFGKSAEPNFLRFGKRIEAN
EPNFLQFGRNASLQNDQEYREGFSRQDRKPNFLRFALWGPMNDKWWQSFSFQFSLRPR
>FLP-1_Onchocerca_volvulus_OVOC4697.1
LKKKHYSDMIILVHAVLLICSLTQISSECCRNAITSDFCTMFNMLSSSQQAEIRQYFVDD
CEGDADEAPKKIEKKKPNFIRFGRAASPIMHGKKDTDPNFLQFERSSSAFTPSGQNFLRF
GRAAEPNFLRFGRVRDSNFLRFGKSAEPNFLRFGKRIEANEPNFLQFGRNASLQNDQEYR
EGFSRQDRKPNFLRFGK
>FLP-1_Parascaris_univalens_PgR028_g030_t01
PLYVALFLIFCSSQVLSECCNDGQTSDFCAVFNILSPTEQAEVRSYLGDNCDGDANEAAK
KIEKRKPNFIRFGRTAPPLTFGKKGSDPNFLRFGRTPSNNFLRFGKSNQAQNFLRFGRNA
EPNFLRFGRLDDPNFLRFGKSAEPNFLRFGKRSEIGINEPNFLRFGRNDQFDREYRKPNF
LRFGK
>FLP-1_Syphacia_muris_SMUV_0000682001-mRNA-1
MQLNISASAFPNATVQSECCYGRSDSAFCKLFQTVSKSEQDAIRYLFKDDCSELEAEYEQ

KEANSAKEVAPITMLIKETIPEVQYDTANNDYIPLNIRSAIATKNDYYTENKGTRNFLRF
GKRDDINFYRYNRDRQASGFLRDVKSAHSSKPNFLRFGKRQFSRRSPFSPSLIRDQKGGP
NFLRFG
>FLP-1_Thelazia_callipaeda_TCLT_0000327701-mRNA-1
MVPVHVTLLIWSLIQVSHGCCMSRSTSDFCSIFNMLSDSEQAEVEQYLGKACKDNAVDDA
IRLVEKRKPNFIRFGRAMLSLNNDRNNINPNFLRFGRIPDSVLGSDSGQNFLRFERETNE
PNFLRFGELSTANVDDEINAWK
>FLP-1_Toxocara_canis_Tcan_16747.1
MSPLHAAILLICCASQVLSECCNDGQTSDFCTVFNMLSPTEQAEVRLYLGEDCEGDVDEA
VKKMEKRKPNFIRFGRAGPLGKKGSDPNFLRFGRTPSNNFLRFGKSNQAQNFLRFGRSAE
PNFLRFGRIADPNFLRFGKSAEPNFLRFGKRSDLGMSEPNFLRFGRNNFLRFGRNENDQF
DREYRKPNFLRFGK
>FLP-1_Wuchereria_bancrofti_maker-PairedContig_3872-snap-gene-2.14-
mRNA-1
MVLIHAILLICSLTQVSSECCRNGITSDYCIIFNMLSSSQQAEIRQYFGHDCQDVDEATR
KIEKRKPNFIRFGRTALPIMYGKKDADPKFLQFGHSSSAFTPSGQNFLRFGREAEPNFLH
FGRVTDPNFLRFGKSAEPNFLRFGKRTEVGDPNFLRFGKNSSFQPTPEYNEGFSRQDRKP
NFLRFGK
>FLP-1_Anisakis_simplex_predicted
ELPCSFLTIKTTNRTAESRIDHYILHSFLGRSADPNFLRFGKSAEPNFLRFGKRPDASMSEPNFLRFG
RNNFLRFGRDNNMMFDREYREPNFLRFGK
>FLP-1_Loa_loa_predicted
KKYSVPKKSGIYFTTRNTISYLHKTNVTFSAPAVVYEKNDVDPKFLQFGHSSSAFTPGRQNFLRFGRA
AEPNFLRFGKTVLPCFCNNSQ
VSSECCKNRITSDFCTIFNMLSSSQQAEIRQYFGNDCQGDTDETARKTEKRKPNFIRFGRSGIFI
>FLP-1_Ancylostoma_caninum_FLP_1_ANCCAN_01462
MPTLLQVGLLLALGAVAQVSAECCSPSDQSDFCMVFNMLSPMEQAEVMSYLGSRHQTGPA
IGYPALGVEGSIDPKRDWNCNPRDACNGDADEALRLIEKRKPNFMRYGRSITFGKKGSDP
NFLRFGRNQPNFLRFGKAAGDPNFLRFGRASADPNFLRFGKRAVDPNFLRFGRKPNFLRF
GK
>FLP-1_Ancylostoma_ceylanicum_FLP_1_Acey_s0394.g621.t1
MPTLLQVGLLLALGAVAQVSAECCSPGDQSDFCMVFNMLSPMEQAEVMSYLGDACNGDAD
EALRLIEKRKPNFMRYGRSITFGKKGSDPNFLRFGRNQPNFLRFGKAAGDPNFLRFGRAS
ADPNFLRFGKRAVDPNFLRFGRKPNFLRFGK
>FLP-1_Ancylostoma_duodenale_ANCDUO_17936
MEKRKPNFMRYGRSITFGKKGSDPNFLRFGRNQPNFLRFGKAAGRASADPNFLRFGKRAV
DPNFLRFGRKPNFLRFGK
>FLP-1_Angiostrongylus_cantonensis_FLP_1_ACAC_0000841601-mRNA-1
MPTLLRAGLLLAVGVSTECCTSGEQSDFCLVYNMLAPVEQAEVMSYLGGTCSGDADEALR
LMEKRKPNFMRFGRSPSFGKKGSDPNFLRFGRSHPNFLRFGKAADNPNFLRFGTNVLGRA
GADPNFLRFGKRPADPNFLRFGRKPNFLRFGK
>FLP-1_Angiostrongylus_costaricensis_FLP_1_ACOC_0001187301-mRNA1
MPTLLRAGLLLALDYFKVSTECCTSGEQSDFCLVYNMLAPVEQAEVMSYLGGTCSGDADE
ALRLMEKRKPNFMRFGRSKLLSLSFGKKGSDPNFLRFGRSHPNFLRFGKAADNPNFLRFG
RAGADPNFLRFGKRPADPNFLRFGRKPNFLRFGR
>FLP-1_Brugia_pahangi_BPAG_0001004901-mRNA-1
MVLIHAILLICSLTQRLAYTHVTLYISHVRKFAFPLLQFLIPYVSGVDVSSECCRNGITS
DYCIIFNMLSSSQQAEIRQYFRHDCQDVDEATRKIEKRKPNFIRFGRTAPPAMHGKKDVD
PKFLQFGHSSSAFTPSGQNFLRFGRAAEPNFLHLGRVTDPNFLRFGKSAEPNFLRFGKRT
EVGDPNFLRFGKNSSFQSTPEYNEGFSRQDRKPNFLRFGK
>FLP-1_Caenorhabditis_angaria_Cang_2012_03_13_00336.g9400.t1
MTLLYQVGLLVAVVATVSAGCCAPGTQSDFCTVFVMLSPTEQNEVVNYIGENCDGDADVA
LQKMEKRKPNFMRYGRSADPNFLRFGRSQPNFLRFGKAAGGDPNFLRFGRSDPNFLRFGK
AAADPNFLRFGKRSADPNFLRFGRSFENFDRESRKPNFLRFGK

>FLP-1_Caenorhabditis_brenneri_CBN10439.1
MKSVPRNRKLFPGENCEGDAEVALQKMEKRKPNFMRYGRSAAVKSLGKKAGSDPNFLRFG
RSQPNFLRFGKASGDPNFLRFGRSDPNFLRFGKAAADPNFLRFGKRSADPNFLRFGRSFE
NFDRESRKPNFLRFGK
>FLP-1_Caenorhabditis_briggsae_CBG21782a.1
MTLLYQVGLLLLVAASFKVSAECCTPGATSDFCTVFSMLSTMEQNEVMSYLGENCEGDAD
VALQKMEKRKPNFMRYGRSAAVKSLGKKAGSDPNFLRFGRSQPNFLRFGKASGDPNFLRF
GRSDPNFLRFGKAAADPNFLRFGKRSADPNFLRFGRSSFDNFDRESRKPNFLRFGK
>FLP-1_Caenorhabditis_elegans_Isoform_A_F23B2.5a.1
MTLLYQVGLLLLVAATYKVSAECCTPGATSDFCTVFSMLSTMEQNEVMNFIGENCDGDAEVALQKMEK
RKPNFMRYGRSA
AVKSLGKKAGSDPNFLRFGRSQPNFLRFGKASGDPNFLRFGRSDPNFLRFGKAAADPNFLRFGKRSAD
PNFLRFGRSFDN
FDRESRKPNFLRFGK
>FLP-1_Caenorhabditis_inopinata_Sp34_40353000.t1
MTLLFQVGLLILVAATYKVSAECCTPGATSDFCTVFSMLSTMEQNEVMSYLGENCDGNAE
AALQKMEKRKPNFMRYGRSADPNFLRFGRSQPNFLRFGKASGDPNFLRFGRSDPNFLRFG
KAAADPNFLRFGKRSADPNFLRFGRSFDNFDRESRKPNFLRFGNETEQNGYERNRGYAAL
TNAVAMETGERCYAPVCESVSIGKAR
>FLP-1_Caenorhabditis_japonica_CJA00035.1
MPLLYQVGLLLLATATYMVSAECCAPGATSDFCTVFSMLSTMEQNEVMSYLGENCDGDAE
VALNTMEKRKPNFMRYGRSTVALGKKASSDPNFLRFGRSQPNFLRFGKASGDPNFLRFGR
SDPNFLRFGKAAADPNFLRFGKRSADPNFLRFGRSFDNFDRESRKPNFLRFGK
>FLP-1_Caenorhabditis_latens_FL83_16558
MTLLYQVGLLLLVAATYKVSAECCTPGATSDFCTVFSMLSTMEQNEVMSYLGENCEGDAE
VALQKMEKRKPNFMRYGRSAAVKSLGKKAGSDPNFLRFGRSQPNFLRFGKASGDPNFLRF
GRSDPNFLRFGKAAADPNFLRFGKRSADPNFLRFGRSFDNFDRESRKPNFLRFGK
>FLP-1_Caenorhabditis_nigoni
MTLLYQVGLLLLVAASFKVSAECCTPGATSDFCTVFSMLSTMEQNEVMSYLGENCEGDAD
VALQKMEKRKPNFMRYGRSAAVKSLGKKAGSDPNFLRFGRSQPNFLRFGKASGDPNFLRF
GRSDPNFLRFGKAAADPNFLRFGKRSADPNFLRFGRSSFDNFDRESRKPNFLRFGK
>FLP-1_Caenorhabditis_remanei_FL81_05841
MTLLYQVGLLLLVAATYKVSAECCTPGATSDFCTVFSMLSTMEQNEVMSYLGENCEGDAE
VALQKMEKRKPNFMRYGRSAAVKSLGKKAGSDPNFLRFGRSQPNFLRFGKASGDPNFLRF
GRSDPNFLRFGKAAADPNFLRFGKRSADPNFLRFGRSFDNFDRESRKPNFLRFGK
>FLP-1_Caenorhabditis_sinica_Csp5_scaffold_00466.g11740.t1
MTLLYQVGLLLLVAATYKVSAECCTPGATSDFCTVFSMLSTMEQNEVMSYLGENCEGDAE
VALQKMEKRKPNFMRYGRSAAVKSLGKKAGSDPNFLRFGRSQPNFLRFGKASGDPNFLRF
VYLQTKIPVIKLVSRSDPNFLRFGKAAADPNFLRFGKRSADPNFLRFGRSFDNFDRESRK
PNFLRFGK
>FLP-1_Caenorhabditis_tropicalis_Csp11.Scaffold629.g9640.t1
MEKRKPNFMRYGRSAAVKSLGKKAGSDPNFLRFGRSQPNFLRFGKASGDPNFLRFGRSDP
NFLRFGKAVADPNFLRFGKRSADPNFLRFGRSFDNFDRESRKPNFLRFGK
>FLP-1_Dictyocaulus_viviparus_nDv.1.0.1.t00713
MLTLLQVGLLLLALGAVAHVTAECCTSGEQTSFCLVYNMLAPVEQAEVMSYLGDACNGDAD
EALRLMEKRKPNFMRFGRSVAFGKKGSXDQHEDLA*DACNGDADEALRLMEKRKPNFMRF
GRRDACNGDADEALRLMEKRKPNFMRFGRSVAFGKKGSDPNFLRFGRNQPNFLRFGKASD
NPNFLRFGRAVGDPNFLRFGKRLDDSPNFLRFGRKPNFLRFGKTARLLGLL
>FLP-1_Diploscapter.coronatus_DCO_023616
MTLLIQVGLLAVLGLAVQAAAKCCSSQDVSHFCTVFHMLSPMEQNEVISYLGEDCSGDAD
EAVKQMDKKRMNFLRFGRSGLGKKAGSDPIFLRFGKRSDSPNFLRFGRSSEFDREVRKPN
FLRFGK
>FLP-1_Diploscapter_pachys_WR25_12872.1
MTLLIQVGLLAVMGLAVQAAAKCCSSQDVSHFCTVFHMLSPMEQNEVISYLGEDCSGDAD

EAVKQMDKKRMNFLRFGRSGLGKKAGSDPIFLRFGKRSDSPNFLRFGRSSEFDREVRKPN
FLRFGK
>FLP-1_Haemonchus_contortus_HCON_00103480
MTTLLQVGLLVALGIVAQAAAECCSNGDQSDFCMVFNMLSPMEQAEVMSYIGETCTGDAD
EALRLMEKRKPNFMRFGRSLTFGSKKGSDPNFLRFGRNQPNFLRFGRNQPNFLRFGKAAG
DPNFLRFGRGAGDPNFLRFGKRGVDPNFLRFGRKPNFLRFGK
>FLP-1_Haemonchus_placei_HPLM_0001038801-mRNA-1
MTTLLQVGLLVALGIVAQAAAECCSNGDQSDFCMVFNMLSPMEQAEVMSYIGETCSGDAD
EALRLMEKRKPNFMRFGRSLTFGSKKGSDPNFLRFGRNQPNFLRFGRNQPNFLRFGKAAG
DPNFLRFGRGAGDPNFLRFGKRGVDPNFLRFGRKPNFLRFGK
>FLP-1_Heligmosomoides_polygyrus_HPOL_0001223701-mRNA-2
MTTLLQVGLLLALGAVAQVAAECCLSGEQSDFCIVFNMLSPVEQAEVMGYLGDSCNGDAD
EAIKLMEKRKPNFMRFGRSPLGLSFGKKGSDPNFLRFGRSQPNFLRFGKAAGDPNFLRFG
RAAGDPNFLRFGKRSVDPNFLRFGRKPNFLRFGK
>FLP-1_Heterorhabditis_bacteriophora_Hba_04845
MFNILISGRNDPNFLRFGKAAADPNFLRFGKRSSDPNFLRFGRSDNFDRENRKPNFLRFG
K
>FLP-1_Mesorhabditis_belari_mbelari.g18410.t1
MPQTAPLAALFAALLIAMVRGDCCPREDRSDFCTIFTMLSPTEQNEVVQYLGDNCQGDAD
AALRLIEKRKPNFMRYGRSPTLELGKKGSDPNFLRFGKRSKPNYLRFGRSDPNFLRFGKK
AAGADPHFLRFGKRSAEPNFLRFGRSSEEFDRELRKPNFLRFGRK
>FLP-1_Micoletzkya_japonica_MicoRS5524-mkr-S105-1.7-mRNA-1
MATLLAILLATIVYKVSAECCVENDSSQFCLVFKMLSPMEQNEVINLIGDSCEGDANEAL
NRLEKRNPNFLRFGRSGMEARLSRSNGPNFLRFGRRDPNFLRFGKAAADPNFLRFGKRAS
GEPNFLRFGRQSQFEREARQPNFLRFGRK
>FLP-1_Necator_americanus_NECAME_08941
MPTLLQVGLFLTLGALAQVRHRSSFAGALFVELSSFRLSYHLSGVSGVLFYLENNQTSAS
FSNMLSPMEQAEVMSYLGETCNGDADEALRLIEKRKPNFMRYGRSITFGKKGSDPNFLRF
GRNQPSFLRFGKAAGDPNFLRFGRATADPNFLRFGKRSVDPNFLRFGRKPNFLRFGK
>FLP-1_Nippostrongylus_brasiliensis_NBR_0000031001-mRNA-1
MTTLLRHVGLLLALGVVAEVCFAFFDLISPIFTFTLLFYAQVQVAAECCVSSDQSEFCVV
FNMLSPMEQAEVMGYLGDTCTGDADEALRLIEKRKPNFMRFGRSKKHRESLSLGKKGADP
NFLRFGRNQPSFLRFGKAAAGDPNFLRFGRRAVDPNFLRFGRKPNFLRFGK
>FLP-1_Oesophagostomum_dentatum_OESDEN_17558
MEKRKPNFMRYGRSITFGKKGSDPNFLRFGRNQPNFLRFGKAAGDPNFLRFGRASADPNF
LRFGKRSVDPNFLRFGRKPNFLRFGK
>FLP-1_Oscheius_tipulae_OTIPU.nOt.2.0.1.t12420
MTGLLSWSICLAALVSQVYSQCCSPTDTSSFCTVFNMLSPMEQGEVMSYLGESCSGDADE
AIKVMEKRKPNFMRYGRSLAMGKKGSDPNFLRFGRSQPNFLRFGKAAGDPNFLRFGRSDP
NFLRFGKAAADPNFLRFGKRSADPNFLRFGRAADGFDREERKPNFLRFGK
>FLP-1_Parapristionchus_giblindavisi_Parapristionchus-mkr-
S_29-2.122-mRNA-1
MVTRFVSLALIAIALPKVVSECCEEGDNSEFCLVFNILSPMEQNEVINLIGDTCEGDANE
ALKKMEKRNPNFLRFGRSDMQMRLSRSNGPNFLRFGRRDPNFLRFGKAATDPNFLRFGKR
SSGEPNFLRFGRPSSFDREARQPNFLRFGRK
>FLP-1_Pristionchus_arcanus_arcanus-mkr-S_700-0.29-mRNA-1
MATLYVGLFFLAISLPTISSQCCSESDNSEFCIVFSMLSPMEQGEVINLLGHRCDGDVQG
ALKKMEKRNPNFLRFGRSQSGDAQLRLSRSNGPNFLRFGRSDPNFLRFGKAAADPNFLRF
GKRASGEPNFLRFGRQSSFDREARQPNFLRFGRK
>FLP-1_Pristionchus_exspectatus_exspectatus-mkr-S_568-0.4-mRNA-1
MATLYVGLFFLALTLPTISSQCCSESDNSEFCIVFSMLSPMEQGEVTNLLGHSCDGDVQG
ALKKMEKRNPNFLRFGRSQSGDAQLRLSRSNGPNFLRFGRSDPNFLRFGKAAADPNFLRF
GKRASGEPNFLRFGRQSSFDREARQPNFLRFGRK
>FLP-1_Pristionchus_fissidentatus_fissidentatus-ag_msk-S837-0.0-

```
mRNA-1
MEQNEVINMLDVQLRLSRSNGPNFLRFGRRDPNFLRFGKASADPNFLRFGKRASGEPNFL
RFGRQTAFDREARQPNFLRFGRK
>FLP-1_Pristionchus_japonicus_japonicus-mkr-S244-1.9-mRNA-1
MASPYMGLILIALSLPTISSECCLESDNSDFCLVFSMLSPMEQNEVTSLLGERCDGDAEG
ALKKMEKRNPNFLRFGRAIGDAQLRLSRSNGPNFLRFGRRDPNFLRFGKASADPNFLRFG
KRASGEPNFLRFGRQSSFDREARQPNFLRFGRK
>FLP-1_Pristionchus_maxplancki_maxplancki-mkr-S329-1.16-mRNA-1
PSIAYTTHSIMAVPYIGLLLAISFPTADNSQFCLVFSMLSPMEQSEVINLLGESCDGDVE
GVLKKMEKRNPNFLRFGRSQGGDGQLRLSRSNGPNFLRFGRRDPSFLRFGKAVADPNFLR
FGKRASGEPNFLRFGRQSSFDREARQPNFLRFGRK
>FLP-1_Pristionchus_mayeri_mayeri-mkr-S1592-0.7-mRNA-1
RKAATDQLRLSRSNGPNFLRFGRRAAAADPNFLRFGKGKRAXXXXXXXXXXXXXXXXXXX
XXXSFFLFSGRKAATDQLRLSRSNGPNFLRFG
>FLP-1_Pristionchus_pacificus_PPA14576.1
MATLYVGLFFLALSLPTCCSESDNSEFCIVFSMLSPMEQGEVTNLLGHSCDGDVQGALKK
MEKRNPNFLRFGRSQSGDAQLRLSRSNGPNFLRFGRSDPNFLRFGKAAADPNFLRFGKRA
SGEPNFLRFGRQSSFDREARQPNFLRFGRK
>FLP-1_Strongylus_vulgaris_SVUK_0001484501-mRNA-1
MLTLLQVGLLVALGAVAQVSAECCAPGDQSDFCVVFNMLSPIEQAEVMSYLVQPLVENQG
KIRKNSGAFGKKGSDPNFLRFGRNQPNFLRFGKASGDPNFLRFGKLAISAVV
>FLP-1_Bursaphelenchus_xylophilus_BXY_0709200.1
MTPHGAGVLLGILTVIYVKIGVEGCSGTGAFCRFYSQLDVMDQALINELMEEVQHAKPQK
RDSKFTGEFGKKGSEPNFLRFGKRAAPAPNAAGANFLRFGKSGADPNFLRFGKRATEFRL
DATEPNFLRFGKRPDPSAMSNNFLRFGKRSSLDTLDREQRQSNNFLRFG
>FLP-1_Halicephalobus_mephisto_MSTRG.4509.1.p1
KKNFRHFVMSRPSHTVAVLLLLSCFVQVFSVCCSSSENSEFCEFFDQLSPKEKNEIKKFF
GEDCESLKNMEKRKPNFIRFGRSDPNFLRFGRSAPQNNFLRFGKSSNTQQNFLRFGRATG
NEPNFLRFGRSTEPSFKRLTKSADPNFLRFGKRADTVADPNFLRFGKSNNFLRFGRSSGD
EDAFEREYRKPNFLRFG
>FLP-1_Panagrellus_redivivus_Pan_g13857.t1
MSLRSWHAVILGLLLGCVVQVFAKCCQPADQSAFCQSFEQLTLDEQREIQELLGDACDGN
ADEAIKSMEKRKPNFIRFGRSDPNFLRFGRAPAQNNFLRFGRTPQGQQNFLRFGKRSNEP
NFLRFGREPSFLRLTKSADPNFLRFGKRAGSDPNFLRFGKSNNNNFLRFGRSPSDDNAFE
REYRKPNFLRFG
>FLP-1_Parastrongyloides_trichosuri_PTRK_0000594500.1
MKNFILLVLLTVSYISSSITAFPDCCKTNQNAEVCLVFNKLSEDEKTFVTTEGVLDEQCE
LPHITPEKRKPNFIRYGRSTGGVPTAVDKKAADPNFLRFGRSSDHQNFLRFGRNLGLNEA
NFLRFGKSSSPDFLRFGKRNAEIKEPNFLRFGKRNNFLRFGRNLIDDQFNREYRKPNFLR
FGKRSNSNTNFLRFGRAPTAVLFDDTFDRNYRQQPDFLRFGK
>FLP-1_Rhabditophanes.sp.KR3021_RSKR_0000837500.1
MLVYTQNIIKLLLCSVVVLTATTTIPNCCKTNPNSAVCSQWTYFTPDEQAMIINENLIDT
DCSVIAMGKEKRKPNFIRYGRSSGQNPTALDKKASDPNFLRFGRSTTGEPNFLRFGRAAG
DNNFLRFGKSASPDFLRFGKRSFDNSKEPNFLRFGKRNNFLRFGRSQGDEDQFNREYRKP
NFLRFGKRSGAATNFLRFGRATAFLENESFARNYRQQPDFLRFGK
>FLP-1_Steinernema_carpocapsae_SC.X.g5249.2
MFPLVACASARLFISLAGSPPVLGECCSAESTSEFCSVFNLLSLPEQSEVRVILGENCDG
DVDEAMKKMDKRKPNFIRFGRSAPIQFGKKGSDPNFLRFGRSTADNNFLRFGKSNSGHQN
FLRFGRSDPNFLRFGRAANDPNFLRFGKSSDPNFLRFGKRQIDDSEPNFLRFGKRNNFLR
FGRAPSDQFDREYRKPNFLRFG
>FLP-1_Steinernema_feltiae_L889_g12017.t1
MFPFVAVPLLLVLVCVSQVFGECCTVDSRSEFCSVFSLLSLPEQAEVREILGENCDGDIE
EAMKKMDKRKPNFIRFGRSAPIQFGKKGSDPNFLRFGRSPTDHNFLRFGKSNAGHQNFLR
FGRSDPNFLRFGRAANNDPNFLRFGKSSDPNFLRFGKRQIDDSEPNFLRFGKRNNFLRFG
```

RAAPSEQFDREYRKPNFLRFG
>FLP-1_Steinernema_glaseri_L893_g3682.t2
MFPFLAVPLLVLVCVSQVFGECCSAESTSEFCSVFKLLSLPEQSEVRTFLGENCDGDLAE
AVKKMDKRKPNFIRKIILAAPIQFGKKGSDPNFLRFGRSTAEHNFLRFGKSNPGQQNFLR
FGRSDPNFLRFGRAANDPNFLRFGKSSDPNFLRFGKRQIDDSEPNFLRFGKRNNFLRFGR
ASAEQFDREYRKPNFLRFG
>FLP-1_Steinernema_monticolum_L898_g13587.t2
MFPFVAVPLLVLVCVSQVLGECCATDSTSEFCSVFNLLSRPEQAEVREILGENCDGDFEE
AMKKMDKRKPNFIRKIILAAPIQFGKKGSDPNFLRFGRSATEHNFLRFGKSNAGHQNFLR
FGRSDPNFLRFGRAATNDPNFLRFGKSSDPNFLRFGKRQIDDSEPNFLRFGKRNNFLRFG
RAPSDQFDREYRKPNFLRFG
>FLP-1_Steinernema_scapterisci_L892_g30548.t1
MFPLVAVLLLVFLSVSQVFGECCSAESTSEFCSVFNLLSLPEQSEVRVILGENCDGDIDE
AVKKMDKRKPNFIRFGRSAPIQFGKKGSDPNFLRFGRSTADNNFLRFGKSNVGHQNFLRF
GRSDPNFLRFGRATNDPNFLRFGKSSSDPNFLRFGKRQIDDSEPNFLRFGKRNNFLRFGR
APSDQFDREYRKPNFLRFG
>FLP-1_Strongyloides_papillosus_SPAL_0000437500.1
MKKYFRKTIFLILLTVFYYSIVAKAFPDCCKANPNAQICLEFDKLSLQEQASIKDSDVLD
DQCQIVTHATPEKRKPNFIRYGRSTGGVVTPAMDKKAADPNFLRFGRSDHQNFLRFGRTL
GIDDANFLRFGKGNSPDFLRFGKRNIEVGKEPNFLRFGKRNNFLRFGRNLVEEQFNREYR
KPNFLRFGKRGTTNTNFLRFGRSPGTLLFNDVFDRNYRQQPDFLRFGK
>FLP-1_Strongyloides_ratti_SRAE_X000064100
MIKYYQKNIFIILLAVNFFSIIINAFPECCKRNVNADICQGFDKLSPEEQASLSAVGVLD
DQCQLITHTIPDKRKPNFIRYGRSLSNMQQSLDKKAADPNFLRFGRSEHQNFLRFGRNLG
GNNANFLRFGKSNSPDFLRFGKRNMESDKEPNFLRFGKRDSFLGYGKNAVEEQFNREYRK
PNFLRFGKRSTTNTNFLRFGRSPAAVLFEDVFERNYRLQPDFLRFGK
>FLP-1_Strongyloides_stercoralis_SSTP_0000669700.1
MIKYCRKSILIVLFTIYCCSIIIKAFPECCKRNVNAEICQGFEKLSLEEQASLSAEGVLD
DQCQLITHTTPEKRKPNFIRYGRSLNVIPQPMDKKAVDPNFLRFGRSEHQNFLRFGRSLG
GNNGNFLRFGKSNSPDFLRFGKKSIEVGKEPNFLRFGKRENFIGFGKSMVEEQFNREYRK
PNFLRFGKRSTTNTNFLRFGRSPAAVFFEDVFDRNYRQQPDFLRFGNPPTEYMISYTVAD
RLGEDLDEYFCVKTETYNTNNSDKQQELLLPSVDRANVLPITAFTPTASLPPSRNINRES
FTGIKAPILRIRNKTDKTKRELESSTIKVYKPQKRTNYSVQPRKVVCLGTQKRTMIPNKI
IPGSTKTVKLSQKQNIYDKKYVFPSQPSPPHSDTSSHGCIDYTRNYCPSGQQYNNDGRIY
IEPRSNSNSSCSNRQTPEVEIDELMPNECNVIDYLDNDIYLDQNYMITEDYPKYEAQNND
VECKYQITDGCFDPSNMYYVTETNGIYNNQETNSNNIYEYVNQDLSLEKLTIQQTPTTISN
NILYKNSTTPVPDSSDSIKGCSLLRGMLENNTSNYNQPKKTINVRVIKSDIKIPQTETAL
INHNRDFKSKSIDTISSSGPSPRCSAEPSSRYSLANDNDNDFASFFIVEDSGNKTVPNES
KEQIEKRFNEDEEFKSKLSNNSLFEKPQLPYAAFVTLILNNLNTFAISVSEVYDGIIFLF
PHFETAYEGWRNSIRHNLSMSRLYEKVEAKVQIGSRKACMWSMVKREEAFNFNEVHKIDA
NMIDHIKSTMRFPQLWDPLVKGNLKLFPLSHSNLKAIVPEFISENEIEEYISYIRKYAFY
MRRLGNKHFMKSPRKTAKGRRKRVKKEDGLEEDCEYGSNINKGNESCTFDHISEPSIKRN
KIDELTFEDYPNNNVAKTDNKMENKQQSKVEESISEFDNFGKALTNDPFCLTPDEEESII
NNYFVSCNGFNDMADTLEDFSFSELHDDIMLENDLGFDDNKENYQSSYNNQTLFDNYEDL
NTMYNDRNKKNDLEEFNGMNNTESSLKDTDQWCLDLI
>FLP-1_Strongyloides_venezuelensis_SVE_1199400.1
MKKYFPKIIFLILLTVFYNSIVAKAFPDCCKANPNAQICLEFDKLSLQEQVSIKNSDVLD
NQCQIITHATPEKRKPNFIRYGRSTGGVVTSAMDKKAADPNFLRFGRSDHQNFLRFGRTL
GIDDANFLRFGKSNSPDFLRFGKRNIEVGKEPNFLRFGKRNNFLRFGRNLVEEQFNREYR
KPNFLRFGKRGTTNTNFLRFGRSPGTLLFNDVFDRNYRQQPDFLRFGK
>FLP-1_Acrobeloides_nanus_ACRNAN_Path_832.g3191.t1
MCPSYSYVLLTSTVFLAIISQVLSECCVSDTTSEFCIAYNVFLTPAEQAEVRKILGEKCE
GDVQEAVKEKRKPNFIRFGKRSVPGTLPFGKKGSDPNFLRFGRSAPANNFLRFGKSPSQQ
NFLRFGKRNADPNFLRFGRANEDSNNFVRLTKSADPNFLRFGKRSSLNEDSAEPNFLRFG

RNSNNFLRFGRAAEKFDQQNFLRFG
>FLP-1_Ditylenchus_destructor_Dd_07428
MSMDSPTKLRPRHIFGGMSSARSSIAIFVTVICLSLQMHLASALVQKLVPSGCCSTPESL
ATNFCKAFTSFNPEDQAELRQLLGSKCDEAAFEQSTVQKRKPNFIRYGKRSTLSSDPNFL
RFGRSNALSTLNDEDIVSILAMDKRVSTAANANFLRFGKSADPNFLRFGKRSPSAGDMTQ
DSAEPNFLRFGRKLPNNNFLRFGRSPSIEPEKFDREYRKPNFLRFG
>FLP-1_Ditylenchus_dipsaci_jg15578
MSMAAPQVGPMLFAGRLLVAVVVLCLSLQINLAESKASECCAAGDQTLFCKAFSSLNSME
QQELRILLGNKCDESAQSMTMEKRKPNFIRFGRSLGNSMHGSDPNFLRFGRSSPMIVEMR
SEQLGDGEAIPVKCQRQFPPIWQIF
>FLP-1_Heterodera_glycines_Hetgly.G000026531
MLCQCHQRHNITICGGGQAPSYVGRSERVLLLALSVCPLAGDMANQCRFQFTQIPKFIVV
SSEATMTGETQQQQNNATRFIRRQHANHGTEHRSLLLFLSLAIGCCALAQSHAADGGTSN
GHLAPMVPNSPISSIQSDPNFLRFGRSNGQLNEFNSASQTPTRTSSNFLRFGKSSMLVSE
PNFLRFGRQKVGGAGGSVDPTFLRFGRAKNNNFLRFGRAVGGDAMLISGDDDETPFTREY
RQANPNFLRFG
>FLP-1_Meloidogyne_arenaria_scaffold2249_cov78.g4396
MKGYCNMTELALFGLVILFIGQMSVLGANSANRNSLLMSGPWALNSWSDADPNFLRFGRS
AASNEEGIKRAAGQSANFLRFGRSAPYDPNFLRFGRQLGNQQQQHNKGLVDQSYLRFGRS
SGGNKGNNFLRFGRGAEDIPSEAEAFEREYRQSNNPNFLRFG
>FLP-1_Meloidogyne_enterolobii_scaffold1885_cov214.g3198
MKGYCNMTEMALFGLVILVVGQMSVLCANSANRNSLLMSGPWALNSWSDADPNFLRFGRS
AASNEEGIKRAAGQSANFLRFGRSAPYDPNFLRFGRQLGNQQQQHNKELVDQSYLRFGRSS
GGNKGNNFLRFGRGAEDIPSEAEAFEREYRQSNNPNFLRFG
>FLP-1_Meloidogyne_floridensis_scf7180000421285.g6595
MKGYCNMTELALFGLLVIFVAQMSVLGANSANRNSLLMSGPWALNSWSDADPNFLRFGRS
AASNEEGIKRAAGQSANFLRFAPYDPNFLRFGRQLGNQQQQHNKGLVGQSYLRFGRSSGG
NKGNNFLRFGRGAEDIPSEAEAFEREYRQSNNPNFLRFG
>FLP-1_Meloidogyne_graminicola_NXFT01002234.1.5946_g
MLQCDSNCIEEYVCCYAFCWTGTFEYENFIQFLGLSDYSNNNAEAEAMKRTAQSANFLRF
GRSMPYDPNFLRFGRQPHKVDQSYLRFGRNTNKGNNFLRFGRSQEQSEAEAFEREYRQSN
NPNFLRFG
>FLP-1_Meloidogyne_hapla_MhA1_Contig108.frz3.gene10
MTVLGANSANRNSLLMSGPWALNSWSEADPNFLRFGRSDPSGQVTSNEGIKRAAQSANFL
RFGKSAPYDPNFLRFGRANNNQQHNKGLVDQSYLRFGRSGAKANNFLRFGRGSEDIPTEA
EAFEREYRQSNNPNFLRFG
>FLP-1_Meloidogyne_incognita_Minc3s00049g02638
MKGYCNMTELALFGFVVLIVGQMSVLGANSANRNSLLMSGPWALNSWSDADPNFLRFGRS
AASNEEGIKRAAGQSANFLRFGRSAPYDPNFLRFGRQLGNQQQQHNKGLVGQSYLRFGRS
SGGNKGNNFLRFGRGAEDIPSEAEAFEREYRQSNNPNFLRFG
>FLP-1_Meloidogyne_javanica_M.Javanica_Scaff14978g071241
MKGYCNMTELALFGLLVIFVAQMSVLGANSANRNSLLMSGPWALNSWSDADPNFLRFGRS
AASNEEGIKRAAGQSANFLRFGRSAPYDPNFLRFGRQLGNQQQQHNKGLVDQSYLRFGRS
SGGNKGNNFLRFGRGAEDIPSEAEAFEREYRQSNNPNFLRFG
>FLP-1_Cylicostephanus_goldi_predicted
IKSIFFLQGDACSGDADEALRLMEKRKPNFMRYGRSELQIFAIRRYKKKNQSLGVTLGKKGSDPNFLR
FGRNQPNFLRFGKAAGDPNFLRFGKFTGMENTLGKGYK
>FLP-1_Pristionchus_entomophagus_predicted
PPPSLLIPFYNLCSSGRRDPNFLRFGKAAADPNFLRFGKRASGEPNFLRFGRQSAFDREARQPNFLRF
GRK
>FLP-1_Globodera_rostochiensis_predicted
SIYSLSRHGEFLTFCQNNFEGKSSASMSTSEPNFLRFGRQKGGVDPTFLRFGRANNNFLRFGRAAGGG
ERGIY
>FLP-1_Parascaris_equorum_predicted

STXXXXXXXXXXXXXXXXXXXXXXXXXCCNDGQTSDFCAVFNILSPTEQAEVRSYLGDNCDGDANEAA
KKIEKRKPNFIRFGRTGNAFRFYNTH
>FLP-2_Plectus_sambesii_PSAMB.scaffold294size58623.g4470.t1
MGPSTGFFLIATAVLCSSAASIFVPNTPYDDGDYSNIGDYQDGPNVPTSNEEDLIAMMKR
YREPVRFGKRAPREPVRFGKRALREPVRFGKRDSREPIRFGKRSADPEHTEKVSVSAPTA
>FLP-2_Ascaris_lumbricoides_ALUE_0000468201-mRNA-1
MIFRSKLLLIACIAFLALYVSIANSEELNQRIESERMARLQGGKRFRGEPIRFGKRAQRE
PIRFGKRVPFDEVPEQSYVYW
>FLP-2_Ascaris_suum_L3E_00746
MIFRSKLLLIACIAFLALYVSIANSEELNQRIESERMARLQGGKRFRGEPIRFGKRAQRE
PIRFGKRVPFDEVPEQSYVYW
>FLP-2_Dracunculus_medinensis_DME_0001024901-mRNA-1
MTFNDNINDNENLSEIKKRLRGEPVRFGKRIREPIRFVDDEYKKRRQPLGTMRFGKRQFS
ENDNFLYSPDSNENDEFFDEILMNKKNTATHPLDTMRFGKRIIFSEIPTTIYEMKDAIND
KRDSPLGTMRFGK
>FLP-2_Parascaris_equorum_PEQ_0000835201-mRNA-1
MARLQGGKRFRGEPIRFGKRAQREPIRFGKRVSFDDVPEQSYIHW
>FLP-2_Toxocara_canis_Tcan_09774.1
MIFRSKLLLIVCIVFFALCVSIAESQALKLRTEDGQVVRLQGGKRFRGEPIRFGKRAQRE
PIRFGKRESFEEVFEQPSLF
>FLP-2_Anisakis_simplex_predicted
SEAAVHERVSEDGIAYMRVPIMKRFRGEPIRFGKRAFREPIRFGKRTAQYPADTADTFLVDDQQQLY
>FLP-2_Parascaris_univalens_predicted
ELEGSQVGAKQSFFFLQPQRTPSLLTEAVLLQTNSEEFNQRIESERMARLQGGKRFRGEPIRFGKRAQ
REPIRFGKRVSFDDVPEQSYIHW
>FLP-2_ANCCAN_02789_peptide:_ANCCAN_02789_pep:protein_coding
MNPRKSLNRRSFISNIFSGIIMISKSLVTLVVLAILSLLTSAVSSQAETAMEARQQFKRF
RGEPIRFGKRVPREPIRFGKRAPLFEPYFDY
>FLP-2_Ancylostoma_ceylanicum_Acey_s0223.g2678.t1
MNPRKSLSRCSFISHIVSGVIMISKSLVTFVVLAILSLLASAVSSQAEAAMEARQQFKRF
RGEPIRFGKRVPREPIRFGKRAPLFEPYFDY
>FLP-2_Ancylostoma_duodenale_ANCDUO_01867
MISKSLVTLVVLAILSLLTSAVSSQAEAAMEARQQFKRFRGEPIRFGKRVPREPIRFGKR
APLFEPYFDY
>FLP-2_Angiostrongylus_costaricensis_ACOC_0000054801-mRNA-1
MASTGHGGPIVMFCDLGFTEAIASSKFYTVLVAAGPSQDENSSKNRQQFKRFRGEPIRFG
KRVPREPISEHIVQVAMFNLNHAHLKQQKNREPTIQRYYATFLQQASRNSRKAGMDASGI
MMSARTFLTIFCILLFTVLYTSSAPNRILMRFGKRATNTVPYAFVPLPDYYRVDLRRLTA
LFTTTLTETFGHKRKYCSVERQKTAHMVIVFHPQIEEVSEDRTEDRKASEQIEKKQLANV
YYSAEELAPLSLSFNPSLMDGNNRTRLINNATIR
>FLP-2_Caenorhabditis_briggsae_CBG01979b.1
MQASAILSALFLVLLAVIVSSFQFVQPKRILPIPTSREQLLRGQLAYLKGTTVAQSNDNQ
LGVFEASMMAKRLRGEPIRFGKRSPREPIRFGKRFNPLPDYDFQ
>FLP-2_C.elegans_,_isoform_a
MQVSGILSALFLVLLAVIVSPFQFVQPKRILPIPTSRDQLLRGQLAYLKGTTVAQPAVNDNTLGIFEA
SAMAKRLRGEPI
RFGKRSPREPIRFGKRFNPLPDYDF
>FLP-2_Caenorhabditis_inopinata_Sp34_X0065500.t1
MQLSGFLTVLFLALFAVIGTAVAQFANDNNLGPFEATLMAKRLRGEPIRFGKRSPREPIR
FGKRFSPLPDYDFQ
>FLP-2_Caenorhabditis.japonica_CJA03637.1
MQLAAILSALLLVLFAAIVSPFQFVQPKRILPTDRDALLRGQLAYLKGTTVAQAVANDNS
IGQFEASLMAKRLRGEPIRFGKRSPREPIRFGKRFNPLPEYDFQ
>FLP-2_Caenorhabditis_latens_FL83_01387

MQFSGILTTLLLVILAVIGTTVAQSANDNNLGPFEASMMAKRLRGEPIRFGKRSPREPIR
FGKRFNPLPDYDFQ
>FLP-2_Caenorhabditis_nigoni_
MQASAILSALFLVLLAVIGTTVAQSNIDNQLGVFEASMMAKRLRGEPIRFGKRSPREPIR
FGKRFNPLPDYDFQ
>FLP-2_Caenorhabditis_remanei_CRE03198.1
MQFSGILTTLLLVILAVIVSPFQFVQPKRILPIPTSRDQLLRGQLAYLKGTTVAQSANDN
NLGPFEASMMAKRLRGEPIRFGKRSPREPIRFGKRFNPLPDYDFQ
>FLP-2_Caenorhabditis_tropicalis_Csp11.Scaffold629.g13664.t1
MQVTGIFTALFLVLLAVIGTTIAQADDNDLAPFEASMMAKRLRGEPIRFGKRSPREPIRF
GKRFNPLPDYDFQ
>FLP-2_Cylicostephanus_goldi_CGOC_0000075501-mRNA-1
MISKSFMTFVVLTILSLLASAVSSQSDAGMEARQQFKRFRGEPIRFGKRAPREPIRFGKR
APLFEPYFDY
>FLP-2_Diploscapter_coronatus_DCO_024594
MDFSRVFAAILLIALSIMSEAVFFGENQIQDAYPRGMAKRLRGEPIRFGKRAPREPIRFG
KRAGETSSFSGFYPYDY
>FLP-2_Diploscapter_pachys_WR25_21230.1
MDFSRVFAAILLIALSIMSEAVFFGENQIQDAYPRGMAKRLRGEPIRFGKRAPREPIRFG
KRAGETSSFSGFYPYDY
>FLP-2_Haemonchus_contortus_HCON_00188000
MISKSLIIVVVLTVLCLLASAVSPQAEAMMESHQGFKRFRGEPIRFGKRVPREPIRFGKR
GPMFEPYFDY
>FLP-2_Haemonchus_placei_HPLM_0000601501-mRNA-1
MFNLIPLTHQETLQSLICLDFSDLLSRISNGAISYRFRFPIKSHISGIMISKSLIIVVVL
TVLCLLASAVSPQAEAMMESHQGFKRFRGEPIRFGKRVPREPIRLPYFDY
>FLP-2_Heligmosomoides_polygyrus_HPOL_0001006001-mRNA-1_
MIGKPLVILVILAVLTLLASAMSPQADGVLENRQQFKRFRGEPIRFGKRVPREPIRFGKR
TSSFQPYFDY
>FLP-2_Heterorhabditis_bacteriophora_Hba_13882
MQVPTMANKSLIAFLLIVILNLLVTALPVNSKSSVEQRQQFKRFRGEPIRFGKRVPREPI
SQPYVSYKNKRVDKLDICDMSAIHLLVTLATLFAIIVGQDYYNEKRSPLGTMRFGKRMAI
DDNYYYLDKKADSTMAETQRSIRTPLGTMRFGKRTSLRLEKRNPLSTMRFG
>FLP-2_Mesorhabditis_belari_mbelari.g9736.t1
MNARNFLLCLFFALFALLTSAYPYEAQSNNLLGQRESKRFRGEPIRFGKRSPREPIRFGK
RAAGIDEIEDYLSQL
>FLP-2_Micoletzkya_japonica_MicoRS5524-mkr-S355-0.26-mRNA-1
MISRFVALLLVVLAFTVMAQNEEEERGGVYHIPLKRFRGEPIRFGKRAPREPVRFGKRAQ
PLEWYYGLDY
>FLP-2_Necator_americanus_NECAME_02416
MVSKSLVTLVVLSILSLLTSAVPLQTESAMETSQQFKRFRGEPIRFGKRVPREPIRFGKR
APLFEPYFDY
>FLP-2_Oesophagostomum_dentatum_OESDEN_15039
MISKSLVTLVVLAILSLLASAVSSQEASLEARQQFKRFRGEPIRFGKRVPREPIRFGKRA
PLFEPYFDY
>FLP-2_Oscheius_tipulae_OTIPU.nOt.2.0.1.t07396
MQVRTVFSLLILALLSILGSTMSAYNRGIDNMPMKRFRGEPIRFGKRSPREPIRFGKRFD
PSGDYY
>FLP-2_Parapristionchus_giblindavisi_Parapristionchus-mkr-S_7-1.13-
mRNA-1
MISRFAALLFFLIATLMIAQAQNLDGISGYEFPMKKRFRGEPIRFGKRAPREPVRFGKRA
SYYPSHGFNAENEY
>FLP-2_Pristionchus_arcanus_arcanus-mkr-S_11-7.12-mRNA-1
MISRHVLILILCITITVFAQAHNYDGPQNTEYLIKKRFRAEPVRFGKRAPREPVRFGKRS

PEHFFYPFELHNQADIEY
>FLP-2_Pristionchus_entomophagus_entomophagus-mkr-S131-0.6-mRNA-1
MLYRHALLLILAFAITLFTQGQTYDETAPAGYEYAMKKRFRAEPVRFGKRAPREPVRFGK
RSSGQRLYPFELPIEY
>FLP-2_Pristionchus_exspectatus_exspectatus-mkr-S_589-0.35-mRNA-1
IADVLSPLNLRLTPLKMISRHVLILILCITITLSAQALNYNGPQNTEYLMKKRFRAEPVR
FGKRAPREPVRFGKRATEHFFRPFELHQNADIDY
>FLP-2_Pristionchus_fissidentatus_fissidentatus-mkr-S6-5.42-mRNA-1
MISRLAALLLLLLATVLIAQAQSFDGGAPGYEYEMKKRFRAEPVRFGKRAPREPVRFGKR
SGQQQQPAFYPFELLNRDVVEY
>FLP-2_Pristionchus_japonicus_japonicus-mkr-S39-1.0-mRNA-1
MISRQTILLFLCITFAIMTEIQAFNEVEFSGHETPMKKRFRAEPVRFGKRAPREPVRFGK
RSSGQLFYPFDLLNNPA
>FLP-2_Pristionchus_maxplancki_maxplancki-mkr-S210-1.15-mRNA-1
MISRCSHLLILAIAFILILHAQSYDEAESAGYEFPIRKRFRAEPVRFGKRAPREPVRFGK
RSTGQLFYPFELFNHPNANY
>FLP-2_Pristionchus_pacificus_PPA39825.1_peptide:_PPA39825.1_pep:pro
tein_coding
MISRHVLILILCITIALFAQAQNYNGPQNTEYLIKKRFRAEPVRFGKRAPREPVRFGKRA
TEHFLHSFKLHQNSDIDY
>FLP-2_Strongylus_vulgaris_SVUK_0000951601-mRNA-1
MISKSMVTFVVLAILSLLASAVSSQSDAAMEARQQFKRFRGEPIRFGKRVPREPIRFGKR
APLFEPYFDY
>FLP-2_Teladorsagia_circumcincta_TELCIR_08098
MISKSLIIVIVLAVLSLLASAVSPQAEAMMESRQQFKRFRGEPIRFGKRVPREPIRFGKR
GPMFEPYLDY
>FLP-2_Caenorhabditis_angaria_predicted
SPFQLMQPKRILHADKNDLFRRQATFLSGGKSVAKPIQPEEPVQPEKEEQKDEPSFLQQLEQASYFEP
YESAIVAKRLRGEPIRFGKRSPREPIR
>FLP-2_Caenorhabditis_brenneri_predicted
LTVSRSQNHLSQTMQVSGILSTLFLVLLAVIVSPFQFVQPKRILPIPTSRDQLLRGQLAYLKGTTVAQ
SVANDNNIGLFEASMMAKRLRGEPIRFGKRSPREPIR
>FLP-2_Caenorhabditis_sinica_predicted
LTVSRCCQNSPPLSQTMQVSGILSALFLVLLAVIVSPFQFVQPKRILPIPTSRDQLLRGQLAYLKGTT
VAQSISNDNNLGVFEASMMAKRLRGEPIRFGKRSPREPIR
>FLP-2_Dictyocaulus_viviparus_predicted
KNFKNHISQVSPALSQEEVIMGNRQQLKRFRGEPIRFGKRVPREPIRYAGINIVFH
>FLP-2_Nippostrongylus_brasiliensis_predicted
AYWSVSISMFILYRSSFIIFKFKLITISKFQKKFQASAVSPQQAEAIMESRQQFKRFRGEPIRFGKRV
PREPIR
>FLP-2_Pristionchus_mayeri_predicted
IDPLRFGKRAPREPVRFGKRSSGASQLYPSERLTNPLEY
>FLP-2_Bursaphelenchus_xylophilus_BXY_1523000.1
MTYALSLMLVVVLVMEMCHSVPEQDFYRSENSNAMPLYMERRFSKFRGEPVRFGKRAYRE
PIRFGKRAYMPNNQISNGGYGQNAEN
>FLP-2_Halicephalobus_mephisto_HMEPH_08203-RA.p1
LGKLTLISKRTTNIINFYSMIFKFLIISTLLFFIFNKNTQALQPSDDNYIPGDYFLQEFT
PVKRYRTEPIRFGKRAAFREPIRFGKRHWSKMNNIFDIQNSNFASRIQNPSQP
>FLP-2_Steinernema_carpocapsae_SC.X.g3562_
MAAYSSIPVQSHMSSSTLSYLFFLLLLVAVFQTSSAYEEKSDYDMAEMTPYELQVKRFRT
EPIRFGKRGPREPIRFGKRAMTNSYAPHRLPGFPAFYGTYNH
>FLP-2_Steinernema_feltiae_L889_g30046.t1
MAAQVLSSSISGQSQMTSSALSYLFIFVLLFNVFQISVSYEEKARYVIPESSIYELQAKR
FRTEPIRFGKRGQREPIRFGKRAVIDFHSPSRLPLYSGFYAKFEQK

```
>FLP-2_Steinernema_glaseri_L893_g22383.t1
MAARSSFASSQSRMTLSTFNCLLVFILLLNLLQSGCCYEEKADYALPASARYELQVKRYR
TEPIRFGKRGQREPIRFGKRIVLNLNIPHRLSGYPPLNSKFQHH
>FLP-2_Steinernema_monticolum_L898_g11232.t1
MFLKFVTAQTSEIRCHLKVAMRSLASYLATRSSFSVSNLKRMAARAISTSSHSPMTSNIF
NYLFSFVLLFSVFQISICYEEKTDYMLPESAPYELQAKRFRTEPIRFGKRGQREPIRFGK
RTVMNFHGLSRLPPLYPSFYSKFEQK
>FLP-2_Steinernema_scapterisci_L892_g1312.t1
MAAACSSSIQLQSHMSSSALSYLFFLLLLVAVFQISTAYEEKSDYNLAEMTPYELQVKRY
RTEPIRFGKRGPREPIRFGKRAMANSYASHRLPAFPAFYGKYDH
>FLP-2_Strongyloides_papillosus_SPAL_0001444700.1
MNRLSLIFFCVFLSILVLTPSSVIGSEIRFPSRDVMKAMLRDPNAYFRGEPIRFGKKSNF
REPIRFGKRGLLHTTEKGDLKEMVNVNNENFDGSQFSPIPLYR
>FLP-2_Strongyloides_ratti_SRAE_X000120400
MNNISLCFYGLLLTILIFTSNNVFGSDFNFPSRDIMKAMLRDPNAYFRGEPIRFGKKSSF
REPIRFGKRNSPNLKKNKGNEIMNFDNENDNFDRKYYIPVPFYNFE
>FLP-2_Strongyloides_stercoralis_SSTP_0001244600.1
MNNLSSFGFFGLLLTILLFTSNNVFGNDFHFPSRDVMKAMLKDPNAYFRGEPIRFGKKSN
FREPIRFGKRNFPNLDKKENNEIMNFNSENGNFYEGYYIPVPFINFE
>FLP-2_Strongyloides_venezuelensis_SVE_0144400.1
MNRLSFIFLCVFLSILVLTPSSVIGSEVRFPSRDVMKAMLRDPNAYFRGEPIRFGKKSNF
REPIRFGKRGLQHTTEKGDSKEIVNVNNENFDGSKFIPIPLYR
>FLP-2_Panagrellus_redivivus_predicted
IDPLRFGKRAPREPVRFGKRSSGASQLYPSERLTNPLEY
>FLP-2_Parastrongyloides_trichosuri_predicted
NVLGNDFRYPSRDVMKAMLRDPKAYFRGEPIRFGKKSSFREPIRFGKRMINNEKLESGEIEKNIDNDE
SHDVPLSYYSFL
>FLP-2_Rhabditophanes_sp_KR3021_predicted
TAMCKDSRLPSIDVMKAMLENPKAYFRGEPLRFGKRSSFREPIRFGKRAYEVPMEVPLQHYYSD
>FLP-2_Acrobeloides_nanus_ACRNAN_Path_1124.g4344.t1
MMMLNKLIFVLIQLIFFVSLLKSCQAHALRYPVQQYLEMEQLNRLRNAAVKRFRSEPVRF
GKRAAREPIRFGKRIPDQFSELQDLLT
>FLP-2_Globodera_pallida_GPLIN_000172000
MPLFNASVLASLLFICVGLLHQPTVDALRQQVETNVKDLSFGSSYPMAKRPYFREPLRFG
KRFGEGSAGGGGLLINPFLYEAYRNGADPFQK
>FLP-2_Globodera_rostochiensis_GROS_g03098.t1
MPLFNASVLASLLFICVGLLHQPTVDALRQQAETNVKDLSFGSSYPMAKRPYLREPLRFG
KRFGEGSAGGGGGLLINPFLYEAYRNGADPFQK
>FLP-2_Heterodera_glycines_Hetgly.G000022202
MLPSNNTTLAILLLIVQCAMLFHQSMVSGVPHRSKTTENELTLGSYPMAKRPYFREPLRF
GKRFSGGSNGGGGFLINPFLYEAYRSENPFEK
>FLP-2_Meloidogyne_arenaria_tig00000327.g59380.t1
MTTKHINLFILLALLMTTAYCFEGIGNKQYKELYPMWKRQMREPLRFGKRQEDKTNYFKN
SNENADNGFPDEAYNGISFQK
>FLP-2_Meloidogyne_enterolobii_scaffold6511_cov291.g9556
MTTKHINLFILLALLITTAYCFEVIGNKQYKEIYPMWKRQMREPLRFGKRQEDKTSYFKN
SNENADNGFPDEAYNEVAFQK
>FLP-2_Meloidogyne_floridensis_maker-nMf.1.1.scaf14423-augustus-
gene-0.8-mRNA-1
MTIKHINLFILLALLMTTAYCFEGIGNKQYKELYPMWKRQMREPLRFGKRQEDKTNYFKN
SNENADNGFPDEAYNGISFQK
>FLP-2_Meloidogyne_graminicola_NXFT01001842.1.4783_g
MVISNSFEELDKQMYPMWQRQMREPLRFGKRLLKQKTVISNCSPVTRQEIIPSLLPVGTL
HESLNDPFLQQQRWLSVRPFKEYPRPHNEELGLLLQYLDSKRNAPILDENVAQLVGESPL
```

GTMRFGKRRNSPPLGTMRFG
>FLP-2_Meloidogyne_incognita_Minc3s00218g07824
MTTKHINLFILLALLMTTAYCFEGIGNKQYKELYPMWKRQMREPLRFGKRQEDKTNYFKN
SNENADNGFPDEAYNGISFQK
>FLP-2_Meloidogyne_javanica_M.Javanica_Scaff996g011868
MTIKHINLFILLALLMTTAYCFEGIGNKQYKELYPMWKRQMREPLRFGKRQEDKTNYFKN
SNENADNGFPDEAYNGISFQK
>FLP-2_Meloidogyne_hapla_predicted
CFEVLGNKQYKEVYPMWKRQMREPLRFGKRLASDTDYFKYEENAGNRLTYDNYKSRISFQNY
>FLP-3_Plectus_sambesii_PSAMB.scaffold294size58623.g4471.t1
MMSGGSAATHFSTVNQANVRRRPFGRQRCLKSAPSAVYSHLLTSHIVMISPTLALLLGLA
AAATAHSFYELDDGYNMRADEDQKRALPGLMRFGKRSPLGVMRFGKRDSFEEPKRAPNLG
TMRFGKRSPLGVMRFGKRSPLGVMRFGKRSPLGTMRFGKRLPLGTMRFGKRAESLPGALR
FGKRESDIDMWYDAAVQEAEKRELNFDENAAHKAEEVMKPIVLQ
>FLP-3_Acanthocheilonema_viteae_nAv.1.0.1.t02960-RA
MKLENELESSDSSLHGKRSNEESKAIHFIRSHLGIMRFGKRIRFDKRIDLDDTMHQNDKN
SDTEHSNDYNDPYGKTSSFDYQLTF
>FLP-3_Anisakis_simplex_ASIM_0001362101-mRNA-1
MSSIVPSNNNNNGNKYESLSNDFNENSAEDEIKKDEVLVPNNGILKRSDYQQRARLGTMR
FGKRDASNPLGTMRFGKRFYQNADQQQLYDNNNVAMLNHAYLYRLINNNVANISPFANND
HFIRYNNNNNNNKRNSPLGTMRFGKKRATGSAVGPLGTMRFGKKR
>FLP-3_Ascaris_lumbricoides_ALUE_0000468301-mRNA-1
MEDGICAEWMPNTKVKSGVKIGEQVGRTRMSTRQWLNGAIDKIGNRMYMTDEDGIWDGIW
ASHVNRANDDNSNTQRSIRTPLGTMRFGKRDANPLGTMRFGKRTSPNDEQLFDEKRNGPL
GTMRFGKRGDPLGTMRFGKRGGPLGTMRFGKR
>FLP-3_Ascaris_suum_AgR027X_g064_t02
MLKKHRTGTNLIIMEDGICAEWMPNTNVKSGVKIGEQVESNEDEYVRRQWLNGAIDKIGN
RMYMTDEDGIWDGIWASHVNRANDDNSNTQRSIRTPLGTMRFGKRDANPLGTMRFGKRTS
PNDEQLFDEKRNGPLGTMRFGKRGDPLGTMRFGKRGGPLGTMRFGKR
>FLP-3_Brugia_malayi_Bm14284.1
MQACARISDRLDYSDISLHGKRSDEESKAIHFIRSHLGIMRFGKRNKAQFDTTHIDNRID
LDHIMRQNDESSIINNSNDYSNSYVKATLFDSIN
>FLP-3_Brugia_pahangi_BPAG_0000905801-mRNA-1
MQACARISDRLDYSDISLHGKRSDEESKAIHFIRSHLGIMRFGKRNEAQFDTTHIDNRID
LDHIMRQNDKSSIINNSNDYSNSYVKATLFDYIN
>FLP-3_Brugia_timori_BTMF_0000976501-mRNA-1
MQACARISDRLDYSDISLHGKRSDEESKAIHFIRSHLGIMRFGKRNEAQFDTTHIDNRID
LDHIMRQNDESSIINNSNDYSNSYVKATLFDSIN
>FLP-3_Gongylonema_pulchrum_GPUH_0000073001-mRNA-1
MFPTENGFGSIWSYEKRAEDETAARSARTPLGTMRFGKRGTVPLGTMRFGKRSELENFLD
ALVPVHPNGDRVDSNSGYVDNFYGKTVYLGPSGKRNSPLGTMRFGK
>FLP-3_Loa_loa_EN70_10045
MIPEYGWECSDSSLHGKRTNEAGKAIHFIRSHLGLMRFGKRNAAQYGIIRFNKRIDLDHL
TQQNDEGNSIDSDNGYRNSYIKAYPFGY
>FLP-3_Onchocerca_ochengi_OOCN_0000410201-mRNA-1
MITCFYSTDILDRSECKSSTCWHEKRSEEKYKTIRFIRSHLGIMRFGKRSDTIRLNNRIE
LNDFLHASAMSRQNDKSGSMDSSNSYGNFYDKTSY
>FLP-3_Onchocerca_volvulus_OVOC5368.1
LSFPYMNPFTTIIIIFFTVSICYMQVSALISDRSECKSSTCWHEKRSEEKYKTIRFIRSH
LGIMRFGKRSDTIRLNNRIELNDFLHASAQPCRDKTIKVAAWTAVTATQSGLADGIDSAL
SCNYDCFILHYAQPIQS
>FLP-3_Parascaris_equorum_PEQ_0001278001-mRNA-1
MNECESNEDEYVRRQWLNGAIDKIGKRMYMMDENGIWDGIRTSPVNRANDDNSNTQRSIR
TPLGTMRFGKRDANPLGTMRFGKRTSPNDEQFFDEKRNGPLGTMRFGKRRGPLGTMRFGK

R
>FLP-3_Parascaris_univalens_PgR074X_g017_t01
LSGYQIRTLNQASKLANKLVRLRFHRSFHLQVEIISVVSTSLMHASIFAFALLTLRMALA
SESNEDEYVRRQWLNGAIDKIGKRMYMMDENGIWDGIRTSPVNRANDDNSNTQRSIRTPL
GTMRFGKRDANPLGTMRFGKRTSPNDEQFFDEKRNGPLGTMRFGKRRGPLGTMRFGKR
>FLP-3_Thelazia_callipaeda_TCLT_0000106101-mRNA-1
MSKDKDGCGETCFNGKQSGNGNKADRFIRSPLGTMRFGKRGNAPFNPMRFGKSYDSDDLS
TILSMTYENDGNNNAGVNNDYPVFEGNPFYLNSYQKRRNPLGTMRFGK
>FLP-3_Toxocara_canis_Tcan_09770.1
MALASESNEEYVRRQLLNGIRMGNRLYTLNEDSGISDGEWAHVKRSTDDNVNQQRAIRTP
LGTMRFGKRDANPLGTMRFGKRTLNDELIEEKRSGPLGTMRFGKRAGPLGTMRFGKR
>FLP-3_Dirofilaria_immitis_predicted
QKEFLDQSECRSNICWHEKRSNEKYKAIRFIRSHLGIMRFGKRSAAAKFGTIRFNKRIDLNDFLHASA
IMQQNDESRSIDSNNDYSNFYGKVSY
>FLP-3_Elaeophora_elaphi_predicted
PAYTSPQFADGSECSDNCLHGKRSNEESKAIHFIRSHLGIMRFGKRSAAQFGTISFDKRIDLDRIIHQ
NDENSNTDSSNDYSISYGKTPSFDLN
>FLP-3_Wuchereria_bancrofti_predicted
SNEESKAIHFIRSHLGIMRFGKRNEAQFDSTRVDKRIDLNYTMRQNDESSIIH
>FLP-3_Caenorhabditis_brenneri_CBN32881.1
MISLNHLILFFCVGCAFLEASDATPKRSPLGTMRFGKRSIGDELAYEEAVFPGAMWKRST
VDSEPVIRDQRTPLGTMRFGKRKPFGTMRFGKRDPENDAPFGTMRFGKRANEDTPFGTMR
FGKRDSEPFGTMRFGKRGDDGTPFGTMRFGKRENDGAPFGTMRFGKRGSDEPLGTMRFGK
RSSDDNAPFGTMRFGKRSSDDNAPFGTMRFGKRTPLGTMRFGK
>FLP-3_Caenorhabditis_briggsae_CBG01978.1
MISPNRLILFFLIGCAVSAASEASPKRSPLGTMRFGKRASLDDVLAFEEEYPGVLWKRST
VDSEPVIRDQRTPLGTMRFGKRSAEPFGTMRFGKRDSEIDAPFGTMRFGKRETVDAPFGT
MRFGKRAAEDAPFGTMRFGKRDPEPFGTMRFGKRSSEPFGTMRFGKRSDEPLGTMRFGKR
SADDTGAPFGTMRFGKRNTLGTMRFGK
>FLP-3_ Caenorhabditis_elegans_
MISPNHLILLFCVNCAFLVASDATPKRSPLGTMRFGKRAIADEMTFEEDGYYPSNVMWKRSTVDSSEP
VIRDQRTPLGTM
RFGKRSAEPFGTMRFGKRNPENDTPFGTMRFGKRASEDALFGTMRFGKREDGNAPFGTMKFGKREAEE
PLGTMRFGKRSA
DDSAPFGTMRFGKRNPLGTMRFGK
>FLP-3_Caenorhabditis_inopinata_Sp34_X0065400.t1
MISLNHLIVLACVGCAFLAALDVTPKRSPLGTMRFGKRSMLDELLVDEDGYSPLGEMWKR
STVDSSEPVIRDQRSPLGTMRFGKRGAEPLGTMRFGKRDLENDVPFATMRFGKRAAGDDA
PFGTMRFGKRDYNEASLGTMRFGKRASDEPLGTMRFGKRSADDSTPFEPMRFGKRNPLGT
MRFGK
>FLP-3_Caenorhabditis_japonica_CJA06079.1
MISLNHLILLFCVGCAIMAASDVTPKRSPLGTMRFGKRSYDSTVGSFDDDVYYPVELYKR
SGDDSDLVIRDQRTPLGTMRFGKRSSEPFGTMRFGKRELESDAPFGTMRFGKRSADEAPF
GTMRFGKRDNEPFGTMRFGKRGAPEEAPFGTMRFGKRDDDGTPFGTMRFGKRASDEPLGT
MRFGKRSVDDSPFQTMRFGKRNPLGTMRFGRK
>FLP-3_Caenorhabditis_latens_FL83_01388
MISVNQLILLSCLGCAVVVASYATPKRSPLGTMRFGKRAVGDELAYEEDGFYPGMMWKRS
TADSEQVIRDQRTPLGTMRFGKRSAEPFGTMRFGKRDPENDAPFGTMRFGKRGADDTPFG
TMRFGKRGDGETPFGTMRFGKRSSEPFGTMRFGKRGDDGTPFGTMRFGKRSSDEPLGTMR
FGKRSSDDSAPFGTMRFGKRTPLGTMRFGK
>FLP-3_Caenorhabditis_nigoni
MISPNRLILLFLFGCAVSAASEASPKRSPLGTMRFGKRAASLDDVLAFEEDYPGVLWKRS
TVDSEPVIRDQRTPLGTMRFGKRSAEPFGTMRFGKRDSETDAPFGTMRFGKRETEDAPFG
TMRFGKRAADDSAPFGTMRFGKRDPEPFGTMRFGKRSSEPFGTMRFGKRSGWFNEDVATV

TLILSDEPLGTMRFGKRSADDTGAPFGTMRFGKRNPLGTMRFGK
>FLP-3_Caenorhabditis_remanei_FL81_01518
MISVNQLILLSCLGCAVVVASDATPKRSPLGTMRFGKRAVGDELAYEEDGFYPGIMWKRS
TADSEQVIRDQRTPLGTMRFGKRSAEPFGTMRFGKRDPESDSPFGTMRFGKRGADDTPFG
TMRFGKRGDGETPFGTMRFGKRSSEPFGTMRFGKRGDDGTPFGTMRFGKRSSDEPLGTMR
FGKRSSGDSAPFGTMRFGKRTPLGTMRFGK
>FLP-3_Caenorhabditis_sinica_Csp5_scaffold_00147.g5616.t1
MISLNHLILLICIGCAFLVASDAPPKRSPLGTMRFGKRAIDDMVAYEEDGYYPGVMWKRS
TVDSEPVIRDQRTPLGTMRFGKRSAEPFGTMRFGKRDPEDAPFGTMRFGKRASDDAPFGT
MRFGKRDNEPFGTMRFGKRGDDATPFGTMRFGKRSSDEPLGTMRFGKRSADGGAPFGTMR
FGKRTPLGTMRFGK
>FLP-3_Diploscapter_coronatus_DCO_022714
MLNRKFTETISHMRLSRLAVSSLVVSLCCVGLTLAKDSDDKRSPLGTMRFGKRDQGEISG
ENMFYELYKRSAGENGLELVEDETPGLLGDEILPLRDVRAPRQNRAAPFGTMRFGKRYGV
GTSSDVVPLGTMRFGKRNPLGTMRFGRK
>FLP-3_Diploscapter_pachys_WR25_21231.1
MLDRKFTEAISHMRLSRLAVSSLVVSLCCVGLTLAKDSDDKRSPLGTMRFGKRDQGEISG
ENMFYELYKRSAGENGLELVEDETPGLLGDEILPLRDVRAPRQNRAAPFGTMRFGKRYGV
GTSSDVVPLGTMRFGKRNPLGTMRFGRK
>FLP-3_Mesorhabditis_belari_mbelari.g9735.t1
MIPAYYLALLAVLCTVTLAQLEPIPENLESSDADEDKRTPLGTMRFGKRAILESADDDFL
DDKLSFNIFDGLLREARSPLGTMRFGKRSGGDKGKALGTMRFGKRGDPLHTMRFGKRQQN
PFSWCVMKNLSWRPMLACRNTMRFGKRDPLGTMRFGRK
>FLP-3_Micoletzkya_japonica_MicoRS5524-mkr-S355-0.25-mRNA-1
MSPLTVSVLIAALVALSLADQVPEGLKRSAAFGTMRFGKRAAAFRMLENYDEDELSRVAR
SGASLGTMRFGKRAMGDAMMAPNDYDLYKRASALGTMRFGR
>FLP-3_Oscheius_tipulae_OTIPU.nOt.2.0.1.t07395
MTSTALLLPFLFLVAVVAGEMANKRAPLGTMRFGKRDYVDDDMDYSGLDKRSLDEETQRS
IRAPLGTMRFGKRAPLGTMRFGKRSYEEGAPFGTMRFGKRDPLGTMRFGK
>FLP-3_Parapristionchus_giblindavisi_Parapristionchus-mkr-S_7-1.12-mRNA-1
MSATTTTFLLCTIILAAALADFNADYKRSAFGTMRFGKRSNGVYEMQGDSLPRLARSGAS
LGTMRFGKRAMSHEIEPEDMEYYSGVKRAAALGTMRFGRR
>FLP-3_Pristionchus_arcanus_arcanus-mkr-S_68-3.71-mRNA-1
MARILLAVLCIAVLCIALSYAGYNEELMKRNAFGTMRFGKRSMQLLESGEAPNLPRLTRS
GASFGTMRFGKRSMASSGEYLQ
>FLP-3_Pristionchus_entomophagus_entomophagus-sn_msk-S131-0.21-mRNA-1
MANTLIAVLILCVALVSAGKFFHCIRNTLNLVTFSRFQNYNEEIIKRNAFGTMRFGKRSM
RLLESGEASGLPRLTRSGASLGTMRFGKRSMASAEYEVR
>FLP-3_Pristionchus_exspectatus_exspectatus-mkr-S_22-3.0-mRNA-1
MAKLLLAVLFLCVALIYAGYNEELMKRNAFGTMRFGKRSLPLFESGEASVTLPRLTRSGA
SFGTMRFGKRSMPSSGEYLQ
>FLP-3_Pristionchus_fissidentatus_fissidentatus-mkr-S6-5.4-mRNA-1
MASTSPDSSLPSLHYFEMANLAAVFLLIVALVAAEYNEEAMKRSNAFGTMRFGKRALRLM
ESGDMTSLPRLTRAGASLGTMRFGKRSMASAEYEPEELDLFGGSKRSAALGTMRFGRR
>FLP-3_Pristionchus_japonicus_japonicus-mkr-S39-1.1-mRNA-1
MANTVAVLILCVALVSAGYTEELMKRNAFGTMRFGKRSMRLLESGEVSANLPRLTRSGAS
LGTMRFGKRSMAVGAYEPEQMAMFHGVKRSAALGTMRFGRR
>FLP-3_Pristionchus_mayeri_mayeri-mkr-S241-5.23-mRNA-1
MAHTLLAVLIICVALVSAEYSDDLTKRNAFGTMRFGKRSMRLVESGEASAGLPRLTRSGA
SLGTMRFGKRAMATAELELPEEMDIFPGAKRSAALGTMRFGRR
>FLP-3_Pristionchus_pacificus_PPA39821.1
MAKVLLSVLILCVALVYADYSEELVKRNAFGTMRFGKRSASDLLESGEDSSNSPRLTRSG

ASLGTMRFGKRSMAVGEYATEEMSRDAKRSAALATMRFGRR
>FLP-3_Heterorhabditis_bacteriophora_predicted
GTSLRLEKRNPLSTMRFG
LLSDYYNEKRSPLGTMRFGKRMAIDDNYYYLDKKADSTMAETQRSIRTPLGTMRFGKRK
>FLP-3_Teladorsagia_circumcincta_predicted
DFPFPGPLGTFRFGKRSGRPERLPLIDQGN
>FLP-3_Caenorhabditis_angaria_predicted
GSVKDQVSLEFKWGPLKNMFFCLDGPLGTMRFGKRSGN
SFLDEKIQQMILDVPLGTMRFGKRSGK
DPRFGKVRQFL
QRAIGKTSSVGNNEMGKTIGRFGRRPSWFQSL
FLLLTLFFLDEPFGTMRFGKRDPLGTMRFGK
TSDDKELSGKRAPLGTMRWGKRSVDLDDDQVGFNLYERDVRSPLGTMRFGKRSGEPRVQVGTS
TIFIDEPFGTMRFGKRSGKLVRKNSKNNFC
ISDEPFGTMRFGKRSGKLNIDK
ISDEPFGTMRFGKRSGKFLKPNSSS
>FLP-3_Caenorhabditis_tropicalis_predicted
YRSITSSSCAVWVARFWRRRVSQWVVVLITRGLVLDATPKRSPLGTMRFGKRANDELMFEDDDYPSLL
WKRSTVDSEPVIRDQRTPLGTMRFGKRSAGRDSL
RELFIVFLEPFGTMRFGKRDSENDTPFGTMRFGKRGDDAPFGTMRFGKRDNGKD
>FLP-3_Bursaphelenchus_xylophilus_BXY_1522900.1
MSGWTVTFFALFLVGCTVAEKTTEDQTAEKRAHPDGQMLRASNWISTRPMMSPDNEIGYL
IQYLEKRASSMENDAESRDIRSPLGTMRFGKRADGPLGTMRFGKRNPLGTMRFGKRNPLG
TMRFGKRAEGPLGTMRFGKRTPDFYDY
>FLP-3_Halicephalobus_mephisto_HMEPH_08204-RA.p1
LEFNMNSQRLFTLVLLVISFIQINAKDESSIAIQKPEDDQDYLETSHFKPSEDILNAYNE
ELQSLIEKRNTNNYQDNNGERNIRSPLGTMRFGKRNPNPLGTMRFGKRNPNPLGTMRFGK
RNPNPLGTMRFGKRNPNPLGTMRFGKRSLEDYELNNTPLGTMRFGKRSFDFYN
>FLP-3_Panagrellus_redivivus_Pan_g5472.t1
MGLYHIHVTLLALCCAAAYATSVNEAAEYDVEEKRSPKLSFKTFSPIVDLESFESDDANL
NKRNVVEMNNQNTQRDTRAPLGTMRFGKRNPLGTMRFGKRASAIGTMRFGKRNPLGTMRF
GKRAVDEFESMKNARFGPMQNYFE
>FLP-3_Parastrongyloides_trichosuri_PTRK_0000786300.1
MHSVSFYSFAFLLVFLVFTTTNVLGNDFRYPSRDVMKAMLRDPKAYFRGEPIRFVTFSSY
LDSAENYTNNMKPMVNPQGIHSRAQNMLLLYPSKSFWALEGANNEAKNLKNTLFSKNKRS
EITNTVTGDNDITFPGRNIRSPLGTMRFGKRDLIQDVPNNYDDMNNWILYYKRSERPSPL
GTMRFGK
>FLP-3_Steinernema_carpocapsae_SC.X.g3561
MNACCFAITLLLCQLVSSETPAKHSPKNELDQEDNDDVRYLLSIVDDWPELQGLKQRTIR
SPLGTMRFGKRDPLGTMRFGKRATLGTMRFGKRNPLGTMRFGKRDPLGTMRFGKRVPLGT
MRFGKRVPLGTIRFGKRDPLGTMRFGKRDPLGVMRFGKRGLPEAAMTPQLAVANAPYSDE
EN
>FLP-3_Steinernema_feltiae_L889_g30047.t1
MNVCCFALTLLLCQLVSAEVPAKHPSNGQVDQDDNDDVRYLLSIVDDWPELEGLHKQRTI
RSPLGTMRFGKRDPLGTMRFGKRDPLGTMRFGKRDPLGTMRFGKRDPLGTMRFGKRDPLG
TMRFGKRVPLGTMRFGKRVPLGTMRFGKRNPLGTMRFGKRSVPLGTMRFGKRSDLRLQLS
KGLNGSKTSVHHQHKTSRFFFFLSNLWSRVILVSLSTSSACSKSN
>FLP-3_Steinernema_glaseri_L893_g22384.t1
MSAVSLSRSFSVNYLPQVTNVPSPHLTRFSEASAKPAAGNELEEDDNDDVRYLLSVVDEW
PQIEGVHQQRSIRSPLGTMRFGKRDPLGTMRFGKRATLGTMRFGKRDPLGTMRFGKRVPL
GTMRFGKRDPLGTMRFGKKRDPLGTMRFGKRASLGTMRFGKRSSEEPASFGQLDGKSSFY
NSEEI
>FLP-3_Steinernema_monticolum_L898_g11233.t1
MNVCCFAFTLLLCHLVSAEAPVKQATNNELDQEDDDVRYLLSVVDEWPELQGLNKQRTIR

SPLGTMRFGKRDPLGTMRFGKRATLGTMRFGKRDPLGTMRFGKRDPLGTMRFGKRVPLGT
MRFGKRDPLGTMRFGKRATLGTMRFGKRDPLGTMRFGKRSVTLGTMRFGKRAIIEPALVQ
>FLP-3_Steinernema_scapterisci_L892_g1313.t1
MNACCFALTLLLYQLVSSETPAKHSPQNELDQEDNDDVRYLLSIVDDWPELQGLNKQRTI
RSPLGTMRFGKRDPLGTMRFGKRATLGTMRFGKRDPLGTMRFGKRDPLGTMRFGKRVPLG
TMRFGKRVPLGTMRFGKRDPLGTMRFGKRDPLGTMRFGKRSLPEPAIASQLALENAPYSN
EEN
>FLP-3_Strongyloides_papillosus_SPAL_0001444800.1
MILFLNKKHSSAFSPFILLLLINVSFSYYFDNTEEYPVNIKPMAVYNKVHSKPQSMYFFY
PYRFGWPMERVNENENKKTITPSKNKRSIINEHKFVDDSEEILNGRNIRSPLGTMRFGKR
GQFNELPLSDEDFSNWILYNKRTMRPNPLGTMRFGK
>FLP-3_Strongyloides_ratti_SRAE_X000120500
MFTLLNNRNSITFLLTIFLIFINVSFAYYFDSFDDYPVKVNQLEMYKKIHSKPQSMYFFY
PARLNWAVDKTNFIENKGDISSNKNKRSNLNENKSDDNESILNGRNIRSPLGTMRFGKRG
KIYDIPMSDEEFNSWIIYNKRTAKPNPLGTMRFGK
>FLP-3_Strongyloides_stercoralis_SSTP_0001244500.1
MYKKLYSKPQNMYFFYPVKLNWALDENTVNKNKRDTSFNKSKKSNFNENKSDNNDSHLNG
RNIRSPLGTMRFGKRGQHNIPTTDEDFKNWIIYNKRTVKPNPLGTMRFGK
>FLP-3_Strongyloides_venezuelensis_SVE_0144300.1
MILFLNEKHSSTFFHFIILLLINVSFTYYFDSTEEYPVNIKPMTVYNKVHSKPQSVYFFY
PYRFGWPMEGVNENKKTITPNKNKRGIINEHKFDDDSEEILNGRNIRSPLGTMRFGKRGQ
LDELPLSDKDFSNWILYNKRTMRPNPLGTMRFGK
>FLP-3_Rhabditophanes_sp_KR3021_predicted
LQGPSRFTNAFTTNYKYPASKDNWIYESDDKIFPFIYDQMISIPNAKRNAPEFAEEPLNKESAISDRD
TRSPLGTMRFGKRSPSLNRLFNEHYEVKRDTHKSNPLGTMRFGRK
>FLP-3_Acrobeloides_nanus_ACRNAN_Path_1124.g4345.t1
MYKARYPDLYKSPVQEDSSRSRRSLIEKEPKDEAQEVANEEDSSFFDILEKRVADLDALR
DSKTPLGTMRFGKRSPNPLGPMRFGKRAPALGPMRFGKRAPTLGPMRFGKRVKALGPMRF
GKRAAAPQFGTMRFGKRTYDDSKIFL
>FLP-3_Ditylenchus_destructor_Dd_08212
MANYSLILTFEFLLIAFFSVFGGHCALVIGNKAKTVYSEPSTGPAQPSISQVSELIQPPF
TYRWLSARPLIQEYYQTHPETDLSYLLQYLEKRSASQLLESPFSEDEAQRDIRSPLGTMR
FGKRNPLGTMRFGKRNGTPLGTMRFGKRQDPGSKENVNYSS
>FLP-3_Globodera_rostochiensis_GROS_g03097.t1
MHLLEMIFKVLPSSLSPFLLLLFLALLFDLSLCQPTSQQKSIQELLNATPLLPSAEQHDS
LGDPFQQQRWLSVRPFQEYRRPQTDELGLLLQYLDNKRNAPSMLTETNGGGGQSSNGSPL
GTMRFGKRKFNSPLGTMRFGKRGHNNAPFGTMRFG
>FLP-3_Heterodera_glycines_Hetgly.G000022201
MQTFVMSKLKVLSSSADFSSFLFLALLFDFSFSQPTIQQKSIQELLNTPTFLPTAEKHDY
FVAPFLQQQKQRWLSVRPFQEFRRSPQTADLGLLLQYLDNKRNAPSLMEDEGNGRHGISG
SPLGTMRFGKRTLNSPLGTMRFGKREYNKSPLGTMRFG
>FLP-3_Meloidogyne_arenaria_M.Arenaria_Scaff14749g085065
XTKVRITSYNVCYTSITTMTIQKISNLIKLFVLIAASFVISNCSPAIPRQETIPSLLPNG
ALHETLTDPFQQQRWISVRPFQDYPRPHNEELGLLLQYLDSKRNAPLLDENVAQLVGESP
LGTMRFGKRRNTSPLGTMRFG
>FLP-3_Meloidogyne_enterolobii_scaffold15333_cov101.g18388
MTNQKISNFIKLFVLIAASFVISNCSPAIPRQETIPSLLPNGALHETLTDPFQQQRWISV
RPFQDYPRPHNEELGLLLQYLDSKRNAPLLDENVAQLVGESPLGTMRFGKRRNTSPLGTM
RFG
>FLP-3_Meloidogyne_floridensis_maker-nMf.1.1.scaf14423-augustus-
gene-0.7-mRNA-1
MSNLIKLFVLIAASFVISNCSPAIPRQETIPSLLPNGALHETLTDPFQQQRWISVRPFQD
YPRPHNEELGLLLQYLDSKRNAPLLDENVAQLVGESPLGTMRFGKRRNTSPLGTMRFG
>FLP-3_Meloidogyne_graminicola_NXFT01001842.1.4783_g

```
MVISNSFEELDKQMYPMWQRQMREPLRFGKRLLKQKTVISNCSPVTRQEIIPSLLPVGTL
HESLNDPFLQQQRWLSVRPFKEYPRPHNEELGLLLQYLDSKRNAPILDENVAQLVGESPL
GTMRFGKRRNSPPLGTMRFG
>FLP-3_Meloidogyne_hapla_MhA1_Contig213.frz3.gene28
MPFTPTQITSTMTIQKVTNPTALFVLFAASIVISNCSPVVPRQELPSLLPTGALHETLND
PFIQQQRWLSVRPFQDYPRPHNEELGLLLQYLDSKRNAPLLDENAAQIVGESPLGTMRFG
KRRNSSPLGTMRFG
>FLP-3_Meloidogyne_incognita_Minc3s00218g07825
MTIQKICNLIKLFVLIAASFVISNCSPAIPRQETIPSLLPNGALHETLTDPFQQQRWISV
RPFQDYPRPHNEELGLLLQYLDSKRNAPLLDENVAQLVGESPLGTMRFGKRRNTSPLGTM
RFG
>FLP-3_Meloidogyne_javanica_scaffold11525_cov308.g15662
MTNQKISNFIKLFVLIAASFVISNCSPAIPRQETIPSLLPNGALHETLTDPFQQQRWISV
RPFQDYPRPHNEELGLLLQYLDSKRNAPLLDENVAQLVGESPLGTMRFGKRRNTSPLGTM
RFG
>FLP-3_Ditylenchus_dipsaci_predicted
ARQKKPTRHHAIRQKIHPLGTMRFGKRGQVNDDDNNQ
GDPRHSLSLGTMRFGKRGSELGKRNPLGTMRFGKRSTH
>FLP-3_Globodera_pallida_predicted
SDLSLCQPTSQQKSIQELLNATPLLPPAEQHDSLGDPFQQQRWLSVRPFQEYRRPQTDELGLLLQYLD
NKRNAPTMLTETDGGGGQSSNGSPLGTMRFGKRKFNSPLGTMRFGKRGHNNAPFGTMRFG
>FLP-3_Haemonchus_contortus_predicted
SSAHGLGAEPGRTVDSYFTSLHFSGPLGTFRFGKRNGGRSTSETLYLNDQFN
>FLP-3_Haemonchus_placei_predicted
SSAHEKGAEPEKSIDSYFTSLHFSGPLGTFRFGKRNGGRSTGETLYLNDQFN
>FLP-4_Romanomermis_culicivorax_predicted
LNHFTGYFEKRKPNFIRFGRGSGNENIKQKFSDKI
KISKKIVRTNYRITNLFFMLDFWDEKRKPTFIRFGRSYYY
TLICMIQLYPKASTYLSVPQKFSTDVRPNFIRFG
>FLP-4_Plectus_sambesii_PSAMB.scaffold42size100754.g952.t1
MKSAAFITGCSCALVALLFAGCCLDEADAFALPRQLEENQQLLRSLFDYEDSLPVRPARQ
WSESLSGRVKGSGKTGKPTFIRFGKRLDTFGKSGKPTFIRFGKRSGLMSRPDLFDDARNE
Q
>FLP-4_Acanthocheilonema_viteae_nAv.1.0.1.t06602-RA
MVQKLLMSYDAVADQKRSYWKTTIEIIKNLSETISPKMAHTQCTGISATIYFALCFGLFI
MTTYTKANVINGEDNEFANENAKDWYAISLRSEPQLRFEHGSRSHGKPTFIRFGKRDYDA
FST
>FLP-4_Anisakis_simplex_ASIM_0002102901-mRNA-1
LLANSMPSLQEAGAGSSGAGSASGVAPPFSRLTRGGKPTFIRFGKRSSGWQQSASAKSN
>FLP-4_Ascaris_lumbricoides_ALUE_0001612501-mRNA-1
MAAFKTTVFSCLVTLMLTLFVVESINAELDNGDITMIEIFQAKDDMLRALYNYILYERKG
AHDFDGFERAQRSGKPTFIRFGKRSANIVTNPENI
>FLP-4_Ascaris_suum_AgR001_g616_t01
MAAFKTTVFSCLVTLMLTLFVVESINAELDNGDITMAKDDMLRALYNYILYERKGAHDFD
GFERAQRSGKPTFIRFGKRSANIVTNPENI
>FLP-4_Brugia_malayi_Bm10067.1
MVQIRCIGTSSIIYFTLCVGLFVLTTFTNANVINEEDDEPISENAKHWYPALLRSEPQFG
FERGSRSHGKPTFIRFGKRYFDAYNPNLFNNK
>FLP-4_Brugia_pahangi_BPAG_0000787401-mRNA-1
MVQIRCIGTSSIIYFTLFVGLFVLTTFTNANVINEEDDEPISENAKHWYPALLRSELQFG
FERGSRSHGKPTFIRFGKRYFDAYNPNLFNNK
>FLP-4_Dirofilaria_immitis_nDi.2.2.2.t00572
MAQNRHIEITITAYSAFFIGFCISAICANASIINEENDEAGNENRKDWYATLLKSKPQFG
FERDSRSHAKPTFIRFGKRDFDAYDPNLFNIK
```

>FLP-4_Elaeophora_elaphi_EEL_0000386201-mRNA-1
MAQTQSNGINSATYFALCVGLFVLTTSCTNANVINEEDNGFNDGNTKTWYALLLRSKPQF
GFERGSRSHGKPTFLRFGKRDFDAYNPDPFKNK
>FLP-4_Onchocerca_flexuosa_X798_01688
FQSFCRKTIRFKIEIIKKFKHEFNLSKTIFLKMAQMRCIGISATACFTVCMGLYILATCA
NATILDDDDDFVNEDAKVWYAALMRSKPQFGFERGSRSHGKPTFIRFGKRDFSTYNPNFF
NK
>FLP-4_Onchocerca_ochengi_OOCN_0000427101-mRNA-1
MAQMRYIRINATACFTVCMGLYILAICANAIIPEDDDYFVNEDTKAWYAALLSSKPQFGF
ERDSRSHGKPTFIRFGKRDFSAYNPNIFNNK
>FLP-4_Onchocerca_volvulus_OVOC10618.1
MARMRYIRINATACFTVCMGLYILAICANAIIPEDDDYFVNEDTKAWYAALLSSKPQFGF
ERDSRSHGKPTFIRFGKRDFSAYNPNIFNNK
>FLP-4_Parascaris_equorum_PEQ_0000455501-mRNA-1
MLYEPKGSHDFDGFERAQRSGKPTFIRFGKRSANIVTNPENI
>FLP-4_Parascaris_univalens_PgR064_g035_t01
TMAAFKITVFSCLITLMLTLYAVESINAELDNGDIRVTKDDILRALYNYMLYEPKGSHDF
DGFERAQRSGKPTFIRFGKRSANIVTNPENI
>FLP-4_Thelazia_callipaeda_TCLT_0000020001-mRNA-1
MVLTRYTRTNVTVRFALYASVFYVLTTYTNAYILNDNDNTIEANHENVKDSYREFLWSRP
QFGYERGARAHGKPTFIRFGKRDSIDHNANFIM
>FLP-4_Toxocara_canis_Tcan_00224.1
MAADNTSFKAMFTFRSVIFSFLGSLTLAILLHTVKAEQDNRQIVITKDDLIRALYAYMLS
EPKLPQDSDEFRRELRSGKPTFIRFGKRIPSLVANSGSN
>FLP-4_Wuchereria_bancrofti_WBA_0000737301-mRNA-1
MWAGSSIQKFIGGLQLSSGPRFVLKTKRFKIEIAKNSSQMISSNMVQTRCIGTSATIYFT
LCVGLFLLTTFTNANVINEEDDEPINENAKPWYPALLRSEPQFRFERGSRSHGKPTFIRF
GKRYFDAYNPNLFNNK
>FLP-4_Dracunculus_medinensis_predicted
IFQVNYQSKFIEFPIIDYDPRMKDFYNRDLRSHGKPTFIRFGKRNENS
>FLP-4_Gongylonema_pulchrum_predicted
HTLYSLSHFINRNFQARENARAWYEVLLKSRPLNGFERGSRSHGKPTFIRFGKRNFYDYNANLFNNR
>FLP-4_Ancylostoma_ceylanicum_Acey_s0533.g3057.t2
MLLLLFDRPCHHSRNAFPVMTLIKLSKGGTEEDMNTQCFVAFCIACIVLVAGFDERVNDA
YEPEPVAADSGFFRNFRSSSNGKPTFIRFGKRAQPSFIRFGRAQPSFIRFGRQATA
>FLP-4_Ancylostoma_duodenale_ANCDUO_14571
MFFPKGYIPDCIPVKASNIAINFINIPERQVFMGSYFGLKGKDSQVPRANFTKDMNMQCF
AAFCIACFVLVAGFDERANDVYEPEAAVADSAFFRNFRSSSNGKPTFIRFGKRAQPSFIR
FGRAQPSFIRFGRQATA
>FLP-4_Angiostrongylus_cantonensis_ACAC_0000690101-mRNA-1
MHWRRYRPGTQYEFKSNFRSAPSGKPTFIRFGKRTQPNFIRFGRAQPSFIRFGKKDATF
>FLP-4_Caenorhabditis_angaria_Cang_2012_03_13_01035.g15443.t1
MSSVQTILLSLIFAILVIFASAQQQQQQAQQQNNLVADDDILTQEASIDGEYTFPLVHLR
GLRSSSNGKPTFIRFGKRASPSFIRFGRK
>FLP-4_Caenorhabditis_ brenneri_CBN06535.1
MNAFSSSLKTFLFSIIFAMLLVLAAGQTSSGEDVADQEPKSLITEFNTPEVVEQTNNFWP
PVHLRGLRSSNGKPTFIRFGKRGSPSFIRFGRK
>FLP-4_Caenorhabditis_briggsae_CBG02703.1
MNAFPSSLKTFLFSILFATLLVYAAGQTPSSEDVEPQEEQQQKELIAPDEYITPEIIEQT
NNFSAVHLRGLRSSNGKPTFIRFGKRASPSFIRFGRK
>FLP-4_Caenorhabditis_elegans_C18D1.3.1
MNAFSSSLKTFIFSLLFATLLALTAAHPPSSGEEIAEQEEKNIASPDELIPEIVEQQNFW
PPVHLRGLRSSNGKPTFIRFGKRASPSFIRFGK
>FLP-4_Caenorhabditis_inopinata_Sp34_20208800.t1

MLLVLSSAQTPSDDVEQLEPKNLEEFETQEVVEPISNFWPPVHLRGLRSSNGKPTFIRFG
KRASPSFIRFGRK
>FLP-4_Caenorhabditis_latens_FL83_12761
MNAFTSSSLKTFLFSILFATLLVLTAGQTPSSAEDVEQIEQQKGFETQDEYITPEIVEQT
NNFWPPVHLRGLRSSNGKPTFIRFGKRASPSFIRFGRK
>FLP-4_Caenorhabditis_nigoni
MNAFPSSLKTFLFSILFATLLVFAAGQTPSGEDVEPQEEQQKGLITTDEYITPEIIEQTN
NFWPLVHLRGLRSSNGKPTFIRFGKRASPSFIRFGRK
>FLP-4_Caenorhabditis_remanei_FL81_08167
MNAFTSSSLKTFLFSILFATLLVLTAGQTPSSGEDVEQIEQQKGFETQDEYITPEIVEQT
NNFWPPVHLRGLRSSNGKPTFIRFGKRASPSFIRFGRK
>FLP-4_Caenorhabditis_sinica_Csp5_scaffold_00609.g13530.t1
MNAFTSLKTLIFSILFATLLVLSAAQTPSTSSAEDVEPEQQLEQQKGLITPEEYITPEII
EQTNHFWPPVHLRGLRSSNGKPTFIRFGKRASPSFIRFGRK
>FLP-4_Caenorhabditis_tropicalis_Csp11.Scaffold629.g7957.t1
MNAFSASLKTLLFSILFAMFLVLTAAETAPSSAEDIVEQDQKSLITTPDEYVQPEVIEQT
NNFWHPMHLRGLRSSNGKPTFIRFGKRASPSFIRFGRK
>FLP-4_Diploscapter_coronatus_DCO_024260
MLYQLFASLCVLALALLVVVSSAATLQNPANVEAQRQPEILEANDDNLPIIEPLEDAISS
VQENQWFDRVDRSGSKPTFIRFGKRAQPAFIRFGKRAAPSFVRFGRK
>FLP-4_Diploscapter_pachys_WR25_20448.1
MLYQLFASLCVLALALLVVVSSAATLQNPANVEAQRQPEILEANDDNLPIIEPLEDAISS
VQENQWFDRVDRSGSKPTFIRFGKRAQPAFIRFGKRAAPSFVRFGRK
>FLP-4_Heligmosomoides_polygyrus_HPBE_0000119601-mRNA-1
MSLEMLIKTTSSSICALVLYVCCVSYAFQCCPPPRDGPKHFGEVKLSWKNADDTQDFEAS
IAFNFRNGKPTFIRFGKRAQPSFIRFGRAQPSFIRFGRRAPDSVMEPNHLRTEVVSSLEN
EILSLSKMGDIPQSSNTSCQ
>FLP-4_Heterorhabditis_bacteriophora_Hba_09074
MLARYFLSLCILFISLKLSTEEQMVPDRLWGQLYKQLPYTLPNDSEGVDDDDGFYRTFRS
ATNGKPTFIRFGKRAQPSFIRFGRAQPSFIRFGKREDTIQTNN
>FLP-4_Mesorhabditis_belari_mbelari.g14243.t1
MIRPFHSSLFLLSFLVLNIESCCDEEKQLKQTLGNRLDELRFQPSFTNQQQIRIPEDVLR
MLAQYDSMRNEQQSPLTRMSRANGKPTFIRFGKRSKPTFIRFGKRSSFKNAEDQWIDGSM
>FLP-4_Parapristionchus_giblindavisi_Parapristionchus-mkr-
S_91-1.169-mRNA-1
MIFKLLVLLLSLIALSFSSTTDNNQYQTGESDIYDDSIISYLVNHPSGMTRMTRSNGKPT
FIRFGKRSSMYSSE
>FLP-4_Pristionchus_arcanus_arcanus-mkr-S_225-0.64-mRNA-1
MLHKLIVLFFALLAAFALASPIDSGYQMGEADSYDGVMPMVQMNGPLARMTRSNGKPTFI
RFGKRSMFNQYE
>FLP-4_Pristionchus_exspectatus_exspectatus-mkr-S_6-4.24-mRNA-1
MLHKLIVLFFALLAAFALASPINSGYQMGEADSYDGVMPMVQMNGPLARMTRSNGKPTFI
RFGKRSMFNQYE
>FLP-4_Pristionchus_fissidentatus_fissidentatus-mkr-S68-4.24-mRNA-1
MLHKMIAFFFALLVASALAGPIDSGYPMVDSDAYDGMMPVMPFNREALSRMTRSNGKPTF
IRFGKRSAYNRFE
>FLP-4_Pristionchus_japonicus_japonicus-mkr-S3-10.64-mRNA-1
MLHKLIVLFLALLAAFALASPIDSGYQMGEADSYDGVIPMVQMNGPLARMTRSNGKPTFI
RFGKRSMFNQYE
>FLP-4_Pristionchus_maxplancki_maxplancki-mkr-S23-2.35-mRNA-1
MLHKLIVLFFALLAALTLASPIDSGYQMGETDSFDGIMPMVQMNGPLARMTRSNGKPTFI
RFGKRSMFNQYE
>FLP-4_mayeri-mkr-S42-2.50-mRNA-1_mayeri-mkr-S42-2.50-mRNA-1
MLHKLIVLFFVLLATLALASPIDSAYQMGEVDSYDAVMPMVMNGGLPRMTRSNGKPTFIR

FGKRSGSMSENDLYE
>FLP-4_Pristionchus_pacificus_PPA39937.1
MLHKLIVLFFALLAAFALASPVDSGYQMSEADSYDGVMPMVQMNGPLARMTRSNGKPTFI
RFGKRSMFNQYE
>FLP-4_Ancylostoma_caninum_predicted
TQPIDSSTYIFLAWNSSPFPYFPFRSSSNGKPTFIRFGKRAQPSFIRFGRAQPSFIRFGRQATA
>FLP-4_Angiostrongylus_costaricensis_predicted
KWCYYRNDLNHYRNFRSASNGKPTFIRFGKRAQPSFIRFGRAQPSFIRFGKKDATF
>FLP-4_Caenorhabditis_japonica_predicted
EDVDRTGFLTPENEITVRDIFSVIVNDIPFSARRTLFLQQLLASSAPSWITVGSFLFFLIFSISVPCS
SSNGKPTFIRFGKRGSPSFIRFGRK
>FLP-4_Cylicostephanus_goldi_predicted
XXXXXXXXXXXXXXXXXHLSRVCIRCSLLFQAIFTVVVFWIIDKIIFKLEERKKCKPAKTSKLNIPHF
RSSNGKPTFIRFGKRSKPTFIRFGRSVPTYSNYESQETQEA
>FLP-4_Micoletzkya_japonica_predicted
IFHCLLSLLFQLVVDSSFSSDNYVLPFNRAPLTRMTRSNGKPTFIRFGKRSSWDA
>FLP-4_Necator_americanus_predicted
NAQNFESRELIISAILLLLFRSNGKPTFIRFGKRAQPSFIRFGRAQPSFIRFGREATA
>FLP-4_Nippostrongylus_brasiliensis_predicted
NAVINMEYYFDFFRSGKPTFIRFGKRAQPSFIRFGRAQPSFIRFGRQSAADVSA
>FLP-4_Oscheius_tipulae_predicted
SLRKYEPLSFRSGAGVASQGKPSFIRFGKRAQPSFIRFGKRAQPSFIRFGRSSSDQY
>FLP-4_Strongylus_vulgaris_predicted
VKYESSSSTNKYEVFLNHSFRMRHTNINLLFSKKIGTLRMLSSRSSNGKPTFIRFGKRAKPTFIRFGR
SGPLSGLENQDLEVA
>FLP-4_Bursaphelenchus_xylophilus_BXY_0584900.1
MKCTVQRVVGILLMLGVLAECISDEETAKRENILKALYRLNDQNSYRLYRASRSNGKPTF
IRFGKRSPPAYDGYEHADFNTQ
>FLP-4_Halicephalobus_mephisto_MSTRG.1068.1.p1
MVRLSSQNMAPLVLVSLVLGLVVAIPVPQQGYQNGEFTNDNELAYMSFLNSRFHDIPYHL
NRASRSNGKPTFIRFGKRSPPDSSRLLEQPSDLEWNMRY
>FLP-4_Panagrellus_redivivus_Pan_g17324.t1
MVHLGTVTTLATVIFSVLIAIISAQGYEPYGYGSGQANFAPDQFFESFREQNPIYNRALR
SNGKPTFIRFGKRSFAYPNKPAYY
>FLP-4_Steinernema_carpocapsae_SC.X.g165
MYVRMLNTIVLFLIGLVLVARCAPVAEENDLFDQLYKRDQLLRSIYQLRADSPYVSSFMR
ASRSNGKPTFIRFGKRSGQPYPAVYDSRA
>FLP-4_Steinernema_feltiae_L889_g24975.t1
MYVRMLNTIALFLIGLVLVANCAPHANLEDDFSDQLFKRDQLLRSIYQLRAASPYVSSFM
RASRSNGKPTFIRFGKRSNPTNAMYDQDV
>FLP-4_Steinernema_glaseri_L893_g16422.t1
MYVRMLNTVALLFIALVLVAKCSPIPEENDSYDQYLMKRDQLLRSIYQLRADSPYVSSFM
RASRSNGKPTFIRFGKRSGSNVDPYDPRA
>FLP-4_Steinernema_monticolum_L898_g13719.t1
MYVSMLNTIALFLIALVLVARCTPNSEENEFYDQLVKRDQLLRSIYQLRAASPYVSSFLR
ASRSNGKPTFIRFGKRSNPDGLYGLQA
>FLP-4_Steinernema_scapterisci_L892_g29948.t1
MYVRMLKTIALFLVGLVLVARCAPVAEENDLFNELYKRDQLLRSIYQLRADSPYVSSFMR
ASRSNGKPTFIRFGKRSGAPYPAYDPRA
>FLP-4_Acrobeloides_nanus_ACRNAN_scaffold9100.g29210.t1
MFFVRAALIDESEPSAYLDKRGDLARALYELQADPRNNYQNYDRALRSNGKPTFIRFGKR
SFLGEYPTQQDKL
>FLP-4_Ditylenchus_destructor_Dd_10388
MIATLISLSILCFVVAAQAHPSGYPVSVQSDDDESPAAVDPALWLRAQYAHRNPETAMRE

AALLRAMYQAAMMRGEPQDEFDIASDPRLYRASRSNGKPTFIRFGKRSATEPKDKKAFEP
>FLP-4_Ditylenchus_dipsaci_predicted
RPAMQIAKPVSAPAQTAPVSSDMKLYRASRSNGKPTFIRFGKRAGAKRSIAH
>FLP-5_Ascaris_lumbricoides_ALUE_0001083601-mRNA-1
MLSRLQLVVLALALCLLQVTTSQLSETFDRDERSPKQKFIRFGRAGSARFIRFGRSSPDL
WDESKGSSEIDEMKRAGPRFIRFG
>FLP-5_Ascaris_suum_L3E_02339 peptide: L3E_02339
XSETFDRDERSPKQKFIRFGRAGSARFIRFGRSSPDLWDESKGSSEIDEMKRAGPRFIRF
G
>FLP-5_Dracunculus_medinensis_DME_0000743901-mRNA-1
MFDTLTESVVHTISALIMSKRMMDAYERDARAPKAKFIRFGRNNNMKFIRFGRSFNDRES
DEDGLAGKNKFDQRMFQR
>FLP-5_Parascaris_equorum_PEQ_0000460101-mRNA-1
MKDSNKQQLHLFLTITSEITQYIEIQTSISDETFDRDERSPKQKFIRFGRAGSARFIRDI
GGAIFIEIGGLLFILWRILIDYCVRRVSSQ
>FLP-5_Thelazia_callipaeda_TCLT_0001024001-mRNA-1
MATSDQWLLLLSKVIIFNLIKQKQNKKGVTFVERNERAPKQRFIRFGRRDLRFPDDSRNS
RIPLDWNGEDLQALNAAEMKRMGSRFIRFG
>FLP-5_Toxocara_canis_Tcan_18845.1
MLSRLQLVVLALALCLVQLATSQLSETFDRDERAPKQKFIRFGRAAGARFIRFGRSYPDQ
WILSETNVNTNRLNAFLRPDRINMQPGRP
>FLP-5_Anisakis_simplex_predicted
CEFRLETYERDLRAPKPKFIRFGRGAGARFIRFGRSSPYDQWVCC
>FLP-5_Gongylonema_pulchrum_predicted
SRFSGATFFDREVRAPKQRFIRFGRSXXXXI
SRFSDATFFDREVRAPKQRFIRFGRSGPRYLEFGSNSPTDWVSKSSRHAFLKKAINCLKYGVFSFCYF
ACLKTTSHKHCSKVRKVF
>FLP-5_Parascaris_univalens_predicted
CFRSETFDRDERSPKQKFIRFGRAGSARFIRFGRSSPDLWVRL
>FLP-5_Ancylostoma_ceylanicum_Acey_s0015.g2820.t1
MWAELGRGAEVTTGPPIIIAAFEMSSRHTYAVLFIASILVLQYVTAQSDETYEFHRDARA
PKFIRFGRGGGAKFIRFGRSGTNTWLHEITCVNVTYLATCCSSELFQENDMTDYEGGLGM
LREDKRAAKFIRFG
>FLP-5_Ancylostoma_duodenale_ANCDUO_17186
MYVMAIFPEEPRDSYETYEFHRDARAPKFIRFGRGGGAKFIRFGRSGTNTWRLTSTHSEI
PLHKYTKLQYPYTTDLPENS
>FLP-5_Angiostrongylus_costaricensis_ACOC_0000269901-mRNA-1
MSSQHTTYAALFIVAIFMLQYATTESDDRYVRQRIARAPKFIRFGRGGGAKFIRFGRSDN
NMKDDEIDEAEYYDDAHQDDKRAAKFIRFG
>FLP-5_Caenorhabditis_angaria_Cang_2012_03_13_00282.g8428.t1
MSSQRTTIAIFFIASLFILQFVSADNSEEEAEYSADKYQRIARAPKPKFIRFGRAGAKFI
RFGRSGANTWDDEYVAPSNEVYVKRGAKFIRFG
>FLP-5_Caenorhabditis_brenneri_CBN09712.1
MRNVPAFQLRLPKSNRLPSNRSFLATMSSRSTTIAFLFLATLLIFQCVSAQSSDEDTDFI
DKYQRIARAPKPKFIRFGRAGAKFIRFGRSGANTWEDGYAAAAPSVNELYVKRGAKFIRF
G
>FLP-5_Caenorhabditis_briggsae_CBG22794.1
MSSRSTTIAFLFIATLLVFQCVSAQLSDEETSFLDQYQRVARAPKPKFIRFGRAGAKFIR
FGRSGANTWEDGYAAAAPSVNDLYVKRGAKFIRFG
>FLP-5_Caenorhabditis_elegans_C03G5.7.1
MSSRSTTIAFLFIATLLVFQCVSAQSSAEDADYLEKYQRIARAPKPKFIRFGRAGAKFIR
FGRSRNTWEDGYASPSVNELYVKRGAKFIRFG
>FLP-5_Caenorhabditis inopinata_Sp34_X0166300.t1
MSSRSTTIACLFIATLLVFQSVSAQLSDEEADYLDKYQRIARAPKPKFIRFGRAGAKFIR

FGRSGGNTWEDGYAAPSVNELYVKRGAKFIRFG
>FLP-5_Caenorhabditis_japonica_CJA09174.1
MSSRSTTIAFLFIATLLVFQCVSAQSAEEADYMADKYERIARAPKPKFIRFGRAGAKFIR
FGRAGPSTWEDGFQVPSANDLYVKRGAKFIRFG
>FLP-5_Caenorhabditis_latens_FL83_02033
MSSRSTTIAFLFIAILLVFQCVSAQSSDEDSEYLDKYQRIARAPKPKFIRFGRAGAKFIR
FGRSGANSWEDGYAAPSVNELYVKRGAKFIRFG
>FLP-5_Caenorhabditis_nigoni_Cni-
MSSRSTTIAFLFIATLLVFQCVSAQMSDEETSFLNQYQRVARAPKPKFIRFGRAGAKFIR
FGRSGANTWEDGYAAAAPSVNDLYVKRGAKFIRFG
>FLP-5_Caenorhabditis_remanei_FL81_03393
MSSRSTTIAFLFIATLLVFQCVSAQSSDEDSEYLDKYQRIARAPKPKFIRFGRAGAKFIR
FGRSGANSWEDGYAAPSVNELYVKRGAKFIRFG
>FLP-5_Caenorhabditis_tropicalis_Csp11.Scaffold629.g11000.t1
MSSRSTIAFLFIATLLAFQCVSAQSSEEDAEYLDKYQRIARAPKPKFIRFGRAGAKFIRF
GRAGANTWEDGYAQPSVNDLYVKRGAKFIRFG
>FLP-5_Diploscapter_coronatus_DCO_024726
MGVARSLVLTIIASLIILDVVYSQSDDEELVRIMRSPERNKFIRFGRGGAKFIRFGRSSS
STWEDPSQLEALDETVEVKRAAGGAKFIRFG
>FLP-5_Haemonchus_contortus_HCON_00164350
MSPRYTYAVVFIASLLVLQYVAAQSDDMYELQRVVRAPKFIRFGRGGGAKFIRFGRSGTN
TWDDDSSDITSYAHQDDKRAAKFIRFG
>FLP-5_Haemonchus_placei_HPLM_0001454701-mRNA-1
MSSWSFFASIGAFADKVGLSAKTWSLFQKIEEFFSDDMYEPQRVVRAPKFIRFGRGGGAK
FIRFGRSGTNTWVSR
>FLP-5_Heligmosomoides_polygyrus_HPOL_0001998501-mRNA-1
MSSRHTYAVLFIASLLVLQYVAAESDDVYEMQRVARAPKFIRFGRGGGAKFIRFGRSGTN
TWVSGLIRRLECDSKVLNEPGG
>FLP-5_Mesorhabditis_belari_mbelari.g16204.t1
MISIRTLLFATLLLVLAVLCDMAAAQSDEETQFLRQFRAPSPKFIRFGRAGQKFIRFGRS
ANTWDDSALDDPEGIDLAMAKRAGQKFIRFG
>FLP-5_Micoletzkya_japonica_MicoRS5524-mkr-S48-10.9-mRNA-1
MAAASSLFRPLCLFLALSCLLQLTAAQGEQLETYVRPMRAPKLIRFGRSGAKLIRFGKRA
DPEYEADDAQLNDDLFDAIKRAGPKVIRFG
>FLP-5_Necator_americanus_NECAME_00547
MWAGRGRDVEMTTGPPIIIAAFEMSSRQTYAVLFIASILVLQYVTAQSDDMYEFQRAARA
PKFIRFGRGGGAKFIRFGRSGSNTWVS
>FLP-5_Oscheius_tipulae_OTIPU.nOt.2.0.1.t04297
MSASQHYLAIFFFCSILVLQMVVAQSSDETEGGPSPRYVRTPKAKFIRFGRAGAGGAKFI
RFGRSGANTWDEDVLETADGLIREEMAKRAGAKFIRFG
>FLP-5_Pristionchus_entomophagus_entomophagus-mkr-S358-0.20-mRNA-1
MSPLPSPLILLLLSSLVLAAFAQSDDDSYYREARAPKLIRFGRAGPKLVRFGKRSDPTMN
AEEYDNTLNEMLAAYKRGGPKLVRFG
>FLP-5_Pristionchus_exspectatus_exspectatus-mkr-S_61-2.12-mRNA-1
MTSSSFHLLSTSYEGGAGIPFPPLHFSEDGSYYRDVRAPKLIRFGRAGPKLVRFGKRSDP
MMNEEYDNVMNEMISEYKRGGPKLVRFG
>FLP-5_Pristionchus_fissidentatus_fissidentatus-mkr-S4-4.28-mRNA-1
MASSSPIVLLLVASLLLHTITAQNYDDFSRDARAPKLIRFGRAGPKLVRFGKRSDPNLNA
EEYDNILNDMYYGSADKRGGPKLVRFG
>FLP-5_Pristionchus_maxplancki_maxplancki-mkr-S2-16.0-mRNA-1
FSSPFGVLPSLYIQTGQSVMSSLPSPIIMLIISSLLFATISAQSEDGSYYRDVRAPKLIR
FGRAGPKLVRFGKRSDPAMNEEYDNALNEMISEYKRGGPKLVRFG
>FLP-5_Pristionchus_mayeri_mayeri-mkr-S517-0.14-mRNA-1
MSPLPSPLIILLISSLLVSVITAQSDDEGYYREARAPKLVRFGRGGPKLVRFGKRSDPSM

NTEEYDDMTAYKRGGPKLVRFG
>FLP-5_Teladorsagia_circumcincta_TELCIR_01782
MSPRYTYAVVFIASLLVLQYVAAQSDDMYELQRVARAPKFIRFGRGGGAKFIRFGRGGTN
TWLLTGGLLDRVEGGLALDVIQALMAIMTNEIQQKCLADLLEIGDTDDDTSDDIGYAHQD
DKRAAKFIRFG
>FLP-5_Ancylostoma_caninum_predicted
CISLFYVHFHTRCTYYFSDETYEFHRDARAPKFIRFGRGGGAKFIRFGRSGTNTWVG
>FLP-5_Caenorhabditis_sinica_predicted
INTPLHFFFSDEETDYLDKYQRVARAPKPKFIRFGRAGAKFIRFGRSGANTWVSDSVAIQRLHKEQKS
KTIAFPVVLVHKALRRLWREILNPI
>FLP-5_Dictyocaulus_viviparus_predicted
TIFIYYSDDNKYDRQRITRTPKFIRFGRGNGAKFIRFGRNDKNTVRSTQ
>FLP-5_Diploscapter_pachys_predicted
SNLSDDEELARIMRSPERNKFIRFGRGGAKFIRFGRSSSSTWVSLKFKPELLIII
>FLP-5_Heterorhabditis_bacteriophora_predicted
KSTRSRTRKGKLLFSEDEFEYQRLARAPKPKFIRFGRGGGAKFIRFGRSVSNAWVGAIATFF
>FLP-5_Nippostrongylus_brasiliensis_predicted
LIHTIYTFSDDMYDMQRVARAPKFIRFGRAGGAKFIRFGRSGSNTWVS
>FLP-5_Oesophagostomum_dentatum_predicted
XXXXXXXXXXXXXXXXXXXXXXXXXXXXXXXXXXXXXXXXXXXXXXXXXXXXXXXXXXXXXXXXXXAA
RAPKFIRFGRGGGAKFIRFGRSGSNTWVSLHEICFIRELNLLLSL
>FLP-5_Pristionchus_arcanus_predicted
LIQYSFHFSEDGSYYRDVRAPKLIRFGRAGPKLVRFGKRSDPTMNEVPILIYFTSSFSTRMNSVLSH
>FLP-5_Pristionchus_japonicus_predicted
VASAAASERETNDTKQQLLILLSRTDHDSNYQNSSFRDDDSSYYRDVRAPKLIRFGRAGPKLVRFGKR
SDPEMNAEVINSI
>FLP-5_Strongylus_vulgaris_predicted
PILTKLKSHYNYFSDEIYEFQREARAPKFIRFGRGGGAKFIRFGRSGTNTWVSLENLIPYLHLNYMQF
PCEGMENMTAAEYSNDYTYIHYLASSCVQ
>FLP-5_Bursaphelenchus_xylophilus_BXY_0123800.1
MVASLRSLCLRLFAVLCLVVLMVSAAPVDESEGYLREARAPKAKFIRFGRAGQKFIRFGR
GGAYGPGTAFPSDLYYQY
>FLP-5_Halicephalobus_mephisto_MSTRG.4679.1.p1
MLGLSSATSRLFSIFCVLFVICCLVAESEAQSLERGVRTPKAKFIRFGRAAGGQKFIRFG
RSSWDYGDGDVPEMELEEPEFVPVLRTFKRAGQKFIRFGRK
>FLP-5_Panagrellus_redivivus_Pan_g3321.t1
MASWIAWHSARQTVSTLCFLVLIAALLLDCNAQSLERGIRAPKNKFIRFGRSGQKFIRFG
RNSAIANDDLEPRRVAYEDNLFEQSPFFKQLNYNKRAGQKFIRFG
>FLP-5_Parastrongyloides_trichosuri_PTRK_0000848300.1
MTVMKYINIFCLLLIIYTTLATCTPISDDFERETRGGQKLIRFGRGGQKLIRFGRSSNPP
GPPSDIEGEEGVVYEDIDPGFFFNEYNPYKRGGQKLIRFGRK
>FLP-5_Rhabditophanes_kr3021_RSKR_0000689800.1
MFFRFNRDQRGGQKLIRFGRAGQKLIRFGRSGRFQGFGDQTEDERQPYEQTYYNPFSTFK
RGGQKLIRFG
>FLP-5_Steinernema_carpocapsae_L596_029879
MSASWIRILLFFVVALVAVQEAVGKNMERDARAPKPKFIRFGRGGGGQKFIRFGRSGAAR
QVLNPQDIYANQFYDPLEVDSAASFLDSAALAAPANDWQQFKRGGPKFIRFG
>FLP-5_Steinernema_feltiae_L889_g21160.t1
MSSSWIRILLLVVVGLLAIQAQSLERNARAPKPKFIRFGRSGSGGQKFIRFGRSGGHHPQ
LSLQDLDANQFYDPLATFDDLSDAEPAPMVAEADDWQQFKRGGPKFIRFG
>FLP-5_Steinernema_glaseri_L893_g17861.t1
MLNSPFARFSALNDHLMFYVAMSFGQKITFPSSSPIPPASKMPSTVSDTLSDPLPNSFGC
GLAARGVVSQRRNPTWSGCKRVYKFVEVRRRSAFSDPLSSHTDYSSLRYEMSSSWIRILL
LVVVALLALGESLERDARAPRQKFIRFGRAGQKFIRFGRSGPQAHQLNLQELLAANQIAE

SLNPVDDLPELSAPAEDDWQQLKHPKFFRFE
>FLP-5_Steinernema_monticolum_L898_g20238.t1
MSSSWIRILFVVVFTLLAIQAHGQSLERNARAPKPKFIRFGRASGGQKFIRFGRSGQQAQ
LNPQDIYANQLYNDPLTSLDDLSEVEAAPVDDWQQFKRGGPKFIRFG
>FLP-5_Steinernema_scapterisci_L892_g13652.t1
MSTSWIRIVLLFVVALVAIQEAQGQNMARDARAPKPKFIRFGRGGGGQKFIRFGRSGAAR
QELDPQDIYANQFYDPLGMDSAFLDSAAPVAPAAGANDWQQFKRGGPKFIRFG
>FLP-5_Strongyloides_papillosus_SPAL_0000432400.1
MIEGVGRETRGGQKLIRFGRGGQKLIRFGRSLNPPTKLDIEKEEGIVYDENEPNYNSEDY
SFFKRGGQKLIRFGRK
>FLP-5_Strongyloides_ratti_SRAE_2000398400
MYVPAIEGIGRESRAGQKLIRFGRGGQKLIRFGRSSNLPPPTDIENEEGTVYDENEPNLY
FGDYNPYKRGGQKLIRFGKK
>FLP-5_Strongyloides_stercoralis_SSTP_0000086700.1
MSIVKFVKLIYLFVLFFIIISTFIPVIESAGRESRAGQKLIRFGRAGQKLIRFGRSLNPP
PPADIANEEGIVYDEHDPDFHFNDYNAFKRGGQKLIRFGKK
>FLP-5_Strongyloides_venezuelensis_SVE_0286600.1
MIEGVGREIRGGQKLIRFGRSGQKLIRFGRSLNPPTKLDLEKEEGIVYDENEPNYNSEDY
SFYKRGGQKLIRFGRK
>FLP-5_Acrobeloides_nanus_ACRNAN_scaffold181.g6722.t1
MFMPAIVRNRNWTSLLIVLFMLVCLFLQTQAQSMERNLRAPQPKFIRFGRAGQRFIRFGR
SSWDYDTEPGVSNVDDDNVPIILKRAGQRFIRFG
>FLP-5_Ditylenchus_destructor_Dd_09925
MYEHEDGAPNSIPNYLVAQLLGIVEFISLKGSMMSISRYFSANSLKNIFPKSTFLWLAIF
AVVILMVGSPSVRAAPTEQQQDAVQELNGFDRDARAPKPKFIRFGRAGQKFIRFGRSGNT
MDYVNDFDPATLTAEAEFLAPQFRQFQTAPKRAQKFIRFGK
>FLP-5_Globodera_rostochiensis_GROS_g13026.t1
LVDLPDVQPIQIHLYVFRIARVIATKQQRITQFSPGDNESRWGGANSLPIGSFARDELAV
VMRMLIASGVTALSRRRKPIAFSQISYPLSTRRYLPSSELTCSTKFILILFLDQKMSCCA
SPANRPFARRFFSEITTFYVIAAILLLCIQSSEQQIRADSDMDGWFDRQLRVPKPKFIRF
GRAGQKLIRFGRSSTAPNYSDELSQLDALVDAVDELYPSSPELRAFKSSPKRAQKFIRFG
>FLP-5_Meloidogyne_arenaria_M.Arenaria_Scaff11047g075794
MALAEKAINEVSSPFKNWFDRDTRSPKPKFIRFGRSAGNNQKFIRFGRTPSLELVGGDPS
TEVENLDDLIDAVEGLYPSERLRQQQESPSMTAVYLTAPKRAQKFIRFGKK
>FLP-5_Meloidogyne_enterolobii_scaffold6910_cov176.g10022
MALAEEAINEVSSPFKNWFDRDTRSPKPKFIRFGHPSTEVENLDDLIDVVEGLYPSERLR
QQQESPAMTAVYLTAPKRAQKFIRFGKK
>FLP-5_Meloidogyne_floridensis_scf7180000424033.g12070
MALAEEAINEVSSPFKNWFDRDTRSPKPKFIRFGRSAGGDPSTEVENLDDLIDAVEGLYP
SERLRQQQESPSMAAVYLTAPKRAQKFIRFGKK
>FLP-5_Meloidogyne_graminicola_NXFT01000452.1.112_g
ALAEESVNDVLSPYKNWFEREARAPKPKFIRFGRASEKNQKFIRFGRSLPSLEEYGEDPT
IQEENLDELIDVVEGLYPSERLRQQQQQLSPLTTVYLTAPKRAQKFIRFG
>FLP-5_Meloidogyne_incognita_Minc3s00929g18976
MALAEEAINEVSSPFKNWFDRDTRSPKPKFIRFGRSAGNNQKFIRFGRTPSLELVGGDPS
TEVENLDDLIDAVEGLYPSERLRQQQESPSMAAVYLTAPKRAQKFIRFGKK
>FLP-5_Ditylenchus_dipsaci_predicted
SRQQSTKAQIHQIGRAGQKFIRFGRSGNSWVSVLLY
>FLP-5_Globodera_pallida_predicted
FRVNLLNHSDMTATSNLLLFSMKYIHIGQFEHCSEYASFFRFDRQLRVPKPKFIRFGRAGQKLIRFGR
SSTAPVLISKLFFVEKFQILELQR
NYSDELSQLDALVDAVDELYPSSPELRAFKSSPKRAQKFIRFG
>FLP-5_Heterodera_glycines_predicted
FSLRIKCKIKQYGDEGINQLDALVDAVDKLYQSSPEIRAFKTAPKRAQKFIRFG

IKACFRYGARQLRAPKPKFIRFGRSGQKMIPFSRSFSTETVILLNNFL
>FLP-5_Meloidogyne_hapla_predicted
ELGDPSTEVENLDDLIDVVEGLYPSERLKQQQESPAMAAVYLTAPKRAQKFIRFGKK
RFDRDTRSPKPKFIRFGRSAGNNQKFIRFGRSSSPSFVIFLFLN
>FLP-6_Plectus_sambesii_PSAMB.scaffold388size53591.g5400.t1
MPQVASLILAAFTVACAFAIPTSDADFNINTVTSICDVYPTFSGCSSSIGEVDKRKSAYM
RFGRAMSADEQSEMEKRKSAYMRFGKREQMDGDEANMAGLAEKRKSAYMRFGKRDVPESD
LSLPEKRKSAYMRFGKRSADNTMTDTDSASFDMEKRKSAYMRFGRR
>FLP-6_Acanthocheilonema_viteae_nAv.1.0.1.t09069-RA
MSSVLSLLMVMMMIIHGGVEIVECIEMYENESEIADRKIRILCSVNPNFDICPEHAMEKR
KSSYMRFGRSYPAMLEVEPNMNEKRKSAYMRFGKRYVDSNDYVKRKSAYMR
>FLP-6_Ascaris_lumbricoides_ALUE_0001429201-mRNA-1
MMLQSALYMTLFGAVCALRVLTKEESEPQLVSDNDPTMICELYPQLELCTDAHLMDKRKS
AYMRFGRSDSSAIRGDAEEVEKRKSAYMRFGKRDDSASSLSDNGQTYDGEIEKRKSAYMR
FGKRKSAYMRFGKRSDEQPTAEIEKRKSAYMRLVITKAT
>FLP-6_Ascaris_suum_AgR023_g177_t03
LSNRSTKLYKWRNNFNISLRRNIEMMLQSALYMTLFGAVCALRVLTKEESEPQLVSDNDP
TMICELYPQLELCTDAHLMDKRKSAYMRFGRSDSSAIRGDAEEVEKRKSAYMRFGKRDDS
ASSLSDNGQTYDGEIEKRKSAYMRFGKRKSAYMRFGKRSDEQPTAEIEKRKSAYMRFGRR
>FLP-6_Brugia_pahangi_BPAG_0000471501-mRNA-1
MPSVPALLTVMMMIIYGGVEVVKCLEMFENDPENVDREIRTLCSLNPTFNICAEQAMDKR
KSSYMRFGRSYPAILETEPHLSEKRKSAYMRFGKRYVDFNDYVKRKSAYMRYYF
>FLP-6_Dirofilaria_immitis_nDi.2.2.2.t10106
MSTLVVSLMVIMMVIYSGVEIVKSLEMYEDDVEMNDEEIRTLCRINPKLNICPVVILLQF
GKRYVDNNNDFTKRKSAYMRFGR
>FLP-6_Elaeophora_elaphi_EEL_0001074401-mRNA-1
MTSVLALVLVVMMVIYGNVEVAECLEMYENDPEIVDREIRILCNLNPAFDICPEHDMEKR
KSSYMRFGRSHPTIVEVEPHLNEKRKSAYMRFGRRSVDSNEYVKRKSAYMR
>FLP-6_Litomosoides_sigmodontis_nLs.2.1.2.t08114-RA
MRFISQII*IQRYQMQSMVVLLMTMMMIIYGGVECLETYEEEPEISDREIRALCNLNPNF
SICFEYALEKEKLNQMRFGRQPFAIQGIEPHLYEKRKSAYMRFGKRYVDFNDDVKRKSAY
MRFGR
>FLP-6_Loa_loa_EN70_5478
MPSVLVLLMVMMVILYGNVEIVECLEMDENDPESIDREIRTMCHLNPTIDICAENAIEKR
KSSYMRFGRLYPAIFAAEPHLSEKRKSAYMRFGKRYVDSNDYVKRKSAYMRFGR
>FLP-6_Onchocerca_flexuosa_X798_00667
MNILYAKVTVYQMPTVVVLLMVTMMVIYSGVEVVESLQMYENDPGMNEGEIRTLCSLNPT
LSFCSEHAMEKRKSSYMRFGRSYPVILDLEPYPLEKRKSAYMRFGKRYTDSNDFTKRKSA
YMR
>FLP-6_Onchocerca_ochengi_OOCN_0000037801-mRNA-1
MPTIVVLLMVTMMAIYSGVEVVESLQMYENDPEMNEGEIRTLCSLNPTLSFCSEHAMEKR
KSSYMRFGRSYPVILDIEPYPFEKRKSAYMRFGKRSTDSNDFTKRKSAYMR
>FLP-6_Onchocerca_volvulus_OVOC6277.1
MPTIVVLLMVTMMAIYSGVEVVESLQMYENDPEMNEGEIRTLCSLNPTLSFCSEHAMEKR
KSSYMRFGRSYPVILDIEPYPFEKRKSAYMRFGKRSTDSNDFTKRKSAYMRFGR
>FLP-6_Parascaris_equorum_PEQ_0001361001-mRNA-1
MIILKRNVEMMLQRALCMTLFGAVCAFRLXXXXRGIGATTCLRKRSYDDLFGKRDDSAPS
LSDDEQTDGGEMEKRKSAYMRFGKRKSAYMRFHKFVFNIVCLIFCCISSIYFVNDLFVSI
>FLP-6_Thelazia_callipaeda_TCLT_0000606901-mRNA-1
MVSMLSPILITMIIFGGLKIGECILMNENESEITEEDLRVLCQLHLLNKLCYEHAVEKRK
SAYMRFGRSYHALPRTDLSAMAKRKSAYMRFGKRLSNYNNEEIHDQVKRKSAYMR
>FLP-6_Toxocara_canis_Tcan_03851.1
MAETSSSTLPSRPLFVGRKMALQMALFVALFGAACALPMVIKEDLAPQVISDDDLVMMCE
FYPQLELCTNAYLMDKRKSAYMRFGRSDPLVMTGDGGEVEKRKSAYMRFGKRDDAPSSLS

DGAPSYDGAEVEKRKSAYMRFGKRKSAYMRFGKRSEDETKAEAEKRKSAYMRLAEVKRMM
R
>FLP-6_Wuchereria_bancrofti_maker-PairedContig_2527-snap-gene-0.2-mRNA-1
MPSVLALLMVMMMIIYDGVEVVKCLEMYENDPENVDREIQTLCNLNPTFNICADQAMDKR
KSSYMRFGRSYPAILETEPHLSEKRKSAYMRFGKRYVDFNDYVKRKSAYMRMQRMKEKKT
QLTIYA
>FLP-6_Anisakis_simplex_predicted
LNQFIAVHSVDKRKSAYMRFGRADPALVDSTDLGIDKRKSAYMRCVAHFIIK
>FLP-6_Brugia_timori_predicted
LMIKFLEQAMDKRKSSYMRFGRSYPAILETEPHLSEKRKSAYMRFSFLFFFNFIF
FISSKNFRFGKRYVDFNDYVKRKSAYMRYCF
>FLP-6_Gongylonema_pulchrum_predicted
VKPILKVSSHASNANLQQLTIKSPNDACQLKCIVIYSIYHSDTEKKSILEHAMDKRKSAYMRFGRSLS
TSSDYDSYPIEKRKSAYMRHILGDFEQTAFCLFKKFSAIKLFS
>FLP-6_Ancylostoma_caninum_ANCCAN_10786
MEKRKSAYMRFGRSDPELADQFLMDKRKSAYMRFGKRSEIPNEESLDMEKRKSAYMRFGK
RKSAYMRYV
>FLP-6_Ancylostoma_ceylanicum_Acey_s0147.g2600.t1
MNRTVVAALLLTFAVAYAFQSDDQFSGMEKRKSAYMRFGRSDPELADQFLMDKRKSAYMR
FGKRSEVPNEESLDMEKRKSAYMRFGKRKSAYMRFGKRFSEFDDGSEPFDMEKRKSAYMR
FGR
>FLP-6_Ancylostoma_duodenale_ANCDUO_00455
MEKRKSAYMRFGRSGPELADQFLMDKRKSAYMRFGKRSEIPNEESLDMEKRKSAYMRFGK
RKSAYMRYV
>FLP-6_Angiostrongylus_cantonensis_ACAC_0000672301-mRNA-1
MEKRKSAYMRFGRSIPDFDNQLTMDKRKSAYMRSASFFQINILVC
>FLP-6_Caenorhabditis_angaria_Cang_2012_03_13_00257.g8008.t1
MSTYQWVLLCVVVTLAAAQFDGEKEKRKSAYMRFGRSDPEMMEKRKSAYMRFGKRSGDEG
VEVEDEDHDMMKRKSAYMRFDACFYFFRFGKRSAAEELDMEKRKSAYMRFGKRSIDDGLE
MEKRKSAYMRFGKRSFEEGVEEPEEREMEKRKSAYMRFGKRSFNEDLDMEKRKSAYMRFG
R
>FLP-6_Caenorhabditis_brenneri_CBN06668.1
MNSRGLILMLGVIVAVALAQQDSDLEREMEKRKSAYMRFGRSDGGNPMEMEKRKSAYMRF
GKRSSGGDEQESAVGAGDDMEMEKRKSAYMRFGKRSETPEEDVMSAEKRKSAYMRFGKRS
SDMEMLGDGYNGEEGHDLFKRKSAYMRFGKRSMGEEEDHDMAKRKSAYMRFGK
>FLP-6_Caenorhabditis_briggsae_CBG19281.1
MNSRGLIILMLGVVIAVAVAQQDSDLEREMEKRKSAYMRFGRSDGGNPMEMEKRKSAYMR
FGKRSGDEQEDIVGAGGDDMEMTKRKSAYMRFGKRSGAPEEDVMSAEKRKSAYMRFGKRS
SDMEILGNEGIDGAAEDESHDLFKRKSAYMRFGKRSMGQEEDHDMMKRKSAYMRFGR
>FLP-6_Caenorhabditis_elegans_F07D3.2.1
MNSRGLILTLGVVIAVAFAQQDSEVEREMMKRKSAYMRFGRSDGGNPMEMEKRKSAYMRF
GKRSSGGDEQELVGGDDIDMEKRKSAYMRFGKRSGPQEDDMPMEKRKSAYMRFGKRSSDM
EVIGNEGVDGDAHDLFKRKSAYMRFGKRSMGEEEDHDMMKRKSAYMRFGR
>FLP-6_Caenorhabditis_inopinata_Sp34_50246500.t1
MNSRGLVLMLGIVVAVALAQESSDLEREMEKRKSAYMRFGRSDGGSAMEKRKSAYMRFGK
RSGSGDEQEELFGGDGIEMEKRKSAYMRFGKRSEIPEEDAMSAEKRKSAYMRFGKRSADP
SDLDFLGREDEPIRDLFKRKSAYMRFGKRSIGMEEDHDMMKRKSAYMRFGR
>FLP-6_Caenorhabditis_japonica_CJA08901.1
MNSRGLNLLIGVIVAVTFAQQISDLEREMEKRKSAYMRFGRSDGGGNPMEMEKRKSAYMR
FGKRSSPNEEEFAGNDDIEMAKRKSAYMRFGKRSETPEDDVLSAEKRKSAYMRFGKRSSD
MDIGAEEAVNGFEDGHDLFKRKSAYMRHYRFGKRSMLEEDEHDMMKRKSAYMRFGK
>FLP-6_Caenorhabditis_latens_FL83_20718
MNSRGLILMLGIVIAVVLAQQDAGEMEKRKSAYMRFGRSDGGNPMEKRKSAYMRFGKRSG

GDLEEVDETGNDMQMMEKRKSAYMRFGKRSALPEEDVMSAEKRKSAYMRFGKRSAVEMDE
GEVQDAHDLFKRKSAYMRFGKRSADMDEEEQQDHDMEKRKSAYMRFGR
>FLP-6_Caenorhabditis_nigoni_Cni-
MNSRGLIILMLGVVIAVAVAQQDSDLEREMEKRKSAYMRFGRSDGGNPMEMEKRKSAYMR
FGKRSSGDEQELVGGDDMEMMEKRKSAYMRFGKRSGGAPEEDVMSAEKRKSAYMRFGKRS
SDMEMLGNEGIDGAAVDETHDLFKRKSAYMRFGKRSMGQEEDHDMMKRKSAYMRFGR
>FLP-6_Caenorhabditis_remanei_FL81_15239
MNSRGLILMLGIVIAVVLAQQDAGEMEKRKSAYMRFGRSDGGNPMEKRKSAYMRFGKRSG
DLEEVDETGNDMQMMEKRKSAYMRFGKRSALPEEDVMSAEKRKSAYMRFGKRSAVEMDEG
EVQDAHDLFKRKSAYMRFGKRSADMDEEEQQDHDMEKRKSAYMRFGR
>FLP-6_Caenorhabditis_sinica_Csp5_scaffold_00030.g1658.t1
MNSRGLILMLGIVVAVAVAQQDSELEREMEKRKSAYMRFGRSDGGNPMEMEKRKSAYMRF
GKRSSGDEQELVGGDDMEMEKRKSAYMRFGKRSGTPEEDVMSAEKRKSAYMRFGKRSSDM
EMLENEGLDGAAASGDEGHDLFKRKSAYMRFGKRSMGQEEDHDMMKRKSAYMRFGR
>FLP-6_Caenorhabditis_tropicalis_Csp11.Scaffold629.g15746.t1
MNSRGLILMLGVIVAVAIAQQDSDLEREMEKRKSAYMRFGRSDGGNPMEMEKRKSAYMRF
GKRSAGDEQESAVSGGGDDMEMEKRKSAYMRFGKRSETPEEDVMSAEKRKSAYMRFGKRS
SDMEMLDGFNGEQGHDLFKRKSAYMRFGKRSMGEEEDHDMMKRKSAYMRFGR
>FLP-6_Cylicostephanus_goldi_CGOC_0000048201-mRNA-1
MNRTVLAAFLLTFAVVYAFQPDDQFSEMEKRKSAYMRFGRSDPELTDQLSLEKRKSAYMS
TFKLIFILRFGKRSEVADEDTLDMEKRKSAYMRFGKRKSAYMRYVFKKEIFIVSEINQAC
EKSTDKRDLY
>FLP-6_Dictyocaulus_viviparus_DICVIV_02824
MKRTIVVTVLLTFAVVYAFQDDYQPSGLEKRKSAYMRFGRSAPELVDQLMLNKRKSAYMR
FGKRSDPLNLDDEALDMEKRKSAYMRFGKRKSAFMRFGKRSSELEDASDAFDVEKRKSAF
MRFGR
>FLP-6_Diploscapter_coronatus_DCO_023173
MKYLAALFVSTLVYISLAQHSDSEFQDMEKRKSAYMRFGRSDPEMIDMEKRKSAYMRFGK
RSSPDLAVAMPEEYDLEAAGDVDKRKSAFMRFGKRKSAYMRFGKRSAPFLTDFGDVYGDD
FLDMEKRKSAYMRFGRK
>FLP-6_Diploscapter_pachys_WR25_23627.1
MKYLAALFVSTLVYISLAQHSDSEFQDMEKRKSAYMRFGRSDPEMIDMEKRKSAYMRFGK
RSSPDLAVAMPEEYDLEAAGDVDKRKSAFMRFGKRKSAYMRFGKRSAPFLTDFGDVYGDD
FLDMEKRKSAYMRFGRK
>FLP-6_Haemonchus_contortus_HCON_00155670
MNRMIAAAILLTFAVVYAFQPDDQFSGMEKRKSAYMRFGRSDPELADQMMMEKRKSAYMR
FGKRSEALEEDPMDVEKRKSAYMRFGKRKSAYMRFGKRSSEVEDSPDAIDMEKRKSAYMR
FGR
>FLP-6_Haemonchus_placei_HPLM_0000766401-mRNA-1
MNRMIAAAILLTFAVVYAYQPDDQFSGMEKRKSAYMRFGRSDPELADQMMMEKRKSAYMR
FGKRSEALEEDPMDVEKRKSAYMRFGKRKSAYMRFGKRSSEVEDSPDPIDMEKRKSAYMR
FVHELLQTYTDLP
>FLP-6_Heligmosomoides_polygyrus_HPOL_0001059001-mRNA-1
MNRTIAAAFLLTIAVVYAFQPDDQFSGMEKRKSAYMRFGRSDPELADQLMMEKRKSAYMR
FGKRSETVDDDMLDMEKRKSAYMRFGKRKSAYMRFGKRSSDFEDNSDGIDMEKRKSAYMR
FGR
>FLP-6_Heterorhabditis_bacteriophora_Hba_18802
MSGVLITLFFLVVVVYASQQEDPLSEVEKRKSAYMRFGRIDPELYGDEMPMEKRKSAYMR
FGKRSEAADDLDIEKRKSAYMRFGKRKSAYMSLYVKFYHLKSKGYYIRFGKRSMEFDEAL
EPVDMEKRKSAYMRLVFLLDKYLRKMKIFIFRFGR
>FLP-6_Mesorhabditis_belari_mbelari.g20297.t1
MNRIILCLAASLSVILAFQQEFAPDTSAFEEMAKRKSAYMRFGRSDPEMGADGMEMEKRK
SAYMRFGKRSDPEQFDEEMMDMEKRKSAYMRFGKRKSAYMRFGKRSPMEQMGEMEPMDME
KRKSAYMRFGK

>FLP-6_Micoletzkya_japonica_MicoRS5524-mkr-S44-3.61-mRNA-1
MKTVLIGFACLAALTLATNSDDAGDVEKRKSAYMRFGRSDPEMIAEMEKRKSAYMRFGKR
SDAEFAPEEVDEDEMEMEKRKSPYMRFGKRKSAYMRFGKRSADEGMDMEKRKSAYMSPCI
VECDCATFTNDRGKRMFRRAAAQFGRVATGLAGVQVAAAATVALSDKELSRKPEWYQQAV
HSVEDALKRLSKYGFIESPLALEEAHDALIKVAEVHNTELLWRVARVLFEKAELSKNHSE
QLHLLHEAKEYAKKALAVEPAKGSAGAHKWYALILQRLAKLEKKHDYEEEAAKHLEKAVA
IDPKDPYALHLLGVHQYNKKDYAAAIATFQKAEAIKAGFSAANLYYLGVALKGAGKKQEA
IEKLKAAVQTHSRNRFDGKARSNAKNALNSLGLKPEEYEIEDL
>FLP-6_Necator_americanus_NECAME_10132
MEKRKSAYMRFGRSDPELTDQLMMDKRKSAYMRFGKRSEVPDEESLDMEKRKSAYMRFGK
RKSAYMRFGKRFSDFDDDSEPMDMEKRKSAYMRFGR
>FLP-6_Nippostrongylus_brasiliensis_NBR_0000196001-mRNA-1
MEKRKSAYMRFGRSDPEQAEQMMMEKRKSAYMRFGKRSEAIDEDTMDMEKRKSAYMRFGK
RKSAYMRFGKRSPDYEDAADLVDMEKRKSAYMR
>FLP-6_Oesophagostomum_dentatum_OESDEN_09208
MNRTVLAALLLTFAVAYAFQPDDQFSGIEKRKSAYMRFGRSDPELTDQLLMDKRKSAYMR
FGKRSEVPDEDTLDMEKRKSAYMRFGKRKSAYMRFGKRSLEFDDSSDPIDMEKRKSAYMR
FGR
>FLP-6_Oscheius_tipulae_OTIPU.nOt.2.0.1.t00028
MHTVQLIFACCFVATIFATQEDESFDMEKRKSAYMRFGRSDPELANGIAMEKRKSAYMRF
GKRSDGGEEDMDMEKRKSAFMRFGKRKSAYMRFGKRSMDIEESLPMDMEKRKSAYMRFGR
>FLP-6_Pristionchus_arcanus_arcanus-mkr-S_77-0.56-mRNA-1
MKSVIALLFIASITITYGQDDFAGDVAKRKSAYMRFGRSDPQMMGEMEKRKSAYMRFGKR
SDPSLVEPMEGEMADSEMVDEMEKRKSPYMRFGKRKSAYMRFGKRSAPEEEGIDMEKRKS
AYMRMFRRAAATFGRVATGVAGVQVAAAATVALTDKELSKKPEWYQIAVRSIEDALKRTS
KYGYIESAASLEEAHEALVKVGDLHNTEIQWRIARILFEKAELSKNHDEILHLLHEAKDH
AKKALAVEPAKGSAGAHKWYALILTRLAKLEKKPEYESEAVKHLEKAVTVDPKDPYALHL
LGVTQYNNKDYSAAVATFQKAEAIKAGFSASNLYYLGAAQKAAGKKNDAIVTLKAAVQAN
PKNRFDGKARSQAKNALAGLGLKPEEYEVED
>FLP-6_Pristionchus_entomophagus_entomophagus-mkr-S22-0.30-mRNA-1
MRFGRSDPDMMAEKRKSAYMRFGKRSDPSLVESMDGEMADSEMVDEMEKRKSPYMRFGKR
KSAYMRFGKRSAPSEEGIDMEKRKSAYMRFGR
>FLP-6_Pristionchus_exspectatus_exspectatus-mkr-S_314-0.75-mRNA-1
SFSIYQAHHQPSIGIHLKMKSVIALLFIASIAITYGQDDLAGDVAKRKSAYMRFGRSDPQ
MMGEMEKRKSAYMRFGKRSDPSLVEPMEGEMADSEMVDEMEKRKSPYMRFGKRKSAYMRF
GKRSAPEEEGIDMEKRKSAYMRMFRRAAATFGRVATGVAGVQVAAVATVALTDKELSKKP
EWYQIAVRSIEDALKRTSKYGYIESAASLEEAHEALVKIGDLHNTEIQWRIARILFEKAE
LSKNHDEVLHLLHEAKDHAKKALAVEPAKGSAGAHKWYALILTRLAKLEKKPEYESEAVK
HLEKAVTVDPKDPYALHLLGVTQYNNKDYSAAVATFQKAEAIKAGFSASNLYYLGAAQKA
AGKKNDAIVTLKAAVQANPKNRFDGKARSQAKNALAGLGLKPEEYEVEDL
>FLP-6_Pristionchus_fissidentatus_fissidentatus-mkr-S78-6.27-mRNA-1
MSSSSAHSGFTLKMQASLVLFCIGLIAICSAQDDFSGDVAKRKSAYMRFGRSDPEMHGEM
EKRKSAYMRFGKRSDPALLESEGEMTDSEMVDEMEKRKSPYMRFGKRKSAYMRFGKRSAP
ADEVLDMEKRKSAYMRMFRRAASTFGRVATSLAGVQVAAAAVALTDKELSKKPEWYQIAV
RSVEDALKRTSKYGYIESGEALEEAHGALTKVGDLHNTEIQWRIARILFEKAELSKNHDE
ILHLLHEAKDHAKKALAVEPAAGSAGAHKWYALILTRLAKLEKKPELEGEATKHLEKAVK
IDSKDPYALHLLGVQQYNSKDYSAAVASLQKAETIKAGFSAANLYYLGAAQKAAGKKNEA
IVNLKAAVQANPKNRFDGKARSQAKSTLAGLGLKPHEYEVEDL
>FLP-6_Pristionchus_japonicus_japonicus-mkr-S20-2.30-mRNA-1
MLIPALSSIYTIVVRYNRVRSETLEIIPSSARESNPSFFLILSHLRLIISLQSGILTMKS
VLVLLFVASIALSYAQDDFAGDVSKRKSAYMRFGRSDPEMVGEMEKRKSAYMRFGKRSDP
SLVEPMDGEVADSEMVDEMEKRKSPYMRFGKRKSAYMRFGKRSAPAEEGIDMEKRKSAYM
RMFRRAAASFGRVATGLAGVQVAAAAVALTDKELSKKPEWYQIAVRSIEDALKRTSKYGY
IESASTLEEAHEALGKVGDLHNTEIQWRIARILFEKAELSKNHDEVLHLLHEAKDHAKKA

LAVEPAKGSAGAHKWYALILTRLAKLEKKPEYESEAAKHLEKAVTVDPKDPYALHLLGVH
QYNNKDYSAAVTTLQKAEAIKAGFSASNLYYLGAAQKAAGKKNDAIVTLKAAVQANPKNR
FDGKARSQAKNALAGLGLKPEEYEIEDL
>FLP-6_Pristionchus_maxplancki_maxplancki-mkr-S1047-0.10-mRNA-1
IGTLVVCRSRLYVAYDILRTNEIIPSSSSFNLSGSSPSIGIQLKMKSVVALLFIASLAIC
YGQDDFAGDVAKRKSAYMRFGRSDPEMIGEMEKRKSAYMRFGKRSDPSLVEPMEGEMADS
EMVDEMEKRKSPYMRFGKRKSAYMRFGKRSAPVEEGIDMEKRKSAYMRMFRRAAATFGRV
ATGVAGVQVAAAATVALTDKELSKKPEWYQIAVRSIEDALKRTSKYGYIESAASLEEAHE
ALGKVGDLHNTEIQWRIARILFEKAELSKNHDEVLHLLHEAKDHAKKALAVEPAKGSAGA
HKWYALILTRLAKLEKKPEYESEAVKHLEKAVTVDPKDPYALHLLGVTQYNNKDYSAAVA
TFQKAEAIKAGFSASNLYYLGAAQKAAGKKNDAIVTLKAAVQANPKNRFDGKARSQAKNA
LAGLGLKPEEYEVEDL
>FLP-6_Pristionchus_mayeri_mayeri-mkr-S145-0.13-mRNA-1
MKSICVFLLIAAIGIVSAQEDYSGDVAKRKSAYMRFGRSDPQMMAEMEKRKSAYMRFGKR
SDPSLVDPIEGEMADSETIDEMEKRKSPYMRFGKRKSAYMRFGKRSAPADEGMDMEKRKS
AYMRMFRRAAASFGRVATGLAGVQVAAAAVALTDKELSKKPEWYQIAVRSIEDALKRTSK
YGYIESTTSLEEAHEALNKVSDIHNTEIQWRIARILFEKAELSKNHDEVLHLLHEAKDHA
KKALAVEPAKGSAGAHKWYALILTRLAKLEKKPEYETEAYKHLEKAIAVDPKDPYALHLL
GVQQFNNKDYSAAVATLQKAEAIKAGFSASNLYYLGAAQKAAGKKNEAIVALKAAARSQA
KNALAGLKVKPEEYELEDL
>FLP-6_Pristionchus_pacificus_PPA28847.1
MKSVIALLFIASIAITYGQDDFAGDVAKRKSAYMRFGRSDPQMMGEMEKRKSAYMRFGKR
SDPSLVEPMEGEMADSEMVNEMEKRKSPYMRFGKRKSAYMRFGKRSAPEEEGIDMEKRKS
AYMRYRMFRRAAATFGRVATGVAGVQVAAAATVALTDKELSKKPEWYQIAVRSIEDALKR
TSKYGYIESAASLEEAHEALVKVGDLHNTEIQWRIARILFEKAELSKNHDEVLHLLHEAK
DHAKKALAVEPAKGSAGAHKWYALILTRLAKLEKKPEYESEAVKHLEKAVTVDPKDPYAL
HLLGVTQYNNKDYSAAVATFQKAEAIKAGFSASNLYYLGAAQKDAGKKNDAIVTLKAAVQ
ANPKNRFDGKARSQAKNALAGLGLKPEEYEVEDL
>FLP-6_Strongylus_vulgaris_SVUK_0001678501-mRNA-1
MDLTIRNKIFLMMNLFHTFLRAVSILDRCLDMNRTVVAAFLLTFAVAYAFQPDDQFSGME
KRKSAYMRFGRSDPELTDQLLLEKRKSAYMRFGKRSEVPDEDSLDMEKRKSAYMRFGKRK
SAYMRSNSLKSKNERLATY
>FLP-6_Teladorsagia_circumcincta_TELCIR_02863
MNGGREAERVPRRFKIYLPVKENKFFRKLLDKLKDTPEGKVQPQLKVHVFPDSIVLGYTG
LEKLSLSMNRMIAAAILLTFAVVYAFQPDDQLSEMEKRKSAYMRFGRSDPQLADQLMMDK
RKSAYMRFGKRSEAIDEDTMDMEKRKSAYMRFGKRKSAYMRFGKRSSEFDDAPDAIDMEK
RKSAYMRFVFKFTDDDPPLLPHSEHRNMNRGSFFRVPSTMIYSKRQDNIPKKRTSIAAQ
>FLP-6_Angiostrongylus_costaricensis_predicted
TLYFRKITALLSEMEKRKSAYMRFGRSIPDLGNQLTIDKRKSAYMRSASFI
>FLP-6_Parapristionchus_giblindavisi_predicted
MEKRKSPYMRFGKRKSAYMRYGNR
LSIDYRFGKRSADEGIDMEKRKSAYMRYGSFPVGTSFDLLIFFSDSENKMLIMRVSPSACLVHIPFVN
SKFFVLSFLVLLIHHGLVHRGRTYKSSF
>FLP-6_BXY_0826700.1 peptide: BXY_0826700.1 pep:protein_coding
MRHSLVLVALATVIAAEFAVAVTDLCDQFPDLAGCIKPEQVKRKSAYMRFGKRSLSLGEP
VEEDDAEPFAAYEKRKSAYMRFGKRSGGMLEVPEDDGVEMEKRKSAYMRFGKRKSAYMRF
GKRSGQDFLDNSDQPMEMHKRKVAALSP
>FLP-6_Halicephalobus_mephisto_MSTRG.363.2.p1
MNQIAVFVFLSVFVAVCLANAGGVEEYEVLCGRFPSLAECQELNTMEKRKSAYMRFGRSA
PDDLMGEMDKRKSAYMRFGKRSESAEPELEGLQEMEKRKSAYMRFGKRKSAYMRFGKRSE
SVEPEFDAPQEMEKRKSAYMRFGKRKSAYMRFGKRSDDFQDSPLDMEKRKSAYMRFGK
>FLP-6_Panagrellus_redivivus_Pan_g5294.t1
MLAVALLGVFATSVLATEKPAASSDDVDLASICEDFPHLVACEEALMNTLDKMEKRKSAY
MRFGRSNGGIEGDEAAINGAMEKRKSAYMRFGKRSGASEAEVVEPEMEKRKSAYMRFGKR

KSAYMRFGKRAVEDFEDAPVQNGLLAPAEKRKSAYMRFGK
>FLP-6_Parascaris_univalens_PgR013_g104_t01
MRNVEMMLQRALCMTLFGAVCAFRLLTKEESAPQLVSENDLTMICELYPQLELCTNAHLM
DKRKSAYMRFGRSDSSTIPDDAEGVEKRKSAYMRFGKRDDSAPSLSDDEQTDGGEMEKRK
SAYMRFGKRKSAYMRFGKRSDEQPAAEIEKRKSAYMRLGRR
>FLP-6_Parastrongyloides_trichosuri_PTRK_0001097800.1
MIKKFVIISLIVIAIIHNTKGIVNYEESNDRGVGMYDLCEVIPTHILCDPDNRFMDIIKD
RKDSQKKVVEDNTKQNVIKRKSAYMRFGKRSGSDDDEMLVPESNEIEKRKSAYMRFGKRK
SAYMRFGKRSDGEYLNDIETFSPYEKRKSAYMR
>FLP-6_Rhabditophanes_kr3021_RSKR_0000497700.1 peptide:
RSKR_0000497700.1 pep:protein_coding
MNILLLICLGGCICGVALGTSHADEPPKDVKEIDEMLLCDYLPSDPLCQPEQQVLTAEKR
KSAYMRFGRSDPEMMQKRKSAYMRFGKRAEVPEQQDNGDVEMEKRKSAYMRFGKRKSAYM
RFGKRSEVGGDEETDQTEYEPVVDKRKSAYMRFGK
>FLP-6_Steinernema_carpocapsae_L596_027509
MRPLCLAVTVSFLAIASGFKPVQRPLAIEGGATSSLDDGLFCENFPKHPACEAQVGQQME
KRKSAYMRFGRSDGGLDMEKRKSAYMRFGKRGADEEVFEAEEPMEKRKSAYMRFGKRKSA
YMRFGKRGDATFDDDSMAMEKRKSAYMRFGKRSDNVEAMSPFEIEKRKSAYMRFGR
>FLP-6_Steinernema_feltiae_L889_g3651.t1
MTSRYGCSSKPIAPANPFRRLFVERHTALLPSPKRFAXXXXRSFRVAFSRFKRDFLVQPH
LGSGKRILSVGSVVRRTPHGASSISKALRSLQRSPSTFVSSKRMRTFCLAVTVCLVAVAS
GFKPVERLSVAEPNPANSVDDDVLFCENFPKHPACEPQVSQQMEKRKSAYMRFGRSDGGA
EMEKRKSAYMRFGKRGDDEEMFEAEGPMEKRKSAYMRFGKRKSAYMRFGKRGFESFGESA
MMLPEKRKSAYMRFGKRSDNVDAMNPFEMEKRKSAYMRFGR
>FLP-6_Steinernema_glaseri_L893_g29653.t2
MPFHSHLRKDNDEAKIEKKKHEKPMNGPSIPKLLSSPLLGLRDCGFYKWPSCLCPSSLLY
FVSPRLSEEAEVLRRGDYPFGRASAWRLAASTWTVCPEDDRKTNLLVGEGLNSEICDDPG
IPGMAEDYGGQRWKCVRSKLFSAVEWIERDSSVRAESSEIDARGMVEGHVQPSFICSKKK
RTKKKEAFLYADCHKSPHIIEPASLAALPFIVFATSPHTIPHNLMAAATVSHSRIIPRDD
VISPPTTSSAMHRVGSSLSVRMRPFSLALTVILGFAAAAMGLTPVQRAPSALEESFFCET
FPKHPACEAQVGQQMEKRKSAYMRFGRSDGGXXXXVLTYSKSTKFGQQMEKRKSAYMRFG
RSDGGLDMEKRKSAYMRFGKRGDDEEMVPVEEPMEKRKSAYMRFGKRKSAYMRFGKREDP
ALEEDMAMEKRKSAYMRFGKRGDDVDAATPFEMEKRKSAYMRFGR
>FLP-6_Steinernema_monticolum_L898_g34587.t1
MRSFFLTLTVTAVAVAAGFQHEQRSSVEAIPSEPIDHMFFCENFPKHPACETQVGQQMEK
RKSAYMRFGRSDGGVDMEKRKSAYMRFGKRGEYADLLEAEEPMEKRKSAYMRFGKRKSAY
MRFGKRGDDSFDENAMAMEKRKSAYMRFGKRSANVDAISPFEVAKRKSAYMRFGR
>FLP-6_Steinernema_scapterisci_L892_g14472.t1
MRPLCLAVTVSILAIASGFKPVQRPSAVEDAAPNPLDDVLFCENFPKHPACETQVGQQME
KRKSAYMRFGRSDGGIDMEKRKSAYMRFGKRGADEEVFDAEEPMEKRKSAYMRFGKRKSA
YMRFGKRAVSTFDDDSMAMEKRKSAYMRFGKRSDNVDAMSPFEMEKRKSAYMR
>FLP-6_Strongyloides_papillosus_SPAL_0001335200.1
MKYFLLITAIFIFTIANIEGNDDNKDSSMVDNINQPLLCEVLPSHELCLEVQSSVHGAPV
KRKSAYMRFGRSDPEVVEKRKSAYMRFGKRSMGSEEIGDDAFTQDNGIEKRKSAYMRFGK
RKSAYMRFGKRDVNEEGDVEFSSPYEKRKSAYMRFGK
>FLP-6_Strongyloides_ratti_SRAE_1000039800
MKYYLLITFLSISYLLNIEGNDEVNDSLSLETNKQLYICDVIPDHYLCSSDETISTPIKR
KSAYMRFGRSDPDVVEKRKSAYMRFGKRSTGNDEIRDETFIPENGIEKRKSAYMRFGKRK
SAYMRFGKRGMELENSADAFSPLEKRKSAYMRFGK
>FLP-6_Strongyloides_stercoralis_SSTP_0000812800.1
MKYYILVVFLSISCLFNVKGNNEIKESSAIENNKQLYLCDIIPEHYLCTSDESLSTPIKR
KSAYMRFGRSDPGEVEKRKSAYMRFGKRSSGNDEIEDEAIIPENGIEKRKSAYMRFGKRK
SAYMRFGKRDMDMESGSDIYSPLEKRKSAYMRFGK
>FLP-6_Strongyloides_venezuelensis_SVE_0332400.1

MDLQKGDGLLFLKKFFTLAFFYKIKIGETSLKMKYFLLITAIFIFTIVNIKGNDENKDSP
VLDNINQPLLCDVIFSHELCIEAQSLVHGVPVKRKSAYMRFGRSDPEVVEKRKSAYMRFG
KRSMGSEEIGDDAFIQDNGIEKRKSAYMRFGKRKSAYMRFGKRDVSEESDNKIFSPYEKR
KSAYMRFGK
>FLP-6_Acrobeloides_nanus_ACRNAN_scaffold663.g24967.t1
MVYAKEEKTPAPVVKSDEQQSKFANLCEEFPLLRDCLEQQMEKRKSAYMRFGRSSPDVDS
YTDDNEDNSGLLDMEKRKSAYMRFGKRFFPANAHGFNDDSSQEVEKRKSAYMRFGKRKSA
YMRFGKRSNDETGEVEKRKSAYMRFGRR
>FLP-6_Ditylenchus_destructor_Dd_00832
MAVSFKIREILQLKLSHRMSLGCFILMAIIATIFAETTPSTPEHLAEAENLDYFYNPAHF
LQVCAQFPELAACQQVGSLVEKRKSAYMRFGRAAPDAVVEVEDEPAMEKRKSAYMRFGKR
SPSTHIMLENEMADEQVHDMAKRKSAYMRFGKRKSAYMRFGKRSSDDAGEMEKRKSAYMR
FGKRAVPSENMPEMEKRKSAYMRFGKRR
>FLP-6_Ditylenchus_dipsaci_jg8206.1
MAQSSVREAFYAVLLAVFAATFATATNTPTTAKDGPQESGLETSEGSYFYSPAEFLHICA
QFPGLAACQQESGSLVEKRKSAYMRFGRSAPEEEQDSPAMEKRKSAYMRFGKRSPSQESV
NEEVGGDVQKRKSAYMRFGKRKSAYMRFGKRSADNNMEKESQPICALENDLYWLMMKTWE
KWRRGNPPTCDLAKDGELKHH
>FLP-6_Globodera_pallida_GPLIN_001273600
MMWSLFFILFSCFIFRPASSDLLSSSTDAQLQQLCARFPGMEVCQPDDEALGAMAKRKSA
YMRFGRAAAEQEPVMEKRKSAYMRFGKRSPEDENVDQLLALNEPDAQQMEKRKSAYMRFG
KRKSAYMRFGRK
>FLP-6_Globodera_rostochiensis_GROS_g08471.t1
MLLNNNTMSCSSSLFMIWSLFVILFSCFIFRPASSDLLSSSTDAQLQQLCARFPGMEVCQ
PDDEVLGAMAKRKSAYMRFGRAAAEEEPAMEKRKSAYMRFGKRSPEDENVDQLLAFNEPD
ALQMEKRKSAYMRFGKRKSAYMRFGRK
>FLP-6_Heterodera_glycines_Hetgly.G000025058
MAMNTNSIHCSSASSSPSVPMMLSMFCFIFAIFLHPASSSDLLSSSDDAKLQQMLCARFP
GLAECQPNDEASRGIMTKRKSAYMRFGRAAAAADPSEEYEMEKRKSAYMRFGKRSPAEEG
ETYQIWAANEPDAQMEKRKSAYMRFGKRKSAYMRFGRK
>FLP-6_Meloidogyne_arenaria_M.Arenaria_Scaff540g011450
MPNISKSNNFNLNQLLIITSLLLINLTIFISANYDETNLESNLAEIKQLCQQFPNLAECR
ILLSPPMQMEKRKSAYMRLGKRKSAYMRFGKRGGVNEANQIHDSEYISIDGLMAENQPME
KRKSAYMRFGKRKSAYMRLG
>FLP-6_Meloidogyne_enterolobii_Meloidogyne_enterolobii
scaffold41449_cov248.g25743
MPNISKSNNFNLNQHLIITSLLLINLTIFISANYDDIENNLAEIKQLCQQFPNLAECRIL
SPPVQMEKRKSAYMRLGKRKSAYMRFGKRGVNEANQIPDSEYISIDGLMAENQPMEKRKS
AYMRFGKRKSAYMRLG
>FLP-6_Meloidogyne_graminicola_NXFT01000189.1.4896_g
MFKILYNFQFNILIITSLFLINLILATTNHDILEQQQQLCKDFPELIECLYYYIPIENKI
KMMEKRKSAYMRLGKRKSAYMRFGKRGAIENNQISNNEYLNNIDKLLEENQPMEKRKSAY
MRFGKRKSAYMRLG
>FLP-6_Meloidogyne_hapla_MhA1_Contig1561.frz3.gene1
XFGKRGVNEDNQIPDSEYTSIDGLMSENQPMEKRKSAYMRFGKRKSAYMR
>FLP-6_Meloidogyne_incognita_Minc3s00085g04057 peptide:
Minc3s00085g04057 pep:protein_coding
MPNISKSNNFNLNQLLIITSLLLINLTIFISANYDETNLESNLAEIKQLCQQFPNLAECR
ILSTPMQMEKRKSAYMRLGKRKSAYMRFGKRGGVNEANQIPDSEYISIDGLMAENQPMEK
RKSAYMRFGKRKSAYMRLG
>FLP-6_Meloidogyne_javanica_M.Javanica_Scaff301g004621
MPNISKSNNFNLNQLLIITSLLLINLTIFISANYDETNLESNLAEIKQLCQQFPNLAECR
ILLSPPMQMEKRKSAYMRLGKRKSAYMRFGKRGGVNEANQIPDSEYISIDGLMAENQPME
KRKSAYMRFGKRKSAYMRLG

>FLP-7_Plectus_sambesii_PSAMB.scaffold74size85960.g1655.t1
METATRRAAGWVFLGSSPFTTIPIHSVSKGGADPTADGVPSPAHAPPIPPSGNGPNSPQS
ALSLPGHDEPFGMIRNTDRAFSMRTKRAPPPPVGALLARHSRAVIDSRLLSELAQQTRLF
DDINDATALLERLPPVLQLIDALPESLPLLDAIYVKCHQLKRFGSSAFPVRALRPENILT
QVLWWLAHAEDECIVVSNQQISRLSSRSSPPYSLLDTSSRRPDTAAFKFAIMLPHLQSLL
RIIHMECGDLLDEVRAERDRLNEALEALGLHGLVNCSGDRLIRLADAARIELTNHHNRLK
SLVGQLEKCTSVIKTKGWSINSSAGRPVPTRASHQRQMTLRVTEVCERMAQNEGIREILT
EAVENNELDSLLRIVGKRVTSDKNLLTRFHALKSSIPPGYAVIETDPVVAKVLVRFKAAY
DKLLTLAPTATPTTRIRADDSGVGLDDSDRGLSCSSSDTIDGPDSPALVVKRRTRLSGPV
AQQLKARNLLLSQPAHHRSLNELPVERSALLKPVKSLSNLAISGTFEMYRQQNGRSTRSG
SSTTENEEETAKLRNHINRLGAERLNFHQVDDPNMRLSLLIVLPLLFYVTAAYYSDGEEV
GEPDIANEKRQMRASMVRFGKRAPSDRSAMVRFGKRAPSDRSAMVRFGRAPSDRSAMVRF
GKRLSNEENEQRYYAHFNQDEQDREVPATNQL
>FLP-7_Ascaris_suum_AgR018X_g099_t04
MHSAPIFGAFRRYICHFKALDFSRARNWSWNWRVRSILTFYQSQMSTRSKRVQMIRFVEL
IAATAILCQLLLAADGNDFRQLYYFDIYPDGGVLDSTEPMKRTPMDRSSMVRFGKRAPMD
RISMVRFGKRAPMDRSSMVRFGKRMPIDRSSMVRFGKRDFPLFTDEAYSA
>FLP-7_Toxocara_canis_Tcan_01443.1
MIRFLALIAATATLCELLPAADDNDYRQLSYFDVHSDGALFGNNEIIKRTPIERSSMVRF
GKRVPMDRSSMIRFNKRAPMDRSSMVRFGKRTPIERSSMVRFGKRNPSLFADTMYSASSS
I
>FLP-7_Ancylostoma_caninum_ANCCAN_11898
MVQVSVLLLAAVCVATQISLPGAQTREEFDNLERFDNFEKRAPMDRSSMVRFGKRAPMDR
SSMVRFGRAPMDRSSMVRFGKRAPMDRSSMVRFGKRAPMDRSSMVRFGKRDSVEAMVI
>FLP-7_Ancylostoma_ceylanicum_Acey_s0001.g488.t1
MFDDFEKRAPMDRSSMVRFGKRAPMDRSSMVRFGRAPMDRSSMVRFGKRAPMDRSSMVRF
GKRAPMDRSSMVRFGKRDSMEAMMI
>FLP-7_Caenorhabditis_angaria_Cang_2012_03_13_00002.g232.t1
MVRFGRSPMDRSAMVRFGKRSPMDRSAMVRFGKRSPMDRSAMVRFGRSPYDRSSMVRLGK
RSPMDRSAMVRFGKRSATETEADDEQ
>FLP-7_Caenorhabditis_briggsae_CBG14771.1
MLGSPRFLLLALGLLVLVLAESSVEQQVQDQTDLDKSGEQLSEEDIIEEQKRSPMERSSM
VRFGRSPMERSSMVRFGKRSPMERSAMVRFGKRSPMDRSAMVRFGKRLPSDRSSMVRLGK
RSPMDRSAMVRFGKRSPMDRSAMVRFGRSSIDRASMVRLGKRTPIQRSSMVRFGKRSADE
TENTNE
>FLP-7_Caenorhabditis_elegans_F49E10.3.1
MLGSRFLLLALGLLVLVLAEESAEQQVQEPTELEKSGEQLSEEDLIDEQKRTPMQRSSMV
RFGRSPMQRSSMVRFGKRSPMQRSSMVRFGKRSPMQRSSMVRFGKRSPMERSAMVRFGRS
PMDRSKMVRFGRSSIDRASMVRLGKRTPMQRSSMVRFGKRSMEFEMQSNEKNIEDSE
>FLP-7_Caenorhabditis_inopinata_Sp34_X0233000.t1
MLGSRFFLLIFGLLFLICSGEYTEKKVQQPTELGMPGEELTEEDLIDEQKRSPMQRSSMV
RFGRSPMQRASMVRFGKRSPMQRSSMVRFGKRSPMERSAMVRFGKRSPMERSAMVRFGRS
PMERSRMVRFGRSSMDRSSMVRLGKRTPMQRSSMVRFGKRSSYASSNINDFETNERSIQY
NE
>FLP-7_Caenorhabditis_japonica_CJA00701.1
MLGPRFFLLALGLLALIRADESTNEQQIQEQEQNELDKSEEQQLTDGDLIDEQKRSPMQR
SSMVRFGRSSNQMSQFEKRSPMQRSSMVRFGKRSPMQRAAMVRFGKRSPMERSAMVRFGR
SAMDRSSMVRFGRSSIDRASMVSSRLGKRSPMQRSSMVRFGKRSTQQSSSEN
>FLP-7_Caenorhabditis_latens_FL83_00448
MLGSRFLLLALGLLVLVWAEESTEQKVQEQTDLDKSGEQLSEEDLIDEQKRNPLQRSSMV
RFGRSPMQRSSMVRFGKRSPMQRSSMVRFGKRSPMQRSSMVRFGKRSPMERSAMVRFGRS
PMDRSKMVRFGRSSIDRASMVRLGKRTPMQRSSMVRFGKRSAPSDINEMQNNKEIQDNE
>FLP-7_Caenorhabditis_nigoni_Cni-
MLGSPRFLLLALGLLVLVMAENSVEQEVQDQTDLDKSGEQLSEEDIIEEQKRSPMERSSM

VRFGRSPMDRSSMVRFGKRSPMERSAMVRFGKRSPMDRSAMVRFGKRLPSDRSSMVRLGK
RSPMDRSAMVRFGKRSPMDRSAMVRFGRSSIDRASMVRLGKRTPIQRSSMVRFGKRSTSS
DEIENTNE
>FLP-7_Caenorhabditis_remanei_FL82_01797
MLGSRFLLLALGLFVLVWAEKSTEQQVQEQTDLDKSGEQLSEEDLIDEQKRNPMQRSSMV
RFGRSPMQRSSMVRFGKRSPMQRSSMVRFGKRSPMQRSSMVRFGKRSPMERSAMVRFGRS
PMDRSKMVRFGRSSIDRASMVRLGKRTPMQRSSMVRFGKRSAPSDINEIQDNEQIRDNE
>FLP-7_Caenorhabditis_sinica_Csp5_scaffold_00459.g11654.t2
MEDKDHFGASKDHMIVVIYSLGQSKMLGSRFLLLALGLLVLVLAQESTEQKVQEQTDLDK
SGEQDSEEDLIDEQKRNPMQRSSMVRFGRSPMQRSSMVRFGKRSPMQRSSMVRFGKRTPM
QRSSMVRFGKRSPMERSAMVRFGRSPMDRSKMVRFGRSSIDRASMVRLGKRTPMQRSSMV
RFGKRADTNEDISLDEE
>FLP-7_Caenorhabditis_tropicalis_Csp11.Scaffold626.g6560.t1
MLGSRFLLLALGLFVLIAAETAEQQIQEQTDLDKSGEQLPEEEDLTDEQKRSPMQRSSMV
RFGRSPMQRSSMVRFGKRSPMQRSSMVRFGKRSPMQRSSMVRFGKRSPMERSAMVRFGRS
PMDRSKMVRFGRSSIDRASMVRLGKRTPVQRASMVRFGKRSDPTDVNEMELNEQSIQEN
>FLP-7_Diploscapter_coronatus_DCO_023570
MLGLSNLFYAMILGLFALAAVTLAELGDDDQLEMMQFEENQMTDPESLENIDKRSPMDRS
SMVRFGKRGPMQRSSMVRFGKRSPMDRSSMVRFGKRSPMQRSSMVRFG
>FLP-7_Diploscapter_pachys_WR25_14541.1
MILGLFALAAVTLAELGDDDQLEMMQFEENQMTDPESLENIDKRSPMDRSSMVRFGKRGP
MQRSSMVRFGKRSPMDRSSMVSYQKEDFNDTFIEKRGTLLRRKEERSRHNPVGAERRKRF
FFSPIKLRDQKSMLNRLIDRLLAISWLLVLSTWAQNFEKSHLGFPEAWDGDKYRELYCPS
KNVPKWLDGYFLCQLSASYGNLSAPPGQKISHMIDAIGAVVFSAQYYPARPYKIWEFYDR
NMSKASVPWAGWSDYNITAMSKWEQVPSNPDSARFHPNLDFWQVGNRIIAGTEAPYWVGY
EFDVRTLQKFKLFPFTEENDIFSRPRPSMIPISMAIHERNDHDGTIWGSFSAMNFEEQRF
YQGIFTVDRNGVRRVVGLYDYGVWDTNACGPNDEYIGDKTLLPGYIHSITSTENYVILPI
TSLLINPCKFKEPPMTNIRSAIQKGGLWGMDFYDMVPMRFLIFDKRTREFTTGKPLEVFP
SMFVTHQLNAFENPDGMIFADMVVYDSHDPYVKYFYTDFLTSQLYPSTARVLRFTLDSKQ
QRVMYNYLIPQETIAADFPQINHQYEQKPYQWAYIVEHPFAAENKIIKINVDDPAGTRNK
EFQSDPQLVLHEPWFVPKPDASKEDDGVLLIRSLDTQENKGVLLIVDAETMIEIGRAYVP
ISIPFGFHNRYFSKKDLGLPEGFQMNQVMSQFRPADKKGQGGFATLPLRKYTVVPPNGLP
ISTHGMTKPTTATAEISTEEGISSESTTSSTTTAATTTTTTTTTAAPSTIASTTTTTKST
TTAKSEKPDTVTTQKWIPISSTPISTTQSTTRLTTTPKIPRWWPLASTEIPWWQKVENGN
QHTLPPLFPISANIPTSTLKVIAASKSGMKPDESQNRVIVPVPSTPSNIDELYEQTINAL
CSWLPRVFTSISPEMCKMQGKRAAKWMAPIASSYAERFRMSRMNRRNHVQNVNVHASLKP
SRYDPNEETTEDRDEVHQRRKRY
>FLP-7_Haemonchus_contortus_HCON_00164220
MPMVQAALLLITMCVVIISTSGSLASGHYDFADSTLTEGKRTPMVRSSMVRFGKRAPMDR
SAMVRFGRAPMDRSAMVRFGKRAPMDRSSMVRFGKRAPMDRSSMVRFGKRMASGEQSQFA
VLYTKPLINLWYQDTWSTDAISITR
>FLP-7_Haemonchus_placei_HPLM_0000496901-mRNA-1
MVRLVHESLWCQDDSQNHSIASSIFRFGKRAPMDRSAMVRFGRAPMDRSAMVRFGKRAPM
DRSSMVRFGKRAPMDRSSMVRFGKRMASGEQSQFAGL
>FLP-7_Heligmosomoides_polygyrus_HPOL_0000333201-mRNA-1
MVQTAILLVTAYVAIVCASEAFDSQYDLPDALLTEEKRAPMDRSSMVRFGKRAPMDRSTM
VRFGRAPMDRSTMVRFGKRAPLERSSMVRFGKRAPMDRASMVRFGKRDASEELAGMIDI
>FLP-7_Heterorhabditis_bacteriophora_Hba_01962
MRRYMKFDQTENDKDDTLRLRLHVNHLKNELAHARKQIAQMMNLQHRGADTSQRIIIIYI
DFFKYLDFIIIYELLFRLWPEPDYPDATSYYNCSSRQDVTNLIGRFSELFIRGRKEIMVA
LDKLPEFVGSEELQLKTILITNSTLNLYFRKTNAAEAIFTSRKFLYFIYLLNVSRTMLGP
STFLVILTACIATMELVQRKSVHEDDIDYNMQEVRININSCLATLYSACFLSLLFVLFSG
INDEGERASVDQLSLTRFGKRVPMNRSAMVRFGRAPMDRSTMVRFGKRAALDRSSMVRFG
KRSPMDRSSMVRFGKRAPMDRSSMVRQTHNLDLVNVLPWTGLQWSVLGNEKAISRSGLVY

IYIYIYIYILYGYFLCQLSASYGNSSAPSGQKLNHMIDAIGAVGSFHISNGQVLFSSQYY
PARPYKIWEFYDRNMSKASVPWAGWSNYNLTAMSRWEQVPVNPDSARFHPNLDFWKVGNR
IVAGTEAPYWVGYEFDVLPDTSFKNHLFILNGVFTNSIKYYLKQNTVIISIFKQFKVFTR
NFNDLITHQFTKIFEKISALNIFYKFSFLRLICHDFRFAETIAADFPQINHAYEQKPYQW
AYIVEHPFAADNKIIKVYTFILQYCFQVLYSLDIVGVLLIVSKIKSK
>FLP-7_Mesorhabditis_belari_mbelari.g1984.t1
MQPLYWLLFLGTAAITFAQDEDNGEAIAILEEINKMNEPAEYDTYYMEKRAPMDRSSMVR
FGKRAPMDRSSMVRFGKRAPMDRSSMVRFGKRAPMDRSSMVRFGKRAPMDRSSMVRFGKR
APMDRSSMVRFGKRAPMDRSSMVRFGKRSSDDSIEPETGPEVLEDVVAAIEAGVEVPPPK
LELGVFTELAVLLSFDQLSSSSSSAPSMITSKSTSNLSTSSGSTSTTSNAFALDSPISPP
PAHFGYTSANPKF
>FLP-7_Micoletzkya_japonica_MicoRS5524-mkr-S43-3.20-mRNA-1
MLGRLTIAVALLSMVALAQYEDDDYEAQLDQVDKDMAVDKRAMMDRASMVRFGKRAPMDR
STMVRFGRAPMDRSTMVRFGKRAPMDRSTMVRFGKRSPMERSSMVRFGKRNMDRASMVRF
GKRAMM
>FLP-7_Necator_americanus_NECAME_16600
MLHDKRAVRAQNHRSPMDAQPTVCAVNCGISRTQFLLPTLPNKFDPSFCPQKDEKLGWPP
VLRNNQPRIHHDLLNGTLLPMYYIWQSQLIRNKAAIFGKRAPMDRSSMVRFGKRALMDRS
SMVRWELTVHIALCSYMSIRLLIEDRKTCKPKKPRTPFFTLSHDRNIIIRGER
>FLP-7_Nippostrongylus_brasiliensis_NBR_0000713701-mRNA-1
MVQAAFLLVTACFAIVSATETSDTHYDLSDGYLTEEKRATMDRSSMVRFGKRAPMDRSTM
VRFGRAPMDRSTMVRFGKRAPMDRSSMVRFGKRAPMDRASMVR
>FLP-7_Oscheius_tipulae_OTIPU.nOt.2.0.1.t00524
MQRFLLVCLVLAVSACLSQEEPLQDVTYEDMEDDSILENNEEKRAPMDRSTMVRFGRSPM
DRSSMVRFGKRAPMDRSSMVRFGKRAPMDRSSMVRFGRTPMDRSSMVRFGKRAPMDRSSM
VRFGKRAPMDRSSMVRFG
>FLP-7_Parapristionchus_giblindavisi_Parapristionchus-sn_msk-
S_73-1.46-mRNA-1
MFIRIALFSLLSIAVIAQFDEDYENNGDMSDIYNEKRATMDRSSMVRFGKRSPMDRSTMV
RFGRAPMDRSSMVRFGKRAPLDRSSMVRFGKRAPLDRSSMVRFGKRAPMDRSSMVRFGKR
AE
>FLP-7_Pristionchus_arcanus_arcanus-ag_msk-S_3-4.95-mRNA-1
MFSYPVRYLILLGLCAAIVASQFRDDGDDEEGLNEIKEEKRAGMDRASMVRFGKRSPIER
SSMVRFGRAPIDRSSMVRFGKRAPMDRSAMVRFGKRAPMDRASMVRFGKRAPMDRASMVR
FGKRAPMDRASMVRFGKRSAMIFPHLIIYRVSPFFPSPLDRLP
>FLP-7_Pristionchus_entomophagus_entomophagus-sn_msk-S66-4.38-mRNA-1
MLSSSLRSFALLALLVALVASQYEDGSDVAEDAEEATISTEKRAVMDRSSMVRFGKRSPM
DRSTMVRFGRAPMDRSSMVRFGKRAPMDRSSMVRFGKRAPMDRSSMVRFGKRAPMDRSSM
VRFGKRAPMDRSSMVRFGKRSAL
>FLP-7_Pristionchus_exspectatus_exspectatus-mkr-S_1-4.59-mRNA-1
MFFSPSIRSLAVLAVVVAVVATQYEEDDGADLAAESEAIIEEKRAGIDRASMVRFGKRSP
MDRSSMVRFGRAPIDRSSMVRFGKRAPLDRSAMVRFGKRAPMDRSAMVRFGKRAPMDRAS
MVRFGKRAPMDRSAMVRFGKRSM
>FLP-7_Pristionchus_japonicus_japonicus-mkr-S236-3.87-mRNA-1
INLLSFSAHTRAHHLHLKMFSSSFRSFFLLALFATFVASQYEDEVGGDVVEDSEEISTEK
RAVMDRSSMVRFGKRSPMDRSTMVRFGRAPMDRSSMVRFGKRAPMDRSSMVRFGKRAPMD
RSSMVRFGKRAPMDRSSMVRFGKRAPMDRSAMVRFGKRSAM
>FLP-7_Pristionchus_maxplancki_maxplancki-mkr-S6-6.31-mRNA-1
METWLFLLISSISAYLLTPGPLHLLNLKMFSSSFRSFFLLALLAALVASQYEDDVGGDVV
EDSDEISTEKRAVMDRSSMVRFGKRSPMDRSTMVRFGRAPMDRSSMVRFGKRAPMDRSSM
VRFGKRAPMDRSSMVRFGKRAPMDRSSMIRFGKRAPMDRSAMVRFGKRSAMIFPSSSE
>FLP-7_Pristionchus_pacificus_PPA22038.1
MFFSPSIRSLAVLAVVVAVVVSQYEEDVSDLAGESEAIIAEKRAGIDRASMVRFGKRSPI
DRSTMVRFGRAPLDRSAMVRFGKRAPMDRSAMVRFGKRAPMDRASMVRFGKRAPMDRSAM

VRFGKRSM
>FLP-7_Strongylus_vulgaris_SVUK_0001023401-mRNA-1
MVQVSVLLLVEACIAAQISLSGAQERENFDIAQRTFGELEKRAPMDRSSMVRFGKRAPMD
RSSMVRFGRTPMDRSTMVRFGKRAPMDRSSMVRFGKRAPMDRSSMVRFGKRDYMEKPDLA
LMI
>FLP-7_Ancylostoma_duodenale_predicted
RHRRFGKRAPMDRSSMVRFGRAPMDRSSMVR
TSGLTSHLLGIEYEKPQHRQHSLLSHAKPNNSVFRFGKRDSVEAMMI
VRTKGHRVVKLLIQNKIQSRLRRFHNRFGKRAPMDRSSMVRFGKRAPMDRSSMVRYAKLPD
>FLP-7_Angiostrongylus_cantonensis_predicted
WILIIGLLLAELPDGNVGLLLGMGDPAGMMNTPIVHRRVTPRNTCQPRRGSEPLGGWRCRASMVRFGR
ISKPRRSS
>FLP-7_Caenorhabditis_brenneri_predicted
VDQLFDSRKIGLYYFYFRIDEQKRSPMQRSSMVRFGRSPMQRSSMVRFGKRSPMERSAMVRFGKRSPM
ERSAMVRFGKRSPMERSAMVRFGKRSPMERSAMVRFGKRSPMERSAMVRFGRSPLDRSKMVRFGRSSI
DRASMVRYVDETVLV
CKKTESIDFRLGKRTPMQRSSMVRFGKRSSDINEIQSNDNE
>FLP-7_Cylicostephanus_goldi_predicted
RLSXXXXXXXXXXXXXXXXXQEVFHSCCRKNLQAFTEKDFLQDFLRTAASSYLTFRYLRFGKRAPMDR
STMVRFGRTPMDRSSMVRYAQSSEPIDRLKTLYS
CQGESTLLTLRFGKRAPMDRSSMVRFGKRAPMDRSSMVRYENHTSIIF
>FLP-7_Oesophagostomum_dentatum_predicted
VTSPRVLHRLADRPPGRRNSNFDYRFGKRAPMDRSSMVRFGKRAPMDRSSMVR
>FLP-7_Pristionchus_mayeri_predicted
CEMILSKIQMTSFFRFGKRSPMDRSTMVRFGRAPMDRSSMVRFGKRAPMDRSSMVR
PSLHQIYHVIVTMESSKMTSSDLFLRFGKRAPMDRSSMVRFGKRAPLDRSSMVR
IMHLSYILIGCAIQLGCNYNCGPRDVGSVVISGLRHNSCISRFGKRAPMDRSSMVRFGKRSE
>FLP-7_Teladorsagia_circumcincta_predicted
VNENIHKFLPCTLEDDCSHLFRFGKRAPMDRSTMVRFGRAPMDRSTMVR
SGRSVMLRLFHRFGKRAPMDRSSMVRFGKRAPMDRSSMVR
SGRSVMLRLFHRFGKRAPMDRSSMVRFGKRAPMDRSSMVR
>FLP-7_Bursaphelenchus_xylophilus_BXY_0630700.1
MLPQFKSLLVPLLLVVFFTGTMAQDILESYYKRAPMDRASMVRFGKRAPMDRASMVRFGK
RDSDDFELEKRAPMDRASMVRFGKRAPMDRASMVRFGKRAPMDRASMVRFGKRSDFDDIE
Y
>FLP-7_Panagrellus_redivivus_Pan_g12109.t1
MSRIAVYVLIVAACATVMAMAEEINEGNAFDRYERSPLARSAMVRFGKRAPLDRSAMVRF
GRAPLDRSAMVRFGKRAPLDRSSMVRFGKRAPLDRSAMVRFGKRDFSAEY
>FLP-7_Parastrongyloides_trichosuri_PTRK_0001640600.1
MAMFKKLLVPIFVILFTVYVLADENEKINLNESNVSNDIEQDQYQHDEPLGYNNQENVYD
NESIEKRAPLDRSSMVRFGKRAQLDRAMVRFGRSPMDRSSMVRFGRAPLDRSSMVRFGRA
PLARTSMVRFGKRAPLDRSSMVRFGKRAPLDRAMVRFGKRAPLDRAMVRFGKRFE
>FLP-7_Rhabditophanes_kr3021_RSKR_0000592300.1
MKFVGKFVIVLLLSTAILAVTSGLEAGKEEVELQEGLGDEVEENGEIGPLEPLNDNNSMY
EGDLNESEMINKRAPLDRSSMVRFGKRAPLDRSSMVRFGRSPVDRTRMVRFGKRSPLGRS
SMVRFGRSPLDRSSMVRFGKRSQVGRSSMVRFGKRTPHGRSSMVRFGRAPLDRSSMVRFG
KRSVN
>FLP-7_Steinernema_carpocapsae_SC.X.g3276
MFLAYGLLLLSSFYLLVSGFVLEEGMINEKRAPMDRSSMVRFGRAPMDRSSMVRFGKRAP
MDRSSMVRFGKRAPMDRSSMVRFGKRAPMDRSAMVRFGKRAPMDRSSMVRFGKRTSDVQS
EK
>FLP-7_Steinernema_feltiae_L889_g32992.t1
MNSSSYASFVHSNGKALLKQHLNSPLKNVILNRDRNLEALSHSVGPDPRRAGLLSWILLS
VSIFRVHTMILAYGLLVLSSICLLVNSFTLEENMMSEDKRAPMDRSSMVRFGRAPMDRSS

MVRFGKRAPMDRSSMVRFGKRAPMDRSSMVRFGKRAPMDRSAMVRFGKRAPMDRSAMVRF
GKRASETHSEE
>FLP-7_Steinernema_glaseri_L893_g27426.t1
MLIYATVVEHLHILPQPSVPVRSPRFRSRLSLLLIRVPTMFLAYGLLLLSSICLLVSSFT
LEDNMMTEEKRAPMDRSSMVRFGRSPMDRSSMVRFGKRAPMDRSSMVRFGKRAPMDRSSM
VRFGKRAPMDRSSMVRFGKRAPMDRSSMVRFGKRASDVSTEE
>FLP-7_Steinernema_monticolum_L898_g20838.t1
MFLAYGLLLLSSICLLVNGFAVDENIMSEDKRAPMDRSSMVRFGRAPMDRSSMVRFGKRA
PMDRSSMVRFGKRAPMDRSSMVRFGKRAPMDRSAMVRFGKRAPMDRSSMVRFGKRTSDVQ
SEE
>FLP-7_Steinernema_scapterisci_L892_g21875.t1
MFLAYGLLLLSSFCLLVSGFVLEDSLINEDKRAPMDRSSMVRFGRAPMDRSSMVRFGKRA
PMDRSSMVRFGKRAPMDRSSMVRFGKRAPIDRSAMVRFGKRAPMDRSSMVRFGKRTSDVQ
SEE
>FLP-7_Strongyloides_papillosus_SPAL_0001323500.1
MFMRLFILLVVACIATAVFGGDEEKYSPNTSSVSLNILPDQFLSDVEELSQYGDEINKED
DEGIEKRAPMDRSSMIRFGKRDQLDRAMVRFGRSPMGRSSMVRFGRAPLDRSAMVRFGRA
PLARTSMVRFGKRAPLNRAMVRFGKRAPLDRAMVRFGKRAPLDRAMVRFGKRSE
>FLP-7_Strongyloides_ratti_SRAE_X000053000
MFRRLFISIVIVCIATIVLGVESEDYPPNTSDILDDFQSDKLSSEHNQILEINGQYNNVE
DDENIEKRAPMDRSSMIKFGKRAQLDRAMVRFGRAPLDRSSMVRFGRAPLTRTSMIRFGK
RAPLDRAMVRFGKRAPLDRAMVRFGKRAPLDRAMVRFGKRAPLDRAMVRFGKRSE
>FLP-7_Strongyloides_stercoralis_SSTP_0000521900.1
MSRRLFTSIVIVCIATIVFGVEKETYPPNTSDVAIDFSENKYLSEVDGIPEYISQGNDVE
DNEAIEKRAPLDRSSMIRFGKRAPLDRAMVRFGRSPIDRSSMVRFGRAPLDRSSMVRFGR
APLARTSMIRFGKRAPLDRAMVRFGKRAPLDRAMVRFGKRAPLDRAMVRFGKRSE
>FLP-7_Strongyloides_venezuelensis_SVE_0688600.1
MTVSEGNEDDKIRKELYHLLILFCRIIDMFMRLFILLVVACIATAVFGGEENKYSPNTSS
VSLNILPDQFLSDVEELSQYGDGINKEDDEDIEKRAPMDRSSMIRFGKRDQLDRAMVRFG
RSPMGRSSMVRFGRAPLDRSAMVRFGRAPLARTSMVRFGKRAPLNRAMIRFGKRAPLDRA
MVRFGKRATLDRDMVRFGKRSE
>FLP-7_Halicephalobus_mephisto_predicted
SLDVFLTLVFRVYIRSEVLLAIIMNRISAYILVTICIMNVLAQSIDETSDEVLNSFDNIYGKRAPLDR
TSLVRFGKRAPLDRTSLVRFGKRAPLDRTSLVRFGKRAPLDRTSLVRFGKRDSSDFE
>FLP-7_Acrobeloides_nanus_ACRNAN_scaffold1810.g30830.t3
MDREVRAPMDRSSMVRFGKRAPMDRSSMVRFGRSSPDDSELVDKRAPMDRSSMVRFGKRA
PMDRSSMVRFGRSSPDDSELVDKRAPMDRSSMVRFGKRAPMDRSSMVRFGRAPMDRSSMV
RFGKRFSDDFSNFFNTNNEDKRR
>FLP-7_Ditylenchus_destructor_Dd_01328
MSIMVRLLSILFAVILFSSFLTDAISADSESKITELSADTNYIPFMNRFYGDVDPLTASY
RLWKRAPMDRSAMVRFGKRAPMDRSAMVRFGKRAPMDRSAMVRFGKRAPMDRSAMVRFGK
RVPDQFYNSVRYDEGVQFAEAV
>FLP-7_Ditylenchus_dipsaci_jg10872
MTQMYSVLISVVSFLLVVLAVVPRGSGSDASENSLADDYSLNNLLLSAEYGTPGLERFMK
RAPMDRSAMVRFGKRAPMDRSAMVRFGKRAPLDRSALVRFGKRAPMDRSAMVRFGKRAPM
DRSAMVRFGKRAPMDRTTMVRFGK
>FLP-7_Heterodera_glycines_Hetgly.G000001235
MAVLCCDAAVCPPPVVRFGVGFPQGQRAIALGCFLRLPPPPPQLLIPSNFLRIVPTSIHF
DSSPFTTFPLTLPFQIIVTHHKLPSQLKIQLSTRLSSAHSRKERYLLLPFVHSSPIRPFP
SQMAQFPVANILLASLLLGFVILNSKTNAQFQYGGGMLAEGPQMEGLLEDYGRVGDYPFE
SLAKRAPLDRSAMARFGKRAPLDRSALARFGKRAPLDRSAIARFGK
>FLP-7_Meloidogyne_arenaria_M.Arenaria_Scaff4717g049832
MSLQDFGDDYNNAIFADIDGDNENSYESMAKRAPLDRSALVRFGKRAPLDRSALVRFGKR
APLDRAAMVRFGKRAPLDRAAMVRFGKRAPLDRAAMVRFGKRAPFDRSSMVRFGKRK

>FLP-7_Meloidogyne_enterolobii_scaffold3590_cov163.g5768
MSLQDFGDDYNNAIFADIDGDNENSYESMAKRAPLDRSALVRFGKRAPLDRSALVRFGKR
APLDRAAMVRFGKRAPLDRAAMVRFGKRAPFDRSSMVRFGKRK
>FLP-7_Meloidogyne_floridensis_genemark-nMf.1.1.scaf02251-processed-gene-0.3-mRNA-1
MSLQDFGDDYNNAIFADIDGDNENSYESMAKRAPLDRSALVRFGKRAPLDRSALVRFGKR
APLDRAAMVRFGKRAPLDRAAMVRFGKRAPFDRSSMVRFGKRK
>FLP-7_Meloidogyne_graminicola FLP=7 NXFT01003587.1.9376_g
TLLFSALMVSHFSNGEISTDFGPQLMAIEDIGDDYNNAVLAEIDGDDENNYESIIKRAPL
DRSALVRFGKRAPLDRAAMVRFGKRAPLDRAAMVRFGKRAPLDRAAMVRFGKRAPFDRSS
MVRFGKRK
>FLP-7_Meloidogyne_incognita_Minc3s00073g03606
MAQIIFNTLLATLIFSAFFISRFSNGQLTNDFGSQLMSLQDFGDDYNNAIFADIDGDNEN
SYESMAKRAPLDRSALVRFGKRAPLDRSALVRFGKRAPLDRAAMVRFGKRAPLDRAAMVR
FGKRAPFDRSSMVRFGKRK
>FLP-7_Meloidogyne_javanica_M.Javanica_Scaff7529g048456
MSLQDFGDDYNNAIFADIDGDNENSYESMAKRAPLDRSALVRFGKRAPLDRSALVRFGKR
APLDRAAMVRFGKRAPLDRAAMVRFGKRAPFDRSSMVRFGKRK
>FLP-7_Globodera_pallida_predicted
TSTARPFCVGTVSNANIFLNFPHAKLHRFGKRAPLDRSAIARFGK
LVPFMGKQSYLIKSTAFDFVNNSFQKKAFHSIILVVIEYSVRFGKRAPLDRSALAR
>FLP-7_Globodera_rostochiensis_predicted
IEYSVRFGKRAPLDRSALAR
FNSIQLFKSNIPSDLVNEHRSTVLRWHGKQCQHFFKFSHVKLHRFGKRAPLDRSAIARFGK
>FLP-7_Meloidogyne_hapla_predicted
RFGKRAPLDRAAMVRFGKRAPFDRSSMVR
>FLP-8_Romanomermis_culicivorax_nRc.2.0.1.t02208-RA
MQISTILGSLCLLSFLAEFRQTAAVNVEESSSLGKCRQDLNNKLNDPILIEELCYLRNRI
DILSENMYQIVYPDKDVVKRKNEFVRFGRSSAPAFVEHPAEKRKNEFVRFGKRKNEFVRF
GRNSEDLIEAEKRKNEFVRFGRK
>FLP-8_Trichinella_britovi_T03_2772.1
MHMYNYFAIQLCIALLINFSLGQYMENTETASLYEACVMLYGSDSELCQNVSPVIQRSPS
FIKKAADKGGKALFVRFGRNMNLDEEKIKRKNEFVRFG
>FLP-8_Trichinella_murrelli_T05_12732.1
MHMYNNFAIQLCIALLINFSLGQYMENTETASLYEACVMLYGSDSELCQNVSPVIQRSPS
FIKKAADKGGKALFVRFGRNMNLDEEKIKRKNEFVRFG
>FLP-8_Trichinella_nativa_T02_5703.1
MHMYNYFAIQLCIALLINFSLGQYMENTETASLYEACVMLYGSDSELCQNVSPVIQRSPS
FIKKAADKGGKALFVRFGRNMNLDEEKIKRKNEFVRFG
>FLP-8_Trichinella_nelsoni_T07_9398.1
MHMYNYFAVQLCIALLINFSLGQYMENTEAASLYEACVMLYGSDSELCQNVSPVIQRSPS
FIKKATDKGGKALFVRFGRNMNLDEEKIKRKNEFVRFG
>FLP-8_Trichinella_patagoniensis_T12_13833.1
MHMYNYFAIQLCIALLINFSLGQYMENTETASLYEACVMLYGSDSELCQNVSPVIQRSPS
FIKKAADKGGKALFVRFGRNMNLDEEKIKRKNEFVRFG
>FLP-8_Trichinella_pseudospiralis_T4A_9392.2
MHMYNYFAIQLCIALLINFSLGQYMENTEAASLYEACVMLYGSDSELCQNVSPVIQRNPS
FIKKATDKGGKALFVRFGRNINLDEEKIKRKNEFVRFG
>FLP-8_Trichinella_sp.T6.fa_T06_1278.2
MHMYNYFAIQLCIALLINFSLGQYMENTETASLYEACVMLYGSDSELCQNVSPVIQRSPS
FIKKAADKGGKALFVRFGRNMNLDEEKIKRKNEFVRFG
>FLP-8_Trichinella_sp.T8_T08_6619.1_
MHMYNYFAIQLCIALLINFSLGQYMENTETASLYEACVMLYGSDSELCQNVSPVIQRSPS
FIKKAADKGGKALFVRFGRNMNLDEEKIKRKNEFVRFG

```
>FLP-8_Trichinella_sp.T9_T09_10772.1
MHMYNYFAIQLCIALLINFSLGQYMENTETASLYEACVMLYGSDSELCQNVSPVIQRSPS
FIKKAADKGGKALFVRFGRNMNLDEEKIKRKNEFVRFG
>FLP-8_Trichinella_spiralis_EFV56802
MHMYNYFAVQLCIALLINFSLGQYMENTEAASLYEACVMLYGSDSELCQNVSPVIQRSPS
FIKKATDKGGKALFVRFGRNMNLDEEKIKRKNEFVRFG
>FLP-8_Trichinella_zimbabwensis_T11_11695.1
MHMYNCFAIQLCIALLINFSLGQYMENTEAASLYEACVMLYGSDSELCQNVSPVIQRSPS
FIKKATDKGGKALFVRFGRNINLDEEKIKRKNEFVRFG
>FLP-8_Trichuris_muris_TMUE_2000009415.1
MLDECILVYGLESPLCQRIAAMQLRQAPMVKKTTERGGKALFVRFGRSDQLEDAKRKRKN
EFVRFG
>FLP-8_Trichuris_suis_M514_27192
MLDECILAYGLESPLCQRIAAMQLRQASIVKKTNERGGKALFVRFGRSGLLEDRKLKRKN
EFIRFG
>FLP-8_Trichinella_papuae_predicted
FYQKLIAFILEEKIKRKNEFVRFG
LSILVDFFLILIIKVNVFLGGKALFVRFGRNINLDGKYKLTCDNFIKS
>FLP-8_Trichuris_trichiura_predicted
ENILKAILLRKFIDRKLKRKNEFIRFG
LSGILIAYIPGGKALFVRFGRSGLLEGKKTFLRLFY
>FLP-8_Plectus_sambesii_PSAMB.scaffold1604size39546.g14051.t1
MRLLAVIVGLGVLAATQAATDLAAICAKAADAPKSEDGKDVYTHLCQLQLELVEVKDVIH
EMLSDRESAAEEMESAIPEEVEAPSADKRKNEFIRFGKRKNEFIRFGKRKNEFIRFGKRK
NEFIRFGRSAANEEPMEKDEMTKRKNEFIRFGKRKNEFIRFG
>FLP-8_Ascaris_lumbricoides_ALUE_0001294401-mRNA-1
MALVSLIVVAVFWTSSSAREEDIQCSLAALVPPQQKFLADRLCRLEDSTIAMEHVLQELM
NRADITMESTASNTATDEDQAIAKRKNEFIRFGKRKNEFIR
>FLP-8_Ascaris_suum_AgR009X_g171_t02
MQRVTVLHLFFKKVFFQLAQPNPSSATIAEWWSVNRSANRKDSLSANMALVSLIVVAVFW
TSSSAREEDIQCSLAALVPPQQKFLADRLCRLEDSTIAMEHVLQELMNRADITMESTASN
TATDEDQAIAKRKNEFIRFGKRKNEFIRFGKRKNEFIRFGKRKNEFIRFGRSGYQNDNNI
VEKRKNEFIRFG
>FLP-8_Brugia_malayi_Bm1088.1
MTCVPSVSNAFIQVLLVMVLVSHVLPLPLAAHLPVYESLYDEPPLADYNFVPYSGLRNNG
DWMNEPFRADKRKNEFIRFGKRDDFMKFNKRKNEFIRFGKRSIN
>FLP-8_Brugia_pahangi_BPAG_0001235501-mRNA-1
MVYLVSRKMSLDDGWERNAERRDHLKANISTVPFHQINFDIQEAMTCVPSVSNAFIQVLL
VMVLVSHVLPLPLAAHLPVYDSLYDEPPLADYNFVPYSGLRNNGDWMNEPFRADKRKNEF
IRFGKRDDFMKYSGVA
>FLP-8_Brugia_timori_BTMF_0001181701-mRNA-1
MSLDDGWERNAERSEIEKGALEWIERWSSRDHLKANISIVPFHQINFDIQEAMTCVPSVS
NAFIQVLLVMVLVSHVLPLPLAAHLPVYESLYDEPPLADYNFVPYSGLRNNGDWMNEPFR
ADKRKNEFIRFGKRDDFMKYSGVA
>FLP-8_Litomosoides_sigmodontis_nLs.2.1.2.t06558-RA
MILVPHMLPLPLAAHLPAHDSLYDEPPLVDYNFMPYPGMRINDDLMDEPFRADKRKNEFI
RFGKRNQFTRFNKRKNEFIRFG
>FLP-8_Loa_loa_EFO18677.1
MVYLASRKVSLNDEWKGEKYADRNKMEKEALEWIERWSSMLLVMVLVPDILPLSLAAHLP
VHDSLYDESPLVDYNILPYSGMRINDDLMNEPFQVDKRKNEFIRFGKRNGFMKFNKRKNE
FIRFGKRSIKLKMF
>FLP-8_Onchocerca_volvulus_OVOC3976.1
MKCLILTDRFYAERTATFKTTAMTFIPTVSNAFIQVLLIMVLVPNVLPLPFAVHLPVHDS
FYDEPPVVDYNFVPYSGLRINDDLMNEPFRVDKRKNEFIRFGKRDDPMKFKRKNEFIRFG
```

KRSVKLKKF
>FLP-8_Toxocara_canis_Tcan_15961.1
MALLSLIVLVVCWLGGSSANDVEENAFCPLTALVPQQQKPLAYSICKLEDSIIVIEHILQ
ELMSRTDITLGTTATDDNDHSMEKRKNEFIRFGKRKNEFIR
>FLP-8_Wuchereria_bancrofti_maker-PairedContig_3974-snap-gene-1.14-mRNA-1
MMGERGMRREDHLKANISVVPFHQINCDIQEAMTCVPSVSNAFIQMLLVMVLVSHVLPLP
LAAHLPVYDSLYDEPPLVDYNFVPYSGLRNNGDWMNEPFRADKRKNEFIRFGKRDDFMNG
IIRFVCRFNKRKNEFIRFEGKRTHELLEPITYNRGNDRELVAKAEEQLLNTLSLLEALAD
DSNQLQMEKRRNKFEFIRFGRR
>FLP-8_Acanthocheilonema_viteae_predicted
FVCRFNKRKNEFIRFGKRSIKLNML
LLIFVGMRLNGDLMNEPFRADKRKNEFIRFGKRNDFMKLASSCHTVFFF
>FLP-8_Anisakis_simplex_predicted
KLFTLRKFEHFYRFGKRKNEFIRFGKRKNEFIRFVTD
RNSGKSLNTDRRLPFRKKRGRLYEYNVFLFEEVRLLGRIARKEQNKVGDGNEQIPRQSAVEKRKNEFI
RFGKRKNEFIRFGKRKNEFIR
>FLP-8_Dirofilaria_immitis_predicted
ILLGMRIDDELLIDEQPFQVDKRKNEFIRFGKRNDLMK
>FLP-8_Dracunculus_medinensis_predicted
NEFINEKICKIASNFASFRFGKRKNEFIRFGKRIQNR
>FLP-8_Elaeophora_elaphi_predicted
ILVGMRTSDDLMNELFRADKRKNEFIRFGKRNDFMKYAHFVMTYHFFF
NLNNISNGAIGIYYFCRFDKRKNEFIRFGKRFIVLKML
>FLP-8_Gongylonema_pulchrum_predicted
DVHMFPGPRTSGSLINSPFLVEKRKNEFIRFGKRKADPMRYESYRXXXXXXXXXXXXXXXXXXXXXXX
XXXXXXXXXX
>FLP-8_Onchocerca_ochengi_predicted
LILNLNPNLNFLIGLRIYDDLMNEPFRVDKRKNEFIRFGKRDDPMKYVCFVVLYCKNFLFLLFKYMFK
LSLSVAI
FVCRFKRKNEFIRFGKRSVKLKKF
>FLP-8_Parascaris_equorum_predicted
XXXXXXXXXXXXXXXXXXXXXXXXXXXXXXXXXXXWKSKDEFKSTLHTIEACEVYRAEISSKTCFR
FGKRKNEFIRFGKRKNEFIR
>FLP-8_Parascaris_univalens_predicted
MSFQITTWKSKDEFKSTLHTIEACEVYRAEISSKTCFRFGKRKNEFIRFGKRKNEFIR
>FLP-8_Thelazia_callipaeda_predicted
NFTEVKISDNAMVNPYLLYKRKNEFIRFGKREDDMRYLFFFFNKHIKYLWKWKKYYYIFHSIYRNLFQ
IWTFILCNFDAKKIVILY
>FLP-8_Ancylostoma_caninum_ANCCAN_08870
MTISSFDMLLVLLGATLIASGFSYECDVGSFPESQRELGRHVCRLENEVSVLEAAVQEML
QRADVVLNSDEPTVEKRKNEFIRFGKRSLNDVKRKNEFIRFGKRKNEFIRFGRSDPLILD
DAAVEKRKNEFIRFG
>FLP-8_Ancylostoma_ceylanicum_maker-ANCCEYDFT_Contig140-augustus-gene-3.17-mRNA-1
MTISSFDMLLVLLGATLIASGFSYECDVGSFPESQRELGRRVCRLEEMLQRTDVVLNSDE
PTLEKRKNEFIRFGKRSLKFGRSDPLILDDAAVEKRKNEFIRFG
>FLP-8_Ancylostoma_duodenale_ANCDUO_14654
MTISSFDMLLVLLGATLIASGFSYECDVGSFPESQRELGRRVCRLENEVGVLEAAVQEML
QRTDVILNSDEPTVEKRKNEFIRFGKRSLNDVKRKNEFIRFGKRKNEFIRRETQERVHPL
RLSNSGLEI
>FLP-8_Angiostrongylus_costaricensis_ACOC_0000299001-mRNA-1
MLLVLLGTCFFAGGVIGYGCDVASFSEEQHGLGRRVCNLENEVDALKSAVQEILQRTDHT
FNSEPAMEKRKNEFIRFGKRSLTDVKRKNEFIRFGKRKNEFIRFGRFDSTLGGSDVVEKR

KNEFIRFG
>FLP-8_Caenorhabditis_angaria_Cang_2012_03_13_00003.g275.t1
MLVGILLAILVAFVSSISAECDVSTIPDAQKDLALRVCHLEDELSVVQRALQEVMQRTDI
TLDDAAPILNKRKNEFIRFGKRSGEVDKRKNEFIRFGKRKNEFIRFGRSDPALYDDVSME
KRKNEFIRFG
>FLP-8_Caenorhabditis_brenneri_CBN08812.1
MLLGVVISIFVLAISANANCDVSALSTETEKELGLRLCHLEAEMQVVQRALQEVMQQTDV
TMYDQEVPVMNKRKNEFIRFGKRSDGMEKRKNEFIRFGKRKNEFIRFGRSDKGLGLDNDN
VSMEKRKNEFIRFG
>FLP-8_Caenorhabditis_briggsae_CBG07605.1
MLLGVVFSIFVLAISAHATCDVSALATESEKELGLRLCRLESEMQVIERALQEVMQQTDV
TSFDQEVPAMNKRKNEFIRFGKRSDGMEKRKNEFIRFGKRKNEFIRFGRSDKGLGLDDNV
SMEKRKNEFIRFG
>FLP-8_Caenorhabditis_elegans_Isoform B F31F6.4b.1
MLSGVLFSIFVLAISANASCDVSALTTENEKELGLRICHLEAEMQVVQRALQEVMQQTDV
TLYDQEVPVMNKRKNEFIRFGKRSDGMEKRKNEFIRFGKRKNEFIRFGRSDKGLGLDDND
VSMEKRKNEFIRFG
>FLP-8_Caenorhabditis_inopinata.fa_Sp34_X0140600.t1
MLLGVFISILVLAASAQASCDVSAFSNDNEKELGIRICHLEAEMQVVQRALQEVLQQADV
TLSEQEVPVMNKHKNEFFRFEKHSDGMEKRKNEFIRFGKRKNEFIRFGRSDKGLYLNTDN
DISTEKRKNEFIRFG
>FLP-8_Caenorhabditis_latens_FL83_01711
MLLGVVISIFVLAISANANCDVSALNTETEKELGLRICHLEAEMQVVQRALQEVMQQTDV
TLYDQEVPVMNKRKNEFIRFGKRSDGMEKRKNEFIRFGKRKNEFIRFGRSDKGLGLDNDV
SMEKRKNEFIRFG
>FLP-8_Caenorhabditis_nigoni_Cni-flp-
MLLGVVFSIFVLAISAHATCDVSALATESEKELGLRLCRLESEMQVIERALQEVMQQTDV
TSFDQEVPVMNKRKNEFIRFGKRSDGMEKRKNEFIRFGKRKNEFIRFGRSDKGLGLDDNV
SMEKRKNEFIRFG
>FLP-8_Caenorhabditis_remanei_FL81_03726
MLLGVVISIFVLAISANANCDVSALNTETEKELGLRICHLEAEMQVVQRALQEVMQQTDV
TLYDQEVPVMNKRKNEFIRFGKRSDGMEKRKNEFIRFGKRKNEFIRFGRSDKGLGLDNDV
SMEKRKNEFIRFG
>FLP-8_Caenorhabditis_sinica_Csp5_scaffold_00203.g6951.t1
MLLGVVFSIFVLAISANANCDVSALATESEKELGLRLCHLEAEMQVIERALQEVMQQTDV
TSFEQEVPVMNKRKNEFIRFGKRSDGMEKRKNEFIRFGKRKNEFIRFGRSDKGLGLDNDV
SMEKRKNEFIRFG
>FLP-8_Caenorhabditis_tropicalis_Csp11.Scaffold629.g10025.t1
MLLGVVISIFVLAISANANCDVSALSTETEKELGLRICHLEAEMQVVQRALQEVMQQTDV
TTYDQDVPVMNKRKNEFIRFGKRSDGVEKRKNEFIRFGKRKNEFIRFGRSDKGLGLDNDD
VSMEKRKNEFIRFG
>FLP-8_Dictyocaulus_viviparus_nDv.1.0.1.t07631
NEIDVLKAAVQELLQRSDLTLNNEFVPAMEKRKNEFIRFGKRSSKIFQIRIIVEFFSIVK
RKNEFIRFGKRKNEFIRFGRFDPSLRDDNAVGKGLC
>FLP-8_Diploscapter_coronatus_DCO_024386
MQLVIAGGLLLVAVASCVFASSDSDCNGPEKERLCRIEGAIVVLENAVQELLDRTDITMA
ETEVPQKRKNEFIRFGKRSINNEKRKNEFIRFGKRKNEFIRFG
>FLP-8_Diploscapter_pachys_WR25_14892.1
MQLVIAGGLLLVAVTSCIFASSDSDCNGPEKERLCRIEGAIVVLENAVQELLDRTDITMA
ETEVPQKRKNEFIRFGKRSINNEKRKNEFIRFGKRKNEFIRFG
>FLP-8_Haemonchus_contortus_HCON_00180390
MLFVLVVATFVASVASYRSDVAAFPEHFRRIGHNEYRVKAESNNNNGVLNPLVQDPLQRT
DTVLNELVEPTLQKRNNDFICSGKRERRNVKRKNEFIRFGKRGDDLVRFGRSDSTETEPH
VAKRKNEFIRFG

>FLP-8_Mesorhabditis_belari_mbelari.g26501.t1
MIRFQRSSCFHHCGLSHCLHHQSWSSPVIKQTRFSDGFAFLRTRSECLETPCKRCSPEQT
SLLRSQSQNRLRRERTSSFDSESDQLSQKTWRWREKKNEFIRFGKRKNEFIRFGKRKNEF
IRFGRSAPEMMDDGNMEKRKNEFIRFG
>FLP-8_Micoletzkya_japonica_MicoRS5524-mkr-S25-2.15-mRNA-1
MLVPVAVLSVLLGLSLSCELSSYPEDQHPLAQRLCQLENEMRVMERVMQEVLQRADLTVY
PEVENLAIAPEKRKNEFIRFGKRSEGETDKRKNEFIRFGKRKNEFIRFGRSLPSEAEMEK
RKNEFIRFG
>FLP-8_Necator_americanus_NECAME_10076
MLVVLLGATLFASGFSYECDVTNFPESQRDLGRRVCRLENEVGMLEAAVGEMLQRTVPAM
NSDDEPNIEKRKNEFIRFGKRSLSDVKRKNEFIRFGKRKNEFIRFGRSDPFFADDATVEK
RKNEFIRFG
>FLP-8_Nippostrongylus_brasiliensis_NBR_0001012101-mRNA-1
MLFGLLVAAIVASGVSYECDLTQLPEQLHDLGRRVCRLENKLGVLDAVAPDALQGSEFVI
NDEEAPNMEKRKNEFIRFGKRSTNDVVKRKNEFIRFGKRKNEFIRYDFASAFDLEGHRTQ
VLAAPVVSELEKRKNEFIRFG
>FLP-8_Oesophagostomum_dentatum_OESDEN_03497
MTISSFDMLLLLLGATFIASSYTFECDVNSFPESQRELGLRVCRLENEVGILEAAVQEML
QRTDITMISGGEPSVEKRKNEFIRFGKRSLNDVKRKNEFIRFGKRKNEFIRFGRSDPLLY
DGAPVEKRKNEFIRFG
>FLP-8_Oscheius_tipulae_OTIPU.nOt.2.0.1.t00401
MQLALALTVLTIASVFAECDLGGLSESQRPMALRVCKLEQEVAVLENAVQELLQRADITL
NEEETAAPQKRKNEFIRFGKRKNEFIRFGKRKNEFIRFGRSDPAFQEDYSMEKRKNEFIR
FG
>FLP-8_Parapristionchus_giblindavisi_Parapristionchus-mkr-
S_210-0.55-mRNA-1
MIAPSALILLSVGVVFSCDLSDVPEASRSLAEKVCQLEKEMKVMESVLQELLQRIDITVL
PDASDISAAPEKRKNEFIRFGKRSSDGEVEKRKNEFIRFGKRKNEFIRFGRSLPMENEMI
KRKNEFIRFG
>FLP-8_Pristionchus_arcanus_arcanus-mkr-S_307-1.1-mRNA-1
MNPIVLVSLLAGVALSCDLTNVPEGSKDLAEKVCTLEKEMRVIEHVLQELLQRADITVSD
EDLPAPEKRKNEFIRFGKRSADGEMEKRKNEFIRFGKRKNEFIRFGRSMPVEGEMLKRKN
EFIRFG
>FLP-8_Pristionchus_entomophagus_entomophagus-mkr-S267-2.15-mRNA-1
MLPIVMLSLLAGAVLSCDLTNVPEGSKELAEKVCTLEQEMRVIERVLQELLQRTDITISD
EDLAAAPEKRKNEFIRFGKRSAPDGDMEKRKNEFIRFGKRKNEFIRFGRKNEFIRFG
>FLP-8_Pristionchus_exspectatus_exspectatus-mkr-S_231-2.55-mRNA-1
MHPIVLVSLLAGVALSCDLTNVPEGSKDLAEKVCTLEKEMRVIERVLQELLQRADITVSD
EDLPAPEKRKNEFIRFGKRSADGEMEKRKNEFIRFGKRKNEFIRFGRSMPVEGEMLKRKN
EFIRFG
>FLP-8_Pristionchus_fissidentatus_fissidentatus-mkr-S246-5.45-mRNA-1
MGGVKAVLGGIKGPRGGVSPHYLTLDRDSRLSLISTLIMISLLLVSLLAASTLACDLTNV
PEANKELAEKVCALEQEMRVAERVLQELLQRADLTVSDEEFTAPQEKRKNEFIRFGKRSA
DGEMEKRKNEFIRFGKRKNEFIRFGRSAPVDGEMTKRKNEFIRFG
>FLP-8_Pristionchus_japonicus_japonicus-ag_msk-S19-5.31-mRNA-1
MLPIVLVSLLAGAALSCDLTNVPEGSKELAEKVCTLEKEMRVIERVLQELLQRADITVSE
EDLPAPEKRKNEFIRFGKRSADGEMEKRKNEFIRFGKRKNEFIRFGRSMPVEGEMLKRKN
EFIRFG
>FLP-8_Pristionchus_maxplancki_maxplancki-mkr-S40-2.51-mRNA-1
MLPIVLVSLLAGAALSCDLTNVPEGSKELAEKVCTLEKEMRVIERVLQELLQRADITVSE
EDLPAPEKRKNEFIRFGKRSADGEMEKRKNEFIRFGKRKNEFIRFGRSMPVEGEMLKRKN
EFIRFG
>FLP-8_Pristionchus_pacificus_PPA07533.1
MHPIVLVSLLAGVALSCDLTNVPEGSKDLAEKVCTLEKEMRVIERVLQELLQRADITVSD

EDLPAPEKRKNEFIRFGKRSADGEMEKRKNEFIRFGKRKNEFIRFGRSMPVEGEMLKRKN
EFIRFG
>FLP-8_Strongylus_vulgaris_SVUK_0000963701-mRNA-1
MTILSFDMLLLLLGATLIVSGFTYECDVTSFPESQRELGQHVCRLENEVGVLEAAVQEML
QRAGMIFEIYNRSGTARYCLSNLEILADITLSNGDEPSIEKRKNEFIRFGKRSLNDVKRK
NEFIRFGKRKNEFIRFGRSDPLLYDGAPVEKRKNEFIRFG
>FLP-8_Angiostrongylus_cantonensis_predicted
STPRALSFILFIVFDTRFKNSWFRFGRFDSTLGDNGVGKTFHENQLTLFFLRVL
QYFFSDVKRKNEFIRFGKRKNEFIR
RCFTNTRKSSVIPDYTFNIEPAMEKRKNEFIRFGKRSLT
KSTNAIFFTSSLKTVFSVEKRKNEFIRFG
>FLP-8_Cylicostephanus_goldi_predicted
AIVTDITLNSSDEPSIEKRKNEFIRFGKRSLK
MPKVYICSLKYVCCFSDVKRKNEFIRFGKRKNEFIR
RGKTNSFDLARERMNLFGEKEPFFAENSRKLVHFRFGRSDPLLYDGAAGRNFFAFFAALTTPSFGLFS
QHLHLVFSVEKRKNEFIRFG
>FLP-8_Heterorhabditis_bacteriophora_predicted
QQTILMLHKCFSDLTLDREPQLVDKRKNEFIRFGKRSMR
IITVFKNKKFHSSDVKRKNEFIRFGKRKNEFIR
SMSKLYLFYKFKEMIVEIVIQTLFLGMEKRKNEFIRFG
VTPLHSSFRFGRSDPTLIDDSLGA
>FLP-8_Teladorsagia_circumcincta_predicted
NYFCNFCKACFTTDLHCLLFSEVKRKNEFIRFGKRKNEFIR
YLNYIFQRKKRVSVAKRKNEFIRFG
>FLP-8_Bursaphelenchus_xylophilus FLP_8 BXY_1650300.1
MSSVLPSLLLVVMVGITVGQKAYPEDELRVCELTDLETLSLSKCELEGRIAELENKIREI
EDTVMSQGNHVHVKGEVAKRKNEFIRFGKRSLATVLDHMPHSRWNPHSDNALRSLSMVKR
KNEFIRFG
>FLP-8_Ditylenchus_destructor_Dd_01519
MSKFENARIAHLEALVDILLPSVASAQTYAEANPSTENLHILRKEKRKNEFIRFGKRANR
ESDRPVKFASLLLVCLFSRFAVCAQKSLPGLEHCNFYKPTRKQHLFRLNLQRREETNSSS
SALVDEDGE
>FLP-8_Ditylenchus_dipsaci_jg11775
MHLPTIASIFLLAFLQISYCSAVDNFNEIGQDQNNNWISLLSLVRNNALQQKDTVEQCPI
TDMELLGLSHCELMQRIVMLEDEIESLTRKSTSEVTDKSGPSRNKRKNEFIRFGKRSAGD
NSALPEMLRFTKTMSPSLSISDMQIPANIQRQSEKLSLKKRKNEFIRFG
>FLP-8_Halicephalobus_mephisto_MEPH_14855-RA.p1
VLNTVSNTMSTVYSLSISFLALLVWASAQDIQLLDQQQNSCEVMDPAMMQLRICELQHQI
EKLENLVVNGFVSASGGDSEAAMKQKRKNEFIRFGKRKNEFIRFGKRKNEFIRFGRSTGG
NIDESGEQFVAKRKNEFIRFG
>FLP-8_Panagrellus_redivivus_Pan_g7133.t1
MSAMNYLLFSTLVCLAIGSAVEFDYEECRIPTEIDEASLHKKLCELEHRVSIMEIRLEFG
KFGTAPPFDTEVSTPPHMQKRKNEFIRFGKRKNEFIRFGRSNGGADDSQFVAKRKNEFIR
FG
>FLP-8_Parastrongyloides_trichosuri_PTRK_0000527600.1
MASLDIGFFKRSFLFLCVIMLIHMTICENSEHSNQDNFYNKNSVFEEGSNFDIPKNIDFE
NIYGESNMQKRKNEFIRFGKRKNEFIRFGKRDNLPSENILNDQLYITYPNHDLFYNNGIH
SAYKRKNEFIRFGKK
>FLP-8_Rhabditophanes_sp.kr3021_RSKR_0000870400.1
MGIKTITLIFIIHYLIFKVHSFGALDFNDLGANAEVGMEENEFVCPSGFACSESGEMETP
INDHNLLETPHDNELPIMKRKNEFIRFGKRKNEFIRFGKRGMRPEDAFRQSVGKRAMRPE
DAFRQSYGKFGYFPFSVKYKHHQKRKNEFIRFG
>FLP-8_Steinernema_carpocapsae_SC.X.g3516
MHPKMLPLLALIIAFPSLAAFSDGSCRTGAPLTTESIGYDLCTLEKKVEMMQIVIQDLLK

HLDIPTSLIEEGDLKQMVSKRKNEFIRFGKRKNEFIRFGKRKNEFIRFGRSADDLSYEPT
VALEKRKNEFIRFG
>FLP-8_Steinernema_feltiae_L889_g30090.t1
MQPKMLPLLALIIASPSFIASSDVSCRTGAGLVVESLGYDLCHLERKIEAMQIMLQELLK
NPELAAPLLQEADLKEVASKRKNEFIRFGKRKNEFIRFGKRKNEFIRFGRSASELEFEPT
AAIEKRKNEFIRFG
>FLP-8_Steinernema_glaseri_L893_g3329.t1
MQPKMLPLLALILASPAFSASSDVSCRSGAALQDGSYELCRLERKLEVLQLVVQDLMKRL
ELASPLLDETDFKEVGSKRKNEFIRFGKRKNEFIRFGKRKNEFIRFGRSASDDLSFEPTV
ALEKRKNEFIRFG
>FLP-8_Steinernema_monticolum_L898_g27589.t1
MHPKMLPLLALIIAAPSFAALSDVSCRDGAAFAVDTIGYDLCHLEKKVEAMQIMLQQLLK
SLEGAAPLPSLKDTVSKRKNEFIRFGKRKNEFIRFGKRKNEFIRFGRSVDDINFEPMAAI
EKRKNEFIRFG
>FLP-8_Strongyloides_papillosus_SPAL_0001329500.1
MVSPQFGCFRYTLVIVCLLTVFKLNSCKNSDDLHQIKEMAKNYIVERDVNNYLPEQYDIY
RNFDVNSLEKRKNEFIRFGKRKNEFIRFGKRNNVNYENGVNDQISRIYPIYDFIDINDVH
KANKRKNEFIRFG
>FLP-8_Strongyloides_ratti_SRAE_X000149400
MASSHVGWFRYTFLIVYLITLLQVNFSKNTDDLRQIKYMTKNFPLDNEINSENLINLPDQ
NDIYKIFDIASLQKRKNEFIRFGKRKNEFIRFKKKNINNFNNELNNKMLPIYSIFDINDN
DDIHTANKRKNEFIRFG
>FLP-8_Strongyloides_stercoralis_SSTP_0000883300.1
MASPHFGWFRYAFFIVYLITLFQVNFSKTTDDLRQLKYMTKNFPLDNEMNNDNFINLPDQ
NDIYKIFDIASLQKRKNEFIRFGKRKNEFIRFKKRNTKTFYNKLNDQILPVVSIYDIIDN
DNIHSTNKRKNEFIRFG
>FLP-8_Strongyloides_venezuelensis_SVE_0100900.1
MVSPQFGCFRYTLVIVCLLTVFKLNLCKNSDDLHQMKEMAKNYIIERDVNNYLPEQYDIY
KNFDVNSLQKRKNEFIRFGKRKNEFIRFGKRNNVNYESGWNDQISQIYPIYDFIDSNNVH
TANKRKNEFIRFG
>FLP-8_Acrobeloides_nanus_ACRNAN_scaffold3634.g17271.t1
MKCCAGVELTVFISLLSIATVFGAEPGKTNVSKPENDQMCEIASLEMLSYTRCDLMNRVK
TLELLVEALLEQLSNELPIVTPERDSTSIEKRKNEFIRFGKRKNEFIRFGKRKNEFIRFG
RSMGHNLDQNEQSNMDKRKNEFIRFG
>FLP-9_Ancylostoma_caninum_ANCCAN_05889
MHSVSNVTLQDPEALIDYCAQPQNREVCEQLLASVIAEEQDSLPQVDKRKPSFVRFGKRA
ALMEKRKPSFVRFGRK
>FLP-9_Ancylostoma_duodenale_ANCDUO_15994
MGQKPSVEHSTATITSEDPEALIDYCAQPQNREVCEQLLASVIAEEQDSLPQVDKRKPSF
VRFGKRAALMEKRKPSFVRFGRK
>FLP-9_Angiostrongylus_cantonensis_ACAC_0001202001-mRNA-1
MRLFLRSWDTRAGVLPFGWQSYFFPHKSAINFLSSRWMEAWCVWCGVEPVPSFQLFSEFW
HISKHSNQWIQFQVVKRKPSFVRFGKRGDVMSKEKRKPSFVRFG
>FLP-9_Caenorhabditis_angaria_Cang_2012_03_13_00316.g9061.t1
MNTQIIAIFVLALIMMASGMQSVVNDEEFEEALQLAELCRTPQLKEICRKLGEQMAEDGM
LEMKEKRKPSFVRFGKRSGIEMEKRKPSFVRFGRK
>FLP-9_Caenorhabditis_brenneri_CBN17451.1
MNQFYALLLIVCVATVAIAFEEPDLDALADFCGKESNRKYCDQIAQLATQHAMDLNQEQV
RMEKRKPSFVRFGKRSGYPLIIDSEELRMDKRKPSFVRFGRK
>FLP-9_Caenorhabditis_briggsae_CBG22307.1
MSRLYALLLIVCIANVASTAPESIPDLDALAEFCAKESNKRYCAQLAQLSLDSAMEANQE
QVIQMEKRKPSFVRFGKRSGYPLIIDNEELRMDKRKPSFVRFGRK
>FLP-9_Caenorhabditis_elegans_C36H8.3.1
MNQFYALFLVACIAAMANAYEEPDLDALAEFCGKESNRKYCDQIAQLATQHAIGINQEQV

RMEKRKPSFVRFGKRSGYPLVIDDEEMRMDKRKPSFVRFGRK
>FLP-9_Caenorhabditis_inopinata_Sp34_40152700.t1
MSQLYAIAFFVCFAVVVASSYQEPDLDALAEFCEKEANRKYCDQIAQLATQRAMELNQEQ
VRMEKRKPSFVRFGKRSGYPLVTDDEEMRMDKRKPSFVRFGRK
>FLP-9_Caenorhabditis_japonica_CJA03095.1
MIAVLAARKTSEAYEYPQGEAEGEAESNYPGQMEKRKPSFVRFGKRSSSDYPLVLNQEEL
QMNKRKPSFVRFGRK
>FLP-9_Caenorhabditis_latens_FL83_13769
MSQLYALLLIVCIASVATAFQEPDLDALAEFCGKESNRKYCDQIAQLATQHAMDVNQEQV
QMMEKRKPSFVRFGKRSGYPLIIDGEEMRMDKRKPSFVRFGRK
>FLP-9_Caenorhabditis_nigoni_Cni-.1
MSRLYALLLIVCMANVASAAPESIPDLDALAEFCAKESNKKYCAQLAQLSLDSAMEANQE
QVQQMEKRKPSFVRFGKRSGYPLIIDNEELRMDKRKPSFVRFGRK
>FLP-9_Caenorhabditis_remanei_CRE10147.1
MSQLYALLLIACIASVATAFQEPDLDALAEFCGKESNRKYCDQIAQLATQHAMDVNQEQV
QMMEKRKPSFVRFGKRSGYPLIIDGEEMRMDKRKPSFVRFGRK
>FLP-9_Caenorhabditis_sinica_Csp5_scaffold_00901.g16596.t1
MSQLYALLLIVCIANVASAFQEPDLDALAEFCGKESNRKYCEQIAQLATQHAMEVNQILD
PSCHRNLPFNPVNVQEPVQQMEKRKPSFVRFGKRSGYPLIIENEELRMDKRKPSFVRFGR
K
>FLP-9_Caenorhabditis_tropicalis_Csp11.Scaffold484.g1903.t2
MNKFYALLLIVCIANVANAYEEPDLDALAEFCGKESNRKYCDQIAHLATQHALDVNQEQV
RMEKRKPSFVRFGKRSGYPLIIDNEEMRMDKRKPSFVRFGRK
>FLP-9_Dictyocaulus_viviparus_nDv.1.0.1.t00298
MRSIVFTILSMILIAVVTKISGQDYEALAEYCSIPQNRDVCEQLLSELVDDEDRLPRIDK
RKPSFVRFGKRVNVAPIEKRKPSFVRFGRK
>FLP-9_Diploscapter_coronatus_DCO_024680
MRCLLYLLFSLLLLIEYSVLQIEALEVRTFCNEPQNEALCEQLIEEIRRSQNDIGQPKLS
VPEKRKPSFVRFGKRSAAPSKDEILLRQMADWN
>FLP-9_Haemonchus_contortus_HCON_00131250
LCTTPFVMKSIKLTILSLISAVVLVKVSAQDQDALADYCAQPQNREVCEQLLSALSARFE
TVPQMDKRKPSFVRFGKRSEGDSAMEKRKPSFVRFGRK
>FLP-9_Heligmosomoides_polygyrus_HPOL_0000342001-mRNA-1
MDYDGVLSWTMDQEALAEYCAQPQNREVCDQLLSALIEGSDTLPQMDKRKPSFVRFGKRS
VGEVEKRKPSFVRFGRK
>FLP-9_Heterorhabditis_bacteriophora_Hba_18025
MAIYYTASTLIQTANSWSQWYWLATILQSPLHHAFWGPPLVVDLLNGIDWQLDTVLCVLT
LNIYYEKLFIFLWFWLLFVAIISTSNSIYWVVNLCVGTKEEGTLIEFCRESQNKELCEQL
LGAYLEQETEHQTHAEKRKPSFVRFGKRSLQAEKRKPSFVRFGRK
>FLP-9_Micoletzkya_japonica_MicoRS5524-mkr-S2-9.0-mRNA-1
MLAVLFLLPLALANVEEQQPPTVEDKITTVDEFCASFPNLYLCHLRDTLEGSLFELANLQ
STLLRHIGEPRVDVSDASASATGAGEVEKRKSSFVRFGKRAAADLDDDAIVEQKRKSSYV
RFGKRSESAPDEFLGDKVDEKRKSSFVRFGKRKSSYVRFG
>FLP-9_Necator_americanus_NECAME_10565
MVNPFLSQSNLRTSSVLSRNVRGSSTAFCHQLQYPCSLQDNEALVNYCAQPQNREVCEQL
LASLILEEQESLPQMDKRKPSFVRFGKRTVAMMEKRKPSFVRFGRK
>FLP-9_Oesophagostomum_dentatum_OESDEN_10632
MNNRAQTRYTDPDALAEFCAQPQNREACEQLLAAALAEEQESLPQIDKRKPSFVRFGKRA
PALEKRKPSFVRFGRK
>FLP-9_Oscheius_tipulae_OTIPU.nOt.2.0.1.t13767
MPEDFEDPDMEDPTLPECSLYSFEQYVDKLMESYIKKLHIPRDSFPTLYTTMKTSQICLL
LCLVVCSFAQSLNSQLIESICALPENEQLCAELVSAAVKEVRRVAEAEDTEPMTMEKRKP
SFVRFGKRSDGQMEKRKPSFVRFGRK
>FLP-9_Ancylostoma_ceylanicum_predicted

YHHFQYTPTSTATLFQVDKRKPSFVRFGKRAALVEKRKPSFVRFGRK
>FLP-9_Angiostrongylus_costaricensis_predicted
FLHNNKHQERAKNKPYAFQLLPKFWHISKLSNQWIQFQVDKRKPSFVRFGKRADAMSKEKRKPSFVRF
GRK
>FLP-9_Diploscapter_pachys_predicted
ARFLRIVSLLNYWFEKEALINFQPEKRKPSFVRFGKRSAAPSRDEILLRQMADWN
>FLP-9_Haemonchus_placei_predicted
TDRGWNPVMSDWNFLFIYNVPRFQMDKRKPSFVRFGKRSEGDSAMEKRKPSFVRFGRK
>FLP-9_Nippostrongylus_brasiliensis_predicted
SLDRACVSSASYHSIRCTPEHMAVIKTQIIFVCFQMDKRKPSFVRFGKRSEDLMMEKRKPSFVRFGRK
>FLP-9_Teladorsagia_circumcincta_predicted
RISAYFATSESVSITTSPLCPSTLSVVPAFKLEFLTASNAAEFQMDKRKPSFVRFGKRSDGEMAMEKR
KPSFVRFGRK
>FLP-10_Caenorhabditis_angaria_Cang_2012_03_13_00005.g433.t1
MQFKCLLFLIFSLIFCLVTSVPINEYHRGGSLSEKRQGKQRSGYIRFGKRSHDPNAELLD
LDQLLF
>FLP-10_Caenorhabditis_brenneri_CBN03474.1
MQLAAVFLLVVLCLAAVFATPIKDASRARREVASEKRQTKSRSGYIRFGKRRVDPNAELL
YLDQLLL
>FLP-10_Caenorhabditis_briggsae_CBG17695.1
MQLSAVFVFLVLCLAAVFAVPLKDASRARREVAPSEKRQTKSRSGYIRFGKRRVDPNAEL
LYLDQLLL
>FLP-10_Caenorhabditis_elegans_T06C10.4.1
MQLSIVFVFFVLCLAAVFAVPISDASRARRQVASEKRQPKARSGYIRFGKRRVDPNAELL
YLDQLLI
>FLP-10_Caenorhabditis_inopinata_Sp34_40101800.t1
MQLSVVVVFLVLCLAAVFAVPVSDVSRARREAASEKRQTKARSGYIRFGKRRVDPNAELL
YLDQLLI
>FLP-10_Caenorhabditis_japonica_CJA05732.1
MQLSVVFLFFVLCLAAVFAVPIKEESRARRSVPGSEKRQTKARSGYIRFGKRRVDPNAEL
LYLDQLLL
>FLP-10_Caenorhabditis_latens_FL83_19196
VYRQSTLLETTMQLSVVFLLVVLCLAAVIAIPISDASRARRDAVSEKRQTKSRSGYIRFG
KRRVDPNAELLYLDQLLL
>FLP-10_Caenorhabditis_nigoni
MQLSAVFVFLVLCLAAVFAVPLKDASRARREVAPSEKRQTKSRSGYIRFGKRRVDPNAEL
LYLDQLLL
>FLP-10_Caenorhabditis_remanei_FL81_05193
MQLSVVFLLVVLCLAAVIAIPISDASRARRDAVSEKRQTKSRSGYIRFGKRRVDPNAELL
YLDQLLL
>FLP-10_Caenorhabditis_tropicalis_Csp11.Scaffold629.g7532.t1
MQLSVVFLLVILCFAAVFATPISDESRARRAVASEKRQTKSRSGYIRFGKRRVDPNAELL
YLDQLLL
>FLP-10_Diploscapter_coronatus_DCO_025220
MKCLSIFFVLLLAILLTNAMPEDRFRRSMAVANAKRQAASANGMRSGYIRFGKRKSNPND
DLLMLDQLIL
>FLP-10_Caenorhabditis_sinica_predicted
ESFFLSIPFPASRARREAAASEKRQTKSRSGYIRFGKRRVDPNAELLYLDQLLL
>FLP-10_Diploscapter_pachys_predicted
NIEILVFIISVLKLQYYSFADRFRRSMAVANAKRQAASANGMRSGYIRFGKRKSNPNDDLLMLDQLIL
>FLP-11_Plectus_sambesii_PSAMB.scaffold74size85960.g1659.t1
MHHSVLALTVGLACLALCFASVELDEDESAVVDKRAMRNALVRFGRGGSMRNAIVRLGKR
AMRNAIVRFGKRDDDSSPEGAAEEDKRSPPQPFVRFGRSGHSEHVNSILDTLRKLNEANE
Y

>FLP-11_Anisakis_simplex_ASIM_0001234201-mRNA-1
MFQWESSVCTMMQRSTLIGLLVCVAFAVVAYAEDDDQMADKRAMRNALVRFGRSGMRNAL
VRFGKRADDNEYSTWDEKRNGAPQPFVRFGRSGRVDHIHDILSTLQRLQLANE
>FLP-11_Ascaris_lumbricoides_ALUE_0000665901-mRNA-1
MVSSPTPLYSRSFSFAQMLRSTVVGLFACIAVAVVASAEDDEQVAEKRAMRNALVRFGRS
GMRNALVRFGKRADDNEYSTLDEKRNGAPQPFVRFGRSGRVDHIHDILSTLQRLQLANE
>FLP-11_Ascaris_suum_AgB05_g330_t01
MVSSPTPLYSRSFSFAQMLRSTVVGLFACIAVAVVASAEDDEQVAEKRAMRNALVRFGRS
GMRNALVRFGKRADDNEYSTLDEKRNGAPQPFVRFGRSGRVDHIHDILSTLQRLQLANE
>FLP-11_Brugia_malayi_Bm13445.1
MHFVHTYFLLFYCFIVQFVTITVASAHNDEALGEKRAIRNALVRFDRSGIRNALVRFGKR
MSDMYFLDAENRRATALNPSGKVSF
>FLP-11_Brugia_pahangi_BPAG_0000991601-mRNA-1
MHFVHTYFLLFYCFIVQFVTTTVASAHNDEALYAIFCFSLDLGEKRAIRNALVRFDRSGI
RNALVRFGKRMSDMYFLDAENRRATALNPSGKVSF
>FLP-11_Brugia_timori_BTMF_0001466401-mRNA-1
LVRFDRSGIRNALVRFGKRMSDMYFLDAENRRATALNPSGKVSF
>FLP-11_Dirofilaria_immitis_nDi.2.2.2.t01765
MANAQDDETLGEKRAMRNALIRFGRSGIRNALIRFGKRMSDMYFSDAENRRTVVLHPSVK
FDRFEHSKHINNILSKI
>FLP-11_Dracunculus_medinensis_DME_0000200901-mRNA-1
MRIRGDEWPWWLENKSSLVVGSVCSMQQITVIALFFSMVLAIFVNGDNVDDLLRDKRAMR
NALVRLGRSSMRNALIRFGKRSVDTDLNDQAYGYKRNGAPQPFVRFGRSQRLHDILSEIE
TAQSTEKR
>FLP-11_Enterobius_vermicularis_EVEC_0000451701-mRNA-1
MYRLAVLLAIVIVAAEAYPMDSDDYVYADKRAMRNALVRLGRSGMRNALVRSELLRALQE
HQVTVELNTERRINDLLFIYYNCFNRNISFKCFERIICLFDLFSCLNLSLRQTQTNSCKK
ASCY
>FLP-11_Onchocerca_ochengi_OOCN_0000197101-mRNA-1
MPLKTDSTKVKLSPHNGSRFCYLMSLSFSADPYSNSLRSPAASTLSPFADDAYGRFFDRK
PTAPRNALVRFG
>FLP-11_Onchocerca_volvulus_OVOC4352.1
MPEIVVQNRNLKKFQYKKLDCAISGKDFAFAVNEKRLHQIAAKASQIIVSMRPIHAYFLL
FYCLIAQFVIIASDQDDETLGQKRAMRNALIRFGRSGIRNALVRFGKRMSDMYFLDPENR
RAAALHPSRSLAALRVRNIAKIFFQKFNDSAVR
>FLP-11_Parascaris_univalens_PgB07_g029_t01
QMLRSTIAGLFACIAVAVVAGAEDDEQLAEKRAMRNALVRFGRSGMRNALVRFGKRADDN
EYSAMDEKRNGAPQPFVRFGRSGRVDHIHDILSTLQRLQLANE
>FLP-11_Syphacia_muris_SMUV_0000858901-mRNA-1
MYRSAVFLICIVIAAEAYPMDADDYFYTDKRAMRNALVRLGRSGMRNALVSKQFKHNRRL
LQLWNISAIFDTKFYIILRLIQKSLHFNCLAKSISTYHGNILMCLIKNVLKYLLYCYIQL
HKFSNIFEVFINDLLG
>FLP-11_Thelazia_callipaeda_TCLT_0000283401-mRNA-1
MKREEKRAVRGNLVRFDRSAMRNALVRFGKRMTNFDITLPENEQIIASHKSVHFDHLQHP
NYMKDFMPKTRRLQWNVDYLR
>FLP-11_Toxocara_canis_TCNE_0000833701-mRNA-1
MLRSTVVGLFVCVALAVFANAEDDEQVADKRAMRNALVRFGRSGMRNALVRFGKRADDNE
YSTWDEKRNGAPQPFVRFGRSGRVDHMHDILSTLQRLQLANE
>FLP-11_Wuchereria_bancrofti_maker-PairedContig_261-snap-gene-1.11-
mRNA-1
MHFVHTYFLLSYCFIVQFVTIAITSVRNDEALGEKRAIRNALVRFDRSGIRNALVRFGKR
TSDTYFLNAESRRPTALNPSVR
>FLP-11_Gongylonema_pulchrum_predicted
GSFFFFRADKRPMRNALVRFGRAGMRNALVR

>FLP-11_Loa_loa_predicted
KHCFRREKRAIRNALVRFGRSGIRNALVRSLSTKSLKQKKKKNNN
>FLP-11_Onchocerca_flexuosa_predicted
VLFSNKKKCFRGEKRTMRNALIRFGRSGIRNALVRLLSIKSLKKVDKKSD
>FLP-11_Parascaris_equorum_predicted
YRAEKRAMRNALVRFGRSGMRNALVR
>FLP-11_Ancylostoma_caninum_ANCCAN_00124
MPSPASTTLCLIAVLVVAALAQDDSSPEKRAMRNALVRFGRAGGGMRNALVRFGKRSSAD
DDYEAAMQDKRNGAPQPFVRFGRSGHLDHMHDILSTLQKLEMANYY
>FLP-11_Ancylostoma_ceylanicum_Acey_s0084.g1709.t1
MPSPASTTLCLIAVLVVAALAQDDSSPEKRAMRNALVRFGRAGGGMRNALVRFGKRSSAD
DDYEAAMQDKRNGAPQPFVRFGRSGHLDHMHDILSTLQKLEMANYY
>FLP-11_Ancylostoma_duodenale_ANCDUO_12265
MPSPASTTLCLIAVLVVAALAQDDSSPEKRAMRNALVRFGRAGGGMRNALVRFGKRSSAD
DDYEAAMQDKRNGAPQPFVRFGRSGHLDHMHDILSTLQKLEMANYY
>FLP-11_Angiostrongylus_cantonensis_ACAC_0000368601-mRNA-1
MLSPVSRTAYFMALLIAYTLAQDDYIPEKRAMRNALVRFGRAGGGMRNALIRFGKRSSVD
DDYDQRKRNGAPQPFVRFGRSSQLDHMHDILSTLQKLEMANYY
>FLP-11_Angiostrongylus_costaricensis_ACOC_0000049001-mRNA-1
MGGPSNKFGFCAAFEVLLLQTSLSLFSPEKRAMRNALVRFGRAGGGMRNALVRFGKRSSV
DDDYDQRKRNGAPQPFVRFGRSGQLDHMHDILSTLQKLEMANYY
>FLP-11_Caenorhabditis_angaria_Cang_2012_03_13_00035.g1991.t1
MMRAAFVLVIVCLAAVFAQYDDISNEKRAMRNALVRFGRASGGMRNAMVRFGKRSAALED
EEEYVSPLAGKRNGAPQPFVRFGRSGQIDHMHDILSTIQKLQFAGN
>FLP-11_Caenorhabditis_brenneri_CBN11115.1
MTQFSALALLLIIFVATTFAQYDDVSAEKRAMRNALVRFGRASGGMRNALVRFGKRSSLD
EEDYAPESPLQGKRNGAPQPFVRFGRSGQLDHMHDILSTLQKLQYAGNK
>FLP-11_Caenorhabditis_briggsae_CBG14520.1
MTKFSALVLILVIFVAASFAQPYDDVSAEKRAMRNALVRFGRASGGMRNALVRFGKRSAM
DEEDFSPESSLQGKRNGAPQPFVRFGRSGQIDHMHDILSTLQKLQYAGNK
>FLP-11_Caenorhabditis_elegans_K02G10.4a.1
MTQFSALALLLIVFVAASFAQSYDDVSAEKRAMRNALVRFGRASGGMRNALVRFGKRSPL
DEEDFAPESPLQGKRNGAPQPFVRFGRSGQLDHMHDLLSTLQKLKFANNK
>FLP-11_Caenorhabditis_inopinata_Sp34_X0251900.t1
MAQSSALAFLLIIFFAAISAQYDDVSAEKRAMRNALVRFGRASGGMRNALVRFGKRSSQD
EEDFAPESSLHGKRNGAPQPFVRFGRSGQLDHMHDILSTLQKLQYAVNK
>FLP-11_Caenorhabditis_japonica_CJA00813.1
MTQFSALALLLIVLVGVTLAQSYEDVSAEKRAMRNALVRFGRASGGMRNALVRFGKRSAL
DEDDFSPESYQGKRNGAPQPFVRFGRSGQIDHMHDILSTLQKLQYAGNN
>FLP-11_Caenorhabditis_latens_FL83_00627
MTKLSALALLLIVFVAATFAQSYEDVSAEKRAMRNALVRFGRASGGMRNALVRFGKRSSL
DEEDFAPESSLQGKRNGAPQPFVRFGRSGQLDHMHDILSTLQKLQYAGNK
>FLP-11_Caenorhabditis_nigoni_Cni-
MTKFSALALILVIFVAASFAQSYDDVSAEKRAMRNALVRFGRASGGMRNALVRFGKRSAM
DEEDFAPESSLQGKRNGAPQPFVRFGRSGQIDHMHDILSTLQKLQYAGNK
>FLP-11_Caenorhabditis_remanei_FL81_02271
MTKLSALALLLIVFVAATFAQSYEDVSAEKRAMRNALVRFGRASGGMRNALVRFGKRSSL
DEEDFAPESSLQGKRNGAPQPFVRFGRSGQLDHMHDILSTLQKLQYAGNK
>FLP-11_Caenorhabditis_sinica_Csp5_scaffold_00053.g2698.t3
MTQFSALALLLVIFVAATFAQSYDDVRSAPCATLWFDSEXXGRASGGMRNALVRFGKRSA
LDEEDFAPESPLQGKRNGAPQPFETPVRRQQVMTEDDRILLEQLLRRIHQL
>FLP-11_Dictyocaulus_viviparus_nDv.1.0.1.t00680
MCPAAIPSLFKGRNHLFSLIMFMLSPVSRTLFLVVLLVASTLAQDIYNSEKRAMRNALVR
FGRAGGGMRNALVRFGKRSGNDDYDTSSQSKRNGAPQPFVRFGRSGQVDHIYDILSTLQK

LETANYH
>FLP-11_Diploscapter_coronatus_DCO_030457
MYFLFISFNLVMLKQGSPSHFRYLQPAAVGRSPIYSLVRMQYPTLVCLLVGLVALFAAAY
GQDDDSYNMDKRAMRNALVRFGRATGMRNALVRFGKRSSPDVEEFGQTLQDKRNAGVPQP
FVRFGRSGQIDHLSDMISTLQRLESVGSK
>FLP-11_Diploscapter_pachys_WR25_02813E.1
MQYPTLVCLFVCLVALFAAAYGQDDDSYNMDKRAMRNALVRFGRATGMRNALVRFGKRSS
PDAEEFGQTLQDKRNAGVPQPFVRFGRSGQIDHLSDMISTLQRLESVGSK
>FLP-11_Haemonchus_contortus_HCON_00176100
MTSSTTIKFCIIAVVIASLRAQDSFALEKRAMRNALVRFGRAGGSMRNALVRFGKRYLAT
DDDYATAAAQGKRNGAPQPFVRFGRSGHLDHIHDILSTLQKLQLANYH
>FLP-11_Haemonchus_placei_HPLM_0001208401-mRNA-1
MTSSTTIKLCIVAVVIASLRAQDNFALEKRAMRNALVRFGRAGGSMRNALVRFGKRYLAT
DDDYATAAAQDKRNGAPQPFVRFGRSGHLDHIHDILSTLQKLQLANYH
>FLP-11_Mesorhabditis_belari_mbelari.g4147.t1
MAPPLAFPFSLFTSSSTSSDSFAPSSLLTAMHPLVSWFLLIVLLFSLSTQAQLEDDDDIS
FEKRAAPMRNALVRFGRAAPNGGLRNALVRFGKRSSVSTQDKRNAQPQPFVRFGRSGNHL
DHMHDILSTLQRLQYANN
>FLP-11_Micoletzkya_japonica_MicoRS5524-mkr-S25-2.8-mRNA-1
MRTSRISPEGTVAYGASLEKTSLSRMHTSLLLGFVLVAAIAYAQYDEDLEFDKRSMRNAL
VRFGRSSGGMRNALVRFGKRSDPDHEGNWQDKRNGMPQPFVRFGRSGQLDHMHDILNTLE
QIHQVHP
>FLP-11_Necator_americanus_NECAME_00060
MSSSASTSLCIIAVLIVVALAQDDSSPDKRAMRNALVRFGRAGGGMRNALVRFGKRSSAD
DDYEAALQDKRNGAPQPFVRFGRSGHLDHMHDILSTLQKLEMANYY
>FLP-11_Nippostrongylus_brasiliensis_NBR_0000793401-mRNA-1
MPSSTSITLCFVVVLVAVVLAQDDSTAEKRAMRNALVRFGRAGGGMRNALVRFGKRSSTT
LDDYEAAAQDKRNGAPQPFGEVRFGRSGQLDHMHDILSTLQKLQMANYY
>FLP-11_Oesophagostomum_dentatum_OESDEN_00051
MPSSATLCLVAVLVVVALAQDDSSVEKRAMRNALVRFGRAGGGMRNALVRFGKRSSADDD
YEAAMQDKRNGAPQPFVRFGRSGQLDHMHDILSTLQKLEMANYY
>FLP-11_Oscheius_tipulae_OTIPU.nOt.2.0.1.t00592
MASTLSVALFALCIVAIMAQYDDIETEKRAMRNALVRFGRASGGARNALVRFGKRSSGDL
YDQYESSMQDKRNGAPQPFVRFGRSGRLEHMHDILSTLQKLEMANSK
>FLP-11_Pristionchus_entomophagus_entomophagus-mkr-S267-2.7-mRNA-1
MNSSLVFALVACVAIVAAQQYDDEEIGFEKRAMRNALVRFGRSSGGMRNALVRFGKRSDP
EMESAQYQDKRNGAPQPFVRFGRSGQTDHMHDILSTLARVQSASSSLFDGKDDSGGTCSQ
DKKTCGGEEKKAPTKKKTAPLDISTMWKKPDPAIALAFKGHYYDEEKYEGKVAQTVLKKL
EEKYRVLEERCSQLSGVCYEVVDRVWRNPEGFETNKWEMLRQVQLKGQIDSHFGRAVLVK
PEVLNWKTFDTRKWEVNKYVVIDREIQSLIGALTGSKAIPREEKNATRDYQILLVGLGTP
VIPNFIEKYHWMDLIVVEREPLFEYLAKKWFNMRKHPNLRILNADPVSFMSFVVAERHEV
DAVLINECKDHVDARPCPANVYLERGTMDVIKKVVRQETGVLGVNSRSSVKTHERSVANA
YEGAFPMCFDYPGDHKQLHFERYKATICFNRKHTPFWLQSSRDFTLAALHDFFFYI
>FLP-11_Pristionchus_exspectatus_exspectatus-ag_msk-S_231-2.21-mRNA-1
MNSSLLFTLLIGVAVVVAQQYDDEEIGFEKRAMRNALVRFGRSSGGMRNALVRFGKRSDP
EMESANYQDKRNGAPQPFVRFGRSGQTDHMHDILSTLARVQSASSVFDGQKKDAGTCSGT
KKTCGDAVASKAASKKAKAPLDVSVMWKKPEPPTKGHYYDEEKYDGRVAQAVLKKLEEKK
YRVLEEKCSQLTGVCYEVVDRVWRNPEGFEQNKWEMIRQIQLKGQVDYHFGRSLLIKPTE
LNWKTYDTKKWVINKYAVLDKEIQSLIGGITGSKAIPRNETDANGKEYQILLVGLGAPII
PNFMEQYHWAKKDPIWNYFAGKWFNMRKHPNLRVLHADPISFMAFIVAQRHEIDAVIINT
CKDYVDPRPCPADVYLERGTIEVIKNVVREHSGMAGVNARSMEGKHPIEVANAYQDSFPM
CFDYPADHKQLTFERYKAAICYNREHPPFWHQSEKQYSTDSFHEFFFYI
>FLP-11_Pristionchus_fissidentatus_fissidentatus-ag_msk-S246-5.22-

mRNA-1
MNSSIVIALLFGAVLVAAQYDEDEIGFEKRAMRNALVRFGRSTGGMRNALVRFGKRSDPE
MESANYQDKRNGAPQPFVRFGRSGNTDHMHDILSTLETLNRVQISSSFLDGKKVDDVNGG
TCSTEKKQCGGGEKKAKKKEPFDISVMWSEKPESNEHHSFDKEGQYDGRVAQAVIKKLGE
KPRVLDTKCSEITGICYEVVDRIWYDNEGFGNNPWEMLRQVQVKGEPEAHFGRATLVKPK
ELNWKTYDTKNWAINDGAVIDRETQALIGGLTGSLAIPRKAVNATRNYYILLVGLGTPTI
TNFLKRYHWMEVLVLDNEPVFEYFARKWFGFRLGPNLNYLQADPISFMSWVVGEKVQYDA
VIINECKDHADPHPCPASVYLERATMDVVRKVVRSKHGVFGVNTRSTEKKYEKEVANAYE
SVFPMCFDYPGEHKKLQFERYKSTICFNRPHRPFWQLSEKQYAYDAHHNFFYYV
>FLP-11_Pristionchus_japonicus_japonicus-mkr-S19-5.0-mRNA-1
MNSSILFALLIGAAVVAAQQYDDEEIGFEKRAMRNALVRFGRSSGGMRNALVRFGKRSDP
EMESANYQDKRNGAPQPFVRFGRSGQTDHMHDILSTLARIQSSLSPSSLLNSKKKDTGTC
SGTKKTCGEKAPAPKKEKVVVDPSVMWKKPEPPTRGHYYDEEKYDGKLAQAVLKKLEEKK
YRVLEEKCSQLTGVCYEVVDRVWRNPEGFEQNKWEMIRQIQLKGEVDYHFGRSLLIKPDE
LTWKTYDTKTWKVNKWAVLDKEIQSLIGGITGSKAIPRNETDANGKEYQILLVGLGAPII
PNFMEQYHWMDIIVVEKDPIFNYFAGKWFNMRKHPNLRVLHADPISFMAFIVAQRHEIDA
VIINTCKDYVDPRPCPADVYLERGTIEVIKNVVREHSGMAGVNARSMIRPVDTHPIEVAN
AYQSSFPMCFDYPADHKQLTFERYKSAICYNREHIPFWHQSEKQYATDSYHEFFFYI
>FLP-11_Pristionchus_maxplancki_maxplancki-mkr-S40-2.45-mRNA-1
SSSGLVPSLSLSLLPIMNSSLLFALLIGVAVVAAQQYDDEEIGFEKRAMRNALVRFGRSS
GGMRNALVRFGKRSDPEMESANYQDKRNGAPQPFVRFGRSGQTDHMHDILSTLARVQSAS
SLFDGQKKEAGTCSGTKKTCGGEAAPIKAKKAPLDVSVMWKKPEPPTKGHYYDEDKYDGK
MAQAVLKKLEEKKYRVLEEKCSQLTGVCYEVVDRVWRNPEGFEWVSWQNKWEMIRQIQLK
GQVDYHFGRSLLVKPTELNWKTYDTKKWVVNKYAVLDKEIQSLIGGITGSKAIPRNETEA
NGREYQILLVGLGAPIIPNFMEQYHWMDMIVVEKDPIWNYLGGKWFNMRKHPNLRVLHAD
PISFMAFIVQQRHEIDAVIINTCKDYVDPRPCPADVYLERGTIDVIKNVVKEHTGMAGVN
ARAMESKHPIEVANAYQDSFPMCFDYPADHKQLTFERYKAAICYNREHPPFWHQSEKQYS
TDSFHEFFFYI
>FLP-11_Pristionchus_mayeri_mayeri-mkr-S427-0.109-mRNA-1
MHSSILFALLFGAVVVASQQFDDDEIGFEKRAMRNALVRFGRSSGGMRNALVRFGKRSDP
EMESANYQDKRNGAPQPFVRFGRSGQTDHMHDILSTLERVQLHLSDAHGERDEYGELQAK
ASPKKPRKEPIDISIMWKKPDPQDMTRYFDADGQWDGKTAQAVLKKLAAKPRVLEEKCSQ
LTGVCYEVVDRVRMNPDGFESNKWEMLRQVQLKGHVDSHFGRATLVKPRELNWKTYDTKK
WEVNKKAVIDKEIQTLIGGLTGSKAIPRNESEAAGREYQILLVGLGTPIIPIFIEQYHWM
DLIVVEKDPIFSYFAKKWFGMREHPNLRILHADSISFMQFAVAQRHEFDAVLINVCKDYV
DERPCPADVYLERSTMEVIRNCVKEETGMAGVNTRSNVKKHEKEVANAYDNVFPMCFDYP
GDHKQLHVERYKSTVCFNREHTPFWLGYERDYTLATYHDWFFYI
>FLP-11_Pristionchus_pacificus_PPA07538.1
AWNRGNEHPNGILSIVCAPDGPLQSSLPTTQSLSLFSPFSIMNSSLLFTLLIGVAIVVSQ
QYDDEEIGFEKRAMRNALVRFGRSSGGMRNALVRFGKRSDPEMESANYQDKRNGAPQPFV
RFGRSGQTDHMHDILSTLARVQSASSVFDGQKKDAGTCSGTKKTCGDAVASKAASKKAKA
PLDVSVMWKKPEPPTKGHYYDEEKYDGRVAQAVLKKLEEKKYRVLEEKCSQLTGVCYEVA
DRVWRNPEGFEQNKWEMIRQIQLKGQVDYHFGRSLLIKPTELNWKTYDTKKWVINKYAVL
DKEIQSLIGGITGSKAIPRNETDANGKEYQILLVGLGAPIIPNFMEQYHWMDMIVVEKDP
IWNYFAGKWFNMRKHPNLRVLHADPISFMAFIVAQRHEIDAVIINTCKDYVDPRPCPADV
YLERGTIEVIKNVVKEHSGMAGVNARSMESKHPIEVANAYQDSFPMCFDYPADHKQLTFE
RYKAAICYNREHPPFWHQSEKQYSTDSFHEFFFYI
>FLP-11_Heterorhabditis_bacteriophora_predicted
FSAEKRAMRNALVRFGRAGGGMRNALVR
TSGDYASNISTYISRFGKRSSYDDDYEAASQDKRNGAPQPFGKIKWEVLIKHA
>FLP-11_Pristionchus_arcanus_predicted
DELKIEELPIFCSFEKRAMRNALVRFGRSSGGMRNALVR
>FLP-11_Bursaphelenchus_xylophilus_BXY_0627300.1
MFSLPFPVYDYSSAITVSGYPSSANPPRLSNNFIEGSLSVPNNKMKLSALILVLALIVCA

VKAQAEFDEENGYAKRSMRNALVRFGRSGMRNALVRFGKRSLGNDYAEAKRAQSAPEPFV
RFGRSAPSHFALYDYYDI
>FLP-11_Halicephalobus_mephisto_HMEPH_15589-RA.p1
FQKPDTGAFAETYVFKMQSTVILCGVALVLCVVSSYAQQSDFDDFTFEKRTPMRNALVRF
GRAGMRNALVRFGKRSAISEEYPVIPEYALEAKRNGAPQPFVRFGRSAGRIDHMHDILNT
LQKLELANSQ
>FLP-11_Panagrellus_redivivus_Pan_g7468.t1
MQASSVLLFGLALAFCVIVSNAQFDEDYAAPTEKRAMRNALVRFGRAAGMRNALVRFGKR
SADEIAVMPDYGNEAKRNGAPQPFVRFGRSAGRIDHMHDILSTLQKIEMANGQ
>FLP-11_Parastrongyloides_trichosuri_PTRK_0000740600.1
MQSYFLLSIILSIFVVALIANGQQYDDVSVGYSPEKKAMRNALVRFGRAGMRNALVRFGK
RSSDNEVQEFALKRNAAPQPFVRFGRSANQGFGEEAYYVPYNVIYANTEA
>FLP-11_Rhabditophanes_kr3021_RSKR_0000528400.1
MQTSFIVAILLSICVASLYVQAHQLDEDSFGFNPDKRAMRNALVRFGRAGMRNALVRFGK
RASDIPIFLTSDYGYKRNGAPQPFVRFGRSGRDGVEDQDLTPQSPSKSSQNSLKEDI
>FLP-11_Steinernema_carpocapsae_SC.X.g4084
MNAYTLFGCLMIIAIVTLVKADVDEESSSVEKRAMRNALVRFGRAGMRNALVRFGKRNQV
DDSSFGANEDKRNGAPQPFVRFGRSGRVDHINDILSTLQKLEMANA
>FLP-11_Steinernema_feltiae_L889_g7374.t1
MNAYTLFGCLIVIAFVSLVKADIDEDSSSVEKRAMRNALVRFGRAGMRNALVRFGKRSQD
ETSFGASEDKRNGAPQPFVRFGRSGRVDHINDILSTLQKLEMANA
>FLP-11_Steinernema_glaseri_L893_g32192.t1
MVKADIDDDSSSVEKRAMRNALVRFGRAGMRNALVRFGKRGQVDDQSFTTTEDKRNGAPQ
PFVRFGRSGRVDHINDILSTLQKLEMANA
>FLP-11_Steinernema_monticolum_L898_g3040.t1
MNAYTLFACLIVVAIVSLANADIDEESSGVEKRAMRNALVRFGRAGMRNALVRFGKRSQD
DSSYGANEDKRNGAPQPFVRFGRSGRVDHINDILSTLQKLEMANA
>FLP-11_Steinernema_scapterisci_L892_g28903.t1
MNMYTLFGCLVIITIVSLANADIDEDSSSMEKRAMRNALVRFGRAGMRNALVRFGKRSQV
DESSFGANEDKRNGAPQPFVRFGRSGRVDHINDILSTLQKLEMANA
>FLP-11_Strongyloides_papillosus_SPAL_0001281500.1
MQSYIVLTILLTVFAAIFIANAQQYDESSIGFSPEKRAMRNALVRFGRAGMRNALVRFGK
RSMDNEMQEFALKRNAAPQPFVRFGRSSNLPIEGYYVPYNSLYGNAEA
>FLP-11_Strongyloides_ratti_SRAE_X000206000
MKSYLILSILLSVFAAVLIANTQEYDENSIGYVPEKRAMRNALVRFGRAGMRNALVRFGK
RSLDNDIQEFALKRNAAPQPFVRFGRSSSLSPDGYFIPYGNIYENNEA
>FLP-11_Strongyloides_stercoralis_SSTP_0000849700.1
MKSSIVLSLLFTVFAAFFIANAQQYDENSLGYLTEKRAMRNALVRFGRAGMRNALVRFGK
RNIDNDIPEFALKRNAAPQPFVRFGRSSNLSPSGYFIPLNNMYDNTEA
>FLP-11_Strongylus_vulgaris_SVUK_0000956801-mRNA-1
MKFCHFFSFYIFSAFSVEKRAMRNALVRFGRAGGMRNALVRFGKRSASDDDYEAAMQDKR
NGAPQPFVRFGRSGQLDHMHDILSTLQKLEMANYY
>FLP-11_Teladorsagia_circumcincta_TELCIR_00302
MYANWEAICSTHLAMILRSAEWFSIVFLLLVTFDAVFSHSRHKRRRHERDGDMRIEHILA
QMDSYEPTVTQQFRPPKYIANNLVMMTLEGRALFKDNSRIAADILIVSEDFTSTESPIIP
LSQDELRARYNTSNLEKRAMRNALVRFGRAGGSMRNALVRFGKRSSTADDDYAAAVAQDK
RNGAPQPFVRFGRSGHLDHIHDILSTLQKLQMANYY
>FLP-11_Acrobeloides_nanus_ACRNAN_scaffold4517.g9432.t1
MRSSILISLIVVLSSVIFTNSQQIDEDVSSGYEKRAAMRNALVRFGRAGGMRNALVRFGK
RSSALEEAAVPDYSEMKRNGAPQPFVRFGRSGGHFDHMHDILSTLQKLEMANS
>FLP-11_Ditylenchus_destructor_Dd_01492
MRLIRRLALLLVMFSLVGLCSGQDSTEDDSEQDYDISLEPLEPYLSAEDDKPDIICEKGG
YCNCIYGDAREGKKEPNVVNCTSASLRGSFLDTAHVVIRAEKFPNFKAEEVRMSHNRIHV
LRKGLIVSGHEDSIISLDFTDNLISDVERYCFDNFPKLVKLKLTNNQLRYEEISDDWLTA

KLGQSLHQLYLDNNRIKNLDDGIFDNLKNLNKLVLDGNKGLRLTPKTFGRGLKRLRILSM
DNCGFDTLDPEVFANLGGLTQLSLSRNRFNSVPLALRNVPRLNLLALSEAFMTHIPQGSF
NNMPALKSIFMRRMPNLTEVMSCAFCGLIDLESVDFSGSKGLGKIDDNAFGLYAKKEVVP
DDFNMLDLRDCNFTGISENLLDWGPPTRLALAGNPLHCDCETMKWLLGNRDVNMVGPVAV
CKSPAALREKPLRKANAKQCGLSAPASSSLFDSNMVLIFLVAVCGIALVVGAGMFAKHKG
VTVSQFLPAGRTQSSQLGYSNLGPSRSGAQNDIDDGDENRLEDDFNRPEFYFRVIHIFKR
NICPIFQLEQIAAVMNPSTILFLLLAALASVIFSSQAQRSPQFDEDFSSMEKRQMRNALV
RFGRSTGMRNALVRFGKRAPIESLADEIKRNGAPQPFVRFGRSTGGRSIDSWQDPSAVQT
VDMPYY
>FLP-11_Ditylenchus_dipsaci_jg25338
MNSSAMLLWLTALVVAVVVCVEAQQPQQFDEELNSSMQKRQMRNALVRFGRSSGMRNALV
RFGKRAQTINDLDDEIKRNGAPQPFVRFGRSSGRPTDPAQDLLSSLQQTEPLY
>FLP-11_Globodera_pallida_GPLIN_000909800
MIISTNAASITDVEDGQQQQQQQQQQPFALFPVASMPAEEFGGDGGGFIGSLPQMAKRRQ
QLLNALLMTRRLGRSPRASVRLPRSTMRNALVRFGKRALLPMMALDDKRNPPQPFVRFGR
SAAANAVENGATDPMFFAVS
>FLP-11_Globodera_rostochiensis_GROS_g04118.t1
MAKLCRFSRFFYALLLLLTLCSILMAEAAIWRMRTDKKAMRNALVRFGKRNAYRSSGEAF
VGAAGFGDSGAHLLRDIGMDDRQTQWAAFGDGGAPRPIKRLLLWPEQ
>FLP-11_Heterodera_glycines_Hetgly.G000027538
MTLAFGISNEIYYRENKRKRDTYYEMASSVSSSSSLAHHHRSHFGILPLIVSPLFLCLSI
FGTDAIPIDGVDDAAFQPFQQRQPFALFAVAQEADENGGIGSASLMAKRHLFEALARQGR
SPRSASSATMRNALVRFGKRALFPSVALDDKRNPPQPFVRFGRSAAEMVDQQQQQPYFPA
L
>FLP-11_Meloidogyne_arenaria_M.Arenaria_Scaff2084g029993
MFNLLILLIFIIIPFNQSILNTNFPSENTINEDEKEVGSFPTLLQLADREAFKSPFERHL
NYLPSSSSTAQKRTMRNALIRFGRSPRSATLRSAMVRFGKRSENSSNNELLESKRNSAPQ
PFVRFGRSFSSIPFNEEFNSPWWIERFAFL
>FLP-11_Meloidogyne_enterolobii_scaffold14079_cov159.g16845
MFNLLILSFFLIIPFNQSILNNFPSENTINEDEKEVGSFPTLLQLADREAFKSPFERHLN
YLPSSSSTAQKRTMRNALIRFGRSPRSATLRSAMVRFGKRSENSSNNELLESKRNSAPQP
FVRFGRSFSSIPFNGQFNSPWWIERFAFL
>FLP-11_Meloidogyne_floridensis_maker-nMf.1.1.scaf18023-augustus-gene-0.4-mRNA-1
LILSIFLIIPFNQSILNNFPSENTINEDEKEVGSFPTLLQLADREAFKSPFERHLNYLPS
SSSTAQKRTMRNALIRFGRSPRSATLRSAMVRFGKRSENSSNNELLESKRNSAPQPFGCF
WLI
>FLP-11_Meloidogyne_graminicola_NXFT01003304.1.8611_g
MNINQLLIFIIPFCQLISSNQFLKENTIKNIEEREEIGSFPTLLQLADREAFKPSFGSDL
FISSSIPSAQKRTMRNALIRFGRSPRSATLRSAMVRFGKRSDNLLNNSILLENKRTSAPQ
PFVRFGRSFPSIPNSYKEQFNTPWWTERFAFL
>FLP-11_Meloidogyne_hapla_MhA1_Contig1393.frz3.gene1
MFNKLILLISLILPFNQSIPSENKVVNDEKEIGSFPTLLQLADREAFKSPFENRHLNYFP
SDSIDSSSSAQKRTMRNALIRFGRSPRSATLRSAMVRFGKRSENSLNNSIMEIKRTSAPQ
PFGKYLFSFRLI
>FLP-11_Meloidogyne_incognita_Minc3s03062g32619
MFNLLILLIFIIIPFNQSILNNFPSENTINEDEKEVGSFPTLLQLADREAFKSPFERHLN
YLPSSSSTAQKRTMRNALIRFGRSPRSATLRSAMVRFGKRSENSSNNELLESKRNSAPQP
FVRFGRSFSSIPFNNEQFNSPWWIEGFAFL
>FLP-11_Meloidogyne_javanica_M.Javanica_Scaff1174g013469
MSILNTNFPSENTINEDEKEVGSFPTLLQLADREAFKSPFERHLNYLPSSSSTAQKRTMR
NALIRFGRSPRSATLRSAMVRFGKRSENSSNNELLESKRNSAPQPFVRFGRSFSSIPFNE
EFNSPWWIERFAFL
>FLP-12_Plectus_sambesii_PSAMB.scaffold1604size39546.g14050.t1

MTTKRDRMRPAFTHLPTRRSPSPVGSLPLAPLFTPHSFHLLPKFVAMHKSIVVVFAVCLI
AAHVDCREEESPLAPKRFDYAFQKRADEKTVQVTEADVINEVQRNLAKSMAYLDEVQNGA
KYNEKRRNKFEFIRFGK
>FLP-12_Anisakis_simplex FLP-16 ASIM_0001212301-mRNA-1
MILLVRKNKGTPELIDPSAYASDGDVLARVQRQLLGAMAMLDALQDGVTKLPEKRRNKFE
FIRFGRR
>FLP-12_Brugia_malayi_Bm17975.1
MTTFISFIIFVSAVFSQKIQGKRTHELLEPIAYNRGSDRELVAKAEEQLLNTLSLLEALV
DDSNQLQMEKRRNKFEFIRFGRR
>FLP-12_Enterobius_vermicularis_EVEC_0001246301-mRNA-1
MEWFVPLIFVCILFQTSDCSRLKKNAASQNYIVETRRARLRRELLEALDELGSLDEGELS
GDLRLWQGKPFVKRRNKFEFIRFGR
>FLP-12_Onchocerca_flexuosa_X798_02278
MLLVMVLAPNVLPNHQRLIITLCHTQFRASKLKYITVTSKLLFKKSDHCANPSECKLNGI
LRKLCDSNQVEMEKRRNKFEFIRFGRR
>FLP-12_Onchocerca_ochengi_nOo.2.0.1.t10732-RA
XSNQFEMEKRRNKFEFIRFGRR
>FLP-12_Syphacia_muris_SMUV_0000616701-mRNA-1
MGWFVLLAFMCILFQDSGCTRLKRDLWGQDYDVPELARSRLRRQLYNPLDDLEGLGGQGK
ALEKRRKFEFIRFGR
>FLP-12_Wuchereria_bancrofti_maker-PairedContig_3974-snap-gene-1.14-
mRNA-1
MMGERGMRREDHLKANISVVPFHQINCDIQEAMTCVPSVSNAFIQMLLVMVLVSHVLPLP
LAAHLPVYDSLYDEPPLVDYNFVPYSGLRNNGDWMNEPFRADKRKNEFIRFGKRDDFMNG
IIRFVCRFNKRKNEFIRFEGKRTHELLEPITYNRGNDRELVAKAEEQLLNTLSLLEALAD
DSNQLQMEKRRNKFEFIRFGRR
>FLP-12_Acanthocheilonema_viteae_predicted
LKHFADSNQLQMEKRRNKFEFIRFGRR
>FLP-12_Ascaris_lumbricoides_predicted
MRICNQDQSLLGTVKLLEKRRNKFEFIRFGRR
>FLP-12_Ascaris_suum_predicted
MRICNQDQSLLGTVKLLEKRRNKFEFIRFGRR
>FLP-12_Brugia_pahangi_predicted
LKHFVDSNQLQVEKRRNKFEFIRFGRR
>FLP-12_Brugia_timori_predicted
LKHFVDSNQLQMEKRRNKFEFIRFGRR
>FLP-12_Dirofilaria_immitis_predicted
NFIDSNQLEMEKRRNKFEFIRFGRR
>FLP-12_Dracunculus_medinensis_predicted
INWIIIDYLLLNSAKISDSKRIEKRRNKFEFIRFGRR
>FLP-12_Elaeophora_elaphi_predicted
VYHYFIKNYMQIDNHSICFLSSDSKLKRFVDSSQLQMDKRRNKFEFIRFGRR
>FLP-12_Gongylonema_pulchrum_predicted
KCTCHNLSLFSGTDQMQMEKRRNKFEFIRFGR
>FLP-12_Litomosoides_sigmodontis_predicted
LFNPISFAYFVDLNQFRMEKKRNKFEFIRFGRR
>FLP-12_Loa_loa_predicted
LYIILMIKVNYLNLCDFYYHFFLSKISIQIDYYNHFILKLFLIKTSLSFLFFSFLFFFFLDSNQLQMD
KRRNKFEFIRFGRR
>FLP-12_Onchocerca_volvulus_predicted
YFLSIYKINFVDSNQFEMEKRRNKFEFIRFGRR
>FLP-12_Parascaris_univalens_predicted
MRICYWDQSLPGTVKLLEKRRNKFEFIRFGRR
>FLP-12_Thelazia_callipaeda_predicted

QVLINSFLDANPIQMEKRRNKFEFIRFGRR
>FLP-12_Toxocara_canis_predicted
SYASMRPMVRRLRFAPLSGPAKLPEKRRNKFEFIRFGRR
>FLP-12_Ancylostoma_caninum_ANCCAN_23342
MRTSTTVLFLVSLLLVSAQKSHSKGAPELIPPMIYENDNSEMLAKVSAQLMNALATIENM
QEGTPIKIAEKRRNKFEFIRFGRK
>FLP-12_Ancylostoma_ceylanicum_Acey_s0354.g3306.t1
MISDSKGAPELITPMIYENDNSEMLAKVSAQLMNALATIENMQESESDGWFHGVPLAFPN
STRKLLGTPIKIAEKRRNKFEFIRFGRK
>FLP-12_Angiostrongylus_costaricensis_ACOC_0000232601-mRNA-1
MEPLMYEGDNSEMLAKISSQLLNTLATIENMQEGRNDCLKIAEKRRNKFEFIRFGRK
>FLP-12_Caenorhabditis_angaria_Cang_2012_03_13_00902.g14850.t1
MNFLFAITLLFVCVIASFAQKSKGSPEYIQQQPAYDSDNNHEALNKVSAQLLTVLAEIEN
MQEGNQQLKMAAEKRRNKFEFIRFGRK
>FLP-12_Caenorhabditis_brenneri_CBN13994.1
MNVRFAIALLLCIIATCVAQKAKGSPEVLPAAMYDGDASHESLNKISSQLLNALAELEAL
QEGSQQLKMAEKRRNKFEFIRFGRK
>FLP-12_Caenorhabditis_briggsae_CBG14501.1
MNSQLVLALLLCFIATSVAQKAKGSPEVLPAAMYDGDASHESLNKISTQLLNALAELEAL
QEGSQQLKMAEKRRNKFEFIRFGRK
>FLP-12_Caenorhabditis_elegans_C05E11.8a.1
MNVQVIIALLFCLIATCATQKVKGSPEVLPAAMYDGELSHESVNKISAQLLNALSELEAL
QEGNQQLKMAEKRRNKFEFIRFGRK
>FLP-12_Caenorhabditis_inopinata_Sp34_X0253300.t1
MNVQLALALFVCLIATSATQKAKGSPELLPAAIYDGDYSHESLNKISSQLLNALAELESL
QEGSQQLKMAEKRRNKFEFIRFGRK
>FLP-12_Caenorhabditis_japonica_CJA35969.1
MNVLFYITLLLLCLIATSAAQKSKGSPEVIPAAMYDGDVSHESLNKISNQLLNALAELEAL
QEGSQQLKMAEKRRNKFEFIRFGRK
>FLP-12_Caenorhabditis_latens_FL83_00641
MNVQFAIALLLCLIATSVAQKAKGSPEVLPAAMYDGDVSHESLNKISSQLLNALAELEAL
QEGSQQLKMAEKRRNKFEFIRFGRK
>FLP-12_Caenorhabditis_nigoni_Cni-
MNSQLVLALLLCLIATSVAQKAKGSPEVLPAAMYDGDASHESLNKISNQLLNALAELEAL
QEGSQQLKMAEKRRNKFEFIRFGRK
>FLP-12_Caenorhabditis_tropicalis_Csp11.Scaffold622.g6272.t1
MNVQFAIALLFCLIATSVAQKAKGSPEVLPAAMYDGDVSHESLNKISTQLLNALAELEAL
QEGSQQLKMAEKRRNKFEFIRFGRK
>FLP-12_Dictyocaulus_viviparus_nDv.1.0.1.t13650
MHTATVVILLLNSLFIGAQKLHDKGTPELMEPLMYESENNEILAKISSQLLNTLATIEDM
QRGIPLKIAEKRRNKFEFIRFGRK
>FLP-12_Diploscapter_coronatus_DCO_024998
MNANVFGLLLVLLLALITDAQKQQHNVDVTQYDDNQASLSKVYDQLLGALAMVDQMQEGN
KITAEKRRNKFEFIRFGRK
>FLP-12_Haemonchus_contortus_HCON_00164300
MRTATVALLLVSALLISAQKSHAKGTPELMQPVVFETDNSDLLAKVSSQLLNALANIESM
QEGTPLKIAEKRRNKFEFIRFGRK
>FLP-12_Haemonchus_placei_HPLM_0000895101-mRNA-1
MIVCFYAKGTPELMQPVVFETDNSDLLAKVSSQLLNALANIESMQEGTPLKIAEKRRNKF
EFIRFGRK
>FLP-12_Heligmosomoides_polygyrus_HPOL_0000334001-mRNA-1
MRTATIVLLLLSSLLVSAQKSHMKGTPELMQPILYESDNSDLLAKVSNQLMSALASIESM
QEGTPIKIAEKRRNKFEFIRFGRK
>FLP-12_Micoletzkya_japonica_MicoRS5524-mkr-S29-1.4-mRNA-1

MIRAGLVVLVCLALATSAQKHQSKDGLMMSEQETLSRVQSQLLGAMALLESLEEPQPDLG
ADGSPLKAEKRRNKFEFIRFGRK
>FLP-12_Necator_americanus_NECAME_09855
FHDLIACRSLPRLLSSLHHITYITLSKTTETKEESNMRTSTTVIFAMSMLLVTAQKSHSK
GAPELIQPVMYDPDNSETLAKVSSQLLTALATIESIQEGGVPPIKIAEKRRNKFEFIRFG
RK
>FLP-12_Nippostrongylus_brasiliensis_NBR_0001490001-mRNA-1
MRSATVVLLLLSVIFVNAQKSHTRQVENVYWLHATNQHSTYLCSFEEVLQQINKLARIDL
KGASELMQPLVYEADNSDLLAKVSSQLLSALANIESLQEGTPVKVAEKRRNKFEFIRFGR
K
>FLP-12_Oesophagostomum_dentatum_OESDEN_00223
MWTSAITLFMAATLLVSAQKPHSKGTPELIQPMIYESDNSEMLAKVSAQLMNALATIENM
QEGTPIKIAEKRRNKFEFIRFGRK
>FLP-12_Parapristionchus_giblindavisi_Parapristionchus-mkr-
S_343-0.3-mRNA-1
MIRPALIALLLCTLTLGEKTQPKSEGVALSESETLARVQGQLLGAMALIESLESPQEYPG
GEPLTAEKRRNKFEFIRFGRK
>FLP-12_Pristionchus_arcanus_arcanus-mkr-S_41-2.44-mRNA-1
MMRHFAILAALCLVFAMGNKNVANQPEGVVVSDQEILARAQSQLLGAMALLETLDNKQEL
TEGEPITAEKRRNKFEFIRFGRK
>FLP-12_Pristionchus_entomophagus_entomophagus-mkr-S533-0.48-mRNA-1
MMRHIALLAAICAILAMGNKNVGKSEGVVVTDQEILARAQSELMGAMALLETLDNKQEFA
GAEPITAEKRRNKFEFIRFGRK
>FLP-12_Pristionchus_exspectatus_exspectatus-mkr-S_5-2.21-mRNA-1
MMRHFAILAALCLVFAMANKNVAAQPEGVVVSDQEILARAQAQLLGAMALLETLDNKQEL
TEGEPITAEKRRNKFEFIRFGRK
>FLP-12_Pristionchus_fissidentatus_fissidentatus-mkr-S51-1.23-mRNA-1
MMRQLVLLSVCFAALSLATKTGGKPEGVIVSDQEILARAQSQLLGAMALLEGLENKQEFV
GEAVDPITAEKRRNKFEFIRFGRK
>FLP-12_Pristionchus_japonicus_japonicus-mkr-S261-1.63-mRNA-1
MMRYIALFAALCAIITMANKNVVKQPEGVMVSDQEILARAQAQLFGAMALLETLDNKQEL
AEGEPITAEKRRNKFEFIRFGRK
>FLP-12_Pristionchus_maxplancki_maxplancki-mkr-S8-7.60-mRNA-1
MMRHFALFVALCAVFAMGNKNVAKQPEGIVVSDQEILARAQAQLLGAMALLETLDNKQEL
AEGEPITAEKRRNKFEFIRFGRK
>FLP-12_Pristionchus_mayeri_mayeri-mkr-S636-0.11-mRNA-1
MMRQLALLAALCALLVTANKNVPKPEGIIISDPEILARAQQQLLGAMALLETLENKQEIS
GADSADPITAEKRRNKFEFIRFGRK
>FLP-12_Pristionchus_pacificus_PPA20651.1
MMRHFAILAALCLVFAMANKNVATQPEGVVVSDQEILARAQAQLLGAMALLETLDNKQEL
TEGEPITAEKRRNKFEFIRFGRK
>FLP-12_Ancylostoma_duodenale_predicted
TLSLPLKTCRKVSLMVGFLMFHWHFPTSTRKLAGTPIKIAEKRRNKFEFIRFGRK
>FLP-12_Caenorhabditis_sinica_predicted
DVXXTLIRITTIRKTKTNISGRDPLRFSNPASYGPEGNTNPPLLIVSFMFTGLSAKGSPEVLPAAMYD
GDVSHESLNKVLTHSREEQLITLCFQISNQLLNALAELEALQEGSQQLKMAEKRRNKFEFIRFGRK
>FLP-12_Diploscapter_pachys_predicted
CSILRKINLLKNCAPAVHWSIETLRVRKYYAHELHLGNKITAEKRRNKFEFIRFGRK
>FLP-12_Heterorhabditis_bacteriophora_predicted
GGQIKMAEKRRNKFEFIRFGRK
>FLP-12_Mesorhabditis_belari_predicted
KKFGIVKLYNYRKFQEVLEEPKKKSNSFNKISKILEPKIAEKRRNKFEFIRFGRK
>FLP-12_Teladorsagia_circumcincta_predicted
VSSQLLNALANIESMQEGMLEVSEQVLLKSTQPLSGTPVKIAEKRRNKFEFIRFGRK

>FLP-12_Bursaphelenchus_xylophilus_BXY_1650400.1
MASRYPGRILGRLLKRPKAWPNRFLSAFTTFFSPKITVTMKSVVASLLAVMVITSLCSAQ
KPHARKGELVESNDEMLARIQGQLLNALEMLQVYQEGPETPQVVGISPSQMKFTEKRRNK
FEFIRFGRR
>FLP-12_Halicephalobus_mephisto_HMEPH_14854-RA.p1
NPISKKSKFSFSTVKTAAVKTIFYKKSKTTIMDKFSISFLLIVLISFANLVVSEKQIQIP
HKGNPEMLETQYYGSNVPSESEILNKIQGQLYNTLELLQTYQEGVPLKFAEKRRNKFEFI
RFGRR
>FLP-12_Panagrellus_redivivus_Pan_g7132.t1
MNPMSQLSLALLLLAVTIATFASAQKNFPSAEVAQPYGNVEEEYLNKVPAQLYNSYGYVQ
AFPDGPHKFAEKRRNKFEFIRFGRR
>FLP-12_Parastrongyloides_trichosuri_PTRK_0000527700.1
MNKFIVLAVLFGLLFATCASSVDTVHDESARATINAGMSDPREVDDFVLRKIYHDLTTSL
DLLEQYAVMNHGVVLSDTHPRLVEKRRNKFEFIRFGRK
>FLP-12_Steinernema_carpocapsae_SC.X.g3517
MTKGHLQSKGSPEMIDPAMVYGSDGEMITRVQSQLLGALQMLQAYQDGPTKFAEKRRNKF
EFIRFGRK
>FLP-12_Steinernema_feltiae_L889_g30089.t1
MIGSALLRSLLLAVLMLFIVSGQKTISQSKGSPEMIEPAMVYGSDGEVISRVQAQLLGAL
QLLEAYQDGPTKLAEKRRNKFEFIRFGRK
>FLP-12_Steinernema_glaseri_L893_g3328.t1
MRHHCVPESKGSPEMLDPALVYGNDGEMITRVQSQLLGALQLLQAYQDGPTKLAEKRRNK
FEFIRFGRK
>FLP-12_Steinernema_monticolum_L898_g27588.t1
MIGSALLRSLFLAVLMVFIVSGQKTIPQSKGSPEMIDPALVYGSDGEVISRVQSQLLGAL
QMLQAYQDGPTKLAEKRRNKFEFIRFGRK
>FLP-12_Steinernema_scapterisci_L892_g4015.t1
MIGSALLRSLLLAVLMLVLVSGQNSVSQSKGSPEMIDPAMVYGSDGEMITRVQSQLLGAL
QMLQAYQDGPTKLAEKRRNKFEFIRFGRK
>FLP-12_Strongyloides_papillosus_SPAL_0001329600.1
MMISKNFIIALLFGFFYITFVVTADTVHDEAARATINAGISDPRENDDFVLRKIYRDLTT
SLELLEQYAVMNHGIFLNENHPRMVEKRRNKFEFIRFGRK
>FLP-12_Strongyloides_ratti_SRAE_X000149300
MNKNIFIAILFGFLYFNCVVAVDTVHDEASRGTINSGISDSRENDDFVLRKIYHDLTSSL
DLLEQYAIMNHGVFLNEGHPRLVEKRRNKFEFIRFGKK
>FLP-12_Strongyloides_stercoralis_SSTP_0000883400.1
MNKNIIIAILFGFLCFNCFAAIDTVHDEASRGVINNGINDQRENDDFVLRKIYRDLMTSI
DLLEQYAVMNHGIFLNEGHPRLVEKRRNKFEFIRFGRK
>FLP-12_Strongyloides_venezuelensis_SVE_0100800.1
MIFSTYIKNGWGRDDKYNLTINRKRTMMSKNFFIALLFGFFYITFVVTADTVHDEAARAT
INAGISDPRENDDFVLRKIYRDLTTSLELLEQYAIMNHGIFLNENHPRMVEKRRNKFEFI
RFGRK
>FLP-12_Rhabditophanes_sp_KR3021_predicted
FKIYDFYFFSASKDISSPNIIDVAMNGEPDIVSAKVRNILKEAIEILNIEDYMSHSAGIVIPDSSSKM
AEKRRNKFEFIRFGRK
>FLP-12_Acrobeloides_nanus_ACRNAN_scaffold3634.g17269.t1
MNPMSQIILLVVLLFGVVFGQKSLPVVKGSPELVEQSPYASDKEVINKVQQQLFNAMELL
QAYEEDAPMKIAEKRRNKFEFIRFGRRR
>FLP-12_Ditylenchus_dipsaci_jg11773
MNPLSAQMIFGTVFLCVFASAAVNAMKPSNKGNPELMENSQVAYGSENELLGKIESQLFG
ALQLLQAYEEAAPMPLKFAEKRRNKFEFIRFGRRRR
>FLP-12_Globodera_pallida_GPLIN_000160300
MIQQIPTALLLVTLATALLMLSGVKTNAQNAHLLVERDFGNAERVNDRPMNGVDGEVIDK
MESRLLGALELLQTYRDAPIVPKFTVKRKNKFEFIRFGKRRR

>FLP-12_Globodera_rostochiensis_GROS_g10245.t1
MHPLTCQPPTRAKLKRSSKTTKMIQQIPTALLLVTLATALLMLSGVKTNAQNAHLLVERD
FGNAERVNDRPMNGVDGDVIDKMESRLLGALELLQTYRDAPIVPKFTVKRKNKFEFIRFG
KRRR*KRREGKRTNGRLGQTRKLVVKGREEDKRTIGPNTEMDEDYARILMR
>FLP-12_Heterodera_glycines_Hetgly.G000019575
MALQISNTAFMLIVLATSLLLLMPSSSSAKVMPQLGFANAEQLMNDNSSPMGAGVEGEVM
DKVEARLLGALELLQSYKEVPIVPKFTEKRKNKFEFIRFGKRRR
>FLP-12_Meloidogyne_arenaria_M.Arenaria_Scaff311g007528
MIYYQQNLFLLVTVCIVSLTLAINVQNMNDLQRNHLIEREFPGENILNAESQLQRQVHTM
DEEMLGRVEAQLMGAMEMLQNYRAASSPAKFTEKRKNNKFEFIRFGR
>FLP-12_Meloidogyne_graminicola_NXFT01001530.1.414_g
MIYYPKNILLFTVFIVATITTSIAINVQNMNDFQRSRLEAREFPGENVLNGESKLQPHTL
DEEMLGRIEVQLMGAMEMLQNYKASSLSPARFTEKRKNNKFEFIRFG
>FLP-12_Meloidogyne_hapla_MhA1_Contig147.frz3.gene38
MMIYYPKNLFLLLTVCIVSVTIAINVQNMNDLQRNHLIEREFPGENILNGESQLQRQVHT
MDEEMLGRVEAQLMGAMEMLQNYRAASSPAKFTEKRKNNKFEFIRFG
>FLP-12_Meloidogyne_incognita_Minc3s00198g07337
MIYYQRNLFLLLTVCIVSLTLAINVQNMNDLQRNHLIEREFPGENILNAESQLQRQVHTM
DEEMLGRVEAQLMGAMEMLQNYRAASSPAKFTEKRKNNKFEFIRFGR
>FLP-12_Meloidogyne_javanica_M.Javanica_Scaff9286g055035
MIYYQQNLFLLVTVCIVSLTLAINVQNMNDLQRNHLIEREFPGENILNAESQLQRQVHTM
DEEMLGRVEAQLMGAMEMLQNYRAASSPAKFTEKRKNNKFEFIRFGR
>FLP-12_Ditylenchus_destructor_predicted
YFNKNAKINCRFGALQLLQAYEEAAPIPAKFAEKRRNKFEFIRFGRRRR
>FLP-12_Meloidogyne_enterolobii_predicted
PSKQLKFIFKLAASSPAKFTEKRKNNKFEFIRFGR
>FLP-12_Meloidogyne_floridensis_predicted
PSKQLKFIFKLAASSPAKFTEKRKNNKFEFIRFGR
>FLP-13_Plectus_sambesii_PSAMB.scaffold8744size5851.g31737.t1
XKSSALGDGKRADNPVIIRFGKRHSLDEDAEDGDLFSVMRRNGKIVDPLIRFGKRGDSIF
SNDPMSFPLTGDMNPAGRFRPTRNQKAQIDPLIRFGKRAAKSADMMIRFGRDPSSRPRRP
DADTMLRFGRAGSSHQPVLMRLGKKSV
>FLP-13_Anisakis_simplex_ASIM_0001211601-mRNA-1
LSRDIRTENFQSAPIIRFGKRMSTQLQGDAKMQSLLRFNRNSQPPHQTARQLLDETVLRF
GRSMSNQAQPLLV
>FLP-13_Ascaris_lumbricoides_ALUE_0000738701-mRNA-1
MAINSISCLITMLAITVAVSTLESSEIRMLESEYGKRGNSYRSFFNPNGELIYNRQIAMN
STYNIILLFYTEFLLSRTIFQGLSSPLIRFGKRTPPEEDLLGRFTRDPQQRIVTDETVLR
FGRSSAPAAA
>FLP-13_Ascaris_suum_AgR036X_g055_t05
ASRFSPKGTFICLVPTVASFSVESLLPPSPSSPAFSLVIALCNKIAFISITNVLPYAPIK
ISVSDGKRGNSYRSFFNPNEYTVVERRDSKLMDPLIRFGKRTNIMGENRLNRNLRAEGLS
SPLIRFGKRTPPEEDLLGRFTRDPQQRIVTDETVLRFGRSSAPAAA
>FLP-13_Brugia_malayi_Bm17974.1
MLTIAFQSMNYLLIALICITLMHSLGSTGIRTFQTEYDITQAERRNQNMMDHLVRDSNGF
GRADYAFGRTARNDAFQTSPLIRFGKRYVMDDEHSEVIAHLLQQYLQQQQQYFDQSNRSP
R
>FLP-13_Brugia_pahangi_BPAG_0001236101-mRNA-1
MMDHLVRDSNGFGRADYAFGRTARNDAFQTSPLIRFGKRYMMDDEHSEVIAHLLQQYLQQ
QQYFDQSNRSPR
>FLP-13_Elaeophora_elaphi_EEL_0000883501-mRNA-1
MTLSLLLLSLSSSIEPIGLLSVLHSSKIFTNKDDIIRAEKRNQNLLDHLMRVSRGFGKSD
RVFGRIARNDAFQTSPLIRFGKRYVVDDGHSEVLAHLLQQYLQQQSNYDESNRYRR
>FLP-13_Litomosoides_sigmodontis_nLs.2.1.2.t08740-RA

MSFKPDLKMPTIAFQPMSYLLTALACITLVHSFELLPTDDVVPPEERNHHLFSYLMRVGR
ALGKSHRTFGGTVRDDAFQASPLIRFGKRYVIDDGRSELLSHLLAQYPQQ
>FLP-13_Loa_loa_EN70_41
MVDIRMDAITESPGRYWPLSHKYDTIDRKYKFFKATNAYRMEAYYPLIRFGRTSYSPAQR
VVTNPKFQETLTTVSDLSDDILLTVFGYCHPIDLIHGCSLVCRRWNYLANYSSLFTEVRV
LVNDFSLEYGSVKMFFHRTSQYLRKLCIDCSLPLPSAEVNALFDICFPNVIHLDLGSFKE
MNTTLLKKLSNCFPNVETLHMEKIERSSRCDDDAQEWEETLKMLFEDESIFPKVQNFFVG
NVTAYCLGNDPKLPVYNRPLNLLHIYSGMDEMEYIGMNEVDFSVIRTSPWRSTLTELHLG
CYITNDGIEYIGLLQNLKVFSWDRSRHIVDEEFAYIKNLYNLEELRVFLGGENCFISSVG
MTALFTLPDEKPEKSFPYKLKHLIIGNYFQTSGDLLQAIDRSCPDLQTLGLPFNEISFFK
DEIMPFIIHNFKHLMFLDLSSLGECYKDEMWNNLNDDDLPNLRLLKLHGNKVNIENLQRL
NLRRPKLLISTRMNHFINWTETENGCVFHDTFDGDIRAIENDLRQIDGLRDFVTKPELYI
YDNFTVNTLERSSSAEFCNEPRSRHPSSDRISRSM
>FLP-13_Onchocerca_volvulus_OVOC3982.1
MIYLVAALACITFTHVESLGIRTLESEYDIAPAERKNENMMDYFVRIGRGSEKPGHIFGR
IARTEAFQTSPLIRFGKRNVMDDARSEVLARLLQYLQQEQPYFDESNRLHH
>FLP-13_Toxocara_canis_Tcan_14437.1
MALNERDVLDGFPLHVIRITAFRLPKQDSAASMTTMWIECAIMWLVFAITINALESSEIR
MLESEYGKRASGYRTLFTPNEYTIIERRDAKLMDPLIRFGKRTTNMGGKQLTRNVRTEDF
QSSPLIRFGKRDLSAEIH
>FLP-13_Wuchereria_bancrofti_WBA_0000537901-mRNA-1
MMDHLLRDSNGFGRADYTFGRTARNDAFQTSPLIRFGKRYVMDDERSELIAHLLHQYLQQ
QQYFDRSNRSPR
>FLP-13_Acanthocheilonema_viteae_predicted
NISISRETEKIFLVLCLIPFIELAIASPLIRFGRNTYPSLYMNLS
>FLP-13_Brugia_timori_predicted
AFQTSPLIRFGKRYVMDDEHSEVIAHLLQQYLQQQQQYFDQSNRSPR
>FLP-13_Dirofilaria_immitis_predicted
AFQTSPLIRFGKRYVMNDGRSEVLARLFQQYPQ
>FLP-13_Dracunculus_medinensis_predicted
MTLDNRIFSIIIGPLIRFGRSKKSGKLISKQ
>FLP-13_Gongylonema_pulchrum_predicted
SFIDKTFCQNFLPNQRSSRLRLQFVALLATIYLADPRYGRLFCWPIPLIRFGRQTKTPLSQFGA
>FLP-13_Onchocerca_flexuosa_predicted
VLQTSPLIRFGKRNVMDDGRSEXXXXXXXXXXXXXXXXXXXXXXXXXXXXXXXXXXXXXXXXXX
XXXXXXXXXXXX
>FLP-13_Parascaris_univalens_predicted
ELTSIRSKINFHLSQSSIDQCERAIFQGLSSPLIRFGKRTPPDEDLLGGFVRDPQQRIATDETVLHFG
RSSAPATA
>FLP-13_Thelazia_callipaeda_predicted
LNSRKLHKFVKIYKSNKKCEIADSLQAFQKSPLIRFGKRFAMEDGLSETLARLQQQQHLLDQEQYLKG
KNRFRR
>FLP-13_Ancylostoma_caninum_ANCCAN_14280
MEGAPLIRFGRTPDAQPLIRFGKRTPDGAPLIRFGRNPEAQPLIRFGKRSQGAPLIRFGR
SVSAPLVRFGRSPEAAPLLRFGRSPEASPLIRFGKTIGAWSIDLTSLLISSHRHPAHHTR
PHHFSIRVPSVLY
>FLP-13_Ancylostoma_ceylanicum_Acey_s0004.g1983.t2
MILLFIETAGPLIRFGKRDVEGAPLIRFGRTPDAQPLIRFGKRTPDGAPLIRFGRNPEAQ
PLIRFGKRSQGAPLIRFGRSVSAPLVRFGRSLEAAPLLRFGRSPEASPLIRFGK
>FLP-13_Ancylostoma_duodenale_ANCDUO_17068
MAFVVSVLNGKRGRGLWFYANDAVEKRSEPFRLPREGRSYDETAGPLIRFGKRDMEGAPL
IRFGRTPDAQPLIRFGKRTPDGAPLIRFGRNPEAQPLIRFGKRSQGAPLIRFGRSVSAPL
VRFGRSLEAAPLLRFGRSPEASPLIRFGKTVLGRSISHRS
>FLP-13_Angiostrongylus_cantonensis_ACAC_0000219901-mRNA-1

MMRTCILVLTVGFAAALDSSETQILYDDYAPDKHSQSQPLLREGRSFDTNAGPLIRFGKR
DPYSAPLIRFGRTPLGQPLIRFGKRSPHESPLIRLPSRLGRNPEASPLIRFGKRSYAAPL
ISSILTLLLCRFGRSPEVSPLIRFGKK
>FLP-13_Angiostrongylus_costaricensis_ACOC_0000366401-mRNA-1
MEWDKYRMQYEGLWAASSHYGCLEEEKKGEAKGEHVNSYQRYIYLNSIAKLQTFQTRPVM
RTCILVLTVGFAAAFDSTETQILYDDYAPDKLSQLQPLLREGRSFDSNAGPLIRFGKRDP
YSAPFIRFGRAPLAQPLIRFGKRSPHSAPLIRYVC
>FLP-13_Caenorhabditis_angaria_Cang_2012_03_13_00256.g7989.t1
MGSPLIRFGKRFETGAPLIRFGKRSPDGAPLIRFGRAPEGSSPLIRFGKRSPGAPLIRFG
RSPSAPLIRFGRSPASAPLIRFGRSSAPLIRFGRK
>FLP-13_Caenorhabditis_brenneri_CBN18134.1
MTSLVFAMMFAVVAVQAFDSSEIRMLDEQYDKSPYFPFLENEKRSDRPTRAMDSPLIRFG
KRAADGAPLIRFGRAPEASPFIRFGKRAADGAPLIRFGRAPEASPFIRFGKRASPSAPLI
RFGRSPSSAPLIRFGRSAGAPLIRFGRASSAPLIRFGRK
>FLP-13_Caenorhabditis_briggsae_CBG05939.1
MMTSLLVIPMMFVVAIQAFDSSEIRMLDEQYETNHPYFPFLEQKRSDRPTRAMDSPLIRF
GKRAADGAPLIRFGRAPEASPFIRFGKRAADGAPLIRFGRAPEASPFIRFGKRAAPSAPL
IRFGRSPSAAPLIRFGRSAAAPLIRFGRASSAPLIRFGRK
>FLP-13_Caenorhabditis_elegans_F33D4.3.1
MMTSLLTISMFVVAIQAFDSSEIRMLDEQYDTKNPFFQFLENSKRSDRPTRAMDSPLIRF
GKRAADGAPLIRFGRAPEASPFIRFGKRAADGAPLIRFGRAPEASPFIRFGKRASPSAPL
IRFGRSPSAVPLIRFGRSAAAPLIRFGRASSAPLIRFGRK
>FLP-13_Caenorhabditis_inopinata_Sp34_40306100.t1
MMVSLLSISMFVIAIQAFESSEIRMIDEQYDTKNPIFEFTEKRSDRPTRAMDSPLIRFGK
RAADGAPLIRFGRAPEASPFIRFGKRGIDGAPLIRFGRAPEASPFIRFGKRASPSAPLIR
FGRSPSSAPLIRFGRSPAAPLIRFGRASSAPLIRFGRK
>FLP-13_Caenorhabditis_japonica_CJA09015.1
MMTSLAVLLVVFVATSEAFDSSEIRMLDEQYDTKNPFLAYLENQKRSDRPTRALDSLDSP
LIRFGKRAADGAPLIRFGRAPEASPFIRFGKRASAVPLIRFGRAPEASPFIRFGKRASPS
APLIRFGRSPAAVPLIRFGRSAAAPLIRFGRK
>FLP-13_Caenorhabditis_latens_FL83_12880
MMTPFLVVSLMFVVSIQAFDSSEIRMLDEQYDSKNPYFPFLENEKRSDRPTRAMDSPLIR
FGKRAADGAPLIRFGRAPEASPFIRFGKRAADGAPLIRFGRAPEASPFIRFGKRASPSAP
LIRFGRSPSAVPLIRFGRSAAAPLIRFGRASSAPLIRFGRK
>FLP-13_Caenorhabditis_nigoni_Cni-.1
MLLLPLSNDYYHLSFSQVVKCCRDSSIVPDPLLKLGPVPIRTMMTSLLVIPMMFVVAIQA
FDSSEIRMLDEQYETNHPFFPFLEQKRSDRPTRAMDSPLIRFGKRAADGAPLIRFGRAPE
ASPFIRFGKRAADGAPLIRFGRAPEASPFIRFGKRAAPSAPLIRFGRSPSAAPLIRFGRS
AAAPLIRFGRASSAPLIRFGRK
>FLP-13_Caenorhabditis_remanei_FL81_05651
MMTPFLVISLMFVVSIQAFDSSEIRMLDEQYEPKNPYFPFLENEKRSDRPTRAMDSPLIR
FGKRAADGAPLIRFGRAPEASPFIRFGKRAADGAPLIRFGRAPEASPFIRFGKRASPSAP
LIRFGRSPSAVPLIRFGRSAAAPLIRFGRASSAPLIRFGRK
>FLP-13_Caenorhabditis_sinica_Csp5_scaffold_00007.g528.t1
MMTSLLVIPLMLVVAIQAFDSSEIRMLDEQFPDDSKNPFYPFLEQKRSDRPTRAMDSPLI
RFGKRAADGAPLIRFGRAPEASPFIRFGKRAADGAPLIRFGRAPEASPFIRFGKRASPSA
PLIRFGRSPSAVPLIRFGRSAGAPLIRFGRASSAPLIRFGRK
>FLP-13_Caenorhabditis_tropicalis_Csp11.Scaffold629.g7970.t1
MMTSLLTISLMFVAAIHAFDSSEIRMLDEQYDAKNPFFPFLDKKRSDRPTRAMDSPLIRF
GKRSADGAPLIRFGRAPEASPFIRFGKRAADGAPLIRFGRAPSASPFIRFGKRASPSAPL
IRFGRSPSSAPLIRFGRSAAAPLIRFGRASSAPLIRFGRK
>FLP-13_Cylicostephanus_goldi_CGOC_0000207701-mRNA-1
MSQTYELNEGFDDIKRLMKVIDFPAEQCQLFLNGKRGRGMWFFGFDPVEKRSDLLHLPRG
ERSYDDNAGPLIRFGKRDPGAPLIRFGRAPDAQPLIRFGKRTPDGAPLIRFGRNPDAQPL

IRFGKRSPAAPLIR
>FLP-13_Dictyocaulus_viviparus_nDv.1.0.1.t03300
PHLPIEETPFDVGINPLVRLGKRTIYSAPFVRFGRTPEVQPIIRFGKRSLDSSSLMRFGS
ELEASPLIRFGKRSPAFHLLRFDRSNAVPFVRFGRSMDAAPLIRFGRTPEVSPLIRFGKK
>FLP-13_Diploscapter_coronatus_DCO_023235
MQLRDWLAVACVAMCGLTIVSAFDSTEIRMLEDELPNADNRIYSRFEDKRSEDTAGMPLI
RFGKRAAQPLIRFGRTPQASPIIRFGKRASPAGSPLIRFGRSAGSPLIRFGRSAGSPLIR
FGRTPSASPLIRFGRK
>FLP-13_Haemonchus_contortus_HCON_00095850
MLRTVVLLLAVSLAVAYDSSELRMLEDEYAMDKRVDTLSRESRSFEENASPLIRFGKRDL
SGAPLIRFGRAPEAHPLIRFGKRAPDSAPLIRFGRDPEASPLIRFGKRSPAAPLIRFGRS
PNASPLIRFGKK
>FLP-13_Haemonchus_placei_HPLM_0001041101-mRNA-1
MRQYNASIFLLSALRSSYHRQSSRFGRAPEAHPLIRFGKRAPDSAPLIRYALLPSLGFAF
FKTSFNVIYPRFGRDPEASPLIRFGKRSPAAPLIRLVISSLAQRLHFG
>FLP-13_Heligmosomoides_polygyrus_HPOL_0001232801-mRNA-1
MLRTCVLLVTVALAAAFDSAELRMLEDEYELEKRSQQMPREERSFDENTGPLIRFGKRDP
GAPLIRFGRAPESAQPLIRFGKRTPDGAPLIRFGRDPDASPLIRFGKRSPAAPLIRFGRS
AAAPLVRFGRSPHAGAPLLRFGRSPDASPLIRFGKK
>FLP-13_Mesorhabditis_belari_mbelari.g18853.t1
MSQFLNGKLELTPSGVKEIRLTSLGYPSLIPNLSSVIERESIKMHNLSLYGIIALSALLI
QIVSAFDSTEIRMLEEEPDTFMKRAASPLIRFGKREFNSSPLIRFGKRSPGAPLIRFGRS
PDAAPLIRFGRSMPMEGPLIRFGRSPNSAPLIRFGKRSPAGPLIRFGRAPEASPLIRFGK
RSAGDEINSNEDEE
>FLP-13_Micoletzkya_japonica_MicoRS5524-mkr-S543-0.26-mRNA-1
MLLVSLLLLLPSLSTAFDSTEIRMLQDEYPLSMRFLPSSFSSPRRYNPRLRSVDSLIPSK
GHLSIGAVISRWNNVHSTEEPSSIPAPSQDKRSAPLIRFGKRSLPMNRDIRQDALGSPLI
RFGKRSNDESEEYQGEEEVEPLERYARAPASAPLIRFGKRAPEPAPLIRFGRAPGSAPLI
RFGKRSSLAPLVRFGRTASAPLIRFGKRSMDD
>FLP-13_Necator_americanus_NECAME_07603
MLRTCVLVLIVGVAAAFDSSEIRMLEDGYHVEKRAAPIHLPREERSFDENAGPLIRFGKR
DSQGAPLIRFGRAPEAQPLIRFGKRSPDSAPLIRFGRDSEAQPLIRFGKRSPAAPFIRFG
RSVSAPLVRFGRSVEAVPLLSLSYVSDHTRKMMMRTVSYVMADHGKHEQSVLGRSHIAID
FLPSSSTLATLEHTSYMSHLFSYSCSIAECC
>FLP-13_Nippostrongylus_brasiliensis_NBR_0000747001-mRNA-1
MLRACVLLLTVALAAAFDSSELRMLEDEYAVEKRSAASSQSLPRDSFDESGSPLMGFRKR
DPGAPLIRFGRAPEAQPLIRFGKRTPEGAPLIRFGRDAEASPLIRFGKRSAPLIRFGRSA
APLVRFGRSPEAAPLLRFGRSPEASPLIRFGKK
>FLP-13_Oesophagostomum_dentatum_OESDEN_15767
MAFVVSVLKRNRGHGLWLFEEDAVEKRADPVRLPREERSFDENAGPLIRFGKRDPQGAPL
IRFGRAPEAQPLIRFGKRTPDSAPLIRFGRNPEAQPLIRFGKRSAAPLIRFGRSMEAAPL
LRFGRSPEASPLIRFGK
>FLP-13_Oscheius_tipulae_OTIPU.nOt.2.0.1.t10391
MAPPVLLFLLGLTSVAAFDSSEIRMLGSDYASSKRSEGFGRVERSLEDGMTGPLIRFGKR
STADGAPLIRFGRAPEASPLIRFGKRNPDGAPLIRFGRAPEASPLIRFGKRTPEGAPLIR
FGRSDGAPLIRFGRSSDYEDLSRDTRAAPLIRFGRK
>FLP-13_Parapristionchus_giblindavisi_Parapristionchus-mkr-
S_359-0.30-mRNA-1
MYTTPLLLLLFSTLIAAFDSTEIRMLQDEYKRSAPLIRFGKRSADSAMQRDERDTMGSPL
IRFGKRSLVDDDEDGVENDEIMERYARAPNSAPLIRFGKRSSQTAPMIRFGRAPSASAPH
IRFGKRSSTAPFVRFGRTASAPLIRFGKRSVDEDQ
>FLP-13_Pristionchus_entomophagus_entomophagus-mkr-S293-2.102-mRNA-1
MERAKHGIRNASTPRGSNSGPHTRSSEMVHYSLALLLTAAVASAFDSTEIRMLQDESGGP
LNDGFGHAAFGPIRLQSREMRDSRIRRQVNDRIREVLGSIRLVKMTVTKDEGDDDSSLED

KRSAPLIRFGKRSVGDEEMARTIRQDALGSPLIRFGKRSAPEEDEELVEEDGDLLERYAR
APASAPLIRFGKRSPDTSPLIRFGRAPSSAPLIRFGKRSSTAPFVRFGRTASAPLIRFGK
RSAM
>FLP-13_Pristionchus_exspectatus_exspectatus-ag_msk-S_571-0.30-mRNA-1
MVHYSLALLLTAALAAAFDSTEIRMLQDESGALPDGFGAAAGFGGPIRLQSRLIGRSDPR
IRRQSSDRIMEALGSIRLVKMTKGNGKHQHQEEEVDDSSLEDKRSAPLIRFGKRSIGDEE
MARTVRQDQGLGSPLIRFGKRSAPEEDEIVEDDGDIMERYARAPASAPLIRFGKRSPDTS
PLIRFGRAPSSAPLIRFGKRSSTAPFVRFGRTASAPLIRFGKRSAM
>FLP-13_Pristionchus_japonicus_japonicus-mkr-S30-4.25-mRNA-1
MKACRYFGVQYIGKEKMVHYSSLLLLLSATLAAAFDSTEIRMLQDESGVLPDGFGAAGFG
GPIRLQSRLIGRSDPRIRRATSDRIMEALGSIRLVKMTKPGKKVEEDDDSSLEDKRSAPL
IRFGKRSIGDEEMARTVRQDQGLGSPLIRFGKRSLPEEEEIVEDDGDVMERYARAPASAP
LIRFGKRSPDTSPLIRFGRAPSSAPLIRFGKRSSTAPFVRFGRTASAPLIRFGKRSAM
>FLP-13_Pristionchus_maxplancki_maxplancki-mkr-S418-0.52-mRNA-1
MVHYSLALLLTAAVASAFDSTEIRMLQDESGVLTDGFGMPGFGGPIRLQSRLIGRSDPRI
RRQRSDRIMEALGSIRLVKMTRPSGKQQPDDEDDSSLEDKRSAPLIRFGKRSIGDEDMAR
TVRQDQGLGSPLIRFGKRSVPEEEEMIEDDGELMERYARAPASAPLIRFGKRSPDTSPLI
RFGRAPSSAPLIRFGKRSSTAPFVRFGRTASAPLIRFGKRSAM
>FLP-13_Pristionchus_mayeri_mayeri-mkr-S16-1.26-mRNA-1
MLAALGSPLIRFGKRSAPDEEDIVDEDGELMERYARAPASAPLIRFGKRSPDTSPLIRFG
RAPSSAPLIRFGKRSSTAPFVRFGRTASAPLIRFGKRSAM
>FLP-13_Pristionchus_pacificus_PPA13711.1
MVHYSLALLLTAALAAAFDSTEIRMLQDESGVLPDGFGAAGGFGGPIRLQSRLIGRSDPR
IRRQSSDRIMEALGSIRLVKMTKANGKQHQEEEEDSSLEDKRSAPLIRFGKRSIGDEEMA
RTVRQDQGLGSPLIRFGKRSAPDEEEIVEDDGDIMERFGRAPSSAPLIRFGKRSSTAPFV
RFGRTASAPLIRFGKRSAM
>FLP-13_Teladorsagia_circumcincta_TELCIR_21494
XAMDKRAEPMPREERSFEEGASPLIRFGKRDFSGAPLIRFGRNPEASPLIRFGKRSPAAP
LIRFGRSPNASPLIRFGKK
>FLP-13_Diploscapter_pachys_predicted
XXXXVACVAMCGLTIVSAFDSTEIRMLEDECEFLASKQLRLSISVPNADNRIYSRFEDKRSEDTAGMP
LIRFGKRAAQPLIRFGRTPQASPIIRFGKRASPAGSPLIRFGRSAGSPLIRFGRSAGSPLIRFGRTPS
ASPLIRFGRK
>FLP-13_Heterorhabditis_bacteriophora_predicted
KSDTASRFGRSPSNAVPLIRFGRSHSSTPFIRYITYNIVD
CIKNAFVRFGRAPDASPLIRFGKK
RFGRTPEATPLIRFGKRSIDGAPFIRFVFKKSSLNAYLD
>FLP-13_Strongylus_vulgaris_predicted
IVLSYLLHCQFQSSLSNRSMFRFGRSVSAPIVRFGRSVDAPLLRQAL
MTAAILRFGRSPEASPLIRFGK
>FLP-13_Bursaphelenchus_xylophilus_XY_0791200.1
MQTSSVLAIVSFLAVLLSCAFAETDSVLDYDDLDEVMAKRAYAGPLIRFGKRSMQRALRN
SPITDPLIRFGKRSDVIESKRPDHSPLIRFGKRAAFRSAPHIRFGKRSDDDALYEW
>FLP-13_Halicephalobus_mephisto_MSTRG.2728.1.p1
MSSINLLILFASICLISICAAFDSSEIRMIEADYEKRNPLNRMFNDYYSETQKRGGLHDP
LIRFGKRSADVEREIRNAMNPLIRFGKRAVPDSAPLIRFGRASLDSAPLIRFGRANFDSA
PLIRFGKRVMDSAPLIRFGKRPNSAPLIRFGKRSSAFSAPHIRFGKRMSEDSLYTRFSRN
RPSPIDQDVMLRFGRSTPATMAEDEEK
>FLP-13_Panagrellus_redivivus_Pan_g23158.t1
MTNITADVDIGALMDVSSILYWSVVSVGLISCPILGRTLWQLFHTYRISQLFSFLLTIVF
LDLIMLGTIIFNVIFEWFPFKGDIVCKLTMVVTNVTACHSNWLWVCMCTQRCVYVFYPMH
RIRRTGFWRIIGNTRRLLVITGMLSLMSQIFWFFLVEEIKVDYNGKQASYCGPDEKVIDA
EMFTYLETLEVLATYFLPFIMTLMADFAVMCKTYDDKRFKTISPKTIRNNQASLGDINVY

LSYKIQSEGGIRAANMRRQRNMRLFLIVATAHGLLNLPYYTGHLVAEYIPNLGGPVFHYV
DTAFYIVYLLQFPFKALFVHWLHKDWCQHLESVGMRKHANSNALKQNQYMDDALIANVRP
GMVSCLRLWFKYKGSRAAQTLYSVQTMLSSGSLVFLFGMLAVVKSFDSSEIRMLEEEYEK
RDPLSQFFDNYAETEKRSPMNGPLVRFGKRSLALATDPLIRFGKRGLDGAPLIRFGKRPD
TSPLIRFGKRSPLSSAPHIRFGKRDIEDAVYSRFSRNRPSPADQDVLLRFGRSTPATMTL
>FLP-13_Parastrongyloides_trichosuri_PTRK_0001632900.1
MIKLTLSLCFLLPITVSGFGIEPAEIRRLDNANIQGKRTPFGDNPLTVIPYTYIIDDMSP
YDNYDKRSPTPLVRFGKREVKNDQLISRELRASMLDSPIVRFGKRSPSGPLVRFGRSSSG
PLVRFGRSPSGPLVRFGRSPSGPLVRFGKRNPGEPLIRFGKRPENLPLIRFGKRSPSGPL
VRFGKRSDSDEEDITQEFEEKEEEF
>FLP-13_Rhabditophanes_kr3021_RSKR_0000737900.1
MIKLVCYLCFLISVCSAFGLEPSQIRQVEAELAEGKRSPLLDGGFLVPYEVLYDEQMEKR
GSYEGKPLIRFGKRASSPMNMNREGRDMLNNPLIRFGKRDPFSTPLVRFGKRDPFSTPLV
RFGKRPESTPLIRFGKRAAGFEGPLVRFGKRSSKFGQPIVRFGRSFGQPIVRFGKRPASF
EGPLVRFGKRNPKFGDPIVRFGRSFGQPIVRFGKRSALPQESEENVQAGEVVEDDDNNAD
IAAATGDEEQ
>FLP-13_Steinernema_carpocapsae_SC.X.g4499.2
MTTSKVFLFAVLAVGAVVAFESSEIRLLEDDLLDDKRSGGGYYGFADEMPIAIERRDGRL
RNPLIRFGKRDAETDEDLSRTARNDFSLNPLIRFGKRSSLNSPLIRFGKRPDQMPLIRFG
KRAPSTAPLIRFGKREVGQEENVFGRSNRAANKPVDDDTILRFGR
>FLP-13_Steinernema_glaseri_L893_g5410.t2
MSADPRSLIAARERSVLVFSGNELGYMATKAEEHIARDGIAKGTLTGSSFFMTKAAALVA
LLAVNVALAFESSEIRLLDDEYDKRGGAGFFGFADDVPIAIERRDGRLQNPLIRFGKRDA
DVEDDFPRTIRNDISLDPLIRFGKRSTLNTPLIRFGKRPDTMPLIRFGKRTPSTAPLIRF
GKRDIEQEESLLGRYSRAAHKPVDADTILRFGR
>FLP-13_Steinernema_monticolum_L898_g12548.t3
MSATRVLIFVLLAIGAGVAFESSEIRLLDDDFLDDKRSGGFYRFGDDLPIAIERRSGHLR
NPLIRFGKRDAEMEDDLSRTVRNDISLDPLIRFGKRSPLNSPLIRFGKRPDTMPLIRFGK
RAPSTAPLIRFGKREVEQDNLFGRYSRAANKPVDADTILRFGR
>FLP-13_Steinernema_scapterisci_L892_g17827.t3
MTTTKTLLFAVLAVGAVVAFESSEIRLLEDDYDKRSGGGYYGFADEMPIAIERRDGRLRN
PLIRFGKRDTESDEDLSRAVRNDYSLNPLIRFGKRSALNSPLIRFGKRPDQMPLIRFGKR
APSTAPLIRFGKRDVGQEENVFGRYNRAANKPVDADTILRFGR
>FLP-13_Strongyloides_papillosus_SPAL_0001499200.1
MIKLLISLSLLIPLTVKGFGIEPAEIRSLDTINVQGKRNLFDSNTFTIVPYEIFFEPVST
YDKRSSIPLVRFGKRSGEFKDNRLITRELRASTLDSPIVRFGKRMSSGPLVRFGRSAPGP
LVRFGRSSAGPLVRFGKRNPKNSPLVRFGRASAGPLVRFGKRSFIEEDELSPDSIGNENE
YI
>FLP-13_Strongyloides_ratti_SRAE_X000047400
MTNMIKQTISLFLLLPLIATSFGIEPAEIRSLDNANVHGKRNILDDESIAVFPYEIFYEP
VLSYDKRSSSPLVRFGKRSSDLKDGSLITRELRASMIDSPLVRFGKRSPSGPLVRFGRSP
AVPLVRFGRSLSGPLVRFGKRNFENLDSQYDKRSGMSPLIRFGRASAGPLVRFGKRSVTE
EDEFMADTMEN
>FLP-13_Strongyloides_stercoralis_SSTP_0000488700.1
MKKKNFYYTYIKIIEIVQNTIKIFQRYYYLFINIKIYILMIKQTISIFFLLLPLIVTSFG
LEPAEIRTLDNTNVQGKRNTLDDDSVAILPYEMFYEPIVSYDKRSSTPLVRFGKRSSDLK
DESLITRELRASMLDSPIVRFGKRSLSGPLVRFGRSPNGPLVRFGRASAGPLVRFGKRSY
LNSLTPFDKRLESSPLIRFGRASSDPLVRFGKRSFMEEDEFLPNIKAD
>FLP-13_Strongyloides_venezuelensis_SVE_1968700.1
MIRLLISLSLLIPLTVKGFGIEPAEIRSLDTINVQGKRNLYDSNTFTIVPYEIVFEPIST
YDKRSSIPLVRFGKRSGELKDNRLITRELRASTLDSPIVRFGKRTSSGPLVRFGRSSAGP
LVRFGKRTLKNSPLVRFGRATSGPLVRFGKRSFIEEDELSPNAIGNENEYI
>FLP-13_Ditylenchus_destructor_Dd_06589
MGAYHTISSLICFSVIFTVSAAFDSSEVRLLEDQNGALKRQAAAYWSYFNDDDALGDASA

VKRSQYAAATSPIIRFGKRSPQYTFGNSMTRHARTQTNMMDSPLIRFGKRSPSTAPLIRF
GKRLPNDPGAPLIRFGKRAFQNSAPHIRFGKRSDSNNLLATDPLTSSEEQLLDSVDPEVA
ADYLYSRFNRNRRPNPVELEDTMLRFG
>FLP-13_Ditylenchus_dipsaci_jg26236
MHSPQSVIDQAAGRPRSCQRQPSYWNYFTTSDNNDGLMKRSQQPYAANTPLIRFGKRSPP
AQFGGMSRQARTQQHMMDPIIRFGKRTPSTAPLIRLENRAGGNFAGSAPHIRFGKRSKEP
LTSSEEIMGYWEPTAADYLYSRINRNTRLNPVDFEDTM
>FLP-13_Globodera_rostochiensis_GROS_g08517.t1
MSRQRPPPIRRIVSRCSLTSPIARRRYLPWPVPTAIPRALFLFLPLLFISSSVSAQSYES
AELRILEEKGGKRMFYQPWSYMKRTPSTEPIIRFEKRSGDDQFMEGFKPLSSADPFRFFK
RSIVVPLMRFGRRPAERAAPLIRFGKRAQIRTANAVPLIRFGRSSEEERQ*MEEKGKERN
GETEDRQREISDHFPFQF
>FLP-13_Heterodera_glycines_Hetgly.G000026186
MFRRCPSLRPLRFFVRPPPPFLSLLSLLFLSLLSVNSQSYESSEMRIMEERGGKRMFYQP
WSFMKRTPSVAKANDFQKRPEGRQFLLRWPSADHFGKRSTVVPLMRFGRRPAERAAPLIR
FGKRAYIRTDAAPLIRFGRSSEERK
>FLP-13_Meloidogyne_arenaria_M.Arenaria_Scaff351g008322
MRQLFNILFPFPPPHYIYSTQTLLTALALHSLLFIVAAALPPHKGIYDSTELTSSEMQSG
KRMSPFISYQPWSYMKRAPTAPIIRFGKRSSWEELIERLNKENEENNFQQQQKRSPNSAP
LIRFGRRLNNAPLIRFGRSWKGNEEKEFNE
>FLP-13_Meloidogyne_enterolobii_scaffold668_cov169.g1168
MRQLFNILFPFPPPHYIYSTQTLLTALALHSLLFIVAAALPPHKGIYDSKELTSSEMQSG
KRMSPFISYQPWSYMKRAPTAPIIRFGKRSNWEELIERLDKENEENNFQQKRSPNSAPLI
RFGRRLNNAPLIRFGRSWKGNEEKEFNE
>FLP-13_Meloidogyne_graminicola_NXFT01003107.1.8088_g
MPFSFYSVYNTQSLLIALVLNSLLFLVTAFPHYSSGIYDTTSEINSSEMQSGKRMSPFIS
YQPWSYMKRAPTAPIIRFGKRSNLEELIERSNENNNQQFYNKRSINSAPLIRFGRRISSA
PLIRFGRSWKGNVGEIINE
>FLP-13_Meloidogyne_hapla_MhA1_Contig1371.frz3.gene13
MMRQFIIPFSSHSIYSTQTLLTAVALPSLFFLVAAFPSHKGIYDSVELSSSEMQSGKRMS
PFISYQPWSYMKRAPTAPIIRFGKRSNWEELIERENKEKINEENNLQQKRSFNSAPLIRF
GRRLNNAPLIRFGRSWKGNEEEKYNE
>FLP-13_Meloidogyne_incognita_Minc3s00008g00527
MRQLFNILFPFPPPHYIYSTQTLLTALALHSLLFIVAAALPPHKGIYDSTELTSSEMQSG
KRMSPFISYQPWSYMKRAPTAPIIRFGKRSSWEELIERLNKENEENNFQQQKRSPNSAPL
IRFGRRLNNAPLIRFGRSWKGNEEKEFNE
>FLP-13_Globodera_pallida_predicted
SLTHFPGAKEATPDGGWDSNPRPSPNPWGAFTPPPPPNSFHLISLQLSSADPFRFFKRSTVVPLIRFG
RRPAERAAPLIRFGKRAQIRTANAVPLIRFGRSSEEERQ
>FLP-13_Soboliphyme_baturini_predicted
KRSTSFTALSLDQLGRMGSSPFIRFGKKSLFILL
>FLP-14_Romanomermis_culicivorax_nRc.2.0.1.t03784-RA
MEIRTVDNNMLIIKIHVVFFPFLFLILNFDVHRFSSSPLTVHGASVGETCEQIENAAEPD
ARKVQICRQKELTETLIDLSTILDASVAAIMPQYGIVPVETDPVVAKRKHEFLRFGKRNQ
QHEFIRFGKRYVVHDEDFENYEKFTSSKTNIFLDEKFKLYLDVPSIRKKIFESLKPGAFL
YRASN
>FLP-14_Soboliphyme_baturini_SBAD_0001074501-mRNA-1
LTLPAVLPSILKALTKQNENLSSLGVSTAELPNYSASEMEKRKHEYLRFGKRKHEYLRFG
KRKHEYLRFAVTHALLNDYCENNYARFTRAIDCRRLEFLFHNSLLTVFG
>FLP-14_Trichinella_britovi_T03_16438.1
MQSIIYIVTFATVICGSLSLEPEEKLCSQVLSSEQLLKEDSGSAQLCRMMQIMTQFQTYV
RLLEETTAQALADRGIIFDIPAEILNDTTFNINKRKHEYLRFGKRKHDYLRFGKRKHEYL
RFG
>FLP-14_Trichinella_murrelli_T05_1139.1

MQSIIYIVTFATVICGSLSLEPEEKLCSQVLSSEQLLKEDSGSAQLCRMMQIMTQFQTYV
RLLEETTAQALADRVYIIVKSQRIIFDIPAEILNDTTFNINKRKHEYLRFGKRKHDYLRF
GKRKHEYLRFG
>FLP-14_Trichinella_nativa_T02_7549.1
LPKDILTWPYRQLKYPSKNWLNKNKMQSIIYIVTFATVICGSLSLEPEEKLCSQVLSSEQ
LLKEDSGSAQLCQMMQIMTQFQTYVRLLEETTAQALADRVYIIVKSQRIIFDIPAEILND
TTFNINKRKHEYLRFGKRKHDYLRFGKRKHEYLRFG
>FLP-14_Trichinella_nelsoni_T07_7856.1
MNILYFKFSTFLFIHRYPSKNKMQSIIYIVTFATVICGSLSLEPEEKLCSQVLSSEQLLK
EDSGSAQLCRMMQIMTQFQTYVRLLEETTAQALADRGIIFDIPAEILNDTTFNINKRKHE
YLRFGKRKHDYLRFGKRKHEYLRFG
>FLP-14_Trichinella_papuae_T10_2079.1
LMPLQLQVWYPSKNWLNKNKMQSIIYVVTFATVVCGSLSLEPEEKLCSQVLSSEQLLKED
SGSAQLCQMMQIMTQFQTYVRLLEETTAQALADRGIIFDIPAEILNDTTFNINKRKHEYL
RFGKRKHDYLRFGKRKHEYLRFG
>FLP-14_Trichinella_patagoniensis_T12_7187.1
MQSIIYIVTFATVICGSLSLEPEEKLCSQVLSSEQLLKEDSGSAQLCRMMQIMTQFQTYV
RLLEETTAQALADRGIIFDIPAEILNDTTFNINKRKHEYLRFGKRKHDYLRFGKRKHEYL
RFG
>FLP-14_Trichinella_pseudospiralis_T4A_11243.1
MYPSKNWLNKNKMQSIIYVVTFATVVCSSLSLEPEEKLCSQVLSSEQLLKEDSGSAQLCR
MMQIMTQFQTYVRLLEETTAQALADRGIIFDIPAEILNDTTFNINKRKHEYLRFGKRKHD
YLRFGKRKHEYLRFG
>FLP-14_Trichinella_sp.T6_T06_8838.1
MYPSKNWLNKNKMQSIIYIVTFATVICGSLSLEPEEKLCSQVLSSEQLLKEDSGSAQLCQ
MMQIMTQFQTYVRLLEETTAQALADRVYIIVKSQRIIFDIPAEILNDTTFNINKRKHEYL
RFGKRKHDYLRFGKRKHEYLRFG
>FLP-14_Trichinella_sp.T8_T08_8720.1
LMPLQLQVWYPSKNWLNKNKMQSIIYIVTFATVICGSLSLEPEEKLCSQVLSSEQLLKED
SGSAQLCRMMQIMTQFQTYVRLLEETTAQALADRVYIIVKSQRIIFDIPAEILNDTTFNI
NKRKHEYLRFGKRKHDYLRFGKRKHEYLRFG
>FLP-14_Trichinella_sp.T9_T09_11701.1
LKQIQVSLVSKDDIQKIIPLPKDILTWPYRQLKYPSKNWLNKNKMQSIIYIVTFATVICG
SLSLEPEEKLCSQVLSSEQLLKEDSGSAQLCRMMQIMTQFQTYVRLLEETTAQALADRVY
IIVKSQRIIFDIPAEILNDTTFNINKRKHENLRFGKRKHDYLRFGKRKHEYLRFG
>FLP-14_Trichinella_spiralis_T01_5322.1
MHFFTFPFIHRYPSKNWLNKNKMQSIIYIVTFATVICGSLCLEPEEKLCSQVLSSEQLLK
EDSGSAQLCRMMQIMTQFQTYVRLLEETTAQALADRGIIFDIPAEILNDTTFNINKRKHE
YLRFGKRKHDYLRFGKRKHEYLRFG
>FLP-14_Trichinella_zimbabwensis_T11_17837.1
MPDMKVSIEELAEQEQDAIYHLRRYFCNCEPEEKLCSQVLSSEQLLKEDSGSAQLCQMMQ
IMTQFQTYVRLLEETTAQALADRGIIFDIPAEILNDTTFNINKRKHEYLRFGKRKHDYLR
FGKRKHEYLRFG
>FLP-14_Trichuris_muris_TMUE_2000007185.1
MMTSATFWIFLAIVILPNVAAQAQEKTCSELLTSALQSQEDRFNAQLCEIMQAAIKMGQT
SSLLEDATEAALQERGVASERSVDSSDRYVMDINKRKHEYLRFGKRKHEYLRFGKRKHEY
LRFG
>FLP-14_Trichuris_suis_D918_06988
MLWIFLAVVIHPSVVGQLQERTCNELLTSQLRSQEDRFNAQLCEIMQAVLKMGRTSALLE
DAIEAALQQRGIANERPVDADERYLVDINKRKHEYLRFGKRKHEYLRFGKRKHEYLRFG
>FLP-14_Trichuris_trichiura_TTRE_0000827001-mRNA-1
LCIANERPVDTDERYLVDINKRKHEYLRFGKRKHEYLRFGKRKHEYLRFG
>FLP-14_Plectus_sambesii_PSAMB.scaffold12452size2758.g34856.t1
MIINVLGVVAALMFLVGTVRADDTATCAQVMATGGDENQTKMLLCQLHESATLLAQLGGL

VSEGLERLMIADGVAPDDDGTGNGIEKRKHEYLRFGKRKHEYLRFGKRKHEYLRFGKRKH
EYLRFGKK
>FLP-14_Acanthocheilonema_viteae_nAv.1.0.1.t08951-RA
MEKAAIIASSTDANDKVLLCQLYESSSLLAELGIFISKDVEKLLANEGVTTDEVQDIEKR
KHEYLRFGKRKHEYLRFGKRKHEYLRFGRK
>FLP-14_Ascaris_lumbricoides_ALUE_0001839401-mRNA-1
MASPGEENHNKMLLCQLFESSTLLAQLGALVSEGLDRLMITQGVVPDVSAEEEGQSIEKR
KHEYLRFGKRKHEYLRFGKRKHEYLRFGRK
>FLP-14_Ascaris_suum_AgR005_g182_t01
MNVQQAGLLLLLGVIIACVSADAPQTCSQVMASPGEENHNKMLLCQLFESSTLLAQLGAL
VSEGLDRLMITQGVVPDVSAEEEGQSIEKRKHEYLRFGKRKHEYLRFGRK
>FLP-14_Brugia_malayi_Bm6384.1
MKGRHHLFSSTLITALIISGCVQYTTGQLQTCSQVFASATEENGKMLLCELYESSSLLAQ
LGTFVSKDVEKLLANEGVTIDDVQDIEKRKHEYLRFGKRKHEYLRFGKRKHEYLRFGRK
>FLP-14_Brugia_pahangi_BPAG_0000853101-mRNA-1
MICCIQVFASSTEENGKMLLCELYESSSLLAQLGTFVSKDVEKLLANEGVTIDDVQDIEK
RKHEYLRFGKRKHEYLRFGKRKHEYLRFGRK
>FLP-14_Brugia_timori_BTMF_0001010801-mRNA-1
MICCIQVFASATEENGKMLLCELYESSSLLAQLGTFVSKDVEKLLANEGVTIDDVQDIEK
RKHEYLRFGKRKHEYLRFGKRKHEYLRFGRK
>FLP-14_Dirofilaria_immitis_nDi.2.2.2.t02505
MKDLLRHLCSSTLITALIISNSVQYTSGQLQTCSQIFASSAEENAKMLLCQLYESSSLLA
QLGIFISKDIEKLLANEGVTTDEVQDIEKRKHEYLRFGKRKHEYLRFGKRKHEYLRFGRK
>FLP-14_Dracunculus_medinensis_DME_0000602901-mRNA-1
ILSNVSYCFTIVFISTAVSLPLSFIFDECILFLFGLETETEIADDGQSIEKRKHEYLRFG
KRKHEYLRFGKRKHEYLRFGRK
>FLP-14_Litomosoides_sigmodontis_nLs.2.1.2.t02031-RA
MQYAAGQLQTCSQIVASSTDENDKVMLCQLYESSSLLAELGTSISKNVENLLANKGVVTA
EDVQDIEKRKHEYLRFGKRKHEYLRFGKRKHEYLRFGRK
>FLP-14_Loa_loa_EN70_3423
MMSSCVQYTTGQLQTCSQVFASSTEENDKVLLCQMYETSSLLAQLGTFVSKDVEKLLANE
GVTMDDVQDIEKRKHEYLRFGKRKHEYLRFGKRKHEYLRFGRK
>FLP-14_Onchocerca_flexuosa_X798_01518
MYNFLYIKSKETFMMYYIFKANRTWLFAAIIIKSSVTVISITHNTVAMKGLLRHLCSSSL
ITALIITISVQYTMGQLQTCSQILASSTEENEKVWLCQLYESSSLLTQLGALVYKDVEKL
LSNEGVITDEVQDIEKRKHEYLRFGKRKHEYLRFGKRKHEYLRFGRK
>FLP-14_Onchocerca_ochengi_nOo.2.0.1.t00427-RA
MKGLLRHLCSSSLITALIITTSVQYTMGQLQTCSQILASSTEENEKVWLCQLYESSSLLT
QLGALVYKDVEKLLSNEGVITDEVQDIEKRKHEYLRFGKRKHEYLRFGKRKHEYIRFGRK
>FLP-14_Onchocerca_volvulus_OVOC8589.1
MKKQHEQFILASSTEENEKVWLCQLYESSSLLTQLGALVYKDVEKLLSNEGVIIDEVQDI
EKRKHEYLRFGKRKHEYLRFGKRKHEYIRFGRK
>FLP-14_Parascaris_univalens_PgR025_g025_t01
VMASPGEENHNRILLCQLFESSTLLAQLGALVSEGLDRLMITQGVVPDVNAEEDGQNIEK
RKHEYLRFGKRKHEYLRFGKRKHEYLRFGRK
>FLP-14_Thelazia_callipaeda_TCLT_0000748501-mRNA-1
MVLASSGEENYNKVLLCQLYESSSLLAQLGALISNGLEKLLINQGSMQEIFAEFNYRLTT
EDDDVQAIEKRKHEYLRFGKRKHEYLRFGKRKHEYLRFGRK
>FLP-14_Toxocara_canis_Tcan_03956.1
MASPGEENHNKMLLCQLYESSTLLAQLGALVSEGLDRLMVTQGVVPDVGAEEEGQGIEKR
KHEYLRFGKRKHEYLRFGKRKHEYLRFGRK
>FLP-14_Wuchereria_bancrofti_maker-PairedContig_1145-snap-gene-1.21-
mRNA-1
MKGRHHLFSSTLITALIISSCVQYTAGQLQTCSQVFATSTEENGKVLLCELYESSSLLAQ

LGTFVSKDVEKLLANEGVTIDDVQDIEKRKHEYLRFGKRKHEYLRFGKRKHEYLRFGRK
>FLP-14_Elaeophora_elaphi_predicted
SNTCVTAQVITGMAIKVIAGVTADDVHDVEKRKHEYLRFGKRKHEYLRFGKRKHEYLRFGRK
>FLP-14_Gongylonema_pulchrum_predicted
KGLSRKINTVTQGCFAGLTADEEDIQGIEKRKHEYLRFGKRKHEYLRFGKRKHEYLRFGRK
>FLP-14_Parascaris_equorum_predicted
EEXXXXXXXXXXXHEYLRFGKRKHEYLRFGKRKHEYLRFGRK
>FLP-14_Ancylostoma_caninum_ANCCAN_11165
MRSELENLWSSGLKLDSFSIGGIAAIEEQAGEVDENGVEKRKHEYLRFGKRKHEYLRFGK
RKHEYLRFGKRKHEYLRFGRK
>FLP-14_Ancylostoma_ceylanicum_maker-ANCCEYDFT_Contig214-augustus-gene-1.14-mRNA-1
MIHWSPIRIAAIEEQSGEVDENGVEKRKHEYLRFGKRKHEYLRFGKRKHEYLRFGKRKHE
YLRFGRK
>FLP-14_Ancylostoma_duodenale_ANCDUO_22440
MGGIAAIEEQSGEVDENGVEKRKHEYLRFGKRKHEYLRFGKRKHEYLRFGKRKHEYLRFG
RK
>FLP-14_Angiostrongylus_cantonensis_ACAC_0000845901-mRNA-1
MTHFELKKKKKKKKATRPRNEIFHHKRCFHGHHPDILSNGGDQQELLLCQLSETSTLLA
QLGILVNEGLDRLMQNQGITSAVDESIEADENGIEKRKHEYLRFGKRKHEYLRFGKRKHE
YLR
>FLP-14_Angiostrongylus_costaricensis_ACOC_0000090701-mRNA-1
MEPCSKKILSNGGDQQELLLCQLSDTSTLLAQLGILVNEGLDRLMQNQGITSAVDESSEA
GENGVEKRKHEYLRFGKRKHEYLRFGKRKHEYLR
>FLP-14_Caenorhabditis_angaria_Cang_2012_03_13_00235.g7585.t1
MVMGQDMTTRWRGQVETSFILANNGDPQDALLCQLSESSMLLAQLGALVSEGVERLVQNH
GIALEDDESNESGEMEKRKHEYLRFGKRKHEYLRFGKRKHEYLRFGKRKHEYLRFGRK
>FLP-14_Caenorhabditis_brenneri_CBN10042.1
MMIGLPTALLLSAFIVAASGQETPAAAAGAVQAPSNPKDCQSILANNGDQQEALLCQLSE
SSMLLAQLGALVSEGVERLVQTHGLALEDETNGENEMEKRKHEYLRFGKRKHEYLRFGKR
KHEYLRFGKRKHEYLRFGRK
>FLP-14_Caenorhabditis_briggsae_CBG15764.1
MMIRLTATALLVLIVAAYGQDAAPGAGAGAVQAAHNPKDCQSILANNGDQQEALLCQLSE
SSMLLAQLGALVSEGVERLVQTHGLALEEETNESDGDMEKRKHEYLRFGKRKHEYLRFGK
RKHEYLRFGKRKHEYLRFGRK
>FLP-14_Caenorhabditis_elegans_Y37D8A.15.1
MMICLPTALLLSAFVVAASGQEAPAGAGASGAAQAPHNPKDCQAILANNGDQQEALLCQL
SESSMLLAQLGALVSEGVERLVQTHGLALEEETNEGDNDMEKRKHEYLRFGKRKHEYLRF
GKRKHEYLRFGKRKHEYLRFGRK
>FLP-14_Caenorhabditis_inopinata_Sp34_30293500.t1
MMISLPTALLLSAFVVVASGQQAGAAASDVQPPQNPRDCQTIVANNGDQQEVLLCQLSES
SMLLAQLGALVSEGVERLVQTHGLALQDETNDAEGDVEKRKHEYLRFGKRKHEYLRFGKR
KHEYLRFGKRKHEYLRFGRK
>FLP-14_Caenorhabditis_japonica_CJA13189a.1
MMICLRTAFVLSAFLAATVAGQAAPASGANAVGPAEPPKDCQSILANNGDQQEALLCQLS
ESSMLLAQLGALVSEGVERLVQGHESDETGGDGGEMEKRKHEYLRFGKRKHEYLRFGKRK
HEYLRFGKRKHEYLRFGRK
>FLP-14_Caenorhabditis_latens_FL83_14515
MMICLPTALLLSAFVVAAYGQEVPAGAGAPAGAVQASHNPKDCQSILANNGDQQEALLCQ
LSESSMLLAQLGALVSEGVERLVQTHGLALEEETNEGENEMEKRKHEYLRFGKRKHEYLR
FGKRKHEYLRFGKRKHEYLRFGRK
>FLP-14_Caenorhabditis_nigoni_Cni-.1
MMIRLTATALLVLIVAAYGQDAAPGAGAGAVQAAHNPKDCQSILANNGDQQEALLCQLSE
SSMLLAQLGALVSEGVERLVQTHGLIPLSESQDEPMIPGLALEEETNESDGDMEKRKHEY

LRFGKRKHEYLRFGKRKHEYLRFGKRKHEYLRFGRK
>FLP-14_Caenorhabditis_remanei_FL81_19395
MMICLPTALLLSAFVVAAHGQEVPAGAGAPAGAVQASHNPKDCQSILANNGDQQEALLCQ
LSESSMLLAQLGALVSEGVERLVQTHGLIPLSSSSSFQNDEDEIPGLALEEETNEGENEM
EKRKHEYLRFGKRKHEYLRFGKRKHEYLRFGKRKHEYLRFGRK
>FLP-14_Caenorhabditis_sinica_Csp5_scaffold_00535.g12566.t1
MIRLPTTLLLLLVVAAYGQETPSAGAPGATGVQAAHNPKDCQSILANNGDQQEALLCQLS
ESSMLLAQLGALVSEGVERLVQTHGLALEDETNEGSVAGADGEMEKRKHEYLRFGKRKHE
YLRFGKRKHEYLRFGKRKHEYLRFGRK
>FLP-14_Caenorhabditis_tropicalis_Csp11.Scaffold629.g16055.t1
MKVVEILANNGDQQEALLCQLSESSMLLAQLGALVSEGVERLVQTHGLALEEETNDGDME
KRKHEYLRFGKRKHEYLRFGKRKHEYLRFGKRKHEYLRFGRK
>FLP-14_Dictyocaulus_viviparus_DICVIV_08607
MTSTEKQLNNLISEATKQKTSSNKLLAVKILSNGADQQELLLCQLSESSTLLAQLGILVS
EGLDRLMQNQGIAVDDSNEVDETGVEKRKHEYLRFGKRKHEYLRFGKRKHEYLSRIFSHM
QIDEGIVEVESYQYRHYAFQTEMNKQITVKDNYSYMMCPEFKSFKGQGYNN
>FLP-14_Diploscapter_coronatus_DCO_023619
MRGLLVALVATVTTTYFVQALDSADTTTADIKDCRYVLANGGEQEELLLCQLQESSQLLT
QLGALVSEGLDRLIQNHGLASSDPSAPAESNGVDKRKHEYLRFGKRKHEYLRFGKRKHEY
LRFGRK
>FLP-14_Diploscapter_pachys_WR25_04893.1
MASGEVLANGGEQEELLLCQLQESSQLLTQLGALVSEGLDRLIQNHGLASSDPSAPAESN
GVDKRKHEYLRFGKRKHEYLRFGKRKHEYLRFGRK
>FLP-14_Haemonchus_contortus_HCON_00084500
MRGLGAGLLLATLSTLLPSSGAAPAETAVAKNCNEILANGGDQQELLLCQLSESSTLLAQ
LGILVSEGLDRLMQNQGIAATEDLDESGVEKRKHEYLRFGKRKHEYLRFGKRKHEYLRFG
KRKHEYLRFGRK
>FLP-14_Haemonchus_placei_HPLM_0000534901-mRNA-1
LGKNRHFQILANGGDQQELLLCQLSESSTLLAQLGILVSEGLDRLMQNQGIAATEDLDES
GVEKRKHEYLRFGKRKHEYLRFGKRKHEYLRFGKRKHEYLRFG
>FLP-14_Heligmosomoides_polygyrus_HPOL_0001716701-mRNA-1
MSATPTSSDFAPAPLRYRIRGTAKWGGWGFGSHIRKMRGLGAGLLLATLSTLLPSSGATA
ADSAVASNCQLRAITNQSNVLIPAIFQILANGGDQQELLLVSAQRESSTLLAQNSRHFSL
SEGLERLMQNQGIAAVEGASEGDDNGIEKRKHEYLRFGKRKHEYLRFGKRKHEYLRFGKR
KHEYLRFGRK
>FLP-14_Heterorhabditis_bacteriophora_Hba_12515
MHYDLFNVCSGFSCTILEIYFPTIYHFCIVHSFSISPILPSIPSSLLLSSDAENLRIISF
FSFRFGKRKHEYLRFGPLVKDSRHWT
>FLP-14_Micoletzkya_japonica_MicoRS5524-mkr-S67-2.53-mRNA-1
MQQSVVAVFVSSLFILSSAEGLTSCAQLSGSSDDNSQKALLCQLAESTQLLAHLGVLVNE
GIERMLQENGLTAQTEDAEEVNKRKHEYLRFGKRKHEYLRFGKRKHEYLRFGRK
>FLP-14_Necator_americanus_NECAME_11622
MISYILCTYEAPKRNQLIQEILSSGGDQQDLLLCQLSESSTLLAQLGILVSEGLDRLMQN
QGIATVEEPAGEVDETGVEKRKHEYLRFGKRKHEYLRFGKRKHEYLRFGKRKHEYLRFGR
K
>FLP-14_Nippostrongylus_brasiliensis_NBR_0001101101-mRNA-1
MFNFDEKRSFRMILILANGGDQQELLLCQLSESSTLLAQLGILVSEGLDRLMQNQGIAAV
ESAAEVDENGIEKRKHEYLRFGKRKHEYLRFGKRKHEYLRFGKRKHEYLRFGRK
>FLP-14_Oesophagostomum_dentatum_OESDEN_19143
IAAVEEPQGVVEENGLEKRKHEYLRFGKRKHEYLRFGKRKHEYLRFGKRKHEYLRFGRK
>FLP-14_Oscheius_tipulae_OTIPU.nOt.2.0.1.t08569
MRTFVGGLLLATLSLFSATLANQQEQTKNCNEILAAGGDQHDLLLCQLSESSTLLAQLGV
LVSEGLDRLIENQGMAAPEAAEGASDVEKRKHEFGKRKHEYLRFGKRKHEYLRFGKRKHE
YLRFGRK

>FLP-14_Parapristionchus_giblindavisi_Parapristionchus-mkr-S_138-1.17-mRNA-1
MQTALSVLLLSTLALNAFADGLASCADLASSGAEDASHKALLCQLAESSQLLAQLGVLVN
EGLERMLQENGLTAQTEEVAEVNKRKHEYLRFGKRKHEYLRFGKRKHEYLRFGRK
>FLP-14_Pristionchus_arcanus_arcanus-mkr-S_722-0.13-mRNA-1
MKSLAIFLISSLAISSLAQTLTSCADITSSTDDSSHKGLLCQLAESSQLLAQLGILVNDG
LERILQENGLTAQTEEGTEVNKRKHEYLRFGKRKHEYLRFGKRKHEYLRFGRK
>FLP-14_Pristionchus_exspectatus_exspectatus-mkr-S_20-3.145-mRNA-1
MKSLAVLFISSLAISSLAQTLTSCADVTSSTDDSSHKGLLCQLAESSQLLAQLGILVNEG
LERILQENGLTAQTEEGTEVNKRKHEYLRFGKRKHEYLRFGKRKHEYLRFGRK
>FLP-14_Pristionchus_fissidentatus_fissidentatus-mkr-S139-4.51-mRNA-1
MKQSLALILLSSLAVSTFAQGLTSCADLSSSSPDDSHKGLLCQLAESSQLLAQLGVLVNE
GLERILQENGLTAQTEEGGDVNKRKHEYLRFGKRKHEYLRFGKRKHEYLRFGRK
>FLP-14_Pristionchus_japonicus_japonicus-mkr-S5-0.34-mRNA-1
MQSSIAILLISSLTVSSLAQSLTSCADITASADDSSHKGLLCQLAESSQLLAQLGVLVNE
GLERILQENGLTAQTEEAGDVNKRKHEYLRFGKRKHEYLRFGKRKHEYLRFGRK
>FLP-14_Pristionchus_maxplancki_maxplancki-mkr-S50-0.19-mRNA-1
VTASSDDSSHKGLLCQLAESSQLLAQLGILVNEGLERILQENGLTAQTDEGGDVNKRKHE
YLRFGKRKHEYLRFGKRKHEYLRFGRK
>FLP-14_Pristionchus_mayeri_mayeri-mkr-S165-0.31-mRNA-1
MQSLALLLLSSLFLSSLAQSLTSCADVTSSPDDSSHKGLLCQLAESSQLLAQLGILVNEG
LERILQENGKGLTAQTEEGAEVNKRKHEYLRFGKRKHEYLRFGKRKHEYLRFGRK
>FLP-14_Pristionchus_pacificus_PPA45403.1
MRTERDMKDTLETIEGTLLEVKLVDENEVTSSTDESSHKGLLCQLAESSQLLAQLGILVN
EGLERILQENGLTAQTEEGTEVNKRKHEYLRFGKRKHEYLRFGKRKHEYLRFGRK
>FLP-14_Strongylus_vulgaris_SVUK_0002021801-mRNA-1
KILINAFFHLGIGAIDEQQQADETGVEKRKHEYLRFGKRKHEYLRFGKRKHEYLRYTGHY
VTSLFFVLKNPFCSGLERESTNIFVLDVSRRVATNGRRLNTDFFCTYLLLSQLRA
>FLP-14_Teladorsagia_circumcincta_TELCIR_03690
MISIGGPALGYRTPGGREATHSGRGFGRHIRKMRGLGAGLLLATLSTLLPSSAAAPADSV
VSKSCNEILANGGDQQELLLCQLSESSTLLAQLGILVSEGLDRLMQNQGIAAAEGTELDE
SGVEKRKHEYLRFGKRKHEYLRFGKRKHEYLRFGKRKHEYLRFGRK
>FLP-14_Cylicostephanus_goldi_predicted
XXXXXXXXXXXXXXXXXXXXXXXXXXXXXXXXETGVEKRKHEYLRFGKRKHEYLRFGKRKHEYLRY
S
NSRFGKRKHEYLRFGRK
>FLP-14_Mesorhabditis_belari_predicted
DSLLSLYKNIVNFKCFRAKNLNLGIADDSEEQSGDMEKRKHEYLRFGKRKHEYLRFGKRKHEYLR
FLIFQFSNNFFRFGKRKHEYLRFGRK
>FLP-14_Bursaphelenchus_xylophilus_BXY_1631500.1
MSRVADPSSLLMANTKIIWSLLGFVFVVQVIEATVAQTQVEVNCGRILANNNLDNEDKQL
LCKIYQQSSSLEQLGAIVSDSLERFMGEATALEEEARPKRKHEYLRFGKRKHEYLRFGKR
KHEYLRFGRK
>FLP-14_Halicephalobus_mephisto_HMEPH_14086-RA.p1
MIFSRIQLALGTFFLVSTACFQASGADSVVQSCNQIMANGAENDEKLLLCQLYESSNLLA
QLGVLVTEGLDQLMLNQGLITEGENEAPDARQKRKHEYLRFGKRKHEYLRFGKRKHEYLR
FGRK
>FLP-14_Panagrellus_redivivus_Pan_g7979.t1
MHITQTYACAGLFVFAAVLMAPNAAADPTAQNCNQILAAGNEGDERVLLCQLYESSSLLA
QLGVLVTEGLDRLMVNQGLTPEVVDGPDSADARQKRKHEYLRFGKRKHEYLRFGKRKHEY
LRFGRK
>FLP-14_Parastrongyloides_trichosuri_PTRK_0000543200.1
MFIPSNGLYAALAVLIVIVGTTLSAADSGAPTCEEIMKTTNELEPKFLTCKVYHDSMASA

VEAIRVTKELEHLLILNGIAIDSNDIESLNGSEDMEESREKRKHEYLRFGKRKHEYLRFG
KRKHEYLRFGRK
>FLP-14_Steinernema_carpocapsae_L596_018036
MVPSSQQFGVLLAGILSTLLVASSVGADAAVAASATTTQCAQILANPEEYSEKTLLCQLY
ESSNLLAELGVLVSEGIDKLMVAQGLSNENEEQATGSGIEKRKHEYLRFGKRKHEYLRFG
KRKHEYLRFGRK
>FLP-14_Steinernema_feltiae_L889_g27993.t1
MVPQRFGVLLAAIVSTLLVSATVGADAAVAAPATTTQCAQILANPDEYSEKTLLCQLYES
SNLLAELGVLVSEGIDKLMLAQGLALENEEQATGSGIEKRKHEYLRFGKRKHEYLRFGKR
KHEYLRFGRK
>FLP-14_Steinernema_glaseri_L893_g25145.t1
MTTHDALLLRFRSTSTASSFAASQWHVECGKYSSGYDTSFSGPLAPLRAAKKRVALGQFC
LKKDSFTPEAVFAPDHSGNXXXXPFLSTTLLPRRRSHFRASYKDFPRYSSGYGTSFSGPL
PPLRAAKKGVALGQSCLKKDSVTPEASAMVPTSQQLGVLLAGILSTLLVSATVGADAAVA
PPATTTQCAQILANPDEYNEKALLCQLYESSNLLAELGALVSEGVDKLMVANGLSAENEE
QASGSGIEKRKHEYLRFGKRKHEYLRFGKRKHEYLRFGRK
>FLP-14_Steinernema_monticolum_L898_g35187.t1
MVPSSQQFGVLLAGILSTLLVVTNVGAEAAVAAPATTTQCAQILANPDEYNEKTLLCQLY
ESSNLLAELGVIVSEGIDKMMVAQGLSTENEEQAAGSGIEKRKHEYLRFGKRKHEYLRFG
KRKHEYLRFGRK
>FLP-14_Steinernema_scapterisci_L892_g26746.t1
MVPSTQQFGVLLAGILSTLLVASSVGAEAAVAASATTTQCAQILANPEEYSEKTLLCQLY
ESSNLLAELGVLVSEGIDKLMVAQGLSNENEEQASGNGIEKRKHEYLRFGKRKHEYLRFG
KRKHEYLRFGRK
>FLP-14_Strongyloides_papillosus_SPAL_0000306700.1
MFISSYGLYVALVVFVAFFGTSCIAADLSSPTCNEIYKTSSELDAKFLACKVYHNSIATS
LEAIRVTKELEQLLVLNGIVVEGNETGIPEIADDGEDEREKRKHEYLRFGKRKHEYLRFG
KRKHEYLRFGRK
>FLP-14_Strongyloides_ratti_SRAE_2000302200
MFISPYGLYVALAVIVAFFGTTCIAADLSTSTCSEIYKTTNELDPKFLACKVYHNSIATS
LEAIRVTKELEQLLILNGIVLEENETNVSESTDEVEEDREKRKHEYLRFGKRKHEYLRFG
KRKHEYLRFGKK
>FLP-14_Strongyloides_stercoralis_SSTP_0000050600.1
MFVSSYGLYVALAVIVAIFGTTCITADLSPSTCSEIYKTSNELDPKFLACKVYHNSIATS
LEAIRVTKELEQLLILNGIIIEGNETGVSDLSNDGEEDREKRKHEYLRFGKRKHEYLRFG
KRKHEYLRFGKK
>FLP-14_Strongyloides_venezuelensis_SVE_1593600.1
MFISSYGLYVALVVFVAFFGTSCIAADLSSPTCNEIYKTSSELDAKFLACKVYHNSIATS
LEAIRVTKELEQLLVLNGIVVEGNETGIPEIADDGEEEREKRKHEYLRFGKRKHEYLRFG
KRKHEYLRFGRK
>FLP-14_Acrobeloides_nanus_ACRNAN_scaffold304.g19871.t1
MSARTGILCFIASVLLSFQLSSADPISQSCSQIVANNPEGDEKVLLCQLYESSSLLAQLG
ALVHDGLERLMLNQGISTDSDSENREKRKHEYLRFGKRKHEYLRFGKRKHEYLRFGRK
>FLP-14_Ditylenchus_destructor_Dd_01791
MTPLNSVSGVAGSLLWILLLMFNNQWRTMAAPTEGDVLLVNKAPATCAQIIGSSNIENDE
KLLLCQLYESSALLTQLGALVSQGLDRLMISQGVQPLPLNGNSDAAVDTEGMESQKDKRK
HEYLRFGKRKHEYLRFGKRKHEYLRFG
>FLP-14_Globodera_rostochiensis_GROS_g08133.t1
MPAPIHAKPSVPTNLLLMLTDSASPPSPAPRPPYSTMTIRSSSCSFLPLSSLLHRGFVVL
CALSVLQFKPSLTTAAAVFAESGPSSVEQFVHSDCAQLAGGDEERLLLCQLYESSALLAQ
LGVLVNEGIGRLAVSQGMGNKFIVADSGREKRKHEYLRFGKRKHEYLRFGRK
>FLP-14_Meloidogyne_arenaria_M.Arenaria_Scaff3291g040356
MQPSNNSFLMVILSLFCVLVCLLQPGLAENGGDNCAQLAGGDEERLLLCQLYESSTLLSQ
LGNFVTEGIERLAATHGLAEKDAGREKRKHEYLRFGKRKHEFVRFGRK

>FLP-14_Meloidogyne_enterolobii_scaffold7403_cov181.g10667
MVILSLFCVLVCLLQPGLAENGGDNCAQLAGGDEERLLLCQLYESSTLLSQLGNFVTEGI
ERLAATHGLAEKDAGREKRKHEYLRFGKRKHEFVRFGRK
>FLP-14_Meloidogyne_floridensis_maker-nMf.1.1.scaf02119-augustus-gene-0.9-mRNA-1
MQPSNNSFLMVILSLFCVLVCLLQPGLAENGGDNCAQLAGGDEERLLLCQLYESSTLLSQ
LGNFVTEGIERLAATHGLAEKDAGREKRKHEYLRFGKRKHEFVRFGRK
>FLP-14_Meloidogyne_graminicola_NXFT01001407.1.3624_g
MQSSNNFKTLLAVFCVLTFLTKSGLAENDSCTQLAGGDEERLLLCQLYESSTLLSQLGNF
VTEGIERLASTHGLIAGDAGREKRKHEYLRFGKRKHEFVRFGKK
>FLP-14_Meloidogyne_hapla_MhA1_Contig147.frz3.gene58
MQSSNNSLIILSLFCGLVCLLKPGLAENGDTCAQLAGGDEERLLLCQLYESSTLLAQLGN
FVTEGIERLAATHGLAGRDAGREKRKHEYLRFGKRKHEFVRFGRK
>FLP-14_Meloidogyne_incognita_Minc3s00396g11593
MQPSNNSFLMVILSLFCVLVCLLQPGLAENGGDNCAQLAGGDEERLLLCQLYESSTLLSQ
LGNFVTEGIERLAATHGLAEKDAGREKRKHEYLRFGKRKHEFVRFGRK
>FLP-14_Meloidogyne_javanica_M.Javanica_Scaff1293g014508
MQPSNNSFLMVILSLFCVLVCLLQPGLAENGGDNCAQLAGGDEERLLLCQLYESSTLLSQ
LGNFVTEGIERLAATHGLAEKDAGREKRKHEYLRFGKRKHEFVRFGRK
>FLP-14_Ditylenchus_dipsaci_predicted
FIQSIHIKNFRYLRFGKRKHEYLRFGKRKHEYLRFG
>FLP-14_Globodera_pallida_predicted
SGENGFWHRPQGMDTDRSTTKSSDHFGDHFQYLRFGKRKHEYLRFGRK
>FLP-15_Plectus_sambesii_PSAMB.scaffold17825size1049.g37415.t1
MLSSVLLVLTVTLVVTIHSTASMPTLALIGYESEDVSSAIKRIPEIARLRELRSSDVPKP
YSLNAVRSGRPNGPLRFGKRRAGPSGPIRF
>FLP-15_Ascaris_lumbricoides_ALUE_0000479901-mRNA-1
MSIPHEFLFFSLLCLLLASSQAQEYDGADFADVEVSNKFKKGRPRGPLRFGKRAVYDWRS
PFNKPYMQRLGSLDLYEER
>FLP-15_Ascaris_suum_AgR001_g577_t01
MSIPHEFLFFSLLCLLLASSQAQEYDGADFADVEVSNKFKKGRPRGPLRFGKRAVYDWRS
PFNKPYMQRLGSLDLYEER
>FLP-15_Gongylonema_pulchrum_GPUH_0002397701-mRNA-1
MKSLSRNFYVSCANTWNLTPTVLQLGKRYISVKIDVNIAHKRKKARPKGPLRFGKRGAGG
LTLPQDWAAYLLRSYPQSNTGRGWNYVD
>FLP-15_Parascaris_univalens_predicted
RFDRFVYFFAAKEVFSVKKGRPRGPLRFGKRAVYEWRSLSNKPYMQRRGSFDLYEER
>FLP-15_Thelazia_callipaeda_predicted
EEITINQAKYNLLFLRSHIMISGKKARPKGPLRFGKRDQTSLILPFRNDQNREYMFRSNPSDYLDWTW
LYNS
>FLP-15_Toxocara_canis_predicted
KLYCFAVKKGRPRGPLRFGKRALQHDWLYYQPYMQRIISPGLYNER
>FLP-15_Ancylostoma_caninum_ANCCAN_05925
MGFEGANFATNDAFDVPFKDLKKAGPQGPLRFGKRRDGPTGPLRFGKRSSLDYSPLAAQQ
PHYFFV
>FLP-15_Ancylostoma_ceylanicum_Acey_s0038.g3614.t1
MQSCCSSDLTRLISEIDDVVSEPEYFVPYLKKAGPQGPLRFGKRRDGPTGPLRFGKRSSL
DYSPLAAQQPHYFFV
>FLP-15_Ancylostoma_duodenale_ANCDUO_05281
MVYILSSCSTKLHTSTDLKKAGPQGPLRFGKRRDGPTGPLRFGKRSSLDYSPLTAQQPHY
FFV
>FLP-15_Angiostrongylus_cantonensis FLP-14 ACAC_0000834801-mRNA-1
MIHGLIDHVSDLKKGGPQGPLRFGKRRSGPSGPLRFGKRSQLDYPFLYGQPSHILMA
>FLP-15_Angiostrongylus_costaricensis_ACOC_0000722201-mRNA-1

MIYDLIDSVSDLKKGGPQGPLRFGKRRSGPSGPLRFGKRSQLNYPLLYGQPPYNLVE
>FLP-15_Caenorhabditis_angaria_Cang_2012_03_13_00287.g8533.t1
MEFSQLIRFVILAVLALTAFAEYEDSFGPSVDVDYFGNYVKKAGPSGPLRFGKRRGPSGP
LRFGKRSGQPAFDSHDWAYNN
>FLP-15_Caenorhabditis_brenneri_CBN07306.1
MQFSTLFRVALLAVLAIAALADYDDNSIGTIPVAVDLDYFSNYVKKGGPQGPLRFGKRRG
PSGPLRFGKRSFHAAPAPEDVMSWYQ
>FLP-15_Caenorhabditis_briggsae_CBG18242.1
MQFSTLFRFAFLAVLAVSAFADYDDSIGTIPVAVDLDYFSNYVKKGGPQGPLRFGKRRGP
SGPLRFGKRSFHTALAPEDVVSFYQ
>FLP-15_Caenorhabditis_elegans_ZK525.1.1
MQFSTLIRVAVFAVLAIATLADYDDNSVGTIPVAVDLDYFSNYVKKGGPQGPLRFGKRRG
PSGPLRFGKRSSFHVAPAAEDVASWYQ
>FLP-15_Caenorhabditis_inopinata_Sp34_30104200.t1
MQISTLLRVTFLAILSISAFADYDDNSIGTIPVAVDLEYFSNYEKKGGPQGPLRFGKRRG
PSGPLRFGKRSFRAAAAAPSEDVVSWYQ
>FLP-15_Caenorhabditis_japonica_CJA09546.1
MHSATLVRFALLAVLAVAVIADYDDPAYAGVVPVAVDVDYFSNYVKKGGPQGPLRFGKRR
GPSGPLRFGKRSFHAAPAPEDVVSWYQ
>FLP-15_Caenorhabditis_latens_FL83_18183
LTEKVGGVLSLAARPPSSCSRELPVASLIFGRRRSLFSFDSHHFSHLVMQFSTLFRVVFL
AVLAIVALADYDDNSVGTVPVAIDLDYFSNYVKKGGPQGPLRFGKRRGPSGPLRFGKRSS
FHAAPAPEDVVSWYQ
>FLP-15_Caenorhabditis_nigoni_Cni-.2
MPIGNYDDSIGTIPVAVDLDYFSNYVKKGGPQGPLRFGKRRGPSGPLRFGKRSFHAAPAP
EDVVSFYQ
>FLP-15_Caenorhabditis_remanei_FL81_19470
MLTEKVGGVLSLAARPPSSCSRELPVASLIFGRRRSLFSFHSHHFSHLVMQFSTLFRVVF
LAVLAIVALADYDDNSVGTIPIAVDLDYFSNYVKKGGPQGPLRFGKRRGPSGPLRFGKRS
SFHAAPAPEDVVSWYQ
>FLP-15_Caenorhabditis_sinica_Csp5_scaffold_00269.g8356.t1
MIMRNPMKSSNTHQEYPVSLPKLHSISAQNTQLFKRYPILSCSNSSSSLHQTMFHSPFST
SPSVQFPQNFPESRALTLKKWAESGAFRQRAPPHSCVPRKPSPCSRNRFTHFLEEEEEGL
FSLFPHSHHFLAPHLVMQFSTLFRVAFLAVLAIAALADYDDSVGTIPVAVDLDYFSNYVK
KGGPQGPLRFGKRRGPSGPLRFGKRSYHAAPAQEDVVSFYQ
>FLP-15_Diploscapter_coronatus_DCO_024838
MPSLIQSLSLFSLLVALIITVIRIESTSASELEDSFLDGGDPYASYVKKSPLGPQGPLRF
GKRSPHGPSGPLRFGKRQAPSFYELY
>FLP-15_Haemonchus_contortus_HCON_00084140
MHGYSIVRLLLVLLAAVCIFAEIEDITDDSKYFVPYLKKAGPQGPLRFGKRRGPSGPLRF
GKRSTFDYPTVFDQQPYYYFV
>FLP-15_Haemonchus_placei_HPLM_0001624501-mRNA-1
LPRAKNILYLISDLKKAGPQGPLRFGKRRGPSGPLRFGKRSTFDYPTVFDQQPYYYFV
>FLP-15_Heligmosomoides_polygyrus_HPOL_0001772001-mRNA-1
MGVALRGLLPLMAFCLRRRTASGTLSPVLMHSYSILRLLLVLLFSVCIFAEIDDVTNDSE
YFVPYLKKAGPQGPLRFGKRRDGPSGPLRFGKRSSQLDYPPPLSQQPYFMLV
>FLP-15_Heterorhabditis_bacteriophora_Hba_13236
MYCYSITRLLLVVLLVVLVVADIEEPANEPEFFVSYLKKAGPQGPLRFGKRRGGPSGPLR
FGKRSDPLQPYPYYFLL
>FLP-15_Mesorhabditis_belari_mbelari.g23225.t1
MGGTSSTLWSPSLVSSLREPPLLHISVKMHSRLSFLVLALFLLMLNFESTEALDQFGEEP
ISYVKKGRPQGPLRFGKRKGPSGPLRFGKRSDGLLLSDGLGWGYLME
>FLP-15_Micoletzkya_japonica_MicoRS5524-mkr-S202-0.5-mRNA-1
MLSLSRLALLFALLVAATLAQYDEPRFVELAMPYVKKARPNGPMRFGKRRGPSGPMRFGK

RSAPVELEDADSLEGY
>FLP-15_Necator_americanus_NECAME_02464
MMKKKVNSRVLLKWRELTRWEKPEGGWAEPCLAVAEPGTSPQAKLTARYTSLMHSYGILR
LLVLLIVAVCVFADIDEVASEPEYFVSCEYLHNLKKAVPQGPLRFGKRRDGPTGPLRFGK
RSSLDFQPMATQQPYYFLV
>FLP-15_Parapristionchus_giblindavisi_Parapristionchus-mkr-
S_279-0.53-mRNA-1
MFSFAHLLFIIALLAAAVTAQYDSDGVEFVNPYVKKARPQGPMRFGKRRGPSGPMRFGKR
SAMVPLDQYYDSAAVADSY
>FLP-15_Pristionchus_arcanus_arcanus-ag_msk-S_470-0.37-mRNA-1
MFSLTRLVALLAILAAFAAAQFDDGAMEYAMPYVKKARPQGPMRFGKRRSGPSGPMRFGK
RSAMVPFDYFYDGEAEQ
>FLP-15_Pristionchus_entomophagus_entomophagus-mkr-S332-2.3-mRNA-1
MFSLTRVIALLLILASFASAQYADDAIEWMPYVKKAKPQGPMRFGKRRAGPSGPMRFGKR
SAMVPFDFYYDGEADQ
>FLP-15_Pristionchus_exspectatus_exspectatus-sn_msk-S_762-0.3-mRNA-1
MEGPKWNVKKARPQGPMRFGKRRSDSSCPPVPFGAGNPPGPAPPGDRATKKQANKCDLFS
KLLVGKKE
>FLP-15_Pristionchus_japonicus_japonicus-mkr-S186-3.19-mRNA-1
MFSLTRLVALLLILGTFAAAQFDDGAIEYVMPYVKKARPQGPMRFGKRRAGPSGPMRFGK
RSAMVPFDYFYDGEIEQ
>FLP-15_Pristionchus_maxplancki_maxplancki-mkr-S217-3.44-mRNA-1
AFSAPSRVLHFRARMFSLARLVALLLVLAAFASAQFDDGAVEYVMNSGPSGPMRFGKRSA
MVPFDYFYEGEAEQ
>FLP-15_Pristionchus_mayeri_mayeri-mkr-S428-0.8-mRNA-1
MFSLTRVLALLLILASVAVAQFDDGALEYAMPYVKKAKPQGPMRFGKRRGPSGPMRFGKR
SAMVPFDFYYDGEADQ
>FLP-15_Pristionchus_pacificus_PPA36868.1
MFSLTRLVALLVILTALAAAQFDDGAIEYAMPYVKKARPQGPMRFGKRRFGPSGPMRFGK
RSAMVPFDYFYDGEAEQ
>FLP-15_Strongylus_vulgaris_SVUK_0000574701-mRNA-1
LQKVLSNQLLVSYIPCVSSFEKDLKKAGPQGPLRFGKRRDGPSGPLRFGKRSLSNSNPIA
QQPYYYLL
>FLP-15_Teladorsagia_circumcincta_TELCIR_02801
MKFACKGQSTHLPTHMNGKIIRLDPKAFSTKRYEDVLPADVGDLKKGGPQGPLRFGKRRG
PSGPLRFGKRSAYDYRALFDQQPYYFV
>FLP-15_Diploscapter_pachys_predicted
ELQHLAKSRLSPPRLLESIFNAFPSHAKHFTGLVKFQVFVISESYFAEHSRKKTISQMSISDVKKSPL
GPQGPLRFGKRSPHGPSGPLRFGKRQAPSFYELY
>FLP-15_Oesophagostomum_dentatum_predicted
LFNFRTSQTDLLALESPAQVLVCLGQRSKNALISAISDLKKAGPSGPLRFGKRRDGPSGPLRFGKRSS
LFDSSLSAQQPYYFLV
>FLP-15_Oscheius_tipulae_predicted
FVESVLFSVQSPVFLESEPPVPLYSHPMFKNHFSDLKKGGPQGPLRFGKRRGGPSGPLRFGKRSSFVD
FEY
>FLP-15_Steinernema_carpocapsae_L596_018859
MLRFSVSTLLLFAFVVLFCIFGSVHSQQDYGLTEVDDEIAIPFKKGRPKGPLRFGKRSPM
APGGPLRFGKRRVGRGDPFQRYLQRVILPKLFEEDAEF
>FLP-15_Steinernema_feltiae_L889_g21305.t1
MRSSLSTLLLFAFVTLFCFIGPVHSQEFHSADVGGFGGLDDEIAIPLKKGPKGPLRFGKR
APGGPLRFGKRRADLFQRYFQRVILPTIFDENQNSEA
>FLP-15_Steinernema_glaseri_L893_g32281.t1
MRSRLSTLLLLAFVALFCVLGSVHSQEYPIAEGFDGLDDAIAVPLKKGRPKGPLRFGKRA
PGGPLRFGKRRADVFQKYLQRVVLPTLFEDADM

>FLP-15_Steinernema_monticolum_L898_g6202.t1
MSTSISALLLLAFVALFAVFGSVHTQEYHLADVGGFSGLDDEIAIPLKKGRPTGPLRFGX
XXXXXXXXXXXXXXXXXXXXXXXXXXXXXXXXXGSAEPIPSRGTSKG
>FLP-16_Plectus_sambesii_predicted
RRHHTLRIAVRRQTFVRFGR
>FLP-16_Anisakis_simplex_ASIM_0001313701-mRNA-1
MTSVFVLFGFVAFIVIMSNAAILNIAKEPVSSDSEAAASENGFSAEDVHAASRPLDKRAQ
TFVRFGKRAQTFIRFGRDAC
>FLP-16_Ascaris_lumbricoides_ALUE_0000746501-mRNA-1
MLGCKTSFQITEMASALAFFGFFGCIVMFSYASVLNIPKDPRPEISDVRSMDEAMQKAYA
QRYRLFLENLLSEAALENRLSAGDVYAASRPLDKRAQTFVRFGKRAQTFVRFGRDASTHP
TEATQM
>FLP-16_Ascaris_suum_GS_15634
MDEAMQKAYAQRYRLFLENLLSEAALENRLSAGDVYAASRPLDKRAQTFVRFGKRAQTFV
RFGRDACKQ
>FLP-16_Brugia_malayi_Bm3396.1
MLFSIALISSTGAAVYSPRRIVLPDDDEIQREMINDLLQREYAERYREYLEKSLAAASKN
NMDDIETHAELHPSNKRGQTFVRFGKRSQSSVRFDN
>FLP-16_Brugia_pahangi_BPAG_0000960501-mRNA-1
MVITTYFLSALMLFSIALISSTGAAVYSPRRIVLPDDDEIQREMINDLLQREYAERYREY
LEKSLAAASKNNMDDIETHAELHPPNKRGQTFVRFGKRSQSSVRFDN
>FLP-16_Brugia_timori_BTMF_0001610201-mRNA-1
MINDLLQREYAERYREYLEKSLAAASKNNMDDIETHAELHPSNKRGQTFVRFGKRSQSSV
RFDN
>FLP-16_Dirofilaria_immitis_nDi.2.2.2.t07136
MVLITYLLSAFMLLSIPLISSTNAAIYNPRRIVLPADEEIQREMINDLLQREYADRYRDY
IEKGLAALSKSNMDDIDALHPNNKRGQTFVRFGKRSQPSVRFGD
>FLP-16_Dracunculus_medinensis_DME_0000259401-mRNA-1
MKIFSDPIIFSFIFKMINSNKIVAKEVSGKLRPLNKRAQTFVRFGKRAQTFVRFGRNQCM
LFFYRIFLDFLLINSRKETAIHFRSIFAAFP
>FLP-16_Gongylonema_pulchrum_GPUH_0000585701-mRNA-1
MLDDLLQREYADRYRDYIESGLSAADAENAVAGGEVRAALRPQDKRAQTFVRFGKRAQTF
VRFGK
>FLP-16_Litomosoides_sigmodontis_nLs.2.1.2.t04255-RA
MFVTTYFLPTLLLSSIALISLTSAAAYSPRRIVLPNNDEEVQRDMLNDLLQREYADRYRE
YIERGLAELTNNLGSIDAHSELRPFKRGQTFVRFGKRSQPSVHLAN
>FLP-16_Loa_loa_EN70_9341
MLFSIALISSTSAGVYSPRRIVLPDDEEMQREMINDLLQRDYADRYRTYLEKGLAAASKN
TIDDIEIHPELHSSNKRGQTFVRFGKRSLTSAGFDK
>FLP-16_Onchocerca_flexuosa_X798_07695
MLQHKLGQQQAIQMVFATCFLPAITLFSISLISSSSATIYNPRRIVLPIDEEMQREMIND
LLLRDYADRYREYIEKALTKNNLDDLETLHSGNKRGQTFVRFGKKSQFSFR
>FLP-16_Onchocerca_ochengi_nOo.2.0.1.t01542-RA
MVFTTFFLPAIMLFSISLISSSNAAIYNPRRIVLPVDEEIQREMINDLLLRDYADRNREY
IEKGLAALAKNNLDDLETLHSGSKRGQTFVRFGKRSQVSFR
>FLP-16_Onchocerca_volvulus_OVOC11158.1
MVFTTFCLPAIMLFSISLISSSNAAIYNPRRIVFPVDEEIQREMINDLLLRDYADRNREY
IEKGLAALAKNNLDDLETLHSGSKRGQTFVRFGKRSQVSFR
>FLP-16_Thelazia_callipaeda_TCLT_0000593301-mRNA-1
MFLQTKINHFIFQIKIFYLRFNLVLCLHSFTANIGAKNGIGGANARPELQFANKRGQTFV
RFGKRSQTFVRFGK
>FLP-16_Toxocara_canis_Tcan_12963.1
MTSALVFFSFFASTVMLSKAAILNVAKEPLPENNDIRNADDALQKAYAQRYRLLLENLPV
EAALENGLSAEDIHVASRPLDKRAQTFVRFGKRAQTFVRFGRDACQLRSSCSIHILHYLP

TLDRNLKHILTSRLQHNLPVIIGYLNL
>FLP-16_Wuchereria_bancrofti_WBA_0000084401-mRNA-1
MVITTYLLSALMLFSIALISSTGAAVYIPRRIVLPDDDEMQREMINDLLQREYAERYREY
LEKGLAAASKNNMDIETHAELHPSNKRGQTFVRFGKRSQSSVRFGN
>FLP-16_Acanthocheilonema_viteae_predicted
SSSNHKNFLCKFVSNLNFDFNFNFAAALKNNVEDTETHGGIHPLKRGQTFVRFGKRSQISN
>FLP-16_Elaeophora_elaphi_predicted
IWISISNFTASLKNNMMEFDTHVEPQPLKRGQTFVRFGKRSLNSARFGN
>FLP-16_Parascaris_equorum_predicted
TLKTTEHDLEAALENRLSAGDVYAASRPLDKRAQTFVRFGKXXXXXXXXXXXXXXXXXXXXXXXXXXX
XXXXXXXXXXXXXXXXXXXXXXXXXXFVAGKHPSWIQQI
>FLP-16_Parascaris_univalens_predicted
TLKTTEHDLEAALENRLSAGDVYAASRPLDKRAQTFVRFGKRAQTFVRFGRDACKQ
>FLP-16_Ancylostoma_caninum_ANCCAN_20772
MNSVELVLLAACTVVLFSLARSSPVSDQRLVEASPEEIERERELYQNLRQAMAESEDGPM
AKRAQTFVRFGKRAQTFVRFGKRAQTFVRFGRSVPEQQ
>FLP-16_Ancylostoma_ceylanicum_Acey_s0006.g2914.t1
MCVTFFLTILACRNPSLLIHSDASLTIRAICFRYNMNSVELVLLAACTVVLFSLARSSPV
SDQRLVEASPEDIERERELYQNLRQALAESDEGPMAKRAQTFVRFGKRAQTFVRFGKRAQ
TFVRFGRSVPEQQ
>FLP-16_Angiostrongylus_cantonensis ACAC_0000739401-mRNA-1
MTQIHFFTEYEPLAKRAQTFVRFGKRAQTFVRFGKRAQTFVRFGRSAPKHNTFFSAHS
>FLP-16_Angiostrongylus_costaricensis_ACOC_0000501101-mRNA-1
MVIGLLGLTIDELQTGSAELSPEDIEQEQEYFQNLRQVPVIGNHYYCQWCHDGKDKKKMA
RQLWSSPQYEPLAKRAQTFVRFGKRAQTFVRFGKRAQTFVRFGRSAPTQM
>FLP-16_Caenorhabditis_brenneri_CBN29085.1
MNFSGFELSSIIAVILIFLQLSSAAVVPVDYASQYGVAAADEMAALPEDGSLFAERPAKR
AQTFVRFGKRAQTFVRFGKRGQTFVRFGRSAPFEQ
>FLP-16_Caenorhabditis_briggsae_CBG02820a.1
MSLSGFEFSSIIAVLLLLIQLSSAAVLPVDYASQYGVASADEMTALPEEGSLFAERPAKR
AQTFVRFGKRAQTFVRFGKRGQTFVRFGRSAPFEQ
>FLP-16_Caenorhabditis_elegans_F15D4.8.1
MNFSGFEFSSIVAFFLLILQLSTAAVLPADYAYGVADEMSALPDSGSLFAEQRPSKRAQT
FVRFGKRAQTFVRFGKRGQTFVRFGRSAPFEQ
>FLP-16_Caenorhabditis_inopinata_Sp34_20323500.t1
MNFPGFEFSSIIAIFLLFLQLSSASVLPADYASQYSVADALALPEDSSLYAERPSKRAQT
FVRFGKRAQTFVRFGKRGQTFVRFGRSAPFEQ
>FLP-16_Caenorhabditis_japonica_CJA17583.1
MNFSGLELSSIVAVFLLLLQLSSAAVLPVDYSGVAEEMALPDDGSLFAERPAKRAQTFVR
FGKRAQTFVRFGKRGQTFVRFGRSAPLEQ
>FLP-16_Caenorhabditis_latens_FL83_10461
MNFSGFELSSIIAIFLLFVQLSSAAVLPVDYASQYGVASADEMTLPEEGSLFAERPSKRA
QTFVRFGKRAQTFVRFGKRGQTFVRFGRSAPLEQ
>FLP-16_Caenorhabditis_nigoni_Cni-.1
MKMSLSGFEFSSIIAVLLLLIQLSSAAVLPVDYASQYGVASADEMTALPEEGSLFAERPA
KRAQTFVRFGKRAQTFVRFGKRGQTFVRFGRSAPFEQ
>FLP-16_Caenorhabditis_remanei_CRE01713.1
MNFSGFELSSIIAVFLLFVQLSSAAVLPVDYASQYGVASADEMTLPEEGSLFAERPSKRA
QTFVRFGKRAQTFVRFGKRGQTFVRFGRSAPFEQ
>FLP-16_Caenorhabditis_tropicalis_Csp11.Scaffold629.g9757.t1
MNFSGFELSSIIAVFLLILQLSSAAVLPADYASQYGVASADEMALPEDGSLFAERPSKRA
QTFVRFGKRAQTFVRFGKRGQTFVRFGRSAPSYEEQ
>FLP-16_Dictyocaulus_viviparus_DICVIV_11320
MNGIELIVLITCFVTLFSVTDSTILKDQRLVELNPNYFERELDYLQNIQQQPELKSEDSP

MQKRAQTFVRFGKRAQTFVRFGKRAQTFVRFGRSTPKNY
>FLP-16_Diploscapter_coronatus_DCO_024376
MNSIQLLAAVLSCLLAVQIVVAAVAETQEKAYSNLASLSEPYGVYDIIVPRDTQRFLSQF
QESNLFDAPQKKAQSFVRFGKRSSDDSFYRPEKRAQSFVRFGRK
>FLP-16_Diploscapter_pachys_WR25_04404.1
MNSIQLLAAVLSCLLAVQIVVAAVAETQEKAYSNLASLSEPYGVYDIIVPRDTQRFLSQF
QESNLFDAPQKKAQSFVRFGKRSSDDSFYRPEKRAQSFVR
>FLP-16_Haemonchus_contortus_HCON_00035475
RYNMNGVELAFLATCAIVLLAFSNASPVNDQRLVEVSPEDIERERELLELLRQEMPAESD
DAPPSMAKRAQTFVRFGKRAQTFVRFGKRAQTFVRFGRSNPEQM
>FLP-16_Haemonchus_placei_HPLM_0000967401-mRNA-1
MHILLPIVSSPIPYDAPPSMAKRAQTFVRFGKRAQTFVRFGKRAQTFVRFGRSNPETSAL
RTYIISTNRMDNIGDLDSGNGPDPNLVVVQNPFGCTSTPP
>FLP-16_Heligmosomoides_polygyrus_HPOL_0000952401-mRNA-1
MNGVELALLATCVAVLLSISSASPSNDQRLVELSPEEVERERELYENLRQELAESDGMPP
AMAKRAQTFVRFGKRAQTFVRFGKRAQTFVRFGRSVPEQM
>FLP-16_Mesorhabditis_belari_mbelari.g1049.t1
MKFSIQILLFALIALIAVSTVTASPLFQSSGQGIVESEFGPADLEALEESMAPMEKRAQT
FVRFGKRAQPFVRFGKRAQTFVRFGRK
>FLP-16_Micoletzkya_japonica_MicoRS5524-mkr-S14-2.67-mRNA-1
MNLLSIILPFLCLVAASFATFLEQGEQQRLAPGQYAGYAEMMEEAPQKRAQTFVRFGKRA
QTFVRFGRSMPARYAMDQE
>FLP-16_Necator_americanus_NECAME_07417
MSSVELVLLVACTTVLFPLAHSSPTSDQRLYEGGPDDTERERELYQSLRQALAESYEPPI
EKRAQTFVRFGKRAQTFVRFGKRAQTFVRFG
>FLP-16_Nippostrongylus_brasiliensis_NBR_0001224201-mRNA-1
MGNFSRYTMNGVELVLLATCAAVLLSVSSASPVNEQRLVELNPEELERERELYENLRQEL
AQSEDIPPSMAKRAQTFVRFGKRAQTFVRFGKRAQTFVRFGRSAAEQQ
>FLP-16_Oscheius_tipulae_OTIPU.nOt.2.0.1.t10082
MNSSQLLAVVVCALLALVYSAPVPESLKSDDLAQYAELAEQERLLEEALREQGYPLYEVD
DVPQPMEKRAQTFVRFGKRSPMEKRAQTFVRFGKRAQTFVRFG
>FLP-16_Pristionchus_arcanus_arcanus-ag_msk-S_56-2.67-mRNA-1
MNFLATLIPLLCIIAACMATLMDTEQQQRQAQAPAGYGYEQYLEEAPVKRAQTFVRFGKR
AQTFVRFGRSMPARYQEEMDQ
>FLP-16_Pristionchus_entomophagus_entomophagus-mkr-S62-1.4-mRNA-1
DCRVTHTSGALPVMNFLATLIPLLCIIAACFATLMDSEQQQQQLRQPIGYGYEQYLDEAP
VKRAQTFVRFGKRAQTFVRFGRSAPRFEEDQ
>FLP-16_Pristionchus_exspectatus_exspectatus-ag_msk-S_163-0.14-
mRNA-1
MNFLATLIPLLCIIAACFATLMDTEQQSRQAQAPAGYGYEQYLEAQAPAGYGYEQYLEEA
PVKRAQTFVRFGKRAQTFVRFGRSMPARYQEEMDQ
>FLP-16_Pristionchus_fissidentatus_fissidentatus-mkr-S82-10.21-
mRNA-1
MNFLATLLPLLCLLAVCFASLVESEQQQPAAYSYDQFLEEAPVKRAQTFVRFGKRAQTFV
RFGRSLPARYETEE
>FLP-16_Pristionchus_japonicus_japonicus-mkr-S213-1.35-mRNA-1
MNFLATLLPLLCILAACFAAIADSAEQQQRPMQLPSGYGYEQYLEEAPQKRAQTFVRFGK
RAQTFVRFGRSMPARYQEEMDQ
>FLP-16_Pristionchus_maxplancki_maxplancki-mkr-S39-0.35-mRNA-1
MNFLATLIPLLCILAACLATLVDTEQQQRQMQAPAGYGYEQYLEEAPVKRAQTFVRFGKR
AQTFVRFGRSMPARYAQEEMDQ
>FLP-16_Pristionchus_mayeri_mayeri-mkr-S738-0.34-mRNA-1
MNFFATLIPLLCILAACFATLLETEQQQRQAPAGYRYEDYLEEAPVKRAQTFVRFGKRAQ
TFVRFGRSAPAARYDEYQ

>FLP-16_Pristionchus_pacificus_PPA35380.1
MNFLATLIPLLCIIAVCFATLIDIEQQSRQAQAPAGYGYEQYLEEAPVKRAQTFVRFGKR
AQTFVRFGRSMPVRYQEEMDQ
>FLP-16_Teladorsagia_circumcincta_TELCIR_04870
MYYDAEWEHDRRRGTPERDNRPNLNYNAPSIPPSSAKYRMMEGAVLVDCYSIRGSYNMNG
VELALLATCTIVLLSCSNASPVNDQRLVEVSPEEIERERELLALLRQEMPESDDTPPSKR
AQTFVRFGKRAQTFVRFGKRAQTFVRFGRSNPEQM
>FLP-16_Ancylostoma_duodenale_predicted
RMIVSDDGPMAKRAQTFVRFGKRAQTFVRFGKRAQTFVRFGRSVPEQQ
>FLP-16_Caenorhabditis_angaria_predicted
ILYCFSEKPQKRAQTFVRFGKRAQTFVRFGKRAQPFVRFVFNCMHEKLCCFRLG
>FLP-16_Caenorhabditis_sinica_predicted
RSIFPDYASQYGVAENEMGALPEEGSLFAERPAKRAQTFVRFGKRAQTFVRFGKRGQTFVRFGRSAPF
EQ
>FLP-16_Heterorhabditis_bacteriophora_predicted
TSASIQNFILNNVLFIEEPMLNKRAQTFVRFGKRAQTFVRFGKRAQTFVRFGRSNQEQM
>FLP-16_Parapristionchus_giblindavisi_predicted
SLIFAEAPAKRAQTFVRFGKRAQTFVRFGRSMPARYAVDEE
>FLP-16_Strongylus_vulgaris_predicted
KSRFLQISDESPISKRAQTFVRFGKRAQTFVRFGKRAQTFVRFG
>FLP-16_Bursaphelenchus_xylophilus_BXY_1577100.1
MNSQTLLMLCCTLALQFAYYVDAAAFLRQDKDKSAPAVVENANQQYISPGQEQVVESAPE
AELPAQKRAQTFVRFGKRAQTFVRFGKRAQTFVRFG
>FLP-16_Halicephalobus_mephisto_HMEPH_03960-RA.p1
MQSTWVILCLAAIAFNYASSAAVQQPGNNPDETLEQQQKSEEVEQQIREYVNQMREREQK
ALLQAMREKALYAAADPRDYYETVNPEEDDLQNPQKRAQSFVRFGKRAQTFVRFGKRAQT
FVRFG
>FLP-16_Panagrellus_redivivus_Pan_g4860.t1
MQVSTVLLATVVVAAVSALTDAGAALKTVDHELPAEKSAAVDHQDEIAAMEDMIRDYVNR
MREKQQLAMLKEMRDKNIMSAVGPREFYEQEVAPSGDDADKPEFNSSPEKRAQSFVRFGK
RGQSFVRFGKRAQDFVRFGKRAQSFVRFGKRAQDFDRFGKRAQDFVRFGKRGQSFVRFG
>FLP-16_Parastrongyloides_trichosuri_PTRK_0001641300.1
MDCRSIISWIFIALLSLSLVSTAALENQQESDLTYPDNIYGLYPIDDIKDVVPSKRAQTF
VRFGKRAQTFVRFGKRAQTFVRFG
>FLP-16_Rhabditophanes_sp.KR3021_RSKR_0000821000.1
MEGRSIISILIALILCIQMISAVSVHSPQEYELGYAADDNYGLYPVNGDNVETRVPSKRA
QTFVRFGKRAQTFVRFGKRAQTFVRFG
>FLP-16_Steinernema_carpocapsae_SC.X.g6657
MNVLIRIALVFISLSHVVFSSTIRNSMNQPNVAVAAANRELERLPSQEELLNQELQNIYL
AELMKQYAAENGVNTNQVEEPVNEQQAYGGMDKRAQTFVRFGRSSSVGASSVEEPVSQQQ
YGGMDKRAQTFVRFGKRGQTFIRFG
>FLP-16_Steinernema_feltiae_L889_g10804.t1
MNIVVRIALLFISLTHIVSSSALRNSINQPSVAMANHLSATDRLAATQEELNQELQNIYL
AELMKEYERSAAQNNVNTIQGDEQQQPLTEMNKRAQTFVRFGKRGQTFIRFG
>FLP-16_Steinernema_glaseri_L893_g33854.t1
MNIVIRILFVFVGLAFVGSSSAIRNSMNQLPQNTAHGRLSGSDMERFDVLNAIGSELQNL
QSILRAELSKPIRKEKSNQEKFIPVDDALMVNHADEKDKRAQTFVRFGKRGQTFIRFG
>FLP-16_Steinernema_monticolum_L898_g17790.t1
MNIVVRIALVFVSLTHIVSSSAIRNSINQPSVAMDTNRLSPDSDRLATQEELNQELQNFY
LTELMKQYERENTVNANQGEEPSMSNQQPLDMNKRAQTFVRFGKRGQTFIRFG
>FLP-16_Steinernema_scapterisci_L892_g28934.t1
MNVIVRIALVFISLSHVVSSSTIRNSMNQPGVAVAAAANREPEQSPSQEELLNQELQNIY
LAELMKQYAAAASENAVNNNQVEEPVSQQQVYGGMDKRAQTFVRFGKRGQTFIRFG
>FLP-16_Strongyloides_papillosus_SPAL_0001324400.1

MDRRSIFTWIITALLSLTLISAVTNENQQENELVYPDGIYEYYPVNDVPQALPSKRAQTF
VRFGKRAQTFVRFGKRAQTFVRFG
>FLP-16_Strongyloides_ratti_SRAE_X000053700
MDCRSIITWIITALLSLSLISSAAIENQQEDSDLVYPYNYYGYIPYNNYIQPSQSKRAQT
FVRFGKRAQTFVRFGKRAQTFVRFG
>FLP-16_Strongyloides_stercoralis_SSTP_0000617100.1
MDCRSIITWIITALLSLSLISPAAIENQQEESDLVYPYNYYGYIPYNNFIQPPQSKRAQT
FVRFGKRAQTFVRFGKRAQTFVRFG
>FLP-16_Strongyloides_venezuelensis_SVE_0689300.1
MDRRSIITWIITALLSLTLISAVTNENQQENELVYPDGIYEYYPVNDVPQTIPSKRAQTF
VRFGKRAQTFVRFGKRAQTFVRFG
>FLP-16_Acrobeloides_nanus_ACRNAN_scaffold5725.g32375.t1
MQATLFFFSALVIYGAELATAALLRPKYDLEKGSDNQQQILEDYPGYQQEIVELQNNPEL
QKAYYDVLSRVQPEIGADVRNNIPSDVPEKRAQTFVRFGKRAQTFVRFGKRAQTFVRFGR
DTNKQHAPVAPSEQ
>FLP-16_Ditylenchus_destructor_Dd_03376
MNPSEFLLILATSLSFLGWHTEALPARDNRPAKALIHGSNLQGEDPSPEIVPLIPEFRLY
DPQPQFFAEAPFPAVPQSSYQPRRDDVEPERIASPMDKRAQTFVRFGKRAQTFVRFGKRA
QTFVRFG
>FLP-16_Globodera_pallida_GPLIN_001364200
MQSLPLLSLLPVRPSNSQMTMSSSTSSSHCAPSSFSRKLFLWLLPATVTLFTTIHQSPFA
AASPFMPQKDDSVVVPMTSFGRPLPLSLVPTPPLYFVFPENVPLERPFDEQNDDGGSEEQ
FAEEAMGTKAKRAQTFVRFGKRAQTFVRFGKRAQTFVRLGRDTQRQFDGKMQSEQQQKKA
>FLP-16_Globodera_rostochiensis_GROS_g00582.t1
MSSSTSSHCVPSSFSRKLFLWLLPATVTLFTTIHQSPFAVASPFIPQKDDSVVVPMTSFG
QPLPPSPLSLVPNPPLYFVFPENLPLERPFDEQNDGSEEELAEEAMGTKAKRAQTFVRFG
KRAQTFVRFGKRAQTFVRLGRDTQRQFDGKMQSEQQQKKA
>FLP-16_Heterodera_glycines_Hetgly.G000016540
MTMFFSSSSSVCVPFSLGHKLFLFLLPILFAVQQFGMTEASPLIQQKDDSIPMPLVFDRP
WQPPISLVPNPPPAYFVIPANVPLERPIYDQNDSAIEDELAEEAKAKRAQTFVRFGKRAQ
TFVRFGKRAQTFVRFGRDAQRQQEMANEQAQKKE
>FLP-16_Meloidogyne_arenaria_M.Arenaria_Scaff1495g023845
MNLKEQQIYLNIQLLFFILAVISFLTTKGSEVKQRENNKLEYNKNEIERQKDQLIRDLIA
LTSERQYSRDWQQSQQQQNFLNSFGPSPHLFPSSGIEWPQQQQKIFLEGEVEEPLEENEK
EKRAQTFVRFGKRAQTFVRFGKRGQTFVRFGRDSKHQQNLSDQKQLKTDKQ
>FLP-16_Meloidogyne_enterolobii_scaffold681_cov388.g1207
MYLKEQQIYLNIQLLFFILAVTSFLTTKGSEVKQRENNKLEYDKNEIERQKEQLIRDLIA
LTRERQYSRDWQQLQQQQNFLKSFGPSPHLFPSSGFEWPQQQQKIFLEEGEVEEPENEKE
KRAQTFVRFGKRAQTFVRFGKRGQTFVRFGRGENLNLNLKLGAGHRRFKTSTKFVRSEAV
NKRQTIKNMANHEE
>FLP-16_Meloidogyne_floridensis_augustus_masked-nMf.1.1.scaf03158-
processed-gene-0.0-mRNA-1
MISHSMRPYEQLIRDLIASLTRERQYSRDWQQSQQQQNFINSFGPSPHLFPSSGIEWPQQ
QQKIFLEEGEVEEPLEENEKEKRAQTFVRFGKRAQTFVRFGKRGQTFVRFGRGEILKNLN
LKLGVGHRLTHP
>FLP-16_Meloidogyne_graminicola_NXFT01000413.1.10828_g
KKENKINEIERQKEQFIDDLIRQKLFSRQQNFLYNPYFNFELPKEQQQIEDKDYYFPLEG
KDKRAQTFVRFGKRGQTFVRFGKRAQTFVRFGKDLQHQQNLLREYKQQ
>FLP-16_Meloidogyne_hapla_MhA1_Contig381.frz3.gene4
MTLKEQIYLNIQLLFFILAISSFLAIKGSEIKQQGENKIENDKNEFERQKEQLIRDLIAL
TRERQISQDWQQLQQQQNFRNSFGPSPHLFPSSGFEWPQKQQQNLLQNYDDEEPLEEKDK
RAQTFVRFGKRAQTFVRFGKRGQTFVRFGRDLKHQQNLADQKQLKNEKQ
>FLP-16_Meloidogyne_incognita_scaffold46398_cov131.g17005
MNLKEQQIYLNIQLLFFILAVSSFLTTKGSEVKQRENNKLEYNKNEIERQKEQLIRDLIA

SLTRERQYSRDWQQSQQQQNFINSFGPSPHLFPSSGIEWPQQQQKIFLEEGEVEEPLEEN
EKEKRAQTFVRFGKRAQTFVRFGKRGQTFVRFGRDSKHQHNLSDQKQLKTDKQ
>FLP-16_Meloidogyne_javanica_M.Javanica_Scaff4438g034603
MNLKEQQIYLNIQLLFFILAVSSFLTTKGSEVKQRENNKLEYDKNEIERQKEQLIRDLIA
SLTRERQYSRDWQQSQQQQNFLNSFGPSPHLFPSSGIEWPQQQQKIFLEGEVEEPLEENE
KEKRAQTFVRFGKRAQTFVRFGKRGQTFVRFGRDSKHQQNLSDQKQLKTDKQ
>FLP-16_Ditylenchus_dipsaci_predicted
RPSRPTPQPLQKRAQTFVRFGKRAQTFVRFGKRAQTFVRFG
>FLP-17_Plectus_sambesii_PSAMB.scaffold2694size21766.g18760.t1
peptide: PSAMB.scaffold2694size21766.g18760.t1
MSNHFFLLTISSLCLQLCCDPALARPQWQSTIADVQRSIDAARDDMECFRTVGESCQDAE
LGLTGQCPCDLGLQCVYNVCVPVYTNNEPSNVEGEYLSVRENRTSSSACPTMYGRLIEVA
FVALSCAQIVLSQDAAAAAVAPEQKTDVKEQEMAELFCQQYNHLNLCKLRETLEGALVEI
QYLLEGDDNASESPLGTSMEKRKSAFVRFGKRKSAFVRFGKRKSAFVRFGRSLEEAENKR
KSSYIRFG
>FLP-17_Ascaris_lumbricoides_ALUE_0000906501-mRNA-1 peptide:
ALUE_0000906501-mRNA-1
MGGVTVIVLFGFVTAICANEIPASLGEEMCKEYPDLHLCRLQSTLEGALAEIQYLVNGDT
VTSTSSTANKRKSAFVRFGKRVDEEEDNEKTGEGVDENVVKEEKREWHCEITEQIQVEGD
NKRNKG
>FLP-17_Ascaris_suum_GS_19665 peptide: GS_19665
MGGVTVIVLFGFVTAICANEIPASLGEEFCKEYPDLHLCRLQSTLEGALAEIQYLVNGDT
VTSTSSTANKRKSAFVRFGKRVDEEEDNEKEIQKRKSAFVRFGRSDSDKRKSSYIRFG
>FLP-17_Parascaris_univalens_PgR144_g014_t01 peptide:
PgR144_g014_t01
MGGVMVIVLFGFVTAICANEIPASLGEEFCKEYPDLHLCRLQSTLEGALAEIQYLVNGET
VASTSGTVSKRKSAFVRFGKRVDEEEDNEKRSIPNSVYATALAVQEVYCKEAILAELDFE
GVELE
>FLP-17_Toxocara_canis_Tcan_05767.1 peptide: Tcan_05767.1
MSTSIGEQFCKEYAHLHLCRLQDNLEGALAEIQYLVNSDVAAPVTSVINKRKSAFVRFGK
RPNNEEDDEKEVEKRKSAFVRFGRSGSEKRKSSYIRFG
>FLP-17_Anisakis_simplex predicted
KLRSFTDCFKKHDFSNKRRLLIKRFSVNFEMNVGYCIRLQEVEKRKSAFVRFGRSDNDKRKSSYIRFG
>FLP-17_Dracunculus_medinensis predicted
EIDKRKSAFVRFGRSKFNEKGKSSYIRFGRK
>FLP-17_Gongylonema_pulchrum predicted
SQPFIPVRSFGQLTVHIKRLAQSFLIFSVPAPEINKRKSAFVRFGKRNEPAEVSQQFLEQKKFFCL
>FLP-17_Parascaris_equorum predicted
LKRMVSIKKRRWFTVFSYLIGLVASTSGTVSKRKSAFVRFGKRVDEEEDNEKVTDFT
>FLP-17_Ancylostoma_caninum_ANCCAN_20066 peptide: ANCCAN_20066
MWCVYFFISLIVCSFAANEDNLSEEFCRQFPSLHLCRLHDNLQGSLVELQYLLQDNNIEI
GNPSAATNPMEKRKSAFVRFGKRAADDAMEVEKRKSAFVRFGRSVPIETPDKRKSQYIRF
GRK
>FLP-17_Ancylostoma_ceylanicum_Acey_s0050.g1896.t2 peptide:
Acey_s0050.g1896.t2
MWCVYFFFSLIVCSFAANEDNLSEEFCRQFPSLHLCRLHDNLQGSLVELQYLLQDSNIEI
AAPVSPVEKRKSAFVRFGKRAADDAMEVEKRKSAFVRFGRSAPIETPEKRKSQYIRFGRK
>FLP-17_Ancylostoma_duodenale_ANCDUO_18171 peptide: ANCDUO_18171
MWCVYFFLSLIVCSFAANEDNLSEEFCRQFPSLHLCRLHDNLQGSLVELQYLLQDNNIEI
GNPSAATNPMEKRKSAFVRFGKRAADDAMEVEKRKSAFVRFGRSVPIETPDKRKSQYIRF
GRK
>FLP-17_Angiostrongylus_cantoniensis_Angca_000478 peptide:
Angca_000478
MYILINVFFNLVMWCALFFVSLAISSIVASANDEHFTQEFCRHFPSLNLCRLRGTLQGSL

LELQYLLQNVSPDNDPNSDNAVDTSPMEKRKSAFVRFGKRS
>FLP-17_Angiostrongylus_costaricensis_ACOC_0000969901-mRNA-1
peptide: ACOC_0000969901-mRNA-1
MWCALFFVSLAISSIVASASDEHPTQEFCRYFPSLNLCRLREALQGSLLELQYLLQDVSP
DNGPNSNNALDTSPMEKHKSAFVRFGKRSFEDASVS
>FLP-17_Caenorhabditis_angaria_Cang_2012_03_13_00517.g11771.t1
peptide: Cang_2012_03_13_00517.g11771.t1
MIAKAIILTVLLSTTIAAATSEEQQANSFCEKFPTLHMCRLRDELTGSLVELQYLLHDNG
EGLPAMPQNEMQKRKSAFVRFGKRSSDDSAAATSAEDETLEMEKRKSAFVRFGRAGPQEK
RKSQYIRFGK
>FLP-17_Caenorhabditis_brenneri_CBN02195.1 peptide: CBN02195.1
MLSKLVLIACCLLISSSNGASMEEIQSEKFCEKFPTLHMCRLKEELTGSLVELQYLLQDG
MNVGGQQQAGVQEVQKRKSAFVRFGKRSAPEEEGMEMEKRKSAFVRFGRSVGMEPQLTEK
RKSQYIRFGK
>FLP-17_Caenorhabditis_briggsae_CBG00359.1 peptide: CBG00359.1
MLFKLVFIALLFASSYGASMEEIQSEKFCEKFPTLHMCRLKEELTGSLVELQYLLQDGIN
VGGQQQAGGVQEVQKRKSAFVRFGKRSASDEEGMEMEKRKSAFVRFGRSIGMEPQFTEKR
KSQYIRFGK
>FLP-17_Caenorhabditis_inopinata_Sp34__40204200.t1 peptide:
Sp34_40204200.t1
MLSKLVLITCLLLASSSNGASMEEIQSEKFCEKFPTLHMCRLKEELTGSLVELQYLLQDG
MSVGGQSQVEAQEVQKRKSAFVRFGKRSASEEEGMEMEKRKSAFVRFGRSAGLEPQITEK
RKSQYIRFGK
>FLP-17_Caenorhabditis_latens_FL83_23278 peptide: FL83_23278
MLSKLVFITFLLFTSSFGASMEEIQSEKFCEKFPTLHMCRLKEELTGSLVELQYLLQDGI
NVGGQQQQVGAQEVQKRKSAFVRFGKRSAPEEEGMEMEKRKSAFVRFGRSLGMEPQLTEK
RKSQYIRFGK
>FLP-17_Caenorhabditis_nigoni_Cni-flp-17 peptide: Cni-flp-17
MLFKLVLIALLFASSYGASMEEIQSEKFCEKFPTLHMCRLKEELTGSLVELQYLLQDGIN
VGGQQQAGGVQEVQKRKSAFVRFGKRSAPEEEGMEMEKRKSAFVRFGRSIGMEPQFTEKR
KSQYIRFGK
>FLP-17_Caenorhabditis_remanei_CRE21839.1 peptide: CRE21839.1
MLSKLVFITFLLFTSSFGASMEEIQSEKFCEKFPTLHMCRLKEELTGSLVELQYLLQDGI
NVGGQQQQVGAQEVQKRKSAFVRFGKRSAPEEEGMEMEKRKSAFVRFGRSLGMEPQLTEK
RKSQYIRFGK
>FLP-17_Caenorhabditis_sinica_Csp5_scaffold_03212.g29437.t1 peptide:
Csp5_scaffold_03212.g29437.t1
MLFKLVFLTCCLLFAGSQGASMEEIQSEKFCEKFPTLHMCRLKEELTGSLVELQYLLQDG
INVGGQQQAGVQEVQKRKSAFVRFGKRSAPEEEGMEMEKRKSAFVRFGRSLGMEPQLTEK
RKSQYIRFGK
>FLP-17_Caenorhabditis_tropicalis_Csp11.Scaffold171.g648.t1 peptide:
Csp11.Scaffold171.g648.t1
MLVKLVLIACCLLFSSSNGASMEEIQSEKFCEKFPTLHMCRLKEELTGSLVELQYLLQDG
INVGGQQQPAGAQEVQKRKSAFVRFGKRSAPEEEGMEMEKRKSAFVRFGRSIGMEPQLTE
KRKSQYIRFGK
>FLP-17_Dictyocaulus_viviparus_DICVIV_07526 peptide: DICVIV_07526
MWCALLFFSSIVYSIAASTNNERSSLEVCRQVPSLYLCRLYGTLQESLYELQDLLQDTEP
ENAAVTNLMHKRKSAFVRFGKRSVDEVPEPEKRKSAFVRFGRSVNIPEKRKSQYIRFGRK
>FLP-17_Diploscapter_coronatus_DCO_022899 peptide: DCO_022899
MAQIWHSILPLFALFGLGCSAIHDRLVADQFCGKFPTLHMCRLRTELEGSLGELQYLLQD
ATNDVQSAEMSGTTSSPDLAGQPEKRKSTYVRFGKRSASDLLEDAPVKRKSTYVRFGKRS
ANEFDDATSELQMEKRKSAFVRFG
>FLP-17_Diploscapter_pachys_WR25_20045.1 peptide: WR25_20045.1
MAQIWHSILPLFALFGLGCSAIHDRLVADQFCGKFPTLHMCRLRTELEGSLGELQYLLQD

ATNDVQSAEMSGTTSSPDLAGQPEKRKSTYVRFGKRSASDLLEDAPVKRKSTYVRFGKRS
ANEFDDATSELQMEKRKSAFVRFG
>FLP-17_Haemonchus_contortus_HCON_00123460-00001 peptide:
HCON_00123460-00001
MWCVLFVFSLIACSFASSRDDQLSEQFCRQFPSLNLCRLHDTLQGSLVELQYLLQDTNLD
SAAPGSSAEKRKSAFVRFGKRAAEESAEIEKRKSAFVRFGRSAEFDMPEKRKSQYIRFGR
K
>FLP-17_Haemonchus_placei_HPLM_0000820801-mRNA-1 peptide:
HPLM_0000820801-mRNA-1
MWCVLFVFSLIACSFASSRDDQLSEQFCRQFPSLNLCRLHDTLQGSLVELQYLLQDTNLD
SAAPGSSAEKRKSAFVRFGKRAAEESAEIEKRKSAFVRFGRSAEFDMPEKRKSQYIRFGR
K
>FLP-17_Heligmosomoides_polygyrus_HPOL_0000658301-mRNA-1 peptide:
HPOL_0000658301-mRNA-1
MWCVLFVLSLIVCSSLAGNEEHLSEQFCRQFPSLHLCRLHDTLQGSLVELQYLLQDTNAE
GAVPLNQMEKRKSAFVRFGKRSADDTMEVDKRKSAFVRFGRSASSMEMPDKRKSQYIRFG
RK
>FLP-17_Heterorhabditis_bacteriophora_Hba_02546 peptide: Hba_02546
MLHYVMLFVFFALSIIVCTFAGNEEILSEEFCRQFPSLHLCRLHDTLQGSLIELQYLLQD
TNVESAIAVTSAPLDKRKSAFVRFGKRSADDIVDMEKRKSAFVRFGRSVADIPDKRKSQY
IRFGKK
>FLP-17_Mesorhabditis_belari_mbelari.g18803.t1 peptide:
mbelari.g18803.t1
MKRSVGSFVIGMLISLLFSPLQADEQSDFCRQFPSLALCKLPTILQGDSPYTAPGSYDEL
QYLLAGVDQPQQAVGHEMITNQALQKRKSAFVRFGKRSDTQIGDTEIPEKRKSAFVRFGR
SDPAMVAEMEKRKSNYVRFG
>FLP-17_Necator_americanus_NECAME_13924 peptide: NECAME_13924
MWCVFFFLSLIVCSFATNEDNLSEEFCRQFPSLHLCRLHDTLQGSLVELQYLLQDNNVEN
AVPVNPMEKRKSAFVRFGKRAANDAAEIEKRKSAFVRFGRSVPVDVPEKRKSQYIRFGRK
>FLP-17_Nippostrongylus_brasiliensis_NBR_0001430501-mRNA-1 peptide:
NBR_0001430501-mRNA-1
MWCAVFFLSLIACTFAGNEEHLAEQFCPAPMMEKRKSAFVRFGKRSDEDVMDVDKRKSAF
VRFGRSAPLDMPEKRKSQYIRFGRK
>FLP-17_Oesophagostomum_dentatum_OESDEN_09593 peptide: OESDEN_09593
MAGTHCTDFYGSEVSAAIPVGPVEKRKSAFVRFGKRAADFSGEVEKRKSAFVRFGRSVPA
ELPEKRKSQYIRFGRK
>FLP-17_Oscheius_tipulae_OTIPU.nOt.2.0.1.t10567 peptide:
OTIPU.nOt.2.0.1.t10567
MLSAIFLVFCLFTYTHAGTNDEVLSEEFCRQFPSLHLCRLHDTLQGSLVELQYLLQDTEG
MTGAVAPGTVQKRKSAFVRFGKRSTETSEENSDSMSMEKRKSAFVRFGRSIGENPDKRKS
QYIRFGRK
>FLP-17_Parapristionchus_giblindavisi_Parapristionchus-mkr-
S_3171-0.5-mRNA-1 peptide: Parapristionchus-mkr-S_3171-0.5-mRNA-1
MFIRYLILLIVISLTTAEDVSQQTLPSPVDEFCSRYPFLNLCQLRSTLEGSLMELAQIQS
HLVVDDSQDAPEDLQMEKRKSTFVRFGKRSAAPATSPEEEPSKMEEKRKSTFVRFGKRKS
SYVRFG
>FLP-17_Pristionchus_arcanus_PARCANUS000017717.t1 peptide:
PARCANUS000017717.t1
QGPIMFTRVAITILCTLSALMAADTEQQTNPSIPETFCERFPALALCQLRNTLDQSILEL
SSLQEHMLKEAADIRTPGAIEKRKSNFVRFGKRSVEPVEMGQMEVPEKVEEKRKSNFVRF
GKRKSNFVRFG
>FLP-17_Pristionchus_entomophagus_PRENTOPHAG000009985.t1 peptide:
PRENTOPHAG000009985.t1
AHSCRRFHKTSKHSSCCVRARDPSDSAHRTPTVYPSQGPTTMHTRAAIALLISVSAVVAA

DAEQQTAPSIPEVFCDRFPALALCQLRYNLDQSILELSTLQEQMIKDAVDIRPPEAIEKR
KSNFVRFGKRSAEPIEVGEMEEPEKVEQKRKSNFVRFGKRKSNFVRFG
>FLP-17_Pristionchus_expectatus_PEXSPEC000008918.t1 peptide: PEXSPEC000008918.t1
MFTRVTTTILFTLSALMAADTKQQTNPSIPEAFCERFPGLALCQLRYNLDQSIHELSSLQ
EHMLKQVADIRPPEAIEKRKSNFVRFGKRSAEPVEMGQMEVPEKVEEKRKSNFVRFGKRK
SNFVRFG
>FLP-17_Pristionchus_fissidentatus_PRIFISSI000006270.t1 peptide: PRIFISSI000006270.t1
KIPEAFCARFPALALCQLRFNLDQAIVELSTVQEQMLSDAADVRAPVAMEKRKSNFVRFG
KRSESPVEPGELEEPEKVEEKRKSNFVRFGKRKSNFVRFG
>FLP-17_Pristionchus_japonicus_japonicus-mkr-S681-0.28-mRNA-1 peptide: japonicus-mkr-S681-0.28-mRNA-1
MSDSLPGSPLLLLFRLLVPNFFFHFSGIRSLLPPKRLPNTRFSWCALVICRTNPDVHFVL
LYFVLHLASASCLILVQIKRVSQDPTMLVRISIALLFTLTAVFAADAEQQTNPSIPEAFC
ERFPALALCQLRYNLDQSILELSTLQEQMLKEAAAVQQPEAIQKRKSNFVRFGKRSVEPV
KIGQMEEPEKVEEKRKSNFVRFGKRKSNFVRFG
>FLP-17_maxplancki-ag_msk-S170-0.7-mRNA-1 peptide: maxplancki-ag_msk-S170-0.7-mRNA-1
MFLMAFLMATFLMAAEAEQQTNPSIPEAFCERFPALALCQLRYNLDQSILELATLQEQML
KEAVDIRPPEAIEKRKSNFVRFGKRSAEPLELGQLEEPEKVEEKRKSNFVRFGKRKSNFV
RFG
>FLP-17_Pristionchus_mayeri_mayeri-mkr-S83-1.10-mRNA-1 peptide: mayeri-mkr-S83-1.10-mRNA-1
FQGPSMLIRVATALLFAIPALMAADSEQQTLPSIPEAFCARFPALALCQLRFDLDQAIVE
LSAIQSQMLKDSADIQPPEAIAKRKSNFVRFGKRSAPEMGEEEEEPEKVEEKRKSNFVRF
GKRKSNFVRFG
>FLP-17_Pristionchus_pacificus_PPA27516.1 peptide: PPA27516.1
MFTRATTTILFTLSALMAADTEQQTNPSIPEAFCERFPALALCQLRYNLDQSIHELSSLQ
EHMLKEVADIRPPEAIEKRKSNFVRFGKRSAEPVEMGQMEMPEKVEEKRKSNFVRFGKRK
SNFVRFG
>FLP-17_Teladorsagia_circumcincta_TELCIR_11623 peptide: TELCIR_11623
MDGAHSRRVDTVSANIIVAGDSGITAQKRKSAFVRFGKRSIDEAGDVEKRKSAFVRFGRS
APFDMPEKRK
>FLP-17_C52D10.11.1 peptide: C52D10.11.1
MLSKLVLTTCLLLTISGSSQAASMEEIQSEKFCEKFPTLHMCRLKEELTGSLVELQYLLQ
DGINNQQQAGAQEVQKRKSAFVRFGKRSAPEEEAMEMEKRKSAFVRFGRSFGMEPQITEK
RKSQYIRFGK
>FLP-17_Caenorhabditis_japonica FLP-17 predicted
KSAFVRFGRSIGLEPQITEKRKSQYIRFGRK
KRKSAFVRFGKR
>FLP-17_Micoletzkya_japonica_Micoletzkya_japonica FLP-17 predicted
LDSGEPRVDVSDASASATGAGEVEKRKSSFVRFGKRAAADLDDDAIVVR
>FLP-17_Bursaphelenchus_xylophilus_BXY_0707100.1 peptide: BXY_0707100.1
MLGLIIVAGLCSQLAFAFLTPQQNAKLSELCSANPNLFVCTLAEYLDESMIDIDSLIPAE
QRDSLQNPAKNFIGSRMQKRKSAFVRFGKRSQGNSEYAPEDLALNFPIDGILNSRPARKS
SYIRFG
>FLP-17_Halicephalobus_mephisto_MSTRG.8673.1.p1 peptide: MSTRG.8673.1.p1
MNVIVVLSFIVAMSVVNANPVTKEDLCSGMFKDLAICKIRGLTEQAVKETEQILSEYYEI
LSRGNFFEKMDQNPTAVNNQKIPFEKRKSAFVRFGKRSEENSNLNSESPRSADLPNLPWF
LSMNFYSNGPRSGSNENHPEISEIEKRKSSYVRFGKREM
>FLP-17_Panagrellus_redivivus_Pan_g17589.t1 peptide: Pan_g17589.t1

MMKLFLLALCFGVASAKQPLDDICTENFADLNICKLRGVLARAYDELNEIIGTYQDMAYA
YHSGESSKDSAEANFLFEKVQSDRINALRTQRAHVARGPMEKRKSAFVRFGKRSELAEPI
AQIFPSNPNAASDNVPEAREVAKRKSSYVRFG
>FLP-17_Parastrongyloides_trichosuri_PTRK_0000745500.1 peptide:
PTRK_0000745500.1
MFKFYSTIFIAFLLVAASKVVLTQSSPTMDNHGIVENTPDQASAYEMFCRDYSHLQLCKL
ELTLQQALAELQYIILNDDPSEEIESFRSKRKSAFVRFGKRSSDDDMFQNFDKRKSAFVR
FGRSIEDPINGQKRKSSYVRFG
>FLP-17_Rhabditophanes.sp.KR3021_RSKR_0000181000.1 peptide:
RSKR_0000181000.1
MATD*TNDDWDRRHNSFEFQALDTKPLIKKADLLGGHLKVLVPKVEPPYLNYVPIDINQI
NEDKVPPGVVVMILSYVGKELNLTYDFEILETGEWGTMADGKWNGAFEKLIDGDAELVAG
GAVVRYDRSLYTDLTFPIHFEVSGMMIRSSNKYRSYTWLIVTDPFSWKVWMIAFCSIIVS
GIYFKINTYFLKKINKETQYSIFECTWIFFSIAMQQALPRQPINTSCRTLLAFWWLSSMT
LLATFTGSLVALFAVNSLVLPFSDFDGMVQSIKSGKYTLLMDINYPSKTEMIAKSRIPSY
KDLWNEMYNNHRVDYVESIKTGVQIVKNNPNFVLLGPKEVLNIYAAIECHMVILKEEILA
SYISIPFKKHSKYAQYFSERIKKYTEYGFIDKWLKDYKTYILSQNENRCNITQTQSKSKV
SLGMEKAQGAFYVYVSGFLISVVVLIVEMVFNVVLVMWTKYKKMAKFYIYLAVIALAIFV
NNGSVAAQSPEDAELVESVPNKLEGFSKFCKMYSHLQLCKIEHNLEVALAELYYIVYGEV
APGSVMLDSVPAPRKRKSAFVRFGKRSLPEDNQEMAFEKRKSAFVRFGRSMDEYFNNGQK
RKSSYVRFG
>FLP-17_Steinernema_carpocapsae_L596_g10349.t1 peptide:
L596_g10349.t1
MWAFLIFATLVALARTESDVAGSAAVVDEFCQQYNQMSLCHLHGTLEQALTELSFLFGED
AASGGGQTIPMMGKRKSAFVRFGKRSADELEQAMEKRKSAFVRFGRSLQPVEQKRKSSYV
RFG
>FLP-17_contig03713.0.2 peptide: contig03713.0.2
MWAIFVFAIFVALSRADDGAVAGNAAVVDEFCQRYSQMSLCHLHGTLEQALTELSFLFGE
DAAPDGSEPLVMGKRKSAFVRFGKRAVSDLEQAMEKRKSAFVRFGRSLQPVEQKRKSSYV
RFG
>FLP-17_Steinernema_glaseri_L893_g30784.t1 peptide: L893_g30784.t1
MLGFVVIATLLALSNAESEPGAAAVVDEFCQRYSQMSLCHLHGTLQQALTELSFLFGEDG
VPTGPNDPILMSKRKSAFVRFGKRAADEGDQVMEKRKSAFVRFGRSLPAVEQKRKNSYVR
FD
>FLP-17_Steinernema_monticolum_L898_g3546.t1 peptide: L898_g3546.t1
MWALFVFAIVVALVRAEEGEPRNSAVVEEFCQRYNQMSLCHLHGTLEQALTELSFLFGED
VNAEGGDPIVMGKRKSAFVRFGKRAAEDVEQAMEKRKSAFVRFGRSLQPVEEKRKSSYVR
FG
>FLP-17_Steinernema_scapterisci_L892_g11917.t1 peptide:
L892_g11917.t1
MWAFVIFATMVVLARSESDVAGSAAVVDEFCQRYNQMSLCHLHGTLEQALTELSFLFGED
ANPATGGGDPIVMGKRKSAFVRFGKRSADELEQAMEKRKSAFVRFGRSLQPMEQKRKSSY
VRFG
>FLP-17_Strongyloides_papillosus_SPAL_0000071600.1 peptide:
SPAL_0000071600.1
MFKYQSCLATVLFLILSISVTLTFSSPTLERHLPSEVTSDQFSAYEVFCREYSHLQLCKL
ESTLQQALAELQYIILNDEPNEESENFRSKRKSAFVRFGKRSPSDDMNFEKRKSAFVRFG
RSIDDSISGQKRKSSYVRFG
>FLP-17_Strongyloides_ratti_SRAE_X000019700.1 peptide:
SRAE_X000019700.1
MFKYPTSLITVFLFFLSIGITFTLSSPTIGSQHVSDISPDQVSAYEMFCKDYSHLQLCKL
EFTLQQALAELQYIILNDDPTDDNQNIKSKRKSAFVRFGKRDLGDEIAFDKRKSAFVRFG
RSIEDSISGQKRKSSYVRFG
>FLP-17_Strongyloides_stercoralis_SSTP_0000486200.1 peptide:

SSTP_0000486200.1
MFKYPTSFITVLFYSLSIVVSLTLSLPAMESHHAPNPTSDQISAYEMFCKDYSHLQLCKL
EFTLQQALAELQYIILNDDPIDDSENFKTKRKSAFVRFGKRSNDDDMLFDKRKSAFVRFG
RSVEDPINGQKRKSSYVRFG
>FLP-17_Strongyloides_venezuelensis_SVE_0907600.1 peptide:
SVE_0907600.1
MLEKIIIYNNNFNLVMMFKYQSYLVTVLFLILSISVTLTFSSPTLERHSPSEVTSDQFSA
YEVFCREYSHLQLCKLESTLQQALAELQYIILNDEPNEESENFRFKRKSAFVRFGKRSPS
DDMNFEKRKSAFVRFGRSIDDSISGQKRKSSYVRFG
>FLP-17_Acrobeloides_nanus_ACRNAN_scaffold2729.g19047.t1 peptide:
ACRNAN_scaffold2729.g19047.t1
MYAPVFALMLFVAYSTAIPTDTNEFCSVYSHLNLCKLKEVLEGALQEVQFVLSAQTAIPP
QVQARTMVDKRKSAFVRFGKRSPASEESFSMP
>FLP-18_Romanomermis_culicivorax_nRc.2.0.1.t24635-RA peptide:
nRc.2.0.1.t24635-RA
MIDNSMQTVPGEQYYNMASNFKASISESLTISTALTIFYGVVPCAFWVYGVLSYHLNYNS
NNLPKIVIFFYVALRRVLEILVLARIWQIKCYTLLSLIVYFLFMCNRSFAARKTSDSLSN
RSDGVNDGDESYEDENDDETYEEKRALPGVLRFGKKAMPGVMRFGKRWSGGAGAALSPYD
GDDDDEAFYNAYKQRWASRIRKAMPGVMRFGKRSAVKRAGGAIAIGGKHSMPRVMRFG
>FLP-18_Soboliphyme_baturiniSoboliphyme_baturiniSoboliphyme_baturini
_SBAD_0001137701-mRNA-1 peptide: SBAD_0001137701-mRNA-1
MFDKCLFVGLRQNSQRFIDPLDSFRQGEAMTPNWKARFCGKVTKHDRSAESTLIRGNDFD
KYYGIAPMAGFKRDSPGLMRFGKRASMMQYEKSQLPGMMRFGRNYVPDTYSDYLNGEALT
NTE
>FLP-18_Trichinella_britovi_T03_13022.1 peptide: T03_13022.1
MKSYISLQSIMFVLGVVFLCSISFGSCTEADMDNFKENIPGRFLWKKYDAPGLMRFGKRV
VQGYDRYDDSAPGLMRFGKRSYSRQLHDKFGDSDLRLRYVT
>FLP-18_Trichinella.murrelli_T05_10777.1 peptide: T05_10777.1
MKSYISLQSIMFVVGVVFLCSISFGSCTEADMDNFKENVLYEDIPGRFLWKKYDAPGLMR
FGKRVVQGYDRYDDSAPGLMRFGKRSYSRQLHDKFGDSDLRLRYVT
>FLP-18_Trichinella_nativa_D917_09615 peptide: D917_09615
MFVVGVVFLCSISFGSCTEADMDNFKENIPGRFLWKKYDAPGLMRFGKRVVQGYDRYDDS
APGLMRFGKRSYSRQLHDKFGDSDLRLRYVT
>FLP-18_Trichinella_nelsoni_T07_7357.1 peptide: T07_7357.1
MFVLGVIFLCSISFGSCTEADMDNFKENVLYEDIPGRFLWKKYDAPGLMRFGKRVVQGYD
RYDDSAPGLMRFGKRSYSRQLHDKFGDSDLRLRYLT
>FLP-18_Trichinella_papuae_T10_4261.1 peptide: T10_4261.1
MFVLGVVFLCSISFGSCTEADMENFKENVPGRFLWKKYDAPGLMRFGKRIVQGYDRYDDS
APGLMRFGKRSYSRQLHDKFGDSDLRLRYVT
>FLP-18_Trichinella_patagoniensis_T12_3663.1 peptide: T12_3663.1
MKSYISLQSIMFVLGVVFLCSISFGSCTEADMDNFKENVLYEDIPGRFLWKKYDAPGLMR
FGKRVVQGYDRYDDSAPGLMRFGKRSYSRQLHDKFGDSDLRLRYVT
>FLP-18_Trichinella_pseudospiralis_T4C_274.2 peptide: T4C_274.2
MKSYISLQSIMLVLGVVFLCSISFGSCTEADMENFKENVPGRFLWKKYDAPGLMRFGKRI
VQGYDRYDDSAPGLMRFGKRSYSRQLHDKFGDSDLRLRYVT
>FLP-18_Tricinella_spiralis_T01_6784.1 peptide: T01_6784.1
MKSYISLQSIMFVLGVVFLCSISFGSCTEADMDNFKENIPGRFLWKKYDAPGLMRFGKRV
VQGYDRYDDSAPGLMRFGKRSYSRQLHDKFGDSDLRLRYVT
>FLP-18_Trichinella_sp.T6_T06_12727.1 peptide: T06_12727.1
MKSYISLQSIMFVVGVVFLCSISFGSCTEADMDNFKENVLYEDIPGRFLWKKYDAPGLMR
FGKRVVQGYDRYDDSAPGLMRFGKRSYSRQLHDKFGDSDLRLRYVT
>FLP-18_Trichinella_sp.T8_T08_16448.1 peptide: T08_16448.1
MKSYISLQSIMFVLGVVFLCSISFGSCTEADMDNFKENIPGRFLWKKYDAPGLMRFGKRV
VQGYDRYDDSAPGLMRFGKRSYSRQLHDKFGDSDLRLRYVT

```
>FLP-18_Trichinella_sp.T9_T09_5865.1 peptide: T09_5865.1
MKSYISLQSIMFVVGVVFLCSISFGSCTEADMDNFKENVLYEDIPGRFLWKKYDAPGLMR
FGKRVVQGYDRYDDSAPGLMRFGKRSYSRQLHDKFGDSDLRLRYVT
>FLP-18_Trichinella_zimbabwensis_T11_18332.1 peptide: T11_18332.1
MKSYISLQSIMFVLGVVFLCSISFGSCTEADMENFKENVPGRFLWKKYDAPGLMRFGKRI
VQGYDRYDDSAPGLMRFGKRSYSRQLHDKFGDSDLRLRYVT
>FLP-18_Trichuris_muris_TMUE_2000008037.1 peptide: TMUE_2000008037.1
MDVMIVMVPPGDGPLAAEGGTRLTIAAEERKTQLIISARSTHWLTVKVLNPRNESSLDFR
SPQRGIGADEYASKFPHFLYKSNHIANDVPGLMRFGKKSVVPSEQLEVLDGSLMA
>FLP-18_Trichuris_suis_D918_07269 peptide: D918_07269
MRFRSASRDVENPVVSSCTCGVVAVGIQSAFTALNLYYSSQIIGYKVQKMKMHSIFGSFK
LLASVVLFGSVLLAAETDGQLVLPENVMEANAQYGEPVLRNQWKKYDGPGMMRFGKRRYY
NAHFAKNGIRYGGLEDILQPNQMYSRINGPITVEPIMSFQETIYRSRK
>FLP-18_Plectus_sambesii_PSAMB.scaffold3601size17630.g22026.t1
peptide: PSAMB.scaffold3601size17630.g22026.t1
MSAATDRAGAFLPFAQSPALFNLSAALIKRSVPHSIHLIPCPFHGVNRERDVRRRRRTVG
DGDDRLTRLSQLQQAKVPSGSPKAAVFLRGLQSMLHRVETMQLSNPFCRFLLVLCSLIAA
SRAAEIEDQEAAEKRAGGFLTANQVLPGNYFGKRVGMIDRLPTSEDYEWLMSAGKRDVEE
AENDSSKRSAEDNTIPGVLRFGKKSNSELPSGLRFGKRAQSFIRFGKRQDKKDMPGVLRF
GKRDDGAMPGVLRFGKKSDIPGVLRFGKRDDGAMPGVLRFGKKSDIPGVLRFGKRGDSDM
PGVLRFGKRDMPGVLRFGKRDDIPGVLRFGRR
>FLP-18_Acanthocheilonema_viteae_nAv.1.0.1.t00368-RA peptide:
nAv.1.0.1.t00368-RA
METAGLMMHMVKFAVIFVTIMTTTTTTTTIPLYVQMDNSSILSFGKHDYKTDPVYKNFIP
HITPFDESIYEKKDVPGVLRFGKREGDIPGVLRFGKRNDDIPGVLRFGKRSPLGVLRFGR
R
>FLP-18_Anisakis_simplex_ASIM_0001471501-mRNA-1 peptide:
ASIM_0001471501-mRNA-1
MVDWTSISSVHLLAFICISVYAEVELPVLSYYPDSAVVDRDDDPYQIAFTKLGRYSPDQE
GVDAESLAGNAAEKRVACLFSKYNDNIHEQSVTSSFPWDHKHCETGFSEEVYIGFGDEAS
IPGVLRFGKRGMPGVLRFGKRDNEKKAVPGVLRFGKRGDVPGVLRFGKRSDMPGVLRFGK
RAMPGVLRFGR
>FLP-18_Ascaris_lumbricoides_ALUE_0001030201-mRNA-1 peptide:
ALUE_0001030201-mRNA-1
LYAYQSKGGSPFSSLLSSISSHHSSSSEALGPLLQGADASNNMVELAAIAVHLFAILCIS
VSAEIELPDKRAQFDDSFLPYYPSSAFMDSDEAIVAVPSSKPGRYYFDQVGLDAENAMSA
AEKRGFGDEMSMPGVLRFGKRGMPGVLRFGKRENEKKAVPGVLRFGKRGDVPGVLRFGKR
SDMPGVLRFGKRSMPGVLRFGRR
>FLP-18_Ascaris_suum_GS_16229 peptide: GS_16229
MDSDEAIVAVPSSKPGRNAVNLMRINSFAISLEGFGDEMSMPGVLRFGKRGMPGVLRFGK
RENEKKAVPGVLRFGKRGDVPGVLRFGKRSDMPGVLRFGKRSMPGVLRFGRR
>FLP-18_Brugia_malayi_Bm18003.1 peptide: Bm18003.1
METTGWLLQMIKLIIIIVTITTTTKTTITAANLYDHLDNSNFYRSLNDVYEPAATELLIK
RDYGDIPGVLRFGKRDYGANYPYKNFVPNVVPFGENDYEKKDVPGVLRFGKREGDIPGVL
RFGKRNDDIPGVLRFGKRSAPSLSRFGLH
>FLP-18_Brugia_pahangi_BPAG_0000590701-mRNA-1 peptide:
BPAG_0000590701-mRNA-1
MQKKLLIKAIHLTYYGDIPGVLRFGKRDYGANYPYKNFVPNVAPFGENDYEKKDVPGVLR
FGKREGDIPGVLRFGKRNDDIPGVLRFGKRSAPSLSRFGLH
>FLP-18_Brugia_timori_BTMF0000381301-mRNA-1 peptide:
BTMF_0000381301-mRNA-1
YGDIPGVLRFGKRDYGANYPYKNFVPNVVPFGENDYEKKDVPGVLRFGKREGDIPGVLRF
GKRNDDIPGVLRFGKRSAPSLSRFGLH
>FLP-18_Dirofilaria_immitis_nDi.2.2.2.t11162 peptide:
```

nDi.2.2.2.t11162
XLFRLGKDNYGTDYVYNQKLIPDTVQFGENGYEKKDVPGVLRFGKREGDIPGVLRFGKRN
DDIPGVLRFGKRSVPGVLRFGRH
>FLP-18_Dracunculus_medinensis_DME_0000800701-mRNA-1 peptide:
DME_0000800701-mRNA-1
LPHHIHNSIYKCCNFRSSIISRRIFSDSIPGVLRFGKRLNLLQGLTKRTIPGVLRFGKRG
DHKRKNDVPGVLEKKDPIPGVLRFGKRDEIPGVLRFGKRSESDGFTLIKKMPGLRF
>FLP-18_Elaeophora_elaphi_EEL_0000518401-mRNA-1 peptide:
EEL_0000518401-mRNA-1
MYTPGVLRFGKHDYGTDPVYKKLIADIMPFDENVYEKKDVPGVLRFGKREGDIPGVLRFG
KRNDDMPGVLRFGKRDEDIPGVLRFGKRSAPGVLRFGRR
>FLP-18_Enterobius_vermicularis_EVEC_0001306101-mRNA-1 peptide:
EVEC_0001306101-mRNA-1
MDFCFTLKLGVTFSIIMMATALEQSNDRLMSSYFPVYIDDETDGGYPFLLKYPYLAYLAP
YDQGMPQLSKRVEKKDVDGKSDLNFRPSAFRFGKRGSVYSDDTAAGIPYLVKQSFYKFPQ
KRNSQSNRLGKKNMPGIIRFGRR
>FLP-18_Gongylonema_pulchrum_GPUH_0001812401-mRNA-1 peptide:
GPUH_0001812401-mRNA-1
LRFGKRIERVLYDSDKRDVPGVLRFGKRDQEKKNVPGVLRFGKREEDIPGVLRFGKREDV
MPGVLRFGKKNDVPGVLRFGKRNEVPGVLRFGKRTMPGMLRFGRR
>FLP-18_Loa_loa_EJD74811.1 peptide: EJD74811.1
METTGWMMHMVKLVIFIVTITATTTTLYVQLDNSNYGDIPGVLRFGKRDYDPDPVYRNP
LPNIMQFNENGYEKKDIPGVLRFGKREGDIPGVLRFGKRNDDIPGVLRFGKRSVPGVLRF
GRR
>FLP-18_Onchocerca_flexuosa_X798_06169 peptide: X798_06169
MVEEYKIGVHGGIPGLLRVGRDSVYNQKFAPDVLQFDEGVEKKSAPGVDIPGVLRFGKRI
EEVPGAVPFEKQTKPGVLRFGRR
>FLP-18_Onchocerca_ochengi_Onchocerca_onchengi_nOo.2.0.1.t11006-RA
peptide: nOo.2.0.1.t11006-RA
XLFRFGKDNSYNQKFAPSIMEFDDEYEKKAVPGVLRFGKREIPGVLRFGRRSEDVPGVLR
FGKRSEPGVLRFGRR
>FLP-18_Onchocerca_volvulus_OVOC3170.1 peptide: OVOC3170.1
METTSLTMHMVKLVIIIITITTITTTFGVQMDNLNSYRPPNDAYEFVGPELLMKKDLFRF
GKDNSYNQKFAPGIVEFDDEYEKKAVPGVLRFGKREIPGVLRYGRRSEDVPGVLRFGKRS
EPGVLIRFGRR
>FLP-18_Parascaris_univalens_PgR002_g209_t01 peptide:
PgR002_g209_t01
MSMPGVLRFGKRGMPGVLRFGKRENEKKAVPGVLRFGKRGDVPGVLRFGKRSDIPGVLRF
GKRSMPGVLRFRRR
>FLP-18_Thelazia_callipaeda_TCLT_0000586401-mRNA-1 peptide:
TCLT_0000586401-mRNA-1
MNEDRDASEILPNHDFNRISGLNYLSRHDYGIYYPGYASDGPGVLRFGKRGQEKKNVPGV
LRFGKRDDDIPGVLRFGKRPVPGVLRFG
>FLP-18_Toxocara_canis_Tcan_03428.1 peptide: Tcan_03428.1
MVTTRQVERHSSSSRATSTSSLLQCATPPNSMVDWATIVVHLFAFLCISVYAEVELPDKR
AQFDDPFISYYPISTFFDREEAAADVPVTLSKPGRYYVDQVGLDAENAMSAAEKRGLGDD
TSMPGVLRFGKRAMPGVLRFGKRDNEKKAVPGVLRFGKRGDVPGVLRFGKRSDMPGVLRF
GKKSMPGVLRFGRR
>FLP-18_Wuchereria_bancrofti_maker-PairedContig_855-snap-gene-0.6-
mRNA-1 peptide: maker-PairedContig_855-snap-gene-0.6-mRNA-1
METTGWLLHMIKLIIIIVTITTTTKTTITTANLYDHLDNSDFYRSLNDVYEPAATELLIK
RDYGDIPGVLRFGKRDYGANYRYKNFVPNIMSFGVLRFGKRNDDIPGVLRFGKRSAPSLS
RFGLH
>FLP-18_Litomosoides_sigmodontis FLP-18 predicted

```
LSGVLRFDKREGDISGVLRFGKRYDDIPGVLRFGKRSASGALRFGRF
>FLP-18_Syphacia_muris FLP-18 predicted
QFSGMPYYIKRLSLNNDFQKRIVSNTRLGKKNMPGIIRFGRR
>FLP-18_Ancylostoma_caninum_ANCCAN_13126 peptide: ANCCAN_13126
MWRVSTAPLILLAVLANAADLEEQVFDLPEGEYPDDKTLLGIVSQAQHVSAKRDFDNGMP
GVLRFGKRESIVKKEVPGVFRFGKRSNKKSVPGVLRFGKRSVPGVLRFGKREMPGVLRFG
KRSTPGVLRFGKRHDIPGVMRYHL
>FLP-18_Ancylostoma_ceylanicum_Acey_s0673.g1392.t1 peptide:
Acey_s0673.g1392.t1
MWRVSTAPLILLAVLANAADLEEQVYDLPEGEYPDDETLLGIVSQAQHVSAKRDFDNGMP
GVLRFGKREDIVKKEVPGVFRFGKRSNKKSVPGVLRFGKRSVPGVLRFGKREMPGVLRFG
KRSTPGVLRYGKRHDIPGVMRYHLKCDN
>FLP-18_Ancylostoma_duodenale_ANCDUO_05551 peptide: ANCDUO_05551
MWRVSTAPLILLAVLANAADLEEQVYDLPEGEYPDDETLLGIVSQAQHVSAKRDFDNGMP
GVLRFGKRDDIVKKEVPGVFRFGKRSNKKSVPGVLRFGKRSVPGVLRFGKREMPGVLRFG
KRSTPGVLRFGKRHDIPGVMRYHL
>FLP-18_Angiostrongylus_cantonensis_Angca_006813 peptide:
Angca_006813
MTVVSHVCSKSTSKGVMWRVSTSLILLTALANAADLEEQEKYTEAMTLLGIIPQDQHIYA
KRDLDGDMPGVLRFGKRRGFEKKEVPGVLRFGKRTNKKSVPGVLRFGKRSVPGVLRFGKR
EMPGMLRFGKRVVPGVLRFGKRHEIPEIIRFDKRSTYDILPMELLDKKSVPGVLRFGK
>FLP-18_Angiostrongylus_costaricensis_ACOC_0000285001-mRNA-1
peptide: ACOC_0000285001-mRNA-1
MWRVSTSIILLTVLANAADLEEQVYDIPEEKHTEAMTLFGIGPQEQRIYAKRDLDGDMPG
VLRFGKRRGFEKKEVPGVLRFGKRTNKKSVPGVLRFGKRSVPGVLRFGKREMPGMLRFDK
RSTYDILPMELLDKKSVPGVLRFGK
>FLP-18_Caenorhabditis_angaria_Cang_2012_03_13_00073.g3466.t1
peptide: Cang_2012_03_13_00073.g3466.t1
MQRWSSVILIVLCCLLREALSYTEPNIDGNMPGVLRFGXXXXTPIDDYEVARKLEQSGRV
FSKRDIDGNMPGVLRFGKRESGVQKKEMPGVLRFGKRNIFDETKRVPGVLRFGKRSDILD
KREMPGVLRFGKRATIQEIFDKRGEVPGVLRFGKRNVPGVLRFGKRSDFGEQYAGILLKK
SAPGVLRFGRK
>FLP-18_Caenorhabditis_brenneri_CBN03736.1 peptide: CBN03736.1
MQRWSSVLLITLCCLLRGALAYTEPIYEIVEDDIPAEDIEVARTNEKQDGRVFSKRDFDG
AMPGVLRFGKRGGVWEKRESSVQKKEMPGVLRFGKRAYFDEKKSVPGVLRFGKRDVPMDK
REIPGVLRFGKRDYMTELFDKRSEVPGVLRFGKRDVPGVLRFGKRSDLEEHYAGVLLKKS
VPGVLRFGRK
>FLP-18_Caenorhabditis_briggsae_CBG14115.1 peptide: CBG14115.1
MQRWSGVIFITLCCLLREASAYTEPIYEIVEDDIPAEDIEVTRGNEKQDGRVFNKRDFDG
AMPGVLRFGKRGGVWEKRESSVQKKEMPGVLRFGKRAYFDEKKSVPGVLRFGKRSYFDEK
KSVPGVLRFGKRDVPMDKREIPGVLRFGKRDYTEEMFDKRSEVPGVLRFGKRDVPGVLRF
GKRSDLEEHYAGVLLKKSVPGVLRFGRK
>FLP-18_Caenorhabditis_inopinata_Sp34__X0277900.t1 peptide:
Sp34_X0277900.t1
MQRWSGVLLITLCCLLRGTLAYTEPIYEIVEEDIPAEDTEFARTNEKQDGRVFSKRDFDG
AMPGVLRFGKRGGIWEKRDSGVQKKEMPGVLRFGKRSFLNEKKSVPGVLRFGKRSYFDDQ
KRTVPGVLRFGKREYIEELFNKRSEVPGVLRFGKRDVPGVLRFGKPFQVFFALVASEFPM
MDNPKCVNNIQMISLFSLQSVSFNSLNHDICLVFTMILYVVNSRAPYD
>FLP-18_Caenorhabditis_japonica_CJA10836.1 peptide: CJA10836.1
MQRWLGVLLIALCCSLQGLSAYTEPIYEIVEEDVPVEDFEALRNNEKPDGRVFNKRDFDG
AMPGVLRFGKRGGVWEKRDSSVQKKEMPGVLRFGKRAYFDEKKSVPGVLRFGKRDVPMDK
REIPGVLRFGKRDYMAELFDKRSEVPGVLRFGKRGVPGVLRFGKRSDFEEHYAGVLLKKS
VPGVLRFGRK
>FLP-18_Caenorhabditis_latens_FL83_00909 peptide: FL83_00909
```

MQRWSGVLLITLCCLLRGALAYTEPIYEIVEDDIPTEDMEISRSNEKQDGRVFSKRDFDG
AMPGVLRFGKRGGAWEKRESSVQKKEMPGVLRFGKRAYFDEKKSVPGVLRFGKRSYFDEK
KSVPGVLRFGKRDVPMDKREIPGVLRFGKRDYMTELFDKRSEVPGVLRFGKRDVPGVLRF
GKRSIEEHYAGVLLKKSVPGVLRFGRK
>FLP-18_Caenorhabditis_nigoni_Cni-flp-18 peptide: Cni-flp-18
MQRWSGVLLITLCCLLRGASAYTEPIYEIVEDDIPAEDIEVARGNEKQDGRVFNKRDFDG
AMPGVLRFGKRGGVWEKRESSVQKKEMPGVLRFGKRAYFDEKKSVPGVLRFGKRSYFDEK
KSVPGVLRFGKRDVPMDKREIPGVLRFGKRDYTEEMFDKRSEVPGVLRFGKRDVPGVLRF
GKRSDLEEHYAGVLLKKSVPGVLRFGRK
>FLP-18_Caenorhabditis_remanei_CRE00446.1 peptide: CRE00446.1
MQRWSGVLLITLCCLLRGALAYTEPIYEIVEDDIPTEDMEISRSNEKQDGRVFSKRDFDG
AMPGVLRFGKRGGAWEKRESSVQKKEMPGVLRFGKRAYFDEKKSVPGVLRFGKRSYFDEK
KSVPGVLRFGKRDVPMDKREIPGVLRFGKRDYMNELFDKRSEVPGVLRFGKRDVPGVLRF
GKRSIEEHYAGVLLKKSVPGVLRFGRK
>FLP-18_Caenorhabditis_sinica_Csp5_scaffold_00046.g2403.t2 peptide:
Csp5_scaffold_00046.g2403.t2
MTSVSEMEAEPQPNGLDMQRWSGVLLITLCCLLRGALAYTEPMYEIVEDDIPSEDIEVVR
GNEKQDGRVFSKRDFDGAMPGVLRFGKRGGVWEKRESSVQKKEMPGVLRFGKRAYFDEKK
SVPGVLRFGKRSYFDEKKSVPGVLRFGKRDVPMDKREIPGVLRFGKRDYMTELFDKRSEV
PGVLRFGKRDVPGVLRFGKRSDMEEHYAGVLLKKSVPGVLRFGRK
>FLP-18_Caenorhabditis_tropicalis_Csp11.Scaffold629.g16550.t1
peptide: Csp11.Scaffold629.g16550.t1
MQRWSSVLLITLCCLLRGALAYTEPIYEIVEDDIPAEDIEVARTNEKQDGRVFSKRDFDG
AMPGVLRFGKRGGVWEKRESSVQKKEMPGVLRFGKRAYFDEKKSVPGVLRFGKRSYFDEK
KSVPGVLRFGKRDVPMDKREIPGVLRFGKRDYMTELFDKRSEVPGVLRFGKRDVPGVLRF
GKRSDMEEHYAGVLLKKSVPGVLRFGRK
>FLP-18_Dictyocaulus_viviparus_DICVIV_09394 peptide: DICVIV_09394
MWRVSTAPLILLAVLANAADPEDEVYDLPDEKYTEAMTLLGISPQAQHIYAKRDLNGDMP
GVLRFGKRQDIEKKVVPGVLRFGKRANKKSVPGVLRFGKRSVPGVLRFGKREMPGVLRFG
KRAVPGMLRFGKRFGERSTYDVLPIELLDKKSVPGVLRFGK
>FLP-18_Diploscapter_coronatus_DCO_022330 peptide: DCO_022330
MQWRLNAISQCVIVLLSIAALAIQADGVVGLSDDFAREYLDNLISAEEHAYAKRDFDGSM
PGVLRFGKRSPYFEDRISKKEVPGVLRFGKRSTNEKKSVPGVLRFGKRDVPGLLRFGKRD
EIPGVLRFGKRSVDFAEDFDGENFVKKSVPGVLRFGRK
>FLP-18_Diploscapter_pachys_WR25_19952.1 peptide: WR25_19952.1
MVQCQEFSDSASDRHISKIGFPRRRQVPGVLRFGKRSTNEKKSVPGVLRFGKRDVPGLLR
FGKRDEIPGVLRFGKRSVDFAEDFDGENFVKKSVPAEAAFSRSSVLYPLHSAMSSSLLDT
AEFAEVDAMLDQILDENTDPCAVEGAYESLKKRASSEKDNVEVLWRLAVCCNELAGRFEK
KDRRKEKIIEGRQYALQAYNTDSKHFLAIKWTAILTGQLTNYVSAKERVELGGEMREYQD
AALAMKPDDFILLHLRGRFAYSIATLSWLEKKAAVLLFGPLPELTMDDALKDFMAAFELK
PDWIENLVYIARCHLDKKNHGCSEFRSAIPTQSCFVHLAIRDYFQAAAMEYYSLAAGKPP
RDAAETEQVEEAKAYIKKHGK
>FLP-18_Haemonchus_contortus_HCON_00164730-00001 peptide:
HCON_00164730-00001
MWRVSTASLILLAAIACAADVEEQVYDIPDEEYSEALTLLGIGPEAQHIYAKRDLDGGMP
GVLRFGKRENGVEKKEVPGVLRFGKRTNKKSMPGVLRFGKRSVPGVLRFGKREMPGVLRF
GKRAMPGVLRFGKRTEIPGMMRFGKRSTYDTIPLELLDKKNVPGVLRFGK
>FLP-18_Haemonchus_placei_HPLM_0000749601-mRNA-1 peptide:
HPLM_0000749601-mRNA-1
LTRHLQPTSAIAFTTIFYISWLPSKNKLFSDLDGGMPGVLRFGKRENGVEKKEVPGVLRF
GKRTNKKSMPGIYKPENRDFST
>FLP-18_Heligmosomoides_polygyrus_HPOL_0001729701-mRNA-1 peptide:
HPOL_0001729701-mRNA-1
MSPYLLGVMWRVSASLVLLAVLTYATEVEENVYDVPEEEYTEALTLLGIGPEAQHIYAKR

DLDGGMPGVLRFGKREGGVEKKEVPGVLRFGKRTNKKSMPGVLRFGKRSVPGVLRFGKRE
MPGVLRFGKRAVPGVLRFGKRREIPGVMRFGKRSTYDYIPLEILDKRAVPGVLRFGK
>FLP-18_Mesorhabditis_belari_mbelari.g10245.t2 peptide:
mbelari.g10245.t2
MLGRQCCVGSSLLIAFASLLILTQADNQKEFGVQGYDQETENLFDQFAQMQQPAGFYDDS
SHLAEKRGDMDGSMPGVLRFGKRAQSFIRFGRSDGDMQKKEMPGVLRFGKRSPQEKKSMP
GVLRFETRSPAFFALERKACLECSVLEESKR
>FLP-18_Nippostrongylus_brasiliensis_NBR_0001332501-mRNA-1 peptide:
NBR_0001332501-mRNA-1
MPGVLRFGKREGVEKKEVPGVLRFGKRTNKKSMPGVLRFGKRGVPGVLRFGKREMPGVLR
FGKRAVPGVLRFGKRHEIPGVMR
>FLP-18_Oesophagostomum_dentatum_OESDEN_00866 peptide: OESDEN_00866
MNFFHLFTKCFLSADMDGGMPGVLRFGKREGFAEKEVPGVLRFGKRSNKKSVPGVLRFGK
RNVPGVLRFGKREIPGVLRFGKRSAPVVFRYNKRQNIPGVMRCDRFQHLTLVKLTEIADS
ELPLTVYYFKHS
>FLP-18_Oscheius_tipulae_OTIPU.nOt.2.0.1.t11414 peptide:
OTIPU.nOt.2.0.1.t11414
MPCYTASFCFLLAVIVSWASAQSTGIDDDEQQQQMLMESLAADHFYAKRDLDGTMPGVLR
FGKRSSYLDGMRDRAMEKKEMPGVLRFGKRSIENEKKSVPGVLRFGKRDPMPGVLRFGKR
EEIPGVLRFGKRGDMPGVLRFGKREDMPGVLRFGKRAVPGVLRFGKRGDMPGVLRFGKRG
FEDYDEIPGILLKKSVPGVLRFGKK
>FLP-18_Parapristionchus_giblindavisi_Parapristionchus-ag_msk-
S_424-0.97-mRNA-1 peptide: Parapristionchus-ag_msk-S_424-0.97-mRNA-1
MMDHSSNHLLLSITLSSILLISTLIIIPSTMNRLNSIKDSLSEQMIEFNEIETSSWSSMR
QLRGYNLIKRMKRNTGAECNCESGSPCPVGPPGRTGQPGIDGTPGIRGEPGVRGEVGKLP
EVRPKVEGGCRQCPPGPKGPPGWQGPPGPPGKIGLEGNRGQNGRPGLIGPIGHPGMQGES
GSIGKDGEKGEKGRDGIKGEKGEKGEDGVKGAIGPKGYPGNPGLNGNTGTIGSHGPQGPT
GIHGPPGYKGHPGPIGEIGQPGSDAGYCKCPDRLNMKIIDSTTTVSTVTQRSTTEGKIES
TTKMERFSVDYIESDDHIDLIPIDKTYLLLVPQTGEIMNSPLMISSILLSLLVIISSSET
TGYDDVSESSLVDEVLSPSYDEFYHITDKRDSMPGVLRFGKRAQSFVRFGKRLSGIEKKE
MPGVLRFGKRSEKKSVPGVLRFGKRSDMPGVLRFGKKSVPGVLRFGRK
>FLP-18_Pristionchus_arcanus_arcanus-sn_msk-S_224-1.26-mRNA-1
peptide: arcanus-sn_msk-S_224-1.26-mRNA-1
MLFTGLDDAESLMIENMYPSYEDYHLQDKRDSMPGVLRFGKRAQAFVRFGKRLSPIEKKE
MPGVLRFGKRNEKKSVPGVLRFGKRSAGMDSDMETLIFKKSVPGVLRFGRK
>FLP-18_Pristionchus_entomophagus_entomophagus-ag_msk-S48-4.79-
mRNA-1 peptide: entomophagus-ag_msk-S48-4.79-mRNA-1
MNLILAYSTSLLSLINHFDSVHNGREFLYSKLSPLTLERIFAIVHNSFVSSCRTTPIEPA
DVPSTENNQSGEVKSLAILPTLEVRAARQKRLLSVAMKRSDRTSGAQLLGRTCPICRRVF
ENRCKLTLHQMEHKREANPYKCPLRGCPAEYPDYRQFDFLLPRLVHGILFIIELITFLLF
IFGIYPEKARLLLPFAVAEVLRVIVLAALIVYYIALNFQNDSFNGMSRAVGHLFFVLGAH
IFIIYLVIATYHTLRNRGSSLYAPPPQVMTTGHHVIVLEQPIEEAMLPTIRDVMSIHMKE
FEHIEKEVYSSVSSLRSPSSSSRIPRQSAPTCNCEGGSPCPAGPPGRPGQSGSDGLPGTP
GQVGEQGAPAAPVESRSKPDRGCRVCPPGPKGPPGYAGPPGALGKHGEPGPQGQSGRNGL
SGPLGPPGIPGDQGPLGKEGEKGEKGREGVRGEKGEQGTVGAHGASGPRGYPGNPGRDGA
AGRPGAIGTQGEAGKQGVPGYKGYPGPMGANGEPGDDAGYCKCPARINGEVSEQRVEATT
QKTEQPWFMKDTSTKSSLSADSSSSDKGVDYVESDRLDDGESLMIEGMYPSYEDYHLQDK
RDSMPGVLRFGKRAQAFVRFGKRLSPIEKKEMPGVLRFGKRNEKKSVPGVLRFGKRSSGM
ESEMEGLIFKKSVPGVLRFGRK
>FLP-18_Pristionchus_fissidentatus_fissidentatus-mkr-S15-1.48-mRNA-1
peptide: fissidentatus-mkr-S15-1.48-mRNA-1
MGLEDSDGLLEIFPSYDEYHLQEKRDSMPGVLRFGKRAQAFVRFGKRLSPIEKKEMPGVL
RFGKRNEKKSVPGVLRFGKRSMDGSDMEGLIFKKSVPGVLRFGRK
>FLP-18_Pristionchus_japonicus_japonicus-mkr-S120-9.63-mRNA-1

peptide: japonicus-mkr-S120-9.63-mRNA-1
MSLARHNYHQLSENGVNDEIYRALHLSYSYLNAACYYDRDDVALPNIQKFFMKLSDEKRE
LAFKLIHYQNERGGKVRLESIGKPDDTSFEHPKDALVMAIQHEKKTNDAFLALCEHAEKN
DDPHLSDFIEEELLEPVVASLKKLSDLHTNAKRVGKGLGEYVFDRESFDDNKTIMVAQSP
SPRPSPTHVPSFKCPSCEAVFLDTIDSLINHYDSVHNGREHLYSKLSPITLERIFAIVHN
SFVSSCRLPYSPSSSSVDSSFPFPAEQNRWKEWFVSSMKTLRCPFTDGLVLPTQNTNQLK
HDDSPPPLGTASVIKTTPIEPADVPSTPTNESGEVNGNGEDKVEKSLIGLLPTLEVRAAR
QKRLLSVAMKRSGSTGAQLLGRTCPICRRVFENRCKLTLHQMEHKKELNPYKLRVHLITS
HNGVMGAAVQMSQPIAPSVPPPIVKEEMISDWLQEPSLAKRYSCNLSMKKTVFVLLLFVL
LLSLGIVQVIFIEKFNFLLPRLVHGVLFIIELITFLLFIFAIYPEKPRLILPFAVAEVLR
VIVLSALIVYYIVLNFQHDTISIMARAVGHLFFVLIAHVFMIHLVIATYYTLRNRGSSLY
APPPQVLSTGHHVIVLEQPIEEAMVYKVIAPFFPYPSIMVGRFLSPLRDEFVVSLTLSSI
VLLATLIAIPSTIGRLNTIRDALTIHMKEFEHIEKEVYSSVSSLRSPPSSTRIPRQSTPE
CNCEGGSPCPSGPPGRSGQSGTDGLPGTPGRVGEAGEPAAPVERRTKPERGCRVCPPGPK
GAPGWTGLPGPLGKHGEPGPQGQSGRPGVSGPLGPPGVPGDQGPLGKEGEMGEKGREGVR
GEKGEQGTVGSPGATGPKGYPGNPGRDGTPGRPGAIGTQGEAGKQGVPGYRGYPGPAGVN
GVPGVDAGYCKCPARMTGEIMEAVAEQVTVAATTQKTEQPWFMKDTTTKATPSSDKGVDY
FNPKLMRLTYLPLIINCELDKCIGTPQDGIVLDRLLRSSLHCSLRPGMEDAESLMIESMY
PSYDEYHLQDKRDSMPGVLRFGKRAQAFVRFGKRLSPIEKKEMPGVLRFGKRNEKKSVPG
VLRFGKRSAGMDSDMETLIFKKSVPGIHYRFLH
>FLP-18_Pristionchus_maxplancki_maxplancki-mkr-S386-1.19-mRNA-1
peptide: maxplancki-mkr-S386-1.19-mRNA-1
RERERREGKRGRGRERAIHATSHLSLPLRLPTMSLARHNYHQLSENGVNDEISRALHLSY
SYLNAACYYDRDDVALPNIHKFFMKLSDEKRELAFKLIHYQNERGGKVRLESVNKPDETT
FEHPKDALIMAIQHEKKTNDSFRALCEHAEKNDDPHLSDFIEEELLEPVVDSLKKLADLH
TNAKRVGKGLGDERNQRMVAQSPSPRPFPTHVPSFKCPSCEAVFLDTIDSLLNHFDSVHN
GREHLYSKLSPLTLERIFAIVHNSFVSSCRLPYCIASTSGEASDFPFPTEQNRWKEWFVS
SMKTLRCPFTDGLVLPTQNNNNAKQHDESPPPLGTASVIKTTPIEPADVPSTPTFESGEV
NEKGDERTLRSLLPTLEVRAARQKRLLSVAMKRSGATGAQLLGRTCPICRRVFENRCKLT
LHQMEHKKELNPYKCPLRDCPSEHPDYRQLRVHLITSHNGVMGAPVQMSQPIAPSAPVMI
SDWLQEPSLAKRYSCNLSMKKTVYVLLLFVLLLSLGIVQVIFIERFNFLLPRLVHGVLFI
IELITFLLFIFAIYPEKPRLLLPFAVAEVLRVIVLSALIVYYIILNFQNDSLNGMARAVG
HLFFVLIAHVFIVHLVIATYHTLRNRGSSLYAPPPQVISTGHHIIVLEQPIEEAIPDEFL
VSLALSSIVLLSTLVAIPSTIGRLNTIRDALTIHMMEFDHIEKEVYSSVSSLRSSPSPHR
PPRQSSPECNCEGGSPCPVGPPGRPGQPGTDGLPGTPGRVGEAGESAVPVERRTKPERGC
RVCPPGPKGPPGWTGPPGALGKHGEPGPQGQSGRPGVSGPLGPPGVPGDQGPFGKEGEKG
EQGREGVRGEKGEAGTVGAHGAAGPKGYPGNPGRDGAPGRPGFMGTQGEAGKQGVPGYKG
YPGPVGTNGVPGIDAGYCKCPDRMTEVFVEVTKVMDEITVPSTTEKTEQPWFMKDTTTKT
TSSSEKGVDYVESESQSDIIPDGIVLGSLLLRAVHCCLRADSMPGVLRFGKRAQAFVRFG
KRLSPIEKKEMPGVLRFGKRNEKKSIRY
>FLP-18_Pristionchus_mayeri_mayeri-mkr-S4-6.54-mRNA-1 peptide:
mayeri-mkr-S4-6.54-mRNA-1
MISDWLQDPSLGTRYPCNLSMKKTIFVLILFVLLLSLGIVQVLRVIVLSALIIYYIVLNF
QNDSFNGMARAVGHLFFVLLAHVYIVFLVMSTYNTLRNRDSSRYAPPPSVLTTGHHIIQT
TNRGSDVRKEIATMTYSAHLSIYPPLIHRNLIDANTDAAKRMVALVDGGIKVAAPSSFHS
PIMGRFLGVLRDEFLLSAVFSSIVLLSSIIVIPSTIGRLNSIRDALNIQMKEFNAIEREV
YSSVSSLRSSSSSPSSSRLPRQSQPECNCEGGSPCPAGPPGRPGQPGTPGLPGTPGRIGE
EGAPAAPYEIRTKAERGCRVCPPGPKGVPGYTGPPGALGKHGEPGPQGEPGRTGVSGPLG
PPGVPGDQGSLGKEGEKGERGREGVRGEKGESGTVGAPGATGPRGYPGNPGRDGSAGGPG
PRGTLGEAGKQGVPGYKGYPGPAGSNGVPGIDAGYCKCPVRMSEVYTEVPEVTEKPTTTT
TEASYYSAATTGVSEQPWFMKDSTTKTAPPSEKGVDYVELEGQTDIDGIILDRRFFPSLH
CSCPGLDDADSLVMENMYPSYEDYHIQDKRDSMPGVLRFGKRAQAFVRFGKRLSPIEKKE
MPGVLRFGKRNEKKSVPALLRKVGKEYGVLRFGKRSPAMDSDIMEGLVFKKSVPGVLRFG
RKFANFALHSWFLLTARGIVGAGKSLTATLLPPITLLSLSLSPPPDKLLLYHPAVWIKD

>FLP-18_Pristionchus_pacificus_PPA24926.1 peptide: PPA24926.1
MRYSTDTGWYRPRCLSPSFSPLVQSYSSHHLNQASFIPCFFLPLQPQEIEWILLDYDRRL
RRKTYLESDLQILFTGLDDAESLMIENMYPSYEDYHLQDKRDSMPGVLRFGKRAQAFVRF
GKRLSPIEKKEMPGVLRFGKRNEKKSVPVEWAREKRIERKGASILALLRKVGREYGVLRF
GKRSVGMDSDMETLIFKKSVPGVLRFGRK
>FLP-18_Strongylus_vulgaris_SVUK_0001445701-mRNA-1 peptide:
SVUK_0001445701-mRNA-1
MWRVSTALFILLVALANAADLQEQVYDIPEGDYVNDLSLLENIPRAKELSDKPDLDSNMS
RVLRYAKSEDDLKKEILGVLRYGKRSNKKSVPGVLRFGKRSVPGVLRFGKRDIPGVLRFG
KKSMPSVLRFGKRHEIPGVMR
>FLP-18_Teladorsagia_circumcincta_TELCIR_06210 peptide: TELCIR_06210
MGMFRTYWPSKETGWRRDTKIPGSEPRRSLVPSSKYTVDLDGGMPGVLRFGKRENGVEKK
EVPGVLRFGKRTNKKSMPGVLRFGKRNVPGVLRFGKREMPGVLR
>FLP-18_Caenorhabditis_elegans_Y48D7A.2.1 peptide: Y48D7A.2.1
MQRWSGVLLISLCCLLRGALAYTEPIYEIVEEDIPAEDIEVTRTNEKQDGRVFSKRDFDG
AMPGVLRFGKRGGVWEKRESSVQKKEMPGVLRFGKRAYFDEKKSVPGVLRFGKRSYFDEK
KSVPGVLRFGKRDVPMDKREIPGVLRFGKRDYMADSFDKRSEVPGVLRFGKRDVPGVLRF
GKRSDLEEHYAGVLLKKSVPGVLRFGRK
>FLP-18_Micoletzkya_japonica_Micoletzkya_japonica FLP-18 predicted
KNLYTPRRLTPESPSFPLLAISYVPSISLIHPSSHLSPLPPLPQSFSPSGVLRFGKRSDNIPGVLRFG
KKSVPGVLRFGRK
>FLP-18_Necator_americanus FLP-18 predicted
IHETFNMNCRFGKRNNGNMISFPVLVKKSVPGMLRFGK
>FLP-18_Pristionchus_exspectatus FLP-18 predicted
RSKITLYSPPSHTLISSHRLTILSPSGYFIEIFDGIPSILQLSSPFSFPFTHSPSFIVSPHSSPIFSS
SGVLRFGKRSAGMDSDMETLIFKKSVPGVLRFGRK
>FLP-18_Cylicostephanus_goldi FLP-18 predicted
XXXXXXXXXXXXXXXXXXXXXXXXXXXXXXXXXXXXXXXXXXXXXXXXXXXXXXXXXXXXXXXXXXXX
QVSCDSGREACRVCCDSEKEKYLVSYGGCFLSILSCYKVVCRFGKRSMPSVLRFGKRREIPGMMR
XXXXXXXXXXXXXXXXXXXXXXXXXXXXXXXXXXXXXXXXXXXXXXXXXXXXXXXXXXXXXXXXXXXS
GVLRFGKRSVPGVLRFGKREIPGVLRWVFFIHFELL
>FLP-18_Heterorhabditis_bacteriophora FLP-18 predicted
KSREQAFRFGKRSLNFDDVPGVLLGKKSVPGVLRFGK
LAGVFRFDKRDMPGVLRFGKREMPGVLR
KFLCSFISRFGKREEIPGVLRFGKRSVPGVLRFGKRTVPGVLRFGKREDIPGVLR
>FLP-18_Bursaphelenchus_xylophilus_BXY_1201400.1 peptide:
BXY_1201400.1
MSNQWLLVCVMGVVGVCTVQAAFFRGQPWKELFIADDSRADKRSVAYLPTLDEFSSYGSY
YYNSPDEVDGYRQAKRDDQAMPGVLRFGKRADIDKKEMPGVLRFGKRSDGEAEKKAVPGV
LRFGKRSDMPGVLRFGKRDGAEMPGVLRFGKKSEMPGVLRFGKRSDMPGVLRFGKRGDMP
GVLRFGRK
>FLP-18_Halicephalobus_mephisto_HMEPH_05621-RA.p1 peptide:
HMEPH_05621-RA.p1
FSFQGQTGRFEMQLRSLLLNHFFLALVITVIIGLVAAEESHDKSKEDGDMEKKMMPFASP
LEFNENIYFPEEEGADELNALDKRSPLSFEGDVPGILRFGKRSQSFVRFGRSGDELSKLV
EKKQVPGVLRFGKRSDMPGVLRFGKRSDMPGVLRFGKRDPDMPGVLRFGKRDADMPGVLR
FGKRDPDMPGVLRFGKRDVDMPGVLRFGKRDQEKKNVPGILRFGKRSDMPGVLRFGKRSD
MPGVLRFGKRDSDIPGVLRFGKRDSDIPGVLRFGKRFEISFLDKKSEMPGVLRFGR
>FLP-18_Panagrellus_redivivus_Pan_g22928.t1 peptide: Pan_g22928.t1
MPILATMKLRTDATCLLVAVIAAAAAMSCAAETSEDSQKGQKRMYPFVPTLGFEDSLTVT
DEPSFLDSEDLYRAAKRSDMADVPGILRFGKRSQSFVRFGRSAADELTRLMEKKQMPGVL
RFGKRSSDGDDDLKNEKKNVPGVLRFGKRSDMPGVLRFGKKRAFEDGAMDMPGVLRFGKR
AGFESGDMPGVLRFGKRGDMPGVLRFGKRGDMPGVLRFSKRASAFENGDMPGVLRFGKRS
DMPGVLRFGKRFELAYLDKKNDIPGVLRFGKK

>FLP-18_Parastrongyloides_trichosuri_PTRK_0000949300.1 peptide: PTRK_0000949300.1
MFLASGNSSSFIFFLLLVVICGIASGAYDDSEFDNIKIFFFIDDYEPNYLDISALNGEDF
AEKRDLEGGVIGIPGIMRYGKRDHPHLVRFGKRETEFMKKEMPGVLRFGKRNFNEQQKKA
VPGVLRFGKRRDMPGVLRFGKRGDMPGVLRFGKRGDMPGLLRFGKRRDMPGVLRFGKRGD
MPGLLRFGKRPSSFENFYLDKKDMPGLLRFGK
>FLP-18_Rhabditophanes.sp.KR3021_RSKR_0000312000.1 peptide: RSKR_0000312000.1
MRIIKQSTNTVFILLFVIVSAAGLLVEENQPCNTERDILLATDPSQNTQTYLKCVPANIG
SIGFWNKLFCAPATKFDFVTQSCIQDLSTNGLGGTFNFNAFKSYTNTQAYKTPPQTKEQL
PLVEPPLKIAILNNSCVNGETCIGNSICDISAGVCKCPYGHTADLSTLSCNEDKPDGGPM
VVSTVQQEDGSFKFTFRSQGPYSFNQQLSDLTMAKPTYKKVDNPEKIEFVGAKYPAEMPT
IKPFAVLKALFAKPGEACTDGKICGGGSYCMAPAMLCLCPGTLIEKHGECVHPVHETPAV
EKVNVGGLCNTYSECQDDSSCVLGRCRCISPLLEIAGKCIFQMAPKEVGPGEVCENGEVC
IKGSICSADIPVCICPVGTDLEDDICVPVLTTDTNTAPTLKPPTTMPQLPDTLPPTTAAL
PTTETSTTSSTTTTTTTTTPIPTTSTTTTIPPETKISTIYTMPTLTHTYPNRPLSAGAKF
LKVSETCSLNTDCMMGAYCNGNTNPPTCQCLSTHVNVNGKCEKVIYPGQYGCKNDIQCSV
SYPGSVCLDRQCNCPLGYRAIEQTCVSETNPFKKTTSIKPCSLTKPSFACYFAKMACPGG
KCSSSFDVNKNLFDSNHRLITLSKRQTKIMSVSRSSSWKGHAIDEDNSVSINSFTCLPTQ
LNCAGDKGLCYEGICYCFDGYFPDYQMKICIKIEERNQGSEGGVYVDVGGECQLYVDRCL
GGSVCSAVNQVCECSEGGRVKNGLCHQFPGGSCVKGESCDNGASCEMGICGCPESHIIVN
KSCILKIVQINEMCMHGERCMPGLVCRFGRCVRDSNLRMKPIEEFKSIPQFMIVAPGEVC
FKDEYCRAGKCIDGVCGCGQDEFLINGKCLKEKDGNVIEETTIDNCDKLECSNGFTCKSN
LCECAQGKQIIGEECLWSGDAEETLNKVLDTLKAQETSRVPISGEACSTLCGSGAFCLSG
ICTCPPLTVVDFLGRCKSSNPVATPSLPIYFKHQSNLAKASNTYHSGIEKELTALPGESC
MEVGTICLNYSFCQDGVCYCGPNKINHNNVCVEANKKLPEIVIKKIEPKMLHDYEDETKS
EENTNSNYVECDQTSFCPGGMQCLNNKCAFSHRLLSPPGGSCDSEETCTGNSVCRNDYCV
CREPNMIVINGLCLTSDILASTKVFETVTTVQTPSSPVYSTIGLKQNIGDRLAETIYQNT
FGPTEPSTIANTPTTQLIPVLANTNGGRKAAPGPLDSCVGGSQCLDNFCICPQGTIADTV
SGRCETAAEEEEDEEAVTEAVEEPRKGTLKQSAIYSFIPTTTSTNQQLINSLIINNQQKT
PILNSISTTKAPNSFKNNLINTDLTELNRLNLINPDSTATNSFNLINTDSTSKIVTSTPL
PLPNPQLYIIRRPATAYNSLTTSFLSKHPGQDIRENIPTIVHHHNISVISLNPDNNNNGD
QPKGDLYTIPFTQTYRSSSKEDFNKDNGINDKDLPTFATLFGGMALTTSNPFNFNKFKNL
NNNGSPSEIIKTEKSLEVPQVKLSSLPSAPSIVNEVVVDECAKQDQSTSYPNESCQDGQA
CVGGSECGPEKVCTCPSSRPIIRDNVCTSVNSTQLEKIAGPGEECSFEVKCSENAECSQS
LCRCNSGFIAISNECMKLPTAQSVSITNNVPSSDLKLQSAGGFGSNSGAKPRIVGAPIRR
TKKIKNGGKGAITTTNEAGICPSGNEPTRDDATNKVISCSGMTPNCPPRSYCYVTGVADG
IFNCYTDADYAPSEELVGLLNGEGDYSTESKRNLEEEDGAGVPALLRFGKRTVHNFYRFG
KRETALMKKDMPGLMRFGKRSNVPGLLRFGRSFDDKKSMPGFGSPMRFGKRGFDEEKSMP
LVMRFGKRSDFQYYNLNKKDMPGNLRFGKKSGMSDSLMLGKKSDMPGKSKVR
>FLP-18_Steinernema_carpocapsae_L596_g2714.t1 peptide: L596_g2714.t1
MLGHLNEIAVVGVFTLCALALVSAEIDSSDAAATASKLEYLVDELPKEDELLLEQLDRPT
WYDPEAYDTVKRAVGDETSMPGVLRFGKRGQAYVPFGRLDKKEMPGVLRFGKRGEKKAVP
GVLRFGKREIPGVLRFGKRDEIPGVLRFGKKSEMPGVLRFGKRNVPGVLRFGRK
>FLP-18_Steinernema_feltiae_L889_g15885.t1 peptide: L889_g15885.t1
MLGHLNEIAVVGIITICCTLVFVSAEVDSSDAAASASSKLEYFVEVPKDGGDELDEEGRA
TWYDPEAYDTVKRAIGEEASMPGVLRFGKRGQSFVRFGRLDKKEMPGVLRFGKRGEKKAV
PGVLRFGKRDIPGVLRFGKRDDIPGVLRFGKKSEMPGVLRFGKRNVPGVLRFGRK
>FLP-18_Steinernema_monticolum_L898_g5958.t1 peptide: L898_g5958.t1
MLGHLNEIVVVGFVALCTFAFVSAEIDSSDAASASKLEYFVDVPKDGGDELLEQDRPTWY
DPDAYDTVKRAMGDEASMPGVLRFGKRGQSFVRFGRLDKKEMPGVLRFGKRGEKKAVPGV
LRFGKREIPGVLRFGKRDEIPGVLRFGKKSEMPGVLRFGKRNVPGVLRFGRK
>FLP-18_Steinernema_scapterisci_L892_g18180.t1 peptide: L892_g18180.t1

MLGHLNEIAVVGVFTLCALALVSAEIDSSDAAAAASKLEYLVDELPKDDELLLEQLDRPT
WYDPEAYDAVKRAVGDETSMPGVLRFGKRGQAYVPFGRLDKKEMPGVFRFGKRGEKKAVP
GVLRFGKREIPGVLRFGKRDEIPGVLRFGKKSEMPGVLRFGKRNVPGVLRFGRK
>FLP-18_Strongyloides_papillosus_SPAL_0001744300.1 peptide:
SPAL_0001744300.1
MFLASGNSFISISIIIIASILGVALGTYDNSAYLEDSSVSVKDDYEPSYVSLADIESGDL
TEKRDYDGEIVGIPGIMRYGKRDYPHLIRFGKRETEYMKKEMPGVLRFGKRDYNGQQKKN
VPGVLRFGKRGDMPGVLRFGKRDQIPGLLRFGKRGDMPGVLRFGKRDQIPGLLRFGKRPS
YESFIIDKKDMPGLLRFGK
>FLP-18_Strongyloides_ratti_SRAE_2000372400.1 peptide:
SRAE_2000372400.1
MFLASGNSFISISLLIIAVILNVTSDAFDDSAYIEDSSVSVKDDYEPNYFGITNNDGGDL
TEKRDIDDGIVGIPGIMRYGKRDYPHLIRFGKRETEFMKKEMPGVLRFGKREYNEQQKKA
VPGVLRFGKRGVPGLLRFGKRDDMPGLLRFGKRDQIPGLLRFGKRGDMPGVLRFGKREQI
PGLLRFGKRPSFENYMLDKKDMPGLLRFGK
>FLP-18_Strongyloides_stercoralis_SSTP_0000782300.1 peptide:
SSTP_0000782300.1
MFLASGNSIIFLSILIIAAILGVTSGTYDDSAYLEDSPVSVKDDYEPNYFGLTNIDSGDL
PEKRDIDEGVLGIPGIIRYGKRDYPHLIKFGKRETEFMKKEMPGVLRFGKRKYNGQQKKA
VPGVLRFGKRGVPGLLRFGKRDDMPGLLRFGKRDQIPGLLRFGKRGDMPGVLRFGKRPSY
DDFLIDKKDMPGLLRFGK
>FLP-18_Strongyloides_venezuelensis_SVE_0560300.1 peptide:
SVE_0560300.1
MFLASGNSFISISIIIIASILGVVSGAYDNSAYLEDSSASVKDDYEPNYVSLADIENGDL
TEKRDYDEGIVGIPGIMRYGKRDYPHLIRFGKRETEYMKKEMPGVLRFGKRDYNGQQKKN
VPGVLRFGKRGDMPGVLRFGKRDQIPGLLRFGKRGDMPGVLRFGKRDQIPGLLRFGKRPS
YESFIIDKKDMPGLLRFGK
>FLP-18_Steinernema_glaseri FLP-18 predicted
CPKPLLLGVLRFGKREIPGVLRFGKRDIPGVLRFGKKSEMPGVLRFGKRNVPGVLRFGRK
>FLP-18_Acrobeloides_nanus_ACRNAN_Path_924.g3554.t2 peptide:
ACRNAN_Path_924.g3554.t2
MDMDKRFEDETSMPGVLRFGKRAQSFVRFGRSIDKGAVIKKEMPGVLRFGKRSSYDEDEI
ESKEKKDVPGILRFGKRDQDIPGVLRFGKRDIDNMPGVLRFGKRGDDIPGVLRFGKRDVD
NMPGVLRFGKRDIDNMPGVLRFGKRSDMPGVLRFGKRGDDIPGVLRFGKRDGIPGVLRFG
KRDDIPGVLRFGKRSNDFESLGVLRFGKKSEMPGVLRFGK
>FLP-18_Ditylenchus_destructor_Dd_02429 peptide: Dd_02429
MPTKCVSPFKDCHRIFIGHMDHFLLLLVLSLGYFTNSLADATSEGHTHEDAVSAEGNTDN
VPRDPFTQSAEKRMNSYVYSYPSADIMDVLRPWKRSADYGDSQMPGVLRFGKRSQSFIRF
GRSADFDDDGTRMLSKKEMPGVLRFGKRGHQEKKAVPGVLRFGKRDDIPGVLRFGKREMD
VPGVLRFGKRDGDMPGVLRFGKRDDIPGVLRFGKRYMEANYLPAGVLRFGKRDGDMPGVL
RFGKKSDMPGVLRFGKRSTDMPGVLRFG
>FLP-18_Ditylenchus_dipsaci_jg1467 peptide: jg1467
MMMDDGASMLSKRNAWSAQVRQERPTRKESCSWCSSLWQKEIRHSWSASIWQRDSDIPGV
LRFGKRGGDMPGVLRFGKRYSSLDNSNYLPAGVLRFGKRSDDIPGVLRFGKKSEMPGVLR
FGKRSSDMPGVLRFGKRSSDMPGVLRFG
>FLP-18_Globodera_pallida_GPLIN_000471600 peptide: GPLIN_000471600
MPSAKGRDSRHQLFDENSPEYFPSACCEFLHVQSALYLASFIQLVFVLLLSLLYLLLEQS
QFINAVDVFRPGVAFVICTNLVGILCALVGVCSHREQYISTQISILIGLVLLSDLFALLI
IFAMAFGARVQLASHLTFQQFLHSFLIDERRWEALLGPFWPYLLAILFHMAAWMFSYADL
MNNFGGPSPLDFVTGNGYYMLDEERPKREDEAVELLTAWKRSSPFRRGFLNGGCSDSVLR
FGKRGPQHEKKAVPGVLRFGKRAEVPGVLRFGKRMPQVLRFG
>FLP-18_Globodera_rostochiensis_GROS_g08903.t1 peptide:
GROS_g08903.t1
MDVTKRMFSYADLMNNFGGPSPLDFVTGNGYYMLDEERPKREDEAVELLTAWKRSSPFRR

GFLNGVQHNYLMKKDEFVAPGVLRFGKRMPGVLRFGKRGPQHEKKAVPGAPLRLMLFLCF
SFPLFHRSDRQMRHSLPFQPSFVASLVSPNIFK
>FLP-18_Heterodera_glycines_Hetgly.G000002830 peptide:
Hetgly.G000002830
MNKLQKQFSFGQKKSEKAFVRAWDGPAKGGTAQQKALAISVLGDGNGKDDGILRLRFSAK
SNLKTIHHELSTTFGLSWFRAWLSRRVVLRPDGRRCRFLFRGSLFLCPSVAMSVSSVLSL
PLLFCVCILQIIATAETEVPSDPPELTKRMFSYADLMNNFGGPSPLDFVSGNSFFILDEE
RPKREDEAVELLSAWKRSSPFQRGFLNGVQRNYLTKKDQFVAPGVLRFGKRMPGVLRFGK
RGPQHEKKAVPGVLRFGKRADVPGVLRFGKRMPQVLRFG
>FLP-18_Meloidogyne_arenaria_M.Arenaria_Scaff9662g071506 peptide:
M.Arenaria_Scaff9662g071506
MICYFQMFILFTSLFFCFGEIIANGEAGHNNEEIGMEKRMLSYADLMNNFGGPSALDLTG
QGLLIDDERPKREQNYNNDDVHIMTLWKRSPSYGPSFFNTAGDLTKKDDFIAPGVLRFGK
RMPGVLRFGVLRFGKRQSQESGAVPGVLRFGKRMPQVLRFGK
>FLP-18_scaffold833_cov169.g1441 peptide: scaffold833_cov169.g1441
MFILLTSLFLCFGEIIANGEAGHNNEEIGMEKRMLSYADLMNNFGGPSALDLAGQGLLID
DERPKREQNYNNDDVHIMTLWKRSPSYGPSFFNTAGDLTKKDDFIAPGVLRFGKRMPGVL
RFGKRDRVVIQEKKAVPGKPLF
>FLP-18_Meloidogyne_floridensis_scf7180000423522.g11204 peptide:
scf7180000423522.g11204
MFILLTSLFLCFGEIIANGEAGHNNEEIGMEKRMLSYADLMNNFGGPSALDLAGQGLLID
DERPKREQNYNNDDVHIMTLWKRSPSYGPSFFNTAGDLTKKDDFIAPGVLRFGKRMPGVL
RFGKRDRVVIQEKKAVPGKPCFNIK
>FLP-18_Meloidogyne_graminicola_NXFT01004063.1.10629_g peptide:
NXFT01004063.1.10629_g
MSIKNNIQTINNFDPNALEFYPSTCCEFLHVKSALYVASFIQANLKIFFTRFTLLYLILE
QSQFINAVDVFRPAIGFIILINLSGIVCALIGVCLQRDFFISVQILLLLALICISDLLAF
VLVFTMAFGSKISLGYSFDAKSLVQSFLVNDQRWESFLGPFWPYLLAIILHMTACSMICF
VAVYRRYRKFLRIFVFEKIITSKQVVFSINTLLSIALRDKVNGEVVNKEHEIDMEKRMFS
YADLMNNFGGPSALELVEPGNNLLLDDERPKREENYNNEDVHIMTLWKRSPSSYLTGDLT
KKDEFIAPGVLRFGKRMPGVLRFGKRDRVVVQEKKAVPGVLRFGKRQAQEGGAVPGVLRF
GKRMPQVLRFGK
>FLP-18_Meloidogyne_incognita_Minc3s00352g10804 peptide:
Minc3s00352g10804
MICYFQMFILLTSLFLCFGEIIANGEAGHNNEEIGMEKRMLSYADLMNNFGGPSALDLAG
QGLLIDDERPKREQNYNNDDVHIMTLWKRSPSYGPSFFNTAGDLTKKDDFIAPGVLRFGK
RMPGVLRFGKRDRVVIQEKKAVPGVLRFGKRQSQESGAVPGVLRFGKRMPQVLRFGK
>FLP-18_Meloidogyne_javanica_M.Javanica_Scaff1641g017255 peptide:
M.Javanica_Scaff1641g017255
MICYFQMFILLTSLFLCFGEIIANGEAGHNNEEIGMEKRMLSYADLMNNFGGPSALDLVG
QGLLIDDERPKREQNYNKDDVHIMTLWKRSPSYGPSFFNTAGDLTKKDDFIAPGVLRFGK
RMPGVLRFGKRDRVVIQEKKAVPGVLRFGKRQSQESGAVPGVLRFGKRMPQVLRFGK
>FLP-18_Meloidogyne_hapla FLP-18 predicted
LKLKSAPLRIFNFLIIQVLRFGKRDRVVQEKKAVPGMSLFNILENQIYNFYLIFKNFKECSVLAKDKL
KRVEQSPACSDSAKECPKFYDLENRSLIPKKTN
GVLRFGKRQAQESGAVPGVLRFGKRMPQVLRFGK
>FLP-19_Plectus_sambesii_PSAMB.scaffold272size59799.g3994.t1
peptide: PSAMB.scaffold272size59799.g3994.t1
MEAKVATLLLVIVTLLSTPITVSAEGELPVICTEAGCFLRLRSASVRPPLLAYNNRNLLH
AAYFSQQPKDRRVSDGAHEDSKLFIPPSGPARLIGDSQTKANQALWVIPIAYRTAAVDVS
PSAVLVDMPQARYFGNFFVLLMLLVLSLASSNGQRTKRQAPSVQADSLALEDMVKRNWAN
KIRFGKREDDEAQKRWASQLRFGKRFTADDNTDTDSPDLENKRGSWASKVRFG
>FLP-19_Acanthocheilonema_viteae_nAv.1.0.1.t10358-RA peptide:
nAv.1.0.1.t10358-RA

MSTGLLFYCILPILALAQNALLDEASEYRQELWPYMNDMEQGVNPNELSNLLYYDPRLKR
WASQLRFGKRANWASKVRFG
>FLP-19_Ascaris_suum_AgR011_g148_t01 peptide: AgR011_g148_t01
MNSRSPLLSAVFLIVVLCGFLSTAESEPEYRGEMMEPDERTYYPFLQWESAMLPHETDGA
ERMKKWASQLRLGKRASWASKVRFG
>FLP-19_Brugia_malayi_Bm5126.1 peptide: Bm5126.1
MFTDLLLYCILPVLTLAQNALLDETNEYRQDLWPYMNDMVQWPNELSNVLYYDPRLKRWA
SQLRFGKRANWASKVRFG
>FLP-19_Brugia_pahangi_BPAG_0000219001-mRNA-1 peptide:
BPAG_0000219001-mRNA-1
MKEAVPQCLLLFSLLITEQKFYFYEQNEKREKLQLYYDPRLKRWASQLRFGKRANWASKV
RFG
>FLP-19_Brugia_timori_BTMF0000944501-mRNA-1 peptide:
BTMF_0000944501-mRNA-1
MFTDLLLYCILPVLTLAQNALLDETNEYRQDLWPYMNDMVQWPNELSNVQNEKREKLQLY
YDPRLKRWASQLRFGKRANWASKVRFG
>FLP-19_Dirofilaria_immitis_nDi.2.2.2.t10238 peptide:
nDi.2.2.2.t10238
VYYDPRLKRWASQLRFGKRANWASKVRFG
>FLP-19_Dracunculus_medinensis_DME_0000672301-mRNA-1 peptide:
DME_0000672301-mRNA-1
MMKRTSKTKKERDDVQDNQESRWASQLRLGKRRRWASQVRFGRKR
>FLP-19_Elaeophora_elaphi_EEL_0000613801-mRNA-1 peptide:
EEL_0000613801-mRNA-1
MFRELLIYCILPVLALAQNTLSDEANEYRQDLWPYMDDLEQWSSPNELSNLLYYYDPRLR
RWASQLKFGKRANWASKVRFG
>FLP-19_Litomosoides_sigmodontis_nLs.2.1.2.t00862-RA peptide:
nLs.2.1.2.t00862-RA
MSKELLLYCILPALALPQQNALPDEGNEYKEKLWPDVDHAEEWPNPNELSNLLGLTDVKR
SDFKSKRRKKIEKLQAFYDPPLKRWASQLRFGKRANWASKVRFG
>FLP-19_Loa_loa_EFO22617.2 peptide: EFO22617.2
MYTELLLYCILPVLALAQNALSDEAYEYRQGLWPIVDDMEEWPNKLSSALYYDPRLKRWA
SQLRLGKRANWASKVRFG
>FLP-19_Onchocerca_flexuosa_X798_04301 peptide: X798_04301
MFAELLFYCILPVLALAQNTLTDEANEYRQNLWPYVDDTEHWPNDLSNFLYYDPPLKRWA
SQLRFGKRANWASKVRFG
>FLP-19_Onchocerca_onchengi_nOo.2.0.1.t05185-RA peptide:
nOo.2.0.1.t05185-RA
MFTELLFYCILPVLALAQNTLTDEANEYRQGLWPYMDNTEQWPNDLSNLLYHDAPLKRWA
SQLRFGKRVNWASKVRFG
>FLP-19_Onchocerca_volvulus_OVOC3513.1 peptide: OVOC3513.1
MFTELLFYCILPVLALAQNTLTDEANEYRQGSWPYMDNTEQWPNDLSNLLYHDAPLKRWA
SQLRFGKRVNWASKVRFG
>FLP-19_Thelazia_callipaeda_TCLT_0000673501-mRNA-1 peptide:
TCLT_0000673501-mRNA-1
MPKKILNDISIILILKFQKIYDDPRFKRWASQLRFGKRITWASKVRYG
>FLP-19_Toxocara_canis_TCNE_0001800601-mRNA-1 peptide:
TCNE_0001800601-mRNA-1
MILAEVVGEVCTTFPGAFSTAEPETEYREEMPADERHYFPQMRWQRPAMLPYEIESADRL
KKWASQLRFGKRASWASKVRFG
>FLP-19_Wuchereria_bancrofti_WBA_0000570601-mRNA-1 peptide:
WBA_0000570601-mRNA-1
MFTDLLLYCILPVLALAQNALLDETNEYRQDLWPYMDDMAQWPNELSNVVYYDPRLKRWA
SQLRFGKRTNWASKVRFG

>FLP-19_Anisakis_simplex FLP-19 predicted
SQLERIHILVFKEWENADRIKKWASKLRFGKRASWASKVRFG
>FLP-19_Ascaris_lumbricoides FLP-19 predicted
NTSVKDTDGAERMKKWASQLRLGKRASWASKVRFG
>FLP-19_Gongylonema_pulchrum FLP-19 predicted
XXXXXXXXXXXXXXXXXXXXXXXXXXXXXXXXXXXXXXXXXXXXXXXXXXXXXXXXXXXXXXXXXXXX
XXXXXXXXXXXXXXXXXXXXXXXXXXXXXXXXXXXXXXXXXXXXQLRFGKRSSWASKVRFG
>FLP-19_Parascaris_equorum FLP-19 predicted
NTSLKETDAAERMKKWASQLRLGKRASWASKVRFG
>FLP-19_Parascaris_univalens FLP-19 predicted
NTSLKETDAAERMKKWASQLRLGKRASWASKVRFG
>FLP-19_Angiostrongylus_costaricensis_ACOC_0000556001-mRNA-1
peptide: ACOC_0000556001-mRNA-1
MSSSLLWSSRKNIQVKQPYQYTFKLRMISWKKKKLENYKRWMQSVLCKYDMIMFKPSFQR
PYQKRWANQVRFGKRANNWASSVRFGR
>FLP-19_Caenorhabditis_angaria_Cang_2012_03_13_00091.g4067.t1
peptide: Cang_2012_03_13_00091.g4067.t1
MSFQLTLFTVLLIFIAIVLGQHVGSQPDSAENEALLAADADEFNSLMEYLQKTNPNAVRS
MQKRWANQVRFGKRASWASSVRFG
>FLP-19_Caenorhabditis_brenneri_CBN21772.1 peptide: CBN21772.1
MSFQLTLFSMLLLLIAVVVGQPIQSQIDDLKVVQAVQDNSPLSMEAFNDDPALYDYLESD
PSLKIMEKKWANQVRFGKRASWASSVRFG
>FLP-19_Caenorhabditis_briggsae_CBG04506.1 peptide: CBG04506.1
MSFQLTLFSMLLLLIAVVVGQPIQSPSGDLRVQAVQDNSPLSMEAFNDDPAVYDYIEQSD
PTFKVMEKKWANQVRFGKRASWASSVRFG
>FLP-19_Caenorhabditis_inopinata_Sp34__X0074200.t1 peptide:
Sp34_X0074200.t1
MLLLLIAVVVGQPVQSPYGDSKVQAEGDSSLPWIQGYVPYAPLYEEQLDKSPLTMEKKWA
NQVRFGKRASWASSVRFG
>FLP-19_Caenorhabditis_japonica_CJA02337.1 peptide: CJA02337.1
MSFQLTLLSILLLLIAGVVGQPIQSHASSLKVETGQDNLPLSMEAFNDDPTLYDYFELVD
PSPLKNKRWANQVRFGKRASWASSVRFG
>FLP-19_Caenorhabditis_latens_FL83_01290 peptide: FL83_01290
MSFQLTLFSMLLLLIAVVVGQPIQSHSDDLKVQAVQDNSPLNMEAFNDDSAYYYYYPEQA
DPSFKDVQKKWANQVRFGKRASWASSVRFG
>FLP-19_Caenorhabditis_nigoni_Cni-flp-19 peptide: Cni-flp-19
MLLLLIAVVVGQPIQSPSGDLRVQAVQDNSPLSMEAFNDDPAAYDYIEQSDPTFKVMEKK
WANQVRFGKRASWASSVRFG
>FLP-19_Caenorhabditis_remanei_CRE07236.1 peptide: CRE07236.1
MSFQLTLFSMLLLLIAVVVGQPIQSHSDDLKVQAVQDNSPLNMEAFNDDSAYYYYYPEQA
DPSFKDVQKKWANQVRFGKRASWASSVRFG
>FLP-19_Caenorhabditis_sinica_Csp5_scaffold_00039.g2135.t1 peptide:
Csp5_scaffold_00039.g2135.t1
MSFQLTLFSMLLLLIAVVVDQITISENSGESNKTRSRGARQPIQSPVGDLRVQAVQDNSP
LNMEAFNDDSALYDYLEGLGSVVDTASVMQKNDELFLSLLLIHFTTLCIISLQNNSGTLL
RRCKTGADECDWMNPLQNQNESSTSALFLQQPDPSLKVMEKKWANQVRFGKRASWASSVR
FG
>FLP-19_Caenorhabditis_tropicalis_Csp11.Scaffold629.g13782.t1
peptide: Csp11.Scaffold629.g13782.t1
MSFQLTLFSMLLLLIAVVVGQPIQSPNGDLKVVQAVQDNSPLSMEAFNDDPALYDYLEQS
EPSLKTIEKKWANQVRFGKRASWASSVRFG
>FLP-19_Dictyocaulus_viviparus_DICVIV_09512 peptide: DICVIV_09512
MHQSTSLRNILLVLSAFLCVLSYSSLEDSTGQKIDTRYGDVWQDFPYFLGQHQKRWANQV
RFGKRASNWASSVRFG

>FLP-19_Diploscapter_coronatus_DCO_024687 peptide: DCO_024687
MLLANILSLFLLLILTLASVYAFPRTNQEQAEADLTSASQEIYPYENQFDSYLASLSDGG
GFGLRPVKKWANQVRFGKRANWASSVRFGRRK
>FLP-19_Haemonchus_contortus_HCON_00175255-00001 peptide:
HCON_00175255-00001
MHRDLFLALLAFICVFAYRPFDGARYGHDGTYEELWHSDLNTRGRNQKRWANQVRFGKRA
SSWASSIRFG
>FLP-19_Heligmosomoides_polygyrus_HPBE_0000412201-mRNA-1 peptide:
HPBE_0000412201-mRNA-1
MLRTFATWIPPPSKKNRVQSQAGGRGPYNDCNHQFQPQFQHFNQKRWANQVRFGKRASSW
ASSVRFG
>FLP-19_Heterorhabditis_bacteriophora_Hba_17684 peptide: Hba_17684
MYIVARPQMDSDNSLMWYDSDDGTLRNSELGMGSLYAQHQKRWANQVRFGKRGNWASSVR
FG
>FLP-19_Mesorhabditis_belari_mbelari.g24225.t3 peptide:
mbelari.g24225.t3
MLLRLFTLIALLFALVCIVYGADEQALESEDADNVAPLRLVRLANSDADIDLPQIYKRWA
TQVRFGKRAANSWASSVRFG
>FLP-19_Micoletzkya_japonica_MIJAPON000017745.t1 peptide:
MIJAPON000017745.t1
NPSSSPIPFIPSAMLHRLSLCILSIALLIASVYGDAQDSHWDDVGPYPQWKLDSFPRSLR
SEADSLSAAKKWANQVRFGKRASWASQVRFGRRR
>FLP-19_Necator_americanus_NECAME_06129 peptide: NECAME_06129
MKAPRKNLTNVMGNNRKDVECIKLGERDALSIESGWKEEMVCVRVLKLGGAASKRLTSPF
HTHSTSNAPAVIHATPPAPSSLRHCMRTSCISVLAYPSLDDAKQDVDVAYLDSWQDTPFF
QGPYQKRWANQVRFGKRASSWASSVRFG
>FLP-19_Nippostrongylus_brasiliensis_NBR_0000715401-mRNA-1 peptide:
NBR_0000715401-mRNA-1
MGQWPETPELQHLQQKRWANQVRFGKRASSWASSVRFG
>FLP-19_Oesophagostomum_dentatum_OESDEN_05620 peptide: OESDEN_05620
MHQPSMLRHLLLILLAIVCVLAYPSLDDARQELDVPYMDPWQDIPAFQGSYQKRWANQVR
FGKRASWASSVRFG
>FLP-19_Parapristionchus_giblindavisi_Parapristionchus-sn_msk-
S_1201-0.16-mRNA-1 peptide: Parapristionchus-sn_msk-S_1201-0.16-
mRNA-1
MFTHHLFILISLLLISLSTLQADEDVQWDDLSPYSQWNYDSFPRSLRSEMSSPSKKWANQ
VRFGKRSGQKSWASQVRFGRR
>FLP-19_Pristionchus_arcanus_Pristionchus_arcanus_arcanus-mkr-
S_240-1.38-mRNA-1 peptide: arcanus-mkr-S_240-1.38-mRNA-1
MLTARLLVCVLLAVLCAVVLSAEYDDYAAPAQFDTFPRALRSELPSYSKKWANQVRFGKR
SGAGQRSWASQVRFG
>FLP-19_Pristionchus_entomophagus_entomophagus-mkr-S220-1.100-mRNA-1
peptide: entomophagus-mkr-S220-1.100-mRNA-1
MLTARVLILLIAVLCTITIRAEDELEDLVDRHPTQWKFDSFPRALRSEMPVYSKKWANQV
RFGKRSAGGQKSWASQVRFG
>FLP-19_exspectatus-ag_msk-S_1165-0.6-mRNA-1 peptide: exspectatus-
ag_msk-S_1165-0.6-mRNA-1
MEAKETSDHCSAYNLFCALENVTESHSKSFVAGQTAAREYDDYEQQRGAAQFDAFPRALR
SELPSYSKKWANQVRFGKRSGAGGQRSWASQVRFG
>FLP-19_Pristionchus_fissidentatus_fissidentatus-mkr-S589-0.16-
mRNA-1 peptide: fissidentatus-mkr-S589-0.16-mRNA-1
PPLHHERDMLTARLFLLLAAAVGTVTAAALRDDDESLAIDFDAPTQWKFDSFPRVLRSDM
GNGYSKKWANQVRFGKRSAGAVGQRSWASQVRFG
>FLP-19_Pristionchus_japonicus_japonicus-mkr-S82-1.1-mRNA-1 peptide:

japonicus-mkr-S82-1.1-mRNA-1
MLTARLVAALLLVAVCAVMVVAEYEEAEYEVRAPAQYEAFSRALRSELPTYSKKWANQVR
FGKRSGAGGQRSWASQVRFG
>FLP-19_Pristionchus_maxplancki_maxplancki-mkr-S15-7.9-mRNA-1
peptide: maxplancki-mkr-S15-7.9-mRNA-1
MLTARLLVVMLLAALCAVVITAEYDDYEPRAAAQFDTFPRALRSELPSYSKKWANQVRFG
KRSAGQRSWASQVRFG
>FLP-19_Pristionchus_mayeri_mayeri-mkr-S379-0.3-mRNA-1 peptide:
mayeri-mkr-S379-0.3-mRNA-1
MLTARLIVLLVATLCTLTYGAEYDDGPSDFESAPKQFDSFPRALRSEMPSYSKKWANQVR
FGKRSAGQRSWASQVRFG
>FLP-19_Pristionchus_pacificus_PPA31028.1 peptide: PPA31028.1
MLTARLLVCVFLAVLCAVVLSAEYDDYEQQRGAPAQFDAFPRALRSELPSYSKKWANQVR
FGKRSSGAGGQRSWASQVRFG
>FLP-19_Teladorsagia_circumcincta_TELCIR_05143 peptide: TELCIR_05143
MDGTVNASSSHKELWVTSIMAYPSLDDARHSNNAAYVEPWQYTPHIHGPYQKRWANQVRF
GKRASSWASSVRFG
>FLP-19_Caenorhabditis_elegans_M79.4.1 peptide: M79.4.1
MSFQLTLFSMLFLLIAVVVGQPIQSQNGDLKMQAVQDNSPLNMEAFNDDSALYDYLEQSD
PSLKSMEKRWANQVRFGKRASWASSVRFG
>FLP-19_Ancylostoma_caninum FLP-19 predicted
SLNISSKQVTSAQLACAQESQKTLIKSTQHLRAFLLHSYTFVPQHMQFQPALQGPYQKRWANQVRFGK
RASWASSVRFG
>FLP-19_Ancylostoma_ceylanicum FLP-19 predicted
TFLSHLYTFVLQHVQFQPALQGPYQKRWANQVRFGKRASWASSVRFG
>FLP-19_Ancylostoma_duodenale FLP-19 predicted
EHFYSIPYTFVLQHVQFQPALQGPYQKRWANQVRFGKRASWASSVRFG
>FLP-19_Angiostrongylus_cantonensis FLP-19 predicted
DLFLGRKKLEYYKRWMQNVLYKYDMIMFKPSFQGPYQKRWANQVRFGKRANNWASSVRFG
>FLP-19_Diploscapter_pachys FLP-19 predicted
ASLSDGGGFGLRPVKKWANQVRFGKRANWVSQNITKNEN
ASSVRFGRRK
>FLP-19_Haemonchus_placei FLP-19 predicted
RGDRVLTLESKVLEFLRGNRYEPTELYSIFNFQLNTRGRNQKRWANQVRFGKRASSWASSIRFG
>FLP-19_Oscheius_tipulae FLP-19 predicted
KVDFQIMTGYKNPEKKWANQVRFGKRASSWASSVRFG
>FLP-19_Halicephalobus_mephisto_HMEPH_01778-RA.p1 peptide:
HMEPH_01778-RA.p1
MWVQVLFVPGPVNWLRRKAVRFFNKNKHKHRVRLKPEPPRLAALTWTICFNMVSTRYLLC
LILGYVALVAAAYQADSEVHYQYVKRLPEYDPQYYPSSLWELDNAVRLKKWASQLRFGKR
SGNSWATQVRFG
>FLP-19_Panagrellus_redivivus_Pan_g13.t1 peptide: Pan_g13.t1
MVWTRIALCLFFAYVAVVANAYQNLDFPVQRPLRYGSLLAKRSSPLMDSESAEVESYNLY
PQNFFNVDNDVRLKKWATQLRFGKRSPWASQVRFG
>FLP-19_Parastrongyloides_trichosuri_PTRK_0000950500.1 peptide:
PTRK_0000950500.1
MVLSNVLIKFSLLLIGIICLINGEKGKIQPEPLTFDLARDGGLIHPSFYYEIKVPHDLSE
EDIPYQYMDQLMSIPRNKKWASQLRYGKRSNWASQLRYGRK
>FLP-19_Steinernema_carpocapsae_L596_g12999.t1 peptide:
L596_g12999.t1
MLTAQLILACLLAVFACALGYSSYPASETEDSRAVPFVPLPYGYWQSPQQFLDTNDELDD
SQRLKRSKWASQIRYGKRASWASQVRFG
>FLP-19_contig00432.0.71 peptide: contig00432.0.71
MLTAQLLLACLLAVFASAYGYSNFPPSDSDDSPVAPKVVPLPYGYWRSPEQPVETADTMP

ENDDSFRLLKRSKWASQIRYGKRGSWASQVRFG
>FLP-19_Steinernema_glaseri_L893_g21176.t1 peptide: L893_g21176.t1
MLGFPQPYLVHDSNSLMGPAAGHDLCPVLIGTSSLALGDIKLVSNDPAWIWIPSRRPETH
PPEKXXXXKLVSNDPAWISWIPSRRPETHPPEKSSLLEYLLSIAFGLTEEDPKANLVIGF
GSSVTIQQHPYLVSDLSLPLRTPKLDLRTDPGHFAMLSAQLTLACLLLLLAVVYGHSNYV
PSDSDDSPVVVVPYPYGPYGSWQPSQLVREDTGSEDDLFRAAKRSKWASQIRYGKRASWA
SQVRFG
>FLP-19_Steinernema_monticolum_L898_g29047.t2 peptide:
L898_g29047.t2
MSAVVKWRQFPSVTFSEPLRFAMLTTQLILACLLAVFAYAYGYSNFPPTESEESPSTQFV
PLPHGFWQQSPQLMADTLQEDDDSLRLKRSKWASQIRYGKRASWASQVRFG
>FLP-19_Steinernema_scapterisci_L892_g30829.t1 peptide:
L892_g30829.t1
MLTAQLILACLLAVFACALGYSSYPAPASETEDSRAAPFVPLPNGYWQSLQQLLDTNDDQ
DDSLRLKRSKWASQIRYGKRASWASQVRFG
>FLP-19_Strongyloides_papillosus_SPAL_0000460300.1 peptide:
SPAL_0000460300.1
MAYSSILIKVTFLIIGFILLISAESIKNVPESEQFNTNEGTELFNPHFYQFKKFQYPIDD
EIFSYYQANEQIPIRDRKWASQLRYGKRSSWASQLRYGKK
>FLP-19_Strongyloides_ratti_SRAE_2000348200.1 peptide:
SRAE_2000348200.1
MAYSSILNKITFLFLGFILLITAELNKNNAVSEQFTANEGVESFNPYLYQIKKFQYPYDD
DMSLYYQMNEQIPIRDRKWASQLRYGKRSSWASQLRYGKK
>FLP-19_Strongyloides_stercoralis_SSTP_0000410900.1 peptide:
SSTP_0000410900.1
MAYSNILTKVTFLLFGFILIITADFNKNNEGLEQFNSNEKIDSFNPYLHLFKKFQYPFDN
DDALLLYQINEQIPIRNRKWASQLRYGKRSSWASQLRYGRK
>FLP-19_Strongyloides_venezuelensis_SVE_1751700.1 peptide:
SVE_1751700.1
MAYSSILIKVTFLIVGFILLISAESSKNAPVSEQFNANEGTELFNPYFYQFKKFQYPTDN
EIFSYYQTNEQIPIRDRKWASQLRYGKRSNWASQLRYGKK
>FLP-19_Bursaphelenchus_xylophilus FLP-19 predicted
FQHAEEPHRMKKWASSLRFGKRGPGWASQVRFG
>FLP-19_Acrobeloides_nanus FLP-19 predicted
VDSNNLRLKKWATQLRYGKRGGPSGWASQVRFG
>FLP-19_Ditylenchus_destructor_Dd_08537 peptide: Dd_08537
MPPVMKPSRTVVWSLRRLFPSLPVSRNALLIFTLFLCVILFVNIGCEAASLSMREIDDYM
APYVDAYAVDNGSWQPESLRLKKWASQLRFGKRAASNGWASQVRFG
>FLP-19_Ditylenchus_dipsaci_jg24717 peptide: jg24717
MSLTKAIRRLILGRFLPASSWQSMFMIAFIFAVIVCVNVTEAALSMEDDYNGKIDSYMAP
YVDAMHEDWQPESLRLKKWASQLRFGKRGNGWASQVRFG
>FLP-19_Globodera_pallida_GPLIN_001242300 peptide: GPLIN_001242300
MNSSRSTVFSTVSLLSLFILSLIVVTTECRAGLIALEELDNQIGPYSVEWQLAPKAAGGD
VGVKDSAEGENWVPELLRLRKRWATQLRFGKKAVAAADGPIRQWASQVRFGRRGEEQRAR
REYIAAMDPSEAPGVNLFGTVIPKDVLFAQLFYLLYFASFGSLFPLLAIYFKQLGLSAAQ
AGILLGSRPLVEFVASPFWGSFADRFRKGKMLLLFSLCSMIVFTLAVGFVQPLTPYCVVL
DQNSTGGECKFLVPASEIISGGALGYIKKVAGIGRRRRDILSIIDLSTFDNKGDLVAGKA
PEYVTKDKVCNYDKEMYGTLVSPPHSTRVYREQAVEQAFMLLWLLISLGEFFASPALALA
DGYTLSLVSDTPKDFGKIRLYGSAGWGIAMFLMGIGLDFSDTFRNHPCPTKNTTERNYTL
NFVTCTLFALAALVVTTQFKFPQPSRQHRPDEIGGLVMDTRVTEVDPTMAQRTRAKQLNT
NSQTEEAPWKSALRAMLKNIHFLAFLLGVVSVGFGAGNIFAFLFWHLQDIGGSPLLFGIA
SITNHAAEIVTDPQAKAQFIKPKFVAGTVHVFGGECFPFSDHFLAFQPVLLQGAVVSILW
ASATSYVSIVSPPHLKGTSQHILALLSANFSRFLCKTLTFRFHGLGRGVGPIIGGFFIRA
HGTRALFLLMAFLTLCVLGVYYAVNLKLKNESVKYSGFDAFNEDTAGGETLAPQGLPMLH

FGDNKITEAFNQTAVSNQNYGAIEENSDEKQRDDAYDRYVTGH
>FLP-19_Globodera_rostochiensis_GROS_g04780.t1 peptide: GROS_g04780.t1
MSSSQSTVFSTVSLLLLLSLFLLSLIVVTTECRAGLIALEELDNQIGPYSVEWQLAPETA
GGNVGVGDSAEGENWVPELLRLRKRWATQLRFGKKAVAAADGPIRQWASQVRFGRK
>FLP-19_Heterodera_glycines_Hetgly.G000004102 peptide: Hetgly.G000004102
MDIPIGASDRNGHSHLSSFAPSADDSLQAAQRVAKPTALLEPVELDELRPIQSQNEVSPQ
QKTVDTRPARETKQSGNENEVVLTGSGTESAANSSMDIGKKQPKEMKNGNLLTTVYPKDV
PFSWISRPTFDVAAMPPILANPQNRLTAAAFADIVRSIQRDSRFVSYALFYSRMVPVWLV
FSIVILLAVLASSPMGGLPVMVFAFVWLFMLVLGLLFCALLQKFIVLALRVLVKEINAIS
QTFRLLVGIQNRGIVFIFYDFDGCREDIARQIRVQQANNLPFSSANCLSDIEITQKADEL
MLAHAQAYLKGLVKHRLRFPTRPSEGVSDFLPRHVAKSHCLCQFVAMEHFHKGPQKGLAP
WYERMFAVLDSLFFVLAFADLAHLGTRSITRLSLAIVFGAVGIGTVRLFTRHAIIGPMPS
AVYFASISLPPTANLHIKAVEFARRCHCHHYLFFYQPFLLYFTVFSTVSLFRLFPFFFVF
IITMTRESRAEMIGLEELDNQIGPYAIEWPLATETTGGEGKDSAGGENWVPELLRLRKRW
STQLRFGKRAFAGPRRQWASQVRFGRK
>FLP-19_Strongylus_vulgaris FLP-19 predicted
PVFHGPYQKRWANQVRFGKRASWASSVRFG
RNDNENLVLISXXXXWANQVRFGKRASWASSVRFG
>FLP-20_Plectus_sambesii_PSAMB.scaffold389size53576.g5306.t1 peptide: PSAMB.scaffold389size53576.g5306.t1
MNQTALAALILLVAVQLAQPYVLEPKRASELFGKRASELFGKRASELFGKRASELFGKRA
SELFGKRSAQPAFISILADRPYAEMIAKRSVYDDDYASAGMTPDMIDEVMDRFETQRGRR
ASELFGR
>FLP-20_Ascaris_lumbricoides_ALUE_0000661901-mRNA-1 peptide: ALUE_0000661901-mRNA-1
MKLVPWNIRAAETCGMRHMRLRDLGNGAVELRGLRYDGDDGSLETAGSDLLSSSYDSDAH
DVSRRAVLRLGKRSLMRLGKRGIMRLGKRAPMRLG
>FLP-20_Ascaris_suum_GS_07434 peptide: GS_07434
MSTLVRYVLAVLLIVDAVHLRRLAIIQGYDGDDGSLETAGSDLLSSSYDSDAHDVSRRAV
LRLGKRSLMRLGKRGIMRLGKRAPMRLG
>FLP-20_Dracunculus_medinensis_DME_0000374201-mRNA-1 peptide: DME_0000374201-mRNA-1
MNTFTTWLLIFVIMVGETSEKPNSFFISKNGPELENFPLEDISDFDLHPYIVAYKRAMMR
LGKRSVFRFGKRALMRLG
>FLP-20_Onchocerca_onchengi_nOo.2.0.1.t12479-RA peptide: nOo.2.0.1.t12479-RA
MALFERGVTREARHLRMRDTQTWVIYRAYNSLHRFDGKYVVYRFGGKHVCHFGGKHVYHFG
SKHVYRFGGKYVYRFGGKHVYCFGGKHVYRFGGKHVYCFGGKNFYSFDLFEAICTL
>FLP-20_Parascaris_univalens_PgR017X_g046_t01 peptide: PgR017X_g046_t01
MLVPPDICVNEMKFLPWNIRTAETCCMRHMRLRDSGNGDVELQGLRYDGDDGSLETAGSD
FLSSSYDSDAHDVSRRAVLRLGKRSLMRLGKRGIMRLGKRAPMRLG
>FLP-20_Toxocara_canis_Tcan_05432.1 peptide: Tcan_05432.1
MTLTKTKCEELQNFAAASALRARMPTFVEYMLIVSLLVDSIYLCRSAVMPDFEVDGVGLE
SADLADTSPAHRNSPLHTVNKRAILRLGKRALMRLGKRAMMRLGKRAPMRLG
>FLP-20_Ancylostoma_ceylanicum_Acey_s0081.g1505.t1 peptide: Acey_s0081.g1505.t1
MGHTSTNARKFITCLCCFILASLVVAFPNQQLKRQSLEDGVVPWRLLQYQSYPSWKTPTM
VEKRAMMRLGKRAMKVLEKRAMMRLGK
>FLP-20_Caenorhabditis_angaria_Cang_2012_03_13_00338.g9431.t1 peptide: Cang_2012_03_13_00338.g9431.t1
XAIVEYEDPAYENVENFLSENEKSDNSQVNKRAMMRFGKRAMMRFGKRAMMRFG

>FLP-20_Caenorhabditis_brenneri_CBN14777.1 peptide: CBN14777.1
MGYSHSRVFVALVLLSMLIAVCMAAPPAAPSQDVDNTSDDYPFFEEDTLGMTDDGSDGPH
EEKRAVFRMGKRAMMRFGKRAVMRFGKRSVFRLG
>FLP-20_Caenorhabditis_briggsae_CBG16156.1 peptide: CBG16156.1
MMGHSRSRFVIALLLLSVLIAICVAAPPAISLQDLPAEDYPFLEEDVLDLPSDGTDAPIA
EKRAVFRMGKRAMMRFGKRAVMRFGKRSVFRLG
>FLP-20_E01H11.3.1 peptide: E01H11.3.1
MLGYTQSRVVITLLLFSVFLAVCMATPSGYPGQELQNVSDDYPIYEEEGLQLSAEGTDEP
HEEKRAVFRMGKRAMMRFGKRAMMRFGKRSVFRLG
>FLP-20_Caenorhabditis_inopinata_Sp34__X0053700.t1 peptide:
Sp34_X0053700.t1
MIGYPQSRVVITLLLLSILVSACLAVPFPNPSWEMQSGSNDYPAEETNEFAGDGTDEQQL
HEEKRAVFRMGKRAMMRFGKRALMRFGKRSVVRLG
>FLP-20_Caenorhabditis_japonica_CJA04591.1 peptide: CJA04591.1
MMGYSCRAVIALLMLSVLIAVCVAVPNGYGSEDVVNEYPMLYDDVDPFDLQIEGTEEPLQ
KRAVMRMGKRAMMRFGKRAMMRFGKRSVFRLG
>FLP-20_Caenorhabditis_latens_FL83_02262 peptide: FL83_02262
MMGYSQSRVTIALLLFAVLIAVCVAAPPAFPAQDLQNEDWPMLEEEALEMPGDGTDGPHE
EKRAVFRMGKRAMMRFGKRALMRFGKRAVMRFGKRSVFRLG
>FLP-20_Caenorhabditis_nigoni_Cni-flp-20.1 peptide: Cni-flp-20.1
MMGHSRSRFVIALLLLSVLIAICVAAPPAISSQDLPAEDYPFLEEDLLELPADGTDAPIA
EKRAVFRMGKRAMMRFGKRAVMRFGKRSVFRLG
>FLP-20_Caenorhabditis_remanei_CRE16761.1 peptide: CRE16761.1
MMGYSQSRVTIAILLFAVLIAVCLAAPPAFPAQDLQNEDWPMLEEEALEMPGDGTDGLHE
EKRAVFRMGKRAMMRFGKRAVMRFGKRSVFRLG
>FLP-20_Caenorhabditis_sinica_Csp5_scaffold_00011.g738.t1 peptide:
Csp5_scaffold_00011.g738.t1
MMGHSQSRFVIALLLLSVLIAICVAAPPAFPGQDLAAEDYPIFAEEALEMPGDGSDGPHE
YGKTRRLPYGQKSYDEIRETRRDEIRKEIRVPTWLSHVNVLHLSILHLTLTSVRALDF
>FLP-20_Caenorhabditis_tropicalis_Csp11.Scaffold629.g13429.t1
peptide: Csp11.Scaffold629.g13429.t1
MMGSQSRIIVAFVLLSVLIAACFAAPPAVPSQELQNGSDNYPIFEDNLMELSNDDEPHEE
KRAVFRMGKRAMMRFGKRAVMRFGKRSVFRLG
>FLP-20_Diploscapter_coronatus_DCO_023826 peptide: DCO_023826
MLGPTTSSSKGFGPIDLCFVLIVLIAILGESDAMQVLYRNNKETGPLPEFDNIKAKSDDT
HKPLDEVNERTSYNLGDRYLNDYDATALARPFIYPAEKRAMMRLGKRSPFRLGKRALMRL
G
>FLP-20_Diploscapter_pachys_WR25_21700.1 peptide: WR25_21700.1
MLGPTTSSIKGFGPIDLCFVLIVLIAILGESDAMQVLYRNNKETGTLPEFDNIKSKSDDT
HKPLDEVNERTSYNLGDRYLDDYDATVLARPFIYPVEKRAMMRLGKRSPFRLGKRALMRL
G
>FLP-20_Heligmosomoides_polygyrus_HPBE_0001731601-mRNA-1 peptide:
HPBE_0001731601-mRNA-1
MVQMSTYARVFLTCFCYVILSSYVIALPDQYRSRPSLDNNLLSWRLSELPQSRILTSSAL
EKRAMMRLGKRVESGYGKRAIMRLGK
>FLP-20_Heterorhabditis_bacteriophora_Hba_13103 peptide: Hba_13103
MTEPGVSFSVHFDVSLKTARDMINLHLIVIFLHLLFISPISANGTLSVTELPYWKENRFI
DCIFCIDYDNFKRISVSGCATSRPMTCKGNICFMRQHKSPQFFLYTSGCLNLTQSEFELI
AHGRSDVDIQAGGKGDETLLCEVSRKTNTCLCSDQTRCNNMTLPSPFSEFHSPSLLSRLD
FRELLHFRLFIPQEPVLRYTDVRQVLELLAYKYQILVQLIIHNGYEAYKTDNTSDTLHTI
IFMILL
>FLP-20_Mesorhabditis_belari_mbelari.g16224.t1 peptide:
mbelari.g16224.t1
MTLRWLGSLLSRWIFLFSISLFLISSQAFPAYDAVLVEELSPQGVQYENLYMMPKRSPLR

LGKRGLMRMGKRGMMRLG
>FLP-20_Micoletzkya_japonica_MIJAPON000014964.t1 peptide:
MIJAPON000014964.t1
LQMQHVSWVLFLVLPLLAVAAVLEPEGELGNYYRSYYDSEAAVAGPESSVEKRAPMRLGK
RAFIYKRAPMRLGKRAALRLGKRSFS
>FLP-20_Oscheius_tipulae_OTIPU.nOt.2.0.1.t00537 peptide:
OTIPU.nOt.2.0.1.t00537
MSGLLRTVCIFILLVVLLNAFPRYGTSADLEIGNDYNSPEMYKSSEREEEPTVHKRALMR
LGKRAMMRLGKRAMMRLG
>FLP-20_Parapristionchus_giblindavisi_Parapristionchus-mkr-
S_103-0.3-mRNA-1 peptide: Parapristionchus-mkr-S_103-0.3-mRNA-1
MQVSWVLLFLCAISVIVSVQSEVIEDDSLSPRNYFGYGAQDDDGQPWGAIIKRAPMRMGK
RAAMRLGKRAPMRMGKRGPFRLGFYGTPWTQMQRREMIRLLSQLGMNTYVYAPKDDVKHR
LQWRDRYNTEEAGELQSLIEYSKSIGLDFIYALSPGIDIEYSNAKDITAVKGKLEQMKNI
GCEGFALLFDDIEYHLSDDDTSQYGTPANAQVAVVNDCYTHLNSPTHFYFCPTEYCSTRG
KPSLLESDYLRTIGDRLHPGIEIFWTGPLVIPKLISSDHSKAVAMVLKRKPLIWDNLYAN
DYDKRRLFLGPFQGRPQSLKSETSGILLNPNCQWELNYPAIHTYAEWFNFEQSPVPVTES
SMDEDSNSSSSSSYVGETALKSVLQGWLKDINVLIPNMPAPRPRLPTVELSLGSSPPGRP
DPCVTPITTISNITNAIKLTNDEGPSTSGSESGLVVNKEEPMDRSPSDESDVSMVDSGRS
TSNFDSNHLTFDQLLTLSHCYYLPFDHGERVKDILKGYMYCFREASLMATFKQPDVPLNT
EAIVDWLSYFSRVTEYIDHITCAFRFLHHNKALQIMGITVDLVKYASEAHGILAVMEGLL
HWIRDGNYSYVTEVRDLRWENDMYGEPWAVFHGFVEDVAYAILPITASAALNRLLSHKSE
ETMLSALRIRQFKADDVSNYSSLRRTVSRDLNEDLLEMKNFFNRFYRPFLDFNPELCFVA
EELTPRNGEVPTTTVCGILNILQFYESYHNSIGDTMIDESAISSAWMPPLPQQSDIQHYT
SFIDLRARCDELDKDASKRLIQMIAVVLSYQGSTGLYTACSPCERDKVNLLLSLQFTELN
STHMDQILLGLLLTDIELNSSDEEEFNVDIDEETEPREKSVEETSKTSVE
>FLP-20_Pristionchus_arcanus_Pristionchus_arcanus_arcanus-mkr-
S_41-4.80-mRNA-1 peptide: arcanus-mkr-S_41-4.80-mRNA-1
MQFSLVLSVILLLATTVAEADYEPSQSRLGFGGYSNWASGASAPARWIDADIIEEEKRAP
MRLGKRAAFRMGKRAPMRLGKRAPMRMGKRGPLRLGFYGTPWSQSQRRDLLRVLSNQGLN
TYVYAPKDDVKHRKEWRDKYGTEEAAELQTLIEYAKSLNINFVYALSPGMDIEYSKVKDI
SAVKAKLEQLKSIGCESFALLFDDIEYKLSDADELQFGSPANAQVAVVNDCYTQLNSPSH
FYFCPTEYCSTRAKPTLQESDYLRTIGDRLHPGIEIFWTGPLVVPENISEDNCRAIGMVL
KRKPLIWDNLYANDYDTKRMFLGPFKGRTQSLKKECAGILLNPNCQFDLNRPAILSYTEW
YHAEEEGSDTDDSSMTTEPPPNRFAPSTSYDRSMTKWIKELFLPFIPTRLRTTENDFGSS
ESSQHTTFSQTTDLSPNEGVMVLEEDKPIEEPMDRSPSCASDISMADSGRFTTILESAPM
TVAQLGALASCYYLPFEHGPIATDILNGILWMYRHANVMAPGYIEEVVRKRSASGNSNGS
DKNIEEIEKTRANWMIQHKKVSDGIELIAGAFRTISANKDHNSDLVQYASDAYGVLVVVE
AVLEWIRQGHISELTEPRERRWTTPEYFGEPWHIFFSFVDNVAYSMLPLSHPHSLIFGHR
DTAPMSAVMHIRGLKSQYYADWQPSLRRTLPDDEVDSMESLDTYKEKYMKPFLEWSEKHC
FVAEEETMTGRVPIASICGVLNILDYYEHHRKLEEERMNSEEEDLTSPAWLPPLPTDPTV
LHFSSFMDIRVKDETDIHAAKRLIQTLAVVLAYDGAVGLYTCCLPSERDKVSLLLSLNFI
ELSSTHTDQILLGIRLVIPDDDAFSEEEEGDDDKEEDGERLLKEKSVEDRGDKDDY
>FLP-20_Pristionchus_entomophagus_entomophagus-mkr-S87-1.33-mRNA-1
peptide: entomophagus-mkr-S87-1.33-mRNA-1
MQFSLVLSALLLIATTITAVDYEPSQSRFAPAAYSNWASRASDPSAWIGNDDVIDEAQKR
APMRLGKRAAFRMGKRAAFRMGKRAPMRLGKRGPLRLESHFTSIGMITTLDICSMDAAAM
EGHPLLRPPSKPFIRGIIEGFYGAPWTQSQRRDLLRTLAQQGLNTYVYAPKDDVKHRKEW
REKYGTEEAAELQSLIEYAKSLNINFVYALSPGIDIEYSRVKDISAVKAKLEQLKSIGCE
SFALLFDDIEYQLSDADESQFGTPANAQVAVVNDCYTQLNSPAHFYFCPTEYCSTRAKPT
LLESDYLRTIGDRLHPGVEIFWTGALVVPESISEDNCRAVSMVLKRKPLIWDNIYANDYD
TKRMFLGPFKGRGQSLKKECCGLLLNPNCQFDLNRPAILSYTEWFNAEEEGSDTDDSSMT
TDPPSSRFVPGHSFERAMTKWMKELFLPFIPTRIRLAEVEFGSSGSSQLTTFSQATDSPN
EGSSEMIMEEPKPNEEPMDRSPSCGSDISMADSGRFTTIIESSPLTIAQLCALADCYYLP

FEHGQTANDILNGILWIGNSNGSDKNFEEIEKTRANWILQHQKITECIELISGAFRALCA
NKIHNPDLVQYASDAYGVLIVVEAVLGWIREGHLSELTEPRERRWTKPEYFGEPWHLSFS
FVENIGYSMLPPSHLHSVLFGHRDVDPMTAILSIRGLRREDTPLLSPLRRTVADDDVDSI
FSLETFVDKYYRPFLDWSEKHCFLAEEETKTGKIPVASVCGVLNIIEYHEHHNKLEADRM
SGEGELPSPPAWMPAIPTEPAIGQFSSFFDLRVKDESDIPAAKRLIQTLAVVMAYNGAVG
IYTCCLPSERDKVSLLLSLNFVELSSTHTDQILLGLRLVMPDVVVSEDEEESEQKEEDSE
REIKEKSVDERGDKDDY
>FLP-20_Pristionchus_exspectatus_exspectatus-mkr-S_5-4.4-mRNA-1
peptide: exspectatus-mkr-S_5-4.4-mRNA-1
MQFSLVLSVILVLAITVGAADYEPSQSRLAFGGYSNWASGASAPARWIDADIIEEEKRAP
MRLGKRAAFRMGKRAPMRLGKRAPMRMGKRGPLRLG
>FLP-20_Pristionchus_fissidentatus_fissidentatus-mkr-S82-1.11-mRNA-1
peptide: fissidentatus-mkr-S82-1.11-mRNA-1
MNYIKQSYVHLADMQSYWLFITILALATLNAVVMAEDYDVPSRLGLGSYSSNWGEKASDG
GRWILDDDDSTAPQKRAPMRMGKRAAFRLGKRAPMRMGKRAPMRMGKRGPLRLVDSALPP
SDPARHHLAMISSLHICNIDLASMQGCLLLRPPPKPFIKGIIEGFYGAPWTQFQRRDLLR
TLAQRGLNTYVYAPKDDLKHRKDWRDKYGTEEAAELQSLIEYAKSLCINFVYALSPGVDI
EYSKVKDIGAVKAKLGQLKSIGCESFALLFDDIEYKLSDADESQFGTPANAQVAVVNDCY
THLNSPAHFYFCPTEYCSTRAKPTLLESDYLRTIGDRLHPGIEIFWTGPLVVPAIISDEN
ARSIAMVLKRKALIWDNLYANDYDTKRIFLGPFKGRSQSLKKECAGILLNPNCQFEVNIP
SIISYSDWLTTDEDGSDTDDSSMTIDQQSNRFIPSTSFERTMAKWLKDLVVPIIPLRIRL
NEIDFNSSISSQQTTYSLITDFSPNEANRELVRNNYIQTSKMILEEQRSLEEPMDRSPSC
DSDISMADSGRFTGIYEGSSLTNVQLCALAHCYYLPFEHGPTAHDILNGVLWMYRHASVM
SAGYVEEVRKRSVSSGSNSSEKEEEAEKTRENWIKQHGKVADAIELIAGAFRVIGANRIH
NPDLVQYASDVYGVLIVVEAVMGWIKEGNLSELTEPRERRWSLDRFGEPWNVSAAFVDDV
AYAMLPPINPLTFIFGHRDREPINGMMTIRRTRRDDLQSMSGLRRTVPDDDIDSIFSTEI
FIEKYYRPFLDWSEKHCFVAEDGMRMRKTPLASICGVPNIIDFYEHHNRIEAEKMNSDDE
CPSVPSWLPPLPTAPAVHHFTSFCDIRAKDEGDIHAAKRLIQTLAVVLAYEGSIGLYTCC
HPCEREKVSMLLSIGFVELNSTHTDQILLGLKFVPPDAHFTDDEDEEDIEEEMEKEAKEE
NVEDRGDQDDY
>FLP-20_Pristionchus_japonicus_japonicus-mkr-S181-1.52-mRNA-1
peptide: japonicus-mkr-S181-1.52-mRNA-1
MQFSLVLSLILVLATTITAADYEPSQSRLAIGGYSNWADGASVPRWIDADIIEESKRAPM
RLGKRAAFRMGKRAPMRLGKRAPMRMGKRGPLRLGFYGTPWTQSQRRDLLRVLAQQGLNT
YVYAPKDDVKHRKEWREKYGTEEAAELQSLIEYSKSLDINFVYALSPGMDIEYSKVKDIS
AVKTKLEQLKSIGCESFALLFDDIEYQLSDADESQFGTPANAQVAVVNDCYTQLNSPAHF
YFCPTEYCSTRAKPTLQESDYLRTIGDRLHPGIEIFWTGPLVVPENISGDNCRAIGMVLK
RKPLIWDNLYANDYDTKRMFLGPFKGRGQSLKKECSGLLLNPNCQFDLNRPAILSYTEWN
NADEESSDTDDSSMTTDPPANRFLPAPSYDRAMAKWMKELFLPFIPTRLRLTENDFGSSG
SSQHTTFSQETDLSPNEGVMILEEEKSVGEPMDRSPSCDSDISMADSGRFTTILESAPMT
IAQLCALASCYYLPFEHGKTATDILNGILWLYRHANVMAPGYIEEVVRKRNASGNSNGSD
KNIEELEKTRANWIMQHRKVTEAIDLIAGAFRTISANKDHNPDLVQYASDAYGVLVVVEA
VLAWIRQGHISELTEPRERRWTTPEFMGEPWHIFFSFVDNVAYSMLPPSHPHPLIFGHRD
TQPMNALMHIRGLRKEDVSNISSLRRTIAEDEIDSMSLVDAFPDKYMRPFLHWSEKHCFI
AEEETMSGRVPLASICGVLNVLDYYEHHRKLEEDRMKSEEEELIAPIAWLPPLPTDPTVL
HFSSFMDIRVKDETDIHAAKRLIQTLAVVLAYDGAVGLYTCCLPCEKEKVNLLLSLNFIE
LESTHTDQILLGLRLVIPDDTISVEEEEEEDNEEELERVRKETSVEDRRDKDDY
>FLP-20_Pristionchus_maxplancki_maxplancki-mkr-S8-7.76-mRNA-1
peptide: maxplancki-mkr-S8-7.76-mRNA-1
MQFSLVLSVILVLATAINAADYEPSQSRLAFGGYSNWAASAPDRWIDDKVIEEEKRAPMR
LGKRAAFRMGKRAPMRLGKRAPMRMGKRGPLRLGFYGTPWNQSQRRDLIRVLSQQGLNTY
VYAPKDDPKHRKEWREKYGTEEAAELQTLIEYAKSLNINFVYALSPGMDIEYSKVKDISA
VKAKLDQLKSIGCDSFALLFDDIEYKLSDADESQFGSPANAQVAVVNDCYTQLNSPSHFY
FCPTEYCSNRAKPTLQESDYLRTIGDRLHPGIEIFWTGPLVVPENISEDNCRAIGMVLKR

KPLIWDNLYANDYDTKRMFLGPFKGRPQALKKECSGILLNPNCQFDLNRPAILSFTDWNN
ADEEASDTDDSSMTTDPPASRFTSVPSYERAMAKWMKELFLPFIPTRLRLTENDFGSSGS
SQQTTFSQATDLSPNEGVMILEDDKEEPMDRSPSCASDISMADSGRFTTILESAPMTVAQ
LCALGSCYYLPFEHGQTATAILNGILWMYRHANVMAPGYIEEVARKRSASGNSNGSGKSV
EESEKTRANWMIQHRNVSDGIDLIAGAFRTIGANRDHNADLVQYASDAYGVLVVVEAVLA
WIRQGHISELTEPRERRWTTPEFFGEPWHIFFSFVDNVAYSMLPPSHPPSLIFGHRDTQP
MSALLHIRGLRNEDVPNISLRRTIGDDENDSMESLDCMRDKYMRPFLEWSEKHCFIAEQE
TMTGRMPLASICGVLNILDYYEHHRKLEEDRMKEEESLPAVAWLPPLPTDPTVLHFSSFM
DIRVKDETEIHAAKRLIQTLAVVLAYEGAVGLYTCCLPSEREKVSLLLSLNFVELSSTHT
DQILLGLRLVIPDDTFSEDEEDGEDNDDEKEKLRKASREKSVEDRGDKDDY
>FLP-20_Pristionchus_mayeri_mayeri-mkr-S241-2.21-mRNA-1 peptide:
mayeri-mkr-S241-2.21-mRNA-1
MMSFSLVLSFLFLLLAAAVSAADYDSSPSRLSLGAYWPSQSEFIIYDTFQTSEKKSLRWI
DDDDVIEEDKRAPMRMGKRAAFRMGKRAPMRMGKRAPMRMGKRGPLRLGDDMEVLSLSVI
EMTSYLDMLYSDQAVMEESIMGPLLRPPSKPFIRGIIEGFYGSPWTQAQRRDLLRTLSQQ
GLNTYVYAPKDDVKHRKEWREKYGTEEAAELQSLIEYAKSLDINFVYALSPGNDIEYSKV
KDISAVKAKLEQLKSIGCESFALLFDDIEYQLSDADESQYGTPANAQVAVVNDCYAQLNS
PTHFYFCPTEYCSTRAKPTLLESDYLRTIGDRLHPGIEIFWTGPLVVPETISEDNCKAFN
MVLKRKPLIWDNLFANDYDTKRAFLGPFRGRGQSLKKECAGILLNPNCQFDLNRPAILSY
TEWYNAEEEASDMDDSSMTTDPPPNRFVPSISFERAMAKWIKELFLPFVPSRLRLNDIEF
DSSGSSQNTNFSTDLSPNEGTVMVLEEPKSVEEPMDRSPSCDSDISMADSGRFTTIIDSG
PLNHAQLCALASCYYLPFEHGQTANEILNGILWMYRHANVMAPGYIEEVRKRSASEGSTG
SGKSFEGVEKTRANWMIQYEKTTEAIRLISGAFRALFANKIHNPDLVQYASDVYGVLVVA
EAVLAWIKEGHLSELTEPRERRWTMPDYFGEPWHISFSFVDNVAYAMLPPSHPHLVIFGH
RDNEPMNALMSIRALKSGDRANLSTLRRTVAADDVDSMFSLETFVEKYLRPFLDWSEKHC
FLAEEETRNGRVPVASICGVLNIFEYYDHHNKVEADRMGGQAEPSSPPAWLPPLPAESAG
QQFSSFFDIRVKDEADLHVAKRLIQTIAVVMAYEGAVGLYTCCHPCERGKVSLLLSLNFA
ELSSTHTDQILLGLQLIVPDAPLSEDEEEENDENEDKSERENKETSVEERGDKDDY
>FLP-20_Pristionchus_pacificus_PPA46569.1 peptide: PPA46569.1
MQFSLVLSVILVLSTTVAAAADYEPSQSRLAFGGYSNWASGASGPDRWIDSDDVIDEEKR
APMRLGKRAAFRMGKRAPMRLGKRAPMRMGKRGPLRLDDE
>FLP-20_Ancylostoma_caninum FLP-20 predicted
CFDFLHREGIVPMFVRINVFRTFSRPATIEKRAMMRLGKRAMKNLEKRAMMRLGK
>FLP-20_Ancylostoma_duodenale FLP-20 predicted
CFDFPQRKGIVPMLVRNNAFRIFSRPTTIEKRAMMRLGKRALKNLEKRAMMRLGK
>FLP-20_Caenorhabditis_sinica FLP-20 predicted
CHQFVFREKRAVFRMGKRAMMRFGKRAVMRFGKRSVFRLG
>FLP-20_Cylicostephanus_goldi FLP-20 predicted
SQLPPKLENSHFSVSNNVFFFRPSTMEKRAMMRLGKRATMHFDKRAMMRLGK
>FLP-20_Haemonchus_contortus FLP-20 predicted
LFRHNVAVKRAMMRLGKRLETYHRKRAIMRLGK
>FLP-20_Haemonchus_placei FLP-20 predicted
SFRHNVAVKRAMMRLGKRLETYHHKRAIMRLGK
>FLP-20_Heterorhabditis_bacteriophora FLP-20 predicted
LGNEILLCISMDLVKYKSRIMFGSNEIYYRQPAIEKRAMMRLGKRAMMRLGK
>FLP-20_Necator_americanus FLP-20 predicted
LIPTLLGGVAIIALVNNICNFRPSIPEKRSIMRLGKRATIHYHKRAMMRLGK
>FLP-20_Nippostrongylus_brasiliensis FLP-20 predicted
VIIFRSSAMEKRAMMRLGKRMLSQFDKRAIMRLGK
>FLP-20_Strongylus_vulgaris FLP-20 predicted
KVIPLLKKPLFRPSMIEKRAIMRLGKRATMEFNKRAMMRLGKXXXXXXXXXXXXXXXXXXXXXXXXXX
XXXXXXXXXXXXXXXXXXXXXDILKTLFKIE
>FLP-20_Teladorsagia_circumcincta FLP-20 predicted
RLVNCLTKNGDARTELFKTDWVIVSLARISSTGQLVTQFNQMPFRSSAAAKRAMMRLGKRVMSHYQKR

AIMRLGK
>FLP-20_Bursaphelenchus_xylophilus_BXY_1046900.1 peptide:
BXY_1046900.1
MMISAYLLGWLLAFIGVFTQAADLRQNYPTYTDAVFGEQPADPSFLSRSPYSTLYYANDN
PQAQAPAQLLGAEAAIPGQKRAYMRLGKRAYMRLGKRADQLQEMMEKRARLRLG
>FLP-20_Panagrellus_redivivus_Pan_g14459.t1 peptide: Pan_g14459.t1
MIVRELVATMLAILLTMTVIDSALGLPLNDDLSLYDDVYESPRQVESDFFTPEKRALMRL
GKRAYMRLGKRTSYFLPSKRAPLRLG
>FLP-20_Parastrongyloides_trichosuri_PTRK_0000469800.1 peptide:
PTRK_0000469800.1
MLFKLGAVLLILIIAINAPGIHSQQNGYQLQSKTLNQDAYDLEYPIYKSSYDDYISDINK
RAYVRLAGKRALVRLGKRAYVRLGKRRNELPEIF
>FLP-20_Rhabditophanes.sp.KR3021_RSKR_0000788550.1 peptide:
RSKR_0000788550.1
MYTNEHIKQNIKPEDSSYLLYANQRNPDVRQESYQKRAYLRLGKRALLRLGKRAYIRLVG
GQNYSNSVVTMDLITKEIKIAGQLLNGRKSHTSSRIDNHAYIWGGYCCKKIERYDVHTNK
TQEIKCRLDNVIDSLSSVVHDSKLFQIGGFSCKKTIDTVKYLDFEKCAFINESKLIVPDM
WHDSIVIDSNIYTTPGAYCKNIQRRDLRERNWQLMRKVIPNKTYRNAVCSLGQTNIVCIG
GWNLDSEVYSDCFVFDIRADDWRQIEPLPKALKAAQACETTKEIIVFGGIQSDHTYDSCI
NEILCYDKNENTWTKLDQNLPFPNGWSTKVTF
>FLP-20_Steinernema_carpocapsae_L596_g14227.t1 peptide:
L596_g14227.t1
MIFASLLAMILLCLQTLVYGYPHSPQSDLALQSYGDYYSPLEFGDNGENMEKRAMMRLGK
RAMMRLGKRSEFYAYEKRAPLRLG
>FLP-20_Steinernema_glaseri_L893_g32181.t1 peptide: L893_g32181.t1
MIIASLISLSILCLYAFVAGFPQVAQPELVPQLYGDYFMPQELTDYSENLVEKRAMMRLG
KRAMMRLGKRSDDSKHQKRAPFRLG
>FLP-20_Steinernema_monticolum_L898_g10355.t1 peptide:
L898_g10355.t1
MIFAAILSLFLVCLQTLVNGFPHVPQSEIGLQSYNDYFSPMEFNVNSENVLEKRAMMRLG
KRAMMRLGKRSEYFKYEKRAPLRLG
>FLP-20_Steinernema_scapterisci_L892_g28894.t1 peptide:
L892_g28894.t1
MIFASLIAVILLCLQTLAQGYPHSPQSDLVLQSYGDYYSPLEFGDNAENVEKRAMMRLGK
RAMMRLGKRSEFYAYEKRAPLRLG
>FLP-20_Strongyloides_papillosus_SPAL_0001260100.1 peptide:
SPAL_0001260100.1
MFAKVFITLIIFVVVINVSGFQSQSRDGQFLSKFAVPDNNDLQLNPYSNHYYGPMSEFKK
RAYVRLAGKRALVRLGKRAYVRLGKRDNQIPYIF
>FLP-20_Strongyloides_ratti_SRAE_X000178200.1 peptide:
SRAE_X000178200.1
MFLKVFLTLITFVIVFYVSGYQLQSTNDQFLSKNIILDNFVSQNLPYNHIYTEQNPKYNK
RAYVRLAGKRALVRLGKRAYVRLGKKNYLLPFDS
>FLP-20_Strongyloides_stercoralis_SSTP_0000265900.1 peptide:
SSTP_0000265900.1
MLPKIFITLITFVTIIYVTGFQLQPTNDQFLSGNLISDDLVLQDYPNVNIYNEQNPDIRK
RAYVRLAGKRALVRLGKRAFVRLGKRNYLLPY
>FLP-20_Strongyloides_venezuelensis_SVE_0825600.1 peptide:
SVE_0825600.1
MKYSFFRFILKYNCHITTMFAKVFITLIIFVVVINVSGFQSQSRDVQFISKFVTPDNNDL
QLNPYSHHYYGPMSEFKKRAYVRLAGKRALVRLGKRAYVRLGKRDNQIPYIF
>FLP-20_Halicephalobus_mephisto FLP-20 predicted
GSSSLFFSLRLPYRALLVESSKIKVILQLRLIYGYFSDPTLYDDFIDLQSLDEQPVLDKRAYMRLGKR
AYMRLGKRAYLRLGKRAPLRLG

>FLP-20_Steinernema_feltiae FLP-20 predicted
IGLFFXXXXXXXXXXXXXXXXXXXXXXXXXXXXXXXXXXXXXXXXXXXXXXXXXXXVENSESPLEKR
AMMRLGKRAMMRLGKRSHFFRYDKRAPLRLG
>FLP-20_Acrobeloides_nanus FLP-20 predicted
NFLALRSLSDSWTSFDTPSNNLRYYALYPVEDEASVSSNIDKRRFLHYIQKRAMMRLGKRSPMRMGKR
APLRLGRK
>FLP-20_Ditylenchus_destructor_Dd_01980 peptide: Dd_01980
MSQLILWNTLLFVLFTMCHADFTGLTYDESRSPDEQQRPLTWVEQNVRRHPTHPRALLID
DGREAILIPDAGRFRTVSDPTRFIDSSMLQWVTKPKRALMRLGKRAPMRLG
>FLP-20_Globodera_pallida_GPLIN_000095700 peptide: GPLIN_000095700
MTRRRMLSFSNLLPFFLLPLLLSLFQASDSFPSPGVFGLLQLNDQIRPPSLSTYAKLSLE
EPWMFVRPTQQRFGDRTRSYLWHKRAMAQVGKRALMRLG
>FLP-20_Globodera_rostochiensis_GROS_g04943.t1 peptide:
GROS_g04943.t1
MSTTRRRMLSFSNLPLLPFFFLPLLLSLFQASDSFPSPGVFGLLQLNDQIRPPPLSTYAK
LSLEEPWMFVRPTQHRFGARTRSYLWHKRAMVQLGKRALMRLG
>FLP-20_Heterodera_glycines_Hetgly.G000000503 peptide:
Hetgly.G000000503
MRFPRGVFSLLYCFNSSSLLPFLLPFPLIFCLLQTCDSFPAPGVFGLLQLNDQIRPPPIS
SYAKLPFDEPWMFFVPTHRFNDRTHLWHKRAMMRLGKRALMRLG
>FLP-20_Meloidogyne_arenaria_M.Arenaria_Scaff2050g029612 peptide:
M.Arenaria_Scaff2050g029612
MTNQRISCTQILLVFHCLSLILFYANSQPIMFAAPTFVKMSNGVIWPSLLLRPTDIDNEN
IFSQNPQEFVKVATPRKRAIMRLGKRALMRLGKK
>FLP-20_Meloidogyne_incognita_Minc3s04733g36903 peptide:
Minc3s04733g36903
MSNGVIWPSLLLRPTDMDTENIFSQNPQEFVKVAAPRKRAIMRLGKRALMRLGKK
>FLP-20_Meloidogyne_javanica_M.Javanica_Scaff923g011230 peptide:
M.Javanica_Scaff923g011230
MTNQRISCTQTLLVFYCLSFILFYANSQPIMFAAPTFVKMSNGVIWPSLLLRPTDMDTEN
IFSQNPQEFVKVAAPRKRAIMRLGKRALMRLGKK
>FLP-20_Meloidogyne_floridensis_maker-nMf.1.1.scaf21123-augustus-gene-0.3-mRNA-1 peptide: maker-nMf.1.1.scaf21123-augustus-gene-0.3-mRNA-1
MTNQRISCTQILLVFHCLSLILFYANSQPIMFAAPTFVKMSNGVIWPSLLLRPTDIDNEN
IFSQNPQEFVKVATPRKRAIMRLGKRALMRLGKK
>FLP-20_Meloidogyne_enterolobii FLP-20 predicted
KFFSRYFSHIFSHFSSQNPQEFVKVATPRKRAIMRLGKRALMRLGKK
>FLP-20_Meloidogyne_graminicola FLP-20 predicted
IFFKLIIFSQNSPEIAEIATPRKRALMRLGKRALMRLGKK
>FLP-20_Meloidogyne_hapla FLP-20 predicted
VVVLFGLHYCCVHQSWIQTTFIGNSVNFFLFFPILHTFLISSQNPEEIVQVATPRKRAIMRLGKRALM
RLGKK
>FLP-21_Plectus_sambesii_PSAMB.scaffold601size46076.g7362.t1
peptide: PSAMB.scaffold601size46076.g7362.t1
MSSLTITLLAFTVALCAIGVPLSAAQASNIENYDDSARLNQEALRTAILGRYYLDKLNEM
AGNDYLDESWLSTLKAQQQEGSSQAKRGLGPRPLRFG
>FLP-21_Ascaris_lumbricoides_ALUE_0001383801-mRNA-1 peptide:
ALUE_0001383801-mRNA-1
MSPFKRASAMRTVCTTFVIVPLLLVAIFGCISSALPSSDADVVYRLRVSTGAEQIFRMLI
SPICWKNRGSMKRGLGPRPLRFG
>FLP-21_Brugia_malayi_Bm17770.1 peptide: Bm17770.1
MNLIVLSILLITLFQQQYFAQVAPPNDLFYEFMNPYMNSLRSPDVNILSSYADERSWKRA
LGPRPLRFG

>FLP-21_Brugia_pahangi_BPAG_0000487201-mRNA-1 peptide:
BPAG_0000487201-mRNA-1
FRNIFFSKTCVVFFFTELFVSSIFIVTFSVLQTTDDESDSTQYPTYYAVPPTIFHPSSTS
SFRWIHLFQPTIICILVLLVFSVNPYMNSLRSPDVNILSSYADERSWKRALGPRPLRFG
>FLP-21_Dracunculus_medinensis_DME_0000042501-mRNA-1 peptide:
DME_0000042501-mRNA-1
MFAIVLSLLAFMIATHGASLSQNSDLDDAASRFLARYLTQLYSDPDGQNMPIIDEARLKK
NGLGLRPLRFG
>FLP-21_Litomosoides_sigmodontis_nLs.2.1.2.t01192-RA peptide:
nLs.2.1.2.t01192-RA
MNLTVFSILLVALLQQQCFAQGALPNEIFYEFMNPYMNSLRSPDANILSAYAEDRSWKRA
LGPRPLRFG
>FLP-21_Loa_loa_EFO25334.1 peptide: EFO25334.1
MDDEKMLSAFHRDFEILSSKARNVFSFIEQLCAPICVITFSVLKISGDESDSVQHSTYRA
IPATLFRTMNPYMNSLRSPDVNILSSYADERSWKRALGPRPLRFG
>FLP-21_Onchocerca_volvulus_OVOC7056.1 peptide: OVOC7056.1
MNLIILSILLIALFQQQYFAQIAPPNDLFYEFMNPYMNSLRSSDVNMLSSYADDRSWKRA
PGPRPLRFG
>FLP-21_Acanthocheilonema_viteae FLP-21 predicted
KNIDCFRSWKRALGPRPLRFG
>FLP-21_Anisakis_simplex FLP-21 predicted
GXFNFRSMKRGLGPRPLRFG
>FLP-21_Ascaris_suum FLP-21 predicted
RDSALVAHEYRNYRSMKRGLGPRPLRFG
>FLP-21_Brugia_timori FLP-21 predicted
VKFDLKNALHFRSWKRALGPRPLRFG
>FLP-21_Dirofilaria_immitis FLP-21 predicted
QYLDFRSWKRARGPRPLRFG
>FLP-21_Elaeophora_elaphi FLP-21 predicted
EIQFKKKSVHYFRSWKRALGPRPLRFG
>FLP-21_Gongylonema_pulchrum FLP-21 predicted
SYCRQHFRSWKRALGPRPLRFG
>FLP-21_Onchocerca_flexuosa FLP-21 predicted
CKERFKKIQIRFIEYRNFRSWKRAPGPRPLRFG
>FLP-21_Onchocerca_ochengi FLP-21 predicted
KYHNFRSWKRAPGPRPLRFG
>FLP-21_Parascaris_equorum FLP-21 predicted
YENKKGSKILLNSFSISAALVPHEYRDYRTMKRGLGPRPLRFG
>FLP-21_Parascaris_univalens FLP-21 predicted
YENKKGSKILLNSFSISAALVPHEYRDYRTMKRGLGPRPLRFG
>FLP-21_Thelazia_callipaeda FLP-21 predicted
KLITRNHIYRSWKRALRPRPLRFG
>FLP-21_Toxocara_canis FLP-21 predicted
CTYLIYRYFRSVKRGLGPRPLRFG
>FLP-21_Wuchereria_bancrofti FLP-21 predicted
KSKMEKKEKVKFDLKNALHFRSWKRALGPRPLRFG
>FLP-21_Ancylostoma_caninum_ANCCAN_04793 peptide: ANCCAN_04793
MRVSGFLVFIACIIAWAFAAPVSDTEAAYRILNKYLQRFGGDDLQDVYLVGDHG
>FLP-21_Ancylostoma_ceylanicum_Acey_s0099.g3158.t1 peptide:
Acey_s0099.g3158.t1
MRVSGFLVFIACIIAWAFAAPVSDTEAAYRILNKYLQRFGGDDLQDVYLVGDHGSIKRGL
GPRPLRFG
>FLP-21_Angiostrongylus_costaricensis_ACOC_0001281501-mRNA-1
peptide: ACOC_0001281501-mRNA-1

MRICGSIVFIVCVVAWAFAAPVSDAEAAYRILNQYLQRFDPEDLQDMYFISDHGSDKRGL
GPRPLRFG
>FLP-21_Caenorhabditis_angaria_Cang_2012_03_13_00318.g9100.t1
peptide: Cang_2012_03_13_00318.g9100.t1
MRFLVILFACILAWVLAAPSVIDREEDAFRLLNNYLQKYGPDENYIYLMDHGSAKRGLGP
RPLRXXXK
>FLP-21_Caenorhabditis_brenneri_CBN03252.1 peptide: CBN03252.1
MRLFILLSCLLAWVLAAPYIDQEDALRVLNAYLEQFGPGGGDKVYYVAEDDHGSMKRGLG
PRPLRFG
>FLP-21_Caenorhabditis_briggsae_CBG20591a.1 peptide: CBG20591a.1
MRLFILLSCLLAWVLAAPYIDQEDALRVLNAYLEQFGPAAEKVYYVAEDDHGSIKRGLGP
RPLRFG
>FLP-21_C26F1.10.1 peptide: C26F1.10.1
MRLFILLSCLLAWVLAAPYIDQEDALRVLNAYLEQFGPGSDRVYYVAEDDHGSMKRGLGP
RPLRFG
>FLP-21_Caenorhabditis_inopinata_Sp34__50086910.t1 peptide:
Sp34_50086910.t1
MRLFILFSCLLAWVLAAPYVDQEDALRVLNAYLEQFGPGADKIYYEAGDHGSMKRGLGPR
PLRFG
>FLP-21_Caenorhabditis_japonica_CJA00820.1 peptide: CJA00820.1
MRLFVLLSCLLAWVLAAPLVDQEDAFRILNTYLQQFGPDDNRNAYYVLNDDHGSMKRGLG
PRPLRFG
>FLP-21_Caenorhabditis_latens_FL83_22388 peptide: FL83_22388
MFQTMRLFILLSCLLAWVLAAPYIDQEDALRVLNAYLEQFGPGADKVYYVAEDDHGFRSM
KRGLGPRPLRFG
>FLP-21_Caenorhabditis_nigoni_Cni-flp-21 peptide: Cni-flp-21
MRLFILLSCLLAWVLAAPYIDQEDALHVLNAYLEQFGPAAEKVYYVAEDDHGLVLVKSRA
IEGFAEGIWHFGGFIGL
>FLP-21_Caenorhabditis_remanei_CRE18947.1 peptide: CRE18947.1
MRLFILLSCLLAWVLAAPYIDQEDALRVLNAYLEQFGPGADKVYYVAEDDHGSMKRGLGP
RPLRFG
>FLP-21_Dictyocaulus_viviparus_DICVIV_02218 peptide: DICVIV_02218
MRISGFIVFITCIVAWTFAAPVSDIEAAYRILNKYLERFGSDELQNAYLIRDNG
>FLP-21_Diploscapter_coronatus_DCO_025274 peptide: DCO_025274
MRSSCRFLVALLAFFLSWVASLPLSDNEAAQLFGRYLQRFSGSEPSGYYYEHESNKRGLG
PRPLRFG
>FLP-21_Haemonchus_contortus_HCON_00158810-00001 peptide:
HCON_00158810-00001
MRVSGFLVFLACIVAWAFATPVSDTEAAYRILNKYLQRFGPDDLQDVYLYGDHGSNKRGL
GPRPLRFG
>FLP-21_Haemonchus_placei_HPLM_0001769801-mRNA-1 peptide:
HPLM_0001769801-mRNA-1
MRVSGFLVFLACIVAWAFATPVSDTEAAYRILNKYLQRFGPDDLQDVYLYGDHGSNKRGL
GPRPLRFG
>FLP-21_Heligmosomoides_polygyrus_HPOL_0000400701-mRNA-1 peptide:
HPOL_0000400701-mRNA-1
MRISGFIVFIACIVAWAFAAPVSDTEAAFRILNKYLQRFGGDDLPDVYLVGEHGSNKRGL
GPRPLRFG
>FLP-21_Nippostrongylus_brasiliensis_NBR_0000622101-mRNA-1 peptide:
NBR_0000622101-mRNA-1
MEERSRAVSETPSAINPLDIAESPKPYLILRMRISGIIVFLACIIAWAFAAPVSDTEAAY
RILNKYLQRFGSDDLQDVYLIGDHGSAPFIMRPPTASTSDYSYNSHAFGLA
>FLP-21_Oesophagostomum_dentatum_OESDEN_18901 peptide: OESDEN_18901
MRVSGFLVFIACIIAWAFATPVSDTETAYRILNKYLQRFGPDDLQDVYLVGDHG

```
>FLP-21_Oscheius_tipulae_OTIPU.nOt.2.0.1.t10058 peptide:
OTIPU.nOt.2.0.1.t10058
MRATSVLLFFACLVACALAVPVSDTEAAFRILNKYLSRFNNDDLQELYYGADHGSIKRGL
GPRPLRFG
>FLP-21_Strongylus_vulgaris_SVUK_0001260801-mRNA-1 peptide:
SVUK_0001260801-mRNA-1
MSTRDLLILKGMRVSGFLVFIACIIAWAFASPVSDTEAAYRILNKYLQRFGGDDLQDVYL
VGDHGIVVEHEHLMSGFLMSL
>FLP-21_Ancylostoma_duodenale FLP-21 predicted
SPMNSYQCPLMLEGGGGRELFTPTPFSPVCIYIFATQVQALDASLRLPRVNASIRVLTVGKATAGSCG
VFTCLPLAIPMMHGASVGRIFRKFSKIHDFRSIKRGLGPRPLRFG
>FLP-21_Angiostrongylus_cantonensis FLP-21 predicted
RLAHADLLWTLSVDVWSVNIYSSCNRFNFLNLNRSHCRSDKRGLGPRPLRFG
>FLP-21_Caenorhabditis_sinica FLP-21 predicted
AVVQRSVGGRSVCERLLSINTYYKNNNPGHSEFFNLRFSNPCYSFRSMKRGLGPRPLRFG
>FLP-21_Caenorhabditis_tropicalis FLP-21 predicted
TIFYSILISFFRSMKRGLGPRPLRFG
>FLP-21_Cylicostephanus_goldi FLP-21 predicted
RHLEKSSFCEIWKVLMHFLKTHDFTATLCFHFSSIKRGLGPRPLRFG
>FLP-21_Diploscapter_pachys FLP-21 predicted
TNRVSFRSNKRGLGPRPLRFG
>FLP-21_Heterorhabditis_bacteriophora FLP-21 predicted
AKQCDDSIFRSIKRGLGPRPLRFG
>FLP-21_Mesorhabditis_belari FLP-21 predicted
AMNQIIGTKHNKAQQTRALLENIVTSIFRSNNKRGHGPRPLRFG
>FLP-21_Necator_americanus FLP-21 predicted
IDVGTSPLHSTLVLSIELLRKFTNFRSYKRGLGPRPLRFG
>FLP-21_Teladorsagia_circumcincta FLP-21 predicted
TCFRSNKRGLGPRPLRFG
>FLP-21_Bursaphelenchus_xylophilus_BXY_0389500.1 peptide:
BXY_0389500.1
MRTDALLKVAIFLFLAILAQLGQAGPILESTNDLTALKMLRNYMERYGEPDGDVGYVVLD
DMRSNKRGMGPRPLRFG
>FLP-21_Halicephalobus_mephisto_MSTRG.13615.1.p1 peptide:
MSTRG.13615.1.p1
VTLHDFVEFAFLRKLLVTQKLLLTKDIITSQDTRETLETMALVCRTQLYPLFLVILATLI
LLMQASEAAPVSDSANAYRILSSYMRQFNDPELDAAGYIVFDDKSSIKRALGPRPLRFG
>FLP-21_Panagrellus_redivivus_Pan_g12093.t1 peptide: Pan_g12093.t1
MSSVASSRFTTLIVATLIAAIALLAVVVPYSEAAPTSDAANAYRILSNYMKQFDDRELDS
AGYIVFDDKSSIKRALGPRPLRFG
>FLP-21_Parastrongyloides_trichosuri_PTRK_0000369900.1 peptide:
PTRK_0000369900.1
MVKRFVSCLFLIGMIILISARNINSSSLEGNGNILLKQYREVPGDEYETRPFYYNGYLIE
NVDPRISYKRAVGPRPLRFG
>FLP-21_Steinernema_carpocapsae_L596_g19959.t1 peptide:
L596_g19959.t1
MVAVVRLLLAAFLVLFSVFVAPARSAPTSVDDQQYRLLSRYLSQFNAPEYDPSGYMYFDQ
RSMKRGLGPRPLRFG
>FLP-21_Steinernema_feltiae_L889_g32029.t1 peptide: L889_g32029.t1
MGDYKGAAIRQLKLFRVNYAMVAFVRLLLAVFFLLVSVFAIPSLSVPTSDSDQQYQLLSR
YLAQYNTPFDLDGYNMYAEPRSIKRGLGPRPLRFG
>FLP-21_Steinernema_glaseri_L893_g17088.t1 peptide: L893_g17088.t1
MVAVVRLLLAVFFVLVSVLAMPSLAAPISDAEQQYRLLSRYMNQYNAPEFDLNGLLFADQ
RSIKRGLGPRPLRFG
```

>FLP-21_Steinernema_monticolum_L898_g16109.t1 peptide:
L898_g16109.t1
MVAIVRFLLAVFFVLVSICSVPSLSAPTSDSEQQYRLLSRYLNQFNTPFELDGYGMYAEP
HSIKRGLGPRPLRFG
>FLP-21_Steinernema_scapterisci_L892_g23780.t1 peptide:
L892_g23780.t1
MVAIVRLLLAAFLVLVSVFAIPARSAPTSVDDQQYRLLSRYLSQFNAPEYDPSGYMVLDQ
RSMKRGLGPRPLRFG
>FLP-21_Strongyloides_papillosus_SPAL_0000319300.1 peptide:
SPAL_0000319300.1
MAKYFLSCLCLIGAIILLSSHCISTIPIDDSGNFLIRQLRDFPVEEFDGRPIYYNGYLIE
NPDSRISFKRAVGPRPLRFG
>FLP-21_Strongyloides_ratti_SRAE_2000441800.1 peptide:
SRAE_2000441800.1
MVKYFSSCLCLIGAIIILSSHYTNTIPIENSENYFIRQLRDFPVEEFDGRPIYYNGYLIE
NTDPRISLKRAVGPRPLRFG
>FLP-21_Strongyloides_stercoralis_SSTP_0000420600.1 peptide:
SSTP_0000420600.1
MVKYFPYCLCLIGIIIILSSNYTYTIPIDNNENYFIRQLREFPVEEFVGRPIYYDGYFIE
NADPRISFKRAVGPRPLRFG
>FLP-21_Strongyloides_venezuelensis_SVE_1580900.1 peptide:
SVE_1580900.1
MAKYFLSCLCLIGVIILLSSHCISTIPIDDSGNFLIRQLREFPVEEFDGRPIYYNGYLIE
NPDSRISFKRAVGPRPLRFG
>FLP-21_Acrobeloides_nanus_ACRNAN_scaffold2000.g28499.t1 peptide:
ACRNAN_scaffold2000.g28499.t1
MMSLKWILSLFVILGISIRAFAAPPSDSENAYRILSNYVNRFNEPEFNFDGFLFADGRSM
KRGLGPRPLRFG
>FLP-21_Ditylenchus_destructor_Dd_00458 peptide: Dd_00458
MRNSAKVMQTLPNNDEDDLTAIQSNRQKASFAFAIPVSSILIIAMICLLLTFSSANAAPT
GTDITSSEEMRSYLKSIDQLPSPYELAHLDIAGSSDGGTSGYVIIEDLKSNKRGLGMGPR
PLRFG
>FLP-21_Globodera_rostochiensis_GROS_g00870.t1 peptide:
GROS_g00870.t1
MYVTGLIWTLYPLLGFFAVLEFTVGLAHVFLCLPFAHFYLSFWSGISAAITSVYALLLDY
PNKCELLLQFVSAFFAFGLSFTALIENVCIRKVHLSTDQLSFCAGLLNRTATKQMQCDRI
LGHLQMHLLQKFSSEPVEQSLHSVRLFVSSLISFCAIAQFFSGLIVLLLSTIHSHYCAPM
FFTNIVPLVGLYSLIFALFPRVSPNSRFAFRQFLAIVGVALATALASICSFSFFCWANRQ
NRTIGLLPYRTKSEGILLDLRRGNEGDGFGVLRFCRMPERTYEHCERVLDFSFPYLDWPM
EQVENEKAVVRIVLHSLLSVCAIGLVSLFMSDAF*KCQTFRMSDRIVPLLLGLSVFVSLS
VALPAPSLQQMPKIRTVFGRIGEILPAGAPSESQIAETPFNTLVNYEEQLNDWNGGGEQP
PASYFLFPEVKRQINSKRGSLGPRPLRFGRR
>FLP-21_Ditylenchus_dipsaci FLP-21 predicted
VMFLVKNYLMNLERLQVPHEDTDYANGYVIIDDLKYVFYSTFHTKLNFSLKICCRSNKRGLGPRPLRF
G
>FLP-22_Plectus_sambesii_PSAMB.scaffold844size40337.g9100.t1
peptide: PSAMB.scaffold844size40337.g9100.t1
MVSMRCKSGLTMLAVVCLLLAVTVRAQPVATDDADVMAEYSPDALSREGRAASGMKWMRF
GKRAQQVKWMRFGKRAQNVKWMRFGKRDGDSQIDSEQQ
>FLP-22_Anisakis_simplex_ASIM_0001017501-mRNA-1 peptide:
ASIM_0001017501-mRNA-1
MTDMDDMNRAIRASNMKWMRFGKRSPSVKWMRFGKRAPNVKWMRFGKRSSSPSSSSRLVD
NFDEFEQ
>FLP-22_Ascaris_lumbricoides_ALUE_0000551301-mRNA-1 peptide:

ALUE_0000551301-mRNA-1
MCCCSQRGQVDTMDGSHITLLLSLLVAFLVATSAQPPLYDGTEERNPTLQQQSGDISDSH
VYSVMSRTVVTGNLVEYSWCGFLPATDDMNRAIRASNMKWMRFGKRSPNVKWMRFGKRAP
NMKWMRFGKRSPSMEGTVDQFEQ
>FLP-22_Ascaris_suum_GS_20131 peptide: GS_20131
MDGSHITLLLSLLVAFLVATSAQPPLYDGTEDDMNRAIRASNMKWMRFGKRSPNVKWMRF
GKRAPNMKWMRFGKRSPSMEGTVDQFEQ
>FLP-22_Brugia_malayi_Bm3942.1 peptide: Bm3942.1
MNFSQSKIIFALFIASTAFILTSSMHLDPLIATSDNDDDDFIRTLRVPNAKWMRFGKRLP
NAKWMRFGKRAQTAKWMRFGKRNDDFYDY
>FLP-22_Brugia_pahangi_BPAG_0001069401-mRNA-1 peptide:
BPAG_0001069401-mRNA-1
MNFSQSKIIFALFIASTAFILTSSMHLDPLIATSDNDDDDFIRTLRVPNAKWMRFGKRLP
NAKWMRFGKRAQTAKWMRFGKRNDDFYDYQ
>FLP-22_Dirofilaria_immitis_nDi.2.2.2.t06146 peptide:
nDi.2.2.2.t06146
MQLNESKLIITLFIASMALILTASMDVNPLIVASDNDYDFIRNLRTPNIKWMRFGKRLPN
TKWMRFGKRVPTAKWMRFGKRNDDFYDSQ
>FLP-22_Dracunculus_medinensis_DME_0000592101-mRNA-1 peptide:
DME_0000592101-mRNA-1
MDTIRDGALLKFEGVTDDLERNHRTSGVKWMRFGKRSPSVKWMRFGKRAPNVKWMRFGRS
DLAMDNQEGNDS
>FLP-22_Elaeophora_elaphi_EEL_0000990001-mRNA-1 peptide:
EEL_0000990001-mRNA-1
MRSLRAPNTKWMRFGKRLPNAKWMRFGKRAPTAKWMRFGKRNDDFYDSQ
>FLP-22_Litomosoides_sigmodontis_nLs.2.1.2.t04942-RA peptide:
nLs.2.1.2.t04942-RA
MQQITDANSRNPNHFPFEKCWRKRWCTKTEMKLFSESTIIFAFFTASMIFVIASSVRVEP
LLVAADNDDDFIRSLRTPNTKWMRFGKRLPNAKWMRFGKRMENAKWMRFGRQNDDFYDSQ
>FLP-22_Loa_loa_EFO22085.1 peptide: EFO22085.1
MKFSRLEIIIALLITSATSVLTTSVGANPLLDVLDYDDDFIRNVRTGNVKWMRLGKRLPD
HKWMHSGRKRAQNVKWMKFGKQNDNFYDYQ
>FLP-22_Onchocerca_ochengi_OOCN_0000580801-mRNA-1 peptide:
OOCN_0000580801-mRNA-1
MAFILAASMNVNPSLVAMDNDDDLHRNLRTVNTKWMRFGKRLPNTKWMRFGKRAQTTKWM
RFGKRNDDFYDSQ
>FLP-22_Onchocerca_volvulus_OVOC176.1 peptide: OVOC176.1
LFIVSMAFILAASMNVNPSLVAMDNDDDLHRNLRTVNTKWMRFGKRLPNTKWMRFGKRAQ
TTKWMRFGKRNDDFYDSQ
>FLP-22_Parascaris_equorum_PEQ_0001370401-mRNA-1 peptide:
PEQ_0001370401-mRNA-1
MDGSHITLLLSLLVALLVATSAQPPLYDDGTEGIAEKNCFQDDMNRAIRASNVKWMRFGK
RSPNVKWMRFGKRASNMKWMRFGKRSPSMEGTVNQFDQ
>FLP-22_Parascaris_univalens_PgR035X_g025_t01 peptide:
PgR035X_g025_t01
MDGSHITLLLSLLVALLVATSAQPPLYDDGTEDDMNRAIRASNVKWMRFGKRSPNVKWMR
FGKRASNMKWMRFGKRSPSMEGTVNQFDQ
>FLP-22_Thelazia_callipaeda_TCLT_0000461301-mRNA-1 peptide:
TCLT_0000461301-mRNA-1
MVALLVFTFTAPTTAQSSIAMDTDGDDDEFLRELRAPNAKWMRFGKRTPNAKWMRFGKRR
PNAKWMRFGKRNDDFYDIQ
>FLP-22_Toxocara_canis_TCNE_0001211201-mRNA-1 peptide:
TCNE_0001211201-mRNA-1
MNGAHITFLLSLLVALLVATSAQPFLNDGVDDDMNRVLRGSNMKWMRFGKRSPNVKWMRF

GKRAPNVKWMRFGKRFSPIEGIDELEQ
>FLP-22_Wuchereria_bancrofti_WBA_0000952901-mRNA-1 peptide: WBA_0000952901-mRNA-1
MKFSQSKIIFALFIASAAFILTSSMHLDPLVATPDNDDDDFTRTLNVPNAKWMRFGKRLP
NAKWMRFGKRAQTAKWMRFGKRDDDFYDYQ
>FLP-22_Acanthocheilonema_viteae FLP-22 predicted
NCKIKLKLFQDYFTKNLHTPNTKWMRFGKRLPNAKWMRFGKKAPTAKWMRFGKRNDDFYDSE
>FLP-22_Gongylonema_pulchrum FLP-22 predicted
NNKLATFALTTNNFVFQDEFVRYLRAPNAKWMRFGKRAPNAKWMRFGKRAPNAKWMRFGKRSDEFYDS
Q
>FLP-22_Onchocerca_flexuosa
DDLRRNLRAVGTKWMRFGKRLPNTKWMRFGKRAQTTKWMRFGKRSDDFYDSQ
>FLP-22_Ancylostoma_caninum_ANCCAN_09946 peptide: ANCCAN_09946
MVTSGTREENVDFSAILLATNSLGQSLQPPYHYFGRRETQPAHISYYSWIFRRLSKGQRS
LNISWGTVPDFFYDAYNNFFGSKNAPSFKRTPSAKWMRFGKRSPNAKWMRFGKRSPEAKW
MRFGKRSDYEFEGDDDLY
>FLP-22_Ancylostoma_ceylanicum_Acey_s0011.g1541.t2 peptide: Acey_s0011.g1541.t2
MQRLVALVLVCLLAVACTAQDVVEDLPVVRARSLNISWGTLPQFFHSAYSIFIGNKNAPS
FKRTPSAKWMRFGKRSPNAKWMRFGKRSPEAKWMRFGKRSDYEFEGDDDLY
>FLP-22_Ancylostoma_duodenale_ANCDUO_16283 peptide: ANCDUO_16283
MVLLAISSLERLFQPPYHYFGRRETQPTHISYYSWIFRRLSEGQNAPSFKRTPSAKWMRF
GKRSPNAKWMRFGKRSPEAKWMRFGKRSDYEFEGDDDLY
>FLP-22_Angiostrongylus_cantonensis_ACAC_0000004301-mRNA-1 peptide: ACAC_0000004301-mRNA-1
MQRLVAFLMICNMLVAVCTSQETDVLPMVREERALKRTPSAKWMRFGKRSPNTKWMRFGK
RSPEAKWMRFGKRGYYDFDGYDDTE
>FLP-22_Angiostrongylus_costaricensis_ACOC_0000814201-mRNA-1 peptide: ACOC_0000814201-mRNA-1
MRFGKRSPDAKWMRFGKRSPEAKWMRFGKRGDYEFDGYDDTE
>FLP-22_Caenorhabditis_brenneri_CBN12719.1 peptide: CBN12719.1
MNRSLICVLLLASLVASQVIDMDGQLTGLVQNDALLMEPQDKRSPSAKWMRFGKRSPSAK
WMRFGKRSPSAKWMRFGKRDGADAEQDY
>FLP-22_Caenorhabditis_briggsae_CBG25200.1 peptide: CBG25200.1
MNRSVICVVLLVSLVSAQVFDLDGQLAALEQQDGALLIEPQVKRSPSAKWMRFGKRSPSA
KWMRFGKRSPSAKWMRFGKRSGAEAIAPEQDY
>FLP-22_Caenorhabditis_inopinata_Sp34__10300700.t1 peptide: Sp34_10300700.t1
MNRSMIALCVVLMVSLVSAQVFDLDDQQLAGLEQNDAILIEPHIKRSPSAKWMRFGKRSP
SAKWMRFGKRSPSAKWMRFGKRSGAEEVAEQDY
>FLP-22_Caenorhabditis_japonica_CJA05306.1 peptide: CJA05306.1
MNKFITATALVLMIALVSAQFNSDSQQEGPIEESSVLRNWETKRSPSAKWMRFGKRSPPA
KWIRFGKRSPNAKWMRFGKRSGEIDSVYEQ
>FLP-22_Caenorhabditis_latens_FL83_06399 peptide: FL83_06399
MNRSMICILLLVSLVSAQVFDLDGQQLAGIEQNDALLIEPHIKRSPSAKWMRFGKRSPSA
KWMRFGKRSPSAKWMRFGKRSGAEGVAEQDY
>FLP-22_Caenorhabditis_nigoni_Cni-flp-22.3 peptide: Cni-flp-22.3
MNRSVICVLLLVSLVSAQVFDLDGQLAALEQQDGALLIEPQVKRSPSAKWMRFGKRSPSA
KWMRFGKRSPSAKWMRFGKRSGAEAVAPEQDY
>FLP-22_Caenorhabditis_remanei_CRE03845.1 peptide: CRE03845.1
MNRSMICILLLVSLVSAQVFDLDGQLAGIEQNDALLIEPHIKRSPSAKWMRFGKRSPSAK
WMRFGKRSPSAKWMRFGKRSGAEGVAEQDY
>FLP-22_Dictyocaulus_viviparus_DICVIV_08883 peptide: DICVIV_08883
MNDTCKKLRTSLLLAFLRLQTSILVLPERVPKRSPSAKWMRFGKRTPNAKWMRFGKRTPD

AKWMRFGKRAYEYEGYDDME
>FLP-22_Diploscapter_coronatus_DCO_024491 peptide: DCO_024491
MQQLMVGLLCSLLVSLALSQSQDNDDSAVLIADEIASQPQDTAAETTFDKRSPSAKWMRF
GKRSPSAKWMRFGKRSPSAKWMRFGKRSDGISNFYEMEY
>FLP-22_Diploscapter_pachys_WR25_08245.1 peptide: WR25_08245.1
MQQLMAGLLCSLLVSLALSQSQDNDDNAVLIADEISNQPQDTAAETTLDKRSPSAKWMRF
GKRSPSAKWMRFGKRSPSAKWMRFGKRSDGVSNFYEMEY
>FLP-22_Haemonchus_contortus_HCON_00012150-00001 peptide:
HCON_00012150-00001
MQRLLALVMVCLLVAVCTAQDVEEIPVGRVERALKRTPSAKWMRFGKRSPNAKWMRFGKR
TPDAKWMRFGKRGEYEFDGYEDME
>FLP-22_Haemonchus_placei_HPLM_0001277501-mRNA-1 peptide:
HPLM_0001277501-mRNA-1
MQVTGNSYKIFQERALKRTPSAKWMRFGKRSPNAKWMR
>FLP-22_Heligmosomoides_polygyrus_HPBE_0000438401-mRNA-1 peptide:
HPBE_0000438401-mRNA-1
LEEFSLKSLKACEGALKRTPSAKWMRFGKRSPNAKWMRFGKRTPDAKWMRFGKRSDYGLE
GYDDME
>FLP-22_Heterorhabditis_bacteriophora_Hba_08014 peptide: Hba_08014
MIDAARNSSMTIVKKHQTSVYQAVRIWHRPLRRKSHARIEYPAYLVKVHLVSMQRLAALA
LLFTILVAVCRSEDIEDAVIQEEAINKRSPSAKWMRFGKRSPNAKWMRFGKRSPDAKWMR
FGKRADYAYEEDY
>FLP-22_Mesorhabditis_belari_mbelari.g25745.t1 peptide:
mbelari.g25745.t1
MWPSMRVLVLLVACLSCALMAIDYEQGTQQLNDKRNEDFEDLAKRSPSAKWMRFGKRSPN
AKWMRFGKRSPSAKWMRFGKRSTDDLPYDSIDY
>FLP-22_Micoletzkya_japonica_MicoRS5524-mkr-S40-2.45-mRNA-1 peptide:
MicoRS5524-mkr-S40-2.45-mRNA-1
MSRVIIATILLGLIVALTSATEDEMTDALVENWKRSPSAKWMRFGKRSPSAKWMRFGKRS
PSAKWMRFGKRSAAVDALEEDDY
>FLP-22_Necator_americanus_NECAME_00821 peptide: NECAME_00821
MYWWQVHRNVPSFKRTPSAKWMRFGKRSPNAKWMRFGKRSPEAKWMRFGKRSDYESDDDL
Y
>FLP-22_Nippostrongylus_brasiliensis_NBR_0000230201-mRNA-1 peptide:
NBR_0000230201-mRNA-1
MDVLLNKDMLAKQQRKLRRGKEKTVDLCFQERSLKRTPSAKWMRFGKRSPNAKWMRFGKR
TPDAKWMRFGKRGDYEFEGYEDLE
>FLP-22_Oesophagostomum_dentatum_OESDEN_08469 peptide: OESDEN_08469
MSSLSLTLAWSSIAAAVPPFPGSYAYVPSFKRTPSAKWMRFGKRSPNAKWMRFGKRSPEA
KWMRFGKRSDYEFEGEDDLY
>FLP-22_Oscheius_tipulae_OTIPU.nOt.2.0.1.t05601 peptide:
OTIPU.nOt.2.0.1.t05601
MNRLVLVALFCALLAVAQAQDSQYNVEDAASPVDELDKRSPNAKWMRFGKRSPNAKWMRF
GKRQNTKWMRFGKRGGLELFELPADY
>FLP-22_Parapristionchus_giblindavisi_Parapristionchus-mkr-
S_93-1.29-mRNA-1 peptide: Parapristionchus-mkr-S_93-1.29-mRNA-1
MSRLIIVIFTILLLSLAVSSQEESSSSLVGSWKRSPSAKWMRFGKRSPNAKWMRFGKRAP
SDKWMRFGKRSEDGAIFDDIEY
>FLP-22_Pristionchus_arcanus_Pristionchus_arcanus_arcanus-mkr-
S_158-1.45-mRNA-1 peptide: arcanus-mkr-S_158-1.45-mRNA-1
MVRLVAFFFLALVTLSFSLESSDSSIENWKRSPSSKWMRFGKRSPNAKWMRFGKRAPSDK
WMRFGKRAALMEDDVEY
>FLP-22_Pristionchus_entomophagus_entomophagus-mkr-S35-7.60-mRNA-1
peptide: entomophagus-mkr-S35-7.60-mRNA-1

MARLLVLFFLALLSISFSMESSDFSIENWKRSPSSGKWMRFGKRSPSNKWMRFGKRSPNS
KWMRFGKRSAITRDDVEY
>FLP-22_Pristionchus_exspectatus_exspectatus-mkr-S_27-3.71-mRNA-1
peptide: exspectatus-mkr-S_27-3.71-mRNA-1
MVRLVAFFFLALVTLSFSLESSDSSIENWKRSPSSKWMRFGKRSPNAKWMRFGKRAPSDK
WMRFGKRPALMEDDVEY
>FLP-22_Pristionchus_fissidentatus_fissidentatus-mkr-S150-5.15-
mRNA-1 peptide: fissidentatus-mkr-S150-5.15-mRNA-1
MSRLFVLFFLALITLSFSQESSDAAIESWKRSPSAKWMRFGKRSPNAKWMRFGKRSPSDK
WMRFGKRSAPLMEDDVEY
>FLP-22_Pristionchus_japonicus_japonicus-mkr-S439-0.3-mRNA-1
peptide: japonicus-mkr-S439-0.3-mRNA-1
MARLVVLFFLALVTLSFSMESSDSSIENWVSIKRSPSSSKWMRFGKRAPSDILSETKRSP
NAKWMRFGKQLGEKLGIRQLEAVSVI
>FLP-22_Pristionchus_maxplancki_maxplancki-mkr-S1006-0.4-mRNA-1
peptide: maxplancki-mkr-S1006-0.4-mRNA-1
MARLVVFFFLALLTLSFSMESSESSIENWKRSPSSKWMRFGKRSPNAKWMRFGKRAPSDK
WMRFGKRSALMENDAEY
>FLP-22_Pristionchus_mayeri_mayeri-mkr-S490-0.8-mRNA-1 peptide:
mayeri-mkr-S490-0.8-mRNA-1
MARLAVLFFLALLTLSFSFESSDSYIENWKRSPSSKWMRFGKRSPNAKWMRFGKRAPSDK
WMRFGKRSAVGDVEY
>FLP-22_Pristionchus_pacificus_PPA35627.1 peptide: PPA35627.1
MVRLVAFFFLALVTLSFSLESSDSSIENWKRSPSSKWMRFGKRSPNAKWMRFGKRAPSDK
WMRFGKRAALMEDDVEY
>FLP-22_Teladorsagia_circumcincta_TELCIR_01869 peptide: TELCIR_01869
MLEITKSSVKKLRSRDRMQRLLALVMVCLLVAVCTSQDVDDLPVERALKRTPSAKWMRFG
KRSPNAKWMRFGKRTPDAKWMRFGKRGEYEFDGYEDLE
>FLP-22_Caenorhabditis_elegans_F39H2.1.1 peptide: F39H2.1.1
MNRSMIALCVVLMVSLVSAQVFDLDGQQLAGLEQNDARLMEQQVKRSPSAKWMRFGKRSP
SAKWMRFGKRSPSAKWMRFGKRSGAEAVSEQDY
>FLP-22_Caenorhabditis_sinica FLP-22 predicted
ISFFQQVKRSPSAKWMRFGKRSPSAKWMR
>FLP-22_Caenorhabditis_tropicalis FLP-22 predicted
SVIKLYLTLTLKTAPSTGKRRKLGILFSKGSSYNPILLTNKMTNCFFLERKKIRSTINRIIFRFGKRS
PSAKWMRFGKRSGAEEVAEQDY
>FLP-22_Strongylus_vulgaris FLP-22 predicted
VYTYQIVYQTINCFKYAPSFKRTPSAKWMRFGKRSPNAKWMR
>FLP-22_Bursaphelenchus_xylophilus_BXY_0887600.1 peptide:
BXY_0887600.1
MGTRFRFRGLNARGRVGRPLSCPNLISCLIVYLPIEVRREGGYDLDTTFALCPAPQAQQK
IFVFYGFCRVWIMVAPMSLLFAFLLVVGHFLGQAQVVSGSDIYQDNLYREARAPAMKWMR
FGKRSPQGKWMRFGKRAPQGKWMRFGKRADDVQDPEYE
>FLP-22_Halicephalobus_mephisto_MSTRG.14074.2.p1 peptide:
MSTRG.14074.2.p1
VYKDFESDFIFQFPLGHVSRMTRVSGLVVFSTVLLALLVLFADAKPYGFDLSSSEEDLFR
QVRAPGVKWMRFGKRTPNAKWMRFGKRAPSAKWMRFGKRSDDEQFDEELYKNLF
>FLP-22_Panagrellus_redivivus_Pan_g19100.t1 peptide: Pan_g19100.t1
MTRFSSYLAIVAVLLSVLVLFADAWDFSSEEDFPRAVRTPGVKWMRFGKRAPNAKWMRFG
KRAPNAKWMRFGKRSDGAANEVPAASESEFAPNVPYF
>FLP-22_Steinernema_carpocapsae_L596_g6927.t1 peptide: L596_g6927.t1
MNTSWTVAVFALFFACLLACSDAAFFDVFSPQLAPMDNMDRAVRAPNVKWMRFGKRAPAA
KWMRFGKRAPAAKWMRFGKRSDSQENFGFPAEAEYNTL
>FLP-22_Steinernema_feltiae_L889_g25784.t1 peptide: L889_g25784.t1

MNTSWSVAVCALLFACLLSCSDAAFFDVFSPQLAPADSMDRAVRAPNVKWMRFGKRAPAA
KWMRFGKRAPAAKWMRFGKRSDQQEGFGGGFAPSVEEYSNL
>FLP-22_Steinernema_glaseri_L893_g2260.t1 peptide: L893_g2260.t1
MPCTQSCVPTRNLGIPDPTSRPDPEASGIPTRNPVPIPKLRDSRPEIPSRSRNFGIPDPK
SRFPNHQSTYLANPKYYISKKALTKTNLTPIMHAWCYKNEDISFTLGCSQLNHVPNPEAS
GIPTRNPVPSRNRDRDTALHVPYCVLLYCACGSWQKAEEVALSSYYTVTLPDQLTHYGTT
SANTPFFCLSHEVVLHPFDKRNKRQSVRASTTTPPSCHHLCGPLNSESTRPTQPTRIVNF
LRCGRLEEAVNAKIKCEIRTERVIPVYSHGTRMTWWEAPPLGGGVSRARFYERAVVGGEL
SMWPGAGTAFRICGSCLISRVSVFSAVVVVCDGSSRTIFVAKSSSVRFCLRPSNMNTSWS
VAVFALFFACLLAFSDAAFFDVFSPQLASDNMDRAVRAPNVKWMRFGKRAPAAKWMRFGK
RAPAAKWMRFGKRSADAQDGFAAGDEYTL
>FLP-22_Steinernema_monticolum_L898_g15669.t1 peptide:
L898_g15669.t1
MNTSWSVAVIALLFACLFAFSNAAFFDVFSPQLAPADSMDRAVRAPNVKWMRFGKRAPAA
KWMRFGKRAPAAKWMRFGKRSGDSEDVGNFGYPVADEYTS
>FLP-22_Steinernema_scapterisci_L892_g9483.t1 peptide: L892_g9483.t1
MNTSWTVAVFALLFACLLACSDAAFFDVFSPQLAPADNMDRAVRAPNVKWMRFGKRAPAA
KWMRFGKRAPAAKWMRFGKRSDSQENFGFPADAEYNTL
>FLP-22_Strongyloides_papillosus_SPAL_0000781800.1 peptide:
SPAL_0000781800.1
MTNFFNLKIVFLFTIALNIFSNFVTPFDVSSDIEGDDNLDFTRVVRAPNVKWMRFGKRAA
YYGIGDKRSPQVKWMRFGKRSDMVNENEIY
>FLP-22_Strongyloides_venezuelensis_SVE_0426600.1 peptide:
SVE_0426600.1
MTNFFNLKIVFLFTIALNIFSNFVTPYDVSSDIEGDDSLDFTRVVRAPNVKWMRFGKRAP
YYGISDKRSPQVKWMRFGKRSDLVNENEIY
>FLP-22_Strongyloides_ratti FLP-22 predicted
TIFFFRFGKRGSYYNDYDKRSPQVKWMRFGKRSDVLNEFQNY
>FLP-22_Strongyloides_stercoralis FLP-22 predicted
GSIMLFFLFLKYIIFRFGKRGSYYNNYDKRSPQVKWMRFGKRYDTSNEIQNY
>FLP-22_Acrobeloides_nanus_ACRNAN_scaffold3183.g32350.t1 peptide:
ACRNAN_scaffold3183.g32350.t1
MAKIAWYYSLFFLIVALAVLFADAAPNADFSEDDIFRVARAPQAKWMRFGKRAPQAKWMR
FGKRAPQAKWMRFGKRDPTFEGEYEVEQ
>FLP-22_Ditylenchus_destructor_Dd_13527 peptide: Dd_13527
MSSVNSSSHSGSVYLRLYSSLFLWVFLATVLVALMQTSNASPPTRFKGAAGQGELFVPSQ
SYYFPASTEDMNRAVRAPGMKWMRFGKRAPNAGKWMRFGKRAPNGGKWMRFGKRTPQDTN
SYGWPSTTR
>FLP-22_Globodera_rostochiensis_GROS_g00946.t1 peptide:
GROS_g00946.t1
MMAASAVLPNTLSPQFSFRWHFLPLLLLALLTISDFASCHSVPTLIAFDPAAYAKLRRYT
PIQHRRGGGRLCAFRSNFCEQQASFLSDDEAAFERAARQPAGGVKWMRFGKRTPQGKWMR
FGKRTMATEGGKWVRFGKRAEEMQNDQ
>FLP-22_Heterodera_glycines_Hetgly.G000002398 peptide:
Hetgly.G000002398
MNASSNWLILCALPSLRPPIPSAIAVTPSSSSSSRPTHRNLPHCLNFFHTLKKGSGRIVR
YSSLFPSIASPIKMVVVLSALSSGTFPPHFSLLCFSADFTNCHSVPSLMAFDPTKLRRLM
PIREFLSDDQLAFERAIRQPAGGVKWMRFGKRTPQGKWMRIGKRKMAIEGGKWVRFGKRA
EEMQGEE
>FLP-22_Meloidogyne_arenaria_M.Arenaria_Scaff387g008957 peptide:
M.Arenaria_Scaff387g008957
MSMDEFDNTDPFYRAARENGVKWMRFGKRVPQQSKWMRFGKRAPSSGKWMRFGKRNSEFM
EEK
>FLP-22_scaffold3339_cov182.g5406 peptide: scaffold3339_cov182.g5406

MLISIILTFLLLINNKNSQVYATQEQQQFSRQVRPYSERAANQREAIAEDVLVGIDSIFA
SPYPKLNLLDGSMSMDEFDNTDPFYRAARENGVKWMRFGKRVPQQSKWMRFGKRAPSSGK
WMRFGKRNSEFMEEK
>FLP-22_Meloidogyne_graminicola_NXFT01000446.1.103_g peptide:
NXFT01000446.1.103_g
MQQKQNKLIKNSSLLFSTKLNKFMLIFNILIIFLLINNEIIQINANQQIKPLKEFNQNLL
DGSMSIDEFDNNDPFYRAARENGVKWMRFGKRVPQQSKWMRFGKRAPSTGKWMRFGKRLS
NDFIENN
>FLP-22_Meloidogyne_hapla_MhA1_Contig396.frz3.gene2 peptide:
MhA1_Contig396.frz3.gene2
MLISIPLIFLLISNENLQVNAIQKQQFSRQVRPYSERATNQNLLDGSMSMDEFDNTDPFY
RAARENGVKWMRFGKRVPQQSKWMRFGKRAPSSGKWMRFGKRNSEFMEEK
>FLP-22_Meloidogyne_incognita_Minc3s00616g15157 peptide:
Minc3s00616g15157
MLISIILTFLLLINNECQVSASQEEQQFSRQVRPYAERAANQNLLDGSMSMDEFDNTDPF
YRAARENGVKWMRFGKRVPQQSKWMRFGKRAPSSGKWMRFGKRNSEFMEEK
>FLP-22_scaffold13611_cov178.g16967 peptide:
scaffold13611_cov178.g16967
MLISTILTFLLLISNECQVSAAQEEQQFSRQVRPYADRAANQNLLDGSMSMDEFDNTDPF
YRAARENGVKWMRFGKRVPQQSKWMRFGKRAPSSGKWMRFGKRNSEFMEEK
>FLP-22_Meloidogyne_floridensis_maker-nMf.1.1.scaf04110-augustus-
gene-0.1-mRNA-1 peptide: maker-nMf.1.1.scaf04110-augustus-gene-0.1-
mRNA-1
NKGRSNVRTTIREHNPQVKSKFINIFKRAPSSGKWMRFGKRNSEXXXXQKIRAPSSGKWM
RFGKRNSEFMEEKLSKQISNKLNKL
>FLP-22_Ditylenchus_dipsaci FLP-22 predicted
IKERLMSSSLLNLLFQNYYWLQPSEDDLTRTARAPGMKWMRFGKRAPNAGKWMRFGKRAPNAGKWMRF
GKRAPESDQF
>FLP-23_Brugia_malayi_Bm14130.1 peptide: Bm14130.1
MKFSLSFILFFLMITTSMTLRGALFRSRKAALESIIRANTGRQDFLRFGRTNEKYNDRHL
IASALLTPVDLK
>FLP-23_Onchocerca_ochengi_OOCN_0000871901-mRNA-1 peptide:
OOCN_0000871901-mRNA-1
MVTTSMELRGALFRSRKTALDSIIQANTGRQDFLRFGRNVANFYDREPKAFETFTPDDFD
>FLP-23_Parascaris_univalens_PgR087_g018_t02 peptide:
PgR087_g018_t02
SLLIKHIRLVIPISMWLHCSNRSYIKVMMLVNRSTLHLLSAPFIMRSALAFLLLLTITSS
TFALRGALYRFRRWSLPNVLHDNIGRQDFLRFGRDAPITFPNGYFEVDKNDATDMTGYAN
>FLP-23_Wuchereria_bancrofti_maker-PairedContig_555-snap-gene-0.8-
mRNA-1 peptide: maker-PairedContig_555-snap-gene-0.8-mRNA-1
MVFFLMITTSMTLRGAMFRSRKAALENIIRANTGRQDFLRFGRANEKYNDRHLIASALLT
PIDLE
>FLP-23_Ancylostoma_ceylanicum_Acey_s0721.g1823.t1 peptide:
Acey_s0721.g1823.t1
MRTILYLLAVLATISSVFGMRGALFRAGRAAPPNYGRQDFLRFGRNAYGSAGQDLFASPW
GEYYAST
>FLP-23_Caenorhabditis_brenneri_CBN02295.1 peptide: CBN02295.1
MLFPKLTILLFLLLVVLQESSAVRGALFRSGRAVPFERVPGQQDFLRFGRAVMASGAGGG
SEGNGPDDMKSFIRINGEPEIIFQ
>FLP-23_Caenorhabditis_briggsae_CBG26101.1 peptide: CBG26101.1
MILPKLLSLLLILMVLKESSAVRGALFRSGRSVPFERAPGQQKSAPICREVQLFVPCLFT
FNRNCMIDKICVHEAPVTTQTPPVSISTPNPKREEKSYAALSLIGPSRRNDFLRFGRASG
AERVGEPQDEIKYGSYGGDADILY
>FLP-23_Caenorhabditis_inopinata_Sp34__30283300.t1 peptide:

Sp34_30283300.t1
MVKRDKRFGPNRESNPGPLAPEARIIPLDHPAAVLHSERCLLSGMLYYKSGTKFAEYSLK
MLLAKLSVLLLILAVVEEAAAVRGALFRSGRAAPFERAVGQQRQAPICREVALFVPCLFT
SQRDCVIDRICVHEKASKLYASSGSPNAKREGSGAESLSQIAPPRRNDFLRFGRAVVASG
VGGGSYAASDDWKTPSLVRIQGEPEMSLQ
>FLP-23_Caenorhabditis_latens_FL83_22794 peptide: FL83_22794
MTLLPKMLTILLILVVLKETAAVRGALFRSGRAVPFERVPGQQKTAPVCREVELYVPCLF
TYNRNCVIDKICVHEAPVTTQTTPISLTTTPNSKREGHRDSLSLFGPPRGNDFLRFGRAG
MASGAGGTSDANGAQDEMKASFLRINGEPEIIFQ
>FLP-23_Caenorhabditis_nigoni_Cni-flp-23 peptide: Cni-flp-23
MILPKLLVLLLILIVLKESSAVRGALFRSGRSVPFERAPGQQKSAPICREVQLFVPCLFT
FNRNCMIDKICVHEAPVTTQTPPVSISTPNPKREEKSYAALSMIGPPRRNDFLRFGRSGM
ASGAERVAEPQDEMKYGSYDGDILY
>FLP-23_Caenorhabditis_remanei_CRE06740.1 peptide: CRE06740.1
MALLPKMLVILLILVVLKETAAVRGALFRSGRAVPFERVPGQQKTAPVCREVELYVPCLF
TYNRNCVIDKICVHESPVATQTTPISLTTTPNSKREGHRDSLSLFGPPRGNDFLRFGRAG
MASGAGGTSDANGAQDEMKASLLRINGEPEIIFQ
>FLP-23_Caenorhabditis_sinica_Csp5_scaffold_03218.g29459.t1 peptide:
Csp5_scaffold_03218.g29459.t1
MKVFFFAIFLLLAVVLRESQAVRGALFRSGRAVXFRSGRAVPFERVPGQQKTAPICREVE
LFVPCLFTVNRDCVIDKICVHEAPVPLKSTTTITSIPVTTPNTKREGSFANLMGPPRGND
FLRFGRAVMASGVGGGSEAGQAARDDFRGEPETIYE
>FLP-23_Caenorhabditis_tropicalis_Csp11.Scaffold630.g18556.t2
peptide: Csp11.Scaffold630.g18556.t2
MFKLSILLLLILVVSKETVAVRGALFRSGRAVPFERVPGQQKLAPKCREVELYVPCLFTY
NRTCVIDKICVHELPVVTTPRPTTPVTTPNTKRENSVEPRRNDFLRFGRAVMASGVGGGS
EGNAPDDMKSFVRINGEPEILFQ
>FLP-23_Heterorhabditis_bacteriophora_Hba_08859 peptide: Hba_08859
MVLSFFLLQTSDSLVIPTSVLVFITHSLAMRGALFRSGRAIRSFDNTKNYIAQDFLRFGR
SPDMHGYEDSIYDWRRMNIPPPDSA
>FLP-23_Pristionchus_arcanus_PARCANUS000037698.t1 peptide:
PARCANUS000037698.t1
LDLSDMRLLVFLVLALMAVHTVFAVRGALFRSRRADPSAWAGREDFLRFGRALSSSEGID
KMDETRGKRFFTDFYADNDDNIDYY
>FLP-23_Pristionchus_expectatus_PEXSPEC000031583.t1 peptide:
PEXSPEC000031583.t1
APKSRPLDLSNMRLLVFLVLALVAFHTVFAVRGALFRSRRADPSAWAGREDFLRFGRALS
SSKEMEQVEDPRGKRFFTDFYADNDDNTEYY
>FLP-23_Pristionchus_fissidentatus_PRIFISSI000028259.t1 peptide:
PRIFISSI000028259.t1
MILPLFLLVLFCSLDSSSAMRGSLMREGRSAPAAFSPYREEFLRFGRAPLRPIESYNGFF
>FLP-23_Caenorhabditis_japonica FLP-23 predicted
PKFSAFQDFLRFGRSAASDGVGDVENLKTMYLRMARDPEVVYR
>FLP-23_Mesorhabditis_belari FLP-23 predicted
CCGLNKVKDFLRFGRSQVDDEVQPISYGRSIGLRYFR
>FLP-23_Oscheius_tipulae FLP-23 predicted
PPLSASEALFSVQDVRILLSEGLTIIVALSLRSVNTLSIVANTKDFLRFGRSAGFDRLYDDQEDFLAH
SRIRVLKEPLQYGFIF
>FLP-23_Parapristionchus_giblindavisi FLP-23 predicted
APTKLRLFQDFLRFGRSPMKPLDHYEYHYGGQTYTIPLRSFLTDGAPFNFNE
>FLP-23_Pristionchus_maxplancki FLP-23 predicted
IIGQKIPVLALVVIHTVSAVRGALFRSRRADPAAWAGREVRLLQSDTNPILFLQDFLRFGRSLSSSDD
KVDDARGKRFFTDFYPNEDNSDYPGY
>FLP-24_Plectus_sambesii_PSAMB.scaffold6152size10059.g27979.t1

peptide: PSAMB.scaffold6152size10059.g27979.t1
MHLQTTFIFICLAAVAALAVSSNEYYAEENDDENIGEDEKRRAGMNADALIRFGKRRMYQ
KQLNEEYDDVLEDLMEKRAPNRADMMMRFGRARSNADKMLRFG
>FLP-24_Acanthocheilonema_viteae_nAv.1.0.1.t00207-RA peptide:
nAv.1.0.1.t00207-RA
MSASKVVIVAVLLVINNFCFIDCKRLQDNDFARQFLFRGGFEPLRYYLSPSDGAYVAKRV
PSAADMMIRFGKRFGNVDAPIGDLNDN
>FLP-24_Ascaris_lumbricoides_ALUE_0001565101-mRNA-1 peptide:
ALUE_0001565101-mRNA-1
MFSLKAIVMIALVVICTFCISESRRFHDDDFSRQFLFRGIDEPLKNYMRLREARILSKRV
PSAADMMIRFGKRSFIEQDME
>FLP-24_Ascaris_suum_AgR005_g139_t01 peptide: AgR005_g139_t01
MFSLKAIVMIALVVICTFCISESRRFHDDDFSRQFLFRGIDEPLKNYMRLREARILSKRV
PSAADMMIRFGKRSFIEQDME
>FLP-24_Brugia_malayi_Bm9731.1 peptide: Bm9731.1
MSTFKVLILAVLLVINHFCNTDSKQLQDSDLFRGGFEPMRYYFNPSDSYIGKRVPNPADM
MIRFGKRSATFDAPTGDLNDE
>FLP-24_Brugia_timori_BTMF0001782301-mRNA-1 peptide:
BTMF_0001782301-mRNA-1
MSTFKVLIVAVLLVINHFCNTDSKQLQDSDLFRGGFEPMRYYFNPSDSYIGKRVPNPADM
MIRFGKRSATFDAPTGDLNDE
>FLP-24_Dirofilaria_immitis_nDi.2.2.2.t00408 peptide:
nDi.2.2.2.t00408
MFAPKIIFVAVLLIINHFCFTDSKRLQDNDFARQFLFRGGFEPMKYYMSTNNDAYIVKRV
PSPADMMIRFGKRFVPHDISFDDLDDN
>FLP-24_Elaeophora_elaphi_EEL_0000036001-mRNA-1 peptide:
EEL_0000036001-mRNA-1
MRQAHFAEKFEMNGTSWKFINLIPVMQRYDEEMLLCFNEDEMNGGFFGCSKCKSSMPFLK
ELYFTMSAPKIIIVAVLLVINHFCFNDSKRLQDNDFARQFLFRGGFEPLRYYLSPNDGLY
ITKRVPSAADMMIRFGKRFARYDVPSEKSDCRDCTSDVVIMNITGTWMCKIKLNSKNSKL
RIFDEVQFEEYIEISVID
>FLP-24_Enterobius_vermicularis_EVEC_0000588301-mRNA-1 peptide:
EVEC_0000588301-mRNA-1
MFRQIIELAVLILLLAAVAQCRYTGLPKEDVDRFNYRVASAPFSGYFREFRDEIPYVAKR
VPNPGDMMVRFG
>FLP-24_Litomosoides_sigmodontis_nLs.2.1.2.t04563-RA peptide:
nLs.2.1.2.t04563-RA
MLASKGVIVAALLVINHFCFNDSERLRDNDFARQFLFRGGLEPLRYYLNSNDDTHIAKRA
PSAADMMIRFGKRFAAYNVPVGDLNDN
>FLP-24_Loa_loa_EFO17253.1 peptide: EFO17253.1
MSTFKILIIALLLVINHFCCSDSKRLQDNDFARQILFRGNFEPMRYYLSPNDAYLVKRVP
SAADMMIRFGKRFTTYDGSIGDINDD
>FLP-24_Onchocerca_onchengi_nOo.2.0.1.t03746-RA peptide:
nOo.2.0.1.t03746-RA
MSASKILIIAVLLIINHFCFNDSKRLQDNDFARQFLFRGGFEPMKYYMSPYDDVYTVKRV
PSAADMMIRFGKRSRTYDVPIGDLNDD
>FLP-24_Onchocerca_volvulus_OVOC8190.1 peptide: OVOC8190.1
LIDQSYFAMSASKILIIAVLLIINHFCFNDSKRLQDNDFARQFLFRGGFEPMKYYMSPYD
DVYTVKRVPSAADMMIRFGKRSRTYDVPIGDLNDD
>FLP-24_Toxocara_canis_Tcan_00436.1 peptide: Tcan_00436.1
MLSLKAIVMIALVVICTFCISESRRFQDDDFARQFLFRGIDEPLRSYMNMRESRFLSKRV
PSAADMMIRFGKRSSFEQDNE
>FLP-24_Wuchereria_bancrofti_WBA_0000218701-mRNA-1 peptide:
WBA_0000218701-mRNA-1

```
MSTLKVLIVAVLLVINHFCNTDSKQLQDSDLFRGGFEPIRYYFSPSGSYIGKRVPNPADM
MIRFGKRSVTFDTPVGDLNDE
>FLP-24_Brugia_pahangi FLP-24 predicted
PSDSYIDKRVPNPADMMIRFGKRSATFNAPIGDLNDE
>FLP-24_Dracunculus_medinensis FLP-24 predicted
IDKNKFQFDENYISKRIPSAADMMIRFGKRFHPLSYEYE
>FLP-24_Gongylonema_pulchrum FLP-24 predicted
NFQPGNDGYMSKRVPSAADMMIRFGKRFAPLDTSGLAYPNFYVD
>FLP-24_Onchocerca_flexuosa FLP-24 predicted
KFDASLIKLFKFKPYDDMYTVKRVPSAADMMIRFGKRSRTYDVPIGDLNDD
>FLP-24_Parascaris_univalens FLP-24 predicted
LQLREPRILSKRVPSAADMMIRFGKRSFIEQDME
>FLP-24_Syphacia_muris FLP-24 predicted
FHDEFPTYIAKRVPNPGDMMVRFG
>FLP-24_Thelazia_callipaeda FLP-24 predicted
NFKLQRDSGIYIAKRVPSAADMMIRFGKRFAPYDQSVRSNDYDNYIA
>FLP-24_Ancylostoma_ceylanicum_Acey_s0266.g712.t1 peptide:
Acey_s0266.g712.t1
MFSLRSIVLAVLSIAFLICVIDARIVDQYDDHMAIPVGAADYRLRGWDDYIVPHKRVPSA
GDMMVRFGKRSV
>FLP-24_Angiostrongylus_cantonensis_ACAC_0000612401-mRNA-1 peptide:
ACAC_0000612401-mRNA-1
MPIFRLAKRSNWRSIVLILWSFVRFIPLVTQQELTPMDDYRLYRFGDYMAMHKRVPSAGD
MMVRFGKRSV
>FLP-24_Angiostrongylus_costaricensis_ACOC_0000026401-mRNA-1
peptide: ACOC_0000026401-mRNA-1
MRIVVNLQIPEKERLAIVLILWRFVSLVTQQELTPMDDYRLYRFGDYLAMHKRVPSAGDM
MVRFGKRSL
>FLP-24_Caenorhabditis_brenneri_CBN08894.1 peptide: CBN08894.1
MISSRSSIILILAILVAIMAVAQCRNIQYDIDEIAPESAFRYAQWGEIPHKRVPSAGDMM
VRFGKRAV
>FLP-24_Caenorhabditis_briggsae_CBG15244.1 peptide: CBG15244.1
MSRTSIILVLAIFVAIAAIAQCRNIQYDVDEISPEAAFRYAQWGEIPHKRVPSAGDMMVR
FGKRSV
>FLP-24_Caenorhabditis_inopinata_Sp34__30329200.t1 peptide:
Sp34_30329200.t1
MPETCPLIFNYLKMLSSRSSIILILAIIVAFMAVAQCRNIQYDVEETAPEAAYRYAQWAD
IPRKRVPSAGDMMVRFGKRSF
>FLP-24_Caenorhabditis_japonica_CJA10537.1 peptide: CJA10537.1
MSRLSFVLLLAVLVAFVCVAQCRNIQYDADEISPEAAFRYAQWGEMPHKRVPSAGDMMEI
CVYTRNDRLRILNLKCPGASGTVFVAMCREEARLGLTATVLVPARPKRIIGSVIGPATPK
PLIGTATVPAGLIPLISTFTVPAIPKPLIDMVTVPARPKPLIGTANIPARLIPLISPFTV
PAWPKPPIGLVIGPATPKPLIGTATIPAMIIPLISTFTAPAWPKPPIGLVIGPATPKPLI
GTATVPAGLIPLISTFTVSAKLNLLIGTATVPVRFGKRSI
>FLP-24_Caenorhabditis_latens_FL83_19071 peptide: FL83_19071
MLSSRTSIVLIFAILVAIMAVAQCRNIQYDIDEITPEAAFRYAQWGEIPHKRVPSAGDMM
VRFGKRSV
>FLP-24_Caenorhabditis_nigoni_Cni-flp-24 peptide: Cni-flp-24
MSRTSIILVLAIFVAIMAIAQCRNIQYDVDEISPEAAFRYAQWGEIPHKRVPSAGDMMVR
FGKRSI
>FLP-24_Caenorhabditis_remanei_CRE25718.1 peptide: CRE25718.1
MLSSRASIVLIFAILVAIMAVAQCRNIQYDIDEITPEAAFRYAQWGEIPHKRVPSAGDMM
VRFGKRSV
>FLP-24_Caenorhabditis_tropicalis_Csp11.Scaffold630.g16685.t1
```

peptide: Csp11.Scaffold630.g16685.t1
MMSSRSSIILVLAIILAIMAVAQCRNIQYDVDEISPEAAFRYAQWGEIPHKRVPSAGDMM
VRFGKRSV
>FLP-24_Diploscapter_coronatus_DCO_025293 peptide: DCO_025293
MFSLRQLAALLVVLFAMFCILEARNMPYEAEALYPGYRFDGWAEAPMHKRVPSAGDMMVR
FGKRSE
>FLP-24_Diploscapter_pachys_WR25_04115.1 peptide: WR25_04115.1
MFSLRQLAALLVVLFAMFCILEARNMPYEAEALYPGYRFDGWAEAPMHKRVPSAGDMMVR
FGKRSE
>FLP-24_Haemonchus_contortus_HCON_00094680-00001 peptide:
HCON_00094680-00001
SGMISVRSIVLAVLFVAFLICAVDAKYLLPYDHEYYQGDYRPESFGDFVAAHKRVPSAGD
MMVRFGKRSV
>FLP-24_Haemonchus_placei_HPLM_0000712801-mRNA-1 peptide:
HPLM_0000712801-mRNA-1
MISVRSIVLAVLFVAFLICAVDAKYLLPYDHEYYQGDYRPESFGDFVAAHKRVPSAGDMM
VRFGKSVPFPGNYGSVADPTVTAAQPCTVPLYYQDSVYAKSSASSCSKSESDVSSDEAYK
RKREKRDRNNEAARTSRLRRKARESQMQKEAEELQQENDALKGEVGELKKVLYSLQDEVR
NRLNVVDIENSENVVQFQNIGYSFHNENL
>FLP-24_Heligmosomoides_polygyrus_HPOL_0001472001-mRNA-1 peptide:
HPOL_0001472001-mRNA-1
MLSLRTIALAVLFVAFLMCVVDARVLQPYDHEPYSGDYRLDAFGDFIAAHKRVPSAGDMM
VRFGKRSV
>FLP-24_Heterorhabditis_bacteriophora_Hba_15417 peptide: Hba_15417
MFSLRSMMWALIALIAFFCVTDARNLQPYGNYQFLDSEGMNLGDYRLEQFGEYINPQKRV
PSAGDMMVRFGKRAD
>FLP-24_Micoletzkya_japonica_MicoRS5524-mkr-S9-4.3-mRNA-1 peptide:
MicoRS5524-mkr-S9-4.3-mRNA-1
MKLLHSLLLLIVALVYFAAARTIYPYANMQELAGAAEEPEWALPSAAKRVPSAGDMMVRF
G
>FLP-24_Necator_americanus_NECAME_08917 peptide: NECAME_08917
MISLRSIVLAVLSIAFLICVIDARIVDQYGLSTFTEDHMAVPYVAGDYRLRAFDDFNFPH
KRVPSAGDMMVRFGKRSV
>FLP-24_Nippostrongylus_brasiliensis_NBR_0001956401-mRNA-1 peptide:
NBR_0001956401-mRNA-1
MQPLANSVIGFYSSSWFERYVIGLMRNESELYPMGDYRLASLGDYVSAHKRVPSAGDMMV
RFGKRSV
>FLP-24_Parapristionchus_giblindavisi_Parapristionchus-mkr-
S_98-1.46-mRNA-1 peptide: Parapristionchus-mkr-S_98-1.46-mRNA-1
MSLYILFLSIKGMKFFILLSIILLSLLFTSSMARTVYSYGQLGEMTDPNVEDNEWIMGSS
AVKRVPSAGDMMVRFG
>FLP-24_Pristionchus_arcanus_Pristionchus_arcanus_arcanus-mkr-
S_209-1.92-mRNA-1 peptide: arcanus-mkr-S_209-1.92-mRNA-1
MKSILIIILFSFVLLLSLSSARNIFPYGPLGQMGETNAEEGEWMIPSMAKRVPSAGDMMVR
FG
>FLP-24_Pristionchus_entomophagus_entomophagus-mkr-S28-4.55-mRNA-1
peptide: entomophagus-mkr-S28-4.55-mRNA-1
MKSVHLILLSLVLLSSLFSLVSSRNIYPYGQTGQMGDMNAEEGGWMIPSMSKRVPSAGDM
MVRFG
>FLP-24_Pristionchus_exspectatus_exspectatus-mkr-S_40-2.105-mRNA-1
peptide: exspectatus-mkr-S_40-2.105-mRNA-1
MKSILIIILLSFILLLSLSSARNIFPYGPVDQMGETNAEEGEWMIPSMAKRVPSAGDMMVR
FG
>FLP-24_Pristionchus_fissidentatus_fissidentatus-mkr-S460-0.36-

mRNA-1 peptide: fissidentatus-mkr-S460-0.36-mRNA-1
MKVLLLLLLSLAMLISFASARNIYPYGQLGQMGEINAEEGDWMVPSMAKRVPSAGDMMVR
FG
>FLP-24_Pristionchus_japonicus_japonicus-mkr-S5-6.49-mRNA-1 peptide:
japonicus-mkr-S5-6.49-mRNA-1
MKSFIVLLLSFILLISLASARNIFPYGAVGQMGETNAEEGEWMVPSMAKRVPSAGDMMVR
FG
>FLP-24_Pristionchus_mayeri_mayeri-mkr-S86-0.2-mRNA-1 peptide:
mayeri-mkr-S86-0.2-mRNA-1
MQSFFFVFFLLLLFSIVSSRSLSPLSPFGQMGEMNVEEGEWMIPSMAKRVPSAGDMMVRF
G
>FLP-24_Pristionchus_pacificus_PPA40082.1 peptide: PPA40082.1
MKSILIILLSFILLLSLSSARNIFPYGPVGQMGEINAEEREWMIPSMAKRVPSAGDMMVR
FG
>FLP-24_Teladorsagia_circumcincta_TELCIR_00856 peptide: TELCIR_00856
MVIWKWRVKSGKIAKERRCEESETLEAVYESRWLKVKPVGLKGTNGTIKSKGKVESIFDT
CPLILFNHLDKELLPFHHPNYTPKAGCRIYEPLTKLIKGRVEVSGNSSKHNCTARCLLPL
KKDRTYNATSWLKVPSPKIFECDIIETQCEKGGVYESFLHTQIYEQRSLPKTIKYLKGEL
GAIQMEFLNKVGDNSRPNGFPLAFGKSIEGGARDLVGLPPLIPDWNDAEICGKYLDQFSY
HLKDYKEMGYKTMIAQDFDVGIIYYPNCLGFNRSEADHIWRPFDLRARESKNFTKSLWQS
CSEPHLEMMDYMEKFMHSYPGTPKIAQIWPTTLAHESLKALYHADKHFHKYFQKNRAVID
RSFMFFMGDHGPRREGIGKVRLGQYENLNPFLMVTIPSAYRNTSLHEQLRQKTFELMTNF
DVHATLMDILKASSIKQLLDHGDILYEVIVYLTPSNGLFSCQELLDDFRCPKKNQYHEDF
PIEPRKRTGRPRTKNTPRAFIRRNESGLALGSGFTRLDRYGRQGDCLVGNNPLRGPYDQL
EERIVSVSYPHHDFRDCSKLCEKETTESRIFAGREWPQWCNGMISLRSIVLAVLFAAFLI
CAVDAKVLLPYDHEFYPGDYRLESFGDFLATHKRVPSAGDMMVRFGKRSV
>FLP-24_Caenorhabditis_elegans_C24A1.1.1 peptide: C24A1.1.1
MLSSRTSSIILILAILVAIMAVAQCRNIQYDVEEMTPEAAFRYAQWGEIPHKRVPSAGDM
MVRFGKRSI
>FLP-24_Ancylostoma_caninum FLP-24 predicted
SRSIPDDHMAIPVGAADYRLRGWEDYIVPHKRVPSAGDMMVRFGKRSV
>FLP-24_Ancylostoma_duodenale FLP-24 predicted
LPFIRVLHANPFPIPDDHMAIPVGAADYRLRGWEDYIVPHKRVPSAGDMMVRFGKRSV
>FLP-24_Caenorhabditis_angaria FLP-24 predicted
KIKGRERILIDFDELSQQSGFPYAQWAELPHKRVPSAGDMMVRFGKRSI
>FLP-24_Caenorhabditis_sinica FLP-24 predicted
HIQNFPDIDEISPEAVFRYAQWGEIPHKRVPSAGDMMVRFGKRSA
>FLP-24_Cylicostephanus_goldi FLP-24 predicted
SHFSISEDSMGLPVTGGDYRLRGWSDYISAHKRVPSAGDMMVRFGKRSV
>FLP-24_Dictyocaulus_viviparus FLP-24 predicted
ICFSHIFVIEHEFILTHDDLHGYENLEIRKRLPSAGDMMVRFGK
>FLP-24_Mesorhabditis_belari FLP-24 predicted
FQDDGIITHKRAPSAGDMMVRFG
>FLP-24_Oesophagostomum_dentatum FLP-24 predicted
GNQSIIDFPDDSLANALGVADYRLRPWGDYMIPRKRVPSAGDMMVRFGKRSV
>FLP-24_Oscheius_tipulae FLP-24 predicted
GDMMVRFGKRGAV
>FLP-24_Strongylus_vulgaris FLP-24 predicted
VFALTHFSDDSMAIPSSVGDYRLRAWNDYYVPHKRVPSAGDMMVRFGKRSV
>FLP-24_Teladorsagia_circumcincta FLP-24 predicted
ASFSYVILPFSDHEFYPGDYRLESFGDFLATHKRVPSAGDMMVRFGKRSV
>FLP-24_Rhabditophanes.sp.KR3021_RSKR_0000069950.1 peptide:
RSKR_0000069950.1
MTSKVFLIFAIVACSLVALTSAAAITPHSNSFNSKDIGENEFGFYYVPRMPMLSPYKENQ

FYAKRAPSAADMMIRFGKRSGVDEITY
>FLP-24_Steinernema_carpocapsae_L596_024451 peptide: L596_024451
MVGFSQLAILFFAVVVALIASNCEARPYDDEFGPSFYRPQYDAMRGFMGSRYLEKRVPNA
ADMMIRFGKRSGF
>FLP-24_Steinernema_feltiae_L889_g15835.t1 peptide: L889_g15835.t1
MTGFTQIAVFVFALFVTLFAVSAEARPYDDELGSSFYRPQYDAMRGFMGSRYLEKRVPNA
ADMMIRFGKRGGF
>FLP-24_Steinernema_glaseri_L893_g17490.t1 peptide: L893_g17490.t1
MVRFAQIVFFVAALLAVLLGASEARPLDDDFGAGFYRPQYDALRGFMTARYMDKRVPNAA
DMMIRFGKRSAF
>FLP-24_Steinernema_monticolum_L898_g8901.t1 peptide: L898_g8901.t1
MSDWVHEENNCTQLSVLNCLYSKEVHFFTSVLRMVGFTQIAVVFALLVAFFGVNSEARPY
EDEFGPSYYRPQYDAIRGFMNSRYLEKRVPNAADMMIRFGKRSGF
>FLP-24_Steinernema_scapterisci_L892_g24575.t1 peptide:
L892_g24575.t1
MDWPQRNECGFAPIRNEEVPWITARGQTGIKGLRRNGERFSPSRISFIKEALRRSRDSSV
SALPNFGHNGSLTRVGARKTYRKHGVLKAKYTDARVLIPESSDVVMLDSLFAQLLPRQSR
GVVLDEHRFLLTVVRRLSESTRGSGGQLDISSASLKTSFCSLSSSAFFGLHRPLPPTAAP
HPTCELVAFSKTKNRRLIAPNGALADLRPADSPTVPSGTIRVSPRPFGTFLSPSTAPLAP
PEIGFLREPKTVARVRKFDFWPISDSREVLLFSLPRMIGFSQLVVFLFAVFVALFATNSE
ARPYDDEFGPSFYRPQYDAMRGFMSSRFLEKRVPNAADMMIRFGKRSGF
>FLP-24_Strongyloides_papillosus_SPAL_0000027100.1 peptide:
SPAL_0000027100.1
MKKTNFVFFTIIFITMLLFIISSNAKSNEYDDNRSLSGSTYNDVFYIPTFRYYNPYGEKI
YDKRAPNKADMMIRFGKRLKPFDAIGIEM
>FLP-24_Strongyloides_ratti_SRAE_2000055800.1 peptide:
SRAE_2000055800.1
MKDIKLRNSGLFLFIFKMRKTKFITFSIIFFTILLFALSSYAKNSEYDENHSLSGSSYND
VFFIPNYKYYNPYGGKIYVKRAPNKADMMIRFGKRLKPFDELGIDV
>FLP-24_Strongyloides_stercoralis_SSTP_0000716000.1 peptide:
SSTP_0000716000.1
MRKTKFITFSIIFFTILIFVLSSYAKNNEYDDSHSISGSLYNDIFYIPNYKYYNPYGGKI
YVKRAPNKADMMIRFGKRFKPFDELGSDA
>FLP-24_Strongyloides_venezuelensis_SVE_1820000.1 peptide:
SVE_1820000.1
MLLFIISSNAKSNEYDDNRSLSGSTYNDVFYIPTYRYYNPYGEKIYDKRAPNKADMMIRF
GKRLKPFDAIGIEM
>FLP-24_Parastrongyloides_trichosuri FLP-24 predicted
YGAKFYVKRAPNKADMMIRFGKRSRTFNEMLNQK
>FLP-25_Ascaris_suum_L3E_03566 peptide: L3E_03566
MAPAAPTHSSSSHRHSLWIFGEGEFSIVHTKKADYDFIRFGRSTPSEKRKGDNSYDYIRF
GKRSLVIVPYWR
>FLP-25_Brugia_pahangi_BPAG_0001313801-mRNA-1 peptide:
BPAG_0001313801-mRNA-1
MNDHRESMLTVIFAILICVISQCSITNAVTYDRFCQHYPPLSQYHSEPHSAKRTNYDFIR
FGRGRRDGPGTYDYIRFGKRLSGLLANQNSEQTNLAK
>FLP-25_Dirofilaria_immitis_nDi.2.2.2.t04908 peptide:
nDi.2.2.2.t04908
MNGNREGILTIIFTFFLLRVVSQRLIANPTSNEQFYHRNPAPLQYQYETYTEKRVPNYDF
VRFGRSGRNDPAYDFIRFGKRFPGLWAKQKFGQIHLVKQSNLAR
>FLP-25_Elaeophora_elaphi_EEL_0000078901-mRNA-1 peptide:
EEL_0000078901-mRNA-1
MNTHRENMLTVILAILLCVVSQCSIANAVSCEQLCHRYSALSQYQSEPHSAKRTNYDFIR
FGRGGKASPATYDYIRFGKRLSEFWANQNSEQANLAR

>FLP-25_Loa_loa_EJD73860.1 peptide: EJD73860.1
MNGHRKSMLTIIFAIFLCVISHYSIANPIKYDELCQQQSPLLLCQFEPNSAKRTNYDFIR
FGRSGRVRPATYDYIRFGKRSTEFRINQDFE
>FLP-25_Onchocerca_onchengi_nOo.2.0.1.t05072-RA peptide:
nOo.2.0.1.t05072-RA
MSGNREGMLIITFTILFCVVTQYSVAATIRYGKFYHPYPAILQQYKSEPPQSAKRTRDYD
FIRFGRSGRSGQDNEYDYIRFGKRFPGSWVNQKFG
>FLP-25_Onchocerca_volvulus_OVOC7604.1 peptide: OVOC7604.1
MSGNREGMLIITFTILFCVVTQYSVAATIRYGKFYHPYLAILQQYKSEPPQSAKRTRDYD
FIRFGRSGRSGQDNKYDYIRFGKRFPGSWVNQEFG
>FLP-25_Wuchereria_bancrofti_maker-PairedContig_958-snap-gene-0.10-
mRNA-1 peptide: maker-PairedContig_958-snap-gene-0.10-mRNA-1
MIATLTDNTKYIKICPMIDHRESMLTVIFAILLCVISQCSIANAITYDRFCQHYPPLSQY
QSEPHSAKRTNYDFIRFGRGRRDGPETYDYIRFGKRLSEFLANQNPEQTNLAR
>FLP-25_Acanthocheilonema_viteae FLP-25 predicted
KVNFIIIMYFTAKRTYYDFIRFGRGGRSSPATYDYIRFGKRLSEFEANQNSEQAELGR
>FLP-25_Anisakis_simplex FLP-25 predicted
TAKVISKKRLVSKFLTISPFCSNFYLLIILTDNYRIIAHLYSLFAVKKADYDFIRFGRTPSSSKRTIG
QSYDYIRFGKRSPSLKMEPYLMQ
>FLP-25_Ascaris_lumbricoides FLP-25 predicted
WKLLAAKKADYDFIRFGRSTPSEKRKGDNSYDYIRFGKRSLVIVPYWR
>FLP-25_Brugia_malayi FLP-25 predicted
KMNFINSTYSIIKLFFFFFFFFTVKRTNYDFIRFGRGRKDGPGTYDYIRFGKRLSGLLANQNSEQTNLA
K
>FLP-25_Gongylonema_pulchrum FLP-25 predicted
NCIGLVRTSCNLFAAKRGGDYDFIRFGRSGTGNQEGAPLTYDYIRFGKRSGSADYLLSS
>FLP-25_Litomosoides_sigmodontis FLP-25 predicted
IISPIHIAFAAMRTNYNFIRFGRGERASPATYDYIRFGKRSEFRHDRIPEQAADFARQSP
>FLP-25_Onchocerca_flexuosa FLP-25 predicted
HLEFSERKPSVQVFQLPLKQQKIIISLIQFIVPNKFFFFFTAKRTHNYDFIRFGRSGRTERSEEDNKY
DYIRFGKRFSGSWVNQKFG
>FLP-25_Parascaris_equorum FLP-25 predicted
WKLLTAKRADYDFIRFGRSTPSEKRKDDNSYDYIRFGKRSLMIVPYWR
>FLP-25_Parascaris_univalens FLP-25 predicted
WKLLTAKRADYDFIRFGRSTPSEKRKDDNSYDYIRFGKRSLMIVPYWR
>FLP-25_Ancylostoma_ceylanicum_Acey_s0008.g230.t1 peptide:
Acey_s0008.g230.t1
MQSISLLLLLLVVVDICYCTLHPCTTPECQQTLEEVLQMEYKPKRNYDFVRFGRSSPAKK
ASYDYIRFGKRSDSRAAAYGGARYALVQL
>FLP-25_Angiostrongylus_cantonensis_ACAC_0001086001-mRNA-1 peptide:
ACAC_0001086001-mRNA-1
LCSCLVKRHYDFVRFGRSDLKKKASYDYIRFGKRRSDVMTT
>FLP-25_Angiostrongylus_costaricensis_ACOC_0000189901-mRNA-1
peptide: ACOC_0000189901-mRNA-1
MKLIRLSLHKSSRGAVAQSAAAVCCLLGSGLKQCRFQENQKIQHTSISFNVFYFSINCVL
QNLIQCYYLVKRDYDFVRFGRPDLKKKASYDYIRFGKRRSDVMTT
>FLP-25_Caenorhabditis_brenneri_CBN11856.1 peptide: CBN11856.1
MMSHNSMIYLLIVAFLISVDGLKQNECPGGDCHEDIPVDLGLVLPPELYESTRLANLLAR
PSSQFKMKRDYDFVRFGRAVPHKKASYDYIRFGRK
>FLP-25_Caenorhabditis_briggsae_CBG10109.1 peptide: CBG10109.1
MSNSMIYLLLVVAVLTTVVDSKKDCSSIRGCDEDVSPVDLGLVLPPELYESTRLANLLAR
PSSQFKMKRDYDFVRFGRSAPMKKASYDYIRFGRK
>FLP-25_Caenorhabditis_inopinata_Sp34__30268200.t1 peptide:
Sp34_30268200.t1

MSHSLMVFYVLATLLLVLPTATSATKECSTDCSEDVPVDLNLVLPPELYEPTRLGRLMQR
PSSQFKMKRNYDFVRFGRSAPSKKATYDYIRFGRK
>FLP-25_Caenorhabditis_japonica_CJA07718.1 peptide: CJA07718.1
MSPVWMILVLIACHLLCGSGASASASDTKNECSLDCDQAPSDLGIMLPPELFDSTRLANL
LAARPALQFKMKRDYDFVRFGRSAPSKKASYDYIRFGRK
>FLP-25_Caenorhabditis_latens_FL83_07430 peptide: FL83_07430
MSHNSMIYLLVVFLILISTITVDAAKKECSGGCEEDVPVDLGLVLPPELYESTRLANLLA
RPSSQFKMKRDYDFVRFGRSAPIKKASYDYIRFGRK
>FLP-25_Caenorhabditis_nigoni_Cni-flp-25.1 peptide: Cni-flp-25.1
MSNSMIYLLLVVAVLTTVVDSKKDCSSIGGCDEDVSPVDLGLVLPPELYESTRLANLLAR
PSSQFKMKRDYDFVRFGRSAPMKKASYDYIRFGRK
>FLP-25_Caenorhabditis_remanei_CRE25593.1 peptide: CRE25593.1
MNRIRFHITVGLNYIDIVIIPFISFSFSPCSFFFFFTNNNSLLHIISFPVVTSLQFVTIM
SHNSMIYLLVVFLILISTITVDAAKKECSGGCEEDVPVDLGLVLPPELYESTRLANLLAR
PSSQFKMKRDYDFVRFGRSAPIKKASYDYIRFGRK
>FLP-25_Caenorhabditis_sinica_Csp5_scaffold_01751.g22916.t1 peptide:
Csp5_scaffold_01751.g22916.t1
MPSSMIYLLVVAVLVITVDASPKKGCSSXXXXXXXXXXXXXXXXXXXXXXXDASPKKVGSS
VDCAQEDAPVDLGLVLPPELYESTRLANLLSRPSSQFKMKRDYDFVRFGRSAPAKKASYD
YIRFGRK
>FLP-25_Dictyocaulus_viviparus_nDv.1.0.1.t12003 peptide:
nDv.1.0.1.t12003
MFPSTYLFLLAVILPINYGNTLPCTSIGCQFYNDIMNPTRFKMKRHYDDVQFGRSDLNKK
ASYDYIRFGKRNENAMMI
>FLP-25_Diploscapter_coronatus_DCO_024153 peptide: DCO_024153
MIRYLFVVLQVLAAGAIAMSLGPSQGDSLAQPGIRTYAAYPPQISRMQFLSDDGYAVHTA
ELKPKRNYDFVRFGRSAPTKKASYDYIRFGKRSSNPLAAFRNVKIQNRRK
>FLP-25_Diploscapter_pachys_WR25_16039.1 peptide: WR25_16039.1
MLRYLFVVLQVLAAGAIAMSLGPSQSDSLVQPGIRTYAAYPPQINRMQFLSDDGYAVHTA
ELKPKRNYDFVRFGRSAPTKKASYDYIRFGKRSSNPLAAFRNVKIQNQRK
>FLP-25_maker-scaffold165-snap-gene-0.6-mRNA-1 peptide: maker-
scaffold165-snap-gene-0.6-mRNA-1
LAPPFVAKKAWSCLQKALRANVTRRQRYSRNMHSTTFVLLFTVLACISYSDSQPCTTAGC
EEAVADIGLPSEYRQKRHYDFVRFGRAAGPEKKASYDYIRFGKRSQIDSDDLNARFSPYQ
FL
>FLP-25_Heligmosomoides_polygyrus_HPOL_0001265601-mRNA-1 peptide:
HPOL_0001265601-mRNA-1
MHHSITVLLLLAALFSCSTSQPCTSPSCERNAEQFSPLDHKVKRHYDFVRFGRSMSPEKK
ASYDYIRFGKRSSWSPAMDGARFSGPLL
>FLP-25_Heterorhabditis_bacteriophora_Hba_20824 peptide: Hba_20824
MGPLVPIITLGLLCVDYSIVLSSTPTECIYTQSMDTYQPLSAEFPKQKRDYDFVRFGRSG
RMKKASYDYIRFGKRSARDYTSEDLLRIRNRQTEEFASVPFKIAT
>FLP-25_Mesorhabditis_belari_mbelari.g22924.t1 peptide:
mbelari.g22924.t1
MEFLQSEAQLVMTNKIPIIPSLLIFTRVTLSEMKSITPFLSLLAFLLLSAQAYEVLLCEQ
QPELCNLLAVERDSRLPISDVPKHAEKRAYDFVRFGRSDPAIGKEKKNSYDYIRFGKRSA
GLLAEDLVKF
>FLP-25_Oscheius_tipulae_OTIPU.nOt.2.0.1.t03826 peptide:
OTIPU.nOt.2.0.1.t03826
MQVLIVVVCFSLFLVQTSLQWTCSGPDCSPAHPLSQQRLRRNNGAYKRRDEYDFVRFGKR
SEAQKKAGYDYVRFGKRSGGSSAWRI
>FLP-25_Teladorsagia_circumcincta_TELCIR_11775 peptide: TELCIR_11775
MANFSANVETALARKIKGNPSRHTVQFCMNDIIKHRITKKETPGNVRTKTRHYDFVRFGR
SVGPEKKASYDYIRFGKRSQGTMDRFDARFPSFQFL

>FLP-25_Caenorhabditis_elegans FLP-25
MSHNSMIYLLVAFLVLLCATTEAKKECSIDCQEDGSAAVDLGLVLPPELYESTRLSNLLARPSSQFKM
KRDYDFVRFGRA
APIKKASYDYIRFGRK
>FLP-25_Ancylostoma_caninum FLP-25 predicted
LARLTETLETHWQFFHKNPRKKDFFSENHGDLSLVSVSAKRNYDFVRFGRSGPAKKASYDYIRFGKRS
DLRAAPYGGARYALAQL
>FLP-25_Caenorhabditis_angaria FLP-25 predicted
IFAVKRDYDFVRFGRSSPPAKKASYDYIRFGRK
>FLP-25_Cylicostephanus_goldi FLP-25 predicted
QNKQKINKHLLTNDFKLFPEPYKLLALLPVSAKRNYDFVRFGRSGSPSKKASYDYIRFGKRSGTREEN
FDTSRYAPIDQQL
>FLP-25_Haemonchus_placei FLP-25 predicted
KKTYPDTLQIADKGTVVGITTYRTLPSHTVIDTDSEKRHYDFVRFGRAAGPEKKASYDYIRFGKRSQI
DSDDLNARFSPYQFL
>FLP-25_Micoletzkya_japonica_Micoletzkya_japonica FLP-25 predicted
AEALGHTSACQCTKMPNTVDFVRFGRGP
>FLP-25_Necator_americanus FLP-25 predicted
NVFSLISLFSAKRNYDFVRFGRSGPAKKASYDYIRFGKRSSDRMDLDSLRSSALEQ
>FLP-25_Nippostrongylus_brasiliensis FLP-25 predicted
INDEVEYRGKYKLQLAMDASSDPNVRIISLFLLTTLNFFTLRQMKSDPCRFLLPHRLCRHMVKQRNLK
KLHIKSPTYQYKFLVKRHYDFVRFGRSGPAKKAASYDYIRFGKRTSSIYIPRFYTNPLV
>FLP-25_Oesophagostomum_dentatum FLP-25 predicted
ENLLRFRFSAKRNYDFVRFGRAARSGPSKKASYDYIRFGKRSVREDSEQDRYDLSGQL
>FLP-25_Strongylus_vulgaris FLP-25 predicted
KSLVSAKRNYDFVRFGRSARSPAKKASYDYIRFGKRNGAQEIHGTSRHDPLNSIYTL
>FLP-25_Panagrellus_redivivus_Pan_g2977.t1 peptide: Pan_g2977.t1
MLARIVFLLTVLPATLAFMMTYGDLCRMDSTLCQKPVSSYAASPVSAEREALFFRDALPR
PSLSEKPTKRGYDFIRFGRSADRQTKKAASTYDYIRFGRR
>FLP-25_Parastrongyloides_trichosuri_PTRK_0001699600.1 peptide:
PTRK_0001699600.1
MTGKVVNLLVFALSIIVIQSSTVLVDEKSPCIPLNYKSQSLDAPVNDCKDDSYMERYIRV
PSFNLKQSILSNRMLFDQLRDPIGNYKIFQKFKNHVDASGRSRYHPAFTKTGFERMIKKT
NFGKYIQTLPGKREYDFVRFGKKNQPNERIQRSDNTEESNSYDFVRFGKRNSPFINTLDD
EMFGDDNKLHDDYSFIRFGRSSEQDKRGTGESMNYDFVRFG
>FLP-25_Rhabditophanes.sp.KR3021_RSKR_0001008500.1 peptide:
RSKR_0001008500.1
MSYWISILFVFVLMICCGGSQPFKDEATLKGEISPCKSAQDINGIFLTKPDDLLHTVITS
DCNPDKKAEKSSFERNSRIPYNNHFKYLSGRLMIEDLLNPMSEKNGKIGKYKDHFDVNNF
VVWDAESVYDRNLRSKIPSKRSSWGKKGEILLASRILRSDPNQLSYDFVRFGKRSGKKVF
QGKVSKTQDVVKRNSQIENSNSHYGYDFVRFGKKSVQIEEAKARNDYDFVRFGKRNPPKE
ETKSRDDYDFARFGKRSIDHLNVHVKTKLEKRSDSKDNYDFVRFD
>FLP-25_Steinernema_carpocapsae_L596_g22503.t2 peptide:
L596_g22503.t2
MSRVPVVADRSSKKPVRSQHDLALPTARMCPRIAAFPTPSALLFSAAVVLLALQPVAEAL
TLSDLCRQDDSLALCQFQEIPQTSETMILNVPKQMRLRRSGGGQSRVRQDEIVKPDGGKR
GYDFIRFGRSPTANRAPLASYDFIRLGRK
>FLP-25_Steinernema_glaseri_L893_g26291.t1 peptide: L893_g26291.t1
MRRAPVVTDQSPQTARSQHDRRRMSPRASTPPLLLSVVVVFALQAAVDALSVQDLCRQAD
SAALCRALQRAPPSLQEADRPDRDHKPDDSEPPAPLRLAFWRYTRLHPRNDPTVAAAASS
FSKNGGKRGYDFIRFGRSANAPLKSYDFIRLGK
>FLP-25_Steinernema_monticolum_L898_g21910.t1 peptide:
L898_g21910.t1
MSLRIASSSCSALLVSFVVVFALQPVAEALTISDLCRQGAVESVALCKIRALQKVTGAPS

TILFDPIQPMQMRSQRSGKNRVEHEEVVKPDETDLPAPLRLVSFRFTRLHPRDERSSAAS
SFSKNGGKRGYDFIRFGRSPSSSNRTPLASYDFIRLGRK
>FLP-25_Steinernema_scapterisci_L892_g30049.t1 peptide:
L892_g30049.t1
MRRVPVVADRSSKKPVRSQHDLASTKARMCPRIVALPSASALLFSAAVVLLALQPVSEAL
TLSDLCHQDNSIALCQLQEVPKSPETMILSAPEPMRLRRSGGGQSRVRQDEIVKSGEPEN
PAPLRLAFWRYTRLHPREDRPAASSISKNGGKRGYDFIRFGRSPATNHAPLASYDFIRLG
RK
>FLP-25_Strongyloides_papillosus_SPAL_0001124200.1 peptide:
SPAL_0001124200.1
MTGKFTNTFILLLFLHVNFCSSLSKASDKSLCNFVVDNIKTGKYTTEELFKACNPSSIHR
NYMHRDTRAPSFSLKHSLFSNRMLFDHLRDSDGINKIFQKFKNHVDASGRNRYHPGFTST
GFDRMMKKTNFNKYMKLKLNKRSDNLPILKKKRDTTDRFQRSDPTDSDGFSYDFVRFGKR
DTPESQDDSLIQSSLGMYGPLYKRSGKYQNLNFGKGGEEGKRGVGESTNYDFVRFG
>FLP-25_Strongyloides_ratti_SRAE_2000220000.1 peptide:
SRAE_2000220000.1
MTGKLITILLFLLFFYGNQSSPLNNDRSLCASIINNIPATFISEEMANDCKPYLKNEKYT
ARDTRTPSFNLKQSIFSNRMLFDQLRDSIGNYRIFQKFKNHVDASGRNRYHPAFTKTGFE
RMIKKTTVNKYKKQPSKKNDDLIIFEKRNDLNGRFQRSESQIPEGFSYDFVRFGKRDPVY
IQKNPSLDNSLEIYKQFYKKSEDNKTFNFRKEGEQDKRGVTESNNYDFVRFG
>FLP-25_Strongyloides_stercoralis_SSTP_0000977400.1 peptide:
SSTP_0000977400.1
MLFDQLRDPAGSYKIFQKFKNHVDASGRNRYHPAFTKTGFERMIKKTAFDKYKKQSIPKR
RGDDLMVFNKRKDPNDRFQRSDKTDPEGFSYDFVRFGKRNPLYYNKKSQPYDDLIGAFNQ
VLHFERGGEQDKRSDLEGTNYDFVRFG
>FLP-25_Strongyloides_venezuelensis_SVE_1829900.1 peptide:
SVE_1829900.1
MTGKFTNTFIFLFFLHVNFCSSLSKTSDKPLCTFVVDNMKTKKYTTEELFKACNPSSTYE
RNMNRDIRAPSFSLKHSIFSNRMLFDHLRDSDGIHKIFQKFKNHVDASGRNRYHPGFTTI
GFDRMTKKTNFNKYMKSRINKRSDNLLILNNRRDTNDRFQRSDPTNYDGFSYDFVRFGKR
DDSESQDDSLIQSSLGMYGPLYKRSEKYQNLNFGRGGEESKRTAIESTNYDFVRFG
>FLP-25_Halicephalobus_mephisto FLP-25 predicted
KFWNLETKISGKRAYDFIRFGRSPERSQSKKASSSYDYIRFGKRSLRR
>FLP-25_Steinernema_feltiae FLP-25 predicted
LISISDGGKRGYDFIRFGRSPSSSNKTPLASYDFIRLGRK
>FLP-25_Acrobeloides_nanus_ACRNAN_scaffold6663.g9391.t1 peptide:
ACRNAN_scaffold6663.g9391.t1
MHYGIGFLAVTIMAGFGWMQEAEGLLTFADLCFQSHDAYSLCKYSHLPKRSISGLYPYPD
SIFRPYNLGFGSSQTLYMLPPDAKQVKSPQKRDYDFVRFGRSDGQKKAAGSYDYIRFGKR
SSYDYIRFGKRSLVHGLDMDLDKLPSQ
>FLP-25_Ditylenchus_destructor_Dd_12079 peptide: Dd_12079
MPEFPVEGPNPDYADRTPNLKVWKQPTSDLLCVQQEVKRSQQKNSQFLLCQPRDSIENNA
VDLGADAMSTIGDQYDVADFPQVMPTGNAAIRLKPEMFKTNQKMSIAQLLGKQKYVLQAP
ERPIPLDYANRASSRIPMKRTTTYDFIRFGKRFPNPPVGSEDGRWKKEDKKANTYDYIRF
GK
>FLP-25_Globodera_pallida_GPLIN_001147600 peptide: GPLIN_001147600
MLFKKVSNYDAIYVFFPHARNEVSPHFRPSALALLCAQSMPSGQLAKVCAHLTGIVLMPS
GDGQTRPILIGPDGIGIGIGMQRKRHFWSARIAALEGRAQHRAGGAIAGTVKRAYDYIRF
GRRSAAVHLAQQQKKNSEHSAGSTYDYIRFG
>FLP-25_Globodera_rostochiensis_GROS_g12968.t1 peptide:
GROS_g12968.t1
MKSMIYAIFDAICFPEMIAGIRCHRRRRRHLPSLITAIGAALCVAAASQPSVNALAAQGS
PQFRPSALALLCAQSMPSGQLAKVCAHLTGIVLMPSGNGQTRPILIGPEGIGIGMQLKRA
YDYIRFGRRSAAVHLAQQQKKKSEHKGSPQFRPSALALLCAQSMPSGQLAKVCAHLTGIV

LMPSGNGQTRPILIGPEGIGIGMQLKRAYDYIRFGRRSAAVHLAQQQKKKSEHIKRAYDY
IRFGRRSAAVHLAQQQKKKSEHSAGGTYDYIRFG
>FLP-25_Heterodera_glycines_Hetgly.G000008458.2 peptide:
Hetgly.G000008458.2
MIVPFDRFRLLPSPFAIGFCFVFAALCVAASPSAKTFLQQQQQQTTLPLVQIVDQQQLDS
DAQTVPAHFRPSSIAFLCANSFPPSPQLAQMCADLTGIVLLPSADDDDGPRRPILIGPEG
IGIGTQRKRHFWSGRIAALEGRAENGAGGTNGGGSAEKRGYDYIRFGRRSSAVAAIPKLA
KQQRKKSADTSGARTAYDYIRFG
>FLP-25_Meloidogyne_graminicola_NXFT01004190.1.10897_g peptide:
NXFT01004190.1.10897_g
MFWLFSLLLILSINVEAFNPSIHQLKEETFNDFKGILNNNNNNLKIYYKIPKYLNIFSIP
IFDQQINKFKLTTKQLKYLCNQFLLKENLKELNFLKNNIKLCEEILISKILKKRSTSSSS
SSSSLYNFIRFGRRNSFQINGNTNKKLNNNGNTYDYIRFG
>FLP-25_Meloidogyne_hapla_MhA1_Contig1000.frz3.gene7 peptide:
MhA1_Contig1000.frz3.gene7
MFSFWVFSLFFITSVSTFSPVRQIKKEIFIPFEEKRSSPSSYDFVRFGRNSFKIDGKTNK
KSNGNNGNTYDYIRFG
>FLP-25_Meloidogyne_incognita_Minc3s01659g25420 peptide:
Minc3s01659g25420
MFLFWMVSLFFIASVSTFSPVHKSEEEIFMPYEDSSLNDLYLITSLNKRPKHLINIFSLP
NQQTNKLQLSPKQLKFLCNKFLKNNSKEYERLLFLSKNINLCKKEEKFITKISENEKRSS
SSYDFVRFGRRNSFQMDGKPNKKSNGNNGNTYDYIRFG
>FLP-25_Meloidogyne_javanica_M.Javanica_Scaff571g007708 peptide:
M.Javanica_Scaff571g007708
MFLFWMVSLFFIASVSTFSPVHKSEEEIFMPYEDSSLNDLYLITSLNKRPKHLINIFSLP
NQQTNKLQLSPKQLKFLCNKFLKNNSKEYERLLFLSKNINLCKKEEKFITKISENEKRSS
SSYDFVRFGRRNSFQMDGKPNKKSNGNNGNTYDYIRFG
>FLP-25_Ditylenchus_dipsaci FLP-25 predicted
TICPNRALIKTFTAKRSTTYDFIRFGRSIPKSLTEKKEDKKASSTYDYIRFGKRSYV
>FLP-25_Meloidogyne_arenaria FLP-25 predicted
FYFKEKRSSSSYDFVRFGRRNSFQMDGKPNKKSNGNNGNTYDYIRFG
>FLP-25_Meloidogyne_enterolobii FLP-25 predicted
FYFKEKRSSSSYDFVRFGRRNSFQMDGKPNKKSNGNNGNTYDYIRFG
>FLP-25_Meloidogyne_floridensis FLP-25 predicted
FYFKEKRSSSSYDFVRFGRRNSFQMDGKPNKKSNGNNGNTYDYIRFG
>FLP-25_Meloidogyne_incognita FLP-25 predicted
FYFKEKRSSSSYDFVRFGRRNSFQMDGKPNKKSNGNNGNTYDYIRFG
>FLP-25_Meloidogyne_javanica FLP-25 predicted
FYFKEKRSSSSYDFVRFGRRNSFQMDGKPNKKSNGNNGNTYDYIRFG
>FLP-26_Ascaris_lumbricoides_ALUE_0000859401-mRNA-1 peptide:
ALUE_0000859401-mRNA-1
MFLDYSNIGNVKIRLLQLVFIFWQELLVVCFQIHLRMMITSSFVAVLVLIIAVHVCAQEE
DEDGVLPERFVRSFPNNYLASDLALRFGKRHLTTLATEQRGTTDKKTEYGKDLLIQFQLN
ALRSWEMNGSSKRRIGINEKRRIDINDLALRFGKRSSYSFDPSNLNLRFGKRSFGGMLEG
FDEKRTFVIPTDLALRFGKK
>FLP-26_Ascaris_suum_AgR034_g029_t04 peptide: AgR034_g029_t04
MMITSSFVAVLVLIIAVHVCAQEEDEDGVLPERFVRSFPNNYLASDLALRFGKRSWEMNG
SSKRRIGINEKRRIDINDLALRFGKRSSYSFDPSNLNLRFGKRSFGGMLEGFDEKRTFVI
PTDLALRFGKK
>FLP-26_Gongylonema_pulchrum_GPUH_0001680701-mRNA-1 peptide:
GPUH_0001680701-mRNA-1
MGTVVMSSVVLSDSKRDYNVNEQQALRFGRSYTFDPSHLNLRFGKRAYESDPMEYANTRL
GKRSSSVLDLLDDDQRRKKAAPAAVYVPLNLADDLALRFGK
>FLP-26_Toxocara_canis_Tcan_10403.1 peptide: Tcan_10403.1

MMIVSSFVATLMLTFAVHVCAQEDEVEDILPDRFIRSMPRNYVPSDLALRFGKRSWATNE
NSKRRIDLNDLALRFGKRNSYVFDPSNLNLRFGKRSLADDFEMKRAFVIPDDLALRFGRK
>FLP-26_Dracunculus_medinensis FLP-26 predicted
LCIYSSPQDAEKRIYNANDLALRFGKRGEFHYDPSSLSLRQILLKNH
>FLP-26_Parascaris_equorum FLP-26 predicted
DXXXXXXXXXXXXXXXXXXXXXXXXXXXXXRRIGSNEKRRFDINDLALRFGKRSSYPFDPSNLNLRSF
FSSNELKTALMLEDIINLKEVTISFDTSS
>FLP-26_Parascaris_univalens FLP-26 predicted
KIEYGKDLPIQFYLNDLRSWEMNGNSRRRIGSNEKRRFDINDLALRFGKRSFYPFDPSNLNLRSFFSS
NELKTALMLEDIINLKEVTISFDTSS
>FLP-26_Ancylostoma_ceylanicum_Acey_s0121.g1013.t1 peptide:
Acey_s0121.g1013.t1
MHPARLLFLISVLIGSACRALYIPHDMQDLFLNTYDKRSRMTLEGVRIPMRYCQSTNHQA
ISSPEGGCIMQLQYGYRPDKREFNADDLTLRFGKRGGEMAFHPNDLALRFGR
>FLP-26_Caenorhabditis_angaria_Cang_2012_03_13_00514.g11740.t1
peptide: Cang_2012_03_13_00514.g11740.t1
MQVGYTFAVVFALLICSIFSYKLPVEFDEEYNPVVSKRHFYNNNKQYKREFNADDLTLRF
GKRSDAMAFSPDQLSLRFGRRR
>FLP-26_Caenorhabditis_brenneri_CBN19897.1 peptide: CBN19897.1
MKAMFVLAILLGTLVATNAFRLPFQFFNSNEDFNSGLAKRNYYESKPYKREFNADDLTLR
FGKRAGAGEQLAFSPDMLSLRFGK
>FLP-26_Caenorhabditis_briggsae_CBG10956a.1 peptide: CBG10956a.1
MKAVLLIAILLGSIAAVSAFRLPFQFFGSQEDFNSGLAKRNYYESKPYKREFNADDLTLR
FGKRAGAGEPLAFSPDMLSLRFGK
>FLP-26_Caenorhabditis_inopinata_Sp34__X0198800.t1 peptide:
Sp34_X0198800.t1
MKVIFMLALLIGSLVVSNAFRLPFQLFDSNENYNNGLSKRNYYETKPYKREFNAEDLTLR
FGKRGGVGETLAFSPDMLSLRFGKK
>FLP-26_Caenorhabditis_japonica_CJA10676.1 peptide: CJA10676.1
MQSIVLAALLLSALAITNAFHIPYGTLESRDDFAGPAKRAAYYVGKPFKREFNADDMMLR
FGKRAGEPVAFSPDMLSLRFGK
>FLP-26_Caenorhabditis_latens_FL83_00252 peptide: FL83_00252
MKAMFVLAILLGSIVAISAFRLPFQFFGSNEDFNSGLAKRNYYESKPYKREFNADDLTLR
FGKRGGAGDTLAFSPDMLSLRFGKK
>FLP-26_Caenorhabditis_nigoni_Cni-flp-26 peptide: Cni-flp-26
MKAVLLIAILLGSIAAISAFRLPFQFFGSQEDFNNGLAKRNYYESKPYKREFNADDLTLR
FGKRAGAGEPLAFSPDMLSLRFGK
>FLP-26_Caenorhabditis_remanei_CRE00570.1 peptide: CRE00570.1
MKAMFVLAILLGSIVAISAFRLPFQFFGSNEDFNSGLAKRNYYESKPYKREFNADDLTLR
FGKRGGAGDTLAFSPDMLSLRFGKK
>FLP-26_Caenorhabditis_tropicalis_Csp11.Scaffold570.g4228.t1
peptide: Csp11.Scaffold570.g4228.t1
MKAMFVLAILLGSLVATNAFRLPFQFFGSNEEFNNGLAKRNYYESKPYKREFNADDLTLR
FGKRGGAGETLAFSPDMLSLRFGK
>FLP-26_Diploscapter_coronatus_DCO_024914 peptide: DCO_024914
MSPSSLLLSLLGLLAFCSTFIHCIYIPDDLQNHRYFFGSIDKRAYLNNQEKREFNGDDLT
LRFGKRSQSMFDPSDSMLRFGRK
>FLP-26_Heligmosomoides_polygyrus_HPOL_0002306401-mRNA-1 peptide:
HPOL_0002306401-mRNA-1
MTSCSSSDYEDIIQSIQSFLLRTELGNFIIGVRTTCTNQKREGNLKNLVVIFAQPNASDL
GEYLQPLLFSIPMMALRFGVDVYAFDYSGYGLSSGKPSEKNIYADIRAMYDYVRLKRSDK
K
>FLP-26_Heterorhabditis_bacteriophora_Hba_16592 peptide: Hba_16592
MNPTWLLLIVFGLLVCVSCGLYIPHDLQDIYMNNYEKRSQFNFPSLRPEKREFNTDDLTL

RFGSATSFHPEDLALRFGRK
>FLP-26_Mesorhabditis_belari_mbelari.g26211.t1 peptide: mbelari.g26211.t1
MRQTIGSSSSFLLQILIALVLLANLASAFYVPYAEDYFMMNNPIMEKRTLGPYGGPRHEK
REFSADDLSLRFGKRSGRLAFNPEDLHLRFGK
>FLP-26_Necator_americanus_NECAME_01178 peptide: NECAME_01178
MGILTAIFPVMYPGRLFLLISVLIGSACRALYIPHDIHSYSILTDPFDIGFEENYHRPRR
IPVKIGYRPDKREFNADDLTLRFGKRSGDMAFHPNDLALRFGR
>FLP-26_Oesophagostomum_dentatum_OESDEN_00079 peptide: OESDEN_00079
MHPARLLFLISVLIGSACRALYIPHDLHDIIANSYDKRSRMVIDGYRPDKREFNADDLTL
RFGKRGDAAFHPDDLALRFGR
>FLP-26_Oscheius_tipulae_OTIPU.nOt.2.0.1.t04110 peptide: OTIPU.nOt.2.0.1.t04110
MRALFFSLLLAFIATLSSAFYVPRDDLYYASSFEKRAKYYEYPRSELSSNHNKRDEFNGD
DLTLRFGKRANFQPEDLALRFGRK
>FLP-26_Pristionchus_entomophagus_entomophagus-mkr-S124-3.8-mRNA-1 peptide: entomophagus-mkr-S124-3.8-mRNA-1
MTRTGSLLTIILLSLLALVAQSYILPYEAYNEDGYYYMPSVAKRSSDMRHPAKREFNVDD
LTLRFGKRSSMSSFGADDLALRFGRRRR
>FLP-26_Pristionchus_fissidentatus_fissidentatus-sn_msk-S148-1.23-mRNA-1 peptide: fissidentatus-sn_msk-S148-1.23-mRNA-1
MCYRKLPKLQKHSHTMTRTGLITMLLLALLTVFSQAYILPYEAYRDDGYYFMPSAAKRSS
HVRHQAKREFNVDDLTLRFGKRSVPVLAFNQDDLALRFGRR
>FLP-26_Pristionchus_maxplancki_maxplancki-mkr-S346-0.5-mRNA-1 peptide: maxplancki-mkr-S346-0.5-mRNA-1
MTRTGQLMIILLSLLALLSQAYILPYEPYAEDGYYYFPSAVKRSSSDHLRHATKREFNVD
DLTLRFGKRSGGLSSFGADDLALRFGRRMNVPLLVLLLAHSVVGQFLKLDLIRHQGCSWK
TSSSWNRKLEFQGGSNKAGAQLTKVDGHAGCYSMHGRVNVHEDVVDEILIFLSVSTTGDT
SRPPDVCRDASPETGCGGVGSCLYCRPCESLGGLSKVLGAQLIVNGKVAGCEPLKKGIYE
DVELRFCLPNVQSLLEWQGISEKAIDNILAASSLDGNGIPKLSLFVTVYVFDKDVRSLLV
SQRKLEERIRKMKHVDVDDQLDSTTYWNLPFNQIIKSQQKFIGCHKLYGTVTISNQHWLQ
SAISYRYLDTMRIQRWSIKPRKPESDVFIVHFEAQTDRTSFNTLRESICILGKALLIVTT
MLALLLVMFYFLVQEQDEVRQEFDRFVVLYGKTYDSAEKEGIAYAFFIENMLELERERSE
QPGKFALGDQSLFPIEGMQMFDGFKENVNRPAEYSLKDKATPVKNQGSCGSCWAFATVAS
VETANVIAGNNLIALSEQEMIECDTRNTGCRGGVRTYAMSFVVQNGLVPESTYPYTAKEG
EQCHIGNATKVFIKDFRLLSTSEDAMADWLFKTGPITFGMNVTRSLYSYRSGVFRPSEAD
CTQKSEGSHALTILGYGSERGQDYWLVKNSWGDYWGDGGYFKLARGANVCGMANNVVAPI
FS
>FLP-26_Pristionchus_mayeri_mayeri-mkr-S639-0.0-mRNA-1 peptide: mayeri-mkr-S639-0.0-mRNA-1
MTSTGLLTIILLSLLALLSQAYILPYEAYSDDGYYYMPSADKRSSDHLRHAAKREFNVDD
LTLRFGKRSGLSSFGSDDLALRFGRHAAPISKPNRIQVMKTVILLLPLLGSVLGQFLKLE
LIRHQGCSWKKSASWNRKLEFQGGSNKAGAQLSRVEGHAGCYSMQGRVTVHEDVTDEILI
FLSVSTTGDTSRPPDVCRDADQETGCGGIGSCLYCRPCESLGGLSKVLGAQLIVNGKKGV
YDEVELRFCLPNVQSLLEWQGISEKAIDNILAASSVDGNGIPKLSLFVTVYVFDKDVRSL
LVSQRKLEDRIRKMKHVEADDQLDSTTYWNLPFNQMIKNQQTFIGCHKLYGTGNPSSLAP
CASLDGTLLRANQRAMSSSFMSSSQGDSQSSDLLRMSYGTMLKVLIVTVVGMALCYAMMT
ALLNERDNVDREFRKFIATYGKSYESIEKMDQAYAFFRENLAELKREMGRQPGVELGFHQ
WMDHHASTFSEKFALGDIIAPPMEGLTTPFTSFEEPVNRPAEFSLRDKATPVKNQGSCGS
CWAFATVASIETLNVIAGNPLVGLSEQEMIECDTRNQGCKGGVRTYAMSFVMQNGLVPES
TYPYTAKEGEQCHIGNATKVFIKDFRLMSTSEDAMADWLFKTGPITFGMNVTKAMYNYRS
GVFRPAEADCAQHSEGSHAMTIMGYGSEQGKDYWLVKNSWGDYWGDGGYFKLARGANVCG
MANNAVAPIFH
>FLP-26_Pristionchus_pacificus_PPA44898.1 peptide: PPA44898.1

MTRTGLLTIILLSLLALLSRAYILPYEAYAEDGYYYFPSVAKRSASDHLRHASKREFNVD
DLTLRFGKRSGGLSSFGADDLALRFGRRR
>FLP-26_Caenorhabditis_elegans FLP-26
MKVMFMLALLFSSLVATSAFRLPFQFFGANEDFNSGLTKRNYYESKPYKREFNADDLTLRFGKRGGAG
EPLAFSPDMLSL
RFGK
>FLP-26_Ancylostoma_caninum FLP-26 predicted
VFQTNVTIRSQPLIACVFSYRPDKREFNADDLTLRFGKRGGEIAFHPNDLALRFGR
>FLP-26_Ancylostoma_duodenale FLP-26 predicted
RSHCLKMLWNNTVSHVLPLSLPDKRHIIRSPPLIACVFSYRPDKREFNADDLTLRFGKRGGEMAFHPN
DLALRFGR
>FLP-26_Caenorhabditis_sinica FLP-26 predicted
FFQFFNSNEDFNGGLAKRNYYDSKALKRQFNADDLTLRFGKRGGAGEPLAFSPDMLSLRFGK
>FLP-26_Cylicostephanus_goldi FLP-26 predicted
XXXXXXXXXXXXXXXXXXXXXXXXXXXXXXXXXXXXXXXXXXXXXXXXXXXXXXXXXXXXXXXXXXXX
XXXXXXXXXXXXXXXXXXXXXXXXXXXXXXXDKREFNADDLTLRFGKRGMDYPNDLALLFERE
>FLP-26_Micoletzkya_japonica_Micoletzkya_japonica FLP-26 predicted
TTVFHCSARPDKREFNVDDLTLRFGKRSSNLANFNPDDLALRFGRR
>FLP-26_Pristionchus_arcanus FLP-26 predicted
NMLFRRHATKREFNVDDLTLRFGKRNGGLSSFGADDLALRFGRRR
>FLP-26_Pristionchus_exspectatus FLP-26 predicted
NASFRRHASKREFNVDDLTLRFGKRSGGLSSFGADDLALRFGRRR
>FLP-26_Pristionchus_japonicus FLP-26 predicted
FRNGSNLDFSKDFCQLLMIPWNKLLTKRKNLTRKIVNICRFISLTPCRPTPRMAITTSHRPSSDRHPI
ICKFEISKYCLFISHTLSPVRAARAKFEIIPINSFRRHATKREFNVDDLTLRFGKRSGGLSSFGADDL
ALRFGRRR
>FLP-26_Strongylus_vulgaris FLP-26 predicted
FSSYKTLGIQVKHMIRDAHCKKIXXXXXXXXXXXXXXXXXXXXXXXXXXXXXXXXFRYRPDKREFNVD
DLTLRFGKRGEMAFHPNDLALRFGRK
>FLP-27_Ancylostoma_caninum_ANCCAN_10892 peptide: ANCCAN_10892
MGLAPGATETHQPHAVVAVTSRQRQPRMQSPNALLVSMLAVLVLATLIPCNAEAQEFRPI
LMNRRDLLPYGEIVSELKGKTMGGRMRFGKRSMNPYQFIPIEAMEAYERQQQI
>FLP-27_Ancylostoma_ceylanicum_maker-ANCCEYDFT_Contig381-
pred_gff_snap-gene-0.5-mRNA-1 peptide: maker-ANCCEYDFT_Contig381-
pred_gff_snap-gene-0.5-mRNA-1
MGLAPGATETHQAHAVVAVTSRQSQPRMQSPNALLVSMFAVLVLATLIPCNAEAQEFRPI
LMNRRDLLPYGEIVSELKGKTMGGRMRFGKRSMNPYQFIPIEALEAYERQQQV
>FLP-27_Caenorhabditis_brenneri_CBN08869.1 peptide: CBN08869.1
MLSIQQVLAFLFVTIALMTFASAQPIDEDRPIFMERREASAFGDIIGELKGKGLGGRMRF
GKRSSAPDMSMAELRAIYGGGPVEYVQL
>FLP-27_Caenorhabditis_briggsae_CBG02476.1 peptide: CBG02476.1
MFSFRKFLAFMLIVIALMASFSSAQPIDEERPIFMERREASAFGDIIGELKGKGLGGRMR
FGKRSSSPSDISMAELRAIYGGGPVEYVQL
>FLP-27_Caenorhabditis_inopinata_Sp34__20114100.t1 peptide:
Sp34_20114100.t1
MSSLQHIIVLLIAAVALITITSAQPIDENRPIFMERREASAFGDIIGELKGKGLGGRMRF
GKRSSSKDVSLPELRAIYGRVPVEYVQL
>FLP-27_Caenorhabditis_latens_FL83_04387 peptide: FL83_04387
MFSFRQTLALMIIAIAIMSYSSAQPIDEERPIYMERREASAFGDIIGELKGKGLGGRMRF
GKRSSSPDISLAELRAIYGGGPVEYVQL
>FLP-27_Caenorhabditis_nigoni_Cni-flp-27 peptide: Cni-flp-27
MFTFQKFLAFVLITVALMASFSSAQPIDEERPIFMERREASAFGDIIGELKGKGLGGRMR
FGKRSSSPSSDISMAELRAIYGGGPVEYVQL
>FLP-27_Caenorhabditis_remanei_CRE11809.1 peptide: CRE11809.1

MFSFRQILALMIVAIAIMSYSSAQPIDQDRPIFMERREASAFGDIIGELKGKGLGGRMRF
GKRSSSPDISLAELRAIYGGGPVEYVQL
>FLP-27_Caenorhabditis_sinica_Csp5_scaffold_00079.g3571.t1 peptide:
Csp5_scaffold_00079.g3571.t1
MTSQSPPVSFLSSFRACSGSSAFQKWAWSSKEMWLFFHKLNSFWCHFRYTNHRLQMFSFR
QFLALLLVAIALMAFSSAQPIDEDRPIFMERREASAFGDIIGELKGKGLGGRMRFGKRSS
SPDISLAELRALYGGGPVEYVQL
>FLP-27_Micoletzkya_japonica_MicoRS5524-mkr-S125-1.83-mRNA-1
peptide: MicoRS5524-mkr-S125-1.83-mRNA-1
MALPLFLRFLLLGLVAILTVSAYDDMQPILMDRRDLSPFGDIVSDLKSKTLGGRMRFGKR
SGPVAPLSAQEWRANYLSGY
>FLP-27_Necator_americanus_NECAME_14245 peptide: NECAME_14245
MGLAPTETHTLNTVSYSRQRQSKMQPQHTLLITVLAIFVLAAFIPSTVEAQEYGPILMNR
RDLLPYGEIVSELKGKTMGGRMRFGKRSGNALHFVPAVVLDAYEHQQQQQQL
>FLP-27_Oesophagostomum_dentatum_OESDEN_13316 peptide: OESDEN_13316
MGLAPTAPETHQLKAVAVVTKRARQSRMQSSPAFLCSMFALLVLAALIPSGTEAQEYRPI
IMNRRDLVPYGEIVSELKGKTMGGRMRFGKRSMSPYLVNPVEAMDAYERQQQF
>FLP-27_Oscheius_tipulae_OTIPU.nOt.2.0.1.t12605 peptide:
OTIPU.nOt.2.0.1.t12605
MLAVQIALALVFSLLLIVEPVQGQDLEKPIQMDRREVGALGDIVTELKGKSMGGRMRFGK
RAMMPHYLPQRPHYVMEEDDFVY
>FLP-27_Pristionchus_arcanus_PARCANUS000017359.t1 peptide:
PARCANUS000017359.t1
FSNRFIRMASLIQMALLLSLLLITLCTVQAFDDYHPILMDRRDSSFNLDDLKSQALGGRM
RFGKRSGGGGGKMFRLNEWRSQLFDRPI
>FLP-27_Pristionchus_entomophagus_entomophagus-mkr-S1-10.51-mRNA-1
peptide: entomophagus-mkr-S1-10.51-mRNA-1
MASLIQMALFLSILLITLCTVQAFDDFRPILMDRRDAPFNELLNGLKTQTIGGRMRFGKR
SGEGEATMRLNQWRPQQFNRPF
>FLP-27_Pristionchus_exspectatus_exspectatus-mkr-S_44-2.1-mRNA-1
peptide: exspectatus-mkr-S_44-2.1-mRNA-1
MASLIQMVLLLSLLLITLCTVQAFDDYHPILMDRRDSSFNLDDLKSQALGGRMRFGKRSG
GGGKMLRLNEWRSQLFDRLI
>FLP-27_Pristionchus_fissidentatus_fissidentatus-mkr-S47-7.3-mRNA-1
peptide: fissidentatus-mkr-S47-7.3-mRNA-1
MPSLLQMVLLISLLLITLSTVASFDEFHPILMDRRDAPFGGLLNDLKTQTLGGRMRFGKR
SGGVESDSHNALQQLHQWRSQLLNRPL
>FLP-27_Pristionchus_japonicus_japonicus-mkr-S154-0.24-mRNA-1
peptide: japonicus-mkr-S154-0.24-mRNA-1
MVAPSKSKDGKKKKDKKEPPSSGGADHKEDAARPITIDCPKSAVAEGKGEGDKGLPNCVS
FFPEVVTYQPENKKQIRNLEVKNKGDRSIMYKMKSTSPGVYRMRPIHFILRPGENKIIKL
SYKGCPDGKAPNLKDRFTVVMAYPPGVESNVKLMWTQKAYVEKIAECTHRKYIKVHFDGY
DVPDKKPASKMVAPPSAPPPAAAAPAPAAPAAAAPPQGGWPGMGGGGGIVYVIYQGDQQG
GGGPPPSEGQDDEKMAPLLSLLLITLCSVQAFDDYHPILMDRRDAPFNLIHDLKSQTLGG
RMRFGKRSGEGGSMLRLNQWRSQLFDRPI
>FLP-27_Pristionchus_maxplancki_maxplancki-mkr-S51-4.39-mRNA-1
peptide: maxplancki-mkr-S51-4.39-mRNA-1
MASLIQMALLLSLLLITFCTVQAFDDYHPILMDRRDAPFNIIHDLKSQTLGGRMRFGKRS
EGGSMLRLNQWRSQLFDRPM
>FLP-27_Pristionchus_pacificus_PPA35587.1 peptide: PPA35587.1
MASLIQMVLLLSLLLITLCTVQAFDDYHPILMDRRDSSFNLDDLKSQALGGRMRFGKRSE
GGKMLRLNEWRSQLFDRPI
>FLP-27_Caenorhabditis_japonica FLP-27 predicted
RLFYVFFYRYWWLKSEKQTNAHHNFSILRFGKRSSSPDMRLAE

>FLP-27_Caenorhabditis_tropicalis FLP-27 predicted
NSKKRRSCHAFLTLFLWILLCFLAVGTYSQDDPTPFPGTNKLIQMNNYRFGKRSSSPDMSLAELRAIY
GGGPVEYVQL
>FLP-27_Caenorhabditis_elegans FLP-27
MFSLTQILTFLLVAITLMTFSSAQPIDEERPIFMERREASAFGDIIGELKGKGLGGRMRFGKRSSSPD
ISLAEMRAIYGG
DQSNIFNFK
>FLP-28_Ancylostoma_caninum_ANCCAN_02791 peptide: ANCCAN_02791
MLSARAVFALLYMLLIAVAVVNSAPNRILMRFGKRTSDPLHVRPMVPVDYYPLELLGSSR
AVDGDM
>FLP-28_Ancylostoma_ceylanicum_Acey_s0223.g2681.t2 peptide:
Acey_s0223.g2681.t2
MLSARAVFALLYMLLIAVAVVNSAPNRILMRFGKRTSDPLHLRPMVPVDYYPLELLGSSR
AVDGDM
>FLP-28_Ancylostoma_duodenale_ANCDUO_01868 peptide: ANCDUO_01868
MMLSARAVFALLYMLLIAVAVVNSAPNRILMRFGKRTSDPLHFRPMVPVDYYPLELLGSS
RAVDGDM
>FLP-28_Angiostrongylus_costaricensis_ACOC_0000054801-mRNA-1
peptide: ACOC_0000054801-mRNA-1
MASTGHGGPIVMFCDLGFTEAIASSKFYTVLVAAGPSQDENSSKNRQQFKRFRGEPIRFG
KRVPREPISEHIVQVAMFNLNHAHLKQQKNREPTIQRYYATFLQQASRNSRKAGMDASGI
MMSARTFLTIFCILLFTVLYTSSAPNRILMRFGKRATNTVPYAFVPLPDYYRVDLRRLTA
LFTTTLTETFGHKRKYCSVERQKTAHMVIVFHPQIEEVSEDRTEDRKASEQIEKKQLANV
YYSAEELAPLSLSFNPSLMDGNNRTRLINNATIR
>FLP-28_Ascaris_lumbricoides_ALUE_0000468401-mRNA-1 peptide:
ALUE_0000468401-mRNA-1
MDSRQLVLLTLLAIMLVTLKSNTVVDAAPNKILMRFGKRTLPLDRNLDEWITEKDLDNLH
NLYYLLRESGEQ
>FLP-28_Ascaris_suum_AgR027X_g063_t01 peptide: AgR027X_g063_t01
NRISMRAHHAPCPQCCQFVIVSLPSVVSETIRPMDSRQLVLLTLLAIMLVTLKSNTVVDA
APNKILMRFGKRTLPLDRNLDEWITEKDLDNLHNLYYLLRESGEQ
>FLP-28_Caenorhabditis_brenneri_CBN32882.1 peptide: CBN32882.1
MPSGSDLLNNTSFLVQKSWNAAPNRVLMRFGKRGGNSEGHLGYRYVPAAASAIAEYIDVD
DMFGGQDRF
>FLP-28_Caenorhabditis_briggsae_CBG01976.1 peptide: CBG01976.1
MFSVRSFVALFCVLILAFSAVNAAPNRVLMRFGKRGGNSEGNLGYRYVPAAAPAIAEYID
VDDVLGGQDRF
>FLP-28_Caenorhabditis_inopinata_Sp34__X0065300.t1 peptide:
Sp34_X0065300.t1
MFSVRSISAIFCVIILVLSTISAAPNRVLMRFGKRGGNSEGHLGYHYAPAAVPAIAEYID
VDDVFGGQDHF
>FLP-28_Caenorhabditis_japonica_CJA18541.1 peptide: CJA18541.1
MFSARSILAIFCALILALSTINAAPNRVLMRFGKRGGASEGHVGYRFVPAAGPAIAEYID
VDDSLLSQDRF
>FLP-28_Parascaris_equorum_PEQ_0001031001-mRNA-1 peptide:
PEQ_0001031001-mRNA-1
MDSRQLAFLSLLAILLVTLKTNTVVDAAPNKILMRFGKRTFPLDRNLDEWITEKVCLFVS
>FLP-28_Toxocara_canis_TCNE_0001561101-mRNA-1 peptide:
TCNE_0001561101-mRNA-1
MNATVDETTRPMNSRHLLLLALFATLLAVFNSKTDVDAAPNKILMRFGKRTYVGDRNPDE
WITEKDIANLRNLYYLMREGGDQQGTR
>FLP-28_Caenorhabditis_elegans FLP-28
MFSVRSIFAIFCVLILALSTINAAPNRVLMRFGKRGGNSEGHLGYRFVPAGAPAIAEYIDVDDVIGGD
DRF

>FLP-28_Anisakis_simplex FLP-28 predicted
LTIFIQLNMVGLMNSRQDNVIVVLFVVTLISVFHSNSMYVECAPNKILMRFGKRTYSLNDINTNNRYL
HESTLTGKVHLFLCYSFLIYYGIHTRFNHNYACISIRELIPSGSVGALIHINQ
>FLP-28_Dracunculus_medinensis FLP-28 predicted
LIIDRLITMNQYFISLFLMIILLINMHMLTAAPNKILMRFGKRNYNYFPDYKTFDKVRFLFLFKLNLT
ENNYEKK
>FLP-28_Parascaris_univalens FLP-28 predicted
NKAKTKVSETTGPMDSRQLAFLSLLAILLVTLKTNTVVDAAPNKILMRFGKRTFPLDRNLDEWITEKV
CLSVS
>FLP-28_Caenorhabditis_latens_FL83_01389 peptide: FL83_01389
MFSVRSILALFCLLILALSTINAAPNRVLMRFGKRGGNSEGHLGYRFVPAAASAYAEYID
VDDVLGGQDRF
>FLP-28_Caenorhabditis_nigoni_Cni-flp-28 peptide: Cni-flp-28
MFSVRSFVALFCVLILAFSAVNAAPNRVLMRFGKRGGNSEGNLGYRYVPAAAPAIAEYID
VDDVLGGQDRF
>FLP-28_Caenorhabditis_remanei_CRE03200.1 peptide: CRE03200.1
MFSVRSILALFCLLILALSTINAAPNRVLMRFGKRGGNSEGHLGYRFVPAAASTYAEYID
VDDVLGGQDRF
>FLP-28_Caenorhabditis_tropicalis_Csp11.Scaffold629.g13663.t1
peptide: Csp11.Scaffold629.g13663.t1
MFSVRSILAIFCVLMLALTTINAAPNRVLMRFGKRGGNSEGHLGYRYVPAVAPAIAEYID
VDDVLGGQDRF
>FLP-28_Cylicostephanus_goldi_CGOC_0000389601-mRNA-1 peptide:
CGOC_0000389601-mRNA-1
MFSARTVFVLFYTLLLAVAIVSAAPNRILMRFGKRADPVQYRPFVPSDYYPVELLGSSRG
ADGDM
>FLP-28_Haemonchus_contortus_HCON_00187985-00001 peptide:
HCON_00187985-00001
MSSTRAVLALFYMLLFSIAIVTSVPNRIFMRFGKRNVDLAEYRNPLPADYFPVELIGTSR
FTDGDM
>FLP-28_Haemonchus_placei_HPLM_0000601601-mRNA-1 peptide:
HPLM_0000601601-mRNA-1
MSSTRAVLALFYMLLFSIAIVTSVPNRIFMRFGKRNVDLAEFRNPLPAEYFPVELIG
>FLP-28_Heligmosomoides_polygyrus_HPBE_0000807001-mRNA-1 peptide:
HPBE_0000807001-mRNA-1
MLSTRAVLALFYMLLFGIVIVTSAPNRILMRFGKRTIRSDSSGQVEDRFGPESPPHSDRQ
RKQVMLKFRLS
>FLP-28_Micoletzkya_japonica_MicoRS5524-mkr-S355-0.24-mRNA-1
peptide: MicoRS5524-mkr-S355-0.24-mRNA-1
MNTRSLLALFMVFVCIAALATAAPNRVLMRFGKRSRAFDDAAMVNAPLAFYGRGPYVPYG
AFEADAE
>FLP-28_Oesophagostomum_dentatum_OESDEN_06709 peptide: OESDEN_06709
MLSARAVLVLLYTLLIAVAVVSAAPNRILMRFGKRTADPQPFQPMVPSEYFPVELLGSSR
AVDTDM
>FLP-28_Parapristionchus_giblindavisi_Parapristionchus-mkr-S_7-1.11-
mRNA-1 peptide: Parapristionchus-mkr-S_7-1.11-mRNA-1
MNSRFVLALLVVVFALVACVTAAPNRVLMRFGKRSPVPLDEADFAPYRYSAYKAMYNPMM
ALGGEAGDLEM
>FLP-28_Pristionchus_arcanus_arcanus-ag_msk-S_11-7.41-mRNA-1
peptide: arcanus-ag_msk-S_11-7.41-mRNA-1
MNARQFIALILAFLYIVTLASAAPSRVLMRFGKRAVAGSRFDFHELPVASYGFLPYGAVN
PQYEGFDESSVDQ
>FLP-28_Pristionchus_entomophagus_entomophagus-mkr-S131-0.7-mRNA-1
peptide: entomophagus-mkr-S131-0.7-mRNA-1

MNTRQFIILLLSIVYLMAALTTAAPSRVLMRFGKRSSAARLDAPEVPAASYGFLPYGALY
PQFEGFEDGIADQ
>FLP-28_Pristionchus_exspectatus_exspectatus-mkr-S_589-0.37-mRNA-1
peptide: exspectatus-mkr-S_589-0.37-mRNA-1
SGTPTVIMNARQFVALVLALIYIVTLASAAPSRVLMRFGKRAVAGSRFDFHELPVASYGF
LPYGAVNPQFEGFEESSVDQ
>FLP-28_Pristionchus_fissidentatus_fissidentatus-ag_msk-S6-5.14-
mRNA-1 peptide: fissidentatus-ag_msk-S6-5.14-mRNA-1
MNTRQFVALLLAVCYLVAAVSAAPSRVMMRFGKRAAGAALLEPQAQHADAAMPYAFLPYG
AMFPQFEGFEDGVLADQ
>FLP-28_Pristionchus_japonicus_japonicus-mkr-S39-1.43-mRNA-1
peptide: japonicus-mkr-S39-1.43-mRNA-1
TPALIMNARKFIALVLTFVYIVALASAAPSRVLMRFGKRAAAASRFDAHEIPAASYGFLP
YGALYPQFEGFEDSAADQ
>FLP-28_Pristionchus_maxplancki_maxplancki-mkr-S210-1.63-mRNA-1
peptide: maxplancki-mkr-S210-1.63-mRNA-1
MNARQFIALVLALVYVVALANAAPSRVLMRFGKRAAAASRFDAHDIPAASYGFLPYGALY
PQFEGFEDSAVDHPKNSSVSMLRLIHSMKDLEPYTSFHISIVMVVGGEIVASRTKAIPVE
NSAHVATIAKCEEC
>FLP-28_Pristionchus_mayeri_mayeri-mkr-S241-5.24-mRNA-1 peptide:
mayeri-mkr-S241-5.24-mRNA-1
VPHTEMNARQLIALLLAFVCLVTPLTTAAPSRVLMRFGKRAAAARFETPEIAAASYGFLP
YGALYPQFEGFEDAPAVDQ
>FLP-28_Pristionchus_pacificus_PPA39822.1 peptide: PPA39822.1
SGTPAVIMNARQFIALVLALLYIVTLASAAPSRVLMRFGKRAVAASRFDFHELPVASYGF
LPYGAVNPQFEGFEESSVDQ
>FLP-28_Strongylus_vulgaris_SVUK_0001106701-mRNA-1 peptide:
SVUK_0001106701-mRNA-1
MKRIGLVAVLPAVTSQQWIQAHSYIAPAVAQESARTKRRIMLSARAIYALLYTLLIAVAI
VSAAPNRILMRFGKRTADPVQYRPFVPADYYPVELLGYVPCFPV
>FLP-28_Angiostrongylus_cantonensis FLP-28 predicted
SLLSISGIMLSARTFLTLFCILLFTVLYTSAAPNRILMRFGKRATNTVPYALVPLPDYYRVDLRRYVQ
QEFLVRN
>FLP-28_Caenorhabditis_angaria FLP-28 predicted
GFPPKKKHTHLTHLPTPPTPTVYKSKHIYAFLFKNKAPPTSSQHGAHLTYIFIPHSCVFLNFSFFREK
NWVIFLGMFSAKSILAFLCIMLVAFVSAAPNRVLMRFGKRAAQSAPLGYRFVPESVPLMEYYE
>FLP-28_Dictyocaulus_viviparus FLP-28 predicted
VTIKVSMLSARTLLIFFCMLLFSLLTIPAAPHRILMRFGKRTHNLMTWQTNDGSRTEFLTSPYYYPAE
VLRYIQYILLFSDLKVVHRFWYSLTSFVFFINLQI
>FLP-28_Diploscapter_coronatus FLP-28 predicted
KLMEFSNFFPKIMCLNFRIFEILKILAASNRVLMRFGKRSPVLLPLSRGMPLNGNNFAEYFE
>FLP-28_Diploscapter_pachys FLP-28 predicted
NFFFSIFEILKILAASNRVLMRFGKRSSVLLPLSRGMPLNGNNFAEYFE
>FLP-28_Heterorhabditis_bacteriophora FLP-28 predicted
KELIFFSFYTSENRDNLYNRFDLIPTYLGIMSARLMVTVFCFLLISMAMVSAAPNRILMRFGKRSSEP
VHLRLTGPGEYYPFEFVGSVPDNINVANIIYVNKILVYTYLTNSINYI
>FLP-28_Mesorhabditis_belari FLP-28 predicted
FQLQPFLHRYPKATPKMLTKALVWLCALLMIVSLVSAAPNRILMRFGKRSAPITGFPYEPVYVYSAEP
R
>FLP-28_Necator_americanus FLP-28 predicted
ADKKHKNHLVPGMLSARVVLALLYMLMITVAIATAAPNRILMRFGKRTADPMHFRPMAPADYYPQELF
G
>FLP-28_Nippostrongylus_brasiliensis FLP-28 predicted
IIHLEKNEKHSNSSTPVRALREQLIKSFLTITTNSFIHSFISEADMLSTRAVLGLLYVLLCGIVLVHS

VPNRILMRFGKRTMDPVNFRAIVPAEYYPVELLG
>FLP-28_Oscheius_tipulae FLP-28 predicted
NKPDWNGRVVFGAVSFQQYFLINQSPQTHSHHSGTMLNIRFLLALLVVLVVCSTVSLAAPNRILMRFG
KRAVGGIPVHLGGEYVPVDYIGWVFWTNVRSAIFSLPYGPDNDF
>FLP-28_Teladorsagia_circumcincta FLP-28 predicted
SISLLIEPYSRTTKPILSGIQPSMRIQLEFGLPVSKSFFLGSMLSTRAVLALFYVLVFGIVIVTSAPN
RIFMRFGKRNIDSAGYRFPLPGEYFPIELLG
>FLP-28_Parastrongyloides_trichosuri_PTRK_0000786400.1 peptide:
PTRK_0000786400.1
MTPIGITKVFILIMIIFMSSSQTTVAAPNRVMMRFGKRFSSFENEHPYNLHPLLQFEGSP
ENIYRLTSYINRNQLYPMIPEIDM
>FLP-28_Rhabditophanes.sp.KR3021_RSKR_0000795400.1 peptide:
RSKR_0000795400.1
MVADNKSSISLKCLVAFMIIFMFTTQLTDAAPSRVMMRFGKRLAPSYGNLFEGNQDYYQG
EPDPEEVMKLHTYLSRNQMFPIVFDD
>FLP-28_Steinernema_carpocapsae_L596_g3143.t1 peptide: L596_g3143.t1
MSPRTLLLAILVLLISSSISSAAPNRILMRFGRSDPNLRPQNAGLPSSFFRSFDASRFGD
DAPSAGGNYLYDPVE
>FLP-28_Steinernema_feltiae_L889_g30048.t1 peptide: L889_g30048.t1
MLYEQEEDQLISLVLNSHSHFLNPKTIISYYDVCNWIVDLHGIKWKQNVCSSSAQDIKVL
LLPFKSLVQSDSRQSKHVISMQQKQLIAVCLILTVICVVESAPNRILMRFGKRVYHQQPA
VGSSDHDIGSDYTYQYHNQPNYEILQNLGR
>FLP-28_Steinernema_monticolum_L898_g3110.t1 peptide: L898_g3110.t1
MSPRAVLLAIFVLLVSSSIGQSAPNRILMRFGRSDPSLRQNLSPNFFKAFEDSRFDSEGP
SADSHYLFEPIE
>FLP-28_Steinernema_scapterisci_L892_g1314.t1 peptide: L892_g1314.t1
MPVYSLLISVFRWCSDFPSIRLRSNSRIPFPPAQKMMLTRKHLLVACLILAAIFSSDAAP
NRILMRFGKRGYQQHSTAAPDYDDGSDYVYHYGTEEEVDNANRY
>FLP-28_Strongyloides_ratti_SRAE_X000120600.1 peptide:
SRAE_X000120600.1
MATIQISYGAPNRVMMRFGKRFAPYVVGHPFYQNSIYQHDDAPDNAYKINSYLNKNQPFP
VVIENN
>FLP-28_Strongyloides_venezuelensis_SVE_0144200.1 peptide:
SVE_0144200.1
MIALMVTFQVSIGAPNRVMMRFGKRFTPIENDRLLSYHRPLLQYEGSSDNNYKFNSYLHK
NPLFPVIIDNN
>FLP-28_Steinernema_glaseri FLP-28 predicted
SENKAPVVSYSTPTATLFVRLLLECSEIRHHPLFSASKQLSPPATVPHFRRSMSPRSILVALLLLLIG
SSIGQAAPNRILMRFGRSDPALRQNVAASQDVFRPFEGAPRFGDSDLAPGNFVFEPVE
>FLP-28_Strongyloides_papillosus FLP-28 predicted
INFLLCFNSFFYRDLKLTKIMASLKFTKTFFIFMIAFMVTFQVSLGAPNRVMMRFGKRFTPLENDRLL
SYHRPLLQYEGSSDNSYKFNSYLHKNPLFPVIIDNN
>FLP-28_Strongyloides_stercoralis FLP-28 predicted
FFFRSKVKTNMTSLRLTIITFIFVFAFMGTLHLSYSAPNRVMMRFGKRFTPYVIDHSSYQYPFFQHEG
FPDNFYKINSYINKNQPFPAVIESN
>FLP-31_Meloidogyne_arenaria_M.Arenaria_Scaff6937g061121 peptide:
M.Arenaria_Scaff6937g061121
MQPFNNNPHLPFSQRQIFTLLFVWFLVISILLTFDGTSALQGASEMETDVLEDDQIVVPW
KKLYRPRGPPRFGKRGLFVMNRQRNFPE
>FLP-31_Meloidogyne_javanica_M.Javanica_Scaff7742g049294 peptide:
M.Javanica_Scaff7742g049294
MQPFNNNPHLPFSQRQIFTLLFVWFLVISILLTFDGTSALQGASEMETDVLEDDQIVVPW
KKLYRPRGPPRFGKRGLFVMNRQRNFPE
>FLP-31_Meloidogyne_enterolobii FLP-31 predicted

KGNNSKNKLLGKKLYRPRGPPRFGKRGLLVMNRQRNFPE
>FLP-31_Meloidogyne_floridensis FLP-31 predicted
IYNSKNKLFSGKKLYRPRGPPRFGKRGLFVMNRQRNFPE
>FLP-31_Meloidogyne_graminicola FLP-31 predicted
SFFLLGKKLYRPRGPPRFGKRGLILMNRHNNYQE
>FLP-31_Meloidogyne_hapla FLP-31 predicted
MCLKTIRLLFHVSVFNFKKFQKTKKFQKKLLGKKLYRPRGPPRFGKRGLLLMNRQRNFPV
>FLP-31_Meloidogyne_incognita FLP-31 predicted
HHYQRIMQPFNNNPHLPFSQRQIFTLLFVWFLVISILLTFDGTSVLQGASEMETDVLEDDQIVVPWKK
LYRPRGPPRFGKRGLLVMNRQRNFPE
>FLP-32_Ancylostoma_ceylanicum_Acey_s0033.g2719.t1 peptide:
Acey_s0033.g2719.t1
MLARSVVFTLLFTILIVDAALPRLRPAKKAMRNSLVRFGKRGDMSDSVFQGESLGPGESD
GLFFEREQQRIPVQYSYY
>FLP-32_Ancylostoma_duodenale_ANCDUO_19190 peptide: ANCDUO_19190
MEECRRVLRPAKKAMRNSLVRFGKRGDMSDSVFQGESLGPGESDGLFFEREQQRIPVQYS
YY
>FLP-32_Angiostrongylus_costaricensis_ACOC_0000284201-mRNA-1
peptide: ACOC_0000284201-mRNA-1
MLFYSISSLGLSDGWAQRSLRVGEGERVGHSPTTIRKAAMLVRSIVLTLFISFLVIDAAV
PRLRPAKKAMRNSLVRFGKRADLSDTVFLGESFGPVDTDGLYFERTQPQNYAQFPYY
>FLP-32_Caenorhabditis_angaria_Cang_2012_03_13_00157.g5854.t1
peptide: Cang_2012_03_13_00157.g5854.t1
MLSFVHTLILALLCSIIFVEAMPRNRPAKKAMRNSLVRFGKRGGGEQLAADDVFLGESYG
PVETYEQFRGDSYANQAPSVIFY
>FLP-32_Caenorhabditis_brenneri_CBN17525.1 peptide: CBN17525.1
MLSFVHTLILALLCSIVFVEAMPSMRPAKKAMRNSLVRFGKRADPIVSDDVFLGESYGSA
DPYEYVPEHMSNRGASPVLLY
>FLP-32_Caenorhabditis_briggsae_CBG07702.1 peptide: CBG07702.1
MLSFVRTLIFALLCSIVFVEAMPSMRPAKKAMRNSLVRFGKRADPIGSDDVFLGESFGSV
GPYEYAPERMSNRGGSPVLLY
>FLP-32_Caenorhabditis_inopinata_Sp34__X0133400.t1 peptide:
Sp34_X0133400.t1
MLSFVHTLILALLCSIVFVEGMPSMRPSKKAMRNSLVRFGKRADPVGSDDVFLGESYGSA
DPYEYVPERLSIREMPSVLLY
>FLP-32_Caenorhabditis_japonica_CJA12197.2 peptide: CJA12197.2
MLSFVHTLILALIFSMVFVEGMPSMRPAKKAMRNSLVRFGKRADTVRSDDVFLGEVYGAP
DSYEYVPKHMSNPAPSVFLY
>FLP-32_Caenorhabditis_latens_FL83_01786 peptide: FL83_01786
MLSFVHTLILALLCSIVFVEAMPSMRPAKKAMRNSLVRFGKRADPVVSDDVFLGESYGSV
DPYVYVPEHMSNRGASSVLLY
>FLP-32_Caenorhabditis_nigoni_Cni-flp-32 peptide: Cni-flp-32
MLSFVRTLILALLCSIVFVEAMPSMRPAKKAMRNSLVRFGKRADPIGSDDVFLGESFGSV
DPYEYAPERMSNRGASPVLLY
>FLP-32Caenorhabditis_ramanei_FLP82_02366 peptide: FL82_02366
RKSGRSLTEAGSLLKAPCQVNHAPRRLHSYDMLSFVHTLILALLCSIVFVEAMPSMRPAK
KAMRNSLVRFGKRADPVVSDDVFLGESYGSVDPYEYVPEHMSNRGASSVLLY
>FLP-32_Dictyocaulus_viviparus_nDv.1.0.1.t02270 peptide:
nDv.1.0.1.t02270
XLLRPAKKAMRNSLVRFGKRADFSNGIILGESFGPIETDGTYFEQARPLNFIQYHPYY
>FLP-32_Diploscapter_coronatus_DCO_025106 peptide: DCO_025106
MQFSLKLFLIALIASCLVILATASPARPIFDKRAMRNSLVRFGKRADPLYQGDDFGPDGY
IESYAGHRMPALYIY
>FLP-32_Heligmosomoides_polygyrus_HPOL_0001698301-mRNA-1 peptide:

HPOL_0001698301-mRNA-1
MLARSLVLTLLISALIVDAALPRLRPAKKAMRNSLVRFGKRADLSDGPVFLGESFGPEES
SGYYSALAPARKPSQYFLY
>FLP-32_Mesorhabditis_belari_mbelari.g6261.t1 peptide:
mbelari.g6261.t1
MAPLRSLLIFLLGLLAISSVFAYPRHFEKRAMRNSLVRFGKRSDILQDSMESPLSSGREI
FVVDPTLLP
>FLP-32_Necator_americanus_NECAME_07322 peptide: NECAME_07322
MLARSIVFTLLFTILIVDAALPRLRPAKKAMRNSLVRFGKRGDVSDNVFLGESFGPGETD
GLYFEREQPKIPVQYSYY
>FLP-32_Nippostrongylus_brasiliensis_NBR_0000718701-mRNA-1 peptide:
NBR_0000718701-mRNA-1
MLARFVVLTLLISVLLVDATLPRLRPAKKAMRNSLVRFGKRADLVEPVYLGKSELFMREA
FGPAESEELYDAINQARKPSPYFYF
>FLP-32_Oesophagostomum_dentatum_OESDEN_01360 peptide: OESDEN_01360
MLARSVVFTLLITILVVDAAIPRLRPAKKAMRNSLVRFGKRSDYPDVFLGESVGPSETDG
LYFERDQQRAPVFHSYY
>FLP-32_Oscheius_tipulae_OTIPU.nOt.2.0.1.t07025 peptide:
OTIPU.nOt.2.0.1.t07025
MASSHVLVFVLVCSLVISEAVPRLRPAKKAMRNSLVRFGKRADYDVLPQDFQGGSYDPSQ
DSLEGFFPDSLHPIIYY
>FLP-32_Pristionchus_fissidentatus_PRIFISSI000018776.t1 peptide:
PRIFISSI000018776.t1
MQFRSFLVFFVIALLAVNAMANRPAKKAMRNSLVRFGKRSAASPDISAELPAYGRWEYLN
RVALDGAD
>FLP-32_Pristionchus_japonicus_TRINITY_c32791_g1_i1_loc1.t1 peptide:
TRINITY_c32791_g1_i1_loc1.t1
PIFSSSSSIMQFRSFIVFLVLALLAVDAMANRPSKKAMRNSLVRFGKRSAAAASDFSAEA
APSYARWEYLNRLAAMDGADGYNL
>FLP-32_Pristionchus_pacificus_PPA20622.1 peptide: PPA20622.1
MQFRSFIVFMVLALLAVDAMANRPSKKAMRNSLVRFGKRSAAAPSDFSAEAAPSYARWEY
LNRLAALDGADNGYNL
>FLP-32_Ancylostoma_caninum FLP-32 predicted
LWEQYNCRLRPAKKAMRNSLVRFGKRGDMSDSVFQGTYSSSLLWIIPK
>FLP-32_Angiostrongylus_cantonensis FLP-32 predicted
CYSLRPAKKAMRNSLVRFGKRTDLSDAVFLGQICINHFDELSFGSQILQQFSHYLLTYLIFT
>FLP-32_Caenorhabditis_sinica FLP-32 predicted
NVSSRSTRVVTNTSQVGVHKENGSFFSMRPAKKAMRNSLVRFGKRADPVGADDVFLGEY
>FLP-32_Caenorhabditis_tropicalis FLP-32 predicted
EVSFFSMRPAKKAMRNSLVRFGKRADPIASDDVFLGKLLAHSTLTNSILSGESYGSADPYEYVPEHMS
NRAASPVLLY
>FLP-32_Cylicostephanus_goldi FLP-32 predicted
XXXXXXXXXXXXXXXXXXXXXXXXXXXXXXXXXXXXXXXXXKKRFEIYFFQRKYFLRLRPSKKAMR
NSLVRFGKRSDVPDNVFLGKNPLFLYIFYFSNWIAISRRIFRAE
>FLP-32_Diploscapter_pachys FLP-32 predicted
MTSIFRIIYAVVYKNSKFFDVWFVKCAQSFARRKKLVVPIAQPLNHAGSPSAVCRRALSIAGARIEIV
MFFEENCKTYNVQNRHLTKEIAKIIIFRPIFDKRAMRNSLVRFGKRADSLYQGKKGK
>FLP-32_Heterorhabditis_bacteriophora FLP-32 predicted
KALAIVQKQNILTFCNLAHFRSRPAKKAMRNSLVRFGKRTDPTDNAVFLGKKTNNEYEIVLLITELVM
TNFLFAAYNYFTLACKV
>FLP-32_Micoletzkya_japonica_Micoletzkya_japonica FLP-32 predicted
RRYFISNSALFFRPSKKAMRNSLVRFGKRAAALPEQGEYVPVDERWAAFLDRAGLVDGEYQY
>FLP-32_Parapristionchus_giblindavisi FLP-32 predicted
LITFHFSPSKKAMRNSLVRFGKRAFPLDGSEELAVDPKWALFYNRNIDGEGYYAQ

>FLP-32_Pristionchus_arcanus FLP-32 predicted
CIRPSKKAMRNSLVRFGKRSGSAPSDFSAE
>FLP-32_Pristionchus_entomophagus FLP-32 predicted
KDAHIEEGGCASLSRRNSEKFLSARILLGLFGAEATLQIYPTSRPAKKAMRNSLVRFGKRAAAPDYSA
E
>FLP-32_Pristionchus_exspectatus FLP-32 predicted
RRFTSGFNINLYKLRVRPSKKAMRNSLVRFGKRSASAPSDFSDE
>FLP-32_Pristionchus_maxplancki FLP-32 predicted
LNVIIRPAKKAMRNSLVRFGKRSASAPSDFSAE
>FLP-32_Pristionchus_mayeri FLP-32 predicted
MWSRNIIIVRPAKKAMRNSLVRFGKRAAPIDYSAE
>FLP-32_Strongylus_vulgaris FLP-32 predicted
FFLKLTKFFRLRPSKKAMRNSLVRFGKRSDVPDNVFLGIILSLL
>FLP-32_Caenorhabditis_elegans FLP-32
MLSFVQTLILALLCSIVFVEAMPSMRPAKKAMRNSLVRFGKRADPVGTDDVFLGESYGSADPYEYVPE
RMSNRGPSSVLL
Y
>FLP-33_Plectus_sambesii_PSAMB.scaffold472size50076.g6039.t1
peptide: PSAMB.scaffold472size50076.g6039.t1
MCSSLRFFLLLLTLSLQCDDTSCFGQWLMVQPFHWRQRGWLSFPATSRMNSKPFMRRDIE
NSYQLDPLTDADNLRTILKGFDQIRKPRFG
>FLP-33_Ascaris_suum_AgR029_g051_t02 peptide: AgR029_g051_t02
MISMLLLVSFFALAASDIADLPSQHVLRRFEFLPLFLQTKYPGEIIPEKRVSIKERRAPI
EGFEDFDGIMRSLDGLQKPRFGRK
>FLP-33_Ancylostoma_caninum_ANCCAN_00652 peptide: ANCCAN_00652
MNFTTLFAFLLSLLTVGAFSVDLPQYYGGYGQLRRGGEGEYVKRMAPMKYRRSPLEGFEE
LSSMMRSIDGIQKPRFGRK
>FLP-33_Ancylostoma_ceylanicum_Acey_s0489.g2376.t1 peptide:
Acey_s0489.g2376.t1
MNFTTLFAFLLSLLTVGAFSVDMPQYYGGYGQLRRSGDSEYVKRMAPMKYRRSPLEGFEE
LSSMMRSIDGIQKPR
>FLP-33_Angiostrongylus_cantonensis_ACAC_0000227401-mRNA-1 peptide:
ACAC_0000227401-mRNA-1
MPMKNRRSPLDGFEDLSSVMRSIDGIQKPRCVSDIFLIPANYYCCKENF
>FLP-33_Angiostrongylus_costaricensis_ACOC_0000733601-mRNA-1
peptide: ACOC_0000733601-mRNA-1
MPKIAAVSERIWSLATTFSNLFRRIMPMKNRRSPLDGFEDLSSVMRSIDGIQKPRYVSDI
FTIPVNYFHYKKSLSYHYMIRNHSKKCLITKKPRLVAVRHPPLSNSLTIY
>FLP-33_Caenorhabditis_brenneri_CBN15572.1 peptide: CBN15572.1
MPSQVVGYSVEPRLASFNDGGELAAQAAVAAAAAQQARSEFMKRFLPAKERRAPLEGFED
MSGFLRTIDGIQKPRFG
>FLP-33_Caenorhabditis_briggsae_CBG08711.1 peptide: CBG08711.1
MRLLIIVAIATIVLATSQVSAYSVEPRLAFVDSGAAEMSQAARQARNNELEFIKRFLPAK
ERRAPLEGFEDMSGFLRTIDGIQKPRFG
>FLP-33_Caenorhabditis_inopinata_Sp34__10037700.t1 peptide:
Sp34_10037700.t1
MRLLLIIVVALISLAVSDRGYAVEPNLYVDANGNGESAAQQRAQNPELEFIKRFLPAKER
RAPLEGFDDMSGFLRTIDGIQKPRFG
>FLP-33_Caenorhabditis_japonica_CJA26229.1 peptide: CJA26229.1
MALLAITITTTTAAPSSHQYPLNYLDAADQDTWTRFLPVKNRRAPLDGFEDMSGFLRTID
GIQKPRFG
>FLP-33_Caenorhabditis_latens_FL83_09229 peptide: FL83_09229
MRLLIIVAAIAFIAISTSQSVAAYSVEPRLATFSDGSAAELARQARNNELEFIKRFLPAK
ERRAPLEGFEDMSGFLRTIDGIQKPRFG

>FLP-33_Caenorhabditis_nigoni_Cni-flp-33 peptide: Cni-flp-33
MRLLIIVAIATIVLATSQASAYSVEPRLAFVDSGAAEMSQAARQARNNELEFIKRFLPAK
ERRAPLEGFEDMSGFLRTIDGIQKPRFG
>FLP-33_Caenorhabditis_remanei_CRE14218.1 peptide: CRE14218.1
MRLLIIVAAIAFIAISSSQSVAAYSVEPRLATFSDGSAAELARQARNNELEFIKRFLPAK
ERRAPLEGFEDMSGFLRTIDGIQKPRFG
>FLP-33_Caenorhabditis_tropicalis_Csp11.Scaffold574.g4340.t1
peptide: Csp11.Scaffold574.g4340.t1
MRLLIIVAVIALVAFSATSAYSTEPRLAFADREMSPAAAQRARNTELEFIKRFLPAKERR
APLEGFEDMSGFLRTIDGIQKPRFG
>FLP-33_Diploscapter_coronatus_DCO_024654 peptide: DCO_024654
MLRRLSSLICLLLISLISAYVAPSIEQPPANSQAANFFYQRQLRDPLYEPYIRRMMPMKE
RRNFEGFEDMGSLMRSIDGIQKPRFG
>FLP-33_Haemonchus_contortus_HCON_00009870-00001 peptide:
HCON_00009870-00001
MNFTALLSFLLSLLVVNAFNIDYPQYGGYGAIRRAVERDYAKRMLPMKNRRSPLEGFEDI
SSMMRSIDEIQKPRFGRK
>FLP-33_Heligmosomoides_polygyrus_HPOL_0002087601-mRNA-1 peptide:
HPOL_0002087601-mRNA-1
MNFTMLLSFLLSLLVVGAFNVDYPQYYGGGYGAMRHALESDYAKRMMPMKNRRSPLEGFD
DISSMMRSIDGIQKPRFGRK
>FLP-33_Necator_americanus_NECAME_00947 peptide: NECAME_00947
MNFTTLLALLLSLLAVGAFNVDFPQYYGGYGQIRRDGEEQYAKRAMPMKFRRSPLEGFEE
FSSMMRSIDGIQKPRFG
>FLP-33_Nippostrongylus_brasiliensis_NBR_0000985601-mRNA-1 peptide:
NBR_0000985601-mRNA-1
MNFTSILSFLLSLVVVGAFNFDYPQYFGGYGAIRRSMLPSKNRRSALEGFDDISSMMRSI
DGIQKPRFGRK
>FLP-33_Oesophagostomum_dentatum_OESDEN_21756 peptide: OESDEN_21756
MKSITVVTFILSLLVLNSMGLETRYGKYGPMRLAGDEQVINSYTPIKYRRASIDAYGDFS
SMMRSMDELQRPRFGRK
>FLP-33_Oscheius_tipulae_OTIPU.nOt.2.0.1.t11991 peptide:
OTIPU.nOt.2.0.1.t11991
MQILVTLGLLFALVCASFAAPFDDFRYFQLPVRRGFPEPDYVKRLMPKERRAPAADFDDM
SSFMRSLDGIQKPRFG
>FLP-33_Pristionchus_entomophagus_PRENTOPHAG000017361.t1 peptide:
PRENTOPHAG000017361.t1
PTHTSRMTSSSSLALICLVTLAVTLVSAWPADAAAARPLLTPAEREYLKELLAERDFVLT
NPAKARRSPGDEGFDFTSALRSIDNIQKPRFGRRR
>FLP-33_Pristionchus_expectatus_PEXSPEC000026662.t1 peptide:
PEXSPEC000026662.t1
TDTHRRMTSSSSLILCFVAVLIAASASAWPAEGGPAAGAAAAPRPLTPLEREYLRELLAA
ERDFVLTNPVKSRRSPSDEGFDFTSALRSIDNIQKPRFGRRR
>FLP-33_Pristionchus_fissidentatus_PRIFISSI000027361.t1 peptide:
PRIFISSI000027361.t1
IQVPTMTSSSSLLVFCTLFILCVLSSAWPTESRLSPEERQLLREVLAERDFVMTNPVKAR
RSPSMEGFDFTSALRSIDNIQKPRFGRRR
>FLP-33_Pristionchus_macplancki_PRIMPAX000017480.t1 peptide:
PRIMPAX000017480.t1
TMTSSSSLVLCFLAVLIAAALPSASAWPAEGGPAAARPLTPVEREYLRELLAERDFVLTN
PVKSRRSPSDEGFDFTSALRSIDNIQKPRFGRRR
>FLP-33_Pristionchus_mayeri_PRIMAYER000032387.t1 peptide:
PRIMAYER000032387.t1
ASAWPADKDAAPRPLTPGEREFLRELLAERDFVLTNPVKTRRSPSDEGFDFTSALRSIDN

IQKPRFGKRR
>FLP-33_Pristionchus_pacificus_PPA04942.1 peptide: PPA04942.1
MTSSSLILCFVAVLIAAAAALPSAWPAEGGPAAGAAAPRPLTPLEREYLRELLAAERDFV
LTNPVKSRRSPSDEGFDFTSALRSIDNIQKPRFGRRR
>FLP-33_Caenorhabditis_elegans FLP-33
MRFLILIVAIVLLSAVHGFSVEPRLAAFADGGAAELAQEARQARNAELEFIKRFLPAKERRAPLEGFE
DMSGFLRTIDGI
QKPRFG
>FLP-33_Caenorhabditis_angaria FLP-33 predicted
SELALLHFQISCRVPSKDRRAPLEGFEDMSGFLRQIDGIQKPRFGRK
>FLP-33_Caenorhabditis_sinica FLP-33 predicted
NLRVVETLTAKSILYLTFRRFLPAKERRSPLEGFEDMSGFLRTIDGIQKPRFG
>FLP-33_Dictyocaulus_viviparus FLP-33 predicted
DFRRVLPMKIRRSPLDGFEDLSSVMRSIDGIQKPRFVRNKKIFLARSITLSVYNLLIFIHKQINNESM
HLLQKSIHIQL
>FLP-33_Panagrellus_redivivus_Pan_g36.t1 peptide: Pan_g36.t1
MAMLSISWIMLGCFLLATVFSAVYGVPVGSSSDNGGHRLPVVPLGWLIPHEPQVVGTWLP
EANRLRRAGPIPFKERRASADTFDMWEGMMDSLDTLQKPRFGR
>FLP-33_Steinernema_carpocapsae_L596_009024 peptide: L596_009024
MQVALFLLVLLVVTVSLSEAMFKHPIGLMAYSAAGGPKPLLRGYRPLLPDEVEIKERRAP
WGEGSEWEGMLQTLDNLRKPRFGK
>FLP-33_Steinernema_feltiae_L889_g32186.t1 peptide: L889_g32186.t1
MQVALFILVLLAATVTFSDAFLKRPVGLMAYGAATGHRPLLRGYRQLLPDEVEMKERRAP
SDPFDEWEGMLQHLDNLRKPRFGK
>FLP-33_Steinernema_glaseri_L893_g4388.t1 peptide: L893_g4388.t1
MQVALFFVLALFAATVSLSDALFKRPIGLMAYGAAGGPKPLLRGYRPLLPDEVEIKERRA
PSDPFDEWEGMLQTLDNLRKPRFGK
>FLP-33_Steinernema_monticolum_L898_g16279.t1 peptide:
L898_g16279.t1
MQVALFILVVLAATVSLSEAFFKRPIGLMTYGAASGSRPLLRGYRPLLPDEVEMKERRAP
SDPFDEWEGMLQTLDNLRKPRFGK
>FLP-33_Steinernema_scapterisci_L892_g10634.t1 peptide:
L892_g10634.t1
MQVALFILVLLFATVSLSEAMFKHPIGLMAYGAAGGPKPLFRGYRPLLPDEVEIKERRAP
SDPYDEWEGMLQTLDNLRKPRFGK
>FLP-33_Ditylenchus_destructor_Dd_00688 peptide: Dd_00688
MHPTFLYRPTVSALTTSSLILCMITILLPQGQCVQAEVADLADEDIPIPAEPAYTPQKSE
QPSKFSTAELSRADMYKLLYTALLRDKERAQWQPSRLRPIAPLQSRTPLSALRHSTDSSW
RMKLAEIPIYEQRRSMKERRAPAPLDHFSEQWGGMMKTMDNLRKPRFG
>FLP-33_Globodera_rostochiensis_GROS_g00355.t1 peptide:
GROS_g00355.t1
MRPIAAVPSPFLPLVVAFGLFSTGSVSYMFLPATRQLFVNPDILSSDAALTPRGFHIADF
LGPNKRFLDGEAPPATIFIGPKERRAPAERSFGEESQWLKAMDNLRKPRFG
>FLP-33_Ditylenchus_dipsaci FLP-33 predicted
TVGGMMKTMDNLRKPRFG
>FLP-34_Plectus_sambesii_PSAMB.scaffold234size63179.g3439.t1
peptide: PSAMB.scaffold234size63179.g3439.t1
MINTQTVAAVIIGTLAVLLATANAAPAVAPAIKKATFNLDDIASAANGASRLRYGKRGGN
RDESYSGIDPELFDTIVSLNNVGKRSGSTADAELFDNLHRFAEAGKRSNLVSDRLSRLAE
ALNGAERPRFG
>FLP-34_Acanthocheilonema_viteae_nAv.1.0.1.t08907-RA peptide:
nAv.1.0.1.t08907-RA
MTTVYWFIGILLLPVLFVCNVPVDPMKRTDKNLEQFTSAINAALRLRYGKRSENPNLFSN
KIVPLPEKDKLSQLANRVVASLNEAERLRFG

>FLP-34_Anisakis_simplex_ASIM_0001685901-mRNA-1 peptide: ASIM_0001685901-mRNA-1
MPFAHVFFDAAIAILIVACAVSALPMHSKKSAHDLQGFASALNGAGRLRYGKRSYDPRLY
PEFFPALDDTGRLTISRLGSDEDMEDPLL
>FLP-34_Ascaris_lumbricoides_ALUE_0001331401-mRNA-1 peptide: ALUE_0001331401-mRNA-1
MSTIQLITAAFVVFLTLTVRGFTIQSKKSAHDLEGFASALNSASRLRYGKRSYDLSLLET
LADLEKFGKRPSTDFDNDDNVEQFIESNKRSSSFLPMDKVVASLNRAERLRFG
>FLP-34_Ascaris_suum_GS_15307 peptide: GS_15307
MSTIQLITAAFVVFLTLTVRGFTIQSKKSAHDLEGFASALNSASRLRYGKRSYDLSLLET
LADLEKFGKRPSTDFDNDDNVEQFIQMQQNEREDHKENAKTSIAFG
>FLP-34_Brugia_malayi_Bm14205.1 peptide: Bm14205.1
MTKVYLFAGTLLLLVLLVSNVQTDRARRSNNDLKKVASAINGALRLRYGKRSDDSELFNK
IILPQEENKLSPLADRVVASLNEAERLRFG
>FLP-34_Brugia_pahangi_BPAG_0000015701-mRNA-1 peptide: BPAG_0000015701-mRNA-1
MTTVYLFAGTLLLLVLLVSNVQTDRTRRSNKDLEQFASAINGALRLRYGKRSHDSELFNK
IILPQEENKLSPLADRVVASLNEAERLRIFNAQEETISDLAEIKIKCLEWLKYKPIFLSP
LSNRLVIIDSILI
>FLP-34_Brugia_timori_BTMF0001533601-mRNA-1 peptide: BTMF_0001533601-mRNA-1
MRNVQTDRARRSNNDLKKVASAINGALRLRYGKRSDDSELFNKIILPQEENKLSPLADRV
VASLNEAERLRICNTQEEIISDLAEIKIKCLEWLKYKPIFLSPLSNRLVIIDSILI
>FLP-34_Dirofilaria_immitis_nDi.2.2.2.t04631 peptide: nDi.2.2.2.t04631
MTRICLFVSTLLLSALLICNVPMDRAERVDRDLGQFTSAINGALRLRYGKRFYNPAIFVP
SQEVGNKYLGPFDDTVEIPIEKQAVIDALVKAINEAEPLRYG
>FLP-34_Elaeophora_elaphi_EEL_0000502501-mRNA-1 peptide: EEL_0000502501-mRNA-1
MTAVYLFIGILLLPVLLVCNAPVDRIQRSDRNLDQFASAINGALRLRYGKRSEDLDLLDK
LVQSPQVKDRYFGPSYNHFNLPM
>FLP-34_Gongylonema_pulchrum_GPUH_0000390201-mRNA-1 peptide: GPUH_0000390201-mRNA-1
MFTVCLFAAAALLIQSASAVPADRGRRSDRDLEQFASALNGALRLRYGKRSYDPNLFDKI
FTLQFVLSIHSATV
>FLP-34_Loa_loa_EFO27505.1 peptide: EFO27505.1
MDRARRSDKALEQIASAINGALRLRYGKRSHNPDLYDKIITPPEEDKLLPLADRIVASLN
RAERLRYAIHFDENLDKNPIYKNFLQYAKYQINMAGLQKGLDETDLI
>FLP-34_Onchocerca_flexuosa_X798_04709 peptide: X798_04709
MTTIYLFVGTLLLPALLVCNVPMDRAERLDRDLEQFTSAINGALRLRYGKRSYDSDMVGK
TVRSQEVGDRSFEPSYDHFYIPIEEQKQLPLVARVIASLNGAERLRYANHFGENLNYKTF
SQKTFCIICKRKMR
>FLP-34_Onchocerca_onchengi_nOo.2.0.1.t01233-RA peptide: nOo.2.0.1.t01233-RA
MTTIYLFAGTLLLPALLVCNVPMDRAERLDKDLEQFTSAINGALRLRYGKRSYDSDVFGK
TVRSQEVGDRYFEPSYDHFYLPIEEQKQLPLADRVIASLNGAERLRFG
>FLP-34_Onchocerca_volvulus_OVOC6916.1 peptide: OVOC6916.1
MTTIYLFAGTLLLPALLVCNVPMDRAERLDKDLEQFTSAINGALRLRYGKRSYDSDVFGK
TVRSQEVGDRYFEPSYDHFYLPIEEQKQLPLADRVIASLNGAERLRFG
>FLP-34_Parascaris_univalens_PgR057_g064_t01 peptide: PgR057_g064_t01
IALNAMSTIRLISAAFVVFLTLTVRGLTSQSKKSVHDLEGFASALNSASRLRYGKRSYDP
SLLETLAELEKSGKYPTTDFDSDNNMELIESKRSSSFLPMDKVVASLNRAERLRFG
>FLP-34_Thelazia_callipaeda_TCLT_0000744801-mRNA-1 peptide:

TCLT_0000744801-mRNA-1
MLKKVVVIIINDPVLMRYGKIFKKITTLYLFYEQLYYCLAESTVCWRNYKLKSSLHLLHQ
ESVPLLHCYQKLYKVKKGQNMIREQYPRESVRNSKYNEKLLLNWHIASTSYINLSILDKF
EQNYVQMITVMKLLSHDMNCSSINDALSVDRSRRMDHDNNLKQFASAINDALRLRYGKRS
YDPNLYDEIATFQQLGKQYYLPYYDDFYLHNQLQKRSLSTDRVIASLNGAERLRFKHEGL
EIRVRTRTHTQTFRTNFIIINSLLILYLKEY
>FLP-34_Toxocara_canis_Tcan_01874.1 peptide: Tcan_01874.1
MVRRQAWALSKAIGKGRHTSDGDGSRKISSAHMFLITALPAQSKKAAHDLQGFASALNGA
SRLRYGKRSYDPYILQTLATQAELEKPSAFDFDNYDNGDQFIESHKRSSLLPIDKVVASL
NGAERLR
>FLP-34_Wuchereria_bancrofti_WBA_0000078701-mRNA-1 peptide:
WBA_0000078701-mRNA-1
MTTVYLFAGTLLLLVLLVSNVQMDRARRSNKDLEQFASAINGALRLRYGKRSHDSELFSK
IILPQEENKLSPLADRVVASLNEAERLRIRKPQEEIISDLAEIKIKCLEWLQIQADLFIT
IIKLYSDYR
>FLP-34_Ancylostoma_caninum_ANCCAN_08994 peptide: ANCCAN_08994
MSDIFWCALREQADYDDPYIYAKRAPLTNKLIQSLNGAERLRFGRK
>FLP-34_Ancylostoma_ceylanicum_Acey_s0089.g2219.t1 peptide:
Acey_s0089.g2219.t1
MEIIDGVSSINIHPLFSLPAPSPTRIKTPEVPPTSLPYLLYPSSSNRCVVRVRCHGIHHI
TINMRSLICYALAVLTVIYSALPVLSFDKKSDMSEFTSAINGASRLRYGKRSFNFGYPLN
ALRDPTDYEDPYIYAKRAPLTNKLIQSLNGAERLRFGRK
>FLP-34_Angiostrongylus_cantonensis_ACAC_0000781201-mRNA-1 peptide:
ACAC_0000781201-mRNA-1
MSEFTSAINGASRLRYGKRNAALDFPLHVLRTYDDYELPYETFEKRAPLTNKFIQSLNGA
ERLRFIITDIAILLKQIIICQGLVENEQRRITDRHDGGDVDEYDDGRL
>FLP-34_Angiostrongylus_costaricensis_ACOC_0001285401-mRNA-1
peptide: ACOC_0001285401-mRNA-1
MQSQPLVVFCFVLLFSIAVASVLPLNKKSDMSEFTSAINGASRLRYGKRNAVLNFPLHEK
RAPLTNKFIQSLNGAERLRFLLIQIVICQGLVGNERQRITNRYDGGDVNEYDDGQWNHGQ
AQMAVLAVMLVTIVVVMVAITTSDFFKRKSRGRGTQHKCKKGILPTTFGPFPPQKCSHFC
C
>FLP-34_Caenorhabditis_angaria_Cang_2012_03_13_00259.g8040.t1
peptide: Cang_2012_03_13_00259.g8040.t1
MLHSLMLFLVVLFVSVLSQPMDKKSDMSGFASALNNAGRLRYGKRSSWDIEQPDAAEFES
YDSYYPELLKRGLNTDSLVASLKGAERLRFGRK
>FLP-34_Caenorhabditis_brenneri_CBN06803.1 peptide: CBN06803.1
MQFKHLMALILVVLALTESVLALPLEKKADISTFASAINNAGRLRYGKRSDPAVWDENSA
IYPAADQYIYTEEGRYPYGLIKRALNTDSLVASLNGAERLRFGRK
>FLP-34_Caenorhabditis_briggsae_CBG17891a.1 peptide: CBG17891a.1
MQFKILMALIVTFVAMNESVLALPLEKKADINTFASAINNAGRLRYGKRSDPSVWEDNMP
IFPAADQYIYTEEGRYPYAQIKRALNTDSLVASLNGAERLRFGRK
>FLP-34_Caenorhabditis_inopinata_Sp34__50000500.t1 peptide:
Sp34_50000500.t1
MMIKIFVPFILTVLMITDSVLSLPLEKKADISTFASAINNAGRLRYGKRSEPDLWEKNNI
GIPAAEEQYVYSGDRYPYSLIKRAFNTDTLVDSLNGAERLRFGRK
>FLP-34_Caenorhabditis_japonica_CJA35737.1 peptide: CJA35737.1
MQFKVFAALILVVCSVADSVFAVPLEKKADISRFTSAINSAGRLRYGKRSDPVTIWEE
>FLP-34_Caenorhabditis_latens_FL83_15258 peptide: FL83_15258
MQSFTTELIIFFTIILSISVLALPLEKKADINTFASAINNAGRLRYGKRSDPAAWEDNTP
IFPSADQYVYSEEGRYPYALIKRALNTDSLVASLNGAERLRFGRK
>FLP-34_Caenorhabditis_nigoni_Cni-flp-34.2 peptide: Cni-flp-34.2
MQFKILMALILAFVAMNESVLALPLEKKADINTFASAINNAGRLRYGKRSDPSVWEDNMP
IFPAADQYIYTEEGRYPYAQIKRALNTDSLVASLNGAERLRFGRK

>FLP-34_GCK72_019108.t1 peptide: GCK72_019108.t1
MQFKFLVSLILVVLALSESVLALPLEKKADINTFASAINNAGRLRYGKRSDPAAWEDNTK
IFPSADQYVYSEEGRYPYALIKRALNTDSLVASLNGAERLR
>FLP-34_Caenorhabditis_sinica_Csp5_scaffold_00112.g4638.t1 peptide:
Csp5_scaffold_00112.g4638.t1
MSKKINIEITVRASKQDVLALPLEKKADINTFASAINNAGRLRYGKRSDPAMWEEKKWSK
PSRFLSFYSDQDSAIFPAADQYVYSEEGRYPYGLIKRALDTDSLVASLNGAQRLRFGRK
>FLP-34_Dictyocaulus_viviparus_DICVIV_12876 peptide: DICVIV_12876
MLSHKSKSPSQLPHVLTVNKKSDMSKFTSAINGASRLRYGKRNIDFGYPLPDKRSALSNK
FIQSLNEAERLSVYFMGAMNSTKQIGLATQEKLQRLDESRSRSIRIVLDVDDGDA
>FLP-34_Diploscapter_coronatus_DCO_024341 peptide: DCO_024341
MQYGSAIECALLFLFSLLLLSVSASPLNVFDKRQDLSAFTSAINGAGRLRYGKRAQPFAF
PIYSDDELNGAEQGWTSNDMLKRSPVASKLIQSLNGVDRLRFGRK
>FLP-34_Diploscapter_pachys_WR25_25817.1 peptide: WR25_25817.1
MQYGLAIECALLFLFSLLLLSVSASPLNVFDKRQDLSAFTSAINGAGRLRYGKRAQPFAF
PIYSDDELNGAEQEWTSNDMLKRSPVASKLIQSLNGVDRLRFGRK
>FLP-34_Haemonchus_contortus_HCON_00140260-00001 peptide:
HCON_00140260-00001
MRSLFCYFVAALTLLYNMLPVSSVDKKSDLSDFASAINSAGRLRYGKRSSDYGYPLSAFN
EPVEFEDPYARMVFEKRAPITSKLIQSLNEAERLRFGRK
>FLP-34_Heligmosomoides_polygyrus_HPOL_0000532101-mRNA-1 peptide:
HPOL_0000532101-mRNA-1
MQHLVLFTVVLCSAAVYSVSSFEKKSDMSEFTSAINGASRLRYGKRSFDVGYPLNALREP
IEYEDPYTQYAKRAPLTNKLIQSLNGAERLRTWGYNMAEQVERAFLKQPTIKLNSRARIL
AGNKRVPRYVRNIGLGFKTPHEASQGSYIDKKCPWTGGNCAIRGNILTGVVLKNKMTRTI
DVQPHCSPAFRDIAPGDLVTIGECRPLSKTVRFNVLKFHKSGSTKKGFAKF
>FLP-34_Heterorhabditis_bacteriophora_Hba_16449 peptide: Hba_16449
MQQFPLAVLVFCAVAILSVLSFEKKSDMSEFASAINGASRLRYGKRSFDYGSFLPMKSIK
DSEDYDEPTYAVYVKRAPLTNKLIQSLNGAERLR
>FLP-34_Mesorhabditis_belari_mbelari.g1519.t1 peptide:
mbelari.g1519.t1
MQASLLSALILAFVAAVSCASLAKKADEVSSFVSGLNGAARLRYGKRSPTDNQESSFLVD
QPMLMDNEYVLQPYGSNGYALYKRSPSAKSLIESLNGAERLRFGRK
>FLP-34_Micoletzkya_japonica_MIJAPON000017614.t1 peptide:
MIJAPON000017614.t1
QVMQGSLALSAVLILAVQLAVASSYEKKSGDLSEFASALNGAGRLRYGKRSMPFLDSFKE
KMARGFDDDDSMAVFTLYDSIPYEKRAPSPDFVTSLNKAERLR
>FLP-34_Necator_americanus_NECAME_08388 peptide: NECAME_08388
MIEFDFSVLSFEKKSDMSEFTSAINGASRLRYGKRNFDFAYPLTALRDPTDFEDPYIYDK
RAPLTNKLIQSLNGAERLRFGRK
>FLP-34_Nippostrongylus_brasiliensis_NBR_0000109201-mRNA-1 peptide:
NBR_0000109201-mRNA-1
MNCSFHLRLFPIAIQPFFSASSLDKKSDMSEFASAINGASRLRYGKRSFEFSYPLNTNTH
SPCVHDLNGARLLGRVPTGLIDTDTLTFSITFYSSYFTLSARSRRFRRTSPIGIGTACIL
LF
>FLP-34_Oesophagostomum_dentatum_OESDEN_20739 peptide: OESDEN_20739
MRALFCYLFAAATVLYSIVPVLSFEKKSDMSEFTSAINGASRLRYGKRSFDFGYPLKLVF
LKIITKLVTITALETLFFGHKKNL
>FLP-34_Oscheius_tipulae_OTIPU.nOt.2.0.1.t00834 peptide:
OTIPU.nOt.2.0.1.t00834
MASPYSTSKHKFSTMQCSILVASLSVAVLFQCVSASGYVKKSDINEFASAINGAGRLRYG
KRGNWDSLYPESLWETEDESENTNNENSNLQYAKRAPVPSNLLQNLKGAERLRFGRK
>FLP-34_PARAPRI000014405.t1 peptide: PARAPRI000014405.t1
MERFYSIFFVLAITLQCISSSSLEKKSDMGDFASALNGASRLRYGKRSGNGGLFFLDSFR

DRVEDDGDYMPMNAEKRSPSTDFIANLNKAERLRY
>FLP-34_Pristionchus_arcanus_PARCANUS000012775.t1 peptide:
PARCANUS000012775.t1
MKQTGLRSTKNGRWLYESGGFLVLYDFSGRRQKESKNKKSVFLPYPNWYCRLSISPARLS
NMQLVILCGIVALACLGEADDFAQFEKKADLGEFASALNGAGRLRYGKRSSFPYIDFRER
MEHGASYDSEPFTFEKRAPSAADFVANLNKAERLR
>FLP-34_Pristionchus_expectatus_PEXSPEC000027703.t1 peptide:
PEXSPEC000027703.t1
MQLVIMCSIVVLACLGEADDFAPFEKKADLGEFASALNGAGRLRYGKRSPVGAFPYIDFR
ERMEHGASYDSEPFTFEKRAPSAADFVANLNKAERLR
>FLP-34_Pristionchus_fissidentatus_PRIFISSI000018650.t1 peptide:
PRIFISSI000018650.t1
MVRCVSALLALALTLDVVLGGASFEKKSDLGEFASALNGAGRLRYGKRSSGGSPYFDALR
ERMMERGFDEDPFMINFEKRAPSPADFVANLNKAERLRFGRK
>FLP-34_Pristionchus_japonicus_TRINITY_c1733_g1_i1_loc1.t1 peptide:
TRINITY_c1733_g1_i1_loc1.t1
MVRCVSAVVVLVLVLDAVLCSPFEKKSDLGEFASALNGAGRLRYGKRSFGGAFPYADFRE
RMEHGAAYDSEPFTFEKRAPSPSADFIANLNKAERLR
>FLP-34_Pristionchus_macplancki_PRIMPAX000018423.t1 peptide:
PRIMPAX000018423.t1
MVRCVSAVLALILVLEAVLCAPFEKKADLGEFASALNGAGRLRYGKRSSGGFPYIDFRER
MEHGAAYDSEPFTFEKRAPSPADFIANLNKAERLR
>FLP-34_Pristionchus_mayeri_PRIMAYER000022629.t1 peptide:
PRIMAYER000022629.t1
MIRHICAVLAFALVAEVVLGAPFEKKADLGEFASALNGAGRLRYGKRSSGGFPYIDFRDR
LEQGAYADTEPFSFEKRAPSPADFVANLNKAERLR
>FLP-34_Pristionchus_pacificus_PPA29058.1 peptide: PPA29058.1
MISGNPFPPILPLIPSKMVRCVSAVVLLVLVLEAVVGLSTMQLVILCGIVALSCLIEADD
FAPFEKKADLGEFASALNGAGRLRYGKRSPVAAFPYIDFRERMEHGASYDSEPFTFEKRA
PSAADFVANLNKAERLRFGRK
>FLP-34_Teladorsagia_circumcincta_TELCIR_01653 peptide: TELCIR_01653
MRSLFCYFIALLTVFYNILPVSSLDKKSDMSEFASAINGASRLRYGKRSFDDGYQLKLPL
LCIAEVVTHQVNVNKLILSGLCITIRLHKLLFKCDAIGGLELIALDNQNI
>FLP-34_Ancylostoma_duodenale FLP-34 predicted
TYFSVLSFEKKSDMSEFTSAINGASRLRYGKRSFNFGYPLK
>FLP-34_Cylicostephanus_goldi FLP-34 predicted
YFSVLSYEKKSDMSEFASAINGASRLRYGKRSYFGYPLK
>FLP-34_Caenorhabditis_tropicalis FLP-34 predicted
IILGHQNTFSVLALPLEKKADISTFASAINNAGRLRYGKRSDPAMWEDK
>FLP-34_Haemonchus_placei FLP-34 predicted
HFSSIFKCKHNFFSEKYHHFSVSSVDKKSDLSDFASAINSAGRLRYGKRSSDYGYPLR
>FLP-34_Pristionchus_entomophagus FLP-34 predicted
FDLRLPGIGDGCIVKDCGVPLENIVRLNNSIIKEKAHSMECPDRILPSSHMHLFRGASFEKKADMGEF
ASALNGAGRLRYGKRSSGGFPFDGPLRYLYGSSRHN
>FLP-34_Strongylus_vulgaris FLP-34 predicted
FFSVLSFDKKSDMSEFASAINGASRLRYGKRSFDFGYPLK
>FLP-34_Caenorhabditis_elegans FLP-34 isoform a
MQFQFLMALIFVALVLTDSVLSLPLEKKADISTFASAINNAGRLRYGKRSDPAMWEENNVIIPSSEDQ
YLYSEGRYPYAL
IKRALNRDSLVASLNNAERLRFGRK
>FLP-34_Bursaphelenchus_xylophilus_BXY_0837900.1 peptide:
BXY_0837900.1
MELKMTNSQALCLVMASLIVLTSTALPLLEDRDFSMEKKSNLNEFMSAMKGAPRLRYGKR
NFDEFVPAYQYVIPQRAVYKRMAAQNLPELIDTLNGAERLRFGRK

>FLP-34_Panagrellus_redivivus_Pan_g22127.t1 peptide: Pan_g22127.t1
MKQTAVFSVAAAAVLLLVIVEPALAQFYFDSYELPEKKSSAALSDFTSALNGAARLRYGK
RSAPSVYSIDPQLRDVIWLQAANKRAPSADKTMPQFLESLNGAERLRFG
>FLP-34_Rhabditophanes.sp.KR3021_RSKR_0000509900.1 peptide:
RSKR_0000509900.1
MTAILHRPCNTLQRKGMYLGKLIIATIILIQAVPSFQGPNLPIKKSPFSFDDLSNQLNSA
SRLRYGKRSGGADLHMLNHVLKLNELGKRSAEGYSDIPEYDSPNKRSSPMVGDAAFFKQL
GKRSSSPMLLSDIEYYEKMGKRSPYLADIASFEQIGKRSGSQLRDLAHFEEVGKRSGKNY
LGDLATFDNIGKKSVPMVNDIAYFGDVGKRSSPLVKDIAYFGDVGKRSVPMVQDIAYLGD
VGKRSAAIPLVHDIVYYDQIGKRSPSHSFFRDAAIYGHAGKRLNNLLQNVAEFQDIGKKS
DRGMLLGDLAHFDNLGKRSSSGDLKNVQYYLDAGKKRSSNWGQLFQDLNDAERLR
>FLP-34_Steinernema_carpocapsae_L596_g21266.t2 peptide:
L596_g21266.t2
MVVLSRFVLIASVVVGLTAIFGTAVPLHEKKSTHDLNEFTMALNGASRLRYGKRSDGSVD
PAALYEQFLAAQQFPYEVQEFPAYTQKRSSIYSDPLALKLVQSLNGAERLRFGRR
>FLP-34_Steinernema_feltiae_L889_g33432.t1 peptide: L889_g33432.t1
MVTVTVILRQIGFALLLAVNLVFARFIDLESSAVPLHEKKSTHDLNEFTMALNGASRLRY
GKRSGGLYDSMVYERSPYDFQDVPSYMGQQKRNAIYSDPLALKLVQSLNGAERLRFGRR
>FLP-34_Steinernema_glaseri_L893_g13884.t1 peptide: L893_g13884.t1
MVSVTVVLRHLALAFFVAVNFAFARFIDFDLDSAALPLHEKKSTHDLNEFAMALNGASRL
RYGKRSFGLYEQLLPQQFDSRQEVPEFTGQKRNIYSDPLALKLVQSLNGAERLRFGRR
>FLP-34_Steinernema_monticolum_L898_g22950.t1 peptide:
L898_g22950.t1
MSRQLFMVCVVFCSIAMPSSAVPLHEKKSTHDLNEFTMALNGASRLRYGKRSEGLNDPNV
YEQFLAQQSPYEFQDVPNYMEPQKRGSAIYSDPLALKLVQSLNGAERLRFGRR
>FLP-34_Steinernema_scapterisci_L892_g23323.t1 peptide:
L892_g23323.t1
MVVLSRFVLLACAIVGLAAILGKSIKIEVTQDDVSTVLDRLKVYTQGLPDFFPSACAFFP
VSNAAHLVFTLATTLGILFHNPGLHNSTSAEPFFQELPFKVPKQSTHFKNMLLVTQSGGN
PGLQQTPLNPQDRRLACKQSSGCPFLSLRGFLSGAGIVKTALMNGNQIKCHESPQMTSSM
QTLLLGTRHKVQPESRPGLNAASGRGCRADFAQSQDGNCPDGTLLIYGWLLLKAGFGDKA
IEKTEGIILRHLLLALFVAVNFAFARFIELDLERNAVPLHEKKSTHDLNEFTMALNGASR
LRYGKRSGSSVDPALYEQFLAAQQFPYDVQDFPAYNQKRSSIYSDPLALKLVQSLNGAER
LRFGRR
>FLP-34_Strongyloides_papillosus_SPAL_0000243300.1 peptide:
SPAL_0000243300.1
MTAIKKSPLDFAAISSQLGSAERMRYGKRNGFDVNMLPHVYKLTEIGKRNANEDVDKRSS
KYMMELGEYQNLGKRSDNFIPEMAKFNEVGKRSLPGYLQEVNEFNKVGKRSNAQFDGDLL
QKYHELGKRYRGNVKEITHFFNLGKRSTLSEINSYNSVGKRSNDFLPHIAEFEKFGKRDS
PYLNEIGKFYELGRRASYNQILPIYEDVGKRSVNPYYPGSETYYYDNVGKRLSSIIKNVA
DYNEVGKRSTGLIPNLALYDDFGKRNDDYENLNYYFQHGKRSPNLNAIVQHLNDVDRLRF
GK
>FLP-34_Strongyloides_ratti_SRAE_1000102000.1 peptide:
SRAE_1000102000.1
MTINYCDIRKICRLIFILSLITVNTNGINNDNYFTAIKKSPLDFAAITSQLGSAERMRYG
KRNGFDVTMLPHVYKLTEVGKRNDNDNINKRSSKYIMELGEYSNLGKRSDNFLPELTKFN
EVGKKSNYYDNLLKKYNPLTKRYRGDVRDITHLYNLGKRPMLSEIGEYENVGKRNSEHIP
HIIEFEKFGKRNSPYLNEIEKFHELGKRTSYNQVIPIYESAGKRSYYPFYDSDTYYYDSV
GKRLSSIIKNVAEYNEVGKRSNSLIPNLVHYNDFGKRNDDYENLNYYFQHGKRSPNLNAI
IQHLNDADRLRFGK
>FLP-34_Strongyloides_stercoralis_SSTP_0000993900.1 peptide:
SSTP_0000993900.1
MTTNYWDLKKISKLIFILSLITGNTNGISNDNYMTAIKKSPLDFAAITSQLGSAERMRYG
KRSMLSEIGEYDNVGKRSNDYIPQIVEFDKFGKRSSYNQVLPIYDNAGKRSYYPFYDSDT

YYYDSIGKRLSSIIKNVAEYNQVGKRSNGLVPHLIHYDDFGKRTDDFDNLNYYYQHGKRS
PNLNAIIQHLNDVDRLRFGK
>FLP-34_Strongyloides_venezuelensis_SVE_0000900.1 peptide:
SVE_0000900.1
MTIINCDVRKISTLIFVLSLVSGSASGFNSDNYITAIKKSPLDFAAISSQLGSAERMRYG
KRNGFDVNMLPHVYKLTEIGKRDVNEDVDKRSSKYIMELGEYQNLGKRSDNFIPEMAKFN
EVGKKSLPGYLQEVNEFNKVGKRNNAQFDGDLLQKYHELGKRSTLSEINDYNSVGKRSND
FLPHIAEFEKFGKRDSSYLREIGKFRELGRRASYNQILPIYENVGKRSVYPFYPGGEAYY
YDSVGKRLSSIIKNVADYNEVGKRSTGLIPNLAHYDDFGKRSDDYENLNYYFQHGKRSPN
LNAIVQHLNDVDRLRFGK
>FLP-34_Halicephalobus_mephisto FLP-34 predicted
KIYFLFFVFCFLKLYDFSYDQIPEKKSHEGLTDFTSALNGASRLRYGKRGYFNLDPEAQEWNQK
KYCRFQEYSKRASSVRDVMPQFVQNLNGAERLRFG
>FLP-34_Acrobeloides_nanus_ACRNAN_scaffold11262.g10622.t1 peptide:
ACRNAN_scaffold11262.g10622.t1
MANTVIFYIFALILAVNCALARFLPDDESFEAPEKKSYDSLTDFTSALNGAARLRYGKRF
WNPYGYPSWSSDHSKRSQFAANLPQFIQNLNGAERLRFGRK
>FLP-34_Ditylenchus_destructor_Dd_09459 peptide: Dd_09459
MMKFTKTFQPSHSCSACVVLLASLAVLVHESLAQPSPHMTDLNRYGPEKKSADLGADFMS
AMNGATRLRYGKRSNGLFYNDLLAAAGPVEVDGQAPFAFRTPSRWATAKRMASFSDNIPA
LVDQLNGAERLRFGRK
>FLP-34_Globodera_pallida_GPLIN_000848700 peptide: GPLIN_000848700
MTQAQSGGNRRMLGEEYLHRLNKRSPMKKSDGLSDFVGSLNGAARLRYGKRSESEGSPAE
EMEDISPQFFWAVRSSNGRWKLAKRASNYAEALPAGLLDQLNGAERLR
>FLP-34_Globodera_rostochiensis_GROS_g08013.t1 peptide:
GROS_g08013.t1
MSLSSANFFPLTTFALFVLLLALHGMTQAQIDGNRRMVVEEYLHRLNKRSPMKKSDGLSD
FVGSLNGAARLRYGKRKALPAGLLDQLNGAERLRFGRK
>FLP-34_Heterodera_glycines_Hetgly.G000024071 peptide:
Hetgly.G000024071
MAQYPSITNVCVKELFVLAIRPFEFRFFPFPWPLPMASHLSVTVLTSIIALFVLLLALHG
TAPNQLKIGENGLFEEYLRQLDKRSPLKKSDGLSDFVGSLNGAARLRYGKRSDGLTAEEI
EQILSLNRWKMPKRVANYAEALPIGLLDQLNGAERLRFGRK
>FLP-34_scaffold4682_cov54.g8296 peptide: scaffold4682_cov54.g8296
MQQTGPFLSICILSFLAIFLSYANSQELGSVAVNDPLLLEEIMLQQMPKRGSMKKADVRD
FIGSINSASRLRYGKRAILEPVIIASPYK
>FLP-34_Meloidogyne_enterolobii_scaffold62222_cov323.g30035 peptide:
scaffold62222_cov323.g30035
MQQTGSFLSICILSFLAIFLSYANSQELGSVAVNEPLLLEEFMLQQMPKRGAMKKADVRD
FIGSINSASRLRYGKRAILEPVIIASPY
>FLP-34_Meloidogyne_floridensis_scf7180000424733.g13855 peptide:
scf7180000424733.g13855
MQQTGPFLSICILSFLAVFLSYANSQELGSVAVNDPLLLEEIMLQQMPKRGSMKKADVRD
FIGSINSASRLRYGKRAILEPVIIASPYK
>FLP-34_Meloidogyne_hapla_MhA1_Contig1860.frz3.gene11 peptide:
MhA1_Contig1860.frz3.gene11
MSYANSQELGSVAINDPILLEEIMLQQMPKRGPMKKADVRDFIGSLNGASRLRYGKRAVL
EPMLIASPYKLEMLAKRASNNYAQALPAGLLDQLNGAERLRFGRK
>FLP-34_Meloidogyne_incognita_Minc3s00058g02988 peptide:
Minc3s00058g02988
MQQTGPFLSICILSFLAIFLSYANSQELGSVAVNDPLLLEEIMLQQMPKRGSMKKADVRD
FIGSINSASRLRYGKRAILEPVIIASPYKLQQMLAKRAPSSYAEALPAGLLDQLNGAERL
RFGRK
>FLP-34_Meloidogyne_javanica_scaffold33649_cov252.g20977 peptide:

```
scaffold33649_cov252.g20977
MQQTGSSLICFLSFLAVFLSYANSQELGSVAVNDPLLLEEIMLQQMPKRGSMKKADVRDF
IGSINSASRLRYGKRAILEPVIIASPY
>FLP-34_Ditylenchus_dipsaci FLP-34 predicted
PSYYSLNLNYYVADLRQRRIVRCRLSSDFMSAMNGATRLRYGKRSGVNYNDLRAASHSKIVSPTTSIR
FKNAQVSLANLL
```
