## Supplementary File 2 for "Exploitation of phylum-spanning omics resources reveals complexity in the nematode FLP signalling system and provides insights into *flp*-gene evolution"

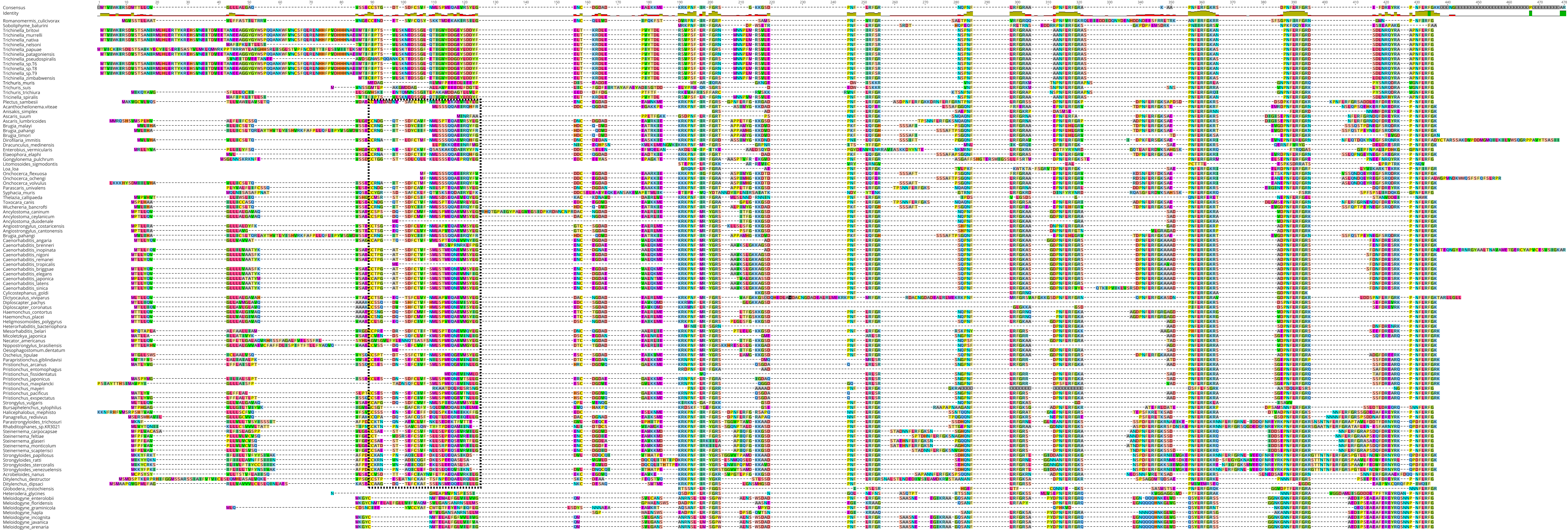

Consensus
Identity
Romanomermis_culicivorax
Soboliphyme_baturini
Trichinella_britovi
Trichinella_murrelli
Trichinella_nativa
Trichinella_nelsoni
Trichinella_papuae
Trichinella_patagoniensis
Trichinella_pseudospiralis
Trichinella_sp.T6
Trichinella_sp.T8
Trichinella_sp.T9
Trichinella_zimbabwensis
Trichuris_muris
Trichuris_suis
Trichuris_trichiura
Tricinella_spiralis
Plectus_sambesii
Acanthocheilonema.viteae
Anisakis_simplex
Ascaris_suum
Ascaris_lumbricoides
Brugia_malayi
Brugia_pahangi
Brugia_timori
Dirofilaria_immitis
Dracunculus_medinensis
Enterobius_vermicularis
Elaeophora_elaphi
Gongylonema_pulchrum
Litomosoides_sigmodontis
Loa_loa
Onchocerca_flexuosa
Onchocerca_ochengi
Onchocerca_volvulus
Parascaris_univalens
Syphacia_muris
Thelazia_callipaeda
Toxocara_canis
Wuchereria_bancrofti
Ancylostoma_caninum
Ancylostoma_ceylanicum
Ancylostoma_duodenale
Angiostrongylus_costaricensis
Angiostrongylus_cantonensis
Brugia_pahangi
Caenorhabditis_angaria
Caenorhabditis_brenneri
Caenorhabditis_inopinata
Caenorhabditis_nigoni
Caenorhabditis_remanei
Caenorhabditis_tropicalis
Caenorhabditis_briggsae
Caenorhabditis_elegans
Caenorhabditis_japonica
Caenorhabditis_latens
Caenorhabditis_sinica
Cylicostephanus_goldi
Dictyocaulus_viviparus
Diploscapter_pachys
Diploscapter_coronatus
Haemonchus_contortus
Haemonchus_placei
Heligmosomoides_polygyrus
Heterorhabditis_bacteriophora
Mesorhabditis_belari
Micoletzkya_japonica
Necator_americanus
Nippostrongylus_brasiliensis
Oesophagostomum.dentatum
Oscheius_tipulae
Parapristionchus.giblindavisi
Pristionchus_arcanus
Pristionchus_entomophagus
Pristionchus_fissidentatus
Pristionchus_japonicus
Pristionchus_maxplancki
Pristionchus_mayeri
Pristionchus_pacificus
Pristionchus_exspectatus
Strongylus_vulgaris
Bursaphelenchus_xylophilus
Halicephalobus_mephisto
Panagrellus_redivivus
Parastrongyloides_trichosuri
Rhabditophanes_sp.KR3021
Steinernema_carpocapsae
Steinernema_feltiae
Steinernema_glaseri
Steinernema_monticolum
Steinernema_scapterisci
Strongyloides_papillosus
Strongyloides_ratti
Strongyloides_stercoralis
Strongyloides_venezuelensis
Acrobeloides_nanus
Ditylenchus_destructor
Ditylenchus_dipsaci
Globodera_rostochiensis
Heterodera_glycines
Meloidogyne_enterolobii
Meloidogyne_floridensis
Meloidogyne_graminicola
Meloidogyne_hapla
Meloidogyne_incognita
Meloidogyne_javanica
Meloidogyne_arenaria

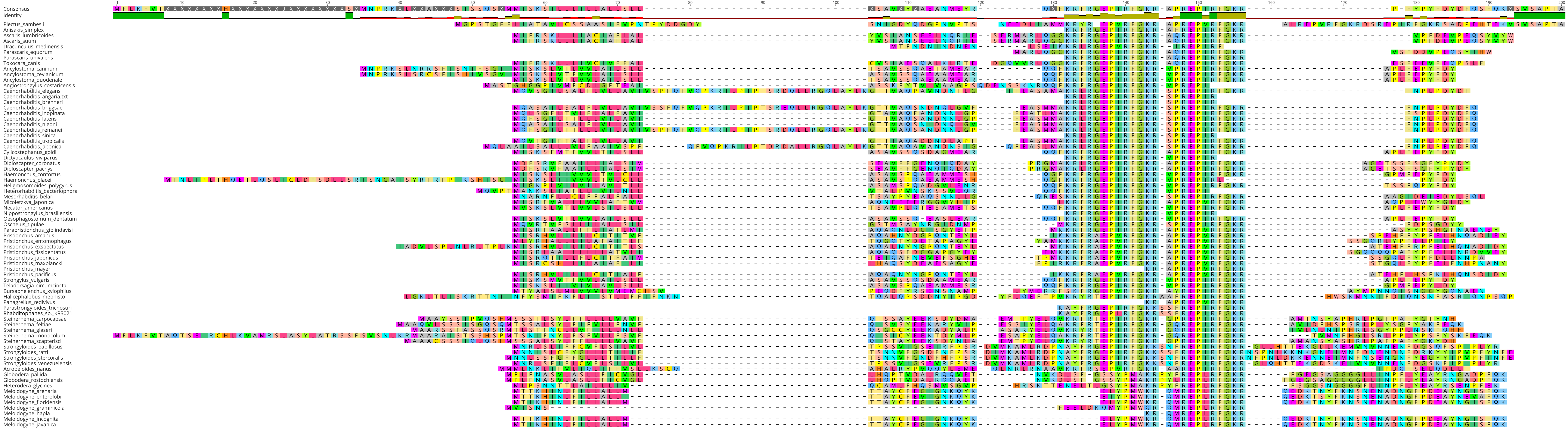
Consensus
Identity
Plectus_sambesii
Anisakis_simplex
Ascaris_lumbricoides
Ascaris_suum
Dracunculus_medinensis
Parascaris_equorum
Parascaris_univalens
Toxocara_canis
Ancylostoma_caninum
Ancylostoma_ceylanicum
Ancylostoma_duodenale
Angiostrongylus_costaricensis
Caenorhabditis_elegans
Caenorhabditis_angaria.txt
Caenorhabditis_brenneri
Caenorhabditis_briggsae
Caenorhabditis_inopinata
Caenorhabditis_latens
Caenorhabditis_nigoni
Caenorhabditis_remanei
Caenorhabditis_sinica
Caenorhabditis_tropicalis
Caenorhabditis_japonica
Cylicostephanus_goldi
Dictyocaulus_viviparus
Diploscapter_coronatus
Diploscapter_pachys
Haemonchus_contortus
Haemonchus_placei
Heligmosomoides_polygyrus
Heterorhabditis_bacteriophora
Mesorhabditis_belari
Micoletzkya_japonica
Necator_americanus
Nippostrongylus_brasiliensis
Oesophagostomum_dentatum
Oscheius_tipulae
Parapristionchus_giblindavisi
Pristionchus_arcanus
Pristionchus_entomophagus
Pristionchus_exspectatus
Pristionchus_fissidentatus
Pristionchus_japonicus
Pristionchus_maxplancki
Pristionchus_mayeri
Pristionchus_pacificus
Strongylus_vulgaris
Teladorsagia_circumcincta
Bursaphelenchus_xylophilus
Halicephalobus_mephisto
Panagrellus_redivivus
Parastrongyloides_trichosuri
Rhabditophanes_sp._KR3021
Steinernema_carpocapsae
Steinernema_feltiae
Steinernema_glaseri
Steinernema_monticolum
Steinernema_scapterisci
Strongyloides_papillosus
Strongyloides_ratti
Strongyloides_stercoralis
Strongyloides_venezuelensis
Acrobeloides_nanus
Globodera_pallida
Globodera_rostochiensis
Heterodera_glycines
Meloidogyne_arenaria
Meloidogyne_enterolobii
Meloidogyne_floridensis
Meloidogyne_graminicola
Meloidogyne_hapla
Meloidogyne_incognita
Meloidogyne_javanica

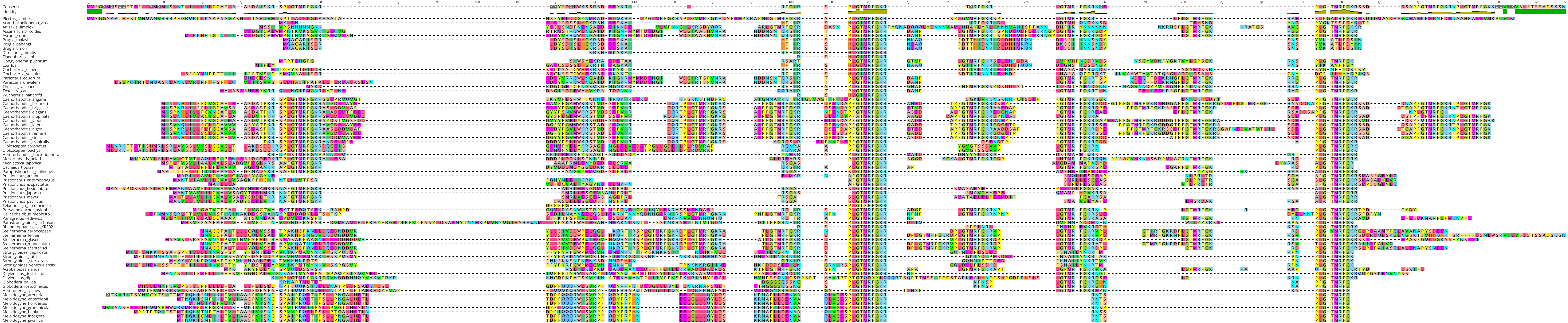
Consensus
Identity
Plectus_sambesii
Acanthocheilonema_viteae
Anisakis_simplex
Ascaris_lumbricoides
Ascaris_suum
Brugia_malayi
Brugia_pahangi
Brugia_timori
Dirofilaria_immitis
Elaeophora_elaphi
Gongylonema_pulchrum
Loa_loa
Onchocerca_ochengi
Onchocerca_volvulus
Parascaris_equorum
Parascaris_univalens
Thelazia_callipaeda
Toxocara_canis
Wuchereria_bancrofti
Caenorhabditis_angaria
Caenorhabditis_brenneri
Caenorhabditis_briggsae
Caenorhabditis_elegans
Caenorhabditis_inopinata
Caenorhabditis_japonica
Caenorhabditis_latens
Caenorhabditis_nigoni
Caenorhabditis_remanei
Caenorhabditis_sinica
Caenorhabditis_tropicalis
Diploscapter_coronatus
Diploscapter_pachys
Heterorhabditis_bacteriophora
Mesorhabditis_belari
Micoletzkya_japonica
Oscheius_tipulae
Parapristionchus_giblindavisi
Pristionchus_arcanus
Pristionchus_entomophagus
Pristionchus_exspectatus
Pristionchus_fissidentatus
Pristionchus_japonicus
Pristionchus_mayeri
Pristionchus_pacificus
Teladorsagia_circumcincta
Bursaphelenchus_xylophilus
Halicephalobus_mephisto
Panagrellus_redivivus
Parastrongyloides_trichosuri
Rhabditophanes_sp._KR3021
Steinernema_carpocapsae
Steinernema_feltiae
Steinernema_glaseri
Steinernema_monticolum
Steinernema_scapterisci
Strongyloides_papillosus
Strongyloides_ratti
Strongyloides_stercoralis
Strongyloides_venezuelensis
Acrobeloides_nanus
Ditylenchus_destructor
Ditylenchus_dipsaci
Globodera_pallida
Globodera_rostochiensis
Heterodera_glycines
Meloidogyne_arenaria
Meloidogyne_enterolobii
Meloidogyne_floridensis
Meloidogyne_graminicola
Meloidogyne_hapla
Meloidogyne_incognita
Meloidogyne_javanica

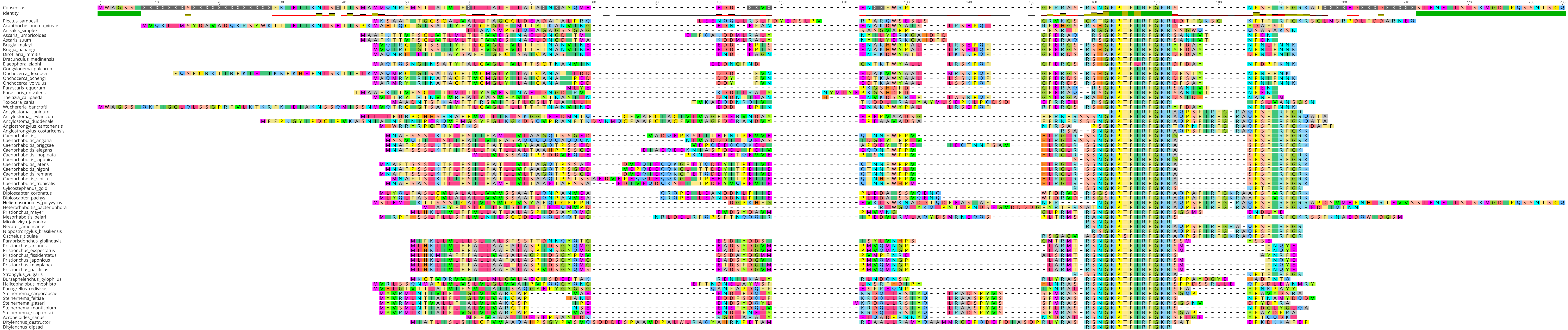

Consensus
Identity
Plectus_sambesii
Acanthocheilonema_viteae
Anisakis_simplex
Ascaris_lumbricoides
Ascaris_suum
Brugia_malayi
Brugia_pahangi
Dirofilaria_immitis
Dracunculus_medinensis
Elaeophora_elaphi
Gongylonema_pulchrum
Onchocerca_flexuosa
Onchocerca_ochengi
Onchocerca_volvulus
Parascaris_equorum
Parascaris_univalens
Thelazia_callipaeda
Toxocara_canis
Wuchereria_bancrofti
Ancylostoma_caninum
Ancylostoma_ceylanicum
Ancylostoma_duodenale
Angiostrongylus_cantonensis
Angiostrongylus_costaricensis
Caenorhabditis_
Caenorhabditis_angaria
Caenorhabditis_briggsae
Caenorhabditis_elegans
Caenorhabditis_inopinata
Caenorhabditis_japonica
Caenorhabditis_latens
Caenorhabditis_nigoni
Caenorhabditis_remanei
Caenorhabditis_sinica
Caenorhabditis_tropicalis
Cylicostephanus_goldi
Diploscapter_coronatus
Diploscapter_pachys
Heligmosomoides_polygyrus
Heterorhabditis_bacteriophora
Pristionchus_mayeri
Mesorhabditis_belari
Micoletzkya_japonica
Necator_americanus
Nippostrongylus_brasiliensis
Oscheius_tipulae
Parapristionchus_giblindavisi
Pristionchus_arcanus
Pristionchus_expectatus
Pristionchus_fissidentatus
Pristionchus_japonicus
Pristionchus_maxplancki
Pristionchus_pacificus
Strongylus_vulgaris
Bursaphelenchus_xylophilus
Halicephalobus_mephisto
Panagrellus_redivivus
Steinernema_carpocapsae
Steinernema_feltiae
Steinernema_glaseri
Steinernema_monticolum
Steinernema_scapterisci
Acrobeloides_nanus
Ditylenchus_destructor
Ditylenchus_dipsaci

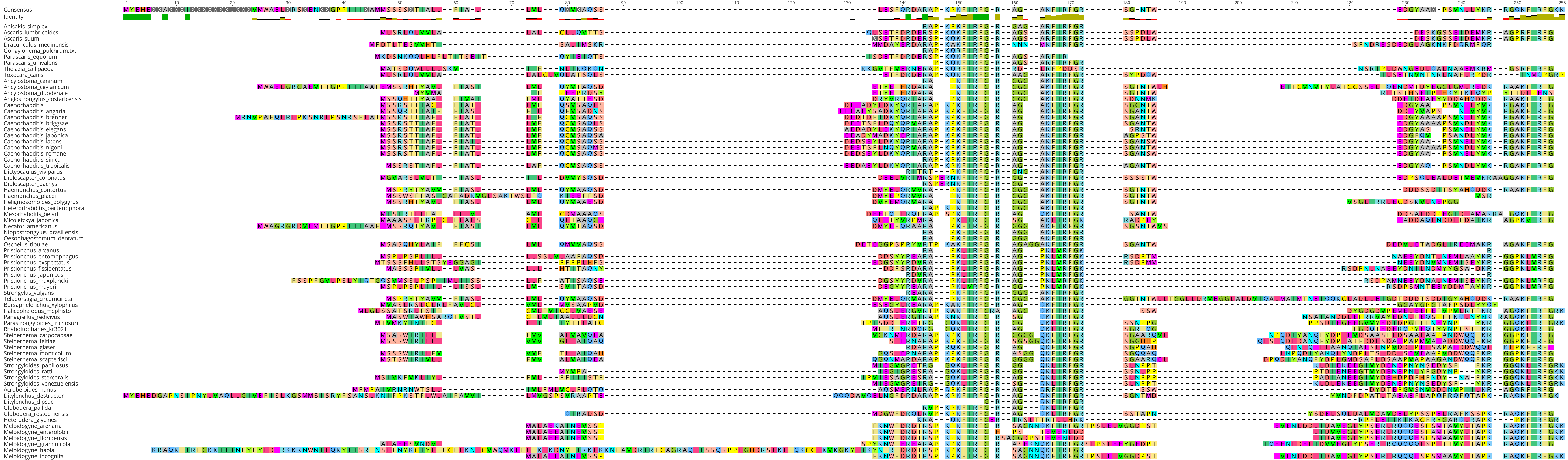

Consensus
Identity
Anisakis_simplex
Ascaris_lumbricoides
Ascaris_suum
Dracunculus_medinensis
Gongylonema_pulchrum.txt
Parascaris_equorum
Parascaris_univalens
Thelazia_callipaeda
Toxocara_canis
Ancylostoma_caninum
Ancylostoma_ceylanicum
Ancylostoma_duodenale
Angiostrongylus_costaricensis
Caenorhabditis
Caenorhabditis_angaria
Caenorhabditis_brenneri
Caenorhabditis_briggsae
Caenorhabditis_elegans
Caenorhabditis_japonica
Caenorhabditis_latens
Caenorhabditis_nigoni
Caenorhabditis_remanei
Caenorhabditis_sinica
Caenorhabditis_tropicalis
Dictyocaulus_viviparus
Diploscapter_coronatus
Diploscapter_pachys
Haemonchus_contortus
Haemonchus_placei
Heligmosomoides_polygyrus
Heterorhabditis_bacteriophora
Mesorhabditis_belari
Nicoletzkya_japonica
Necator_americanus
Nippostrongylus_brasiliensis
Oesophagostomum_dentatum
Oscheius_tipulae
Pristionchus_arcanus
Pristionchus_entomophagus
Pristionchus_exspectatus
Pristionchus_fissidentatus
Pristionchus_japonicus
Pristionchus_maxplancki
Pristionchus_mayeri
Strongylus_vulgaris
Teladorsagia_circumcincta
Bursaphelenchus_xylophilus
Halicephalobus_mephisto
Panagrellus_redivivus
Parastrongyloides_trichosuri
Rhabditophanes_kr3021
Steinernema_carpocapsae
Steinernema_feltiae
Steinernema_glaseri
Steinernema_monticolum
Steinernema_scapterisci
Strongyloides_papillosus
Strongyloides_ratti
Strongyloides_stercoralis
Strongyloides_venezuelensis
Acrobeloides_nanus
Ditylenchus_destructor
Ditylenchus_dipsaci
Globodera_pallida
Globodera_rostochiensis
Heterodera_glycines
Meloidogyne_arenaria
Meloidogyne_enterolobii
Meloidogyne_floridensis
Meloidogyne_graminicola
Meloidogyne_hapla
Meloidogyne_incognita

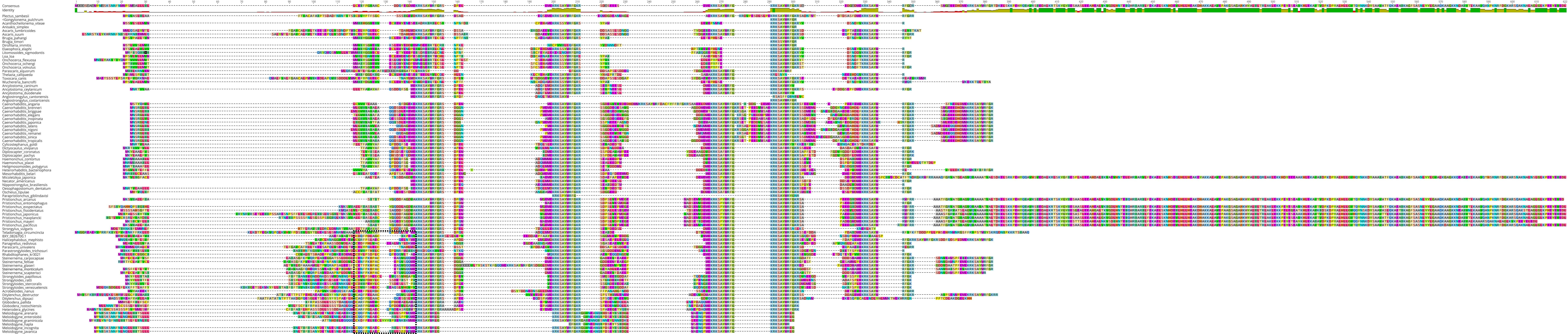
Consensus
Identity
Plectus_sambesii
<Gongylonema_pulchrum
Acanthocheilonema_viteae
Anisakis_simplex
Ascaris_lumbricoides
Ascaris_suum
Brugia_pahangi
Brugia_timori
Dirofilaria_immitis
Elaeophora_elaphi
Litomosoides_sigmodontis
Loa_loa
Onchocerca_flexuosa
Onchocerca_ochengi
Onchocerca_volvulus
Parascaris_equorum
Thelazia_callipaeda
Toxocara_canis
Wuchereria_bancrofti
Ancylostoma_caninum
Ancylostoma_ceylanicum
Ancylostoma_duodenale
Angiostrongylus_cantonensis
Angiostrongylus_costaricensis
Caenorhabditis_angaria
Caenorhabditis_brenneri
Caenorhabditis_briggsae
Caenorhabditis_elegans
Caenorhabditis_inopinata
Caenorhabditis_japonica
Caenorhabditis_latens
Caenorhabditis_nigoni
Caenorhabditis_remanei
Caenorhabditis_sinica
Caenorhabditis_tropicalis
Cylicostephanus_goldi
Dictyocaulus_viviparus
Diploscapter_coronatus
Diploscapter_pachys
Haemonchus_contortus
Haemonchus_placei
Heligmosomoides_polygyrus
Heterorhabditis_bacteriophora
Mesorhabditis_belari
Micoletzkya_japonica
Necator_americanus
Nippostrongylus_brasiliensis
Oesophagostomum_dentatum
Oscheius_tipulae
Parapristionchus_giblindavisI
Pristionchus_arcanus
Pristionchus_entomophagus
Pristionchus_exspectatus
Pristionchus_fissidentatus
Pristionchus_japonicus
Pristionchus_maxplancki
Pristionchus_mayeri
Pristionchus_pacificus
Strongylus_vulgaris
Teladorsagia_circumcincta
BXY_0826700.1
Halicephalobus_mephisto
Panagrellus_redivivus
Parascaris_univalens
Parastrongyloides_trichosuri
Rhabditophanes_kr3021
Steinernema_carpocapsae
Steinernema_feltiae
Steinernema_glaseri
Steinernema_monticolum
Steinernema_scapterisci
Strongyloides_papillosus
Strongyloides_ratti
Strongyloides_stercoralis
Strongyloides_venezuelensis
Acrobeloides_nanus
Ditylenchus_destructor
Ditylenchus_dipsaci
Globodera_pallida
Globodera_rostochiensis
Heterodera_glycines
Meloidogyne_arenaria
Meloidogyne_enterolobii
Meloidogyne_graminicola
Meloidogyne_hapla
Meloidogyne_incognita
Meloidogyne_javanica

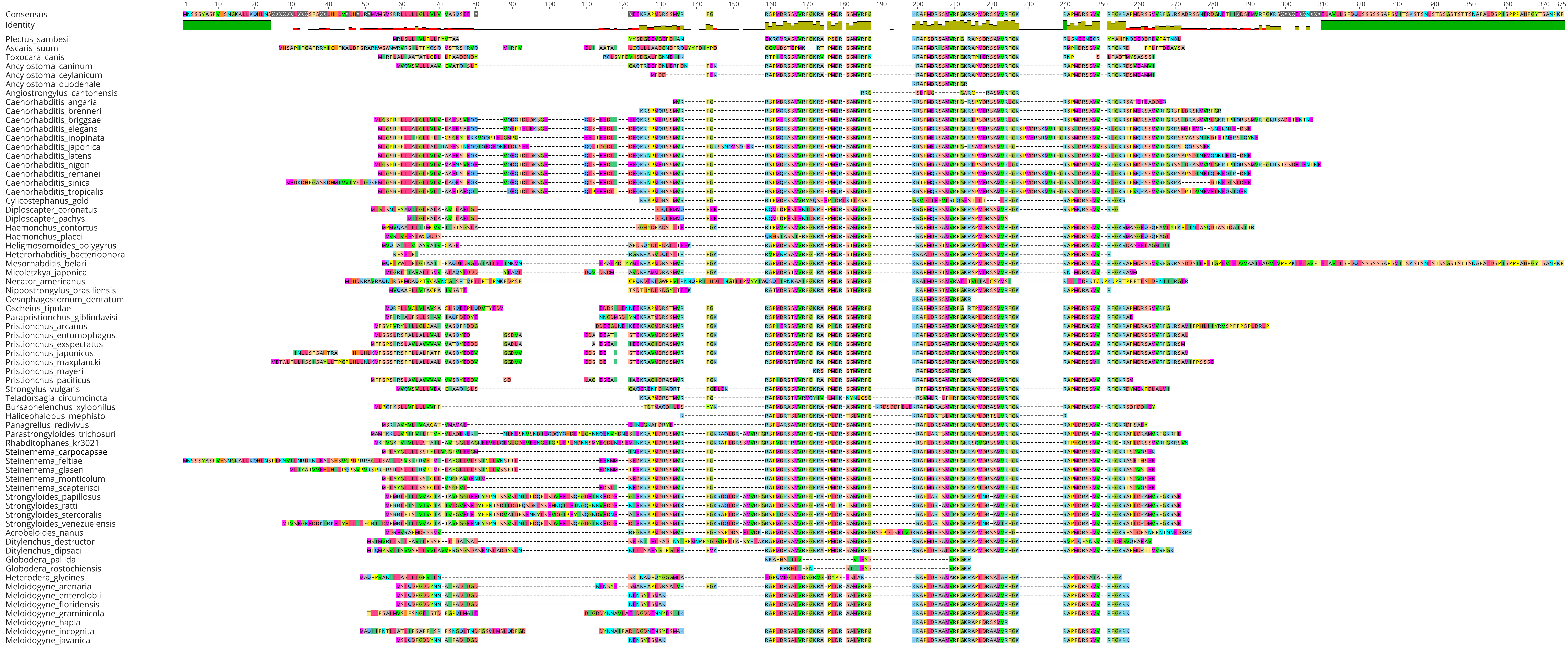
1 10 20 30 40 50 60 70 80 90 100 110 120 130 140 150 160 170 180 190 200 210 220 230 240 250 260 270 280 290 300 310 320 330 340 350 360 370 372
Consensus
Identity
Plectus_sambesii
Ascaris_suum
Toxocara_canis
Ancylostoma_caninum
Ancylostoma_ceylanicum
Ancylostoma_duodenale
Angiostrongylus_cantonensis
Caenorhabditis_angaria
Caenorhabditis_brenneri
Caenorhabditis_briggsae
Caenorhabditis_elegans
Caenorhabditis_inopinata
Caenorhabditis_japonica
Caenorhabditis_latens
Caenorhabditis_nigoni
Caenorhabditis_remanei
Caenorhabditis_sinica
Caenorhabditis_tropicalis
Cylicostephanus_goldi
Diploscapter_coronatus
Diploscapter_pachys
Haemonchus_contortus
Haemonchus_placei
Heligmosomoides_polygyrus
Heterorhabditis_bacteriophora
Mesorhabditis_belari
Micoletzkya_japonica
Necator_americanus
Nippostrongylus_brasiliensis
Oesophagostomum_dentatum
Oscheius_tipulae
Parapristionchus_giblindavisi
Pristionchus_arcanus
Pristionchus_entomophagus
Pristionchus_exspectatus
Pristionchus_japonicus
Pristionchus_maxplancki
Pristionchus_mayeri
Pristionchus_pacificus
Strongylus_vulgaris
Teladorsagia_circumcincta
Bursaphelenchus_xylophilus
Halicephalobus_mephisto
Panagrellus_redivivus
Parastrongyloides_trichosuri
Rhabditophanes_kr3021
Steinernema_carpocapsae
Steinernema_feltiae
Steinernema_glaseri
Steinernema_monticolum
Steinernema_scapterisci
Strongyloides_papillosus
Strongyloides_ratti
Strongyloides_stercoralis
Strongyloides_venezuelensis
Acrobeloides_nanus
Ditylenchus_destructor
Ditylenchus_dipsaci
Globodera_pallida
Globodera_rostochiensis
Heterodera_glycines
Meloidogyne_arenaria
Meloidogyne_enterolobii
Meloidogyne_floridensis
Meloidogyne_graminicola
Meloidogyne_hapla
Meloidogyne_incognita
Meloidogyne_javanica

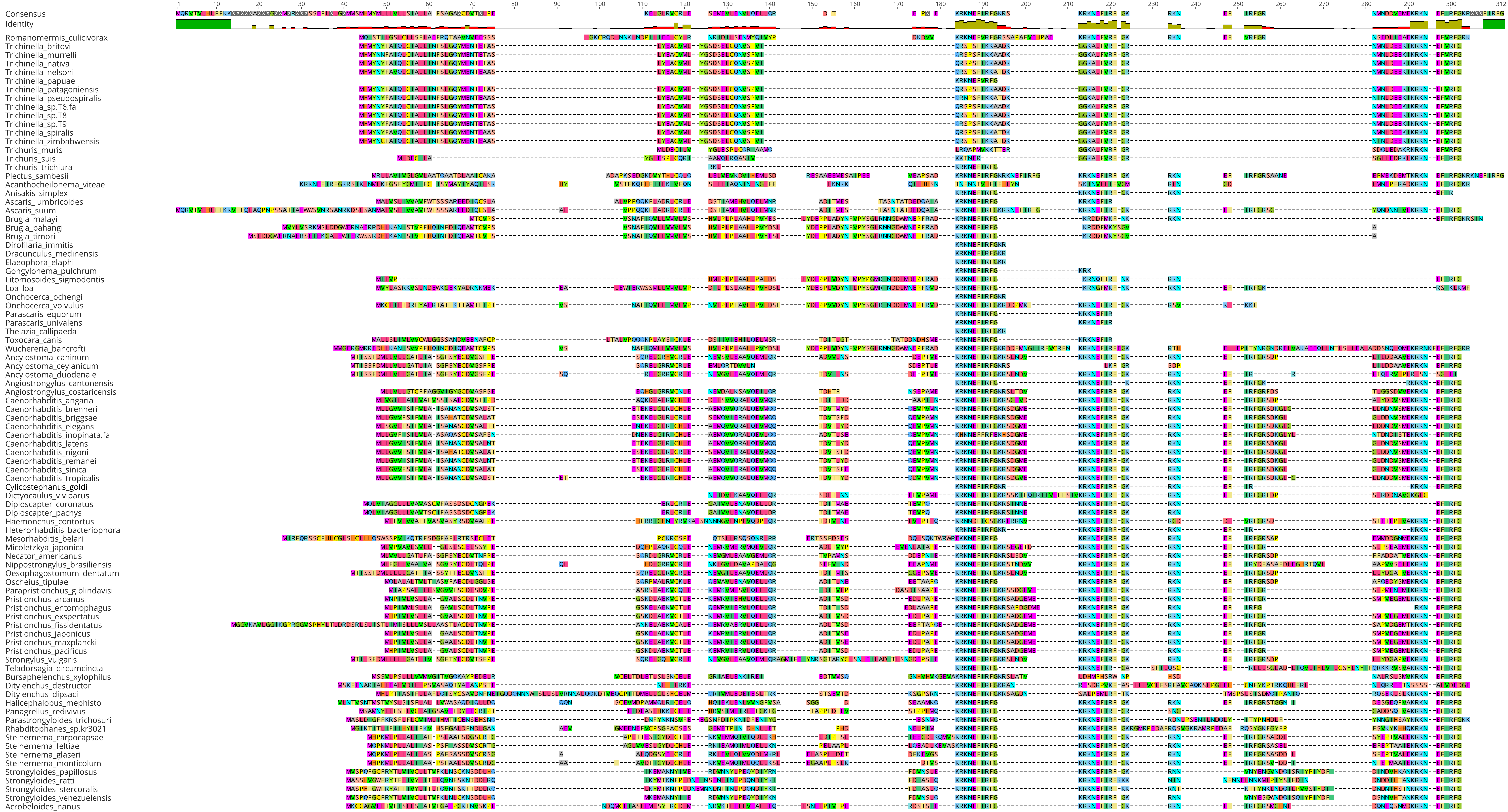
Consensus
Identity
Romanomermis_culicivorax
Trichinella_britovi
Trichinella_murrelli
Trichinella_nativa
Trichinella_nelsoni
Trichinella_papuae
Trichinella_patagoniensis
Trichinella_pseudospiralis
Trichinella_sp.T6.fa
Trichinella_sp.T8
Trichinella_sp.T9
Trichinella_spiralis
Trichinella_zimbabwensis
Trichuris_muris
Trichuris_suis
Trichuris_trichiura
Plectus_sambesii
Acanthocheilonema_viteae
Anisakis_simplex
Ascaris_lumbricoides
Ascaris_suum
Brugia_malayi
Brugia_pahangi
Brugia_timori
Dirofilaria_immitis
Dracunculus_medinensis
Elaeophora_elaphi
Gongylonema_pulchrum
Litomosoides_sigmodontis
Loa_loa
Onchocerca_ochengi
Onchocerca_volvulus
Parascaris_equorum
Parascaris_univalens
Thelazia_callipaeda
Toxocara_canis
Wuchereria_bancrofti
Ancylostoma_caninum
Ancylostoma_ceylanicum
Ancylostoma_duodenale
Angiostrongylus_cantonensis
Angiostrongylus_costaricensis
Caenorhabditis_angaria
Caenorhabditis_brenneri
Caenorhabditis_briggsae
Caenorhabditis_elegans
Caenorhabditis_inopinata.fa
Caenorhabditis_latens
Caenorhabditis_nigoni
Caenorhabditis_remanei
Caenorhabditis_sinica
Caenorhabditis_tropicalis
Cylicostephanus_goldi
Dictyocaulus_viviparus
Diploscapter_coronatus
Diploscapter_pachys
Haemonchus_contortus
Heterorhabditis_bacteriophora
Mesorhabditis_belari
Micoletzkya_japonica
Necator_americanus
Nippostrongylus_brasiliensis
Oesophagostomum_dentatum
Oscheius_tipulae
Parapristionchus_giblindavisi
Pristionchus_arcanus
Pristionchus_entomophagus
Pristionchus_exspectatus
Pristionchus_fissidentatus
Pristionchus_japonicus
Pristionchus_maxplancki
Pristionchus_pacificus
Strongylus_vulgaris
Teladorsagia_circumcincta
Bursaphelenchus_xylophilus
Ditylenchus_destructor
Ditylenchus_dipsaci
Halicephalobus_mephisto
Panagrellus_redivivus
Parastrongyloides_trichosuri
Rhabditophanes_sp.kr3021
Steinernema_carpocapsae
Steinernema_feltiae
Steinernema_glaseri
Steinernema_monticolum
Strongyloides_papillosus
Strongyloides_ratti
Strongyloides_stercoralis
Strongyloides_venezuelensis
Acrobeloides_nanus

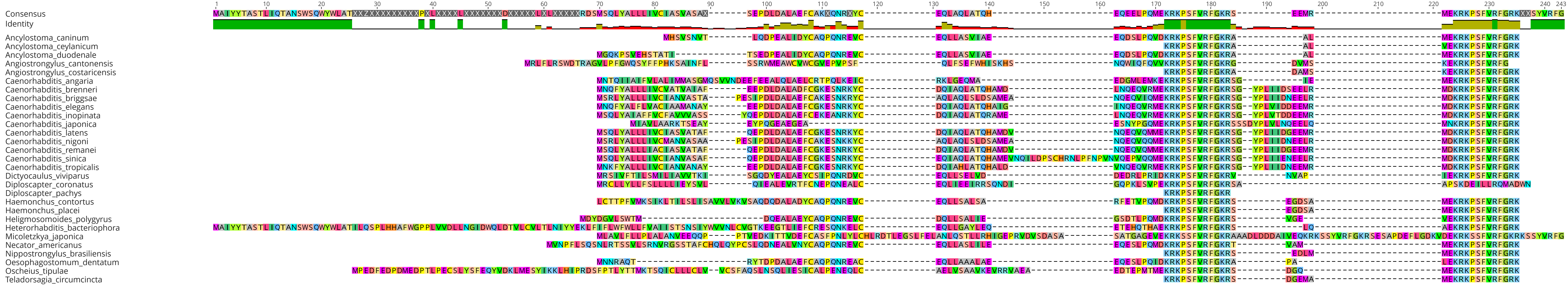
Consensus
Identity
Ancylostoma_caninum
Ancylostoma_ceylanicum
Ancylostoma_duodenale
Angiostrongylus_cantonensis
Angiostrongylus_costaricensis
Caenorhabditis_angaria
Caenorhabditis_brenneri
Caenorhabditis_briggsae
Caenorhabditis_elegans
Caenorhabditis_inopinata
Caenorhabditis_japonica
Caenorhabditis_latens
Caenorhabditis_nigoni
Caenorhabditis_remanei
Caenorhabditis_sinica
Caenorhabditis_tropicalis
Dictyocaulus_viviparus
Diploscapter_coronatus
Diploscapter_pachys
Haemonchus_contortus
Haemonchus_placei
Heligmosomoides_polygyrus
Heterorhabditis_bacteriophora
Micoletzkya_japonica
Necator_americanus
Nippostrongylus_brasiliensis
Oesophagostomum_dentatum
Oscheius_tipulae
Teladorsagia_circumcincta

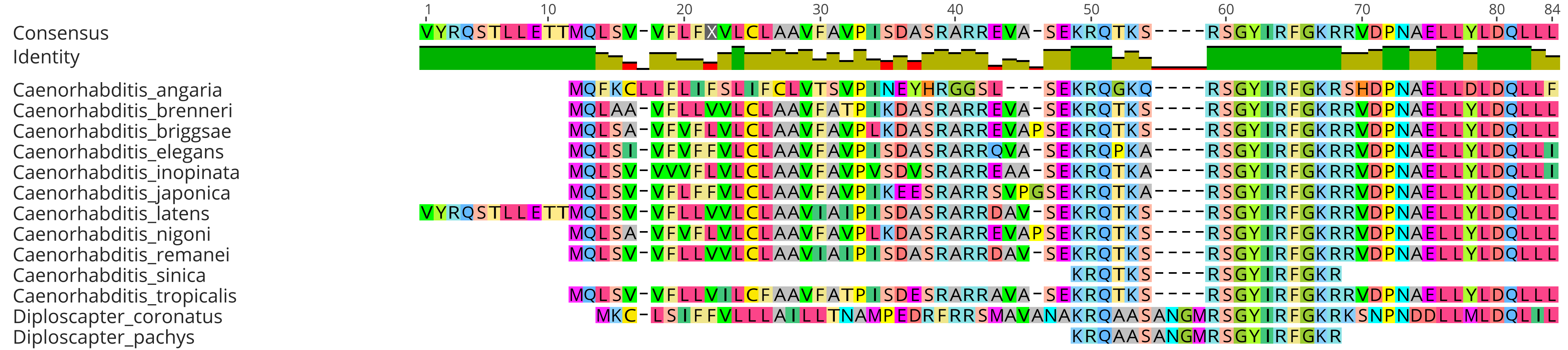

1 10 20 30 40 50 60 70 80 84
Consensus VYRQSTLLETTMQLSV-VFLFXVLCLAAVFAVPISDASRARREVA-SEKRQTKS----RSGYIRFGKRRVDPNAELLYLDQLLL
Identity
Caenorhabditis_angaria MQFKCLLFLIFSLIFCLVTSVPINEYHRGGSL---SEKRQGKQ----RSGYIRFGKRSHDPNAELLDLDQLLF
Caenorhabditis_brenneri MQLAA-VFLLVVLCLAAVFATPIKDASRARREVA-SEKRQTKS----RSGYIRFGKRRVDPNAELLYLDQLLL
Caenorhabditis_briggsae MQLSA-VFVFLVLCLAAVFAVPLKDASRARREVAPSEKRQTKS----RSGYIRFGKRRVDPNAELLYLDQLLL
Caenorhabditis_elegans MQLSI-VFVFFVLCLAAVFAVPISDASRARRQVA-SEKRQPKA----RSGYIRFGKRRVDPNAELLYLDQLLI
Caenorhabditis_inopinata MQLSV-VVVFLVLCLAAVFAVPVSDVSRARREAA-SEKRQTKA----RSGYIRFGKRRVDPNAELLYLDQLLI
Caenorhabditis_japonica MQLSV-VFLFFVLCLAAVFAVPIKEESRARRSVPGSEKRQTKA----RSGYIRFGKRRVDPNAELLYLDQLLL
Caenorhabditis_latens VYRQSTLLETTMQLSV-VFLLVVLCLAAVIAIPISDASRARRDAV-SEKRQTKS----RSGYIRFGKRRVDPNAELLYLDQLLL
Caenorhabditis_nigoni MQLSA-VFVFLVLCLAAVFAVPLKDASRARREVAPSEKRQTKS----RSGYIRFGKRRVDPNAELLYLDQLLL
Caenorhabditis_remanei MQLSV-VFLLVVLCLAAVIAIPISDASRARRDAV-SEKRQTKS----RSGYIRFGKRRVDPNAELLYLDQLLL
Caenorhabditis_sinica KRQTKS----RSGYIRFGKR
Caenorhabditis_tropicalis MQLSV-VFLLVILCFAAVFATPISDESRARRAVA-SEKRQTKS----RSGYIRFGKRRVDPNAELLYLDQLLL
Diploscapter_coronatus MKC-LSIFFVLLLAILLTNAMPEDRFRRSMAVANAKRQAASANGMRSGYIRFGKRKSNPNDDLMLDQLIL
Diploscapter_pachys KRQAASANGMRSGYIRFGKR

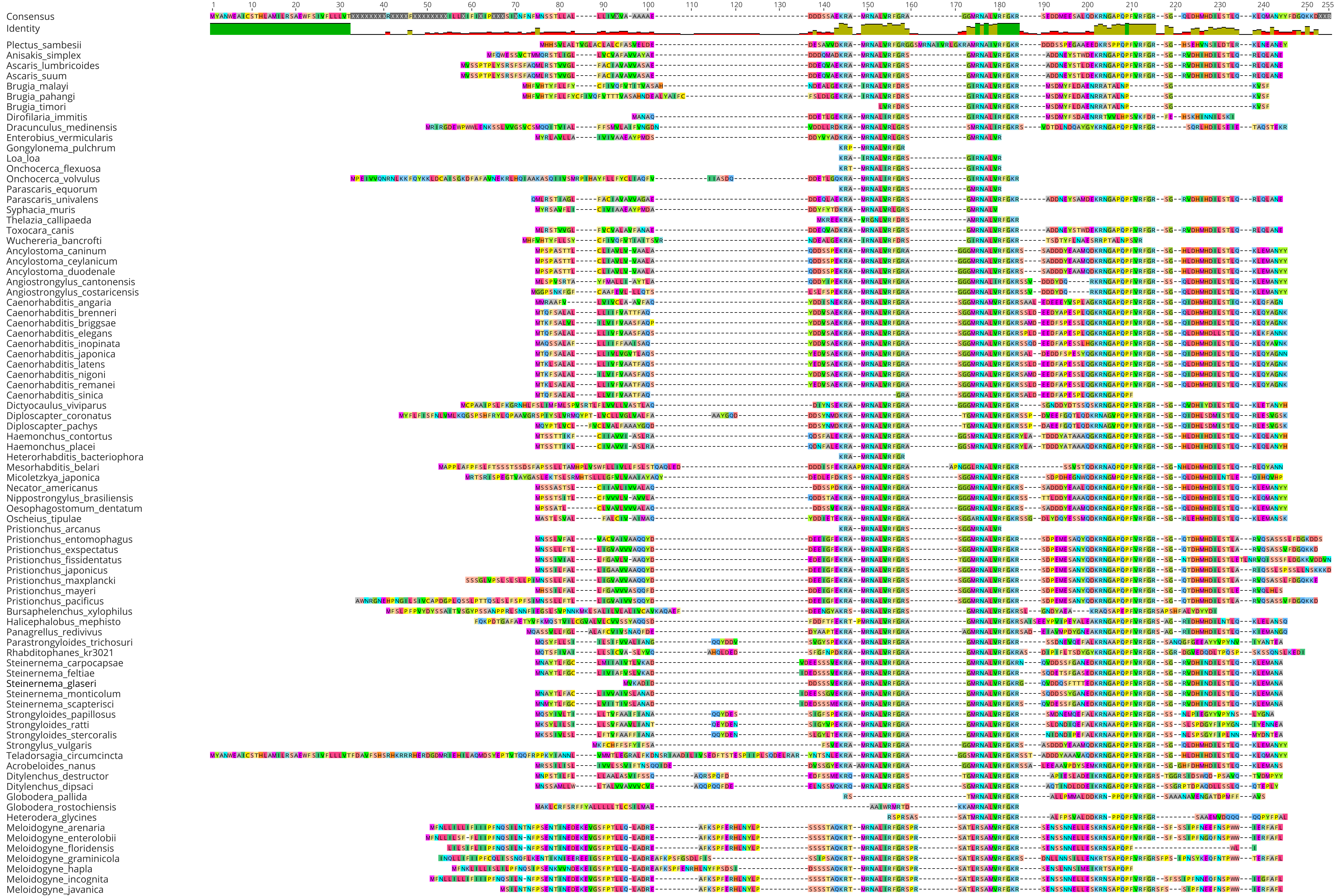
Consensus
Identity
1 10 20 30 40 50 60 70 80 90 100 110 120 130 140 150 160 170 180 190 200 210 220 230 240 250 255
Plectus_sambesii
Anisakis_simplex
Ascaris_lumbricoides
Ascaris_suum
Brugia_malayi
Brugia_pahangi
Brugia_timori
Dirofilaria_immitis
Dracunculus_medinensis
Enterobius_vermicularis
Gongylonema_pulchrum
Loa_loa
Onchocerca_flexuosa
Onchocerca_volvulus
Parascaris_equorum
Parascaris_univalens
Syphacia_muris
Thelazia_callipaeda
Toxocara_canis
Wuchereria_bancrofti
Ancylostoma_caninum
Ancylostoma_ceylanicum
Ancylostoma_duodenale
Angiostrongylus_cantonensis
Angiostrongylus_costaricensis
Caenorhabditis_angaria
Caenorhabditis_brenneri
Caenorhabditis_briggsae
Caenorhabditis_elegans
Caenorhabditis_inopinata
Caenorhabditis_japonica
Caenorhabditis_latens
Caenorhabditis_nigoni
Caenorhabditis_remanei
Caenorhabditis_sinica
Dictyocaulus_viviparus
Diploscapter_coronatus
Diploscapter_pachys
Haemonchus_contortus
Haemonchus_placei
Heterorhabditis_bacteriophora
Mesorhabditis_belari
Micoletzkya_japonica
Necator_americanus
Nippostrongylus_brasiliensis
Oesophagostomum_dentatum
Oscheius_tipulae
Pristionchus_arcanus
Pristionchus_entomophagus
Pristionchus_exspectatus
Pristionchus_fissidentatus
Pristionchus_japonicus
Pristionchus_maxplancki
Pristionchus_mayeri
Pristionchus_pacificus
Bursaphelenchus_xylophilus
Halicephalobus_mephisto
Panagrellus_redivivus
Parastrongyloides_trichosuri
Rhabditophanes_kr3021
Steinernema_carpocapsae
Steinernema_feltiae
Steinernema_glaseri
Steinernema_monticolum
Steinernema_scapterisci
Strongyloides_papillosus
Strongyloides_ratti
Strongyloides_stercoralis
Strongylus_vulgaris
Teladorsagia_circumcincta
Acrobeloides_nanus
Ditylenchus_destructor
Ditylenchus_dipsaci
Globodera_pallida
Globodera_rostochiensis
Heterodera_glycines
Meloidogyne_arenaria
Meloidogyne_enterolobii
Meloidogyne_floridensis
Meloidogyne_graminicola
Meloidogyne_hapla
Meloidogyne_incognita
Meloidogyne_javanica

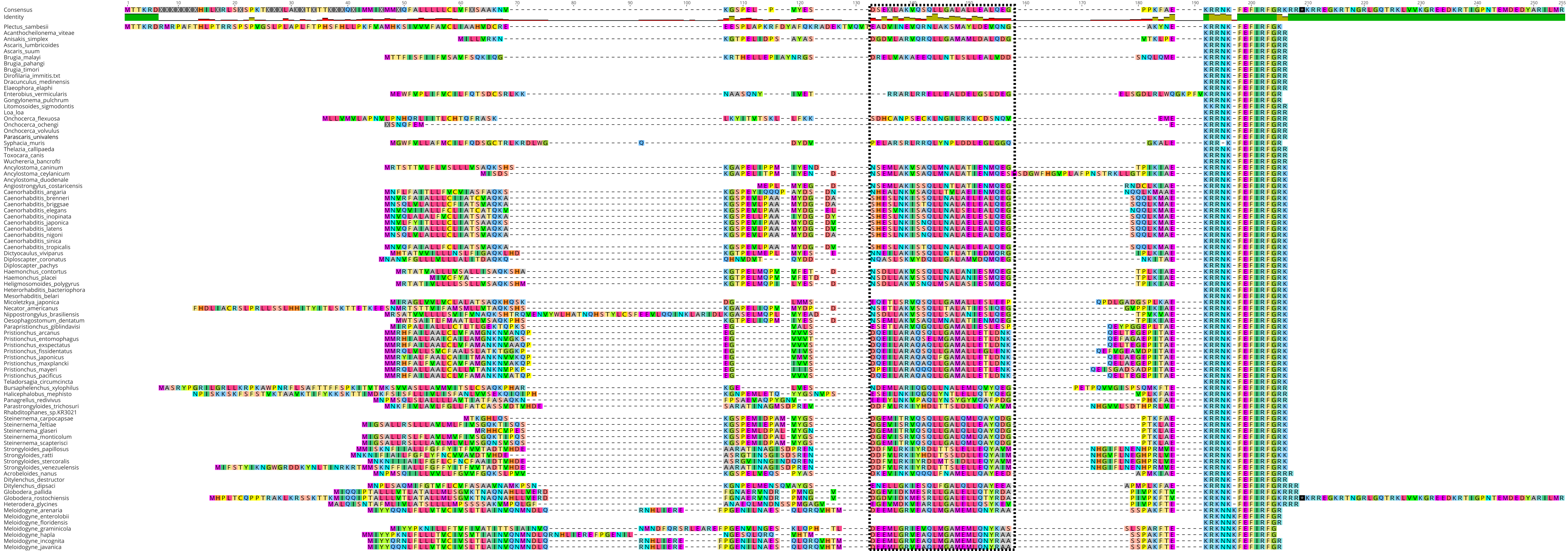
Consensus
Identity
Plectus_sambesii
Acanthocheilonema_viteae
Anisakis_simplex
Ascaris_lumbricoides
Ascaris_suum
Brugia_malayi
Brugia_pahangi
Brugia_timori
Dirofilaria_immitis.txt
Dracunculus_medinensis
Elaeophora_elaphi
Enterobius_vermicularis
Gongylonema_pulchrum
Litomosoides_sigmodontis
Loa_loa
Onchocerca_flexuosa
Onchocerca_ochengi
Onchocerca_volvulus
Parascaris_univalens
Syphacia_muris
Thelazia_callipaeda
Toxocara_canis
Wuchereria_bancrofti
Ancylostoma_caninum
Ancylostoma_ceylanicum
Ancylostoma_duodenale
Angiostrongylus_costaricensis
Caenorhabditis_angaria
Caenorhabditis_brenneri
Caenorhabditis_briggsae
Caenorhabditis_elegans
Caenorhabditis_inopinata
Caenorhabditis_japonica
Caenorhabditis_latens
Caenorhabditis_nigoni
Caenorhabditis_sinica
Caenorhabditis_tropicalis
Dictyocaulus_viviparus
Diploscapter_coronatus
Diploscapter_pachys
Haemonchus_contortus
Haemonchus_placei
Heligmosomoides_polygyrus
Heterorhabditis_bacteriophora
Mesorhabditis_belari
Micoletzkya_japonica
Necator_americanus
Nippostrongylus_brasiliensis
Oesophagostomum_dentatum
Parapristionchus_giblindavisi
Pristionchus_arcanus
Pristionchus_entomophagus
Pristionchus_exspectatus
Pristionchus_fissidentatus
Pristionchus_japonicus
Pristionchus_maxplancki
Pristionchus_mayeri
Pristionchus_pacificus
Teladorsagia_circumcincta
Bursaphelenchus_xylophilus
Halicephalobus_mephisto
Panagrellus_redivivus
Parastrongyloides_trichosuri
Rhabditophanes_sp.KR3021
Steinernema_carpocapsae
Steinernema_feltiae
Steinernema_glaseri
Steinernema_monticolum
Steinernema_scapterisci
Strongyloides_papillosus
Strongyloides_ratti
Strongyloides_stercoralis
Strongyloides_venezuelensis
Acrobeloides_nanus
Ditylenchus_destructor
Ditylenchus_dipsaci
Globodera_pallida
Globodera_rostochiensis
Heterodera_glycines
Meloidogyne_arenaria
Meloidogyne_enterolobii
Meloidogyne_floridensis
Meloidogyne_graminicola
Meloidogyne_hapla
Meloidogyne_incognita
Meloidogyne_javanica

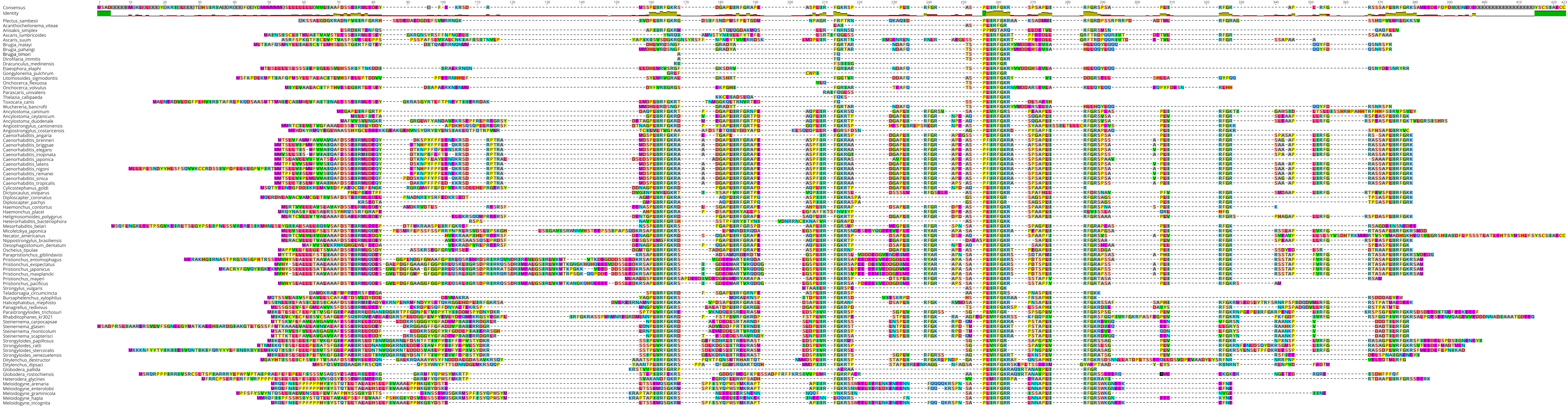

Consensus
Identity
Plectus_sambesii
Acanthocheilonema_viteae
Anisakis_simplex
Ascaris_lumbricoides
Ascaris_suum
Brugia_malayi
Brugia_pahangi
Brugia_timori
Dirofilaria_immitis
Dracunculus_medinensis
Elaeophora_elaphi
Gongylonema_pulchrum
Litomosoides_sigmodontis
Onchocerca_flexuosa
Onchocerca_volvulus
Parascaris_univalens
Thelazia_callipaeda
Toxocara_canis
Wuchereria_bancrofti
Ancylostoma_caninum
Ancylostoma_ceylanicum
Ancylostoma_duodenale
Angiostrongylus_cantonensis
Angiostrongylus_costaricensis
Caenorhabditis_angaria
Caenorhabditis_brenneri
Caenorhabditis_briggsae
Caenorhabditis_elegans
Caenorhabditis_inopinata
Caenorhabditis_japonica
Caenorhabditis_latens
Caenorhabditis_nigoni
Caenorhabditis_remanei
Caenorhabditis_sinica
Caenorhabditis_tropicalis
Cylicostephanus_goldi
Dictyocaulus_viviparus
Diploscapter_coronatus
Diploscapter_pachys
Haemonchus_contortus
Haemonchus_placei
Heligmosomoides_polygyrus
Heterorhabditis_bacteriophora
Mesorhabditis_belari
Micoletzkya_japonica
Necator_americanus
Nippostrongylus_brasiliensis
Oesophagostomum_dentatum
Oscheius_tipulae
Parapristionchus_giblindavisi
Pristionchus_entomophagus
Pristionchus_exspectatus
Pristionchus_japonicus
Pristionchus_maxplancki
Pristionchus_mayeri
Pristionchus_pacificus
Strongylus_vulgaris
Teladorsagia_circumcincta
Bursaphelenchus_xylophilus
Halicephalobus_mephisto
Panagrellus_redivivus
Parastrongyloides_trichosuri
Rhabditophanes_kr3021
Steinernema_carpocapsae
Steinernema_glaseri
Steinernema_monticolum
Steinernema_scapterisci
Strongyloides_papillosus
Strongyloides_ratti
Strongyloides_stercoralis
Strongyloides_venezuelensis
Ditylenchus_destructor
Ditylenchus_dipsaci
Globodera_pallida
Globodera_rostochiensis
Heterodera_glycines
Meloidogyne_arenaria
Meloidogyne_enterolobii
Meloidogyne_graminicola
Meloidogyne_hapla
Meloidogyne_incognita

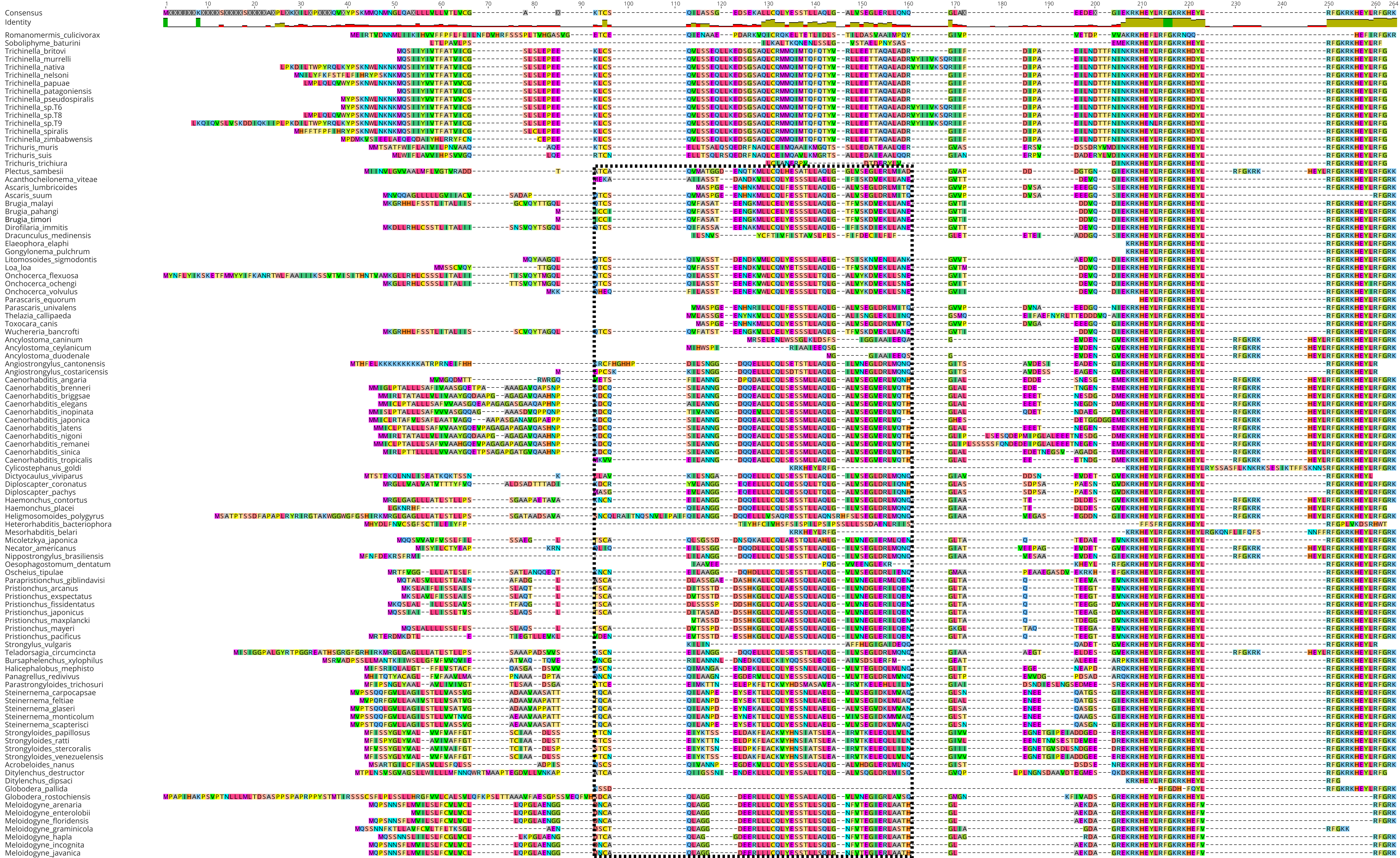

Consensus
Identity
1
10
20
30
40
50
60
70
80
90
100
110
120
130
140
150
160
170
180
190
200
210
220
230
240
250
260
Romanomermis_culicivorax
Soboliphyme_baturini
Trichinella_britovi
Trichinella_murrelli
Trichinella_nativa
Trichinella_nelsoni
Trichinella_papuae
Trichinella_patagoniensis
Trichinella_pseudospiralis
Trichinella_sp.T6
Trichinella_sp.T8
Trichinella_sp.T9
Trichinella_spiralis
Trichinella_zimbabwensis
Trichuris_muris
Trichuris_suis
Trichuris_trichiura
Plectus_sambesii
Acanthocheilonema_viteae
Ascaris_lumbricoides
Ascaris_suum
Brugia_malayi
Brugia_pahangi
Brugia_timori
Dirofilaria_immitis
Dracunculus_medinensis
Elaeophora_elaphi
Gongylonema_pulchrum
Litomosoides_sigmodontis
Loa_loa
Onchocerca_flexuosa
Onchocerca_ochengi
Onchocerca_volvulus
Parascaris_equorum
Parascaris_univalens
Thelazia_callipaeda
Toxocara_canis
Wuchereria_bancrofti
Ancylostoma_caninum
Ancylostoma_ceylanicum
Ancylostoma_duodenale
Angiostrongylus_cantonensis
Angiostrongylus_costaricensis
Caenorhabditis_angaria
Caenorhabditis_brenneri
Caenorhabditis_briggsae
Caenorhabditis_elegans
Caenorhabditis_inopinata
Caenorhabditis_japonica
Caenorhabditis_latens
Caenorhabditis_nigoni
Caenorhabditis_remanei
Caenorhabditis_sinica
Caenorhabditis_tropicalis
Cylicostephanus_goldi
Dictyocaulus_viviparus
Diploscapter_coronatus
Diploscapter_pachys
Haemonchus_contortus
Haemonchus_placei
Heligmosomoides_polygyrus
Heterorhabditis_bacteriophora
Mesorhabditis_belari
Micoletzkya_japonica
Necator_americanus
Nippostrongylus_brasiliensis
Oesophagostomum_dentatum
Oscheius_tipulae
Parapristionchus_giblindavisi
Pristionchus_arcanus
Pristionchus_exspectatus
Pristionchus_fissidentatus
Pristionchus_japonicus
Pristionchus_maxplancki
Pristionchus_mayeri
Pristionchus_pacificus
Strongylus_vulgaris
Teladorsagia_circumcincta
Bursaphelenchus_xylophilus
Halicephalobus_mephisto
Panagrellus_redivivus
Parastrongyloides_trichosuri
Steinernema_carpocapsae
Steinernema_feltiae
Steinernema_glaseri
Steinernema_monticolum
Steinernema_scapterisci
Strongyloides_papillosus
Strongyloides_ratti
Strongyloides_stercoralis
Strongyloides_venezuelensis
Acrobeloides_nanus
Ditylenchus_destructor
Ditylenchus_dipsaci
Globodera_pallida
Globodera_rostochiensis
Meloidogyne_arenaria
Meloidogyne_enterolobii
Meloidogyne_floridensis
Meloidogyne_graminicola
Meloidogyne_hapla
Meloidogyne_incognita
Meloidogyne_javanica

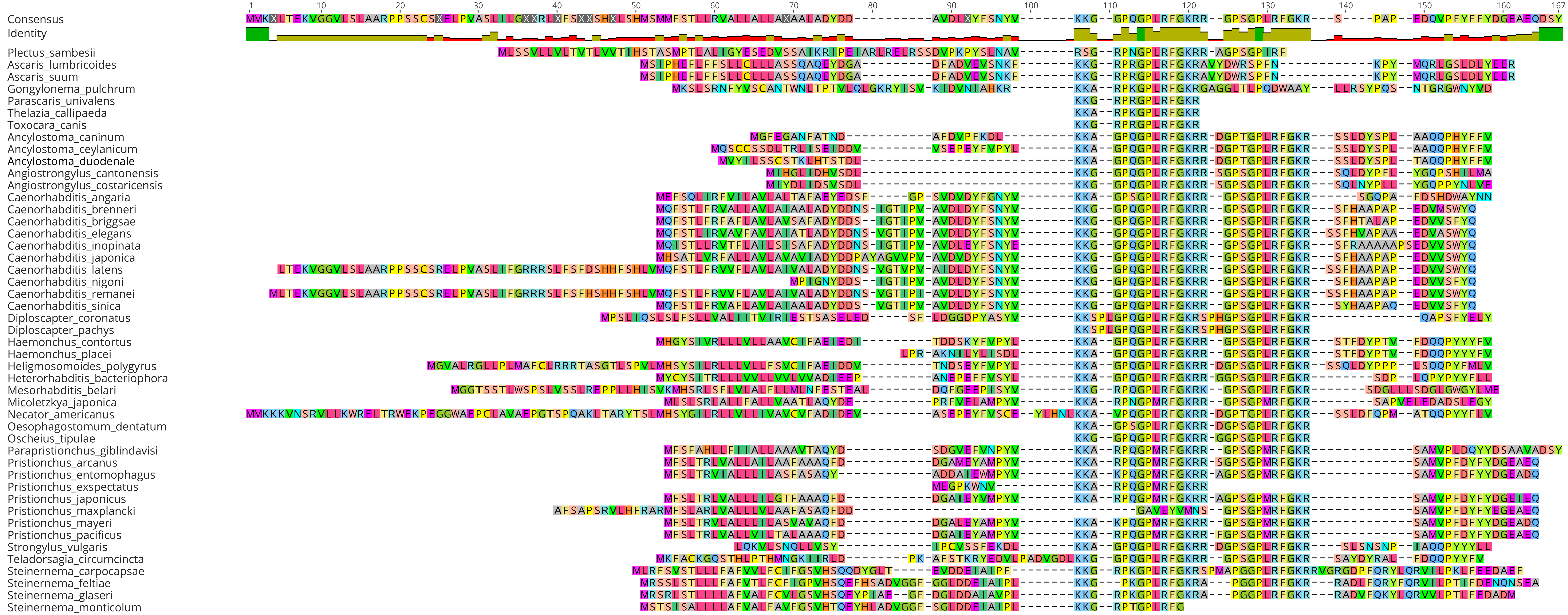

Consensus
Identity
1
10
20
30
40
50
60
70
80
90
100
110
120
130
140
150
160
167
Plectus_sambesii
Ascaris_lumbricoides
Ascaris_suum
Gongylonema_pulchrum
Parascaris_univalens
Thelazia_callipaeda
Toxocara_canis
Ancylostoma_caninum
Ancylostoma_ceylanicum
Ancylostoma_duodenale
Angiostrongylus_cantonensis
Angiostrongylus_costaricensis
Caenorhabditis_angaria
Caenorhabditis_brenneri
Caenorhabditis_briggsae
Caenorhabditis_elegans
Caenorhabditis_inopinata
Caenorhabditis_japonica
Caenorhabditis_latens
Caenorhabditis_nigoni
Caenorhabditis_remanei
Caenorhabditis_sinica
Diploscapter_coronatus
Diploscapter_pachys
Haemonchus_contortus
Haemonchus_placei
Heligmosomoides_polygyrus
Heterorhabditis_bacteriophora
Mesorhabditis_belari
Micoletzkya_japonica
Necator_americanus
Oesophagostomum_dentatum
Oscheius_tipulae
Parapristionchus_giblindavisi
Pristionchus_arcanus
Pristionchus_entomophagus
Pristionchus_exspectatus
Pristionchus_japonicus
Pristionchus_maxplancki
Pristionchus_mayeri
Pristionchus_pacificus
Strongylus_vulgaris
Teladorsagia_circumcincta
Steinernema_carpocapsae
Steinernema_feltiae
Steinernema_glaseri
Steinernema_monticolum

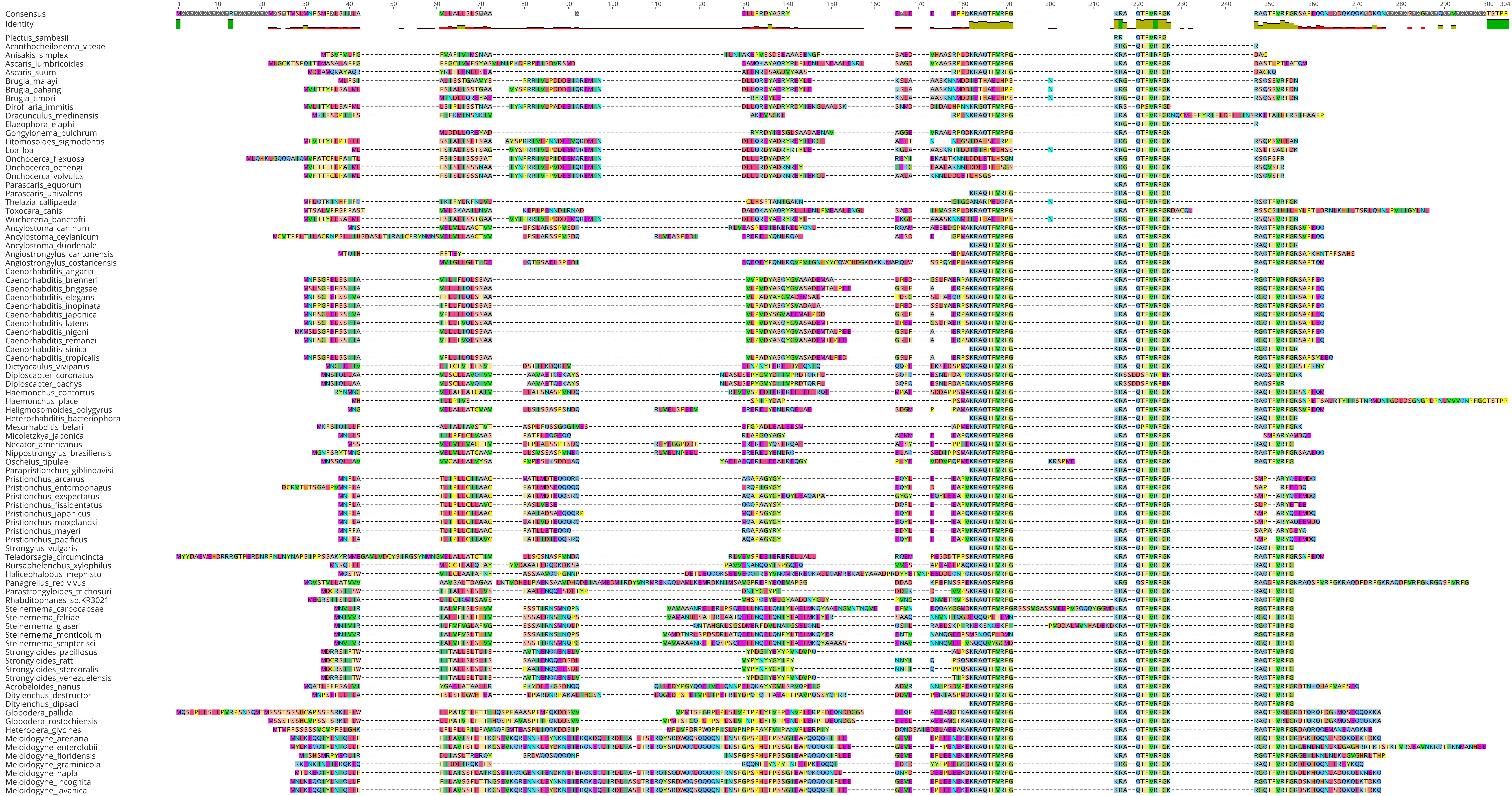

Consensus
Identity
Plectus_sambesii
Acanthocheilonema_viteae
Anisakis_simplex
Ascaris_lumbricoides
Ascaris_suum
Brugia_malayi
Brugia_pahangi
Brugia_timori
Dirofilaria_immitis
Dracunculus_medinensis
Elaeophora_elaphi
Gongylonema_pulchrum
Litomosoides_sigmodontis
Loa_loa
Onchocerca_flexuosa
Onchocerca_ochengi
Onchocerca_volvulus
Parascaris_equorum
Parascaris_univalens
Thelazia_callipaeda
Toxocara_canis
Wuchereria_bancrofti
Ancylostoma_caninum
Ancylostoma_ceylanicum
Ancylostoma_duodenale
Angiostrongylus_cantonensis
Angiostrongylus_costaricensis
Caenorhabditis_angaria
Caenorhabditis_brenneri
Caenorhabditis_briggsae
Caenorhabditis_elegans
Caenorhabditis_inopinata
Caenorhabditis_japonica
Caenorhabditis_latens
Caenorhabditis_nigoni
Caenorhabditis_remanei
Caenorhabditis_sinica
Caenorhabditis_tropicalis
Dictyocaulus_viviparus
Diploscapter_coronatus
Diploscapter_pachys
Haemonchus_contortus
Haemonchus_placei
Heligmosomoides_polygyrus
Heterorhabditis_bacteriophora
Mesorhabditis_belari
Micoletzkya_japonica
Necator_americanus
Nippostrongylus_brasiliensis
Oscheius_tipulae
Parapristionchus_giblindavisi
Pristionchus_arcanus
Pristionchus_entomophagus
Pristionchus_exspectatus
Pristionchus_fissidentatus
Pristionchus_japonicus
Pristionchus_maxplancki
Pristionchus_mayeri
Pristionchus_pacificus
Strongylus_vulgaris
Teladorsagia_circumcincta
Bursaphelenchus_xylophilus
Halicephalobus_mephisto
Panagrellus_redivivus
Parastrongyloides_trichosuri
Rhabditophanes_sp.KR3021
Steinernema_carpocapsae
Steinernema_feltiae
Steinernema_glaseri
Steinernema_monticolum
Steinernema_scapterisci
Strongyloides_papillosus
Strongyloides_ratti
Strongyloides_stercoralis
Strongyloides_venezuelensis
Acrobeloides_nanus
Ditylenchus_destructor
Ditylenchus_dipsaci
Globodera_pallida
Globodera_rostochiensis
Heterodera_glycines
Meloidogyne_arenaria
Meloidogyne_enterolobii
Meloidogyne_floridensis
Meloidogyne_graminicola
Meloidogyne_hapla
Meloidogyne_incognita
Meloidogyne_javanica

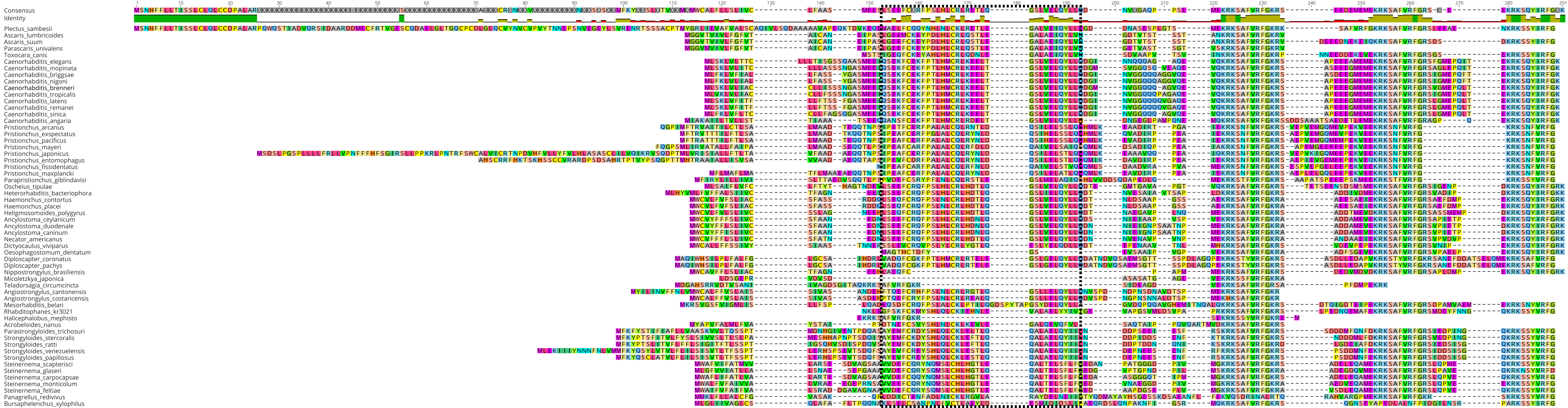
Consensus
Identity
Plectus_sambesii
Ascaris_lumbricoides
Ascaris_suum
Parascaris_univalens
Toxocara_canis
Caenorhabditis_elegans
Caenorhabditis_inopinata
Caenorhabditis_briggsae
Caenorhabditis_nigoni
Caenorhabditis_brenneri
Caenorhabditis_tropicalis
Caenorhabditis_latens
Caenorhabditis_remanei
Caenorhabditis_sinica
Caenorhabditis_angaria
Pristionchus_arcanus
Pristionchus_exspectatus
Pristionchus_pacificus
Pristionchus_mayeri
Pristionchus_japonicus
Pristionchus_entomophagus
Pristionchus_fissidentatus
Pristionchus_maxplancki
Parapristionchus_giblindavisi
Oscheius_tipulae
Heterorhabditis_bacteriophora
Haemonchus_contortus
Haemonchus_placei
Heligmosomoides_polygyrus
Ancylostoma_ceylanicum
Ancylostoma_duodenale
Ancylostoma_caninum
Necator_americanus
Dictyocaulus_viviparus
Oesophagostomum_dentatum
Diploscapter_coronatus
Diploscapter_pachys
Nippostrongylus_brasiliensis
Micoletzkya_japonica
Teladorsagia_circumcincta
Angiostrongylus_cantonensis
Angiostrongylus_costaricensis
Mesorhabditis_belari
Rhabditophanes_kr3021
Halicephalobus_mephisto
Acrobeloides_nanus
Parastrongyloides_trichosuri
Strongyloides_stercoralis
Strongyloides_ratti
Strongyloides_venezuelensis
Strongyloides_papillosus
Steinernema_scapterisci
Steinernema_glaseri
Steinernema_carpocapsae
Steinernema_monticolum
Steinernema_feltiae
Panagrellus_redivivus
Bursaphelenchus_xylophilus

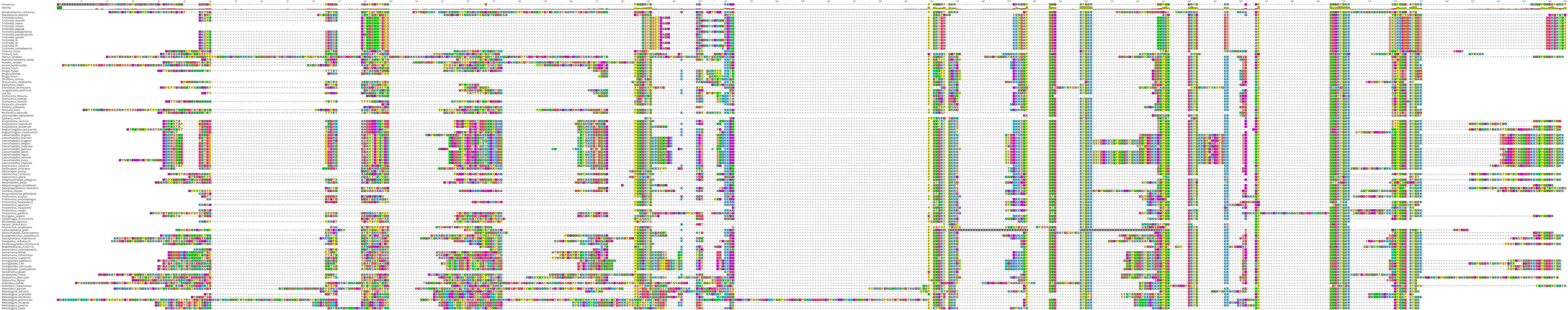
Consensus
Identity
Romanomermis_culicivorax
Soboliphyme_baturini
Trichinella_britovi
Trichinella_murrelli
Trichinella_nativa
Trichinella_nelsoni
Trichinella_papuae
Trichinella_patagoniensis
Trichinella_pseudospiralis
Trichinella_spiralis
Trichinella_t6
Trichinella_t8
Trichinella_t9
Trichinella_zimbabwensis
Trichuris_muris
Trichuris_suis
Plectus_sambesii
Acanthocheilonema_viteae
Anisakis_simplex
Ascaris_lumbricoides
Ascaris_suum
Brugia_malayi
Brugia_pahangi
Brugia_timori
Dirofilaria_immitis
Dracunculus_medinensis
Elaeophora_elaphi
Enterobius_vermicularis
Gongylonema_pulchrum
Loa_loa
Onchocerca_flexuosa
Onchocerca_ochengi
Onchocerca_volvulus
Parascaris_univalens
Thelazia_callipaeda
Toxocara_canis
Wuchereria_bancrofti
Litomosoides_sigmodontis
Syphacia_muris
Ancylostoma_caninum
Ancylostoma_ceylanicum
Ancylostoma_duodenale
Angiostrongylus_cantonensis
Angiostrongylus_costaricensis
Caenorhabditis_angaria
Caenorhabditis_brenneri
Caenorhabditis_briggsae
Caenorhabditis_elegans
Caenorhabditis_inopinata
Caenorhabditis_japonica
Caenorhabditis_latens
Caenorhabditis_nigoni
Caenorhabditis_remanei
Caenorhabditis_sinica
Caenorhabditis_tropicalis
Dictyocaulus_viviparus
Diploscapter_coronatus
Diploscapter_pachys
Haemonchus_contortus
Haemonchus_placei
Heligmosomoides_polygyrus
Mesorhabditis_belari
Nippostrongylus_brasiliensis
Oesophagostomum_dentatum
Oscheius_tipulae
Parapristionchus_giblindavisi
Pristionchus_arcanus
Pristionchus_entomophagus
Pristionchus_fissidentatus
Pristionchus_japonicus
Pristionchus_maxplancki
Pristionchus_mayeri
Pristionchus_pacificus
Strongylus_vulgaris
Teladorsagia_circumcincta
Micoletzkya_japonica
Necator_americanus
Pristionchus_exspectatus
Cylicostephanus_goldi
Heterorhabditis_bacteriophora
Bursaphelenchus_xylophilus.fa
Halicephalobus_mephisto.fa
Panagrellus_redivivus.fa
Parastrongyloides_trichosuri.fa
Rhabditophanes_kr3021.fa
Steinernema_carpocapsae.fa
Steinernema_feltiae
Steinernema_monticolum
Steinernema_scapterisci
Strongyloides_papillosus
Strongyloides_ratti
Strongyloides_stercoralis
Strongyloides_venezuelensis
Steinernema_glaseri
Acrobeloides_nanus
Ditylenchus_destructor
Ditylenchus_dipsaci
Globodera_pallida
Globodera_rostochiensis
Heterodera_glycines
Meloidogyne_arenaria
Meloidogyne_enterolobii
Meloidogyne_floridensis
Meloidogyne_graminicola
Meloidogyne_incognita
Meloidogyne_javanica
Meloidogyne_hapla

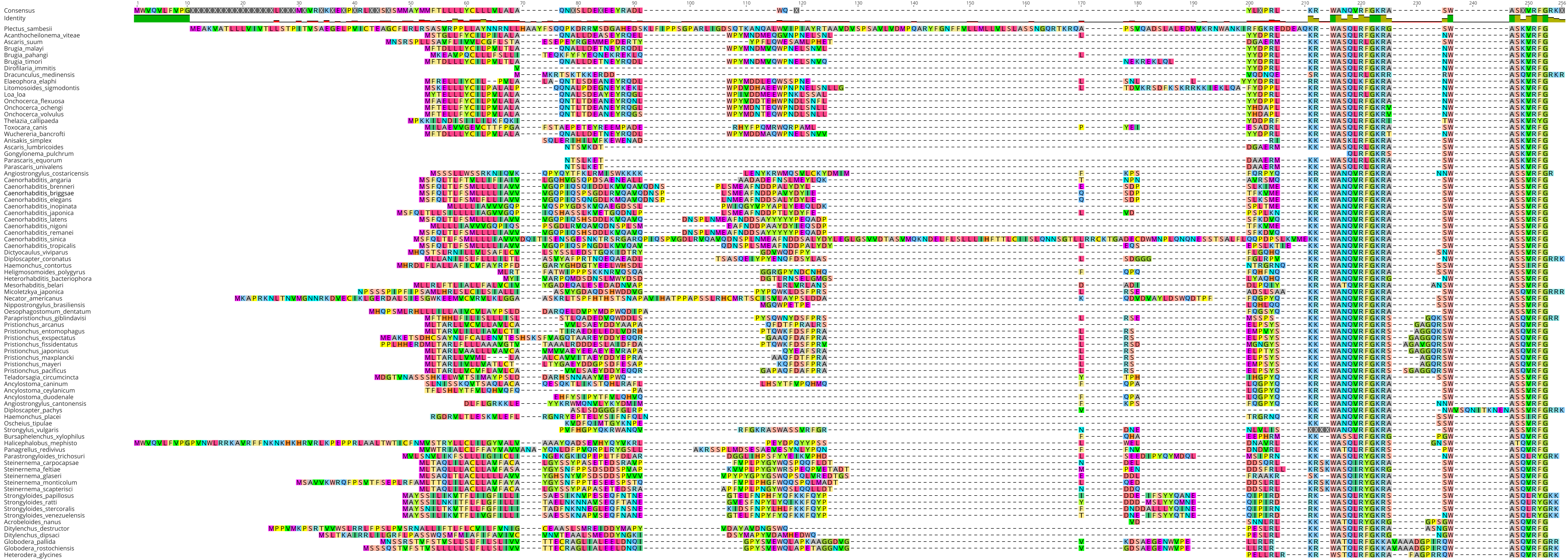
Consensus
Identity
Plectus_sambesii
Acanthocheilonema_viteae
Ascaris_suum
Brugia_malayi
Brugia_pahangi
Brugia_timori
Dirofilaria_immitis
Dracunculus_medinensis
Elaeophora_elaphi
Litomosoides_sigmodontis
Loa_loa
Onchocerca_flexuosa
Onchocerca_ochengi
Onchocerca_volvulus
Thelazia_callipaeda
Toxocara_canis
Wuchereria_bancrofti
Anisakis_simplex
Ascaris_lumbricoides
Gongylonema_pulchrum
Parascaris_equorum
Parascaris_univalens
Angiostrongylus_costaricensis
Caenorhabditis_angaria
Caenorhabditis_brenneri
Caenorhabditis_briggsae
Caenorhabditis_elegans
Caenorhabditis_inopinata
Caenorhabditis_japonica
Caenorhabditis_latens
Caenorhabditis_nigoni
Caenorhabditis_remanei
Caenorhabditis_sinica
Caenorhabditis_tropicalis
Dictyocaulus_viviparus
Diploscapter_coronatus
Haemonchus_contortus
Heligmosomoides_polygyrus
Heterorhabditis_bacteriophora
Mesorhabditis_belari
Micoletzkya_japonica
Necator_americanus
Nippostrongylus_brasiliensis
Oesophagostomum_dentatum
Parapristionchus_giblindavisi
Pristionchus_arcanus
Pristionchus_entomophagus
Pristionchus_exspectatus
Pristionchus_fissidentatus
Pristionchus_japonicus
Pristionchus_maxplancki
Pristionchus_mayeri
Pristionchus_pacificus
Teladorsagia_circumcincta
Ancylostoma_caninum
Ancylostoma_ceylanicum
Ancylostoma_duodenale
Angiostrongylus_cantonensis
Diploscapter_pachys
Haemonchus_placei
Oscheius_tipulae
Strongylus_vulgaris
Bursaphelenchus_xylophilus
Halicephalobus_mephisto
Panagrellus_redivivus
Parastrongyloides_trichosuri
Steinernema_carpocapsae
Steinernema_feltiae
Steinernema_glaseri
Steinernema_monticolum
Steinernema_scapterisci
Strongyloides_papillosus
Strongyloides_ratti
Strongyloides_stercoralis
Strongyloides_venezuelensis
Acrobeloides_nanus
Ditylenchus_destructor
Ditylenchus_dipsaci
Globodera_pallida
Globodera_rostochiensis
Heterodera_glycines

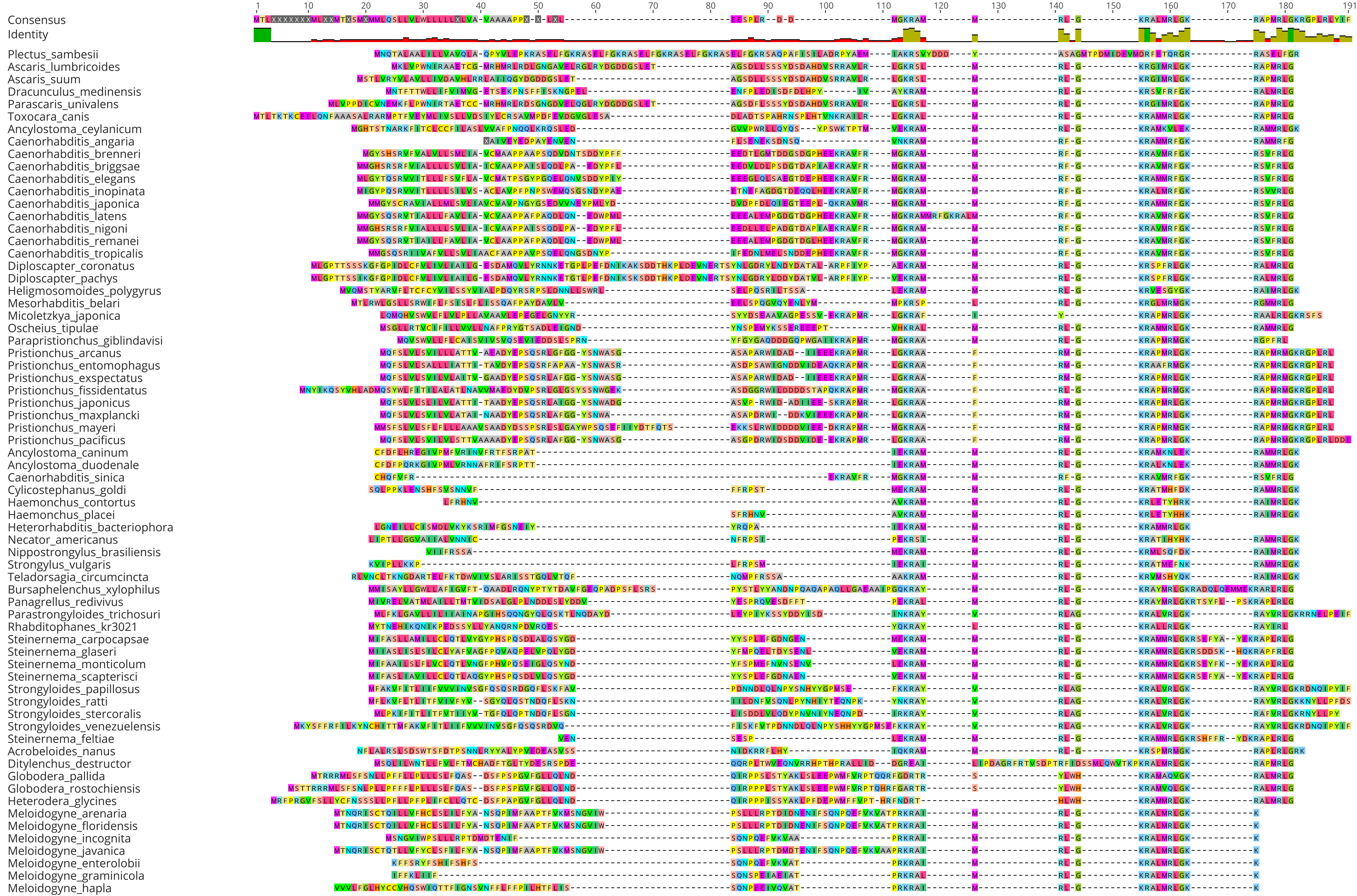

1
10
20
30
40
50
60
70
80
90
100
110
120
130
140
150
160
170
180
191
Consensus
Identity
Plectus_sambesii
Ascaris_lumbricoides
Ascaris_suum
Dracunculus_medinensis
Parascaris_univalens
Toxocara_canis
Ancylostoma_ceylanicum
Caenorhabditis_angaria
Caenorhabditis_brenneri
Caenorhabditis_briggsae
Caenorhabditis_elegans
Caenorhabditis_inopinata
Caenorhabditis_japonica
Caenorhabditis_latens
Caenorhabditis_nigoni
Caenorhabditis_remanei
Caenorhabditis_tropicalis
Diploscapter_coronatus
Diploscapter_pachys
Heligmosomoides_polygyrus
Mesorhabditis_belari
Micoletzkya_japonica
Oscheius_tipulae
Parapristionchus_giblindavisi
Pristionchus_arcanus
Pristionchus_entomophagus
Pristionchus_exspectatus
Pristionchus_fissidentatus
Pristionchus_japonicus
Pristionchus_maxplancki
Pristionchus_mayeri
Pristionchus_pacificus
Ancylostoma_caninum
Ancylostoma_duodenale
Caenorhabditis_sinica
Cylicostephanus_goldi
Haemonchus_contortus
Haemonchus_placei
Heterorhabditis_bacteriophora
Necator_americanus
Nippostrongylus_brasiliensis
Strongylus_vulgaris
Teladorsagia_circumcincta
Bursaphelenchus_xylophilus
Panagrellus_redivivus
Parastrongyloides_trichosuri
Rhabditophanes_kr3021
Steinernema_carpocapsae
Steinernema_glaseri
Steinernema_monticolum
Steinernema_scapterisci
Strongyloides_papillosus
Strongyloides_ratti
Strongyloides_stercoralis
Strongyloides_venezuelensis
Steinernema_feltiae
Acrobeloides_nanus
Ditylenchus_destructor
Globodera_pallida
Globodera_rostochiensis
Heterodera_glycines
Meloidogyne_arenaria
Meloidogyne_floridensis
Meloidogyne_incognita
Meloidogyne_javanica
Meloidogyne_enterolobii
Meloidogyne_graminicola
Meloidogyne_hapla

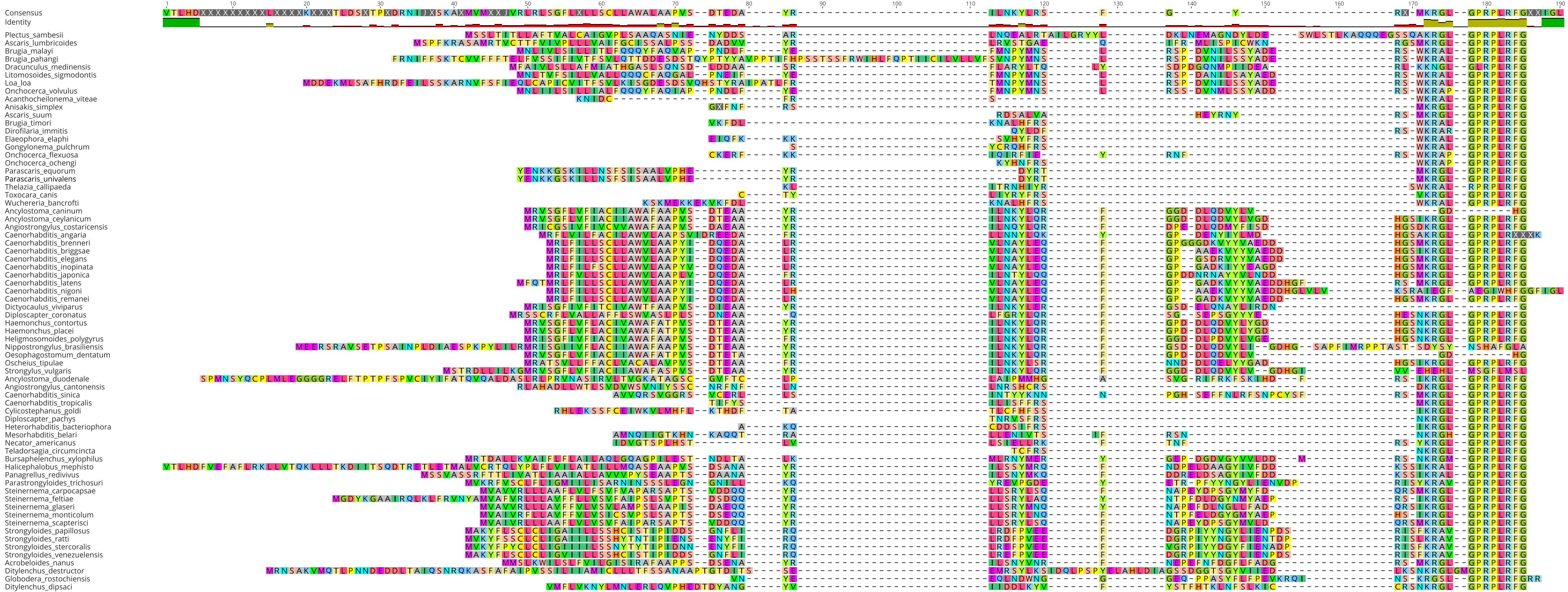

Consensus
Identity
Plectus_sambesii
Ascaris_lumbricoides
Brugia_malayi
Brugia_pahangi
Dracunculus_medinensis
Litomosoides_sigmodontis
Loa_loa
Onchocerca_volvulus
Acanthocheilonema_viteae
Anisakis_simplex
Ascaris_suum
Brugia_timori
Dirofilaria_immitis
Elaeophora_elaphi
Gongylonema_pulchrum
Onchocerca_flexuosa
Onchocerca_ochengi
Parascaris_equorum
Parascaris_univalens
Thelazia_callipaeda
Toxocara_canis
Wuchereria_bancrofti
Ancylostoma_caninum
Ancylostoma_ceylanicum
Angiostrongylus_costaricensis
Caenorhabditis_angaria
Caenorhabditis_brenneri
Caenorhabditis_briggsae
Caenorhabditis_elegans
Caenorhabditis_inopinata
Caenorhabditis_japonica
Caenorhabditis_latens
Caenorhabditis_nigoni
Caenorhabditis_remanei
Dictyocaulus_viviparus
Diploscapter_coronatus
Haemonchus_contortus
Haemonchus_placei
Heligmosomoides_polygyrus
Nippostrongylus_brasiliensis
Oesophagostomum_dentatum
Oscheius_tipulae
Strongylus_vulgaris
Ancylostoma_duodenale
Angiostrongylus_cantonensis
Caenorhabditis_sinica
Caenorhabditis_tropicalis
Cylicostephanus_goldi
Diploscapter_pachys
Heterorhabditis_bacteriophora
Mesorhabditis_belari
Necator_americanus
Teladorsagia_circumcincta
Bursaphelenchus_xylophilus
Halicephalobus_mephisto
Panagrellus_redivivus
Parastrongyloides_trichosuri
Steinernema_carpocapsae
Steinernema_feltiae
Steinernema_glaseri
Steinernema_monticolum
Steinernema_scapterisci
Strongyloides_papillosus
Strongyloides_ratti
Strongyloides_stercoralis
Strongyloides_venezuelensis
Acrobeloides_nanus
Ditylenchus_destructor
Globodera_rostochiensis
Ditylenchus_dipsaci

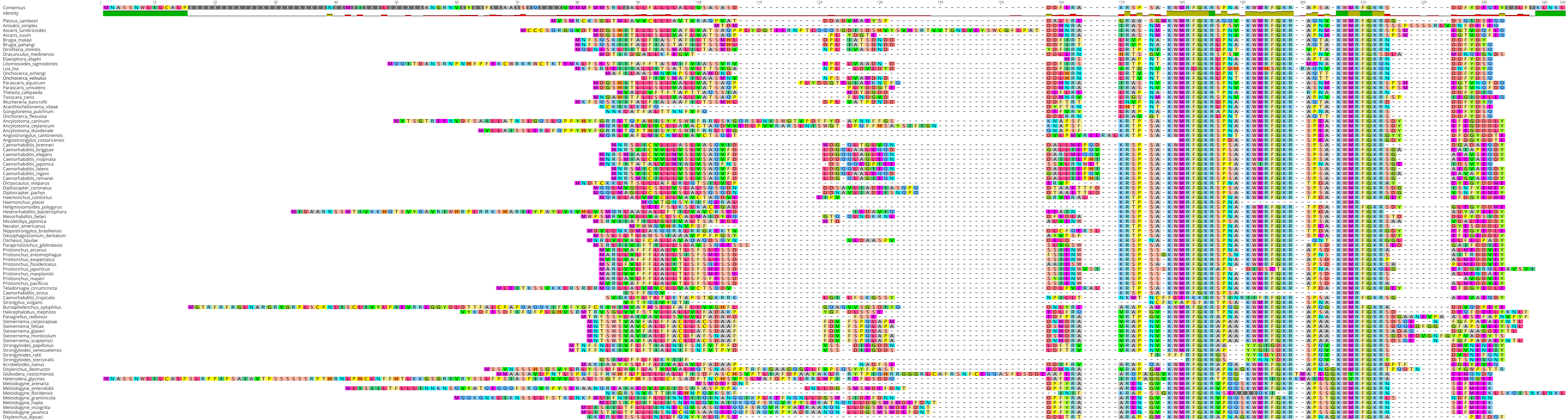
Consensus
Identity
Plectus_sambesii
Anisakis_simplex
Ascaris_lumbricoides
Ascaris_suum
Brugia_malayi
Brugia_pahangi
Dirofilaria_immitis
Dracunculus_medinensis
Elaeophora_elaphi
Litomosoides_sigmodontis
Loa_loa
Onchocerca_ochengi
Onchocerca_volvulus
Parascaris_equorum
Parascaris_univalens
Thelazia_callipaeda
Toxocara_canis
Wuchereria_bancrofti
Acanthocheilonema_viteae
Gongylonema_pulchrum
Onchocerca_flexuosa
Ancylostoma_caninum
Ancylostoma_ceylanicum
Ancylostoma_duodenale
Angiostrongylus_cantonensis
Angiostrongylus_costaricensis
Caenorhabditis_brenneri
Caenorhabditis_briggsae
Caenorhabditis_elegans
Caenorhabditis_inopinata
Caenorhabditis_japonica
Caenorhabditis_latens
Caenorhabditis_nigoni
Caenorhabditis_remanei
Dictyocaulus_viviparus
Diploscapter_coronatus
Diploscapter_pachys
Haemonchus_contortus
Haemonchus_placei
Heligmosomoides_polygyrus
Heterorhabditis_bacteriophora
Mesorhabditis_belari
Micoletzkya_japonica
Necator_americanus
Nippostrongylus_brasiliensis
Oesophagostomum_dentatum
Oscheius_tipulae
Parapristionchus_giblindavisi
Pristionchus_arcanus
Pristionchus_entomophagus
Pristionchus_exspectatus
Pristionchus_fissidentatus
Pristionchus_japonicus
Pristionchus_maxplancki
Pristionchus_mayeri
Pristionchus_pacificus
Teladorsagia_circumcincta
Caenorhabditis_sinica
Caenorhabditis_tropicalis
Strongylus_vulgaris
Bursaphelenchus_xylophilus
Halicephalobus_mephisto
Panagrellus_redivivus
Steinernema_carpocapsae
Steinernema_feltiae
Steinernema_glaseri
Steinernema_monticolum
Steinernema_scapterisci
Strongyloides_papillosus
Strongyloides_venezuelensis
Strongyloides_ratti
Strongyloides_stercoralis
Acrobeloides_nanus
Ditylenchus_destructor
Globodera_rostochiensis
Heterodera_glycines
Meloidogyne_arenaria
Meloidogyne_enterolobii
Meloidogyne_floridensis
Meloidogyne_graminicola
Meloidogyne_hapla
Meloidogyne_incognita
Meloidogyne_javanica
Ditylenchus_dipsaci

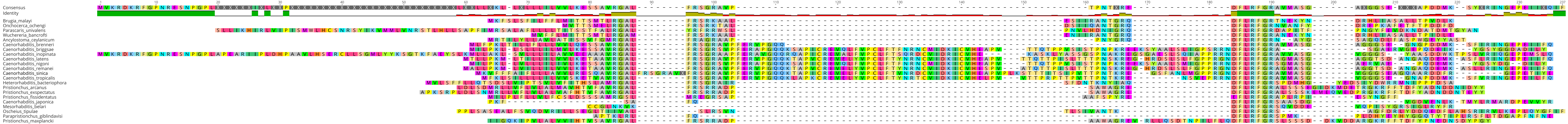
Consensus
Identity
Brugia_malayi
Onchocerca_ochengi
Parascaris_univalens
Wuchereria_bancrofti
Ancylostoma_ceylanicum
Caenorhabditis_brenneri
Caenorhabditis_briggsae
Caenorhabditis_inopinata
Caenorhabditis_latens
Caenorhabditis_nigoni
Caenorhabditis_remanei
Caenorhabditis_sinica
Caenorhabditis_tropicalis
Heterorhabditis_bacteriophora
Pristionchus_arcanus
Pristionchus_exspectatus
Pristionchus_fissidentatus
Caenorhabditis_japonica
Mesorhabditis_belari
Oscheius_tipulae
Parapristionchus_giblindavisi
Pristionchus_maxplancki

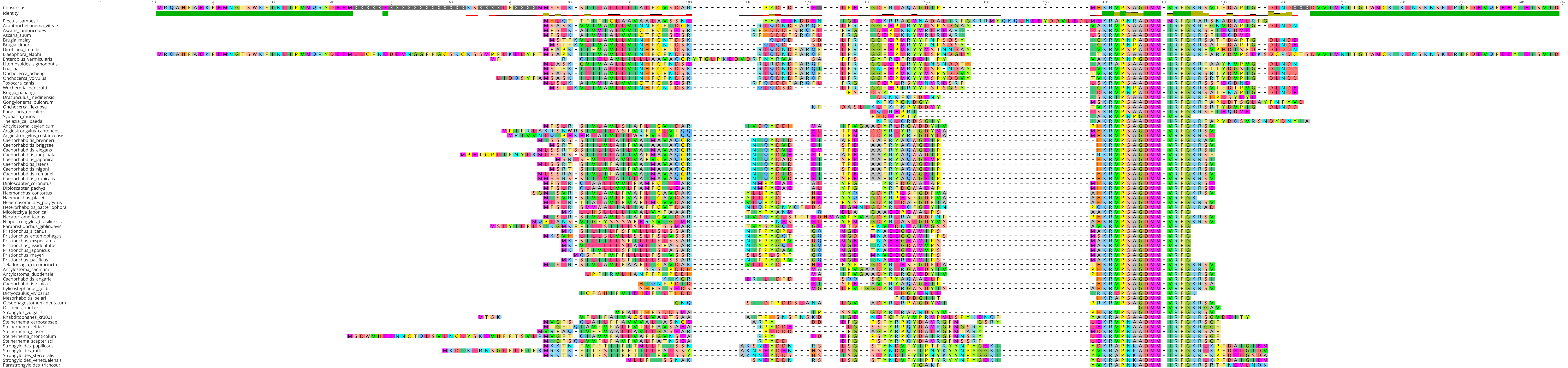
Consensus
Identity
Plectus_sambesii
Acanthocheilonema_viteae
Ascaris_lumbricoides
Ascaris_suum
Brugia_malayi
Brugia_timori
Dirofilaria_immitis
Elaeophora_elaphi
Enterobius_vermicularis
Litomosoides_sigmodontis
Loa_loa
Onchocerca_ochengi
Onchocerca_volvulus
Toxocara_canis
Wuchereria_bancrofti
Brugia_pahangi
Dracunculus_medinensis
Gongylonema_pulchrum
Onchocerca_flexuosa
Parascaris_univalens
Syphacia_muris
Thelazia_callipaeda
Ancylostoma_ceylanicum
Angiostrongylus_cantonensis
Angiostrongylus_costaricensis
Caenorhabditis_brenneri
Caenorhabditis_briggsae
Caenorhabditis_elegans
Caenorhabditis_inopinata
Caenorhabditis_japonica
Caenorhabditis_latens
Caenorhabditis_nigoni
Caenorhabditis_remanei
Caenorhabditis_tropicalis
Diploscapter_coronatus
Diploscapter_pachys
Haemonchus_contortus
Haemonchus_placei
Heligmosomoides_polygyrus
Heterorhabditis_bacteriophora
Micoletzkya_japonica
Necator_americanus
Nippostrongylus_brasiliensis
Parapristionchus_giblindavisi
Pristionchus_arcanus
Pristionchus_entomophagus
Pristionchus_exspectatus
Pristionchus_fissidentatus
Pristionchus_japonicus
Pristionchus_mayeri
Pristionchus_pacificus
Teladorsagia_circumcincta
Ancylostoma_caninum
Ancylostoma_duodenale
Caenorhabditis_angaria
Caenorhabditis_sinica
Cylicostephanus_goldi
Dictyocaulus_viviparus
Mesorhabditis_belari
Oesophagostomum_dentatum
Oscheius_tipulae
Strongylus_vulgaris
Rhabditophanes_kr3021
Steinernema_carpocapsae
Steinernema_feltiae
Steinernema_glaseri
Steinernema_monticolum
Steinernema_scapterisci
Strongyloides_papillosus
Strongyloides_ratti
Strongyloides_stercoralis
Strongyloides_venezuelensis
Parastrongyloides_trichosuri

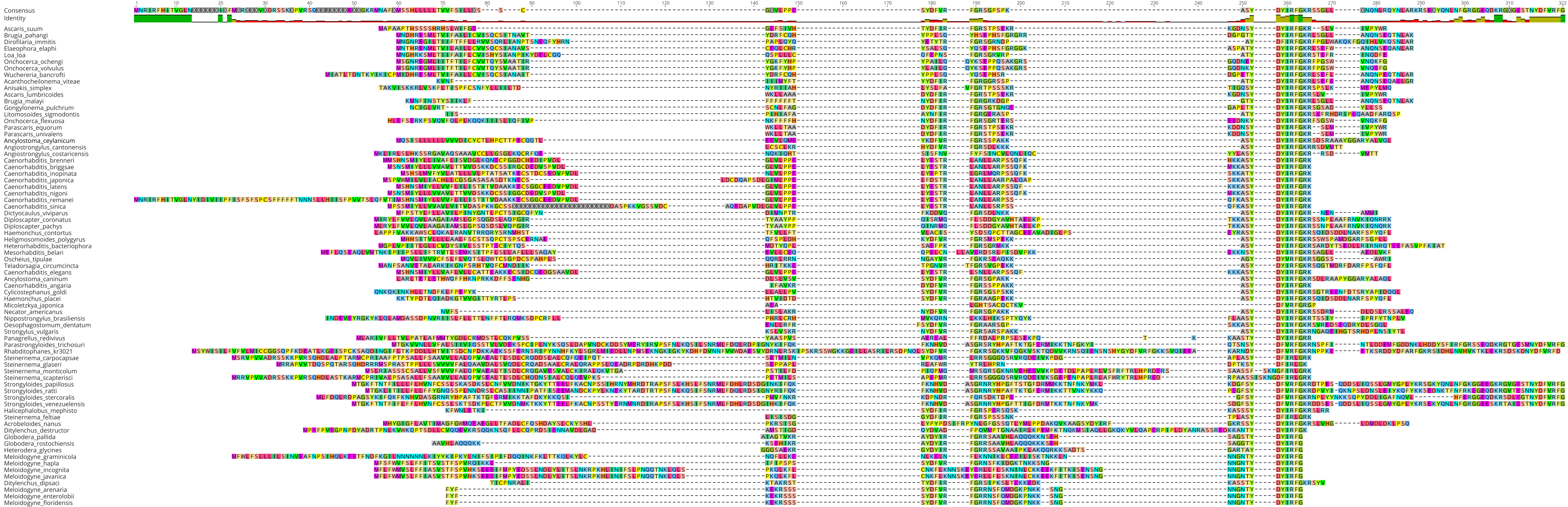

Consensus
Identity
Ascaris_suum
Brugia_pahangi
Dirofilaria_immitis
Elaeophora_elaphi
Loa_loa
Onchocerca_ochengi
Onchocerca_volvulus
Wuchereria_bancrofti
Acanthocheilonema_viteae
Anisakis_simplex
Ascaris_lumbricoides
Brugia_malayi
Gongylonema_pulchrum
Litomosoides_sigmodontis
Onchocerca_flexuosa
Parascaris_equorum
Parascaris_univalens
Ancylostoma_ceylanicum
Angiostrongylus_cantonensis
Angiostrongylus_costaricensis
Caenorhabditis_brenneri
Caenorhabditis_briggsae
Caenorhabditis_inopinata
Caenorhabditis_japonica
Caenorhabditis_latens
Caenorhabditis_nigoni
Caenorhabditis_remanei
Caenorhabditis_sinica
Dictyocaulus_viviparus
Diploscapter_coronatus
Diploscapter_pachys
Haemonchus_contortus
Heligmosomoides_polygyrus
Heterorhabditis_bacteriophora
Mesorhabditis_belari
Oscheius_tipulae
Teladorsagia_circumcincta
Caenorhabditis_elegans
Ancylostoma_caninum
Caenorhabditis_angaria
Cylicostephanus_goldi
Haemonchus_placei
Micoletzkya_japonica
Necator_americanus
Nippostrongylus_brasiliensis
Oesophagostomum_dentatum
Strongylus_vulgaris
Panagrellus_redivivus
Parastrongyloides_trichosuri
Rhabditophanes_kr3021
Steinernema_carpocapsae
Steinernema_glaseri
Steinernema_monticolum
Steinernema_scapterisci
Strongyloides_papillosus
Strongyloides_ratti
Strongyloides_stercoralis
Strongyloides_venezuelensis
Halicephalobus_mephisto
Steinernema_feltiae
Acrobeloides_nanus
Ditylenchus_destructor
Globodera_pallida
Globodera_rostochiensis
Heterodera_glycines
Meloidogyne_graminicola
Meloidogyne_hapla
Meloidogyne_incognita
Meloidogyne_javanica
Ditylenchus_dipsaci
Meloidogyne_arenaria
Meloidogyne_enterolobii
Meloidogyne_floridensis

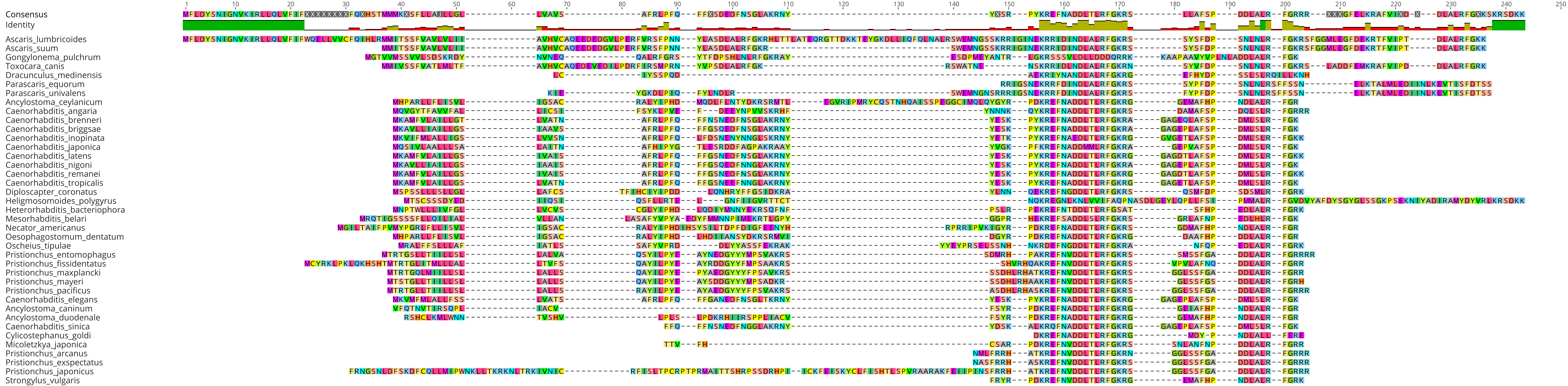

Consensus
Identity
Ascaris_lumbricoides
Ascaris_suum
Gongylonema_pulchrum
Toxocara_canis
Dracunculus_medinensis
Parascaris_equorum
Parascaris_univalens
Ancylostoma_ceylanicum
Caenorhabditis_angaria
Caenorhabditis_brenneri
Caenorhabditis_briggsae
Caenorhabditis_inopinata
Caenorhabditis_japonica
Caenorhabditis_latens
Caenorhabditis_nigoni
Caenorhabditis_remanei
Caenorhabditis_tropicalis
Diploscapter_coronatus
Heligmosomoides_polygyrus
Heterorhabditis_bacteriophora
Mesorhabditis_belari
Necator_americanus
Oesophagostomum_dentatum
Oscheius_tipulae
Pristionchus_entomophagus
Pristionchus_fissidentatus
Pristionchus_maxplancki
Pristionchus_mayeri
Pristionchus_pacificus
Caenorhabditis_elegans
Ancylostoma_caninum
Ancylostoma_duodenale
Caenorhabditis_sinica
Cylicostephanus_goldi
Micoletzkya_japonica
Pristionchus_arcanus
Pristionchus_exspectatus
Pristionchus_japonicus
Strongylus_vulgaris

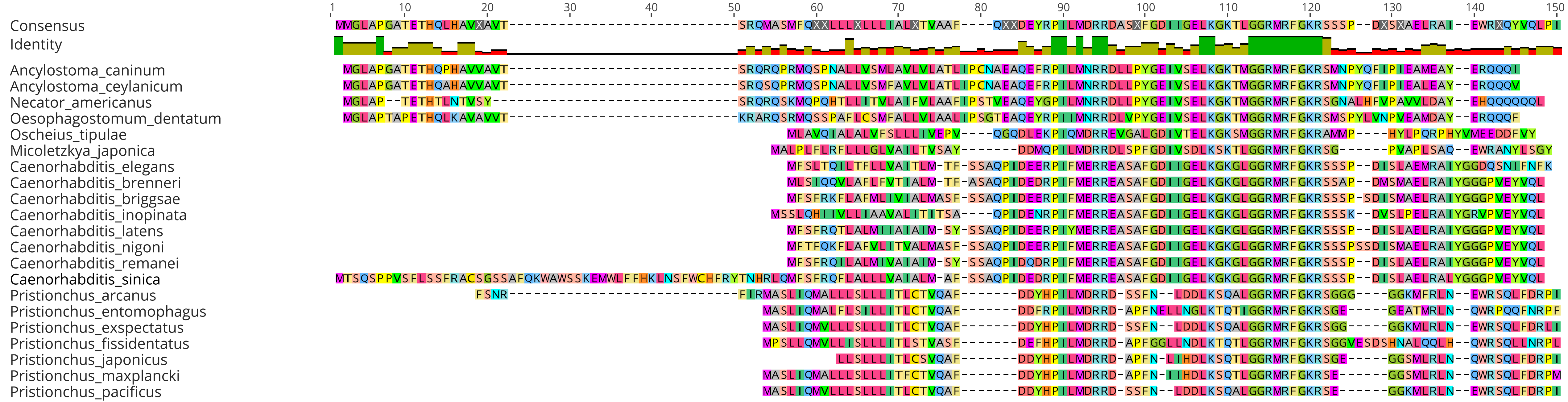
1
10
20
30
40
50
60
70
80
90
100
110
120
130
140
150
Consensus
Identity
Ancylostoma_caninum
Ancylostoma_ceylanicum
Necator_americanus
Oesophagostomum_dentatum
Oscheius_tipulae
Micoletzkya_japonica
Caenorhabditis_elegans
Caenorhabditis_brenneri
Caenorhabditis_briggsae
Caenorhabditis_inopinata
Caenorhabditis_latens
Caenorhabditis_nigoni
Caenorhabditis_remanei
Caenorhabditis_sinica
Pristionchus_arcanus
Pristionchus_entomophagus
Pristionchus_exspectatus
Pristionchus_fissidentatus
Pristionchus_japonicus
Pristionchus_maxplancki
Pristionchus_pacificus

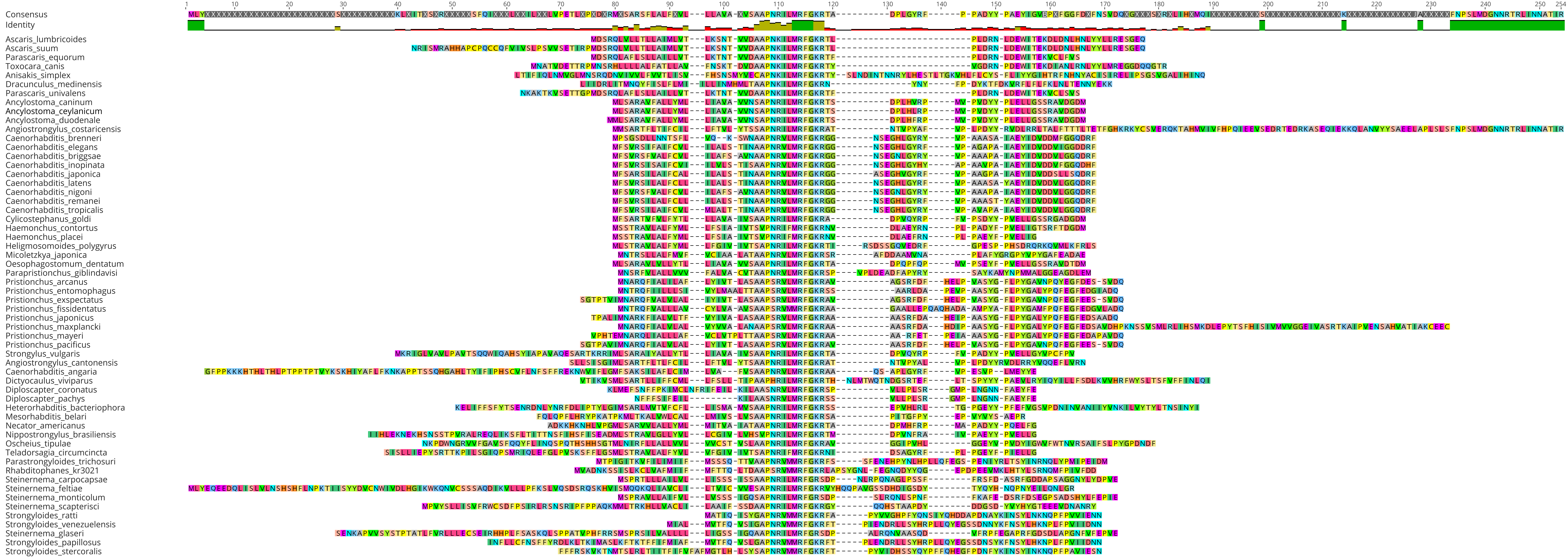

Consensus
Identity
Ascaris_lumbricoides
Ascaris_suum
Parascaris_equorum
Toxocara_canis
Anisakis_simplex
Dracunculus_medinensis
Parascaris_univalens
Ancylostoma_caninum
Ancylostoma_ceylanicum
Ancylostoma_duodenale
Angiostrongylus_costaricensis
Caenorhabditis_brenneri
Caenorhabditis_elegans
Caenorhabditis_briggsae
Caenorhabditis_inopinata
Caenorhabditis_japonica
Caenorhabditis_latens
Caenorhabditis_nigoni
Caenorhabditis_remanei
Caenorhabditis_tropicalis
Cylicostephanus_goldi
Haemonchus_contortus
Haemonchus_placei
Heligmosomoides_polygyrus
Micoletzkya_japonica
Oesophagostomum_dentatum
Parapristionchus_giblindavisi
Pristionchus_arcanus
Pristionchus_entomophagus
Pristionchus_exspectatus
Pristionchus_fissidentatus
Pristionchus_japonicus
Pristionchus_maxplancki
Pristionchus_mayeri
Pristionchus_pacificus
Strongylus_vulgaris
Angiostrongylus_cantonensis
Caenorhabditis_angaria
Dictyocaulus_viviparus
Diploscapter_coronatus
Diploscapter_pachys
Heterorhabditis_bacteriophora
Mesorhabditis_belari
Necator_americanus
Nippostrongylus_brasiliensis
Oscheius_tipulae
Teladorsagia_circumcincta
Parastrongyloides_trichosuri
Rhabditophanes_kr3021
Steinernema_carpocapsae
Steinernema_feltiae
Steinernema_monticolum
Steinernema_scapterisci
Strongyloides_ratti
Strongyloides_venezuelensis
Steinernema_glaseri
Strongyloides_papillosus
Strongyloides_stercoralis

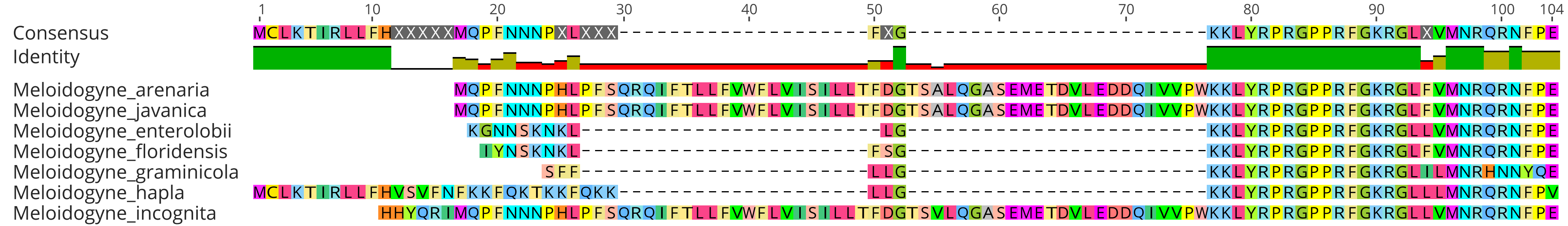
1
10
20
30
40
50
60
70
80
90
100
104
Consensus
Identity
Meloidogyne_arenaria
Meloidogyne_javanica
Meloidogyne_enterolobii
Meloidogyne_floridensis
Meloidogyne_graminicola
Meloidogyne_hapla
Meloidogyne_incognita

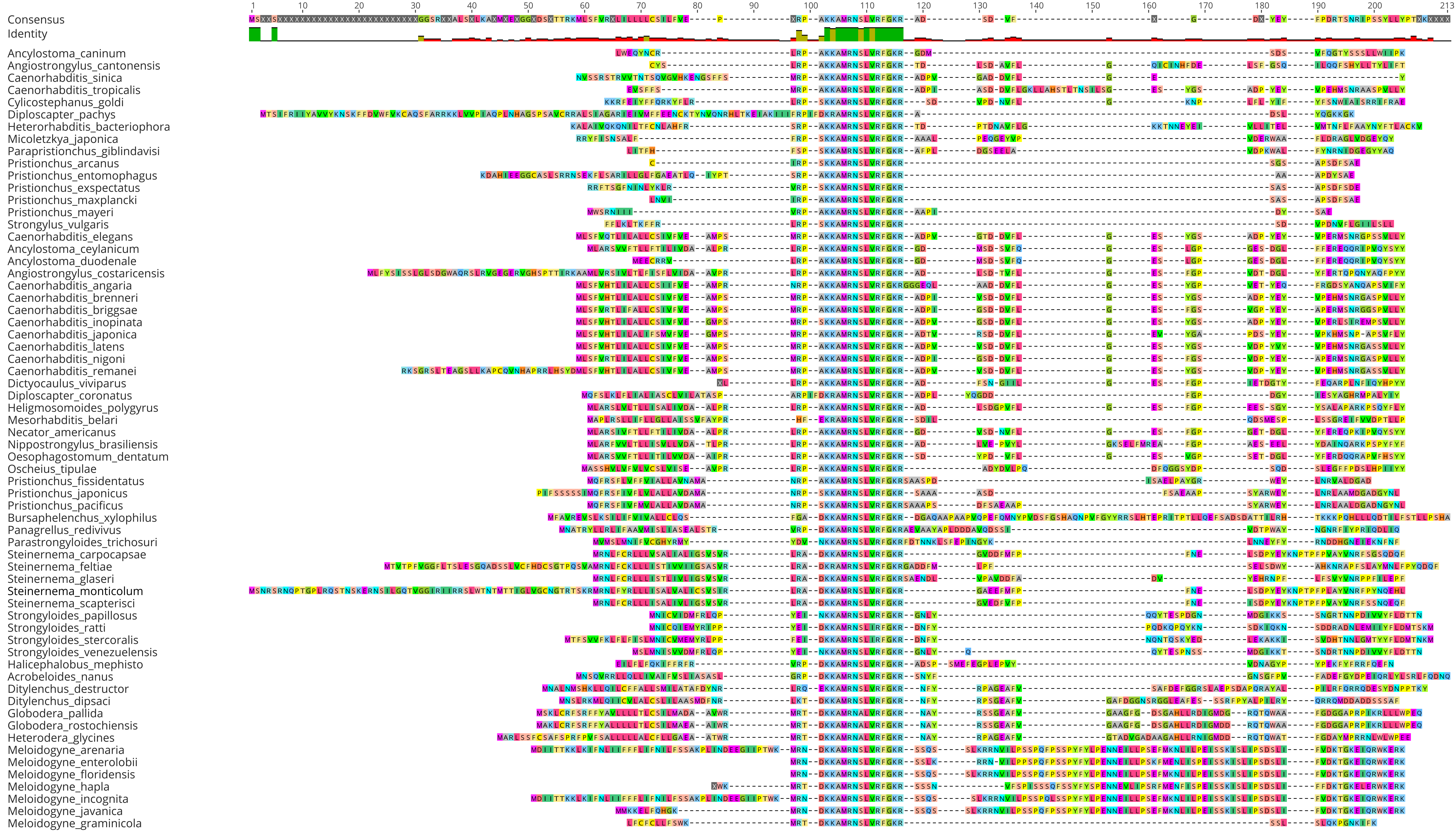

Consensus
Identity
1
10
20
30
40
50
60
70
80
90
100
110
120
130
140
150
160
170
180
190
200
213
Ancylostoma_caninum
Angiostrongylus_cantonensis
Caenorhabditis_sinica
Caenorhabditis_tropicalis
Cylicostephanus_goldi
Diploscapter_pachys
Heterorhabditis_bacteriophora
Micoletzkya_japonica
Parapristionchus_giblindavisi
Pristionchus_arcanus
Pristionchus_entomophagus
Pristionchus_exspectatus
Pristionchus_maxplancki
Pristionchus_mayeri
Strongylus_vulgaris
Caenorhabditis_elegans
Ancylostoma_ceylanicum
Ancylostoma_duodenale
Angiostrongylus_costaricensis
Caenorhabditis_angaria
Caenorhabditis_brenneri
Caenorhabditis_briggsae
Caenorhabditis_inopinata
Caenorhabditis_japonica
Caenorhabditis_latens
Caenorhabditis_nigoni
Caenorhabditis_remanei
Dictyocaulus_viviparus
Diploscapter_coronatus
Heligmosomoides_polygyrus
Mesorhabditis_belari
Necator_americanus
Nippostrongylus_brasiliensis
Oesophagostomum_dentatum
Oscheius_tipulae
Pristionchus_fissidentatus
Pristionchus_japonicus
Pristionchus_pacificus
Bursaphelenchus_xylophilus
Panagrellus_redivivus
Parastrongyloides_trichosuri
Steinernema_carpocapsae
Steinernema_feltiae
Steinernema_glaseri
Steinernema_monticolum
Steinernema_scapterisci
Strongyloides_papillosus
Strongyloides_ratti
Strongyloides_stercoralis
Strongyloides_venezuelensis
Halicephalobus_mephisto
Acrobeloides_nanus
Ditylenchus_destructor
Ditylenchus_dipsaci
Globodera_pallida
Globodera_rostochiensis
Heterodera_glycines
Meloidogyne_arenaria
Meloidogyne_enterolobii
Meloidogyne_floridensis
Meloidogyne_hapla
Meloidogyne_incognita
Meloidogyne_javanica
Meloidogyne_graminicola

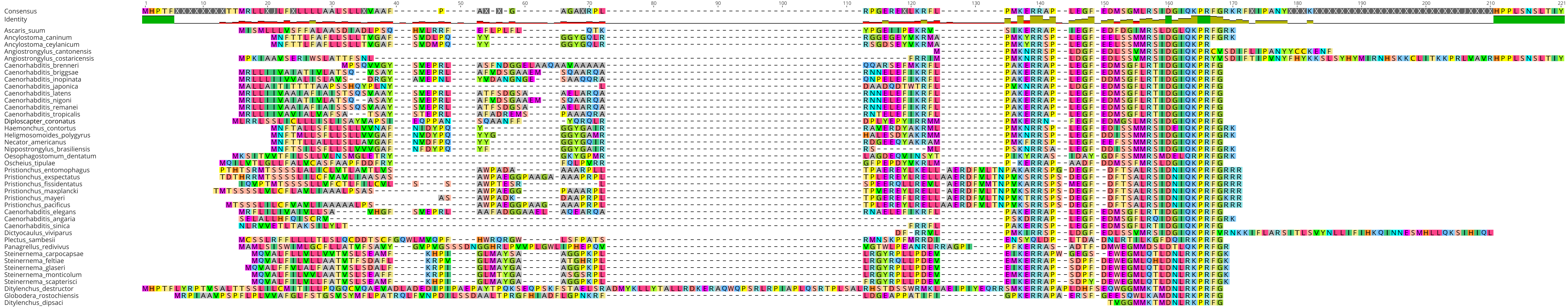
1
10
20
30
40
50
60
70
80
90
100
110
120
130
140
150
160
170
180
190
200
210
Consensus
Identity
Ascaris_suum
Ancylostoma_caninum
Ancylostoma_ceylanicum
Angiostrongylus_cantonensis
Angiostrongylus_costaricensis
Caenorhabditis_brenneri
Caenorhabditis_briggsae
Caenorhabditis_inopinata
Caenorhabditis_japonica
Caenorhabditis_latens
Caenorhabditis_nigoni
Caenorhabditis_remanei
Caenorhabditis_tropicalis
Diploscapter_coronatus
Haemonchus_contortus
Heligmosomoides_polygyrus
Necator_americanus
Nippostrongylus_brasiliensis
Oesophagostomum_dentatum
Oscheius_tipulae
Pristionchus_entomophagus
Pristionchus_exspectatus
Pristionchus_fissidentatus
Pristionchus_maxplancki
Pristionchus_mayeri
Pristionchus_pacificus
Caenorhabditis_elegans
Caenorhabditis_angaria
Caenorhabditis_sinica
Dictyocaulus_viviparus
Plectus_sambesii
Panagrellus_redivivus
Steinernema_carpocapsae
Steinernema_feltiae
Steinernema_glaseri
Steinernema_monticolum
Steinernema_scapterisci
Ditylenchus_destructor
Globodera_rostochiensis
Ditylenchus_dipsaci

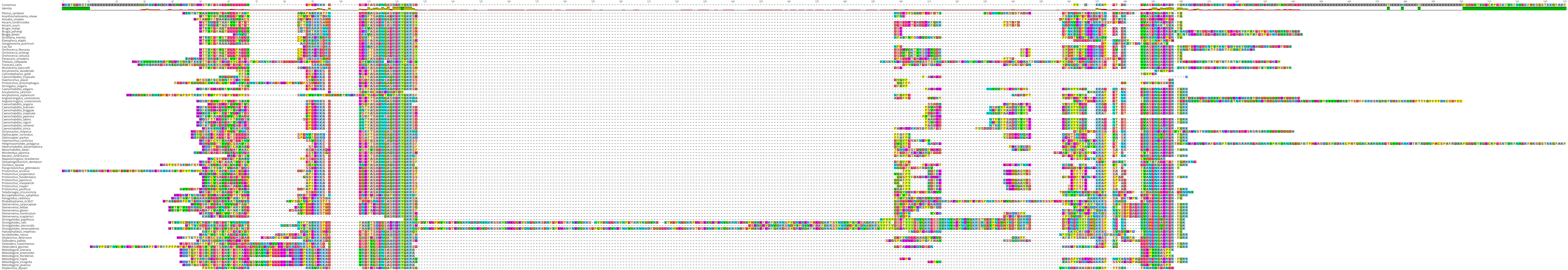
Consensus
Identity
Plectus_sambesii
Acanthocheilonema_viteae
Anisakis_simplex
Ascaris_lumbricoides
Ascaris_suum
Brugia_malayi
Brugia_pahangi
Brugia_timori
Dirofilaria_immitis
Elaeophora_elaphi
Gongylonema_pulchrum
Loa_loa
Onchocerca_flexuosa
Onchocerca_ochengi
Onchocerca_volvulus
Parascaris_univalens
Thelazia_callipaeda
Toxocara_canis
Wuchereria_bancrofti
Ancylostoma_duodenale
Cylicostephanus_goldi
Caenorhabditis_tropicalis
Haemonchus_placei
Pristionchus_entomophagus
Strongylus_vulgaris
Caenorhabditis_elegans
Ancylostoma_caninum
Ancylostoma_ceylanicum
Angiostrongylus_cantonensis
Angiostrongylus_costaricensis
Caenorhabditis_angaria
Caenorhabditis_brenneri
Caenorhabditis_briggsae
Caenorhabditis_inopinata
Caenorhabditis_japonica
Caenorhabditis_latens
Caenorhabditis_nigoni
Caenorhabditis_remanei
Caenorhabditis_sinica
Dictyocaulus_viviparus
Diploscapter_coronatus
Diploscapter_pachys
Haemonchus_contortus
Heligmosomoides_polygyrus
Heterorhabditis_bacteriophora
Mesorhabditis_belari
Micoletzkya_japonica
Necator_americanus
Nippostrongylus_brasiliensis
Oesophagostomum_dentatum
Oscheius_tipulae
Parapristionchus_giblindavisi
Pristionchus_arcanus
Pristionchus_exspectatus
Pristionchus_fissidentatus
Pristionchus_japonicus
Pristionchus_maxplancki
Pristionchus_mayeri
Pristionchus_pacificus
Teladorsagia_circumcincta
Bursaphelenchus_xylophilus
Panagrellus_redivivus
Rhabditophanes_kr3021
Steinernema_carpocapsae
Steinernema_feltiae
Steinernema_glaseri
Steinernema_monticolum
Steinernema_scapterisci
Strongyloides_papillosus
Strongyloides_ratti
Strongyloides_stercoralis
Strongyloides_venezuelensis
Halicephalobus_mephisto
Acrobeloides_nanus
Ditylenchus_destructor
Globodera_pallida
Globodera_rostochiensis
Heterodera_glycines
Meloidogyne_arenaria
Meloidogyne_enterolobii
Meloidogyne_floridensis
Meloidogyne_hapla
Meloidogyne_incognita
Meloidogyne_javanica
Ditylenchus_dipsaci
