## Supplementary Figure 1 for "Exploitation of phylum-spanning omics resources reveals complexity in the nematode FLP signalling system and provides insights into *flp*-gene evolution"

### FLP-1 peptide alignment

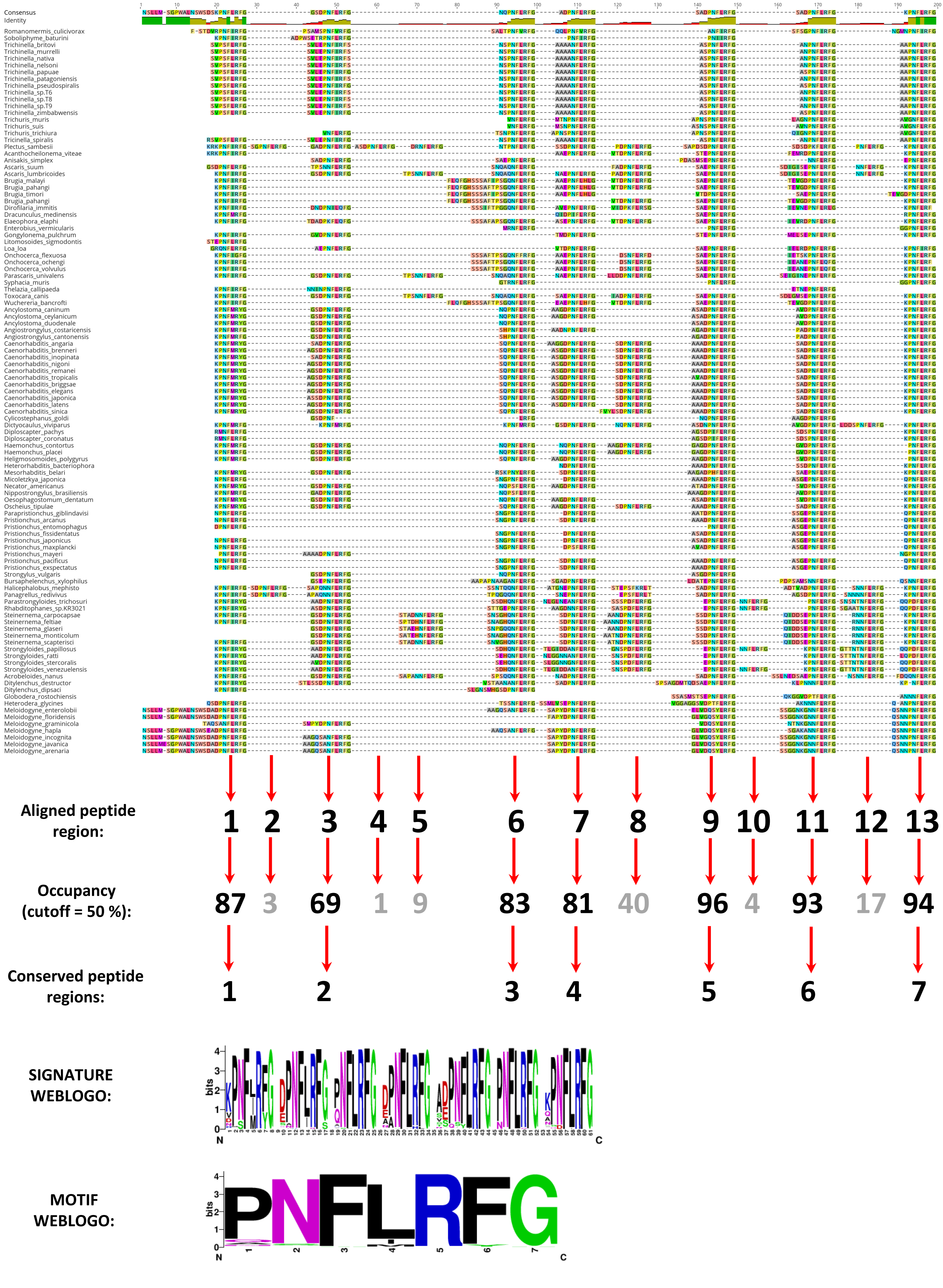

### FLP-2 peptide alignment

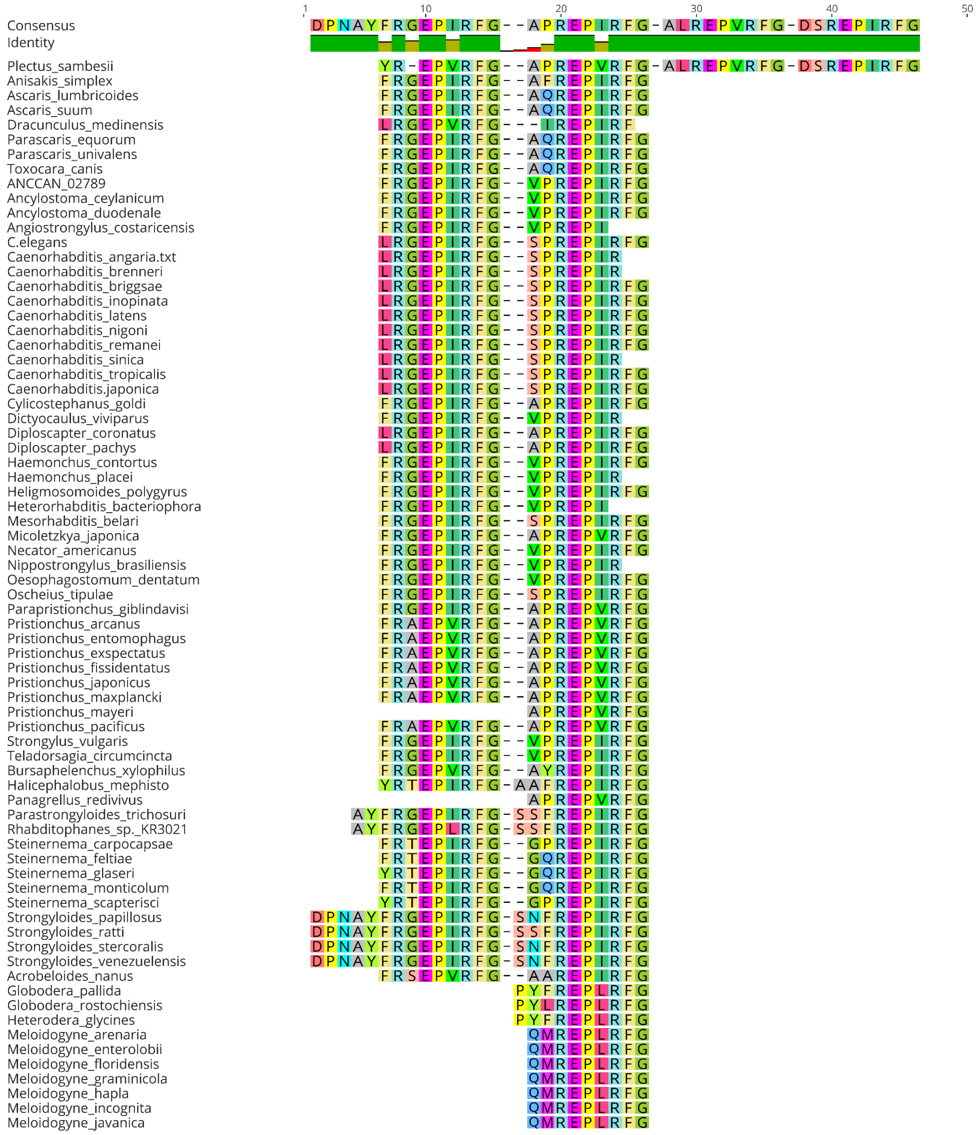


| | | | | |
|---|---|---|---|---|
| **Aligned peptide region:** | **1** | **2** | **3** | **4** |
| **Occupancy (cutoff = 50 %):** | **84** | **100** | **1** | **1** |
| **Conserved peptide regions:** | **1** | **2** | | |

**SIGNATURE WEBLOGO:**

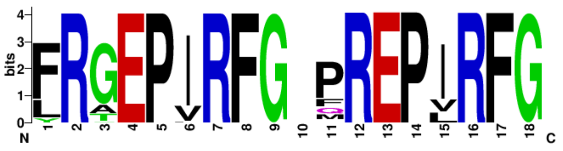

**MOTIF WEBLOGO:**

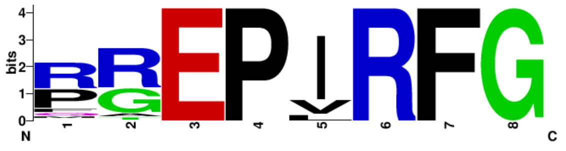

### FLP-3 peptide alignment

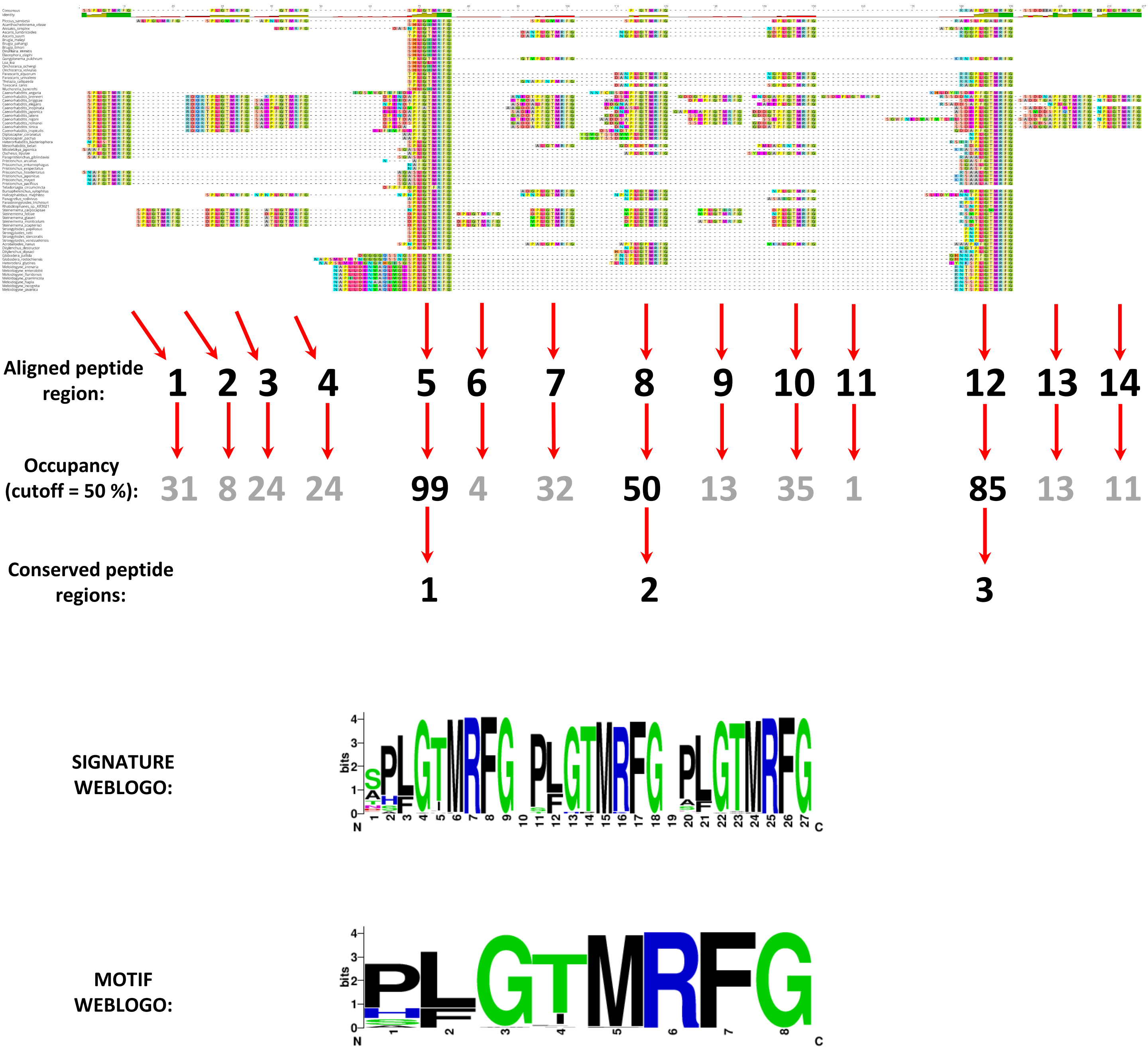

### FLP-4 peptide alignment

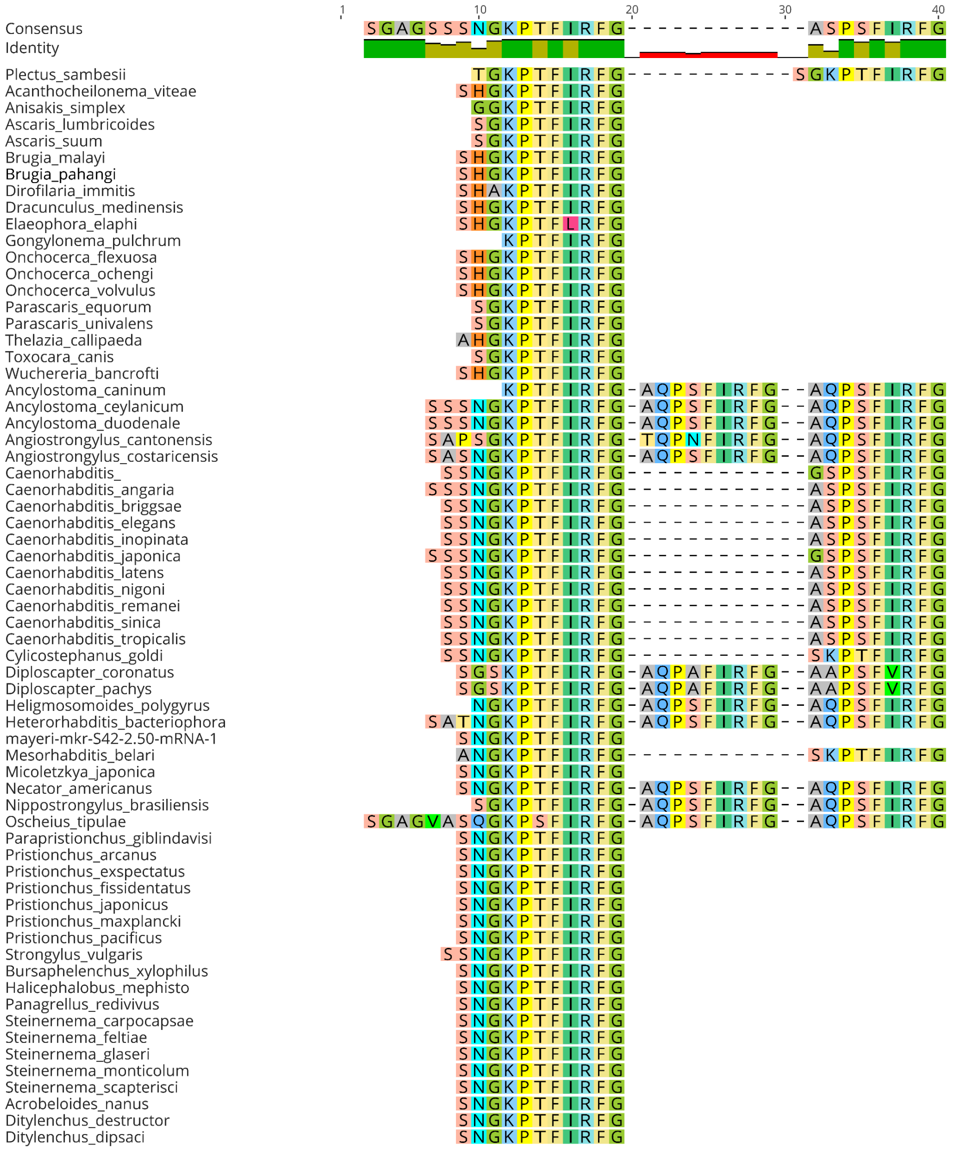


| | | | |
|---|---|---|---|
| **Aligned peptide region:** | **1** | **2** | **3** |
| **Occupancy (cutoff = 50 %):** | **84** | **18** | **40** |
| **Conserved peptide regions:** | **1** | | |

**SIGNATURE/MOTIF WEBLOGO:**

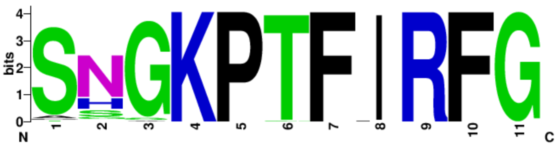

### FLP-5 peptide alignment

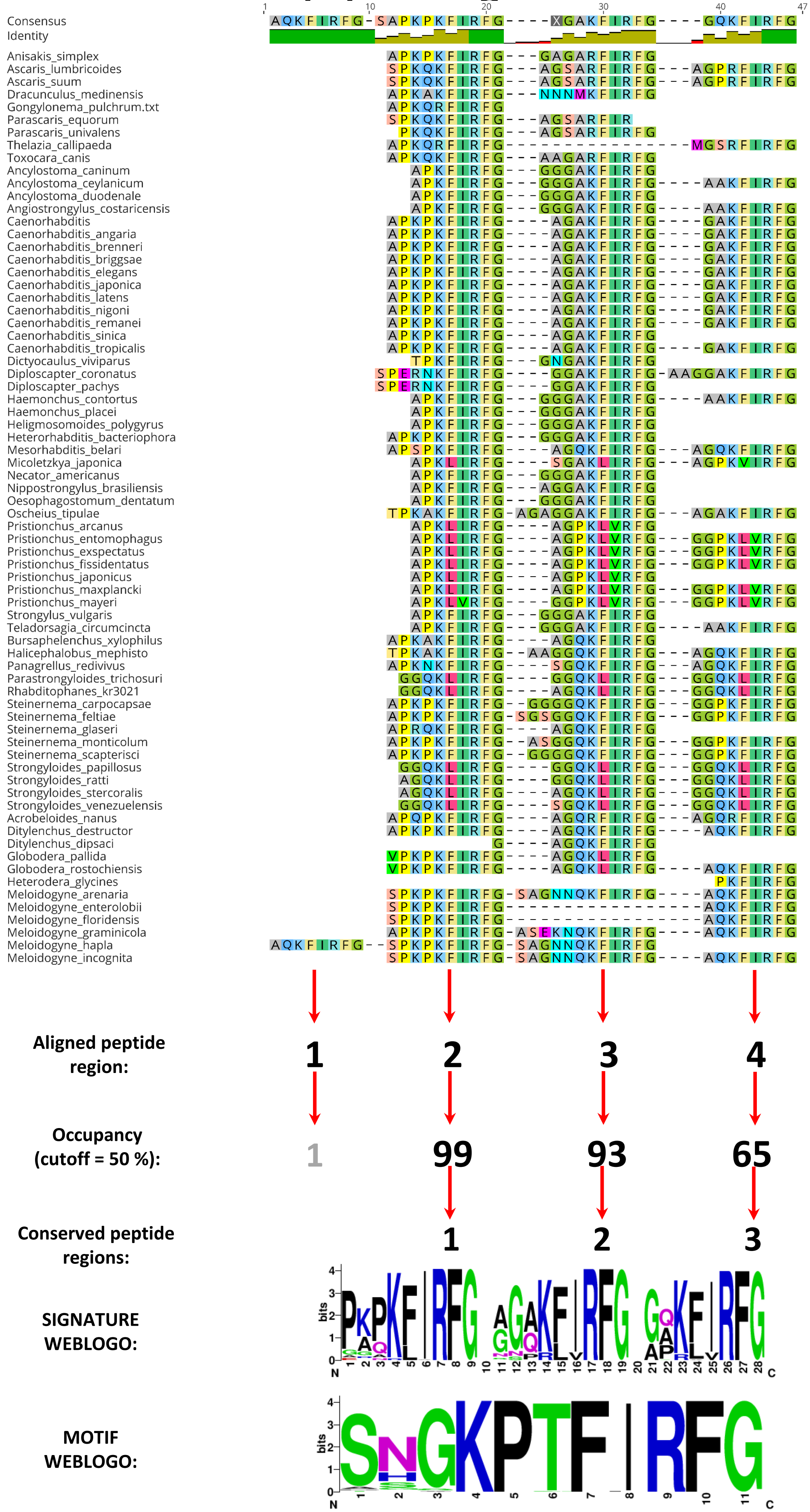

### FLP-6 peptide alignment

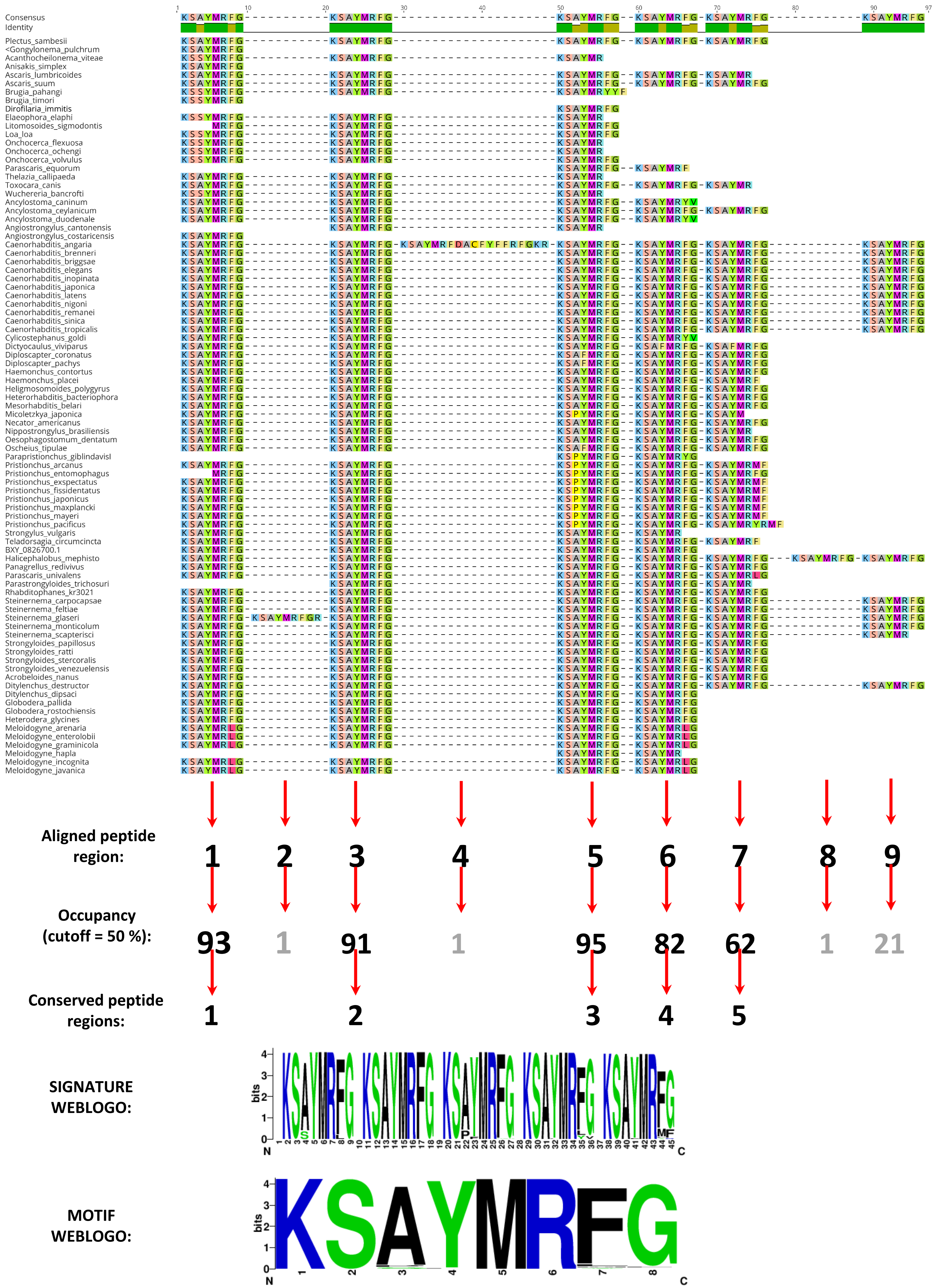

### FLP-7 peptide alignment

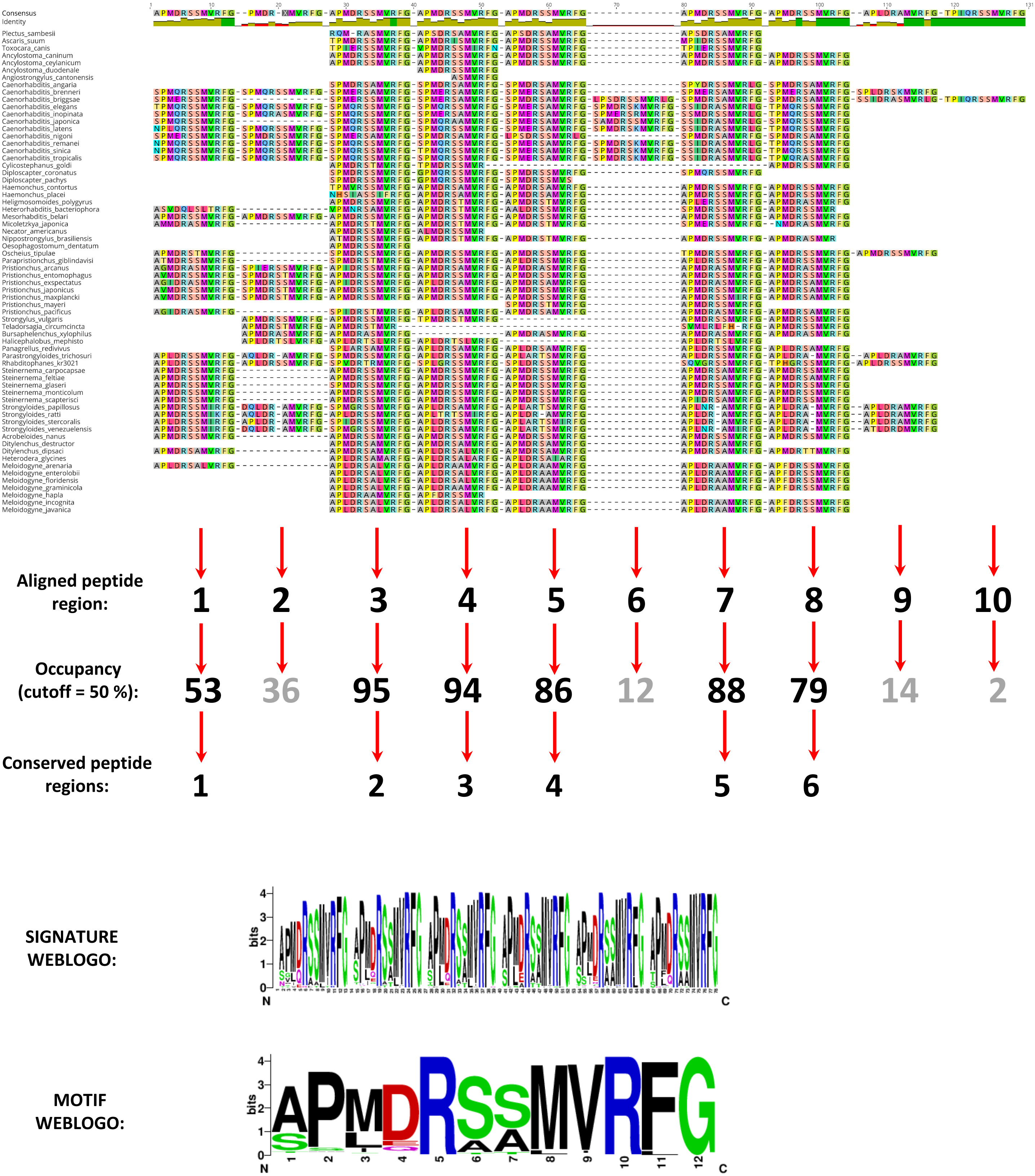

### FLP-8 peptide alignment

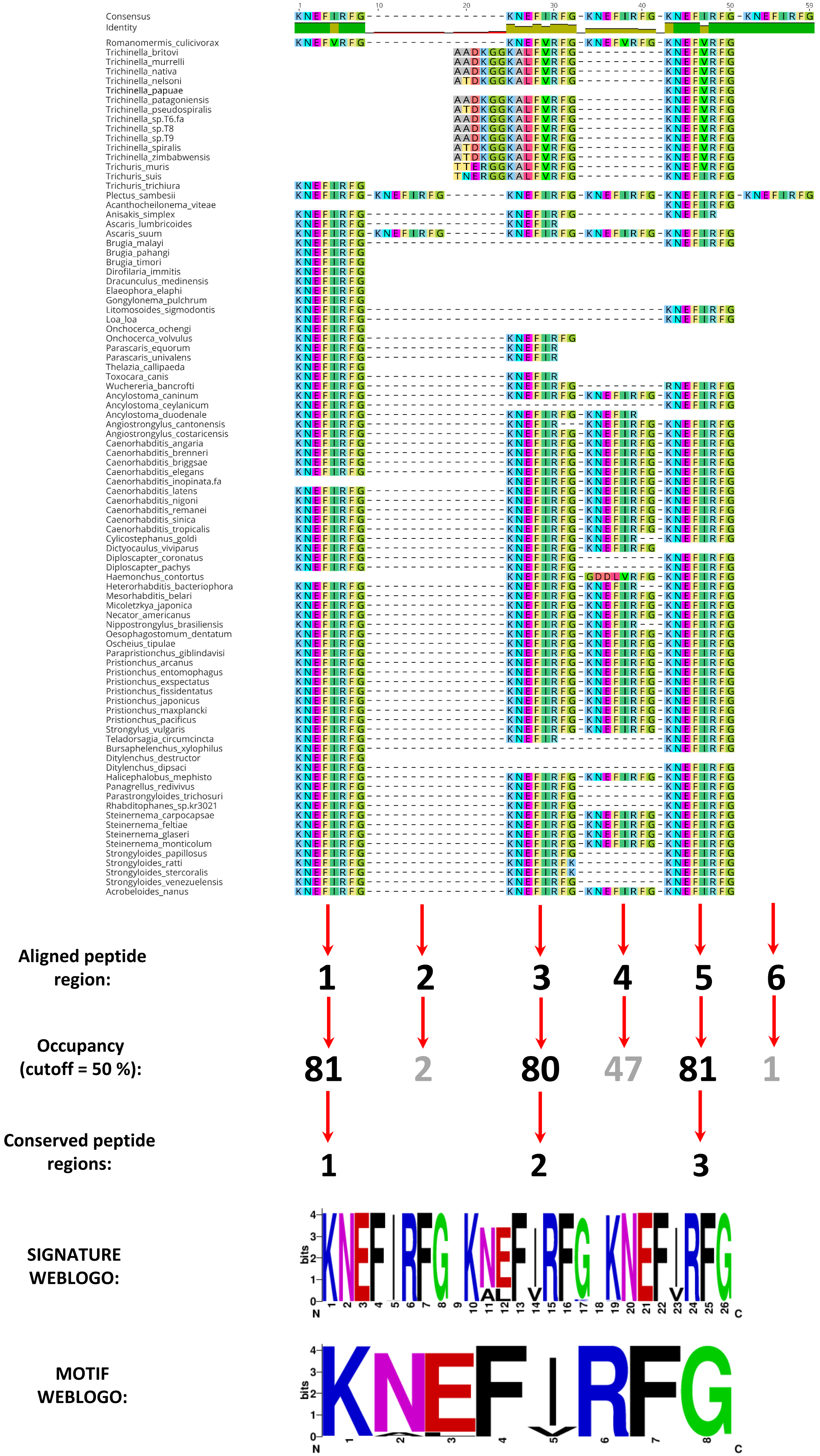

### FLP-9 peptide alignment

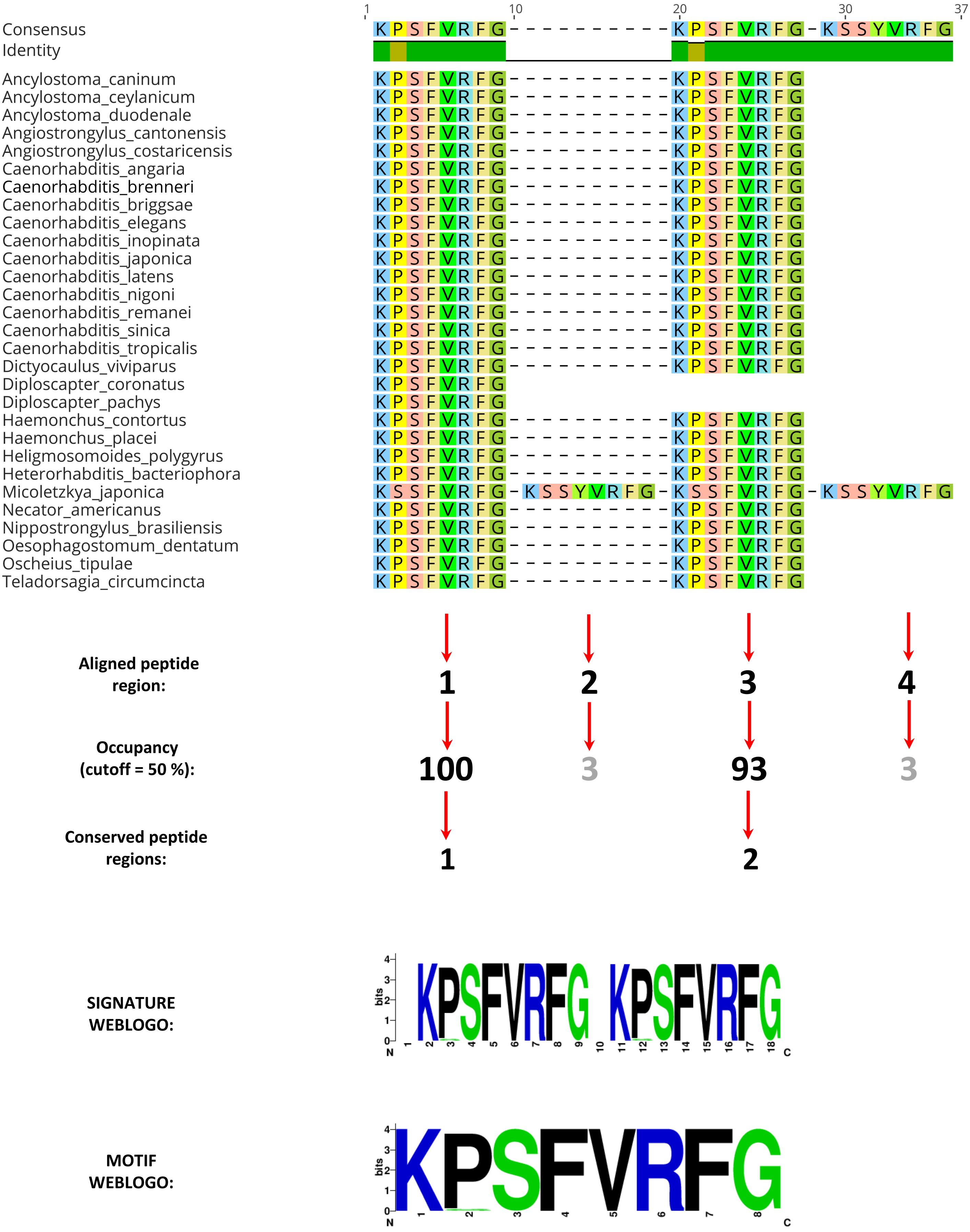

### FLP-10 peptide alignment

| | 1 | 10 | 16 |
|---|---|---|---|
| Consensus | QAASQTKSRSGYIRFG | | |
| Identity | | | |

| Species | Sequence |
|---|---|
| Caenorhabditis_angaria | QGKQRSGYIRFG |
| Caenorhabditis_brenneri | QTKSRSGYIRFG |
| Caenorhabditis_briggsae | QTKSRSGYIRFG |
| Caenorhabditis_elegans | QPKARSGYIRFG |
| Caenorhabditis_inopinata | QTKARSGYIRFG |
| Caenorhabditis_japonica | QTKARSGYIRFG |
| Caenorhabditis_latens | QTKSRSGYIRFG |
| Caenorhabditis_nigoni | QTKSRSGYIRFG |
| Caenorhabditis_remanei | QTKSRSGYIRFG |
| Caenorhabditis_sinica | QTKSRSGYIRFG |
| Caenorhabditis_tropicalis | QTKSRSGYIRFG |
| Diploscapter_coronatus | QAASANGMRSGYIRFG |
| Diploscapter_pachys | QAASANGMRSGYIRFG |

**Aligned peptide region:** 1

**Occupancy (cutoff = 50 %):** 100

**Conserved peptide regions:** 1

**SIGNATURE/MOTIF WEBLOGO:**

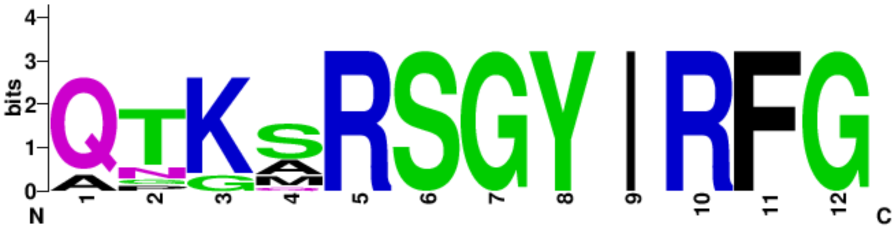

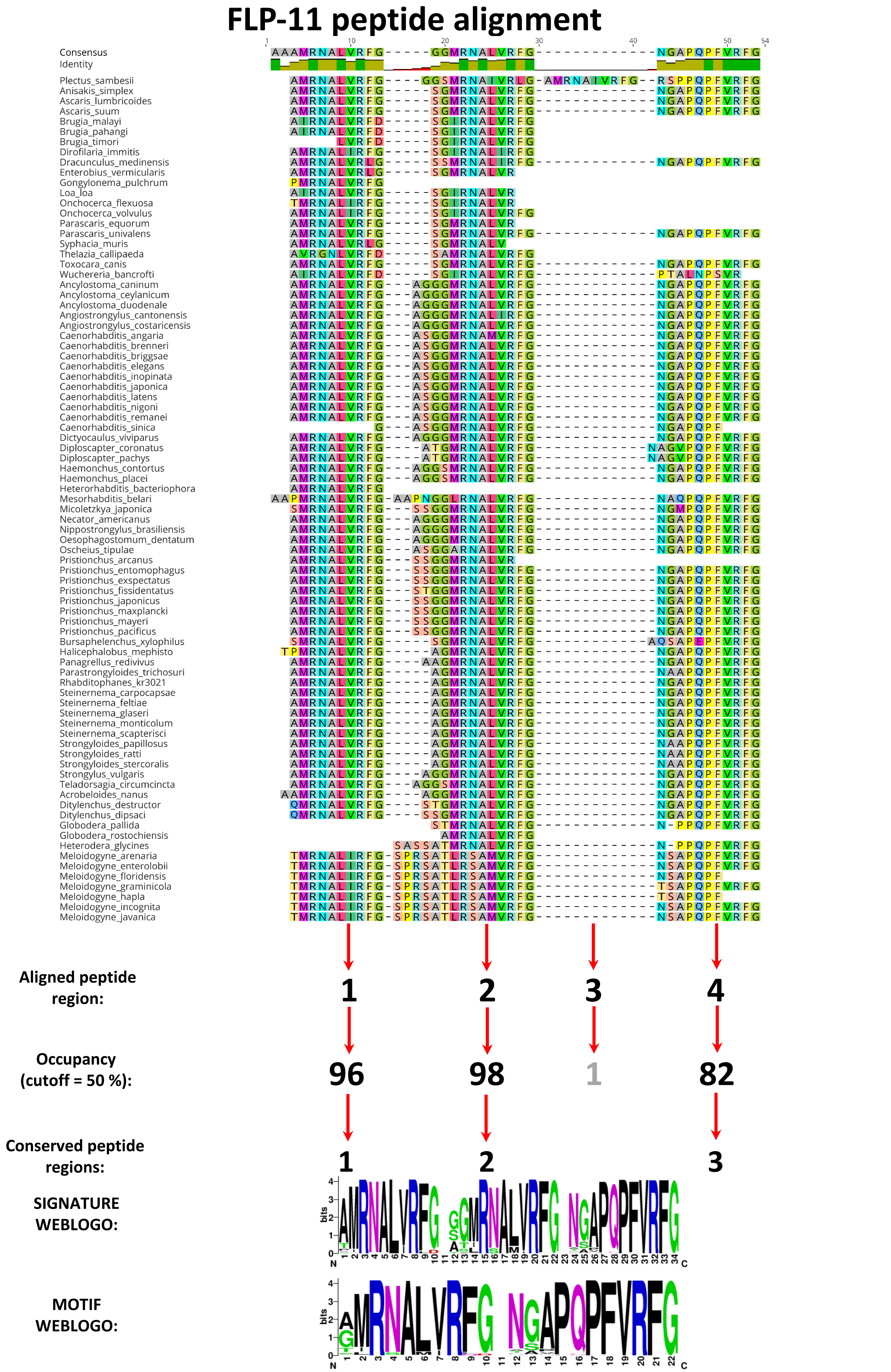

FLP-11 peptide alignment
Consensus
Identity
Plectus_sambesii
Anisakis_simplex
Ascaris_lumbricoides
Ascaris_suum
Brugia_malayi
Brugia_pahangi
Brugia_timori
Dirofilaria_immitis
Dracunculus_medinensis
Enterobius_vermicularis
Gongylonema_pulchrum
Loa_loa
Onchocerca_flexuosa
Onchocerca_volvulus
Parascaris_equorum
Parascaris_univalens
Syphacia_muris
Thelazia_callipaeda
Toxocara_canis
Wuchereria_bancrofti
Ancylostoma_caninum
Ancylostoma_ceylanicum
Ancylostoma_duodenale
Angiostrongylus_cantonensis
Angiostrongylus_costaricensis
Caenorhabditis_angaria
Caenorhabditis_brenneri
Caenorhabditis_briggsae
Caenorhabditis_elegans
Caenorhabditis_inopinata
Caenorhabditis_japonica
Caenorhabditis_latens
Caenorhabditis_nigoni
Caenorhabditis_remanei
Caenorhabditis_sinica
Dictyocaulus_viviparus
Diploscapter_coronatus
Diploscapter_pachys
Haemonchus_contortus
Haemonchus_placei
Heterorhabditis_bacteriophora
Mesorhabditis_belari
Micoletzkya_japonica
Necator_americanus
Nippostrongylus_brasiliensis
Oesophagostomum_dentatum
Oscheius_tipulae
Pristionchus_arcanus
Pristionchus_entomophagus
Pristionchus_exspectatus
Pristionchus_fissidentatus
Pristionchus_japonicus
Pristionchus_maxplancki
Pristionchus_mayeri
Pristionchus_pacificus
Bursaphelenchus_xylophilus
Halicephalobus_mephisto
Panagrellus_redivivus
Parastrongyloides_trichosuri
Rhabditophanes_kr3021
Steinernema_carpocapsae
Steinernema_feltiae
Steinernema_glaseri
Steinernema_monticolum
Steinernema_scapterisci
Strongyloides_papillosus
Strongyloides_ratti
Strongyloides_stercoralis
Strongylus_vulgaris
Teladorsagia_circumcincta
Acrobeloides_nanus
Ditylenchus_destructor
Ditylenchus_dipsaci
Globodera_pallida
Globodera_rostochiensis
Heterodera_glycines
Meloidogyne_arenaria
Meloidogyne_enterolobii
Meloidogyne_floridensis
Meloidogyne_graminicola
Meloidogyne_hapla
Meloidogyne_incognita
Meloidogyne_javanica
Aligned peptide region:
1
2
3
4
Occupancy (cutoff = 50 %):
96
98
1
82
Conserved peptide regions:
1
2
3
SIGNATURE WEBLOGO:
MOTIF WEBLOGO:
bits
N
C

### FLP-12 peptide alignment

| Sequence | 1–11 |
|---|---|
| Consensus | R-NKFEFIRFG |
| Identity | |
| Acanthocheilonema_viteae | R-NKFEFIRFG |
| Anisakis_simplex | R-NKFEFIRFG |
| Ascaris_lumbricoides | R-NKFEFIRFG |
| Ascaris_suum | R-NKFEFIRFG |
| Brugia_malayi | R-NKFEFIRFG |
| Brugia_pahangi | R-NKFEFIRFG |
| Brugia_timori | R-NKFEFIRFG |
| Dirofilaria_immitis.txt | R-NKFEFIRFG |
| Dracunculus_medinensis | R-NKFEFIRFG |
| Elaeophora_elaphi | R-NKFEFIRFG |
| Enterobius_vermicularis | R-NKFEFIRFG |
| Gongylonema_pulchrum | R-NKFEFIRFG |
| Litomosoides_sigmodontis | R-NKFEFIRFG |
| Loa_loa | R-NKFEFIRFG |
| Onchocerca_flexuosa | R-NKFEFIRFG |
| Onchocerca_ochengi | R-NKFEFIRFG |
| Onchocerca_volvulus | R-NKFEFIRFG |
| Parascaris_univalens | R-NKFEFIRFG |
| Syphacia_muris | R--KFEFIRFG |
| Thelazia_callipaeda | R-NKFEFIRFG |
| Toxocara_canis | R-NKFEFIRFG |
| Wuchereria_bancrofti | R-NKFEFIRFG |
| Ancylostoma_caninum | R-NKFEFIRFG |
| Ancylostoma_ceylanicum | R-NKFEFIRFG |
| Ancylostoma_duodenale | R-NKFEFIRFG |
| Angiostrongylus_costaricensis | R-NKFEFIRFG |
| Caenorhabditis_angaria | R-NKFEFIRFG |
| Caenorhabditis_brenneri | R-NKFEFIRFG |
| Caenorhabditis_briggsae | R-NKFEFIRFG |
| Caenorhabditis_elegans | R-NKFEFIRFG |
| Caenorhabditis_inopinata | R-NKFEFIRFG |
| Caenorhabditis_japonica | R-NKFEFIRFG |
| Caenorhabditis_latens | R-NKFEFIRFG |
| Caenorhabditis_nigoni | R-NKFEFIRFG |
| Caenorhabditis_sinica | R-NKFEFIRFG |
| Caenorhabditis_tropicalis | R-NKFEFIRFG |
| Dictyocaulus_viviparus | R-NKFEFIRFG |
| Diploscapter_coronatus | R-NKFEFIRFG |
| Diploscapter_pachys | R-NKFEFIRFG |
| Haemonchus_contortus | R-NKFEFIRFG |
| Haemonchus_placei | R-NKFEFIRFG |
| Heligmosomoides_polygyrus | R-NKFEFIRFG |
| Heterorhabditis_bacteriophora | R-NKFEFIRFG |
| Mesorhabditis_belari | R-NKFEFIRFG |
| Micoletzkya_japonica | R-NKFEFIRFG |
| Necator_americanus | R-NKFEFIRFG |
| Nippostrongylus_brasiliensis | R-NKFEFIRFG |
| Oesophagostomum_dentatum | R-NKFEFIRFG |
| Parapristionchus_giblindavisi | R-NKFEFIRFG |
| Pristionchus_arcanus | R-NKFEFIRFG |
| Pristionchus_entomophagus | R-NKFEFIRFG |
| Pristionchus_exspectatus | R-NKFEFIRFG |
| Pristionchus_fissidentatus | R-NKFEFIRFG |
| Pristionchus_japonicus | R-NKFEFIRFG |
| Pristionchus_maxplancki | R-NKFEFIRFG |
| Pristionchus_mayeri | R-NKFEFIRFG |
| Pristionchus_pacificus | R-NKFEFIRFG |
| Teladorsagia_circumcincta | R-NKFEFIRFG |
| Bursaphelenchus_xylophilus | R-NKFEFIRFG |
| Halicephalobus_mephisto | R-NKFEFIRFG |
| Panagrellus_redivivus | R-NKFEFIRFG |
| Parastrongyloides_trichosuri | R-NKFEFIRFG |
| Rhabditophanes_sp.KR3021 | R-NKFEFIRFG |
| Steinernema_carpocapsae | R-NKFEFIRFG |
| Steinernema_feltiae | R-NKFEFIRFG |
| Steinernema_glaseri | R-NKFEFIRFG |
| Steinernema_monticolum | R-NKFEFIRFG |
| Steinernema_scapterisci | R-NKFEFIRFG |
| Strongyloides_papillosus | R-NKFEFIRFG |
| Strongyloides_ratti | R-NKFEFIRFG |
| Strongyloides_stercoralis | R-NKFEFIRFG |
| Strongyloides_venezuelensis | R-NKFEFIRFG |
| Acrobeloides_nanus | R-NKFEFIRFG |
| Ditylenchus_destructor | R-NKFEFIRFG |
| Ditylenchus_dipsaci | R-NKFEFIRFG |
| Globodera_pallida | K-NKFEFIRFG |
| Globodera_rostochiensis | K-NKFEFIRFG |
| Heterodera_glycines | K-NKFEFIRFG |
| Meloidogyne_arenaria | KNNKFEFIRFG |
| Meloidogyne_enterolobii | KNNKFEFIRFG |
| Meloidogyne_floridensis | KNNKFEFIRFG |
| Meloidogyne_graminicola | KNNKFEFIRFG |
| Meloidogyne_hapla | KNNKFEFIRFG |
| Meloidogyne_incognita | KNNKFEFIRFG |
| Meloidogyne_javanica | KNNKFEFIRFG |

**Aligned peptide region:** 1

**Occupancy (cutoff = 50 %):** 100

**Conserved peptide regions:** 1

**SIGNATURE/MOTIF WEBLOGO:**

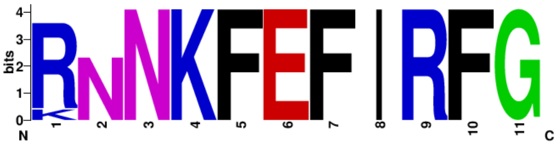

### FLP-13 peptide alignment

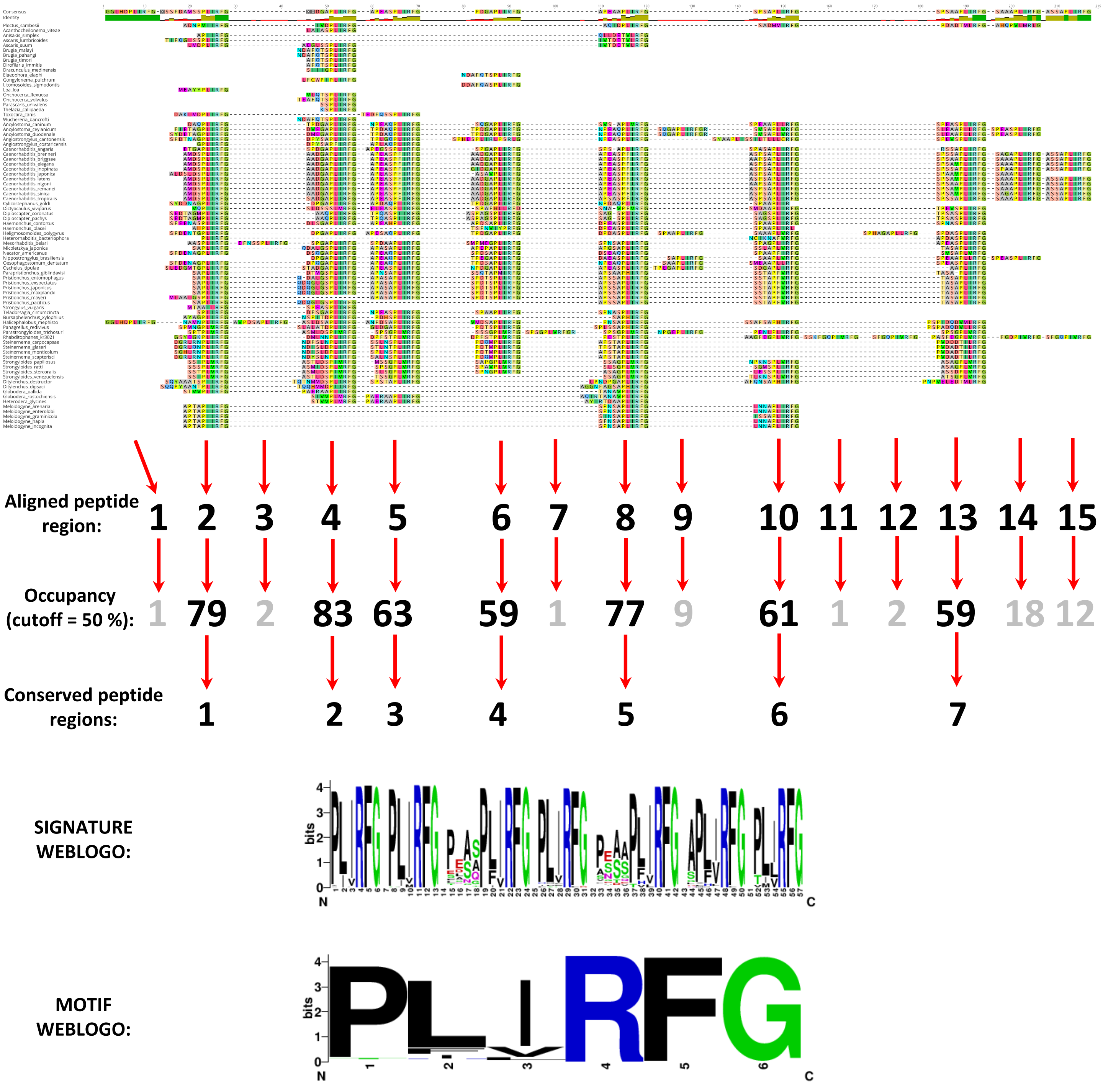

### FLP-14 peptide alignment

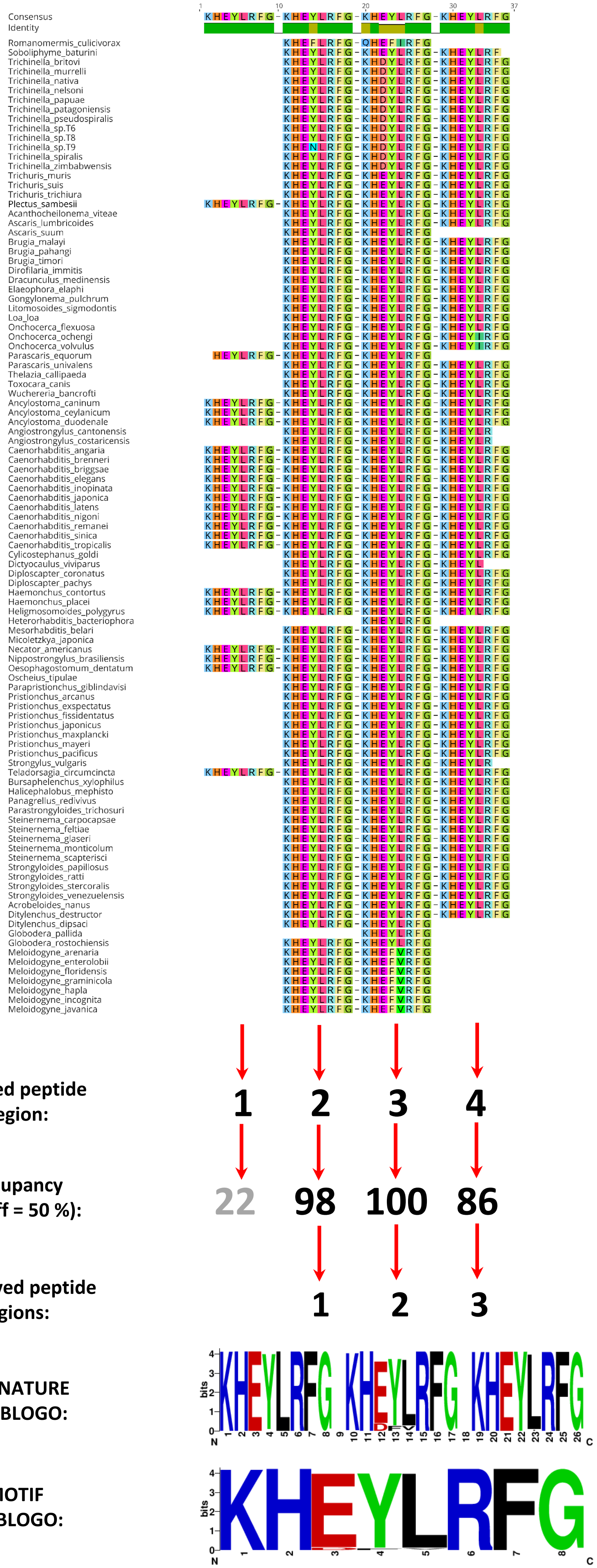

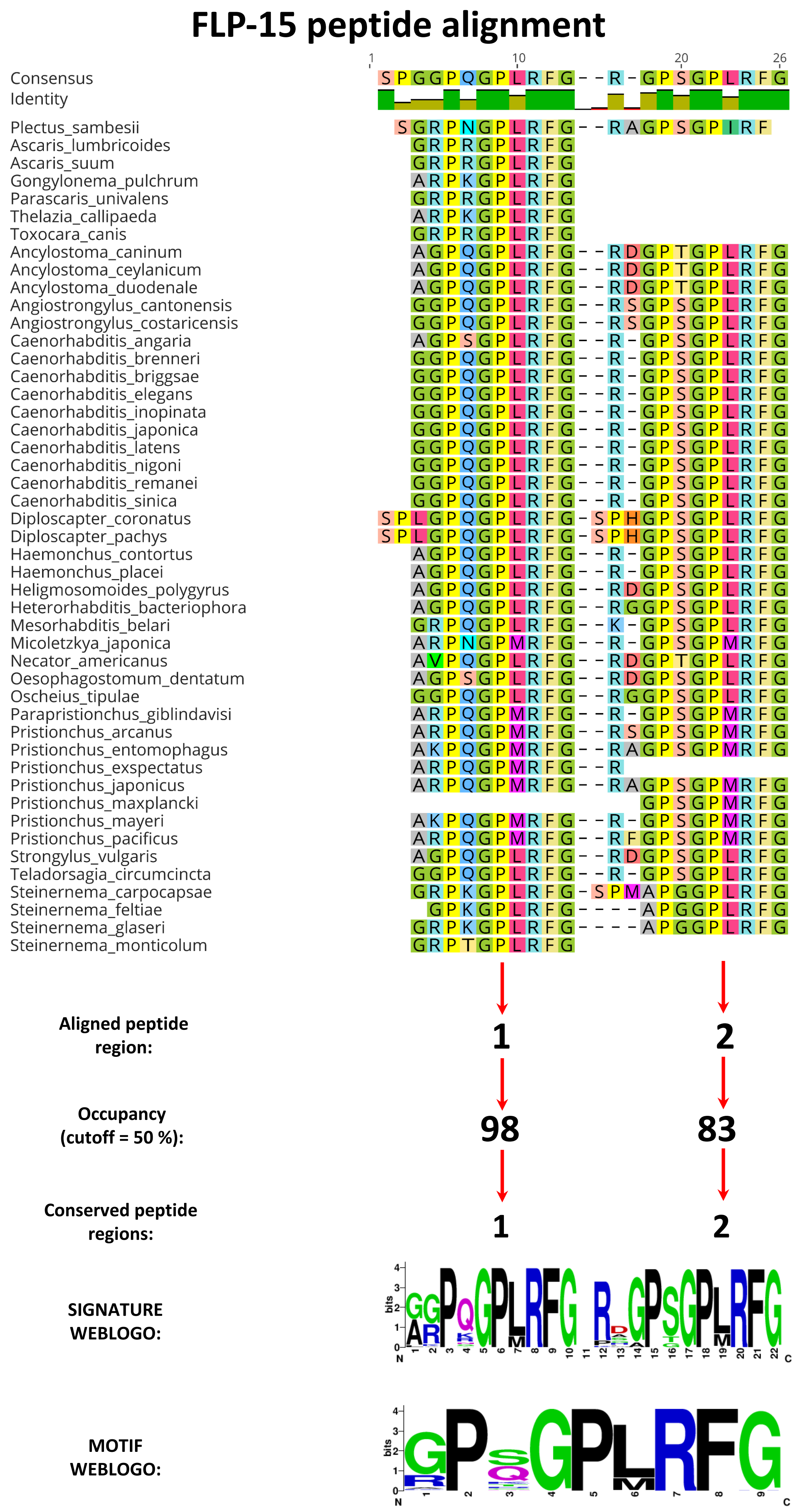
FLP-15 peptide alignment
1
10
20
26
Consensus SPGGPQGPLRFG--R-GPSGPLRFG
Identity
Plectus_sambesii SGRPNGPLRFG--RAGPSGPIRF
Ascaris_lumbricoides GRPRGPLRFG
Ascaris_suum GRPRGPLRFG
Gongylonema_pulchrum ARPKGPLRFG
Parascaris_univalens GRPRGPLRFG
Thelazia_callipaeda ARPKGPLRFG
Toxocara_canis GRPRGPLRFG
Ancylostoma_caninum AGPQGPLRFG--RDGPTGPLRFG
Ancylostoma_ceylanicum AGPQGPLRFG--RDGPTGPLRFG
Ancylostoma_duodenale AGPQGPLRFG--RDGPTGPLRFG
Angiostrongylus_cantonensis GGPQGPLRFG--RSGPSGPLRFG
Angiostrongylus_costaricensis GGPQGPLRFG--RSGPSGPLRFG
Caenorhabditis_angaria AGPSGPLRFG--R-GPSGPLRFG
Caenorhabditis_brenneri GGPQGPLRFG--R-GPSGPLRFG
Caenorhabditis_briggsae GGPQGPLRFG--R-GPSGPLRFG
Caenorhabditis_elegans GGPQGPLRFG--R-GPSGPLRFG
Caenorhabditis_inopinata GGPQGPLRFG--R-GPSGPLRFG
Caenorhabditis_japonica GGPQGPLRFG--R-GPSGPLRFG
Caenorhabditis_latens GGPQGPLRFG--R-GPSGPLRFG
Caenorhabditis_nigoni GGPQGPLRFG--R-GPSGPLRFG
Caenorhabditis_remanei GGPQGPLRFG--R-GPSGPLRFG
Caenorhabditis_sinica GGPQGPLRFG--R-GPSGPLRFG
Diploscapter_coronatus SPLGPQGPLRFG-SPHGPSGPLRFG
Diploscapter_pachys SPLGPQGPLRFG-SPHGPSGPLRFG
Haemonchus_contortus AGPQGPLRFG--R-GPSGPLRFG
Haemonchus_placei AGPQGPLRFG--R-GPSGPLRFG
Heligmosomoides_polygyrus AGPQGPLRFG--RDGPSGPLRFG
Heterorhabditis_bacteriophora AGPQGPLRFG--RGGPSGPLRFG
Mesorhabditis_belari GRPQGPLRFG--K-GPSGPLRFG
Micoletzkya_japonica ARPNGPMRFG--R-GPSGPMRFG
Necator_americanus AVPQGPLRFG--RDGPTGPLRFG
Oesophagostomum_dentatum AGPSGPLRFG--RDGPSGPLRFG
Oscheius_tipulae GGPQGPLRFG--RGGPSGPLRFG
Parapristionchus_giblindavisi ARPQGPMRFG--R-GPSGPMRFG
Pristionchus_arcanus ARPQGPMRFG--RSGPSGPMRFG
Pristionchus_entomophagus AKPQGPMRFG--RAGPSGPMRFG
Pristionchus_exspectatus ARPQGPMRFG--R
Pristionchus_japonicus ARPQGPMRFG--RAGPSGPMRFG
Pristionchus_maxplancki GPSGPMRFG
Pristionchus_mayeri AKPQGPMRFG--R-GPSGPMRFG
Pristionchus_pacificus ARPQGPMRFG--RFGPSGPMRFG
Strongylus_vulgaris AGPQGPLRFG--RDGPSGPLRFG
Teladorsagia_circumcincta GGPQGPLRFG--R-GPSGPLRFG
Steinernema_carpocapsae GRPKGPLRFG-SPMAPGGPLRFG
Steinernema_feltiae GPKGPLRFG----APGGPLRFG
Steinernema_glaseri GRPKGPLRFG----APGGPLRFG
Steinernema_monticolum GRPTGPLRFG
Aligned peptide region:
1
2
Occupancy (cutoff = 50 %):
98
83
Conserved peptide regions:
1
2
SIGNATURE WEBLOGO:
MOTIF WEBLOGO:

### FLP-16 peptide alignment

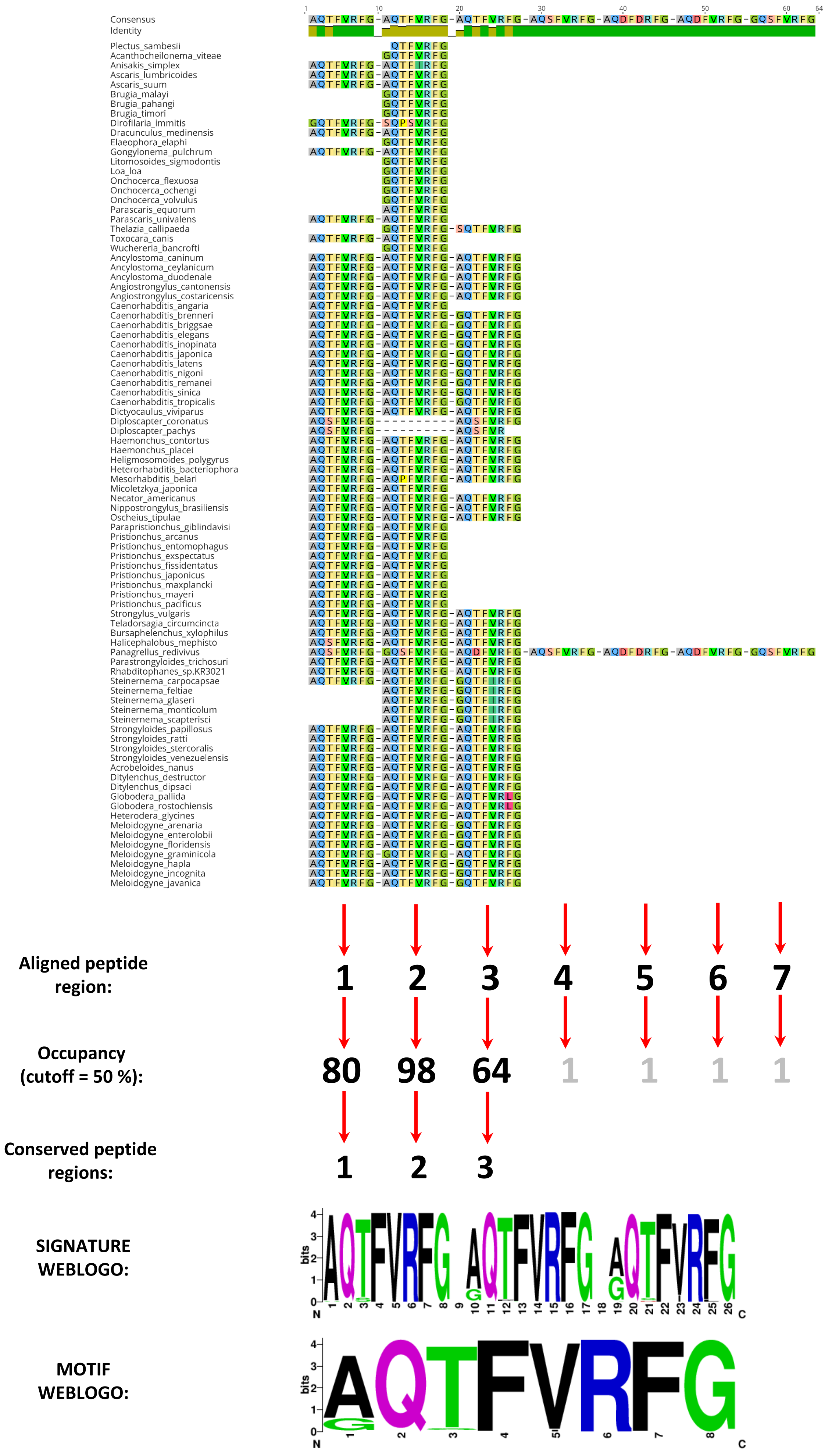

### FLP-17 peptide alignment

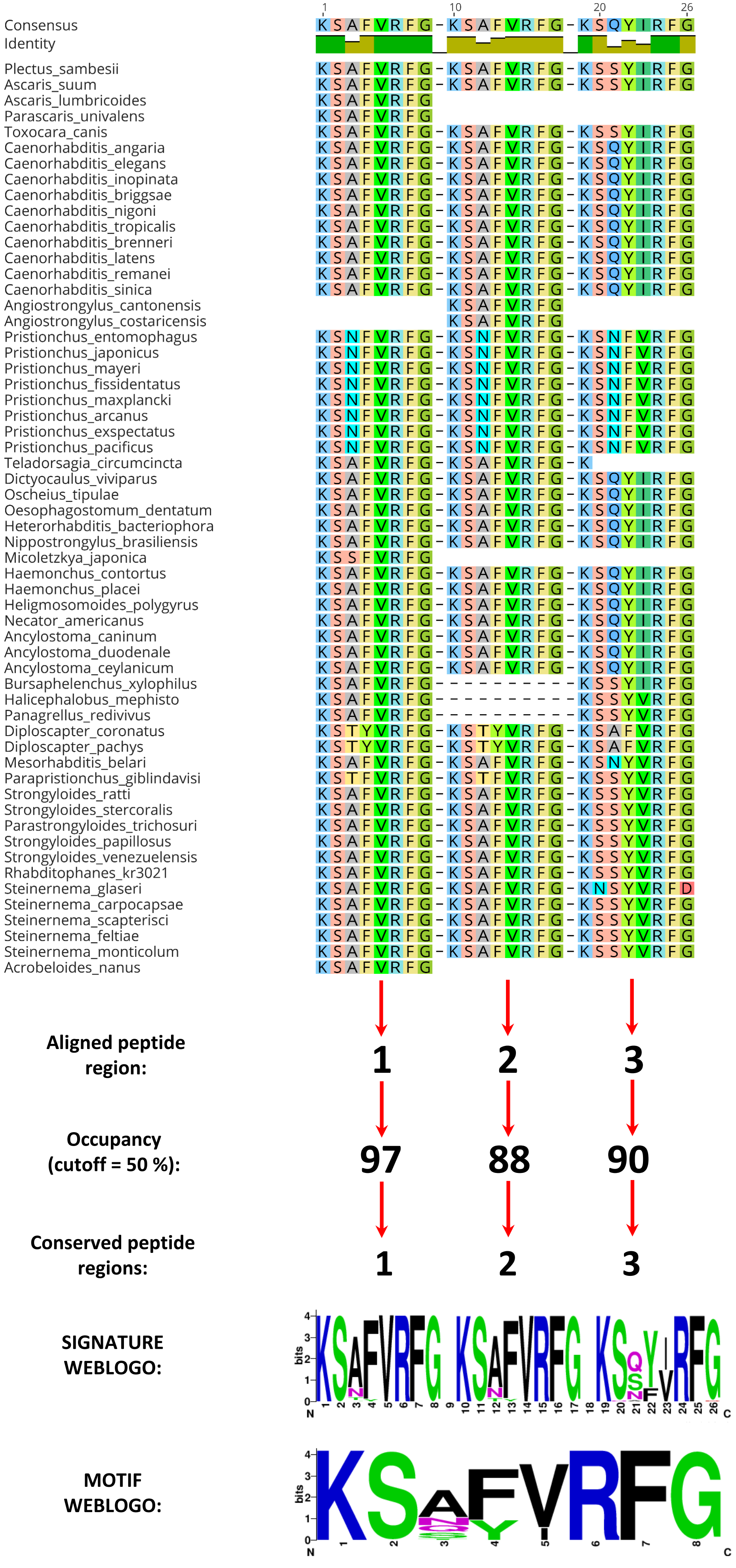

### FLP-18 peptide alignment

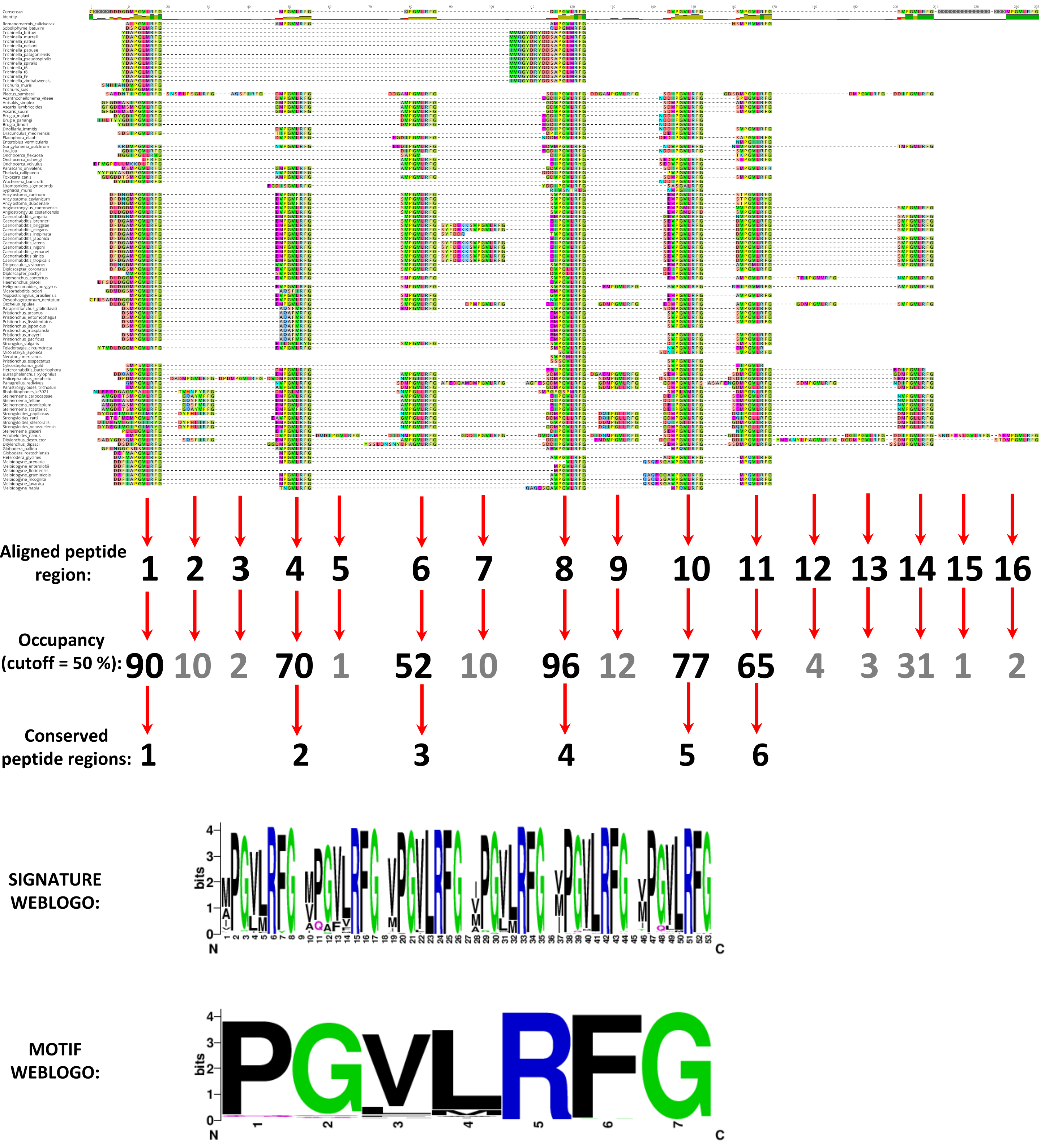

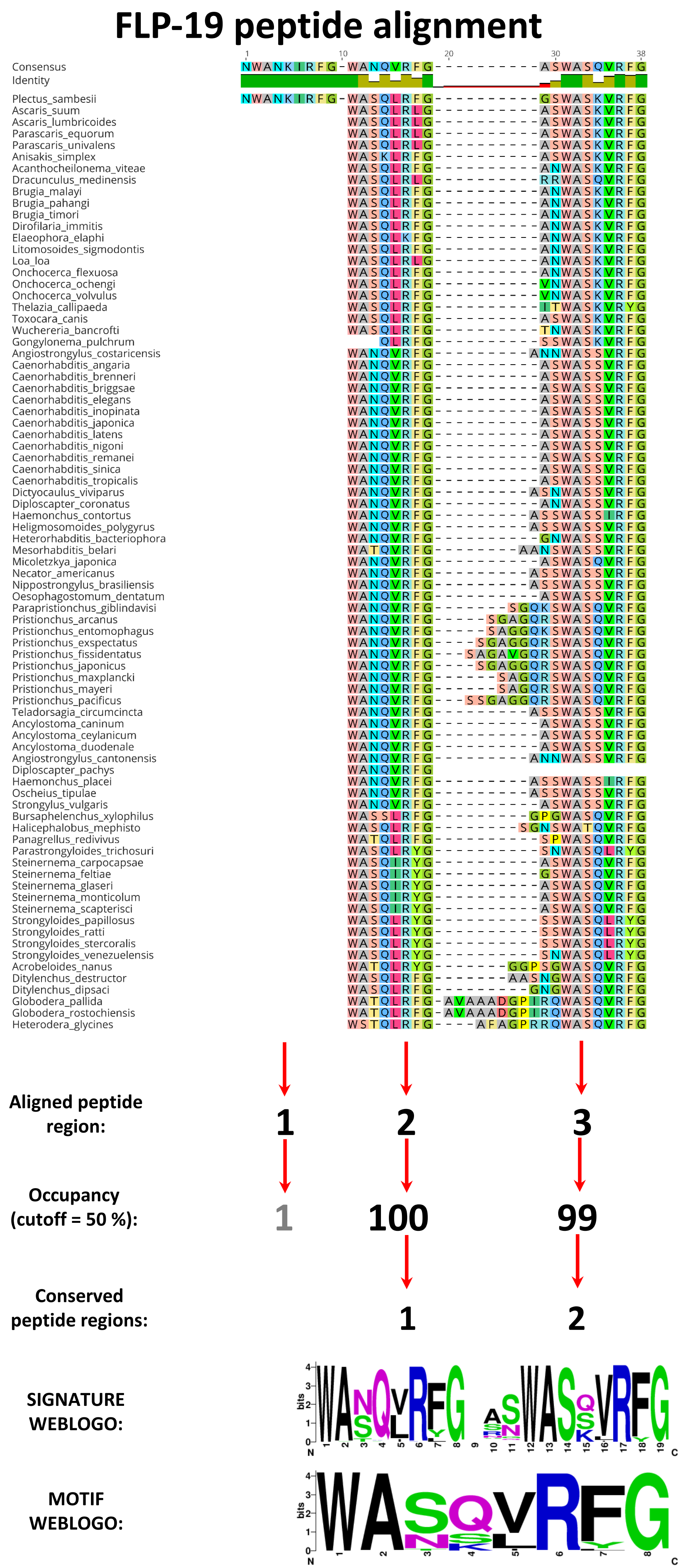

FLP-19 peptide alignment
Consensus
Identity
Plectus_sambesii
Ascaris_suum
Ascaris_lumbricoides
Parascaris_equorum
Parascaris_univalens
Anisakis_simplex
Acanthocheilonema_viteae
Dracunculus_medinensis
Brugia_malayi
Brugia_pahangi
Brugia_timori
Dirofilaria_immitis
Elaeophora_elaphi
Litomosoides_sigmodontis
Loa_loa
Onchocerca_flexuosa
Onchocerca_ochengi
Onchocerca_volvulus
Thelazia_callipaeda
Toxocara_canis
Wuchereria_bancrofti
Gongylonema_pulchrum
Angiostrongylus_costaricensis
Caenorhabditis_angaria
Caenorhabditis_brenneri
Caenorhabditis_briggsae
Caenorhabditis_elegans
Caenorhabditis_inopinata
Caenorhabditis_japonica
Caenorhabditis_latens
Caenorhabditis_nigoni
Caenorhabditis_remanei
Caenorhabditis_sinica
Caenorhabditis_tropicalis
Dictyocaulus_viviparus
Diploscapter_coronatus
Haemonchus_contortus
Heligmosomoides_polygyrus
Heterorhabditis_bacteriophora
Mesorhabditis_belari
Micoletzkya_japonica
Necator_americanus
Nippostrongylus_brasiliensis
Oesophagostomum_dentatum
Parapristionchus_giblindavisi
Pristionchus_arcanus
Pristionchus_entomophagus
Pristionchus_exspectatus
Pristionchus_fissidentatus
Pristionchus_japonicus
Pristionchus_maxplancki
Pristionchus_mayeri
Pristionchus_pacificus
Teladorsagia_circumcincta
Ancylostoma_caninum
Ancylostoma_ceylanicum
Ancylostoma_duodenale
Angiostrongylus_cantonensis
Diploscapter_pachys
Haemonchus_placei
Oscheius_tipulae
Strongylus_vulgaris
Bursaphelenchus_xylophilus
Halicephalobus_mephisto
Panagrellus_redivivus
Parastrongyloides_trichosuri
Steinernema_carpocapsae
Steinernema_feltiae
Steinernema_glaseri
Steinernema_monticolum
Steinernema_scapterisci
Strongyloides_papillosus
Strongyloides_ratti
Strongyloides_stercoralis
Strongyloides_venezuelensis
Acrobeloides_nanus
Ditylenchus_destructor
Ditylenchus_dipsaci
Globodera_pallida
Globodera_rostochiensis
Heterodera_glycines
Aligned peptide region:
1
2
3
Occupancy (cutoff = 50 %):
1
100
99
Conserved peptide regions:
1
2
SIGNATURE WEBLOGO:
MOTIF WEBLOGO:

### FLP-20 peptide alignment

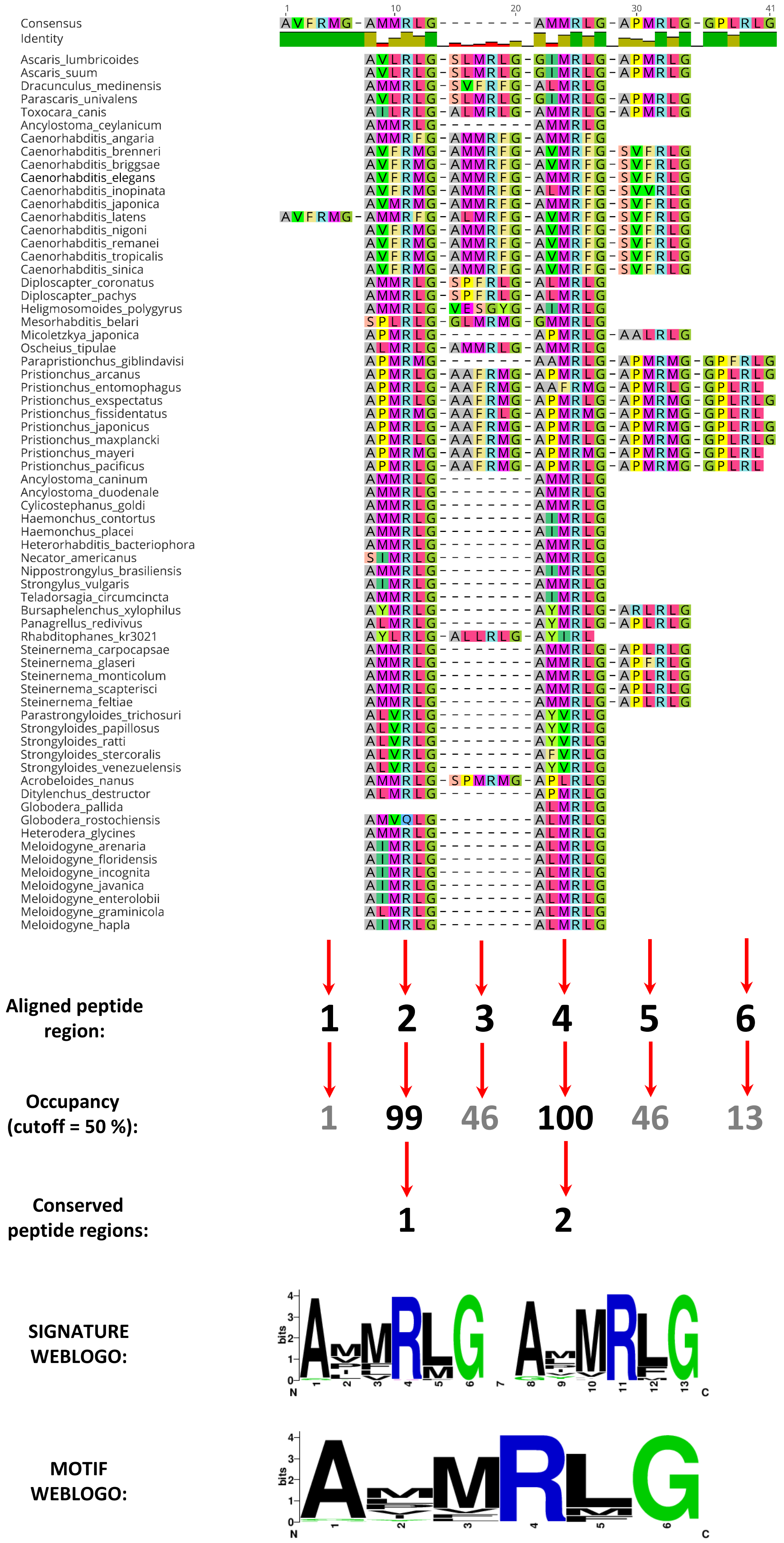

### FLP-21 peptide alignment

| Sequence | Alignment (positions 1–12) |
|---|---|
| Consensus | GXGLGPRPLRFG |
| Identity | |
| Plectus_sambesii | GLGPRPLRFG |
| Ascaris_lumbricoides | GLGPRPLRFG |
| Brugia_malayi | ALGPRPLRFG |
| Brugia_pahangi | ALGPRPLRFG |
| Dracunculus_medinensis | NGLGLRPLRFG |
| Litomosoides_sigmodontis | ALGPRPLRFG |
| Loa_loa | ALGPRPLRFG |
| Onchocerca_volvulus | APGPRPLRFG |
| Acanthocheilonema_viteae | ALGPRPLRFG |
| Anisakis_simplex | GLGPRPLRFG |
| Ascaris_suum | GLGPRPLRFG |
| Brugia_timori | ALGPRPLRFG |
| Dirofilaria_immitis | ARGPRPLRFG |
| Elaeophora_elaphi | ALGPRPLRFG |
| Gongylonema_pulchrum | ALGPRPLRFG |
| Onchocerca_flexuosa | APGPRPLRFG |
| Onchocerca_ochengi | APGPRPLRFG |
| Parascaris_equorum | GLGPRPLRFG |
| Parascaris_univalens | GLGPRPLRFG |
| Thelazia_callipaeda | ALRPRPLRFG |
| Toxocara_canis | GLGPRPLRFG |
| Wuchereria_bancrofti | ALGPRPLRFG |
| Ancylostoma_ceylanicum | GLGPRPLRFG |
| Angiostrongylus_costaricensis | GLGPRPLRFG |
| Caenorhabditis_angaria | GLGPRPLR |
| Caenorhabditis_brenneri | GLGPRPLRFG |
| Caenorhabditis_briggsae | GLGPRPLRFG |
| Caenorhabditis_elegans | GLGPRPLRFG |
| Caenorhabditis_inopinata | GLGPRPLRFG |
| Caenorhabditis_japonica | GLGPRPLRFG |
| Caenorhabditis_latens | GLGPRPLRFG |
| Caenorhabditis_remanei | GLGPRPLRFG |
| Diploscapter_coronatus | GLGPRPLRFG |
| Haemonchus_contortus | GLGPRPLRFG |
| Haemonchus_placei | GLGPRPLRFG |
| Heligmosomoides_polygyrus | GLGPRPLRFG |
| Oscheius_tipulae | GLGPRPLRFG |
| Ancylostoma_duodenale | GLGPRPLRFG |
| Angiostrongylus_cantonensis | GLGPRPLRFG |
| Caenorhabditis_sinica | GLGPRPLRFG |
| Caenorhabditis_tropicalis | GLGPRPLRFG |
| Cylicostephanus_goldi | GLGPRPLRFG |
| Diploscapter_pachys | GLGPRPLRFG |
| Heterorhabditis_bacteriophora | GLGPRPLRFG |
| Mesorhabditis_belari | GHGPRPLRFG |
| Necator_americanus | GLGPRPLRFG |
| Teladorsagia_circumcincta | GLGPRPLRFG |
| Bursaphelenchus_xylophilus | GMGPRPLRFG |
| Halicephalobus_mephisto | ALGPRPLRFG |
| Panagrellus_redivivus | ALGPRPLRFG |
| Parastrongyloides_trichosuri | AVGPRPLRFG |
| Steinernema_carpocapsae | GLGPRPLRFG |
| Steinernema_feltiae | GLGPRPLRFG |
| Steinernema_glaseri | GLGPRPLRFG |
| Steinernema_monticolum | GLGPRPLRFG |
| Steinernema_scapterisci | GLGPRPLRFG |
| Strongyloides_papillosus | AVGPRPLRFG |
| Strongyloides_ratti | AVGPRPLRFG |
| Strongyloides_stercoralis | AVGPRPLRFG |
| Strongyloides_venezuelensis | AVGPRPLRFG |
| Acrobeloides_nanus | GLGPRPLRFG |
| Ditylenchus_destructor | GLGMGPRPLRFG |
| Globodera_rostochiensis | GSLGPRPLRFG |
| Ditylenchus_dipsaci | GLGPRPLRFG |

**Aligned peptide region:** 1

**Occupancy (cutoff = 50 %):** 100

**Conserved peptide regions:** 1

**SIGNATURE/MOTIF WEBLOGO:**

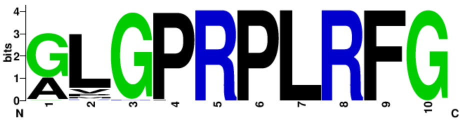

### FLP-22 peptide alignment

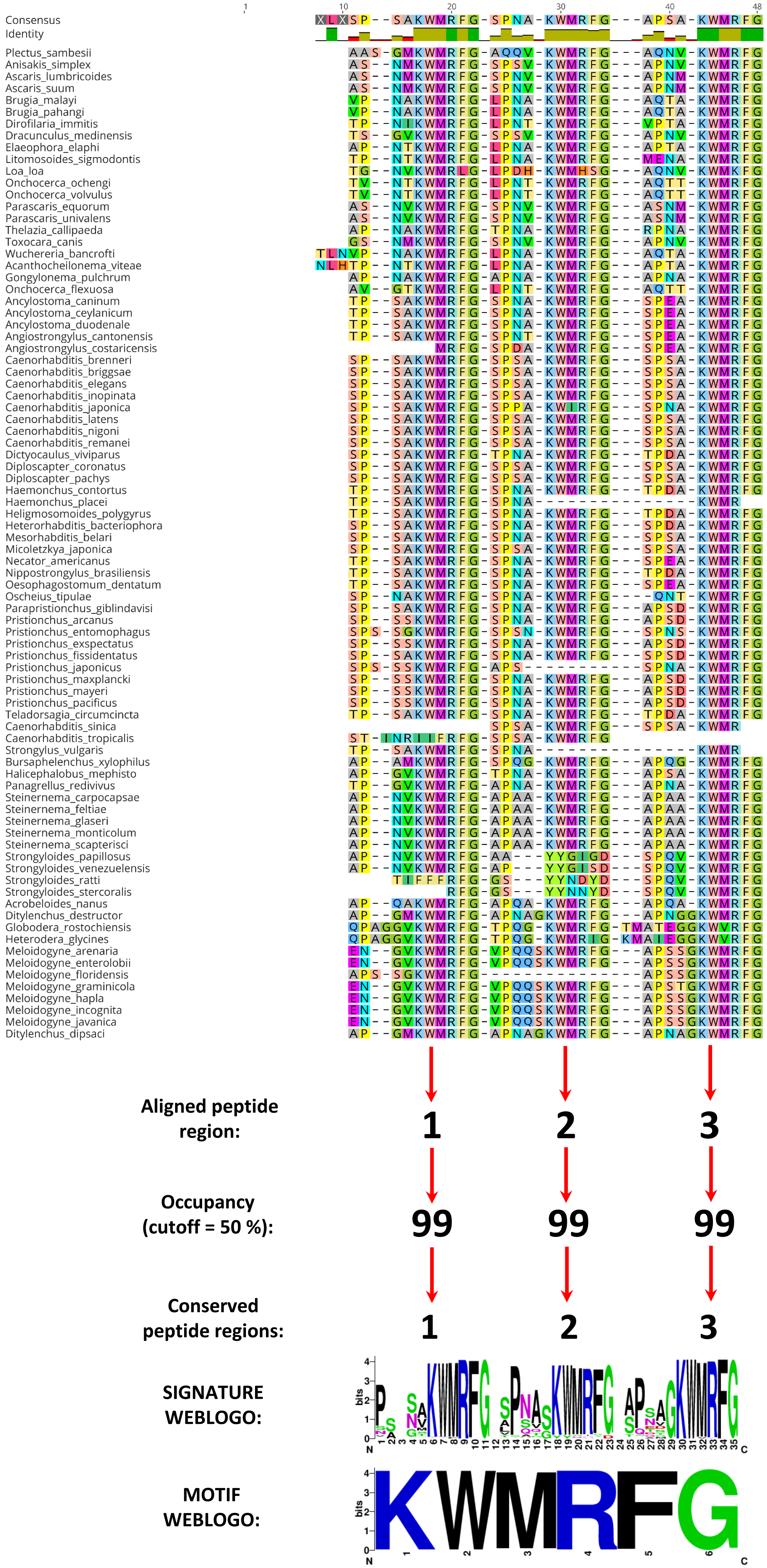

### FLP-23 peptide alignment

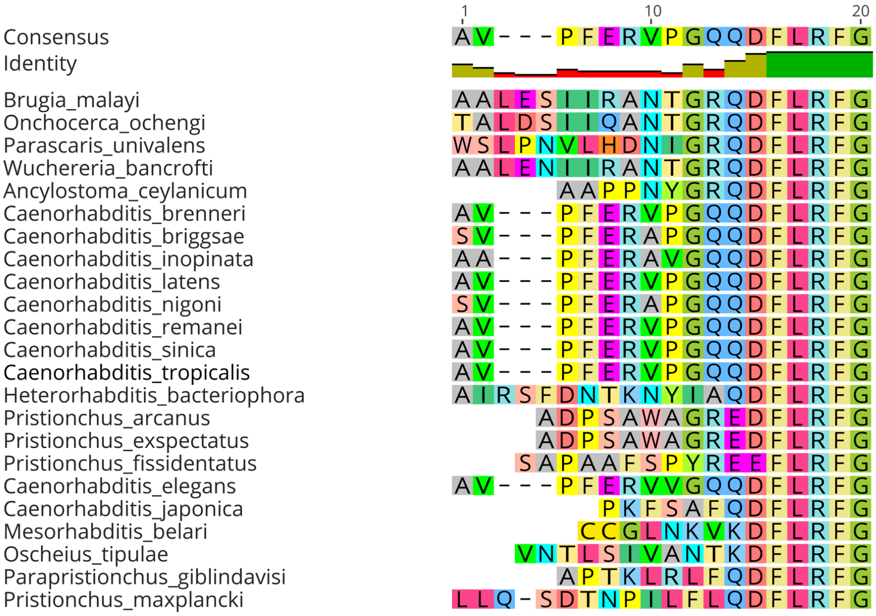


**Aligned peptide region:** 1

**Occupancy (cutoff = 50 %):** 100

**Conserved peptide regions:** 1

**SIGNATURE/MOTIF WEBLOGO:**

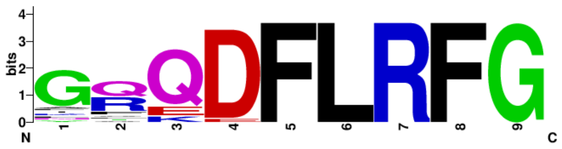

### FLP-24 peptide alignment

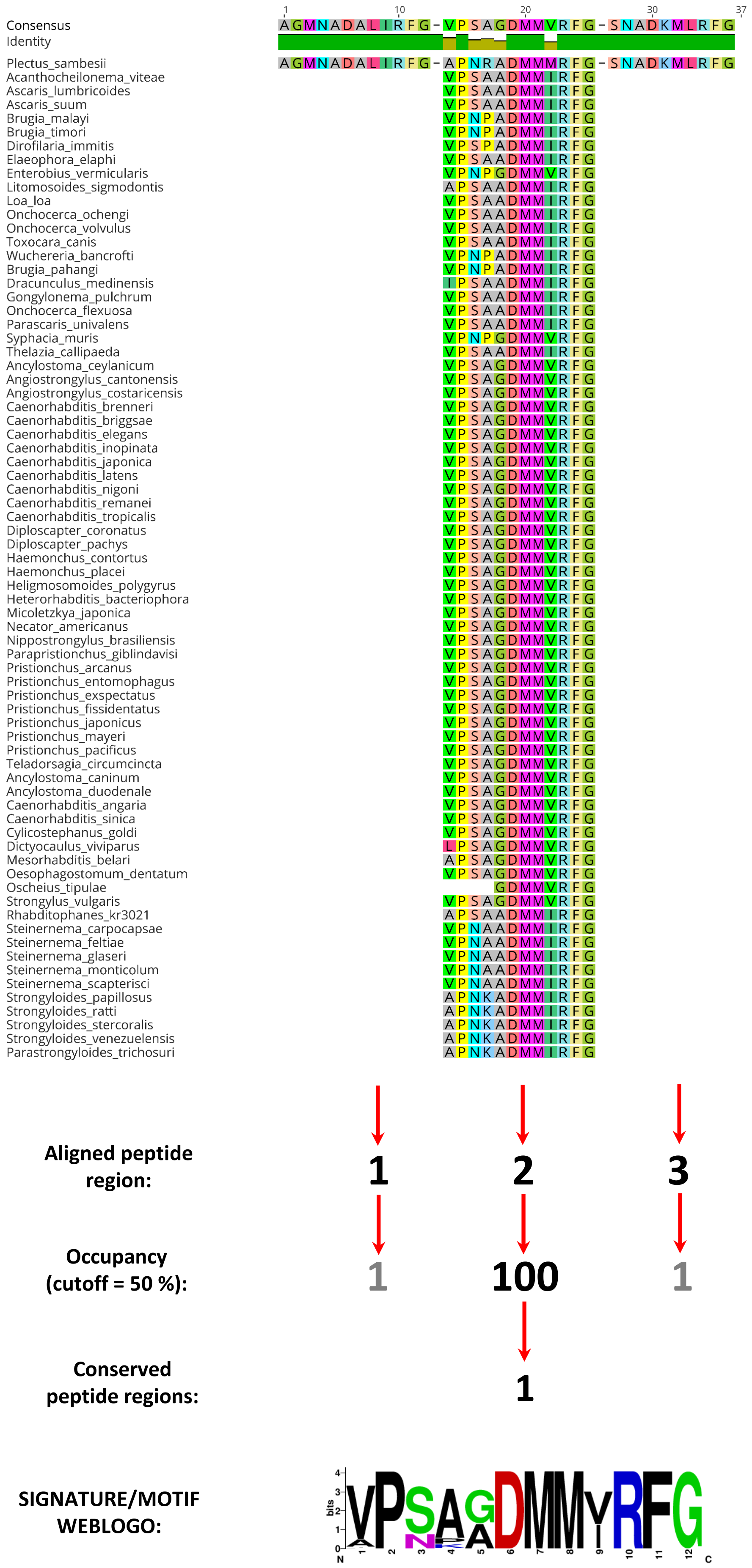

### FLP-25 peptide alignment

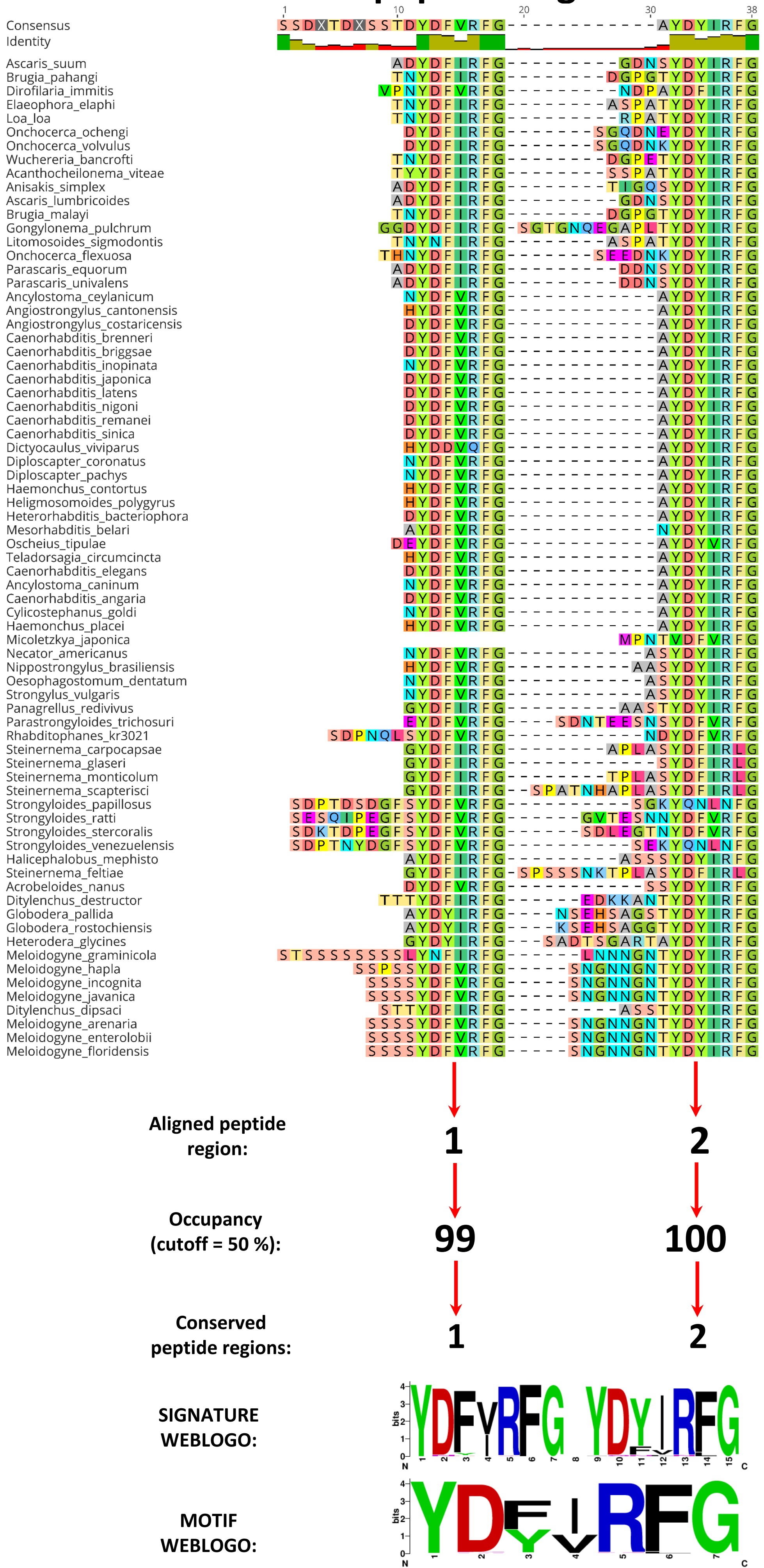

### FLP-26 peptide alignment

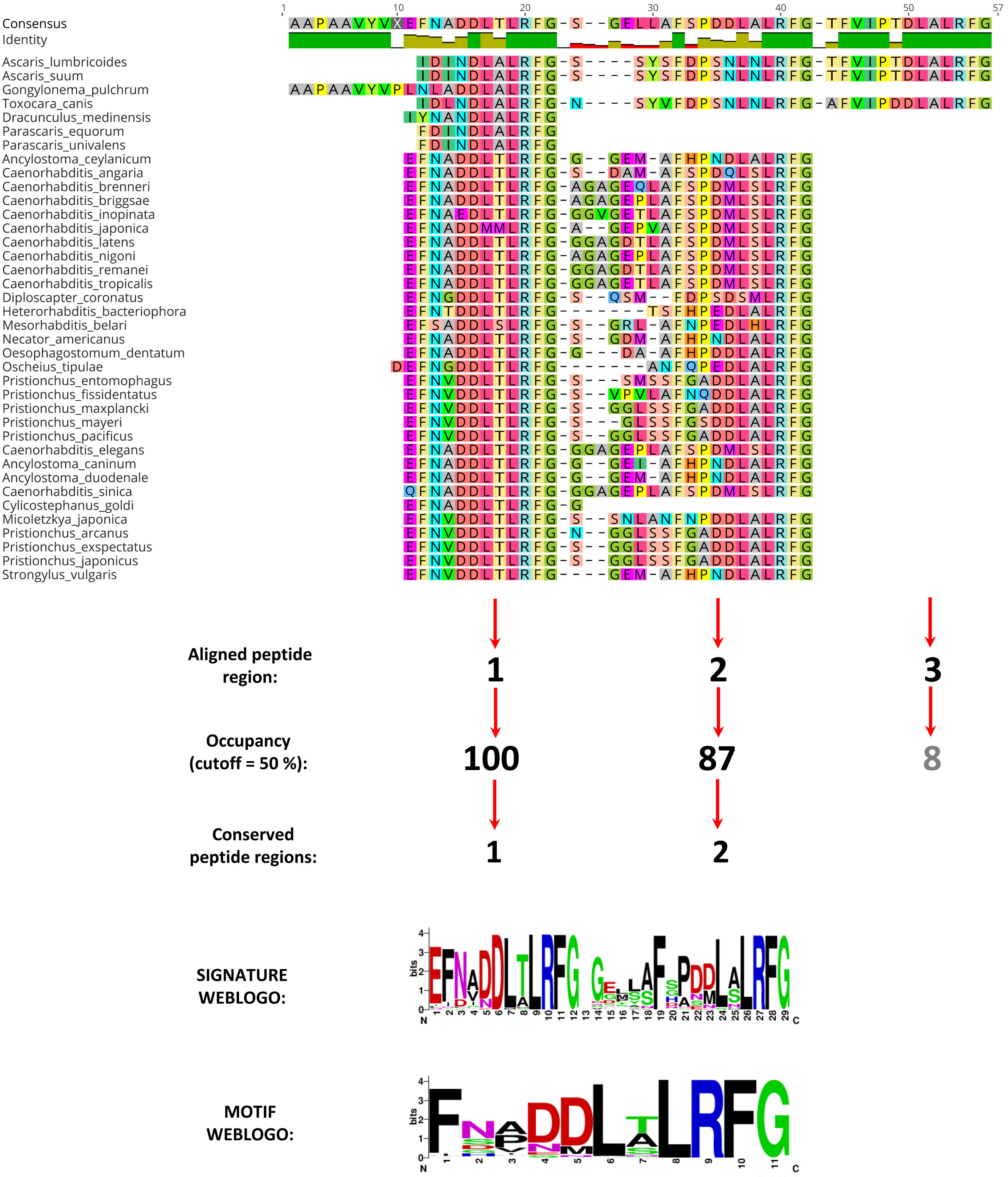

### FLP-27 peptide alignment

| | Sequence |
|---|---|
| Consensus | SQTLGGRMRFG |
| Identity | |
| Ancylostoma_caninum | TMGGRMRFG |
| Ancylostoma_ceylanicum | TMGGRMRFG |
| Caenorhabditis_brenneri | GLGGRMRFG |
| Caenorhabditis_briggsae | GLGGRMRFG |
| Caenorhabditis_inopinata | GLGGRMRFG |
| Caenorhabditis_latens | GLGGRMRFG |
| Caenorhabditis_nigoni | GLGGRMRFG |
| Caenorhabditis_remanei | GLGGRMRFG |
| Caenorhabditis_sinica | GLGGRMRFG |
| Micoletzkya_japonica | TLGGRMRFG |
| Necator_americanus | TMGGRMRFG |
| Oesophagostomum_dentatum | TMGGRMRFG |
| Oscheius_tipulae | SMGGRMRFG |
| Pristionchus_arcanus | SQALGGRMRFG |
| Pristionchus_entomophagus | TQTIGGRMRFG |
| Pristionchus_exspectatus | SQALGGRMRFG |
| Pristionchus_fissidentatus | TQTLGGRMRFG |
| Pristionchus_japonicus | SQTLGGRMRFG |
| Pristionchus_maxplancki | SQTLGGRMRFG |
| Pristionchus_pacificus | SQALGGRMRFG |
| Caenorhabditis_elegans | GLGGRMRFG |

**Aligned peptide region:** 1

**Occupancy (cutoff = 50 %):** 100

**Conserved peptide regions:** 1

**SIGNATURE/MOTIF WEBLOGO:**

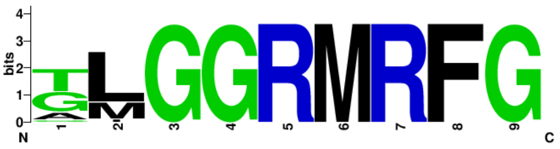

### FLP-28 peptide alignment

| | 1 … 10 |
|---|---|
| Consensus | APNRILMRFG |
| Identity | |
| Ascaris_lumbricoides | APNKILMRFG |
| Ascaris_suum | APNKILMRFG |
| Parascaris_equorum | APNKILMRFG |
| Toxocara_canis | APNKILMRFG |
| Anisakis_simplex | APNKILMRFG |
| Dracunculus_medinensis | APNKILMRFG |
| Parascaris_univalens | APNKILMRFG |
| Ancylostoma_caninum | APNRILMRFG |
| Ancylostoma_ceylanicum | APNRILMRFG |
| Ancylostoma_duodenale | APNRILMRFG |
| Angiostrongylus_costaricensis | APNRILMRFG |
| Caenorhabditis_brenneri | APNRVLMRFG |
| Caenorhabditis_elegans | APNRVLMRFG |
| Caenorhabditis_briggsae | APNRVLMRFG |
| Caenorhabditis_inopinata | APNRVLMRFG |
| Caenorhabditis_japonica | APNRVLMRFG |
| Caenorhabditis_latens | APNRVLMRFG |
| Caenorhabditis_nigoni | APNRVLMRFG |
| Caenorhabditis_remanei | APNRVLMRFG |
| Caenorhabditis_tropicalis | APNRVLMRFG |
| Cylicostephanus_goldi | APNRILMRFG |
| Haemonchus_contortus | VPNRIFMRFG |
| Haemonchus_placei | VPNRIFMRFG |
| Heligmosomoides_polygyrus | APNRILMRFG |
| Micoletzkya_japonica | APNRVLMRFG |
| Oesophagostomum_dentatum | APNRILMRFG |
| Parapristionchus_giblindavisi | APNRVLMRFG |
| Pristionchus_arcanus | APSRVLMRFG |
| Pristionchus_entomophagus | APSRVLMRFG |
| Pristionchus_exspectatus | APSRVLMRFG |
| Pristionchus_fissidentatus | APSRVMMRFG |
| Pristionchus_japonicus | APSRVLMRFG |
| Pristionchus_maxplancki | APSRVLMRFG |
| Pristionchus_mayeri | APSRVLMRFG |
| Pristionchus_pacificus | APSRVLMRFG |
| Strongylus_vulgaris | APNRILMRFG |
| Angiostrongylus_cantonensis | APNRILMRFG |
| Caenorhabditis_angaria | APNRVLMRFG |
| Dictyocaulus_viviparus | APHRILMRFG |
| Diploscapter_coronatus | ASNRVLMRFG |
| Diploscapter_pachys | ASNRVLMRFG |
| Heterorhabditis_bacteriophora | APNRILMRFG |
| Mesorhabditis_belari | APNRILMRFG |
| Necator_americanus | APNRILMRFG |
| Nippostrongylus_brasiliensis | VPNRILMRFG |
| Oscheius_tipulae | APNRILMRFG |
| Teladorsagia_circumcincta | APNRIFMRFG |
| Parastrongyloides_trichosuri | APNRVMMRFG |
| Rhabditophanes_kr3021 | APSRVMMRFG |
| Steinernema_carpocapsae | APNRILMRFG |
| Steinernema_feltiae | APNRILMRFG |
| Steinernema_monticolum | APNRILMRFG |
| Steinernema_scapterisci | APNRILMRFG |
| Strongyloides_ratti | APNRVMMRFG |
| Strongyloides_venezuelensis | APNRVMMRFG |
| Steinernema_glaseri | APNRILMRFG |
| Strongyloides_papillosus | APNRVMMRFG |
| Strongyloides_stercoralis | APNRVMMRFG |

**Aligned peptide region:** 1

**Occupancy (cutoff = 50 %):** 100

**Conserved peptide regions:** 1

**SIGNATURE/MOTIF WEBLOGO:**

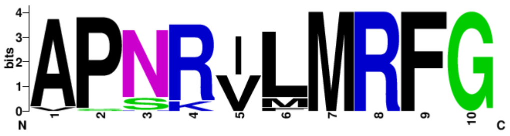

### FLP-31 peptide alignment

| | 1 – 11 |
|---|---|
| Consensus | L Y R P R G P P R F G |
| Identity | |
| Meloidogyne_arenaria | L Y R P R G P P R F G |
| Meloidogyne_javanica | L Y R P R G P P R F G |
| Meloidogyne_enterolobii | L Y R P R G P P R F G |
| Meloidogyne_floridensis | L Y R P R G P P R F G |
| Meloidogyne_graminicola | L Y R P R G P P R F G |
| Meloidogyne_hapla | L Y R P R G P P R F G |
| Meloidogyne_incognita | L Y R P R G P P R F G |

**Aligned peptide region:** 1

**Occupancy (cutoff = 50 %):** 100

**Conserved peptide regions:** 1

**SIGNATURE/MOTIF WEBLOGO:**

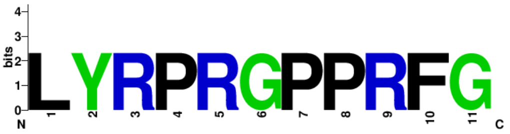

### FLP-32 peptide alignment

| | 1 … 10 |
|---|---|
| Consensus | AMRNSLVRFG |
| Identity | |
| Ancylostoma_caninum | AMRNSLVRFG |
| Angiostrongylus_cantonensis | AMRNSLVRFG |
| Caenorhabditis_sinica | AMRNSLVRFG |
| Caenorhabditis_tropicalis | AMRNSLVRFG |
| Cylicostephanus_goldi | AMRNSLVRFG |
| Diploscapter_pachys | AMRNSLVRFG |
| Heterorhabditis_bacteriophora | AMRNSLVRFG |
| Micoletzkya_japonica | AMRNSLVRFG |
| Parapristionchus_giblindavisi | AMRNSLVRFG |
| Pristionchus_arcanus | AMRNSLVRFG |
| Pristionchus_entomophagus | AMRNSLVRFG |
| Pristionchus_exspectatus | AMRNSLVRFG |
| Pristionchus_maxplancki | AMRNSLVRFG |
| Pristionchus_mayeri | AMRNSLVRFG |
| Strongylus_vulgaris | AMRNSLVRFG |
| Caenorhabditis_elegans | AMRNSLVRFG |
| Ancylostoma_ceylanicum | AMRNSLVRFG |
| Ancylostoma_duodenale | AMRNSLVRFG |
| Angiostrongylus_costaricensis | AMRNSLVRFG |
| Caenorhabditis_angaria | AMRNSLVRFG |
| Caenorhabditis_brenneri | AMRNSLVRFG |
| Caenorhabditis_briggsae | AMRNSLVRFG |
| Caenorhabditis_inopinata | AMRNSLVRFG |
| Caenorhabditis_japonica | AMRNSLVRFG |
| Caenorhabditis_latens | AMRNSLVRFG |
| Caenorhabditis_nigoni | AMRNSLVRFG |
| Caenorhabditis_remanei | AMRNSLVRFG |
| Dictyocaulus_viviparus | AMRNSLVRFG |
| Diploscapter_coronatus | AMRNSLVRFG |
| Heligmosomoides_polygyrus | AMRNSLVRFG |
| Mesorhabditis_belari | AMRNSLVRFG |
| Necator_americanus | AMRNSLVRFG |
| Nippostrongylus_brasiliensis | AMRNSLVRFG |
| Oesophagostomum_dentatum | AMRNSLVRFG |
| Oscheius_tipulae | AMRNSLVRFG |
| Pristionchus_fissidentatus | AMRNSLVRFG |
| Pristionchus_japonicus | AMRNSLVRFG |
| Pristionchus_pacificus | AMRNSLVRFG |
| Bursaphelenchus_xylophilus | AMRNSLVRFG |
| Panagrellus_redivivus | AMRNSLVRFG |
| Parastrongyloides_trichosuri | AMRNSLVRFG |
| Steinernema_carpocapsae | AMRNSLVRFG |
| Steinernema_feltiae | AMRNSLVRFG |
| Steinernema_glaseri | AMRNSLVRFG |
| Steinernema_monticolum | AMRNSLVRFG |
| Steinernema_scapterisci | AMRNSLVRFG |
| Strongyloides_papillosus | AMRNSLVRFG |
| Strongyloides_ratti | AMRNSLIRFG |
| Strongyloides_stercoralis | AMRNSLIRFG |
| Strongyloides_venezuelensis | AMRNSLVRFG |
| Halicephalobus_mephisto | AMRNSLVRFG |
| Acrobeloides_nanus | AMRNSLVRFG |
| Ditylenchus_destructor | AMRNSLVRFG |
| Ditylenchus_dipsaci | AMRNSLVRFG |
| Globodera_pallida | AMRNALVRFG |
| Globodera_rostochiensis | AMRNALVRFG |
| Heterodera_glycines | AMRNALVRFG |
| Meloidogyne_arenaria | AMRNSLVRFG |
| Meloidogyne_enterolobii | AMRNSLVRFG |
| Meloidogyne_floridensis | AMRNSLVRFG |
| Meloidogyne_hapla | AMRNSLVRFG |
| Meloidogyne_incognita | AMRNSLVRFG |
| Meloidogyne_javanica | AMRNSLVRFG |
| Meloidogyne_graminicola | AMRNSLVRFG |

**Aligned peptide region:** 1

**Occupancy (cutoff = 50 %):** 100

**Conserved peptide regions:** 1

**SIGNATURE/MOTIF WEBLOGO:**

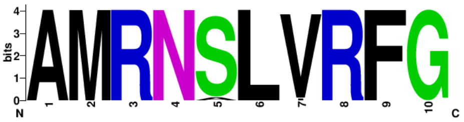

### FLP-33 peptide alignment

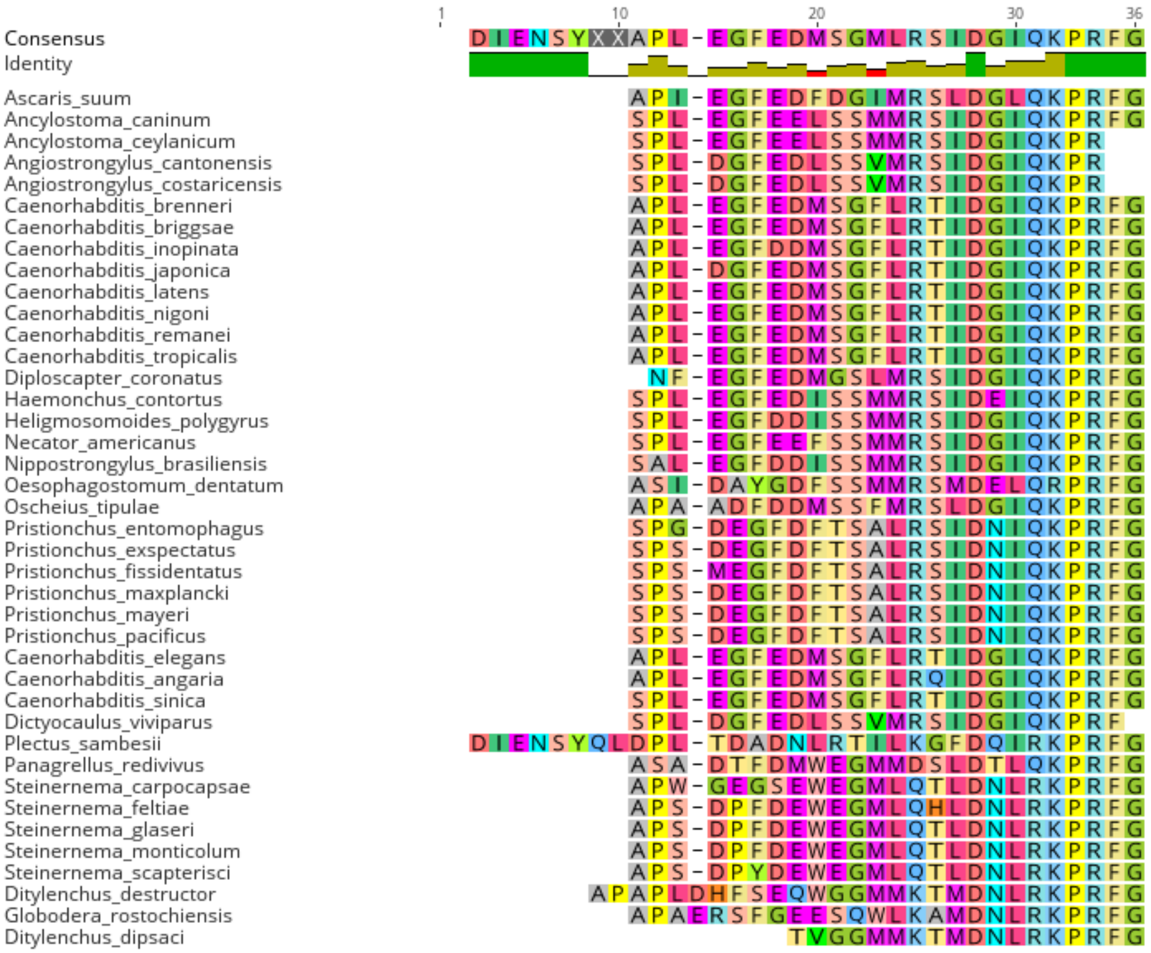


**Aligned peptide region:** **1**

**Occupancy (cutoff = 50 %):** **100**

**Conserved peptide regions:** **1**

**SIGNATURE/MOTIF WEBLOGO:**

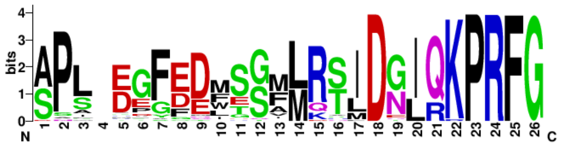

### FLP-34 peptide alignment

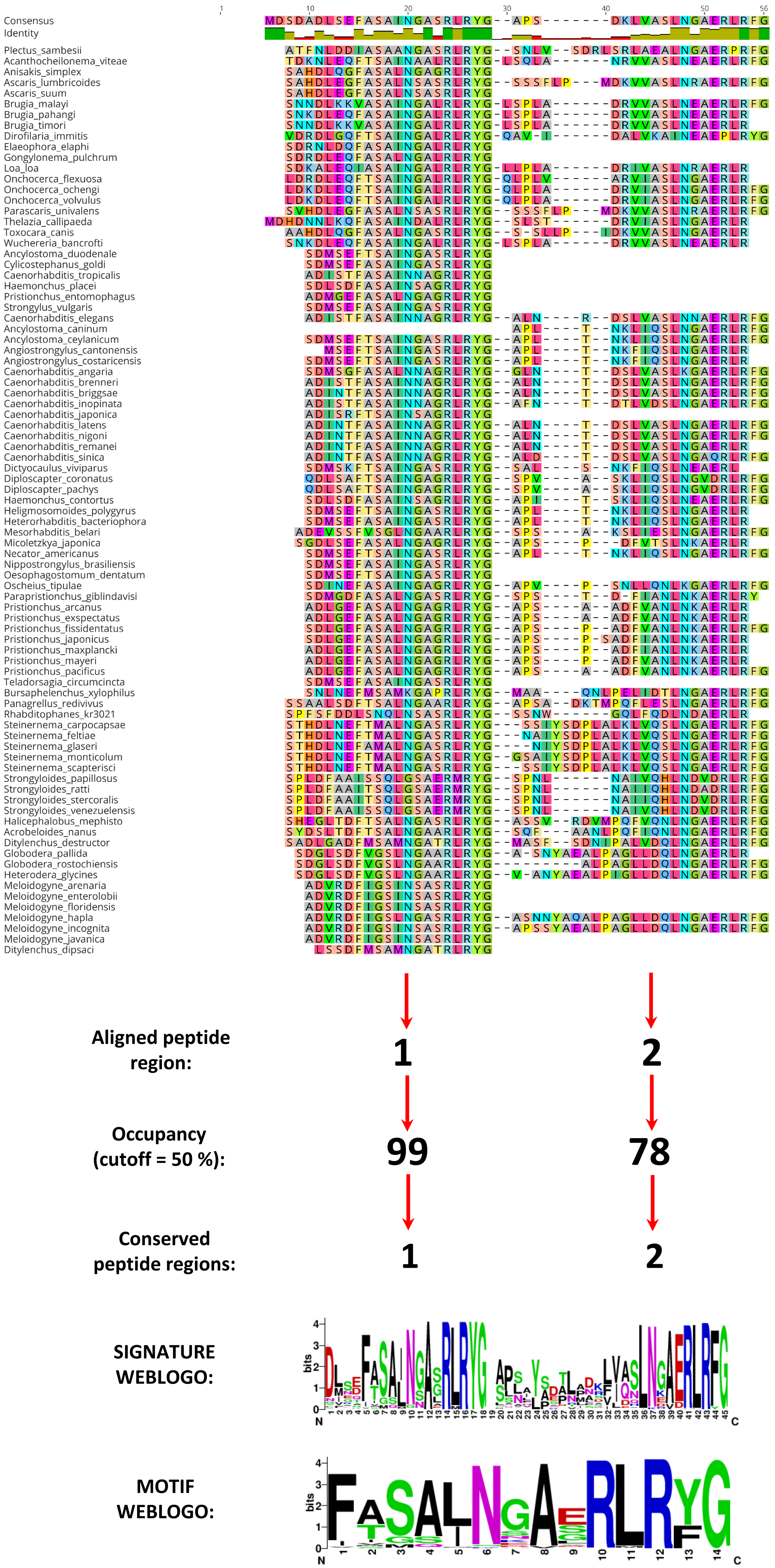
