## Supplementary figures and images for "Exploitation of phylum-spanning omics resources reveals complexity in the nematode FLP signalling system and provides insights into *flp*-gene evolution"

### Supplementary Figure 2

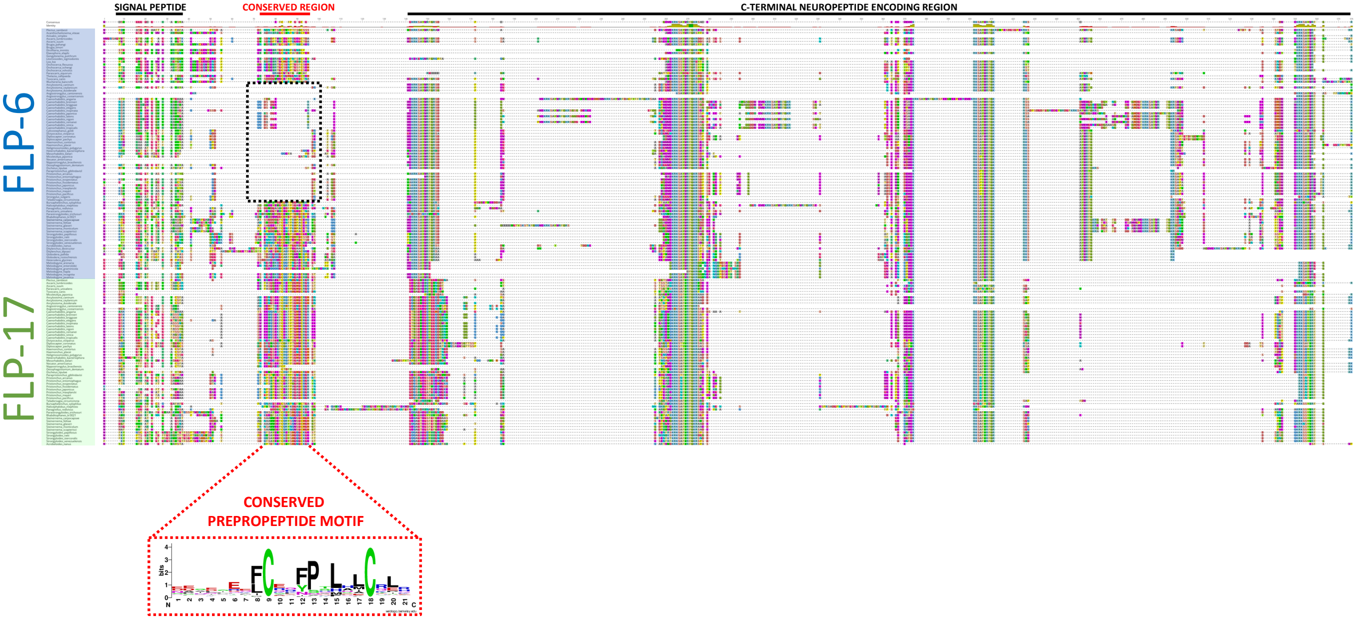

### Supplementary Figure 3

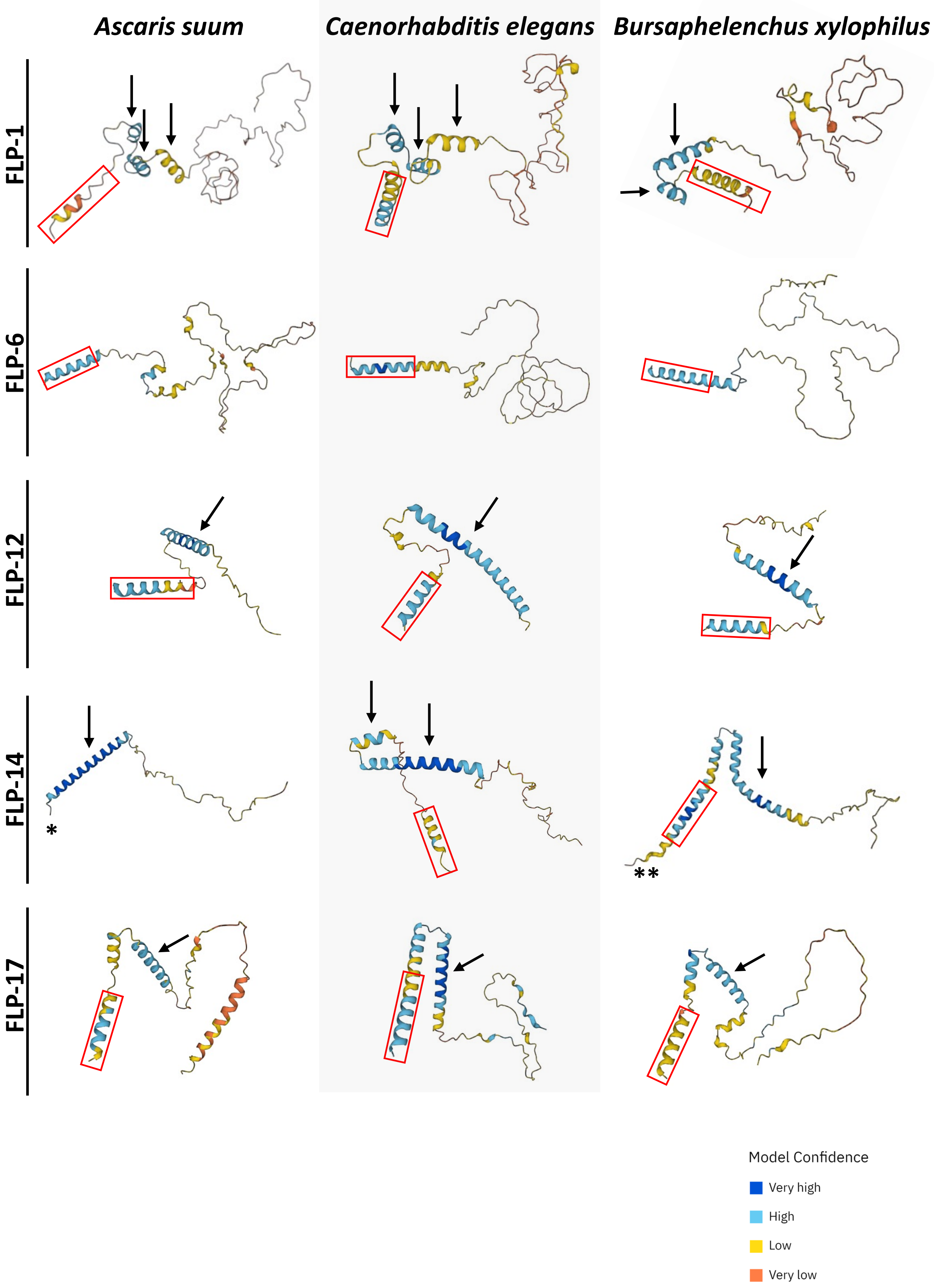

### Supplementary Figure 4

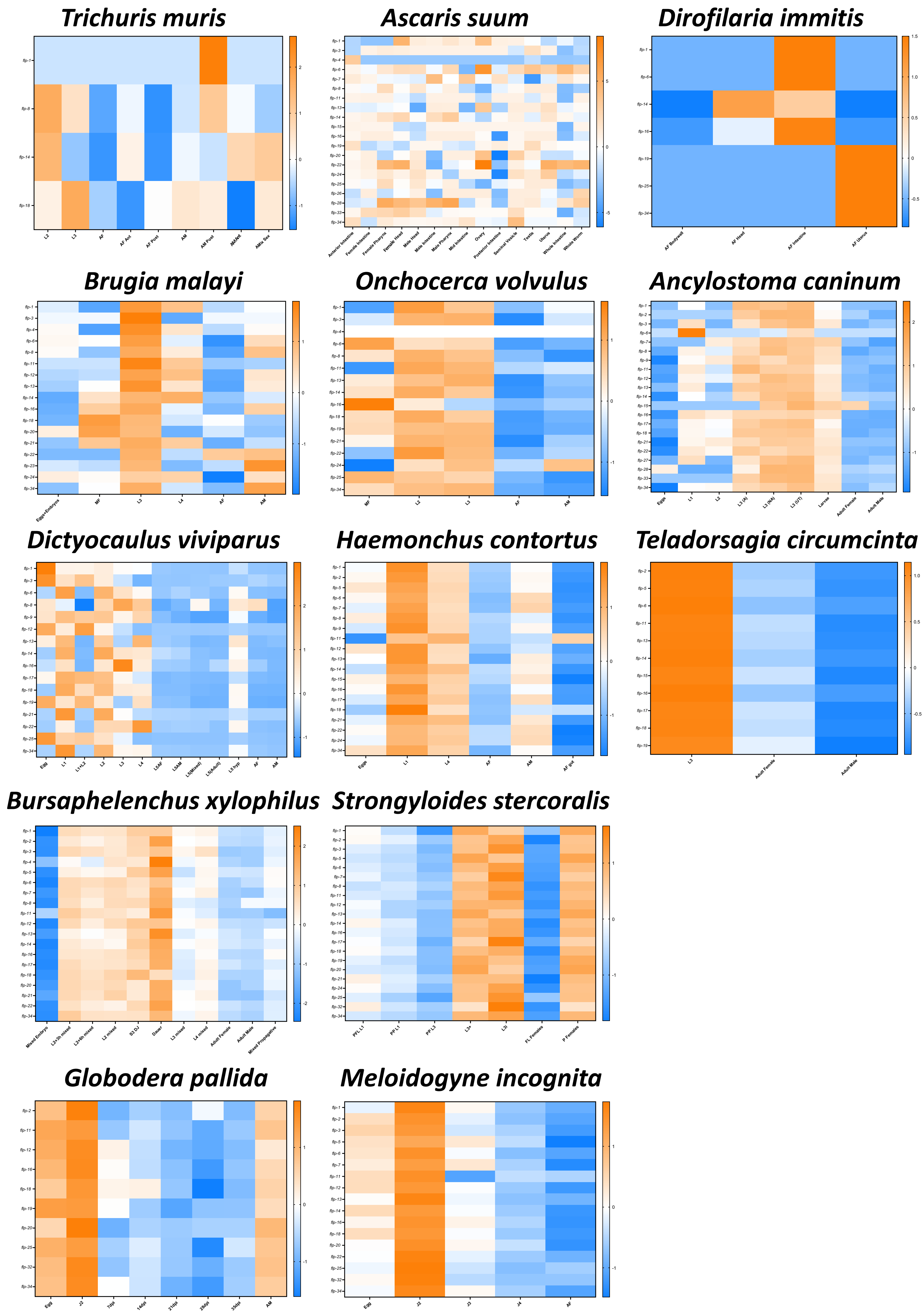

### Supplementary Figure 5

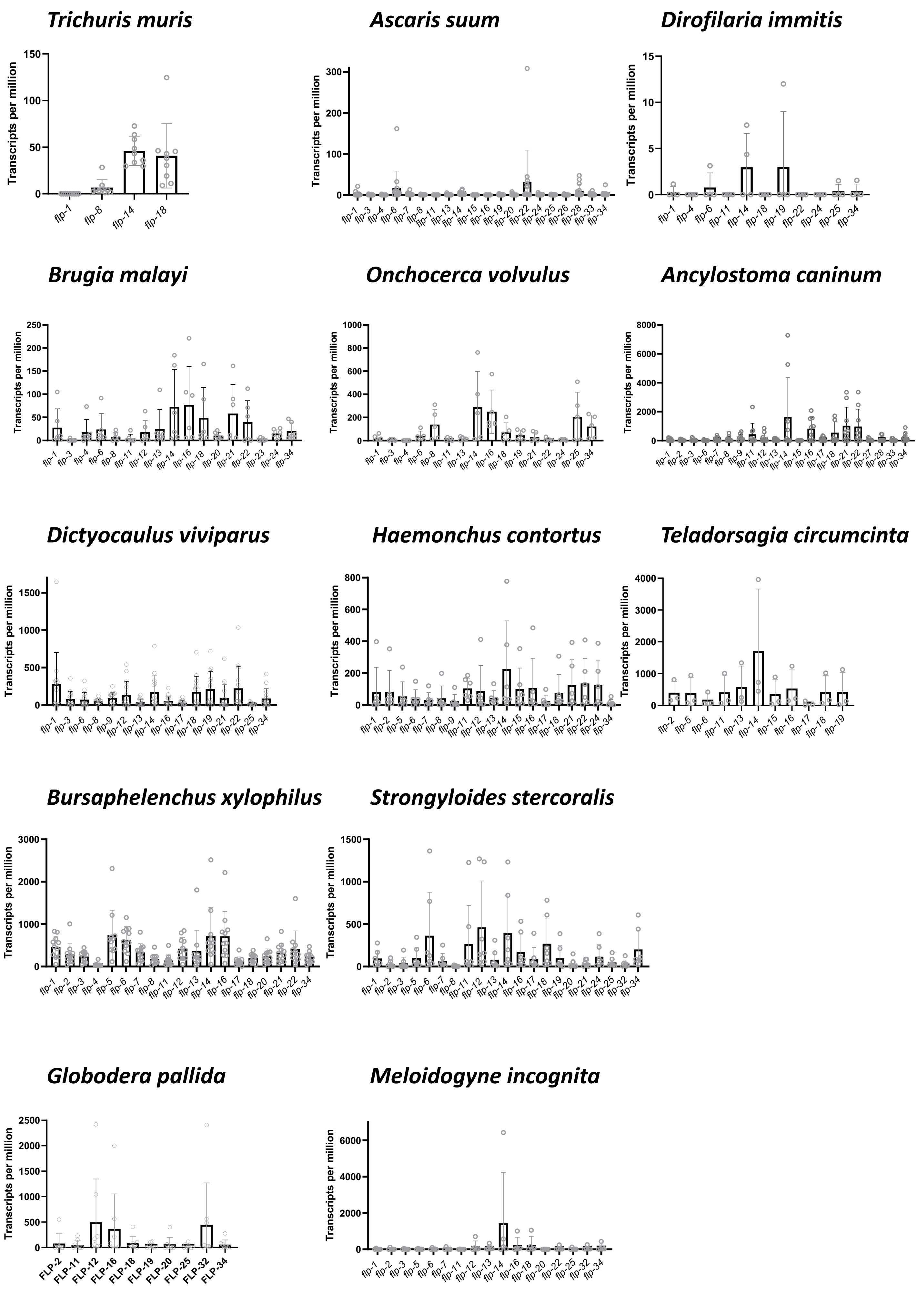
